# Widespread Presence of Catalytic RNAs in All Domains of Life

**DOI:** 10.64898/2026.10.26.761763

**Authors:** Fei Qi, Yue Chen, Yiting Lin, Zipeng Chen, Yuhe Zhong, Fuchun Zhang, Kejia Li, Yonghong Zhao, Meng Li, Siqi Chen, Ilia Kruglikov, Wenjing Yin, Qinling Tang, Junjie Chen, Wenzhui Zhang, Lin Huang, David MJ Lilley, Philipp Kapranov

## Abstract

Very few ribozymes have been found in complex eukaryotes. Here, we developed a novel method, AbRiDGed, for highly accurate and sensitive *ab initio* self-cleaving ribozyme discovery. Our results indicate the existence of millions of self-cleaving ribozymes in humans and other species. We characterized ∼150 novel classes of self-cleaving ribozymes, including one class which exhibits noncanonical biochemical properties and widespread presence in all domains of life and appears to play specific functions in complex organisms. These results revealed that the true complexity of the “ribozyme-ome” is several orders of magnitude higher than previously estimated. We demonstrate that newly identified self-cleaving ribozymes influence organismal fitness by regulating gene expression and enhancing transcriptome complexity *in vivo*. Moreover, these ribozymes hold promise as foundational tools for biotechnological applications, such as direct RNA detection technologies. Overall, catalytic RNAs are prevalent and diverse, and may represent a hidden regulatory layer in various organisms.

## Introduction

The groundbreaking discovery of catalytic RNAs, or ribozymes^1,2^, led to the realization that enzymes, previously thought to be exclusively represented by protein-based entities, can be completely made of RNA. It also significantly expanded the repertoire of the roles played by RNA in the cell beyond being a template for protein synthesis. However, despite the wide array of catalytic activities obtained for artificial ribozymes generated using *in vitro* evolution and selection strategies^3^, a handful of naturally occurring ribozyme classes found in the genomes of modern organisms stand in stark contrast with the vast diversity of known protein-based enzymes represented by thousands of different classes^4^. The paucity of naturally occurring ribozymes is particularly striking in complex organisms, especially in mammals where known naturally occurring ribozymes are very rare and limited to very few classes existing ubiquitously in nature. Overall, until very recently^5^, no classes of ribozymes that emerged recently in the evolution of eukaryotic species have been found. These observations led to an assumption that ribozymes represent vestiges of the “RNA world” where they were quite common in the primordial organisms but then got mostly replaced by protein-based enzymes^4,6,7^. As such, the emergence of novel classes of ribozymes in complex organisms is expected to happen extremely rarely if at all.

The scarcity of naturally occurring ribozymes is especially conspicuous considering their potential for not only enzymatic activities, but also regulation of gene expression. Such a function is performed by the *glmS* class of self-cleaving ribozymes which sense a metabolite and regulate the expression level of the enzyme synthesizing the metabolite in a negative feedback loop^8,9^. Furthermore, the ability of self-cleaving ribozymes to break the 3′,5′-phosphodiester bonds in RNA at specific sites also represents a theoretically useful RNA processing solution, especially in cells of complex eukaryotes where alternative RNA processing is an extremely important part of cellular physiology^10^.

In addition to the ribozymes discovered using traditional approaches in the process of investigating specific biological phenomena, such as splicing or RNA processing, most known ribozymes were identified using bioinformatics-based genome-wide surveys for sequences that either (1) resemble known ribozymes and/or (2) have signatures of evolutionary conservation of RNA structures^11^. While successful in finding multiple members of the known ribozyme classes and a few novel classes, such surveys cannot identify novel ribozymes that either have evolved recently and/or are limited to very few species.

Our discovery of the hovlinc class of ribozymes in humans challenged some of the assumptions above and opened a number of new and exciting possibilities that call for high-throughput ribozyme discovery efforts using *ab initio* genome-wide screenings unbiased by prior estimations of sequence-structure patterns or conservation properties of these catalytic RNA molecules^5^. Furthermore, our previous study suggested that further discovery of self-cleaving ribozymes would require novel approaches with significantly improved sensitivity and accuracy^5^. Therefore, in this study we developed a novel genome-wide screening method for self-cleaving ribozymes, named AbRiDGed (*ab initio* ribozyme discovery genome-wide), and by applying it, discovered and characterized a multitude of novel self-cleaving ribozymes belonging to distinct novel classes. One such novel class of ribozymes is present abundantly in all domains of life and exhibits a striking pattern of enrichment in genomes of complex organisms indicative of its biological functions. Our results also show that the ribozymes discovered in this study are active *in vivo* and contribute to the transcriptomic complexity. Novel self-cleaving ribozymes appear to be refractory to sequence changes that abolish their activities. These results suggest that catalytic RNAs represent a prominent and yet mostly untouched area of biology of eukaryotic cells. Based on one novel ribozyme, we developed a prototype of protein-free method for direct RNA detection, demonstrating that novel ribozymes can also provide solutions for novel biotechnological applications.

## Results

### Discovery of numerous self-cleaving ribozymes in the human genome using AbRiDGed

A self-cleaving ribozyme catalyzes intramolecular transesterification reaction resulting in the cleavage of its own RNA molecule at a specific position — the cleavage site^12,13^. The AbRiDGed method identifies genomic positions of ribozyme cleavage sites with nucleotide-level precision by taking advantage of the unique property of the downstream cleavage products — their 5′-OH termini (Figures 1A and S1). This method starts from the generation of a genome-wide library of *in vitro* transcription (IVT) products that are subjected to the self-cleavage reaction followed by treatments with the RppH and XRN-1 enzymes to enrich the downstream cleavage products possessing 5′-OH termini (Figures 1A and S1; see Methods for details). The 5′-OH termini of the downstream cleavage products are then tagged by either the ligation of synthetic RNA oligonucleotides using the RtcB ligase^14^ in the L1/L2 libraries or the addition of homopolymeric tails to the corresponding cDNAs using terminal transferase in the TA/TC libraries (Figures 1A and S1). The consistency of detection of a potential cleavage site across the 4 different library types represents the key metric for ribozyme discovery in the AbRiDGed method.

**Figure 1.**
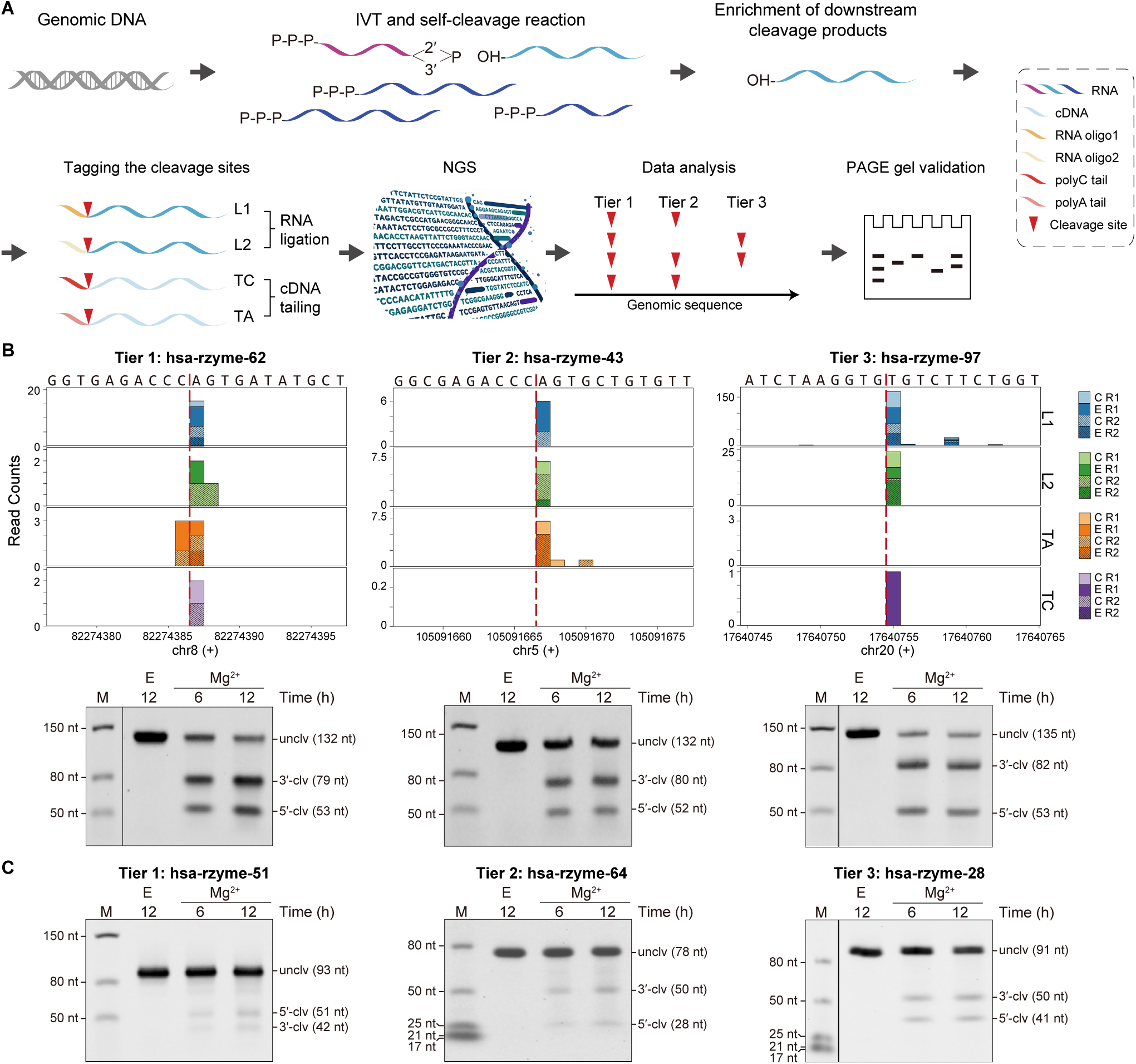
Discovery of novel human self-cleaving ribozymes via AbRiDGed. (A) Schematic representation of the ribozyme discovery pipeline (see Figure S1 for detailed steps). IVT, *in vitro* transcription; NGS, next-generation sequencing; PAGE, polyacrylamide gel electrophoresis. (B) Representative examples of the novel self-cleaving ribozymes. Top, the read counts (Y-axes) from the NGS data in the genomic vicinities of the estimated cleavage sites marked by the red hashed lines. The X-axes denote the genomic coordinates, and the corresponding sequences are shown on the top. “L1”, “L2”, “TA” and “TC” denote the library types; “R1” and “R2” denote the replicates; “E” and “C” denote samples that were respectively treated or not with the RppH and XRN-1 enzymes. Bottom, the corresponding validation gels confirming the self-cleaving activity of the 3 ribozymes. (C) Validation gels for the 3 novel self-cleaving ribozymes found in the randomly selected human genomic regions. (B and C) The uncleaved RNAs (“unclv”) and the upstream and downstream cleavage products (“5′-clv” and “3′-clv”, respectively) are marked with their expected sizes shown in the parentheses. Note, when calculating the expected sizes, the 1–3 guanosines potentially added during sequence construction were included. The “E” lanes contain the control reactions with 6 mM EDTA and no Mg^2+^; the “M” lanes contain the size standards. “Time” shows the duration of the cleavage reaction in hours. See Figures S5 and S6 for the 2 replicates of the validation gels for all 105 novel human ribozymes.

We applied the AbRiDGed method to survey the human genome for novel self-cleaving ribozymes. To characterize the performance of the method, we added spike-ins based on known self-cleaving ribozymes of 9 classes and proven self-cleavage activity^15^ (Figure S1 and Table S1). The cleavage sites of all the 20 spike-ins and the 4 known endogenous human ribozymes (the HH9, HH10, CPEB3 and hovlinc ribozymes) were detected in all 4 types of libraries (Figures S2 and S3). The only exception was the failure to detect several HDV or HDV-like ribozymes in the ligation-based libraries (Figures S2 and S3), consistent with the previous results that such ribozymes are poorly detectable using ligation-mediated methods^16^. Nonetheless, the cleavage sites of the HDV and HDV-like ribozymes were detected in the tailing-based libraries (Figures S2 and S3). Thus, the AbRiDGed method is sufficiently robust and sensitive to readily detect the cleavage sites of various types of self-cleaving ribozymes.

Then, based on the consistency of detection across multiple library types, the candidate ribozyme cleavage sites were classified into 3 Tiers as follows: the Tier 1 candidates were detected in all 4 library types, while the Tiers 2 and 3 candidates were detected in respectively 3 and 2 library types (Figure 1A). Overall, we found 60, 3629 and 246,794 candidate cleavage sites from the human genome in respectively Tiers 1–3 (Table S2). To further filter the candidates of Tiers 2 and 3, we calculated signal-to-local-background ratios for the candidate cleavage sites (Methods). Finally, a total of 300 candidate cleavage sites, including the top 170 sites from Tier 2 and top 70 sites from Tier 3, as well as all the 60 sites from Tier 1, were selected for individual validation using a traditional biochemical approach (Methods).

As exemplified in Figure 1B, true self-cleaving ribozymes were defined as those which yielded two clear major cleavage products of the expected sizes in the +Mg^2+^ reactions but not in the +EDTA controls. Using these criteria, we validated 20/60 (33.3%), 67/170 (39.4%) and 15/70 (21.4%) of the respectively Tier 1–3 candidates (Table S2; see Figure S4 for plots of all their cleavage sites, and Figures S5 and S6 for the validation gels of all the newly discovered human ribozymes). Thus, the overall validation rate reached 34% (102 out of 300 tested candidates) which was significantly higher than that in our previous method — 3.6%^5^. In addition, we tested the top 50 candidates found only in a single library type and obtained no positive (Table S3), confirming that the consistent detection across multiple library types is an important feature of accurate ribozyme discovery.

### Vast unannotated self-cleaving ribozyme-ome in the genomes of humans and other eukaryotes

The results above indicated that self-cleaving ribozymes are far more common in the human genome than previously anticipated. To further confirm this, we then randomly selected 3333 regions of 300 nt from both strands of the human genome (Table S4). These regions were amplified from human genomic DNAs, and then subjected to essentially the same AbRiDGed pipeline as the one used in the above genome-wide screening.

We could successfully obtain high-throughput sequencing data for 3191/3333 (95.7%) regions (Table S4), and upon analysis of the signal profiles in the 4 types of libraries, we selected 405 candidate cleavage sites (Methods) for the biochemical validation (Table S4). In total, we could confirm that at least 3/3191 random regions contained true self-cleaving ribozymes (Figures 1C, S4, S5 and S6, and Table S4). Considering that the size of the double-stranded human genome is ∼6 billion nucleotides and it contains ∼20 million non-overlapping 300-nt regions, these results translated into estimated >10 thousand self-cleaving ribozymes present in the genome. This estimate exceeds the 4 previously known human self-cleaving ribozymes by several orders of magnitude.

The multitude of newly discovered self-cleaving ribozymes in the human genome posed a natural question: whether their existence is a common phenomenon shared by many eukaryotes. Therefore, we performed ribozyme screenings in the genomes of a wide range of eukaryotes from relatively simple to complex model organisms, including fungi (*Aspergillus nidulans* and *Trichoderma guizhouense*), plants (*Arabidopsis thaliana* and *Oryza sativa*), fly (*Drosophila hydei*), zebrafish (*Danio rerio*) and mouse (*Mus musculus*). We used the complete AbRiDGed pipeline on the relatively complex genomes of mouse, zebrafish and rice, while for the smaller genomes of fungi, *D. hydei* and *A. thaliana*, we applied a simplified version that lacked the L2 libraries (see Methods for details). Overall, we generated 147,314 candidates among the 7 species, of which we then tested a very small fraction, 230 candidates (0.16%), using the biochemical assay to provide a proof-of-principle answer to the aforementioned question. As the result, we confirmed that 54/230 (23.5%) candidates represented true self-cleaving ribozymes (Figures 2, S7 and S8, and Table S5), including 4 previously predicted but unconfirmed HDV ribozyme variants from *A. nidulans*^17^. For comparison, only 6 ribozymes have been experimentally confirmed previously in these 7 species: the HH9, CPEB3, CLEC2d and CLEC2e ribozymes in mouse^18–20^, and Osa-1-2 and Osa-1-7 in rice^21^. Thus, considering the very small fraction of candidates tested, multiple undiscovered ribozymes do exist in different eukaryotic species.

**Figure 2.**
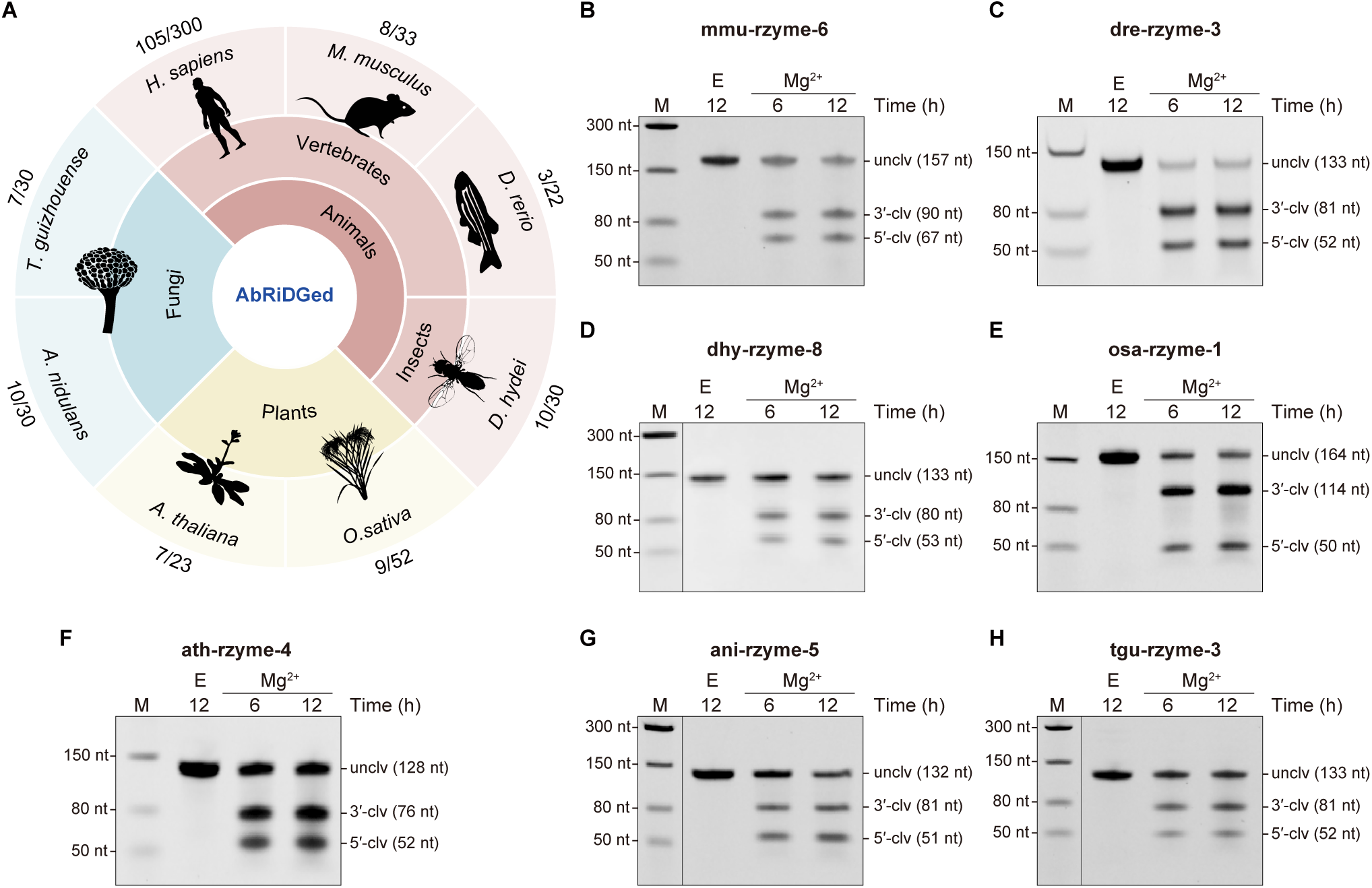
Discovery of novel self-cleaving ribozymes in multiple eukaryotic genomes via AbRiDGed. (A) The species used for the ribozyme discovery in this study and their classifications. The numbers of identified novel ribozymes and tested candidates for each species are annotated in the outer ring. (B–H) Representative examples of validation gels confirming the self-cleaving activity of the novel ribozymes identified from *M. musculus* (B), *D. rerio* (C), *D. hydei* (D), *O.sativa* (E), *A. thaliana* (F), *A. nidulans* (G) and *T. guizhouense* (H). The validation gel annotations are as described in the legend to Figures 1B and 1C. See Figure S8 for the 2 replicates of the validation gels for all 54 novel ribozymes from the 7 non-human species.

### Most of the newly discovered ribozymes represent unique members of novel classes

To investigate the sequence-structure relationships of the newly discovered self-cleaving ribozymes with the known ribozymes and among themselves, we performed a clustering analysis on their sequences together with the sequences representing previously identified or predicted ribozymes from the human and the 7 eukaryotes (Table S6). At the 40% sequence identity threshold, corresponding sequences of previously identified or predicted ribozymes were clustered together, and additionally, 2 clusters containing only 11 or 2 newly discovered human ribozymes were generated (Table S6). The remaining newly discovered ribozymes shared no significant sequence similarity with each other or the previously identified or predicted ribozymes. This result indicated that most of the newly discovered self-cleaving ribozymes represent distinct sequences.

We then performed an *in silico* search against all RNA families from the Rfam and ZWD databases^22,23^. In this search, all previously identified or predicted ribozymes (including the 4 *A. nidulans* HDV ribozyme variants) matched their respective classes (Table S7). However, none of the newly discovered ribozymes resulted in hits to any RNA family (Table S7), strongly suggesting that all these sequences represent novel self-cleaving ribozymes from yet undiscovered classes.

### Genomic distribution and evolutionary conservation of human ribozymes

Of the 109 human self-cleaving ribozymes (Figure 3A) — 105 novel ones discovered in this study and 4 identified previously — 30 (27.5%) and 12 (11.0%) mapped to the sense strands of annotated protein-coding and long non-coding (lnc) RNA genes, respectively (Figure 3B and Table S8). Similar to the known human self-cleaving ribozymes, HH9, HH10 and CPEB3, many (25) of the newly discovered ribozymes also mapped to the introns of protein-coding genes (Figure 3B). Strikingly, we found that the cleavage sites of 2 novel self-cleaving ribozymes (hsa-rzyme-50 and hsa-rzyme-14) were located in the coding sequences (CDSs) of genes encoding the insulin like growth factor 2 receptor (IGF2R) and laminin subunit gamma 1 (LAMC1), respectively (Figure 3C). To our knowledge, this is the first instance where self-cleaving ribozymes map to the CDSs in a eukaryotic genome.

**Figure 3.**
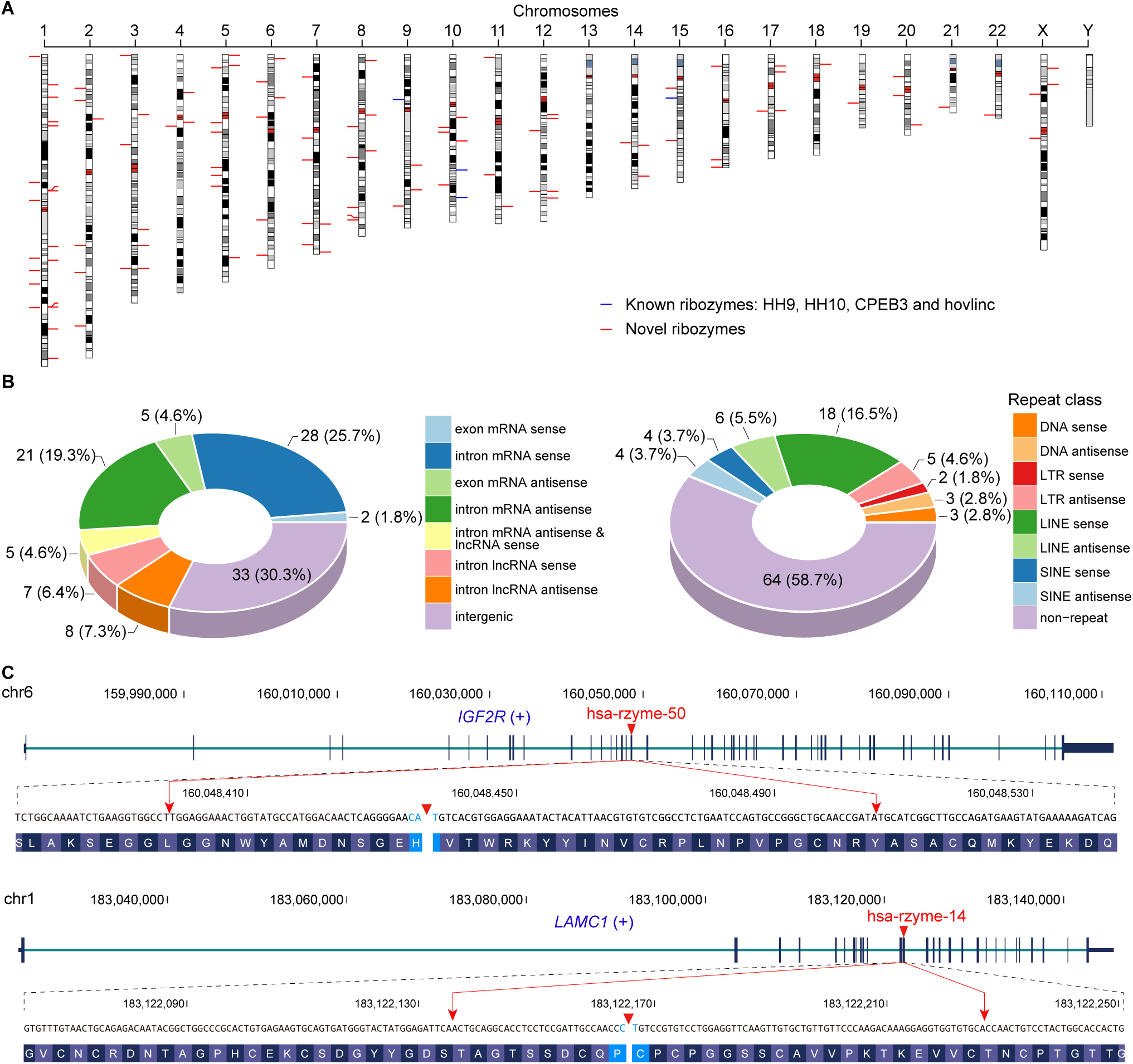
Genomic distribution of human self-cleaving ribozymes. (A) Genomic positions of the 4 known (blue) and the 105 novel human self-cleaving ribozymes (red) found in this study. Bars on the left and right of the chromosomes mark the ribozymes on the plus and minus strand of the genome, respectively. Note, the human chromosome Y was not covered by the screenings in this study, since it is absent from the genome of K562 cell line. (B) Distribution of the human ribozymes among the different genomic elements (left) and repeats (right). (C) The novel ribozymes hsa-rzyme-50 (top) and hsa-rzyme-14 (bottom) located in the protein-coding sequences of the *IGF2R* and *LAMC1* genes, respectively. The positions of the ribozymes in the exons are demarcated by the red arrows and the cleavage sites are marked by the red triangles.

Of the remaining 67 self-cleaving ribozymes, 34 mapped antisense to protein-coding or lncRNA genes and 33 were found outside of annotated transcripts (Figure 3B and Table S8). Since most of the human genome is transcribed and the current lncRNA annotations are incomplete, combined with the fact that antisense transcription is very common in mammalian genomes^24,25^, the 67 ribozymes are very likely transcribed as part of yet unannotated lncRNAs. For example, 7 of the novel ribozymes as well as the hovlinc ribozyme mapped to the class of very long intergenic non-coding (vlinc) RNAs^26^ that are not yet present in public databases (Table S9).

Overall, 45/109 (41.3%) of the human self-cleaving ribozymes were found in the repeats (Figure 3B and Table S10), of which majority (24/45 or 53.3%) mapped to the LINE retrotransposons, including all members of the aforementioned 2 clusters containing 11 or 2 newly discovered ribozymes. Since the ribozymes belonging to the 11-member cluster located in the L1M1, L1MA1 or L1MA2 LINE-1 subfamilies^27^, while the ribozymes in the other 2-member cluster were from the L1MB5 and L1MB7 LINE-1 subfamilies (Table S10), we will refer to them as respectively the L1MA and L1MB clusters.

While ribozymes are often assumed to represent vestiges of the “RNA World”, the discovery of the recently emerged human hovlinc ribozyme has challenged this assumption and suggested that ribozymes can evolve *de novo* in complex organisms^5^. Consistent with these findings, the sequence identities between the sequences of the human ribozymes and their homologs across 179 species (Table S11) dropped significantly outside of primates (Figure S9A and Table S12). In fact, for most of the ribozymes, the sequence identities dropped markedly in prosimians and their homologs could not even be found outside of mammals (Figure S9 and Table S12).

The activity of the hovlinc ribozyme emerged in the hominid lineage ∼13–10 million years ago (Ma) while being subsequently lost in the gorillas due to a single nucleotide change^5^. In depth analysis of evolutionary histories of 7 novel ribozymes in primates were also consistent with these observations. First, while active in the most closely related species, no active homologs were found in prosimians for these 7 ribozymes (Figures S9B and S10, and Table S12). Second, for each of the 7 ribozymes, we also identified single nucleotide variants that totally or almost totally abolished the self-cleavage activity in a primate homolog (Figure S11 and Table S13). This is similar to many other examples of known self-cleaving ribozymes where single substitutions drastically affect activity via perturbing the RNA folding or altering the crucial nucleotide bases involved in the catalysis^5,28–31^.

By performing a more extended test of the hsa-rzyme-97 homologs from 53 species in 9 mammalian orders, the self-cleavage activity was found only in 4 primates — chimpanzee, bonobo, gibbon and Sumatran orangutan, indicating that its activity was gained ∼25–18 Ma and then lost in the gorillas (Figures S9B and S10). However, while consistent with the recent occurrence of novel ribozymes in the human lineage, these results do not exclude independent emergence of some ribozymes in certain distant species or their deep evolutionary conservation.

### Human LINE-1 elements host multiple novel self-cleaving ribozymes

Considering that thousands of LINE-1 elements exist in the human genome, the discovery of the L1MA and L1MB ribozyme clusters suggested that many additional similar self-cleaving ribozymes might also exist, prompting us to analyze the L1MA ribozymes in more detail. We first confirmed the presence of the self-cleavage activity in a 76-nt region for all the L1MA ribozymes (Figure 4A). The multiple sequence alignment (MSA) analysis of the 11 ribozyme-containing LINE-1 elements and a full-length LINE-1 element showed that the L1MA ribozymes mapped to the LINE-1 open reading frame 2 (ORF2; Figure 4B). Subsequently, we used the MSA based on the sequences of all the 11 members of the cluster to generate the consensus sequence and predict the RNA secondary structure of the L1MA ribozymes (Figure 4C and 4D). The MSA and RNA secondary structure were used to build a covariance model which was then employed to conduct a search for homologs of the L1MA ribozymes in the human genome using the Infernal software^32^. This search resulted in 3449 hits with E-values <0.01 (Table S14). We then confirmed the self-cleavage activity of the top hit (chrX:78256308-78256233, minus strand, E-value = 3.10E−20) which was also located in an L1MA2 LINE-1 element (Figures 4A and 4B). This result indicated that additional L1MA ribozymes exist in the human LINE-1 elements and are yet to be discovered.

**Figure 4.**
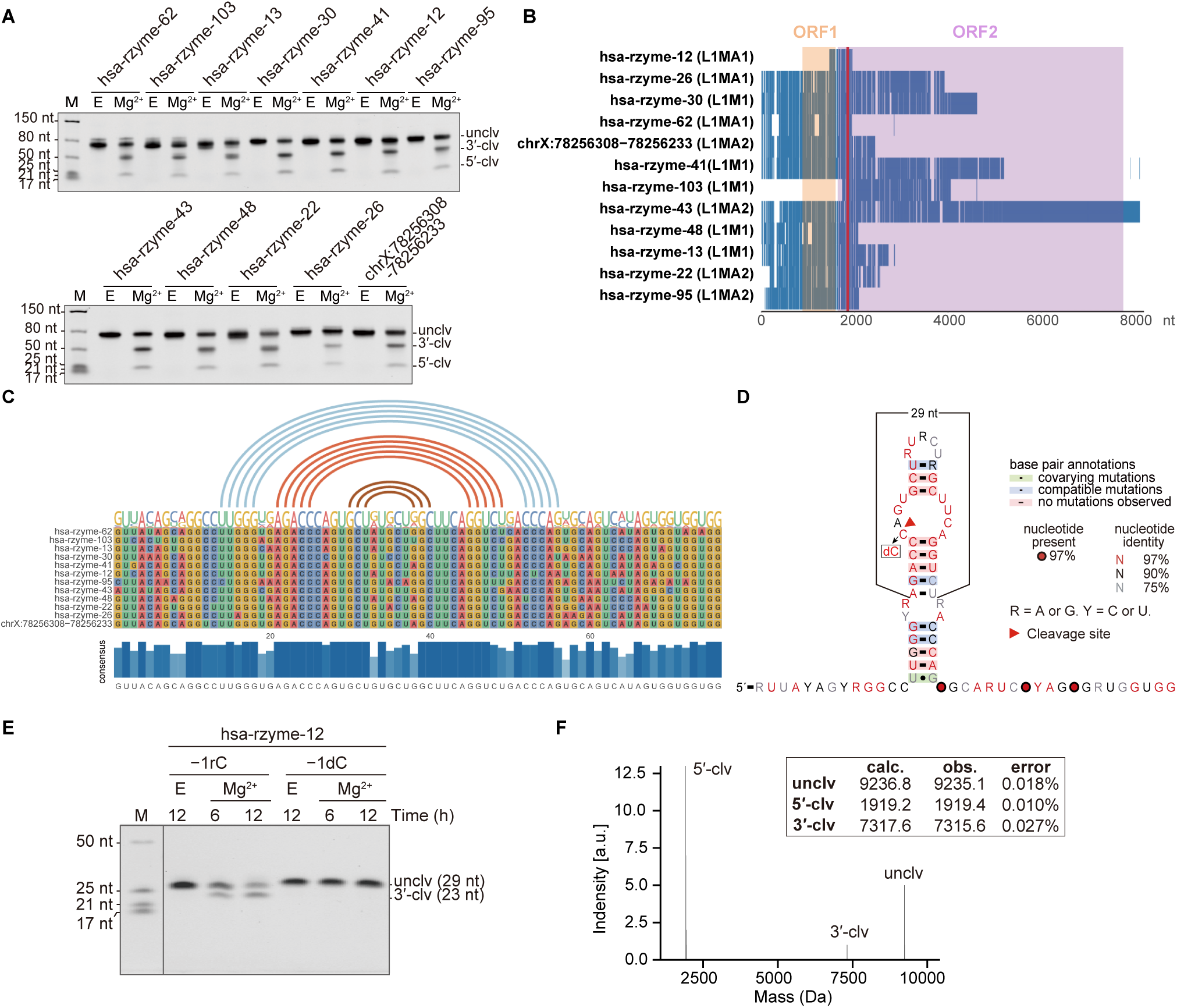
The L1MA class of ribozymes in human LINE-1 elements. (A) Self-cleaving activities of the L1MA ribozymes (6-hour cleavage reactions) including the 11 discovered via AbRiDGed and the one (chrX:78256308-78256233) found in the bioinformatic search. (B) The locations of the L1MA ribozymes in the corresponding LINE-1 elements and relative to the ORF1 (orange) and ORF2 (purple). The cleavage sites are indicated by the red line. (C) The multiple sequence alignment analysis of the L1MA ribozymes. The predicted base pairings are indicated by lines. (D) Predicted RNA secondary structure of the L1MA ribozymes. (E) The self-cleavage activity of the 29-nt L1MA sequences marked in (D) based on the hsa-rzyme-12 ribozyme (−1rC), and the inactivity of the hsa-rzyme-12 sequence possessing a cytidine to 2′-deoxycytidine substitution at the −1 position of the cleavage site (−1dC) as marked in (D). Note, the 6-nt upstream cleavage product is too short for detection. (F) Mass spectrum analysis of the cleavage reaction products of the 29-nt hsa-rzyme-12 ribozyme. (A and E) The annotations are as described in the legend to Figures 1B and 1C.

Based on the predicted RNA secondary structure, we further delimited the active region of the L1MA ribozymes to a 29-nt sequence (Figures 4D and 4E). The RNA secondary structure of the minimal region consists of two stems separated by an internal loop containing the cleavage site at the 5′ side, and a terminal loop (Figure 4D). Mass spectrometry (MS) confirmed the expected cleavage site, as well as the 2′,3′-cyclic phosphate and 5′-OH termini of the respectively 3′ end of the upstream and 5′ end of the downstream cleavage products (Figure 4F). Moreover, an analog molecule of the 29-nt hsa-rzyme-12 ribozyme lacking the 2′ oxygen of the cytidine upstream of the cleavage site showed no cleavage activity (Figures 4D and 4E). These results demonstrated that the cleavage reaction of this ribozyme happens through an internal phosphoester transfer mechanism commonly exploited by known self-cleaving ribozymes^12,13^.

### Activities of self-cleaving ribozymes vary over a wide range

The above results demonstrated that our findings greatly expanded not only the number but also the classes of discovered self-cleaving ribozymes. This enabled us to further investigate their catalytic properties on the vast collection of self-cleaving ribozymes. We measured the first-order rate constants (*k*_obs_) of the self-cleavage reactions for all the novel ribozymes found in this study and 14 sequences from 8 known ribozyme classes (Table S15). The experiments were conducted under two cleavage conditions — pH 7.5 and 10 mM Mg^2+^ with and without 150 mM K^+^, since monovalent cations can inhibit activities of some ribozymes by competing with divalent cations while enhancing the activities of others^33^. The *k*_obs_ measurements were performed as illustrated in Figure S12A for the hsa-rzyme-97 ribozyme.

We obtained the *k*_obs_ for 167/169 and 130/169 ribozymes in the conditions with and without K^+^, respectively (Table S16). Strikingly, we found that 39 ribozymes (all from the newly discovered ones) exhibited no measurable activity under the no K^+^ condition (Table S16). To our knowledge, this is the first description of self-cleaving ribozymes that appear to have a strict requirement for a monovalent ion in addition to a divalent ion for activity under such a condition. Only 2 ribozymes (ath-rzyme-2 and hsa-rzyme-53) were inactive in both conditions (Table S16), presumably because they require higher pH for the activity. The *k*_obs_ of self-cleaving ribozymes fell into wide but similar ranges: 81.46–0.01 h^−1^ and 80.88–0.007 h^−1^ under the respectively plus and minus K^+^ conditions (Figure 5A and Table S16). While the majority of the novel ribozymes represented the ribozyme classes with relatively low *k*_obs_, an appreciable fraction had *k*_obs_ within the range of those measured for the members of known ribozyme classes (Figure 5A and Table S16). The relatively low activity might be a reason that they were not discovered before, and also indicated the high sensitivity of the AbRiDGed method for ribozyme discovery.

**Figure 5.**
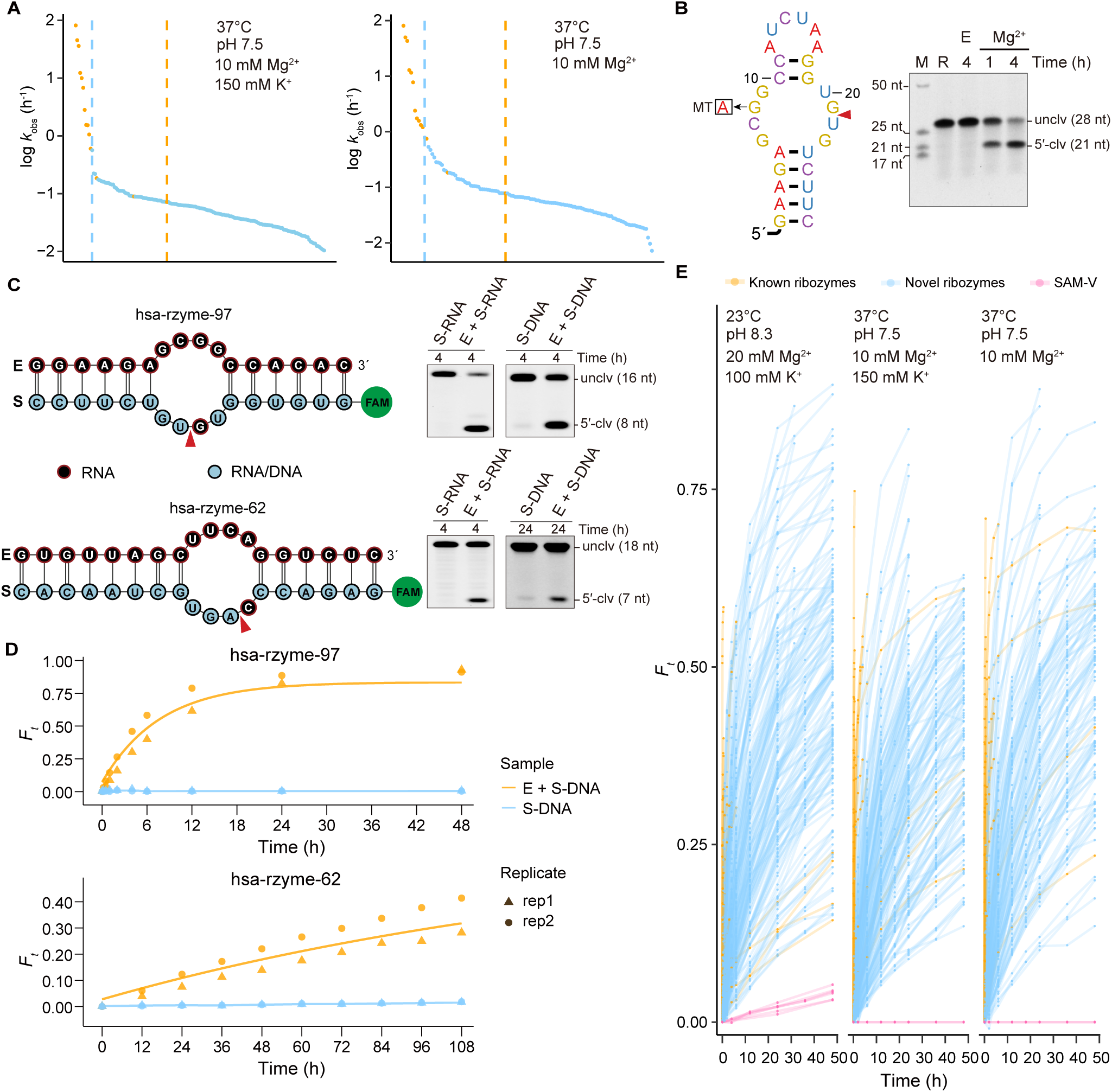
Biochemical analysis of novel self-cleaving ribozymes. (A) The *k*_obs_ (log_10_ values, Y-axes) of the known (orange dots) and novel (blue dots) self-cleaving ribozymes under the 2 indicated conditions. The dashed lines indicate the minimum and maximum *k*_obs_ values for the known (orange) and novel (blue) ribozymes, respectively. (B) Left, predicted RNA secondary structure of the 28-nt hsa-rzyme-97 sequence. “MT” denotes the G>A substitution in gorilla (Figure S11). Right, validation of the self-cleavage activity of the sequence. Note, the 7-nt downstream cleavage product is too short for detection. The “R” lane contains the intact 28-nt RNAs as the size reference. (C) The *trans*-cleavage of hsa-rzyme-97 and hsa-rzyme-62 ribozymes. Left, the *trans*-acting form of each ribozyme. Note, the substrates consist either entirely of RNA (S-RNA) or contain only one RNA nucleotide before the cleavage site (S-DNA) as indicated. Right, *trans*-cleavage reactions of either S-RNA or S-DNA substrates with and without the corresponding enzymes. (D) The cleavage fractions (Y-axes) of the S-DNA substrates with and without the corresponding enzymes in trans versus the reaction time (X-axes). Each reaction was performed in 50 mM Tris-HCl (pH 8.3), 100 mM KCl and 20 mM MgCl_2_ at 23°C for 10 ng substrate with or without 100 ng enzyme. (E) The cleavage fractions (Y-axis) of the active known (orange) and novel (blue) ribozymes as well as the SAM-V sequence (magenta) are plotted versus the reaction time (X-axis; 0–48 hours). (B and C) The gel annotations are as described in the legend to Figures 1B and 1C. The red triangles mark the cleavage sites.

### Biochemical properties of novel ribozymes

To understand the biochemical properties of the novel ribozymes, we conducted a detailed study for the hsa-rzyme-97 ribozyme (Figure S12). The activity of this ribozyme increased log-linearly with increasing pH (Figure S12B) and linearly with increasing Mg^2+^ concentration (Figure S12C). The calculated p*K*_a_ of this ribozyme was 9.46. Like other known self-cleaving ribozymes, the hsa-rzyme-97 required divalent cations, such as Mg^2+^, Mn^2+^, Ca^2+^ or Co^2+^, for cleavage (Figure S12D). The hsa-rzyme-97 ribozyme was inactive in cobalt (III) hexammine (Co(NH_3_)_6_^3+^), suggesting that the inner-sphere interaction with the divalent cation mediated by water molecules is important for the activity (Figure S12D). This ribozyme was also inactive in high concentrations of monovalent salt LiCl (Figure S12D). Altogether, hsa-rzyme-97 possessed a unique combination of biochemical properties compared with the known ribozymes.

We could localize the hsa-rzyme-97 ribozyme activity to a 28-nt sequence consisting of two helices separated by an internal loop, and a terminal loop (Figure 5B). The internal loop contained the cleavage site at the 3′ side and the single G>A substitution that inactivated the gorilla homolog (Figure 5B and Table S13). The location of the cleavage site as well as the presence of the 2′,3′-cyclic phosphate and 5′-OH termini on the cleavage products were also confirmed by the MS (Figure S12E). The predicted RNA secondary structure of the hsa-rzyme-97 ribozyme shared certain similarities with that of the L1MA ribozymes; however, the two classes of ribozymes shared no significant sequence similarity and had different locations of the cleavage sites (Figures 4D and 5B).

To further investigate the catalytic mechanisms of these 2 classes of ribozymes, we split each sequence into 2 parts — the enzyme (E) and the substrate (S) (Figure 5C). As indicated in Figure 5C, significant cleavage of the S sequences was only observed after the addition of the corresponding E part for each ribozyme showing that the ribozymes can cleave in *trans*, and thus confirming the RNA secondary structure predictions. This assay further enabled us to investigate the 2 ribozymes’ catalytic rate enhancement (CRE) which is an innate property of any enzyme and, in the case of a ribozyme, represents the increase in the rate of the transesterification reaction caused by the ribozyme catalysis. We tested the *trans*-cleavage of the 2 ribozymes with the S molecules consisting of DNAs except for the nucleotides immediately preceding the cleavage sites, while the E parts remained all RNAs (Figure 5C). This excludes the effect of the local RNA folding on the stability of a phosphodiester bond^12,34^. While the cleavage activities of such molecules were reduced by an order of magnitude, under various reaction conditions, the cleavage fractions of the S molecules were always significantly higher in the presence of the E parts than in their absence (Figure 5C). Therefore, both novel ribozymes significantly enhance the rates of the transesterification reaction, although the exact enhancement fold cannot be estimated due to the extremely slow reaction of the S molecules in the absence of the E parts.

In addition to the proper RNA folding facilitating favorable in-line geometry required for the transition state, all known self-cleaving ribozymes require additional interactions around the cleavage site to achieve the observed rates of the catalysis^12,34^. To explore whether a similar situation might exist for the novel self-cleaving ribozymes, we compared the cleavage rates of all ribozymes studied in this work to that of the S-adenosylmethionine (SAM) riboswitch SAM-V. Breaker and co-workers^35^ found that a strong site of cleavage was created in the SAM-V riboswitch upon the binding of SAM. As later determined based on the crystal structure, ligand binding creates a major-groove triple helix, in the course of which a cytosine is bulged out with an excellent geometry for in-line attack by the 2′-hydroxyl group^36^. the SAM-V riboswitch thus provides a good system to evaluate the effect of in-line geometry on phosphodiester bond stability in the absence of other factors.

While the expected site-specific cleavage of the SAM-V riboswitch was reproduced under the condition previously used to detect it^35^, we could not reliably measure its exact *k*_obs_ due to the very slow rate of this reaction (Figure S13A). However, according to the fractions of cleaved molecules, all the self-cleaving ribozymes showed significantly faster cleavage than the SAM-V riboswitch under this condition (Figures 5E and S13B). Moreover, under the above 2 conditions which we used to measure the activities of self-cleaving ribozymes, the SAM-V riboswitch showed no detectable cleavage (Figures 5E and S13C), presumably because of the lower pH and Mg^2+^ concentration (Methods). Altogether, these results indicated that the cleavage rates of these self-cleaving ribozymes are far beyond what could be achieved purely by the RNA folding and additional catalytic strategies do contribute significantly to their activities.

All the results above demonstrated that the novel self-cleaving ribozymes share significant similarities in the catalytical and biochemical properties with the previously identified ribozymes. Further understandings of the catalytic mechanisms of these novel self-cleaving ribozymes would require elucidation of their RNA structures at atomic level resolution.

### A novel noncanonical class of self-cleaving ribozymes abundant in all domains of life

In the aforementioned ribozyme validation efforts, we unexpectedly uncovered 15 additional ribozymes from multiple species with similar self-cleavage in either the presence or absence of Mg^2+^ as exemplified in Figure S14A (Table S17). We therefore hypothesized that these ribozymes do not require Mg^2+^ for activity and potentially need only the low concentration (150 mM) of K^+^ present in both conditions. Indeed, we found that each of these 15 ribozymes was comparably active in 150 mM of K^+^ as well as 5 other monovalent cations — Na^+^, Li^+^, Rb^+^, Cs^+^ and NH_4_^+^ (Figures 6A and S14B). This property was quite different from the previously characterized ribozymes — 10 ribozymes from 10 known classes were inactive in 5 out of the 6 cations at such a low concentration and only 4 of them were active in Li^+^ (Figures 6A and S15). Furthermore, the self-cleaving activity of the 15 ribozymes could all be localized to a very similar short stem-loop structure (Figure S16). All together, these results indicated that these ribozymes all belong to one unique novel class, named by us MINDS (Monovalent Ion Dependent Self-cleaving), with distinct noncanonical biochemical properties and RNA structure.

**Figure 6.**
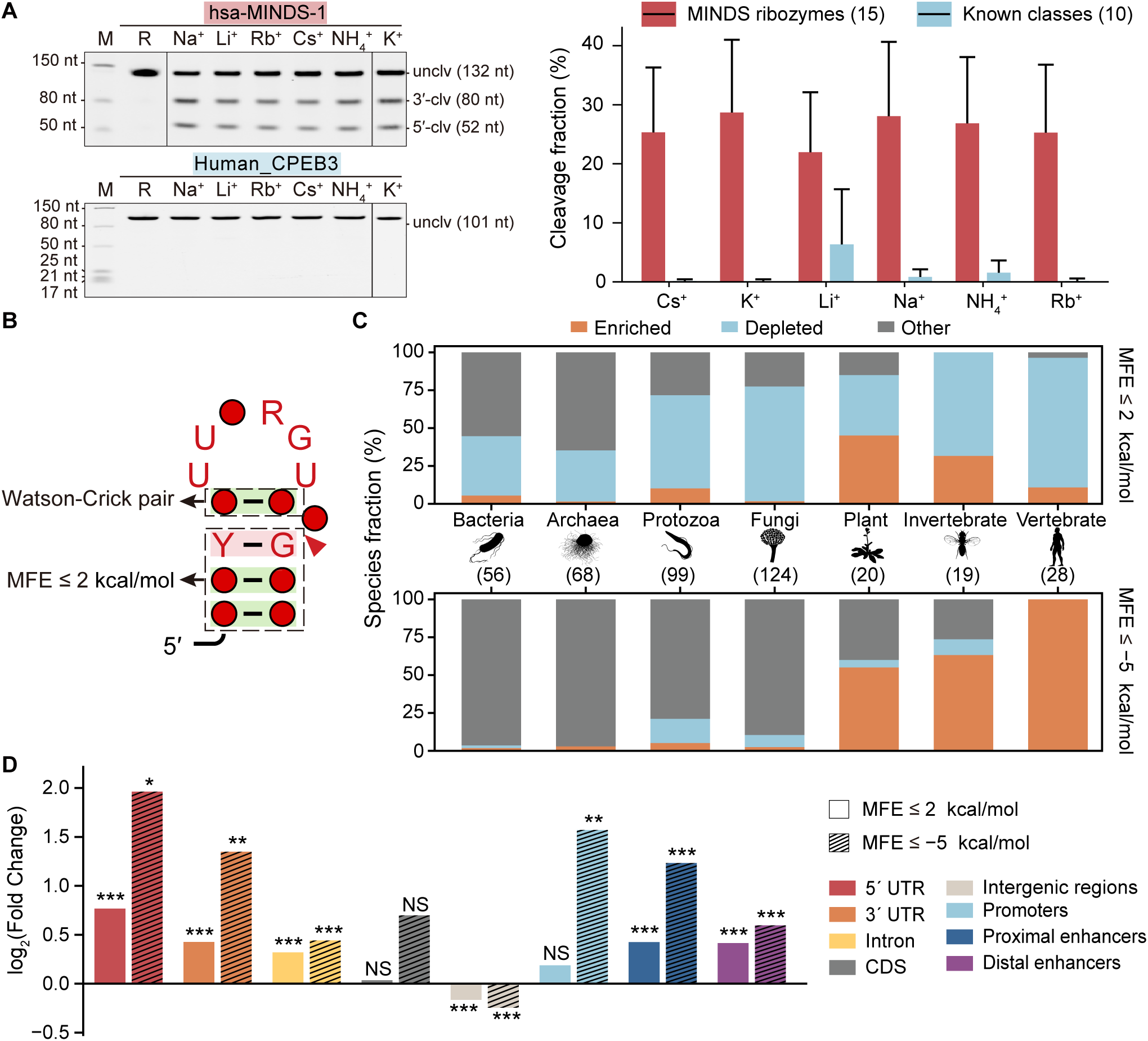
Properties and distribution of the MINDS ribozymes. (A) Left, self-cleaving activity of the hsa-MINDS-1 and the human CPEB3 ribozymes in a low concentration of various monovalent ions. The “M” lanes contain the size standards. The “R” lanes contain the IVT products of the ribozymes without any incubation. Right, the average cleavage fractions (Y-axis) of the 15 MINDS ribozymes and 10 different known classes of self-cleaving ribozymes (Tables S1 and S17) in various monovalent ions (X-axis). The error bars represent the standard deviations. The numbers in the parentheses are the numbers of ribozymes included in the test. The reactions were conducted in 150 mM of the indicated monovalent ions, 30 mM Tris-HCl (pH 8) for 12 h at 37°C. (B) Sequence-structure motif of the MINDS ribozymes, the annotations are described in Figure 4D. (C) The fractions of species (Y-axis) from 7 clades in the genomes of which the MINDS ribozymes are either significantly enriched, depleted or show no significant enrichment or depletion (“Other”) under the two MFE thresholds (X-axis). The numbers in the parentheses are the numbers of species from each clade included in the analysis. (D) The enrichment or depletion of the human MINDS ribozymes in various genomic elements. \**p* < 0.05, \*\**p* < 0.01, \*\*\**p* < 0.001, NS, non-significant.

Then, using the 25-nt sequence of the hsa-MINDS-1 ribozyme we further characterized the biochemical properties of this novel class of ribozyme (Figure S17). The expected cleavage site and the presence of the expected 2′,3′-cyclic phosphate and 5′-OH termini at the respectively upstream and downstream cleavage products were confirmed by MS (Figure S17A). The RNA to DNA single nucleotide substitution preceding the cleavage site abolished the activity consistent with the cleavage requiring nucleophilic attack on the 2′ OH group (Figure S17A). In addition to the 6 monovalent ions mentioned above, the MINDS ribozymes were also active in a wide array of cations: various divalent metal ions (Mg^2+^, Ca^2+^ and Mn^2+^, but not Co^2+^), Co(NH_3_)_6_^3+^ or amino groups (spermidine), as illustrated in Figure S17B. To our knowledge, no other characterized class of self-cleaving ribozymes exhibits such a combination of cation requirements, further supporting the uniqueness of the MINDS class. As shown in Figure S17C, the *k*_obs_ of the hsa-MINDS-1 ribozyme is 0.121 h^−1^ at pH 8, 150 mM K^+^ and 0.098 h^−1^ at pH 8, 6 mM Mg^2+^, while in the presence of both K^+^ and Mg^2+^ the *k*_obs_ is 0.207 h^−1^ (pH 8, 150 mM K^+^ and 6 mM Mg^2+^). Similar to other ribozymes found in this study, the *k*_obs_ of the MINDS ribozyme increased with the increase in cation concentration or pH (pKa = ∼8.7) (Figures S17D, S17E and S17F). The MINDS ribozymes showed significantly higher self-cleavage activity compared with the SAM-V riboswitch in all 3 conditions tested (Figure S17G).

In-depth analysis of self-cleavage activities of various synthetic sequence variants revealed a sequence-structure motif of MINDS ribozymes with the following 4 salient features: (1) a highly conserved 6-nt loop, (2) a single Watson-Crick base pair at the base of the loop, (3) a cleavage site located between a bulged base and a base-paired invariant G residue and (4) a relatively stable stem with a variable sequence (Figures 6B and S18). The minimal functional sequence of the MINDS ribozyme was only 15 nt, making it the shortest self-cleaving ribozyme discovered so far (Figures 6B and S18).

The activity of the MINDS ribozymes increased linearly with the stability of the stem-loop structure (Figure S18F), mostly contributed by the stem, as measured by the minimal free energy (MFE). Little activity was observed in sequences with MFE > 2 kcal/mol; therefore, we used the sequence-structure motif shown in Figure 6B with MFE ≤ 2 kcal/mol for all subsequent analyses of the MINDS ribozymes unless otherwise specified.

Through a bioinformatic analysis, we uncovered that the MINDS ribozyme exists in all domains of life — it was found in each of the tested 824 prokaryotic and eukaryotic species, as well as in 11/145 tested viral genomes (Table S18). Overall, ∼5 million MINDS ribozymes were found in these species (Table S18). We then explored whether the MINDS ribozymes were enriched or depleted in each genome sequence relative to expected by chance. Strikingly, we found the highly active subset of the MINDS ribozymes (MFE ≤ −5 kcal/mol) to be significantly enriched in all the 28 tested vertebrates from coelacanth to human (Figure 6C and Table S19). The enrichment was found only for the most active MINDS ribozymes — when considering all MINDS ribozymes identified with the MFE ≤ 2 kcal/mol threshold we did not observe such a pattern, in fact 24/28 of the tested vertebrate genomes showed significant depletion of these ribozymes (Figure 6C and Table S19). No significant trend of enrichment or depletion was observed in archaea, bacteria, fungi, protozoa, invertebrate or plants at either threshold of MFE (Figure S19 and Table S19).

The intra-genomic distribution of the MINDS ribozymes was also non-random in complex organisms. The human MINDS ribozymes were enriched in introns, 5′ and 3′ untranslated regions (UTRs), but not the CDS, and were depleted in the intergenic space (Figure 6D). Furthermore, we observed significant overlap between the MINDS ribozymes and the human *cis*-regulatory genomic elements — promoters and enhancers (Figure 6D), which are well-known to associate with multiple prominent types of transcripts^24,25^. These patterns also became more pronounced for the highly active subset of these ribozymes (Figure 6D). Altogether, the above results indicate that in complex organisms the MINDS ribozymes are under an evolutionary pressure selecting for highly active entities located at specific functional elements, strongly suggesting that they play specific functions in those organisms as has been shown previously for the polyproline motifs involved in the regulation of the translation elongation^37^.

### *In vivo* activity and physiological relevance of the novel ribozymes

Ostensibly, novel self-cleaving ribozymes could function inside a cell via two non-mutually exclusive mechanisms: (1) creation of novel functional transcripts from the longer precursors and (2) regulating the expression of the host transcripts. To further investigate these possibilities, we focused on the hsa-rzyme-14 ribozyme located in an exon of the human *LAMC1* gene (Figure 3C). First, we explored *in vivo* existence of novel transcripts derived from its cleavage by detecting the 5′ termini of the downstream cleavage products utilizing the 5′ RACE (Rapid Amplification of cDNA Ends) technology in combination with high-throughput sequencing^38,39^ (Figure 7A). For the human CPEB3 ribozyme, the 5′ RACE indeed revealed a strong signal coinciding with its cleavage site in human brain RNAs (Figure 7B), consistent with its *in vivo* activity generating relatively stable cleavage products and implicated in the control of expression of the host CPEB3 transcript in neuronal cells^40^. Similarly, the 5′ RACE for the hsa-rzyme-14 ribozyme also detected a strong signal in the human 293FT cells shortly adjacent to its cleavage site (Figure 7B). The imprecise correspondence between the RACE signal and the cleavage site might be due to the certain imprecision of the method itself, as well as potential degradation of the cleavage-generated transcripts from the 5′ termini in the cell. Overall, the above results confirmed the *in vivo* cleavage activity of the hsa-rzyme-14 ribozyme in its native sequence context — the LAMC1 transcript — as well as the presence of a novel transcript generated by the ribozyme activity.

**Figure 7.**
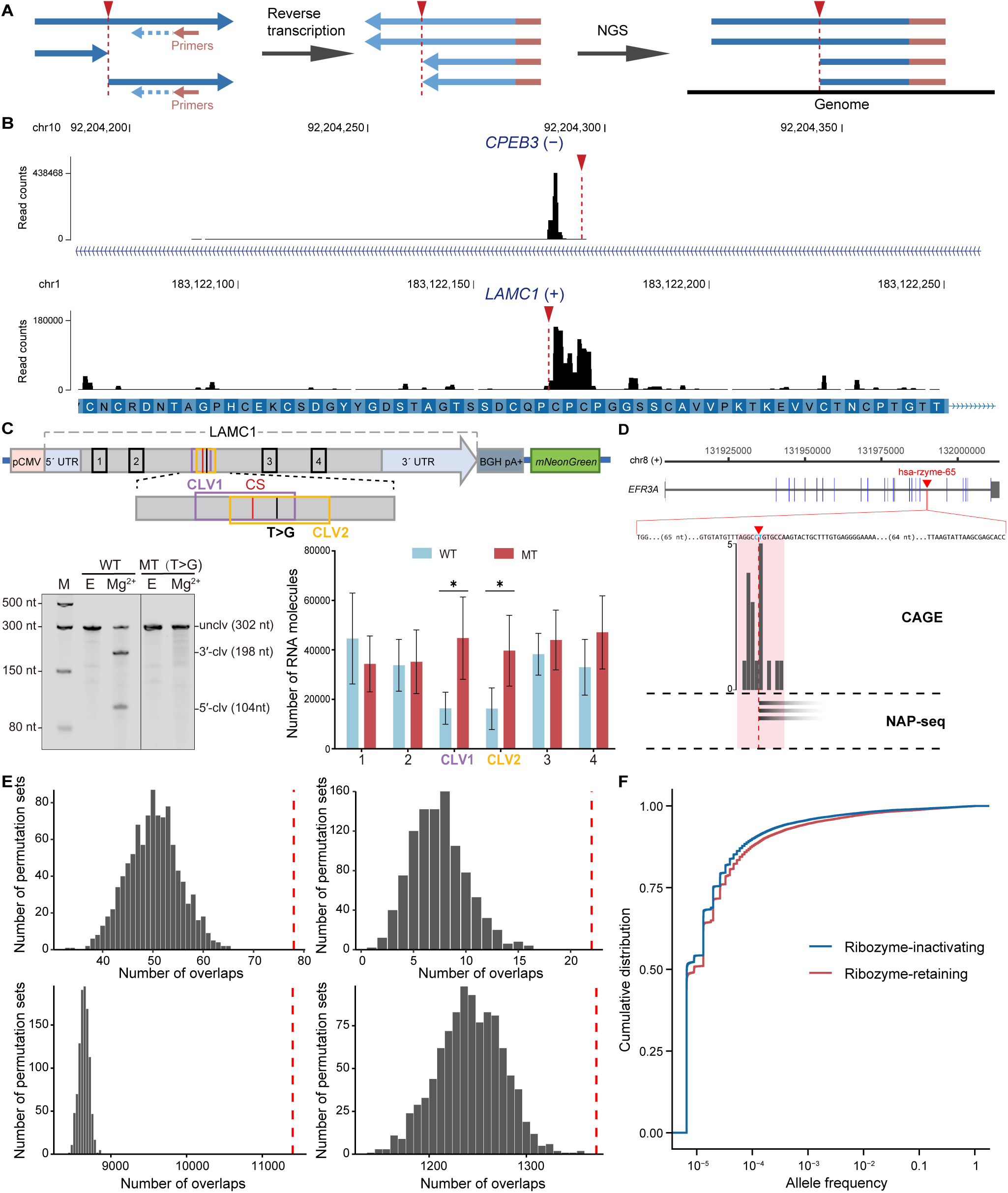
Novel human self-cleaving ribozymes are active and functional *in vivo*. (A) Schematics of 5′ RACE in combination with NGS to detect 5′ termini of the downstream cleavage products. (B) Wiggle plots of the 5′ RACE for the CPEB3 (top) and hsa-rzyme-14 (bottom) ribozymes. Note, a Cytosine-homopolymer immediately downstream the CPEB3 ribozyme cleavage site prevents precise mapping of the RACE signal due to a polyC-tailing step employed in the RACE procedure. (C) Top, schematics of the constructs that overexpress the LAMC1 mRNAs containing either the wild-type (WT) hsa-rzyme-14 ribozyme or the T>G substitution (MT). Bottom-left, inactivation of the hsa-rzyme-14 ribozyme by the T>G substitution. The annotations are as described in the legend to Figures 1B and 1C. Bottom-right, the RNA molecule numbers measured by RT-qPCR using various primer pairs (X-axis) shown in the top diagram. Note, the CLV1 and CLV2 primer pairs measure only the uncleaved LAMC1 mRNA, while the pairs 1–4 measure both cleaved and uncleaved transcripts. The error bars represent standard deviations of 4 biological replicates. \**p* < 0.05 (2-sided Student’s *t*-test). (D) Overlap between the hsa-rzyme-65 ribozyme cleavage site and transcript boundaries datasets. Only the data within ±5 nt of the 5′-end of the downstream cleavage product are shown (pink box). (E) The permutation test results for overlaps between the Mg^2+^-dependent (left) and the MINDS ribozyme cleavage sites (right) with the CAGE (top) and NAP-seq (bottom) datasets. Each histogram shows the distribution of the numbers of overlaps between the 1000 sets of random genomic positions and the transcript boundaries. The actual numbers of overlaps are marked by the red dashed lines. (F) The cumulative distributions of the allele frequency of the human genomic variants that either inactivate or retain the human MINDS ribozyme activity.

We then investigated whether the *in vivo* activity of the hsa-rzyme-14 ribozyme could affect the expression level of the host LAMC1 transcript. We transiently over-expressed the entire ∼7.9 kb LAMC1 mature mRNA with either the active hsa-rzyme-14 ribozyme (WT) or a mutated inactive version (MT) in human 293FT cells. The inactive version of the ribozyme was generated by a synonymous T>G substitution at the same position as the T>C substitution shown in Figure S11 (U48C in the hsa-rzyme-14 RNA). This synonymous T>G substitution minimizes the differences in the codon frequencies, and totally abolished the ribozyme activity (Figure 7C). Following the transfection of the two plasmids expressing the WT and MT LAMC1 mRNAs, the levels of the LAMC1 transcripts were measured using RT-qPCR with different primer pairs (Figure 7C). The total amount of the cleaved and uncleaved LAMC1 mRNAs was measured by 4 independent primer pairs up-or downstream of the hsa-rzyme-14 ribozyme cleavage site, while other two independent primer pairs spanning the cleavage site quantified only the uncleaved LAMC1 mRNA (Figure 7C). Furthermore, the expression level of the *mNeonGreen* gene was used for the normalization for the transfection efficiencies.

In cells transfected with the WT plasmid, the uncleaved LAMC1 mRNA represented on average ∼40% of the total LAMC1 transcript population (Figure 7C). In contrast, this fraction was restored to ∼100% in the cells expressing the MT sequence while the overall levels of the LAMC1 transcripts remained similar (Figure 7C). While again confirming the *in vivo* activity of the hsa-rzyme-14 in the context of the native ∼7.9 kb LAMC1 mRNA, these results indicated that the *in vivo* cleavage of the ribozyme leads to a considerable change in the expression level of the full-length LAMC1 transcript and results in novel transcripts stably present in the cell.

To study the *in vivo* activity of all the novel ribozymes, we took advantage of 2 public human high-throughput datasets that mapped transcript boundaries with nucleotide-level precision in a large and diverse collection of samples. The FANTOM consortium CAGE (cap analysis gene expression) dataset provides the positions of the 5′ ends of capped transcripts^41^. While the self-cleavage of ribozymes generates 5′-OH termini, theoretically they can be converted into 5′-monophosphates via a 5′ RNA kinase activity present in mammalian cells^42^ to which the cap structure can then be added^43–45^. In contrast, the NAP-seq dataset contains the positions of the 5′ and 3′ ends of non-capped human transcripts with various types of termini, including the 5′-OH and 2′,3′-cyclic phosphate that could be generated by the self-cleavage of ribozymes^46^.

The analysis of the 109 human ribozymes requiring Mg^2+^ for their activity, including 105 novel and 4 known ones, and 19,221 human MINDS ribozymes revealed that the cleavage sites of 78 (71.56%) and 22 (20.18%) Mg^2+^-dependent ribozymes overlapped the transcript boundaries from respectively the CAGE and NAP-seq datasets within ±5 nt genomic windows as exemplified in the Figures 7D (Table S20). Notably, the cleavage sites of the hsa-rzyme-14 and hsa-rzyme-50 ribozymes located in the human CDS overlapped with transcript boundaries in a large number of samples from both the CAGE and NAP-seq datasets. The corresponding numbers for the MINDS ribozymes were 11,401 (59.32%) and 1372 (7.14%) (Table S21). The overlap between the cleavage sites of both types of ribozymes and the transcript boundaries in either CAGE or NAP-seq dataset were statistically significant as shown by the permutation tests (Figure 7E, and Tables S20 and S21).

Finally, to directly evaluate the effect of loss of ribozyme activity on fitness, we compared the allele frequencies (AF) of short human genomic variants that either inactivate or have no effect on the ribozyme activities. We classified the variants from the Genome Aggregation Database (gnomAD, v4.1.0)^47^ that map to the MINDS ribozyme into “ribozyme-inactivating” and “ribozyme-retaining” based on their predicted effect on the self-cleaving activity (Table S22). We found that the ribozyme-inactivating variants tend to have significantly lower AF than the ribozyme-retaining variants (Figure 7F; K-S test, *p* < 2.2E−16). This result suggests that the MINDS ribozymes are active *in vivo* and the variants that inactivate the ribozymes are depleted in the population due to their deleterious effects. Indeed, we found many ribozyme-inactivating variants that are associated with clinical traits, according to the ClinVar^48^ and GWAS catalog^49^ databases (Table S22). Taken together, these results suggest that the human self-cleaving ribozymes are active *in vivo,* and their activities contribute to the transcriptome diversity and regulation of gene expression and have physiological and clinical significance.

### Novel ribozymes can be used for direct RNA detection

The importance of direct detection of RNA viruses, various alternative transcript isoforms and short RNAs, such as miRNAs, in diagnostics led to a significant interest in the development of methods that do not rely on reverse transcription or PCR. For example, developing such approaches based on the CRISPR suite of tools has drawn a lot of attention^50^. However, the repertoire of such methods is still limited, and they have not yet gained widespread practical application. In this regard, direct RNA detection methods based on ribozymes have at least two major advantages: (1) they do not require additional protein components such as Cas12 or Cas13 that are essential for all CRISPR-based methods; and (2) they can detect short RNAs which are problematic for the traditional reverse transcription and PCR-based assays, yet highly relevant for detecting functional short RNAs or degraded transcripts.

Therefore, in this work we explored the potential of novel ribozymes as the basis for direct RNA detection techniques. We took advantage of the *trans*-acting form of the hsa-rzyme-97 ribozyme, in which we further localized the catalytic core to 12 nucleotides consisting of 5′-UGUGUC on the S strand and 5′-GAGCGG on the E strand (Figure 8A). Sequences flanking the core are variable as long as they could mediate the base pairing of the E and S strands (Figure 8A). Therefore, the cleavage of designed S molecules could serve as the basis for detecting the presence of any specific target RNAs containing the 5′-GAGCGG motif and acting as the E part (Figure 8A).

**Figure 8.**
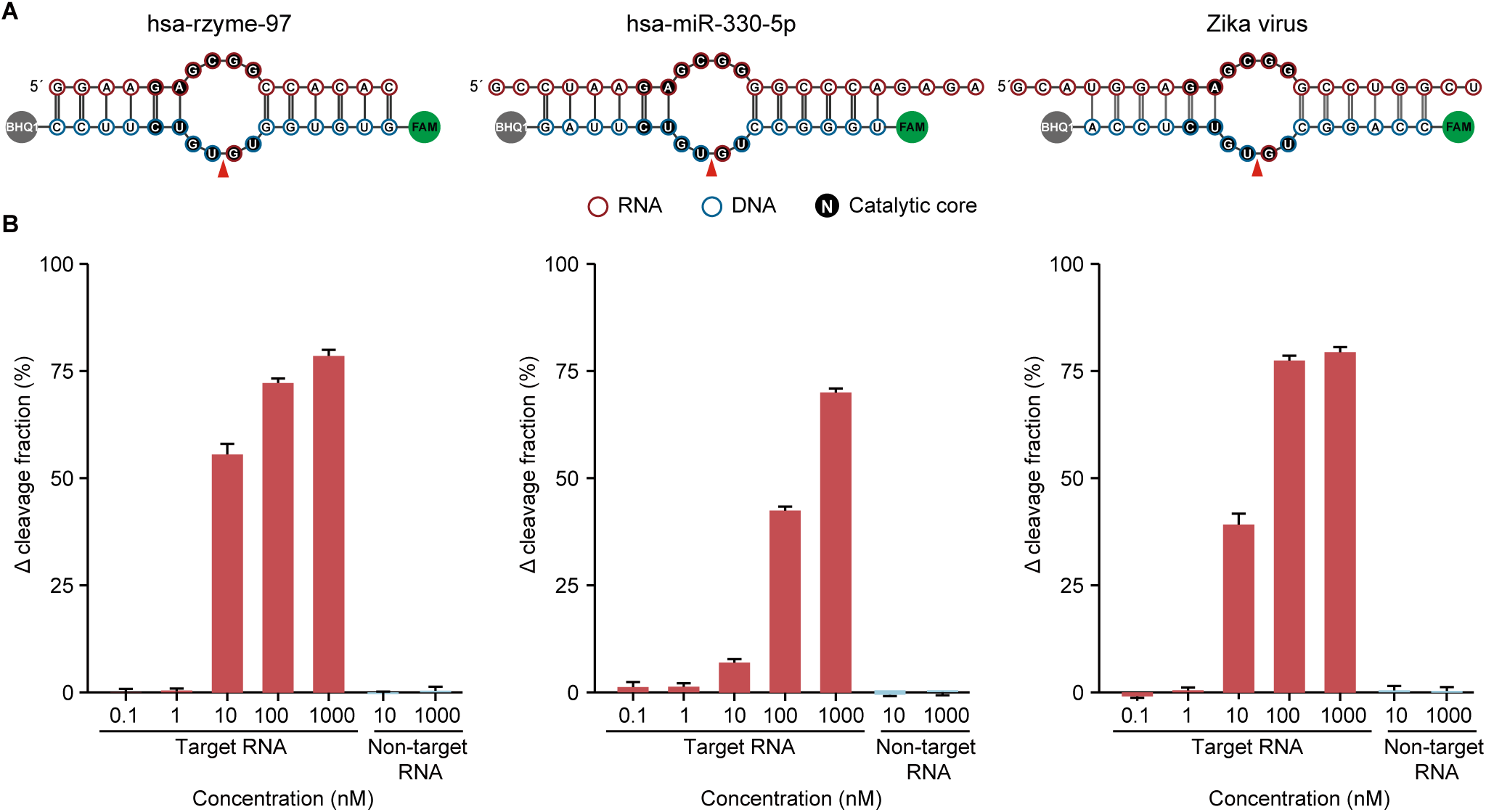
Direct RNA detection using a novel human ribozyme. (A) The secondary structures and sequences of each indicated target sequence and the corresponding substrate. (B) The cleavage fractions (Y-axes) of the substrate for each concentration (X-axes) of the corresponding target sequence and the non-target control. The cleavage fractions of the substrate-only reactions containing no target sequences performed in parallel were subtracted. Each reaction was performed in 30 mM Tris-HCl (pH 9.0) and 20 mM Mg^2+^ with 10 ng of a substrate for 48 h at 37°C and analyzed by denaturing PAGE. The error bars represent standard deviations of 3 replicates.

We verified the sensitivity of this method with variable (0.1–1000 nM) concentrations of the E molecules. In the assays, the S molecule contained BHQ1 at its 3′ end to decrease the background (Figure 8A). We found the detection limit in such method to be in the range of 1–10 nM in gel-based assays (Figure 8B), which was comparable to that reported for the CRISPR-based methods without pre-amplification (0.1–10 nM)^51^. Using the same principle, we could also design and achieve detections for the other 2 targets with a similar sensitivity — one from Zika virus genome and one representing the human miRNA hsa-miR-330-5p (Figure 8A). In addition, the specificity of the detection achieved by the designed base pairings between the S and E parts at the sequences flanking the catalytic core was also confirmed experimentally — unmatched S and E parts resulted in no detection (Figure 8B).

These results provide a proof-of-principle demonstration for the potential of the novel ribozymes in direct RNA detection. The examples shown here based on the catalytic core of the hsa-rzyme-97 with a relatively simple design and assay already achieved decent sensitivity and specificity, which could be further improved by identification of novel catalytic cores with higher activities and wider arrays of target sequences, and more sophisticated sequence design and detection methodologies beyond the basic gel-based assay.

## Discussion

Non-protein-coding functions of RNAs have been shown to be crucial in many biological processes, and the ever-expanding array of roles played by RNA in the cell suggests that some still await discovery. One such function, RNA catalysis, has been known for more than 40 years. However, so far, few classes of naturally occurring catalytic RNAs in general, and self-cleaving ribozymes in particular, have been found in eukaryotes, especially mammals. While some classes of self-cleaving ribozymes, such as the HDV and hammerhead, are present in all clades of life, each of them is represented by variations of the same sequence-structure pattern^52–55^. This led to the general dogma that nearly all RNA-based catalysis has been replaced by protein-based catalysis and become extremely rare in modern complex organisms, while the few remaining ribozymes represent the remnants of the “RNA world”^4,6,7^.

However, in this work, we showed that the complexity of “ribozyme-ome” is several orders of magnitude higher than previously estimated. For example, in the human genome, previously thought to contain only 4 self-cleaving ribozymes, we identified 105 novel Mg^2+^-dependent ribozymes that belong to 103 novel classes. Using structure-sequence properties of just one such class, we found additional 3449 hits, consistent with our estimates that ∼10K Mg^2+^-dependent ribozymes exist in the human genome and remain undiscovered. Furthermore, we discovered 9 human instances of another class of self-cleaving ribozymes, the MINDS ribozyme, which has a unique noncanonical ion dependence and exist ubiquitously and abundantly in all domains of life. Based on the structure-sequence motif of the MINDS ribozyme, we identified additional ∼48K entities in the human genome. Therefore, the total number of ribozymes in the human genome discovered in this study alone can be estimated to be in tens of thousands. Similar conclusion can be reached for other eukaryotic genomes tested in this work. Extending this analysis to all species tested, including prokaryotic and viral genomes, resulted in millions of novel self-cleaving ribozymes. Furthermore, this work focused only on one type of catalytic activity and limited number of reaction conditions, thus opening a real possibility that a much larger array of catalytically functional RNA sequences does exist.

We found that despite their relatively low *in vitro* activities, the novel ribozymes function at least in part by contributing to the diversity and complexity of the *in vivo* transcriptome. The very complex nature of transcriptome in mammals as well as other eukaryotes has been an active field of investigation in the post-genomic era^24,25,56–59^. In part, this complexity is characterized by the presence of many different RNA molecules *in vivo* apparently derived from the same transcript post-transcriptionally^43,58,60–62^. Such RNAs might possess various types of termini and play important roles in a variety of biological processes^43,46,60,61,63^. It has been generally assumed that these RNAs are generated by endonuclease cleavage^62^, however, the results presented here suggest that ribozyme cleavage can also contribute to their biogenesis. It is quite possible that both the original transcripts and post-transcriptionally processed RNA molecules have function, since both are frequently detected in the cell^43,58,62^, and the balance between them is important. In such cases, the complete and fast cleavage of the ribozyme-containing RNA molecules might not always be desired.

We also show that another function of self-cleaving ribozymes is to regulate the copy number of the corresponding RNA molecules, in which case the fast rate of cleavage might also not be universally desired since it would lead to quick depletion of the ribozyme-containing RNA molecules. In fact, considering that the average half-lives of mammalian RNAs range from ∼10 hours^64^ to >20 hours^65^ and some RNAs persist for years^66^, the self-cleavage rates observed in this study are quite consistent with physiological relevance of the novel ribozymes.

Besides contributing to the transcriptome diversity and regulation of gene expression, the prominence of catalytic RNAs has additional major implications. First of all, as exemplified by the apparently deleterious effect of sequence variants that inactivate MINDS ribozyme on fitness, the impact on the ribozyme activity might provide an additional explanation of how a sequence variation translates into a phenotype. A nucleotide change that significantly affects ribozyme activity could be a reason behind a phenotypic effect of intraspecies variation or an answer to the long-standing dilemma of how the very small difference between the genomes of closely related species, like human and chimpanzee, could result in significant biological and behavioral differences. It is conceivable that emergence, loss or fine-tune of the RNA catalytic activity could be achieved by very small sequence changes and thus might contribute significantly to the process of speciation.

Second, our results provide an additional mechanism for the phenotypic effects of synonymous substitutions that while previously widely assumed to be neutral are currently believed to have phenotypic effects^67^. As exemplified by the synonymous substitution in the LAMC1 ribozyme that upregulates the level of the full-length mRNA, other synonymous substitutions could very likely exert their effects by modulating ribozyme activities.

Third, the identification of multiple self-cleaving ribozymes in the still enigmatic lncRNAs, the so-called RNA “dark matter”^24,25^ which is especially prevalent in complex organisms^68^, suggests that these transcripts might function by carrying catalytic RNA domains. This notion is consistent with the inter and intra-genomic distribution of the MINDS ribozymes which supports the functionality of these catalytic RNAs in complex organisms. Fourth, our results suggest that naturally occurring RNAs catalyzing other reactions might also exist, which requires the development of novel high-throughput assays for their discovery.

The above functional scenarios would be very different from the functions of multiple known self-cleaving ribozymes. For example, many such ribozymes participate in the rolling circle replication of various simple (e.g., viral or viroid) genomes where a very high self-cleavage activity is required to cleave the oligomeric RNA species quickly and completely into the monomeric genomes^11^. Therefore, depending on the biological functions and evolutionary pressures, the catalytic rate of self-cleaving ribozymes should vary in a wide range which is consistent with what we and others have found. For example, the *k*_obs_ of naturally occurring self-cleaving ribozymes belonging to the same class were found to vary by ∼5000 fold (0.08–387 h^−1^)^52^.

The varying rate of self-cleavage might also indicate the diversity of RNA folding kinetics and/or catalytic strategies used by different ribozymes. Conceivably, novel ribozymes might use less sophisticated mechanism(s) of catalysis. For example, they might function as metalloenzymes using bound cations, rather than nucleobase-mediated general acid-base catalysis. Furthermore, it is important to emphasize that *in vitro* measurements may underestimate *in vivo* ribozyme activities due to the inherent lack of the following in such assays: (1) critical long-range structural interactions formed by distal sequences^18,69–71^; (2) critical co-factors or ligands exemplified by the glucosamine-6-phosphate required for the *glmS* ribozyme activity^8,72^; and (3) specific RNA binding proteins^73^ and/or RNA modifications^74^ crucial for proper RNA folding. In this regard, it is noteworthy that the change in copy number of the full-length LAMC1 mRNA *in vivo* attributable to the hsa-rzyme-14 activity was significantly higher than would be expected based on the *in vitro* results. The existence of ribozymes with different biochemical requirements is supported by the discovery of the monovalent cation dependent MINDS ribozymes. *In vivo* activity of these ribozymes and the expression of their host transcripts could therefore be regulated by the changes in intracellular concentration of monovalent cations, such as K^+^ and Na^+^, that happen in response to cellular activity or environmental stimuli.

Our work clearly demonstrates that much remains unknown about the RNA-based catalysis, which in turn calls for further studies from multiple disciplines to investigate the RNA structures, reaction mechanisms, biological roles, and evolutionary contributions of ribozymes. In addition to their biological importance, ribozymes also have many high value biotechnological applications in multiple areas, such as diagnostics, synthetic biology^75^, gene editing^76,77^ and gene therapy^78,79^, that could be greatly facilitated by deeper understandings of the true diversity of the ribozyme structures and reaction mechanisms. We have shown that novel ribozymes could provide alternative approach for direct RNA detection. The approach we present here at the proof-of-principle level could be used to detect small or degraded RNAs that can be challenging to the conventional methods.

The method developed in this work, AbRiDGed, for the first time achieved efficient discovery of self-cleaving ribozymes at the genome-wide level and is sufficiently sensitive to discover self-cleaving ribozymes with relatively low *in vitro* activity, which can definitely facilitate further endeavors in ribozyme biology and relating biotechnological applications. Overall, our results suggest that even in modern complex eukaryotes, catalytic RNAs are actually very common and diverse, and might be involved in a variety of biological processes and contribute significantly to the process of speciation and the complexity of life.

## Methods

### AbRiDGed

#### Generation of genomic IVT template libraries

A genomic IVT template library was prepared as described previously^5^ by a limited DNase I digestion of the corresponding genomic DNA to a 200–1,000 bp size range followed by ligation of adapters and PCR amplification to generate a DNA library of IVT templates which contained DNA molecules flanked by the T7 promoter and the common adapter sequences at the 5′ and 3′ end respectively. The IVT templates for the 3333 randomly selected human genomic regions were amplified from the human genomic DNA using PCR with 10 primer pairs per reaction. The primers contained the T7 promoter at the 5′ end of each F (forward) primer and the common adapter adapter2_R at the 5′ end of each R (reverse) primer and were synthesized by Genewiz (Suzhou, China). The oligonucleotides, PCR primers and synthetic RNAs used in this study are listed in Table S23. The PCR products were purified and pooled to generate libraries of up to 960 products per pool.

#### IVT

Each library of genomic or random region IVT templates (200 ng) was combined with 0.02 ng of each of the 20 spike-in IVT templates (Table S1) and used for IVT which was performed in a 20 µl reaction containing 100 U of T7 RNA polymerase (NEB, no. M0251), 1× T7 RNA polymerase buffer (40 mM Tris-HCl pH 7.9, 6 mM MgCl_2_, 2 mM spermidine, 1 mM DTT), 0.625 mM of each ATP, UTP, GTP and CTP (NEB, no. N0450), and 37.5 mM GMP (Sangon Biotech, A600493) for 1 h at 37°C. The GMP was included to compensate for any potential inefficiencies of the following RppH treatment since adding it in an IVT reaction has been shown to result in a significant fraction of IVT products with 5′-monophosphates termini^80^. Then, 1 µl of 2 U of Turbo DNase (ThermoFisher, AM2238) was added to remove the DNA templates, followed by purification with 3× volumes of VAHTS RNA Clean Beads (Vazyme, no. N412). At least 5 IVT reactions were performed in parallel on the same IVT template library and combined for the following steps.

#### Self-cleavage reaction

The purified IVT products (1 µg) were subjected to denaturation for 2 min at 85°C and renaturation by slow (15 min) cooling to 25°C in 30 mM Tris-HCl pH 8.0 (Invitrogen, 15568025) and 150 mM KCl (Sigma, 60135). Compared with our previous method^5^, we modified the condition of the self-cleavage step by increasing the time of the incubation from 1 to 2 h, pH from 7.0 to 8.0 and the concentration of MgCl_2_ (VWR, no. E525) from 2 to 20 mM while keeping the concentration of KCl at 150 mM. The self-cleavage reactions were performed in 20 µl at 37℃ and, after the incubation, purified with 3× volumes of VAHTS RNA Clean Beads.

#### RppH and XRN-1 treatments

RppH converts 5′-triphosphates present on the nascent IVT product to 5′-monophoshates. The RNA molecules containing such 5′ ends are then selectively degraded by the XRN-1 exonuclease which is inactive on RNAs with 5′-OH termini. We increased the duration of the RppH and XRN-1 treatments and the amount of these enzymes compared with the previous method^5^. The purified self-cleavage products (1 µg) were incubated with 10 U of RppH (NEB, no. M0356) in 1× NEBuffer 2 in 20 µl volume at 37℃ for 2 h, followed by addition of 20 µl containing 10 U of RppH and 1× NEBuffer 2 and further incubation at 37℃ for 2 h. The RNA products were purified with 3× volumes of VAHTS RNA Clean Beads, eluted in 40 µl of water and incubated with 5 U XRN-1 (NEB, no. M0338) in 1× NEBuffer 3 in 50 µl volume for 4 h at 37℃, followed by purification with 3× volumes of VAHTS RNA Clean Beads.

#### Ligation and cDNA synthesis (L1 and L2 libraries)

The purified products of the self-cleavage reactions (≤200 ng) with or without the RppH and XRN-1 treatment were ligated by RtcB ligase (NEB, no. M0458) separately to the RNA_oligo1 or RNA_oligo2 oligonucleotides which had monophosphates at both 5′ and 3′ ends and synthesized by Genewiz (Table S23). The ligations were performed in 20 µl reactions containing 200 ng self-cleavage products, 50 pmol RNA_oligo1 or RNA_oligo2, 15 pmol RtcB ligase, 0.1 mM GTP, 1 mM MnCl_2_ and 1× RtcB Reaction Buffer at 37℃ for 2 h, followed by purification with 3× volumes of VAHTS RNA Clean Beads. The purified ligation products were then reverse transcribed using the common 3′ end primer adapter2_R (Table S23) by PrimeScript II kit (Takara, No. 6210A) as follows. The RNA was first denatured at 65°C for 5 min in a 15 µl volume containing 2 µl of 1 µM adapter2_R and 1 µl of 10 mM dNTP mix, followed by rapid snap-cooling on ice for 2 min. Then, 4 µl of 5× PrimeScript II buffer and 1 µl of PrimeScript II RTase (200 U/µl) were added, and the reaction was incubated at 50°C for 50 min, followed by 70°C for 15 min to inactivate the RTase. The cDNA was purified with 2× volumes of VAHTS DNA Clean Beads and eluted in 14 µl of water.

#### cDNA synthesis and tailing (TA and TC libraries)

The purified products of the self-cleavage reactions (<1 μg) with or without the RppH and XRN-1 treatment were reverse transcribed using the common 3′ end primer adapter2_R (Table S23) by PrimeScript II kit using the same reaction and purification conditions as above. The purified cDNAs (100 ng) were then tailed at 3′ ends by polyC or polyA tails separately as follows. First, the purified cDNA was denatured in a 19 µl reaction containing 2 µl of 10× TdT buffer and 2 µl of 2.5 mM CoCl_2_ for 5 min at 95°C and immediately put on ice for 2 min. 5 U of TdT (NEB, no. M0315) and 2 µl of 10 mM dCTP (Takara, 4028) or dATP (Takara, 4026) were then added to the denatured cDNA and the reaction was incubated at 37°C for 30 min. Then, 2 µl of 10 mM ddNTP (Roche, 03732738001) was added to the reaction mixture and the incubation was continued at 37°C for additional 30 min to block the free 3′-OH ends, followed by incubation at 70°C for 10 min to inactivate TdT. The reaction was purified with 1.5× volumes of VAHTS DNA Clean Beads and eluted in 14 µl of water.

#### Library construction for the high-throughput sequencing (L1, L2, TA and TC libraries)

The entire purified cDNA products (14 µl) for each of the four library types were PCR amplified using the common P7_RZ-adpR primer and one of the following primers depending on the library type: P5_N10G10 for the polyC-tailed cDNAs, P5_N10T12 for the polyA-tailed cDNAs, P5_oligo1_F for the cDNAs from the RNA_oligo1 ligation products, and P5_oligo2_F for the cDNAs from the RNA_oligo2 ligation products (Table S23). The 1^st^ round PCR was performed in a 20 µl volume containing 1× Taq buffer, 0.25mM dNTP mix, 0.5 µM each of primers and 1 U of Taq DNA polymerase (Tiangen, no.ET101). The cycling conditions were as follows: initial denaturation at 94°C for 3 min, followed by 30 cycles of denaturation at 94°C for 30 s, annealing at 55°C for 30 s and extension at 72°C for 1 min, then a final extension at 72°C for 10 min. The reactions were then purified with 1.5× volumes of VAHTS DNA Clean Beads and eluted in 20 µl of water. Then, 4 µl of the purified 1^st^ round PCR products were used as the input to the 2^nd^ round of PCR using common primers Illumina_P5 and Illumina_P7 for all library types (Table S23). The cycling conditions were as follows: initial denaturation at 94°C for 3 min, followed by 20 cycles of denaturation at 94°C for 30 s, annealing at 60°C for 30 s and extension at 72°C for 1 min, then a final extension at 72°C for 10 min. Amplified DNA was purified with 1.2× volumes of VAHTS DNA Clean Beads and eluted in 20 µl of water.

The purified 2^nd^ round PCR products were used directly for the high-throughput sequencing on the Illumina Novaseq 6000 platform (PE150; Novogene Corporation, Beijing). The sequencing scales were 25 giga base (GB) and 5 GB for respectively the human genome and random human regions libraries, 20 GB for the *M. musculus* libraries, 10 GB for the *D. rerio*, *O. sativa*, *A. thaliana* or *D. hydei* libraries, and 2 GB for the *A. nidulans* or *T. guizhouense* libraries.

#### Replicates, additional controls and library types used for different species

Each of the 4 types (L1, L2, TA or TC) of human, mouse, zebrafish and rice libraries was represented by 2 replicates of 2 samples (treated and not treated with RppH and XRN-1) — 4 samples per each library type. The fungi, *D. hydei* and *A. thaliana* libraries had 3 types (L1, TA and TC) with 3 replicates per library type. Also, for these four species, an additional control sample was generated for each library type to remove false positives produced by mis-annealing of the primers used in the library construction to endogenous genomic sequences. For the L1 library, this “endogenous control” was generated using the same procedure but with the omission of the ligation step, while for the TC and TA libraries such control libraries had respectively the polyC or polyA tailing steps omitted. For every species, each replicate was performed independently starting with the IVT step (Figure S1).

### Identification of candidate ribozymes

The reference genomes used in this study were listed in the Table S18. Only read pairs in which both reads started with the expected adapter sequences were included in the analysis. The adapter sequences were trimmed using the Cutadapt software (version 3.4). The read pairs originating from the genome or the spike-ins were separated by the adapter sequences at the 5′ ends of the reads 2. Any read pair in which either read 1 or 2 was shorter than 30 nt after the adapter trimming was excluded from further analysis. The processed read pairs were then aligned to either the genomic or spike-ins sequences using the Bowtie2 software (version 2.4.4). For the read pairs from the genome, only alignments with the highest mapping quality scores (MAPQ = 42) and genomic distances between reads 1 and 2 shorter than 2000 nt were kept. The genomic positions of the first nucleotides of reads 1 from each aligned read pair were used for subsequent analysis. The read count for each genomic position was normalized to the total sequencing depth of the corresponding library by calculating the CPM (counts per million) metric.

For each type of libraries, single-nucleotide peaks were identified as genomic positions detected in at least one sample from each replicate and possessing the highest mean CPM values among such positions in the ±6 nt region around them. To remove potential false positives resulting from the non-specific annealing of primers to the endogenous genomic sequences, the peaks that were preceded by sequences with significant matches to 3′ ends of the oligonucleotides used to construct the libraries were removed.

Subsequently, for each of the remaining peaks, a signal-to-local-background ratio was calculated as its corresponding read count divided by the sum of the read counts in the upstream 100 nt (the denominator was incremented by 1 to avoid division by zero). The ratio was calculated separately in each sample of the corresponding library type (including the samples where the read count was 0 at the corresponding genomic position), and then averaged.

To identify candidate cleavage sites, the peaks from the different types of libraries were merged using the following logic:

1. Since the TA or TC peaks might shift from the actual positions of cleavage sites due to the respective U or G bases immediately following the cleavage sites, the peaks from the TA and TC libraries were merged by allowing a maximum distance of 5 nt between a TA and a TC peak. The resulting merged TA-TC peak pair represented the 2 peaks and the positions in-between.
2. Expected to precisely represent the positions of the cleavage sites, the peaks from the L1 and L2 libraries were merged only if an L1 and an L2 peak were either overlapping or adjacent to each other.
3. Merged L1-L2 peak pairs or individual (remaining not merged) L1 or L2 peaks were merged with overlapping or adjacent individual (remaining not merged) TA or TC peaks or merged TA-TC peak pairs.

If multiple peaks from the same type of libraries were merged, then only the peak with the highest mean CPM value was kept. The genomic positions of the candidate cleavage sites were defined based on the read counts from the sequencing data as illustrated in Figures S4 and S7. Sometimes we observed disagreements between the different types of libraries in terms of the cleavage site locations, which were typically small (1–6 nt) and mainly due to the G or U bases immediately following the cleavage site and thus undetectable using respectively the polyC- or polyA-tailing approaches. In such cases, we used the ligation-based libraries (L1 and L2) to call the cleavage sites since ligation of the oligonucleotide takes place prior to subsequent processing steps, such as cDNA synthesis, and should theoretically represent the most precise location of the cleavage site. The distributions of the candidate cleavage sites along the genome were plotted and manually checked. Potential problematic regions were identified as short genomic regions containing abnormally numerous candidate sites. Candidate cleavage sites from such regions were then excluded from the subsequent analysis.

The remaining candidate cleavage sites were categorized into 3 tiers — Tiers 1, 2 and 3 represented sites formed by the merger of peaks from respectively 4, 3 and 2 types of libraries. The candidate cleavage sites were then ranked in each tier separately by the sum of the signal-to-local-background ratios of the peaks used to generate the sites divided by 4 — the number of library types.

The candidate cleavage sites from the randomly selected regions of the human genome were identified using a procedure identical to the one mentioned above with 3 exceptions: (1) the read pairs were aligned to the sequences of the random regions instead of the whole reference genome sequence; (2) a genomic position was defined as a peak only when its CPM was higher than those of the adjacent up and downstream nucleotides and the average CPM of the whole region; and (3) the signal-to-local-background ratio of each peak was calculated as its corresponding read count divided by the average read count of the whole region.

For *D. hydei*, *A. thaliana*, *A. nidulans* and *T. guizhouense* which had the additional “endogenous control” and lacked the L2 libraries, the analytic procedure was changed as follows: (1) peaks that were also detected in the “endogenous control” for the corresponding type of libraries were excluded from analysis; (2) to identify candidate cleavage sites, after the merger of TA and TC peaks, only L1 peaks were merged when overlapping or being adjacent to an individual TA or TC peak or a merged TA-TC peak pair; (3) candidate cleavage sites were categorized into 2 tiers with Tiers 1 and 2 representing sites formed by the merger of peaks from respectively 3 and 2 types of libraries; (4) when ranking the candidate cleavage sites, the sum of the signal-to-local-background ratios of the peaks used to generate the sites was divided by 3 instead of 4.

### Validation of ribozyme candidates

The IVT templates for each ribozyme candidate region were generated by PCR amplification either from genomic DNA or using pairs of overlapping synthetic oligonucleotides. In either case, the F primers always contained the T7 promoter sequence at the 5′ end (Table S23). The primers were made by Sangon Biotech (Shanghai, China) or Genewiz (Suzhou, China). The IVT was performed in 40 µl reaction volume with 200 U of T7 RNA polymerase (Vazyme, no. DD4101-01), 1× T7 RNA polymerase buffer (40 mM Tris-HCl pH 8, 20 mM MgCl_2_, 2 mM spermidine, 2.5 mM TCEP), 1.25 mM each of UTP (Vazyme, no. DD4105-PA-01), ATP (Vazyme, no. DD4106-PA-01), CTP (Vazyme, no. DD4107-PA-01) and GTP (Vazyme, no. DD4108-PA-01) for 2 h at 37°C. Then, 1 µl of 2 U of Turbo DNase was added to remove DNA templates for 30 min at 37°C. The full-length, uncleaved IVT products were purified on 6% PAGE gels containing 7 M urea.

For the self-cleavage test, the PAGE-purified full-length IVT products were first heated to 85°C for 2 min and then slowly cooled to 25°C for 15 min in 30 mM Tris-HCl (pH 8.0) and 150 mM KCl. Then, the reaction was split into 2 parts. MgCl_2_ was added to one part to the final concentration of 20 mM and the reaction was incubated at 37°C in a total volume of 10 µl. After 6 h and 12 h, aliquots of the reaction were taken and quenched by the addition of 1× volume of RNA loading buffer containing 88% formamide, 50 mM EDTA, 0.02% xylene cyanol and 0.02% bromophenol blue, and either stored at −80°C immediately or resolved by denaturing PAGE. In parallel, EDTA was added to the other part to the final concentration of 6 mM and incubated at 37°C in a total volume of 5 µl for 12 h as a control followed by the addition of the RNA loading buffer and processed as above. Finally, the products of the reactions in the RNA loading buffer were denatured at 70°C for 10 min, snap-cooled on ice and resolved using 6% denaturing PAGE gels containing 7 M urea and stained by 3× SYBR™ Gold (Thermo Fisher, S11494). Gels were imaged on either the 3500R or MINI Space 1000 systems (both manufactured by Tanon, Shanghai, China).

### Biochemical characterization of the self-cleaving ribozymes

Biochemical characterizations of the Mg^2+^-dependent ribozymes were performed on the uncleaved ribozyme sequences that were prepared by IVT and PAGE-purified as described above or synthetic RNAs (Table S23) made by custom oligonucleotide synthesis by Accurate Biotechnology Co., Ltd (Changsha, China). The time courses of cleavage were performed under 3 cleavage conditions for each ribozyme and also the SAM-V sequence: (1) 30 mM Tris-HCl pH 7.5, 10 mM MgCl_2_ and 150 mM KCl at 37°C, (2) 30 mM Tris-HCl pH 7.5 and 10 mM MgCl_2_ at 37°C, and (3) 50 mM Tris-HCl pH 8.3, 20 mM MgCl_2_ and 100 mM KCl at 23°C. For each condition, the purified uncleaved IVT products were denatured at 85°C for 2 min in the corresponding cleavage reaction buffer without MgCl_2_ and then slowly cooled to 25°C for 15 min. Cleavage was then initiated by the addition of MgCl_2_ to the appropriate final concentration. In the case of the SAM-V riboswitch sequence, cleavage was initiated by the addition of both SAM to the final concentration of 4 mM and MgCl_2_. The reactions at each time point were quenched by the addition of 1× volume of RNA loading buffer and kept at −80°C until resolved by PAGE. In parallel, for each ribozyme, the control reaction containing the same ingredients with the exception that MgCl_2_ was substituted with 6 mM EDTA and incubated for the longest time period. For SAM-V, an additional control reaction that contained MgCl_2_ but lacked SAM was also incubated for the longest time period. The products of the reactions in the RNA loading buffer were denatured at 70°C for 10 min, snap-cooled on ice and resolved on 6% denaturing PAGE gels containing 7 M urea. The sequences of the SAM-V riboswitch used on this study was identical to the 62 nt *metY* sequence (*metY*_62nt in Table S23) published by Poiata et al.^35^. The RNAs were made by custom oligonucleotide synthesis by Accurate Biotechnology Co., Ltd (Changsha, China).

The *k*_obs_ values were determined by nonlinear fitting to the equation *F_t_* = *F*_∞_(1 − *e*^−*kt*^), where *F_t_* and *F_∞_* are the cleavage fractions of the ribozyme incubated for time *t* and at the endpoint, respectively, and *k* is *k*_obs_. The p*K*_a_ values were determined by fitting the pH-*k*_obs_ profile data to the equation *k*_obs_ = *k*_clv_⁄(1 + 10^(p*K_a_*−pH)^), where *k*_clv_ is the intrinsic rate of cleavage. The fitting to equations were performed using the *nls* function in the R environment. All experiments were done in duplicates.

To determine the dependence of the activity of the hsa-rzyme-97 ribozyme on pH and Mg^2+^ concentration, the self-cleavage reactions were performed essentially the same as above with the following changes. To estimate the effect of pH, the reactions were performed at 37°C in 10 mM MgCl_2_ and 30 mM MES (pH 6.0 or 6.5), Tris-HCl (pH 7.0, 7.5, 8, 8.5 or 9.0) or CAPS (pH 9.5). To estimate the effect of Mg^2+^ concentration, the reactions were performed at 37°C in 30 mM Tris-HCl pH 7.5 and in the presence of 1 mM, 10 mM, 20 mM, 50 mM or 80 mM MgCl_2_.

Tests for the self-cleavage activities of the homologous sequences and the sequences with nucleotide substitutions were performed using uncleaved IVT products obtained as described above. The self-cleavage reactions for the homologs of the hsa-rzyme-100 ribozyme were performed in 30 mM Tris-HCl pH 8, 150 mM KCl and 50 mM MgCl_2_ at 37°C for 2 h, and the cleavage conditions for the homologs of hsa-rzyme-12 were the same, with the exception of the incubation time which was extended to 12 h. The self-cleavage reactions for all other sequences were performed in 30 mM Tris-HCl pH 8, 150 mM KCl and 20 mM MgCl_2_ at 37°C for 6 h. All reactions had the cognate EDTA controls, and their products were resolved on denaturing PAGE as described above.

The biochemical characterization of the MINDS ribozymes was performed essentially the same as above with the exception that the standard cleavage reaction contained 100 ng RNA in 30 mM Tris-HCl (pH 8), 150 mM KCl and no Mg^2+^. To investigate the dependence of self-cleavage on different monovalent or divalent metal ions, the 150 mM KCl in the standard reaction was replaced with one of the following: 150 mM NaCl (Macklin, N775944), 150 mM LiCl (Aladdin, no. L433504), 150 mM RbCl (Macklin, no. R817218), 150 mM CsCl (Macklin, no. C804680), 150 mM NH_4_Cl (Sinopharm Chemical Reagent Co., Ltd., no. 10001508), 6 mM MgCl_2_ (Sigma, no. 63069), 6mM MnCl_2_ (Sigma, no. SLMW2106), 6 mM CaCl2 (VWR, no. E506), 6 mM CoCl2 (Aladdin, no. C118625), 6 mM cobalt hexammine (Aladdin, no. H157071) or 2 mM spermidine (Psaitong, no. S50085). To estimate the synergistic effect of K^+^ and Mg^2+^, 6 mM MgCl_2_ was added to the standard reaction. Unless otherwise indicated, the reactions were always incubated at 37°C.

All gels were imaged on either the 3500R or MINI Space 1000 systems (both manufactured by Tanon, Shanghai, China). Image processing, including cleavage fraction calculations, were performed using the ImageJ software^81^.

### Mass spectrum analysis of cleavage products

To obtain the cleavage products, the synthetic RNAs (Table S23) representing the 34 nt hsa-rzyme-97 (hsa-rzyme-97_34nt), the 29-nt hsa-rzyme-12 (hsa-rzyme-12_29nt), and the 25-nt hsa-MINDS-1 ribozymes were incubated at 37°C in the following conditions respectively: (1) 30 mM Tris-HCl pH 7.5 and 1 mM MgCl_2_ for 12 h, (2) 30 mM Tris-HCl pH 7.5, 150 mM KCl and 10 mM MgCl_2_ for 12 h, or (3) 30 mM Tris-HCl pH 8, 150 mM KCl and 6 mM MgCl_2_ for 6 h. The cleavage products were purified using ethanol and detected using Matrix Assisted Laser Desorption Ionization Time of flight Mass Spectrometry (MALDI-TOF MS). The synthesis of RNAs and MALDI-TOF MS detection were both performed by Accurate Biotechnology Co., Ltd (Changsha, China).

### *In vivo* analysis of the LAMC1 ribozyme activity

The human LAMC1 mRNA sequence containing the WT ribozyme was synthesized based on the NCBI RefSeq NM_002293.4. All sequences were confirmed by Sanger sequencing. The WT and MT versions of LAMC1 mRNA were cloned into the pHS-AVC-LW2008 vector under the control of the CMV promoter. The vector also contains the *mNeonGreen* reporter gene under the control of *EF-1α* core promoter. The sequences and plasmids were commercially synthesized by Beijing Syngenbio Co., Ltd.

The 293FT cells were cultured at 37°C in a humidified atmosphere with 5% CO_2_ in DMEM (Gibco, Cat. No. 11965092) complete medium supplemented with 10% FBS (Vazyme, Cat. No. F101-01), 1× penicillin–streptomycin (Thermo Fisher Scientific, Cat. No.15070-063). Cells were transfected using the Lipofectamine™ 3000 transfection kit (Thermo Fisher, Cat. No. L3000015) with Opti-MEM (Gibco, Cat. No. 31985070) as the transfection medium when they reached approximately 70% confluence (14–16 h after seeding) according to the manufacturer’s instructions. After 6 h, the transfection medium was replaced with fresh DMEM complete medium. After 48 h, mNeonGreen-positive cells were collected using the CytoFLEX SRT Cell Sorter and the total RNA was extracted using TRIzol reagent (TianGen, Cat. No. DP424) and further purified using the E.Z.N.A.® Total RNA Kit (Omega, Cat. No R6834-02.).

The total RNA (1 µg) was reverse transcribed with the HiScript IV All-in-one Ultra RT SuperMix (Vazyme, Cat. No. R433) and the cDNA was used for the RT-qPCR with SYBR PCR master mix (Vazyme, Cat. No. Q412) on the Bio-Rad CFX96 Touch Real-Time PCR Detection System. The mNeonGreen mRNA level was used as internal references to normalize the expression levels of LAMC1. Absolute quantification of WT and MT LAMC1 expression levels for each of the six LAMC1 primer pairs was conducted based on the standard curves generated using serial dilutions of the corresponding plasmids. The initial plasmid concentrations were estimated using the Equalbit dsDNA BR Assay Kit (Vazyme, Cat. No. EQ122-01) on the Qubit 4 Fluorometer (Thermo Fisher Scientific, Cat. No. Q33238).

### Ribozyme sequence analysis

The nucleotide sequences of the newly discovered self-cleaving ribozymes and previously identified or predicted ribozymes from human and the 7 eukaryotes were clustered using the VSEARCH software (version 2.22.1)^82^ with the threshold of 40% sequence identity. The covariance models of 4654 RNA families were downloaded from the Rfam (release 14.10) and ZWD (https://bitbucket.org/zashaw/zashaweinbergdata; commit 241d692) databases. The nucleotide sequences of all the newly discovered human ribozymes were compared to these models using the Infernal software (version 1.1.4).

The sequences of the 11 L1MA ribozymes were aligned using the MAFFT software (version 7.505)^83^ and the resulting MSA was used to predict the RNA structure. Based on the MSA and the predicted RNA secondary structure of the 11 L1MA ribozymes, a covariance model was constructed and then employed to conduct a search against the human genome using the Infernal software to identify additional instances of L1MA ribozymes. The LINE-1 elements containing the L1MA ribozymes and a full-length LINE-1 element (GenBank accession: AF148856.1) were aligned using the MAFFT software. The RNA secondary structures of the hsa-rzyme-97 ribozyme and the MINDS ribozymes were predicted using the RNAfold program from the ViennaRNA package (version 2.5.1)^84^, while the L1MA ribozymes’ secondary structure was predicted using the RNAalifold program the ViennaRNA package. The RNA secondary structures were plotted using the R2R software (version 1.0.6)^85^.

The MFE for the MINDS ribozymes was calculated using the RNAeval program from the ViennaRNA package with the energy parameters of Andronescu model^86^. The MINDS ribozyme motif search in various genomes was performed using the RNAmotif program^87^. The sequence of each genome was shuffled 100 times at the chromosome level using the *sample* function of the R environment, while preserving the overall base composition and avoiding the N bases. The *p*-value for the enrichment or depletion of MINDS ribozyme in a species was calculated as N/100 where N is the number of shuffled sequences in which the number of the MINDS ribozymes was larger or smaller respectively than that in the original genome of the species.

Homologous sequences of the newly discovered human ribozymes were identified based on the pairwise whole-genome alignments retrieved from the Ensembl database^88^ using the Ensembl REST server. For each ribozyme, the homologous sequences were aligned using the MAFFT software, and the normalized sequence identities were calculated using the MView software (version 1.67)^89^. The evolutionary relationships between species were generated based on the NCBI Taxonomy database^90^.

The overlaps between the cleavage sites of the human self-cleaving ribozymes and the annotated transcripts (GENCODE database^91^, release 44), repetitive sequences (Repbase database^92^, downloaded from the UCSC Genome Browser, table “rmsk”) and vlincRNAs (taken from previous publications^26,93^) were generated using the Bedtools software (version 2.30.0). Each ribozyme was counted only once. The overlaps with transcripts were generated using the following successive steps: (1) ribozymes whose cleavage sites overlapped introns or exons of any annotated transcripts on either strand were classified as “genic”, while the other ribozymes were classified as “intergenic”; (2) a “genic” ribozyme would be assigned to the overlapping transcript on the same strand if such transcript existed, otherwise it would be assigned to the transcript on the opposite strand; (3) “genic” ribozymes whose cleavage sites were located in exons of lncRNAs on the same strand were further checked for overlap with any protein-coding mRNAs on the opposite strand. The analysis of enrichment or depletion of the human MINDS ribozymes in the human UTRs, introns, CDS or intergenic regions (GENCODE database, release 44) and promoters or enhancers (ENCODE Consortium^94^) was performed using the *regioneR* package^95^ in the R environment.

### Overlap between the cleavage site of human ribozymes and the transcriptome datasets

The CAGE data from 1829 human samples were downloaded from the FANTOM5 database^41^ (https://fantom.gsc.riken.jp/5/datafiles/latest/basic/); and the NAP-seq dataset was downloaded from the NCBI Gene Expression Omnibus (GEO) database (accession GSE192632). To generate the overlap with the CAGE dataset, the genomic coordinates of the cleavage sites of the human ribozymes were converted to the GRCh37 genome version using the *liftover* function from the *rtracklayer* package^96^ in R environment. The overlaps were generated using the Bedtools software allowing ≤5 nt between a cleavage site and a transcript boundary. To perform the permutation tests, the genomic coordinates of the cleavage sites of the human self-cleaving ribozymes were shuffled 1000 times using Bedtools, yielding 1000 sets of random genomic positions. The overlaps between the 1000 sets of random genomic positions and the CAGE and NAP-seq datasets were also generated using Bedtools allowing no more than 5-nt distance.

## Data availability

All high-throughput sequencing data are accessible under Sequence Read Archive (SRA) accession number PRJNA1119018 with reviewer link https://dataview.ncbi.nlm.nih.gov/object/PRJNA1119018?reviewer=8om9aa9a180htdj66f4qdmnsj9. All processed data mentioned in the manuscript are supplied in Supplementary Tables.

## Code availability

The custom code developed in this work has been deposited in Github (https://github.com/qifei9/rz_discovery_supplements) and Zenodo (https://doi.org/10.5281/zenodo.12875675).

## Acknowledgments

This study was funded by the Research Fund for International Senior Scientists from the National Natural Science Foundation of China (Grant No. 32150710525 to P.K) and National Natural Science Foundation of China (Grant No. 32170619 to P.K). F.Q. was supported by the National Natural Science Foundation of China (Grant No. 32571494), the National Key Research & Developmental Program of China (Grant No. 2024YFF1206802), the Natural Science Foundation of Xiamen, China (Grant No. 3502Z202473014), the Scientific Research Foundation of State Key Laboratory of Vaccines for Infectious Diseases, Xiang An Biomedicine Laboratory (Grant No. 2024XAKJ0102005) and the Fundamental Research Funds for the Central Universities (Grant No. 20720250096). Y.C. was supported by the National Natural Science Foundation of China (Grant No. 32201055), the Fundamental Research Funds for the Central Universities (Grant No. ZQN-1214) and the Scientific Research Funds of Huaqiao University (Grant No. 21BS127). We wish to thank Prof. Zhenzhong Yu from Nanjing Agricultural University for kindly providing the genomic DNAs of *Aspergillus nidulans* and *Trichoderma guizhouense*.

## Author Contributions

Conceptualization: P.K. and D.M.J.L. with input from F.Q. and Y.C. Methodology: P.K., D.M.J.L., F.Q., and Y.C. Software: F.Q. Validation: Y.L., Z.C., Y.Z., F.Z., K.L., M.L., S.C., I.L., W.Y., Q.T., W.Z., and L.H. Formal analysis: F.Q with input from Y.Z. Investigation: Y.C., F.Q., Y.L., Z.C., Y.Z., F.Z., K.L., M.L., S.C., I.L., W.Y., Q.T. and J.C. Data Curation: F.Q., Y.Z, J.C., W.Z., and Y.C. Writing – original draft preparation: P.K., F.Q., and Y.C. Writing – review and editing: P.K., D.M.J.L., F.Q., and Y.C. Supervision: P.K., F.Q., and Y.C. Project administration: P.K. Funding acquisition: P.K.

## References

1. Kruger, K., Grabowski, P.J., Zaug, A.J., Sands, J., Gottschling, D.E., and Cech, T.R. (1982). Self-splicing RNA: autoexcision and autocyclization of the ribosomal RNA intervening sequence of Tetrahymena. Cell 31, 147–157. 10.1016/0092-8674(82)90414-7.

2. Guerrier-Takada, C., Gardiner, K., Marsh, T., Pace, N., and Altman, S. (1983). The RNA moiety of ribonuclease P is the catalytic subunit of the enzyme. Cell 35, 849–857. 10.1016/0092-8674(83)90117-4.

3. Muller, S., Appel, B., Balke, D., Hieronymus, R., and Nubel, C. (2016). Thirty-five years of research into ribozymes and nucleic acid catalysis: where do we stand today? F1000Research 5. 10.12688/f1000research.8601.1.

4. Wilson, T.J., and Lilley, D.M.J. (2021). The potential versatility of RNA catalysis. Wiley interdisciplinary reviews. RNA 12, e1651. 10.1002/wrna.1651.

5. Chen, Y., Qi, F., Gao, F., Cao, H., Xu, D., Salehi-Ashtiani, K., and Kapranov, P. (2021). Hovlinc is a recently evolved class of ribozyme found in human lncRNA. Nat Chem Biol 17, 601–607. 10.1038/s41589-021-00763-0.

6. Gilbert, W. (1986). Origin of life: The RNA world. Nature 319, 618–618. 10.1038/319618a0.

7. Breaker, R.R. (2020). Imaginary Ribozymes. ACS Chemical Biology 15, 2020–2030. 10.1021/acschembio.0c00214.

8. Winkler, W.C., Nahvi, A., Roth, A., Collins, J.A., and Breaker, R.R. (2004). Control of gene expression by a natural metabolite-responsive ribozyme. Nature 428, 281–286. 10.1038/nature02362.

9. McCown, P.J., Winkler, W.C., and Breaker, R.R. (2012). Mechanism and distribution of glmS ribozymes. Methods Mol Biol 848, 113–129. 10.1007/978-1-61779-545-9_8.

10. Licatalosi, D.D., and Darnell, R.B. (2010). RNA processing and its regulation: global insights into biological networks. Nat Rev Genet 11, 75–87. 10.1038/nrg2673.

11. Weinberg, C.E., Weinberg, Z., and Hammann, C. (2019). Novel ribozymes: discovery, catalytic mechanisms, and the quest to understand biological function. Nucleic Acids Res 47, 9480–9494. 10.1093/nar/gkz737.

12. Lilley, D.M.J. (2019). Classification of the nucleolytic ribozymes based upon catalytic mechanism. F1000Research 8. 10.12688/f1000research.19324.1.

13. Jimenez, R.M., Polanco, J.A., and Luptak, A. (2015). Chemistry and Biology of Self-Cleaving Ribozymes. Trends Biochem Sci 40, 648–661. 10.1016/j.tibs.2015.09.001.

14. Chakravarty, A.K., Subbotin, R., Chait, B.T., and Shuman, S. (2012). RNA ligase RtcB splices 3’-phosphate and 5’-OH ends via covalent RtcB-(histidinyl)-GMP and polynucleotide- (3’)pp(5’)G intermediates. Proc Natl Acad Sci U S A 109, 6072–6077. 10.1073/pnas.1201207109.

15. Qi, F., Chen, J., Chen, Y., Sun, J., Lin, Y., Chen, Z., and Kapranov, P. (2024). Evaluating Performance of Different RNA Secondary Structure Prediction Programs Using Self-cleaving Ribozymes. Genom Proteom Bioinf, qzae043. 10.1093/gpbjnl/qzae043.

16. Olzog, V.J., Gartner, C., Stadler, P.F., Fallmann, J., and Weinberg, C.E. (2021). cyPhyRNA-seq: a genome-scale RNA-seq method to detect active self-cleaving ribozymes by capturing RNAs with 2’,3’ cyclic phosphates and 5’ hydroxyl ends. RNA Biol 18, 818–831. 10.1080/15476286.2021.1999105.

17. Weinberg, Z., Kim, P.B., Chen, T.H., Li, S., Harris, K.A., Lunse, C.E., and Breaker, R.R. (2015). New classes of self-cleaving ribozymes revealed by comparative genomics analysis. Nat Chem Biol 11, 606–610. 10.1038/nchembio.1846.

18. Martick, M., Horan, L.H., Noller, H.F., and Scott, W.G. (2008). A discontinuous hammerhead ribozyme embedded in a mammalian messenger RNA. Nature 454, 899–902. 10.1038/nature07117.

19. de la Pena, M., and Garcia-Robles, I. (2010). Intronic hammerhead ribozymes are ultraconserved in the human genome. EMBO Rep 11, 711–716. 10.1038/embor.2010.100.

20. Salehi-Ashtiani, K., Luptak, A., Litovchick, A., and Szostak, J.W. (2006). A genomewide search for ribozymes reveals an HDV-like sequence in the human CPEB3 gene. Science 313, 1788–1792. 10.1126/science.1129308.

21. Roth, A., Weinberg, Z., Chen, A.G., Kim, P.B., Ames, T.D., and Breaker, R.R. (2014). A widespread self-cleaving ribozyme class is revealed by bioinformatics. Nat Chem Biol 10, 56–60. 10.1038/nchembio.1386.

22. Kalvari, I., Nawrocki, E.P., Ontiveros-Palacios, N., Argasinska, J., Lamkiewicz, K., Marz, M., Griffiths-Jones, S., Toffano-Nioche, C., Gautheret, D., Weinberg, Z., et al. (2021). Rfam 14: expanded coverage of metagenomic, viral and microRNA families. Nucleic Acids Res 49, D192–D200. 10.1093/nar/gkaa1047.

23. Weinberg, Z., Lunse, C.E., Corbino, K.A., Ames, T.D., Nelson, J.W., Roth, A., Perkins, K.R., Sherlock, M.E., and Breaker, R.R. (2017). Detection of 224 candidate structured RNAs by comparative analysis of specific subsets of intergenic regions. Nucleic Acids Res 45, 10811–10823. 10.1093/nar/gkx699.

24. Mattick, J.S., Amaral, P.P., Carninci, P., Carpenter, S., Chang, H.Y., Chen, L.L., Chen, R., Dean, C., Dinger, M.E., Fitzgerald, K.A., et al. (2023). Long non-coding RNAs: definitions, functions, challenges and recommendations. Nat Rev Mol Cell Biol 24, 430–447. 10.1038/s41580-022-00566-8.

25. St Laurent, G., Wahlestedt, C., and Kapranov, P. (2015). The Landscape of long noncoding RNA classification. Trends Genet 31, 239–251. 10.1016/j.tig.2015.03.007.

26. St Laurent, G., Shtokalo, D., Dong, B., Tackett, M.R., Fan, X., Lazorthes, S., Nicolas, E., Sang, N., Triche, T.J., McCaffrey, T.A., et al. (2013). VlincRNAs controlled by retroviral elements are a hallmark of pluripotency and cancer. Genome Biol 14, R73. 10.1186/gb-2013-14-7-r73.

27. Khan, H., Smit, A., and Boissinot, S. (2006). Molecular evolution and tempo of amplification of human LINE-1 retrotransposons since the origin of primates. Genome Res 16, 78–87. 10.1101/gr.4001406.

28. Wilson, T.J., Liu, Y., Li, N.S., Dai, Q., Piccirilli, J.A., and Lilley, D.M.J. (2019). Comparison of the Structures and Mechanisms of the Pistol and Hammerhead Ribozymes. Journal of the American Chemical Society 141, 7865–7875. 10.1021/jacs.9b02141.

29. Lafontaine, D.A., Norman, D.G., and Lilley, D.M. (2001). Structure, folding and activity of the VS ribozyme: importance of the 2-3-6 helical junction. The EMBO journal 20, 1415–1424. 10.1093/emboj/20.6.1415.

30. Liu, Y., Wilson, T.J., McPhee, S.A., and Lilley, D.M. (2014). Crystal structure and mechanistic investigation of the twister ribozyme. Nat Chem Biol 10, 739–744. 10.1038/nchembio.1587.

31. Kobori, S., and Yokobayashi, Y. (2018). Analyzing and Tuning Ribozyme Activity by Deep Sequencing To Modulate Gene Expression Level in Mammalian Cells. ACS Synth Biol 7, 371–376. 10.1021/acssynbio.7b00367.

32. Nawrocki, E.P., and Eddy, S.R. (2013). Infernal 1.1: 100-fold faster RNA homology searches. Bioinformatics 29, 2933–2935. 10.1093/bioinformatics/btt509.

33. Johnson-Buck, A.E., McDowell, S.E., and Walter, N.G. (2011). Metal ions: supporting actors in the playbook of small ribozymes. Met Ions Life Sci 9, 175–196. 10.1039/9781849732512-00175.

34. Bevilacqua, P.C., Harris, M.E., Piccirilli, J.A., Gaines, C., Ganguly, A., Kostenbader, K., Ekesan, S., and York, D.M. (2019). An Ontology for Facilitating Discussion of Catalytic Strategies of RNA-Cleaving Enzymes. ACS Chem Biol 14, 1068–1076. 10.1021/acschembio.9b00202.

35. Poiata, E., Meyer, M.M., Ames, T.D., and Breaker, R.R. (2009). A variant riboswitch aptamer class for S-adenosylmethionine common in marine bacteria. RNA 15, 2046–2056. 10.1261/rna.1824209.

36. Huang, L., and Lilley, D.M.J. (2018). Structure and ligand binding of the SAM-V riboswitch. Nucleic Acids Res 46, 6869–6879. 10.1093/nar/gky520.

37. Qi, F., Motz, M., Jung, K., Lassak, J., and Frishman, D. (2018). Evolutionary analysis of polyproline motifs in Escherichia coli reveals their regulatory role in translation. PLOS Computational Biology 14, e1005987. 10.1371/journal.pcbi.1005987.

38. Xu, D., Tang, L., Zhou, J., Wang, F., Cao, H., Huang, Y., and Kapranov, P. (2023). Evidence for widespread existence of functional novel and non-canonical human transcripts. BMC Biol 21, 271. 10.1186/s12915-023-01753-5.

39. Tang, L., Xu, D., Luo, L., Ma, W., He, X., Diao, Y., Ke, R., and Kapranov, P. (2024). A novel human protein-coding locus identified using a targeted RNA enrichment technique. BMC Biol 22, 273. 10.1186/s12915-024-02069-8.

40. Chen, C.C., Han, J., Chinn, C.A., Rounds, J.S., Li, X., Nikan, M., Myszka, M., Tong, L., Passalacqua, L.F.M., Bredy, T., et al. (2024). Inhibition of Cpeb3 ribozyme elevates CPEB3 protein expression and polyadenylation of its target mRNAs and enhances object location memory. Elife 13. 10.7554/eLife.90116.

41. FANTOM, Consortium, and, the, RIKEN, PMI, and, CLST, and (DGT) (2014). A promoter-level mammalian expression atlas. Nature 507, 462–470. 10.1038/nature13182.

42. Weitzer, S., and Martinez, J. (2007). The human RNA kinase hClp1 is active on 3’ transfer RNA exons and short interfering RNAs. Nature 447, 222–226. 10.1038/nature05777.

43. Affymetrix, E.T.P., and Cold Spring Harbor Laboratory, E.T.P. (2009). Post-transcriptional processing generates a diversity of 5’-modified long and short RNAs. Nature 457, 1028–1032. 10.1038/nature07759.

44. Otsuka, Y., Kedersha, N.L., and Schoenberg, D.R. (2009). Identification of a cytoplasmic complex that adds a cap onto 5’-monophosphate RNA. Mol Cell Biol 29, 2155–2167. 10.1128/MCB.01325-08.

45. Mukherjee, C., Patil, D.P., Kennedy, B.A., Bakthavachalu, B., Bundschuh, R., and Schoenberg, D.R. (2012). Identification of cytoplasmic capping targets reveals a role for cap homeostasis in translation and mRNA stability. Cell Rep 2, 674–684. 10.1016/j.celrep.2012.07.011.

46. Liu, S., Huang, J., Zhou, J., Chen, S., Zheng, W., Liu, C., Lin, Q., Zhang, P., Wu, D., He, S., et al. (2024). NAP-seq reveals multiple classes of structured noncoding RNAs with regulatory functions. Nat Commun 15, 2425. 10.1038/s41467-024-46596-y.

47. Chen, S., Francioli, L.C., Goodrich, J.K., Collins, R.L., Kanai, M., Wang, Q., Alföldi, J., Watts, N.A., Vittal, C., Gauthier, L.D., et al. (2024). A genomic mutational constraint map using variation in 76,156 human genomes. Nature 625, 92–100. 10.1038/s41586-023-06045-0.

48. Landrum, M.J., Lee, J.M., Riley, G.R., Jang, W., Rubinstein, W.S., Church, D.M., and Maglott, D.R. (2013). ClinVar: public archive of relationships among sequence variation and human phenotype. Nucleic Acids Res 42, D980–D985. 10.1093/nar/gkt1113 %J Nucleic Acids Research.

49. Cerezo, M., Sollis, E., Ji, Y., Lewis, E., Abid, A., Bircan, Karatuğ O., Hall, P., Hayhurst, J., John, S., Mosaku, A., et al. (2024). The NHGRI-EBI GWAS Catalog: standards for reusability, sustainability and diversity. Nucleic Acids Res 53, D998–D1005. 10.1093/nar/gkae1070 %J Nucleic Acids Research.

50. Kaminski, M.M., Abudayyeh, O.O., Gootenberg, J.S., Zhang, F., and Collins, J.J. (2021). CRISPR-based diagnostics. Nat Biomed Eng 5, 643–656. 10.1038/s41551-021-00760-7.

51. Jiao, C., Peeck, N.L., Yu, J., Ghaem Maghami, M., Kono, S., Collias, D., Martinez Diaz, S.L., Larose, R., and Beisel, C.L. (2024). TracrRNA reprogramming enables direct PAM-independent detection of RNA with diverse DNA-targeting Cas12 nucleases. Nat Commun 15, 5909. 10.1038/s41467-024-50243-x.

52. Webb, C.H., Riccitelli, N.J., Ruminski, D.J., and Luptak, A. (2009). Widespread occurrence of self-cleaving ribozymes. Science 326, 953. 10.1126/science.1178084.

53. Hammann, C., Luptak, A., Perreault, J., and de la Pena, M. (2012). The ubiquitous hammerhead ribozyme. RNA 18, 871–885. 10.1261/rna.031401.111.

54. Webb, C.H., and Luptak, A. (2011). HDV-like self-cleaving ribozymes. RNA Biol 8, 719–727. 10.4161/rna.8.5.16226.

55. Perreault, J., Weinberg, Z., Roth, A., Popescu, O., Chartrand, P., Ferbeyre, G., and Breaker, R.R. (2011). Identification of hammerhead ribozymes in all domains of life reveals novel structural variations. PLoS Comput Biol 7, e1002031. 10.1371/journal.pcbi.1002031.

56. Kapranov, P., Drenkow, J., Cheng, J., Long, J., Helt, G., Dike, S., and Gingeras, T.R. (2005). Examples of the complex architecture of the human transcriptome revealed by RACE and high-density tiling arrays. Genome Res 15, 987–997. 10.1101/gr.3455305.

57. Carninci, P., Kasukawa, T., Katayama, S., Gough, J., Frith, M.C., Maeda, N., Oyama, R., Ravasi, T., Lenhard, B., Wells, C., et al. (2005). The transcriptional landscape of the mammalian genome. Science 309, 1559–1563. 10.1126/science.1112014.

58. Kapranov, P., Cheng, J., Dike, S., Nix, D.A., Duttagupta, R., Willingham, A.T., Stadler, P.F., Hertel, J., Hackermuller, J., Hofacker, I.L., et al. (2007). RNA maps reveal new RNA classes and a possible function for pervasive transcription. Science 316, 1484–1488. 10.1126/science.1138341.

59. Mudge, J.M., Frankish, A., and Harrow, J. (2013). Functional transcriptomics in the post-ENCODE era. Genome Res 23, 1961–1973. 10.1101/gr.161315.113.

60. Mercer, T.R., Wilhelm, D., Dinger, M.E., Solda, G., Korbie, D.J., Glazov, E.A., Truong, V., Schwenke, M., Simons, C., Matthaei, K.I., et al. (2011). Expression of distinct RNAs from 3’ untranslated regions. Nucleic Acids Res 39, 2393–2403. 10.1093/nar/gkq1158.

61. Mercer, T.R., Dinger, M.E., Bracken, C.P., Kolle, G., Szubert, J.M., Korbie, D.J., Askarian-Amiri, M.E., Gardiner, B.B., Goodall, G.J., Grimmond, S.M., and Mattick, J.S. (2010). Regulated post-transcriptional RNA cleavage diversifies the eukaryotic transcriptome. Genome Res 20, 1639–1650. 10.1101/gr.112128.110.

62. Tuck, A.C., and Tollervey, D. (2011). RNA in pieces. Trends Genet 27, 422–432. 10.1016/j.tig.2011.06.001.

63. Wang, X., Yu, D., Yu, J., Hu, H., Hang, R., Amador, Z., Chen, Q., Chai, J., and Chen, X. (2024). Toll/interleukin-1 receptor (TIR) domain-containing proteins have NAD-RNA decapping activity. Nat Commun 15, 2261. 10.1038/s41467-024-46499-y.

64. Yang, E., van Nimwegen, E., Zavolan, M., Rajewsky, N., Schroeder, M., Magnasco, M., and Darnell, J.E., Jr. (2003). Decay rates of human mRNAs: correlation with functional characteristics and sequence attributes. Genome Res 13, 1863–1872. 10.1101/gr.1272403.

65. Wang, C., and Liu, H. (2022). Factors influencing degradation kinetics of mRNAs and half-lives of microRNAs, circRNAs, lncRNAs in blood in vitro using quantitative PCR. Sci Rep 12, 7259. 10.1038/s41598-022-11339-w.

66. Zocher, S., McCloskey, A., Karasinsky, A., Schulte, R., Friedrich, U., Lesche, M., Rund, N., Gage, F.H., Hetzer, M.W., and Toda, T. (2024). Lifelong persistence of nuclear RNAs in the mouse brain. Science 384, 53–59. 10.1126/science.adf3481.

67. Zhang, J., and Qian, W. (2025). Functional synonymous mutations and their evolutionary consequences. Nature Reviews Genetics 26, 789–804. 10.1038/s41576-025-00850-1.

68. Mattick, J.S. (2004). RNA regulation: a new genetics? Nat Rev Genet 5, 316–323. 10.1038/nrg1321.

69. Khvorova, A., Lescoute, A., Westhof, E., and Jayasena, S.D. (2003). Sequence elements outside the hammerhead ribozyme catalytic core enable intracellular activity. Nat Struct Biol 10, 708–712. 10.1038/nsb959.

70. Penedo, J.C., Wilson, T.J., Jayasena, S.D., Khvorova, A., and Lilley, D.M. (2004). Folding of the natural hammerhead ribozyme is enhanced by interaction of auxiliary elements. RNA 10, 880–888. 10.1261/rna.5268404.

71. Murchie, A.I., Thomson, J.B., Walter, F., and Lilley, D.M. (1998). Folding of the hairpin ribozyme in its natural conformation achieves close physical proximity of the loops. Mol Cell 1, 873–881. 10.1016/s1097-2765(00)80086-6.

72. Klein, D.J., and Ferre-D’Amare, A.R. (2006). Structural basis of glmS ribozyme activation by glucosamine-6-phosphate. Science 313, 1752–1756. 10.1126/science.1129666.

73. Semrad, K. (2011). Proteins with RNA chaperone activity: a world of diverse proteins with a common task-impediment of RNA misfolding. Biochem Res Int 2011, 532908. 10.1155/2011/532908.

74. Tanzer, A., Hofacker, I.L., and Lorenz, R. (2019). RNA modifications in structure prediction - Status quo and future challenges. Methods 156, 32–39. 10.1016/j.ymeth.2018.10.019.

75. Dykstra, P.B., Kaplan, M., and Smolke, C.D. (2022). Engineering synthetic RNA devices for cell control. Nat Rev Genet 23, 215–228. 10.1038/s41576-021-00436-7.

76. Tiyaboonchai, A., Vonada, A., Posey, J., Pelz, C., Wakefield, L., and Grompe, M. (2022). Self-cleaving guide RNAs enable pharmacological selection of precise gene editing events in vivo. Nat Commun 13, 7391. 10.1038/s41467-022-35097-5.

77. Liu, Z.X., Zhang, S., Zhu, H.Z., Chen, Z.H., Yang, Y., Li, L.Q., Lei, Y., Liu, Y., Li, D.Y., Sun, A., et al. (2024). Hydrolytic endonucleolytic ribozyme (HYER) is programmable for sequence-specific DNA cleavage. Science 383, eadh4859. 10.1126/science.adh4859.

78. Zhong, G., Wang, H., He, W., Li, Y., Mou, H., Tickner, Z.J., Tran, M.H., Ou, T., Yin, Y., Diao, H., and Farzan, M. (2020). A reversible RNA on-switch that controls gene expression of AAV-delivered therapeutics in vivo. Nat Biotechnol 38, 169–175. 10.1038/s41587-019-0357-y.

79. Wang, L.L., Wu, C.Q., Zhang, Q.L., Wang, Y., Liu, Y., Yang, W.J., Ye, S.L., Tian, Y., and Xu, L. (2024). Chemically Cross-Linked Hammerhead Ribozyme as an Efficient RNA Interference Tool. Journal of the American Chemical Society 146, 6665–6674. 10.1021/jacs.3c12702.

80. Jiang, X., Diwa, A., and Belasco, J.G. (2000). Regions of RNase E important for 5’-end-dependent RNA cleavage and autoregulated synthesis. J Bacteriol 182, 2468–2475. 10.1128/JB.182.9.2468-2475.2000.

81. Schindelin, J., Arganda-Carreras, I., Frise, E., Kaynig, V., Longair, M., Pietzsch, T., Preibisch, S., Rueden, C., Saalfeld, S., Schmid, B., et al. (2012). Fiji: an open-source platform for biological-image analysis. Nat Methods 9, 676–682. 10.1038/nmeth.2019.

82. Rognes, T., Flouri, T., Nichols, B., Quince, C., and Mahe, F. (2016). VSEARCH: a versatile open source tool for metagenomics. PeerJ 4, e2584. 10.7717/peerj.2584.

83. Katoh, K., Misawa, K., Kuma, K., and Miyata, T. (2002). MAFFT: a novel method for rapid multiple sequence alignment based on fast Fourier transform. Nucleic Acids Res 30, 3059–3066. 10.1093/nar/gkf436.

84. Lorenz, R., Bernhart, S.H., Honer Zu Siederdissen, C., Tafer, H., Flamm, C., Stadler, P.F., and Hofacker, I.L. (2011). ViennaRNA Package 2.0. Algorithms for molecular biology: AMB 6, 26. 10.1186/1748-7188-6-26.

85. Weinberg, Z., and Breaker, R.R. (2011). R2R - software to speed the depiction of aesthetic consensus RNA secondary structures. BMC Bioinformatics 12, 3. 10.1186/1471-2105-12-3.

86. Andronescu, M., Condon, A., Hoos, H.H., Mathews, D.H., and Murphy, K.P. (2007). Efficient parameter estimation for RNA secondary structure prediction. Bioinformatics 23, i19–28. 10.1093/bioinformatics/btm223.

87. Macke, T.J., Ecker, D.J., Gutell, R.R., Gautheret, D., Case, D.A., and Sampath, R. (2001). RNAMotif, an RNA secondary structure definition and search algorithm. Nucleic Acids Res 29, 4724–4735. 10.1093/nar/29.22.4724.

88. Zerbino, D.R., Achuthan, P., Akanni, W., Amode, M R., Barrell, D., Bhai, J., Billis, K., Cummins, C., Gall, A., Girón, C.G., et al. (2017). Ensembl 2018. Nucleic Acids Res 46, D754–D761. 10.1093/nar/gkx1098.

89. Brown, N.P., Leroy, C., and Sander, C. (1998). MView: a web-compatible database search or multiple alignment viewer. Bioinformatics 14, 380-381. 10.1093/bioinformatics/14.4.380.

90. Schoch, C.L., Ciufo, S., Domrachev, M., Hotton, C.L., Kannan, S., Khovanskaya, R., Leipe, D., Mcveigh, R., O’Neill, K., Robbertse, B., et al. (2020). NCBI Taxonomy: a comprehensive update on curation, resources and tools. Database 2020. 10.1093/database/baaa062.

91. Frankish, A., Diekhans, M., Ferreira, A.M., Johnson, R., Jungreis, I., Loveland, J., Mudge, J.M., Sisu, C., Wright, J., Armstrong, J., et al. (2019). GENCODE reference annotation for the human and mouse genomes. Nucleic Acids Res 47, D766–D773. 10.1093/nar/gky955.

92. Jurka, J. (2000). Repbase update: a database and an electronic journal of repetitive elements. Trends Genet 16, 418–420. 10.1016/s0168-9525(00)02093-x.

93. Caron, M., St-Onge, P., Drouin, S., Richer, C., Sontag, T., Busche, S., Bourque, G., Pastinen, T., and Sinnett, D. (2018). Very long intergenic non-coding RNA transcripts and expression profiles are associated to specific childhood acute lymphoblastic leukemia subtypes. PLoS One 13, e0207250. 10.1371/journal.pone.0207250.

94. Consortium, E.P., Moore, J.E., Purcaro, M.J., Pratt, H.E., Epstein, C.B., Shoresh, N., Adrian, J., Kawli, T., Davis, C.A., Dobin, A., et al. (2020). Expanded encyclopaedias of DNA elements in the human and mouse genomes. Nature 583, 699–710. 10.1038/s41586-020-2493-4.

95. Gel, B., Díez-Villanueva, A., Serra, E., Buschbeck, M., Peinado, M.A., and Malinverni, R. (2015). regioneR: an R/Bioconductor package for the association analysis of genomic regions based on permutation tests. Bioinformatics 32, 289–291. 10.1093/bioinformatics/btv562 %J Bioinformatics.

96. Lawrence, M., Gentleman, R., and Carey, V. (2009). rtracklayer: an R package for interfacing with genome browsers. Bioinformatics 25, 1841–1842. 10.1093/bioinformatics/btp328 %J Bioinformatics.

