## Supplementary Figures for "Widespread Presence of Catalytic RNAs in All Domains of Life": Figure S3.pdf

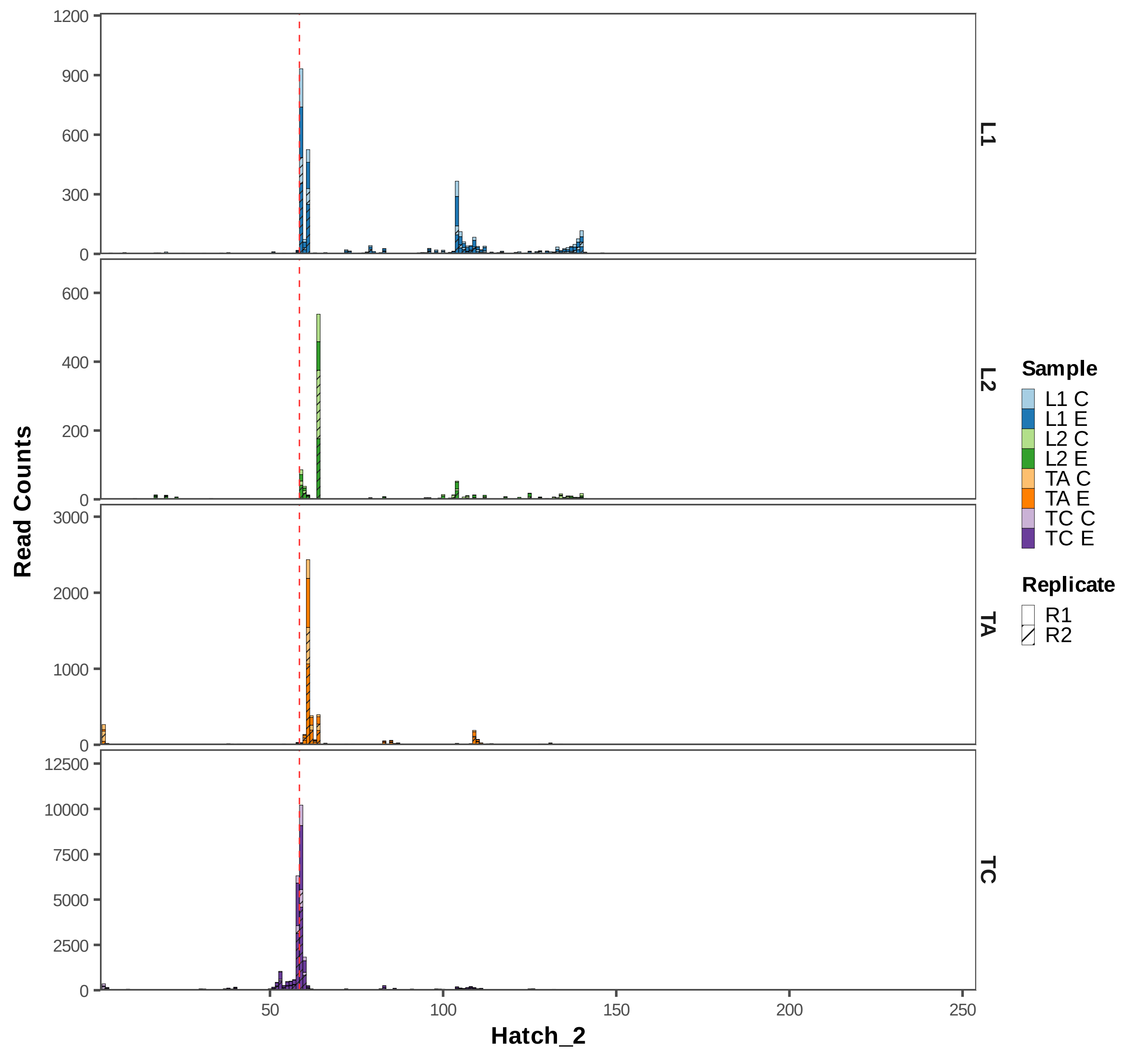

Read Counts
Hatch_2
L1
L2
TA
TC
1200
900
600
300
0
600
400
200
0
3000
2000
1000
0
12500
10000
7500
5000
2500
0
50
100
150
200
250
Sample
L1 C
L1 E
L2 C
L2 E
TA C
TA E
TC C
TC E
Replicate
R1
R2

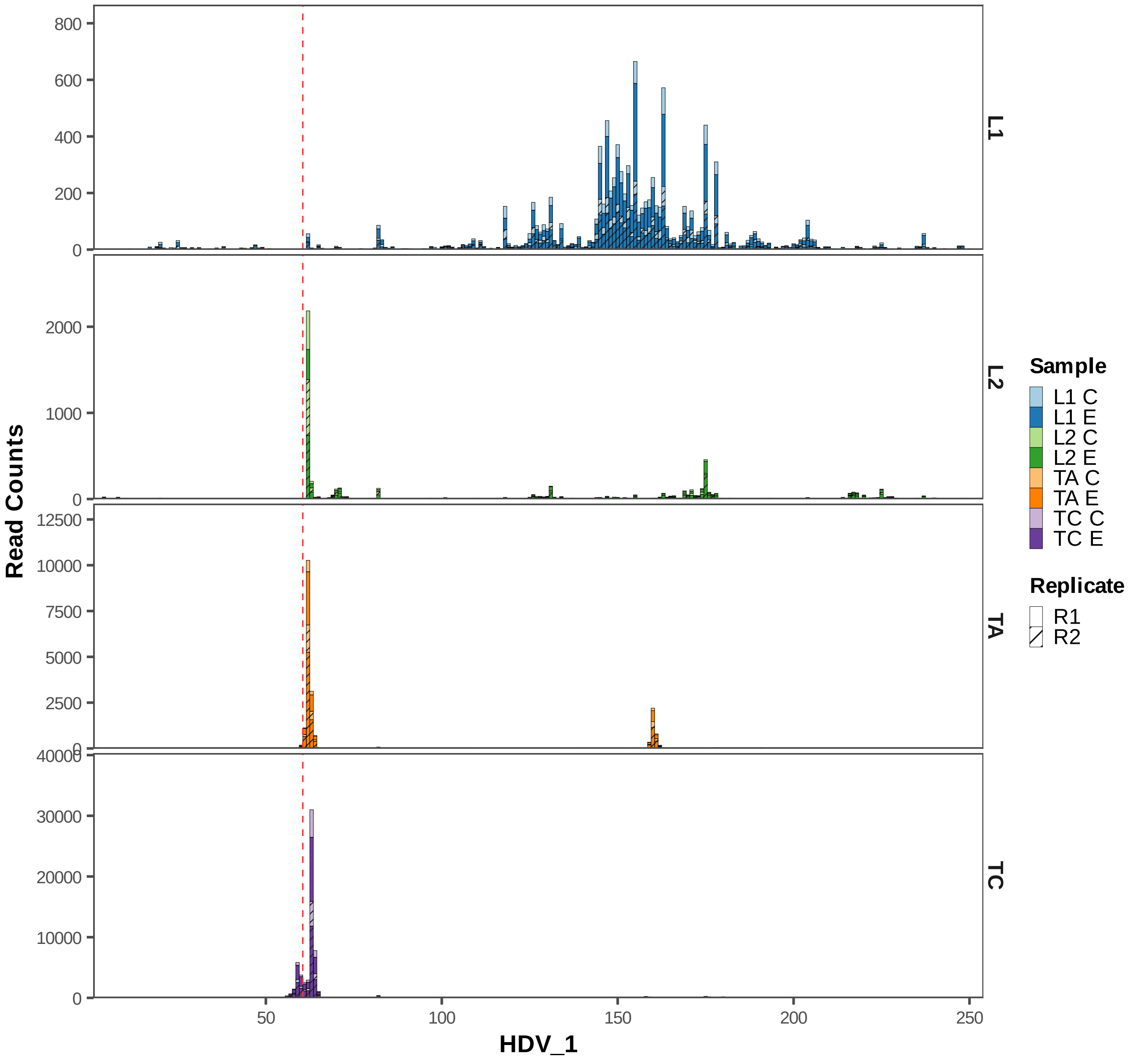

Read Counts
HDV_1
L1
L2
TA
TC
Sample
L1 C
L1 E
L2 C
L2 E
TA C
TA E
TC C
TC E
Replicate
R1
R2
800
600
400
200
0
2000
1000
12500
10000
7500
5000
2500
40000
30000
20000
50
100
150
200
250

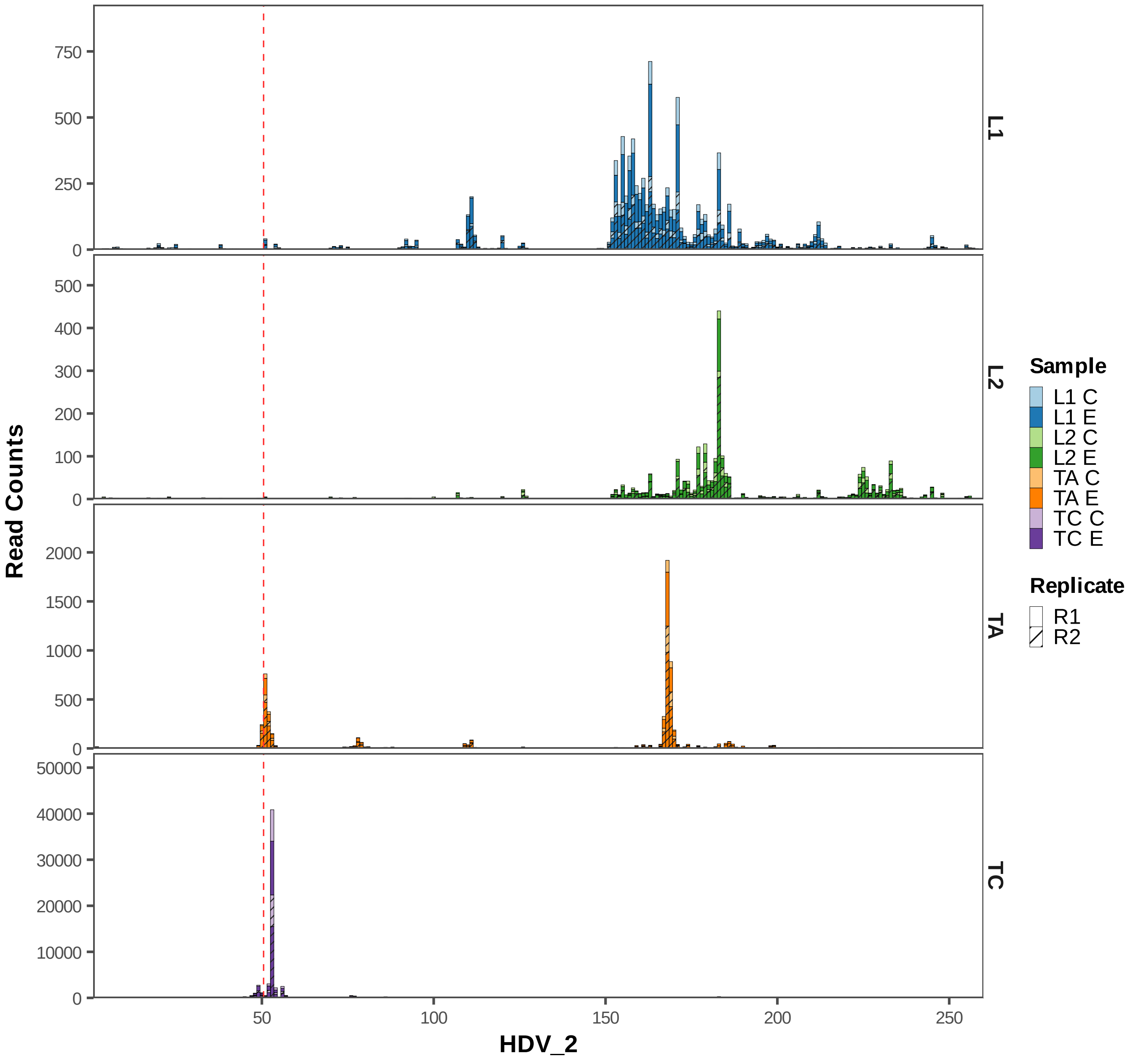

Read Counts
HDV_2
L1
L2
TA
TC
750
500
250
0
400
300
200
100
2000
1500
1000
50000
40000
30000
20000
10000
50
150
Sample
L1 C
L1 E
L2 C
L2 E
TA C
TA E
TC C
TC E
Replicate
R1
R2

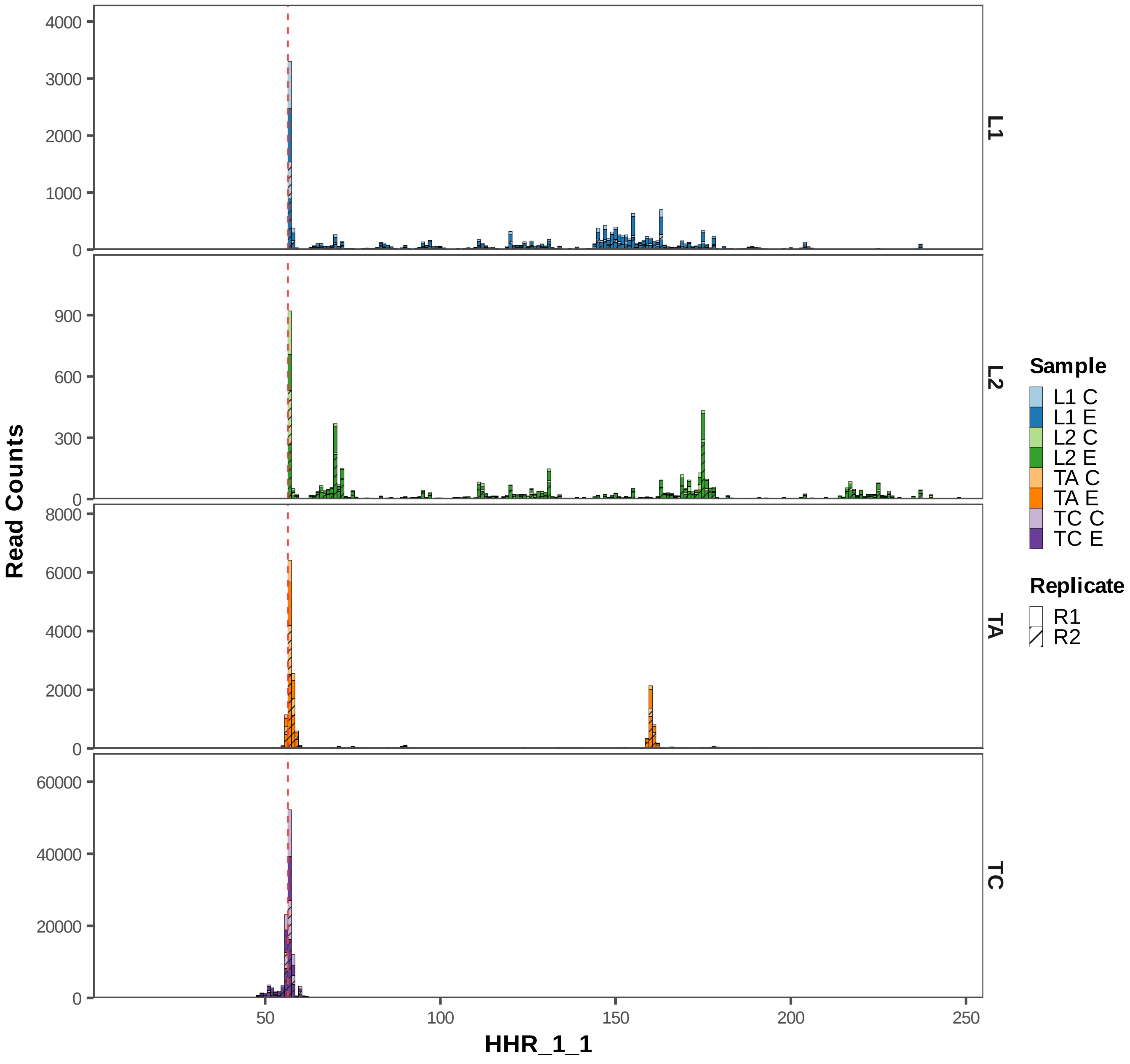

Read Counts
HHR_1_1
L1
L2
TA
TC
4000
3000
2000
1000
0
900
600
300
0
8000
6000
4000
2000
0
60000
40000
20000
0
50
100
150
200
250
Sample
L1 C
L1 E
L2 C
L2 E
TA C
TA E
TC C
TC E
Replicate
R1
R2

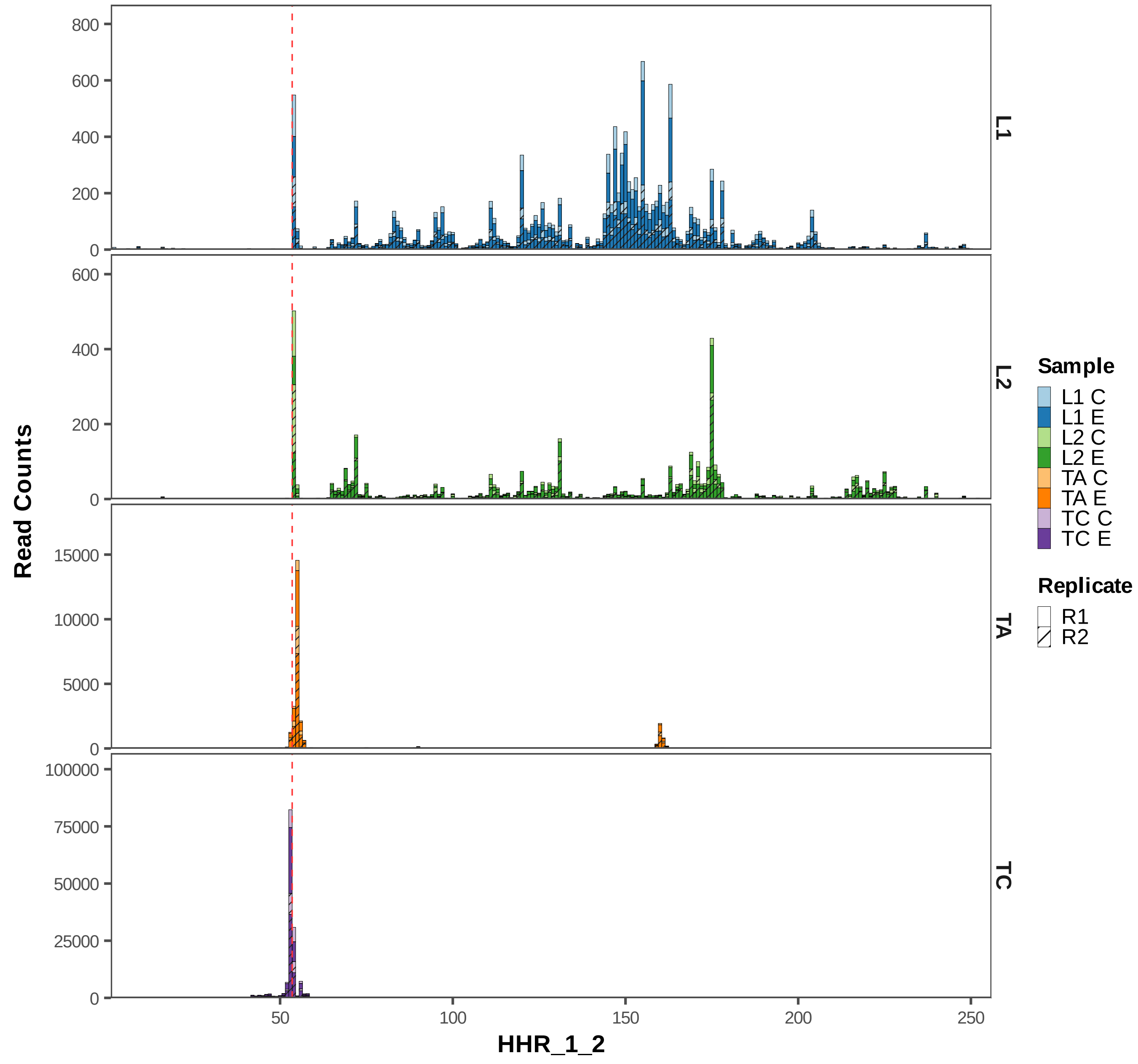
Read Counts
HHR_1_2
L1
L2
TA
TC
800
600
400
200
0
15000
10000
5000
100000
75000
50000
25000
50
100
150
200
250
Sample
L1 C
L1 E
L2 C
L2 E
TA C
TA E
TC C
TC E
Replicate
R1
R2

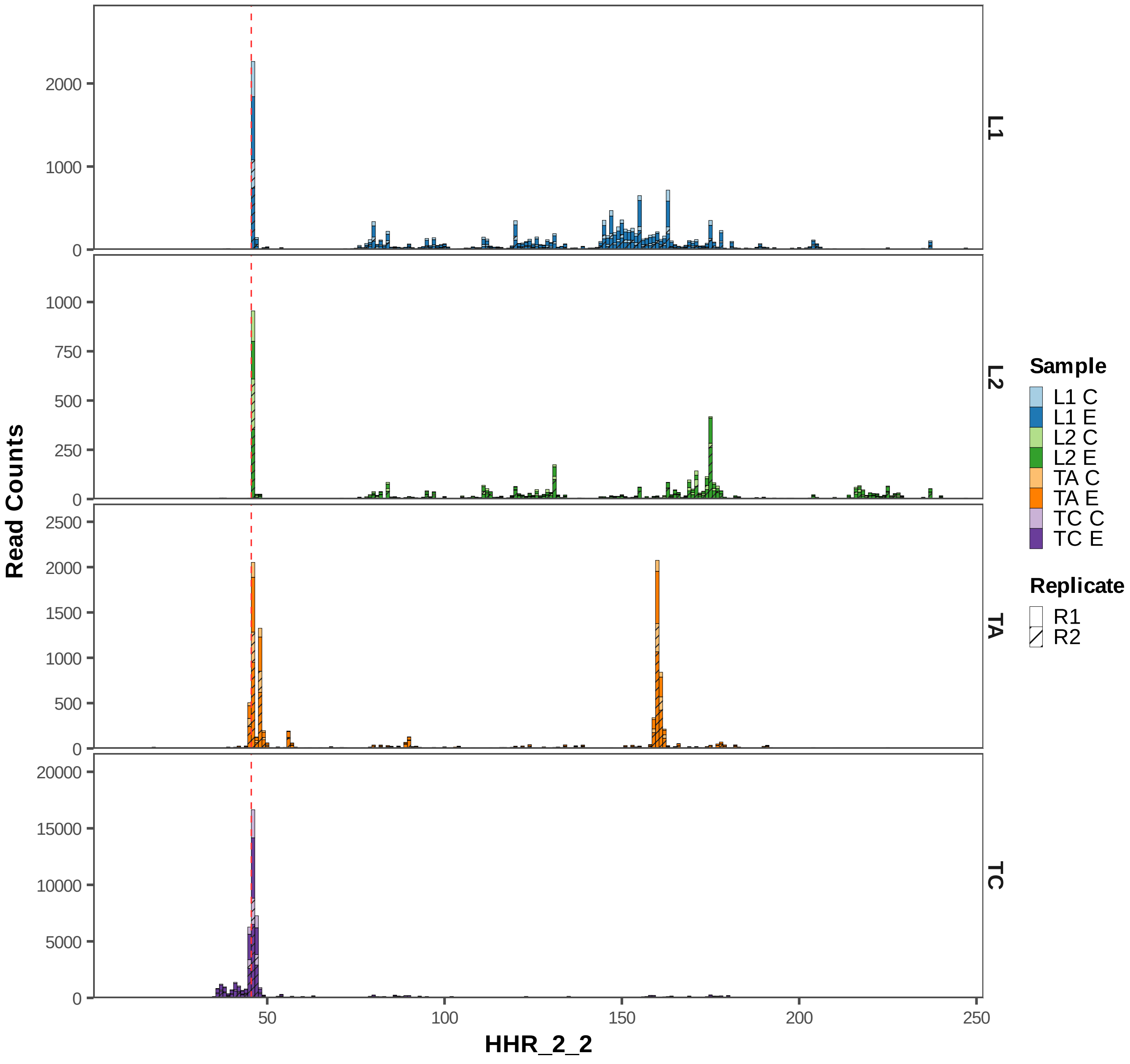

Read Counts
HHR_2_2
L1
L2
TA
TC
0
1000
2000
250
500
750
1500
2500
5000
10000
15000
20000
50
100
150
200
Sample
L1 C
L1 E
L2 C
L2 E
TA C
TA E
TC C
TC E
Replicate
R1
R2

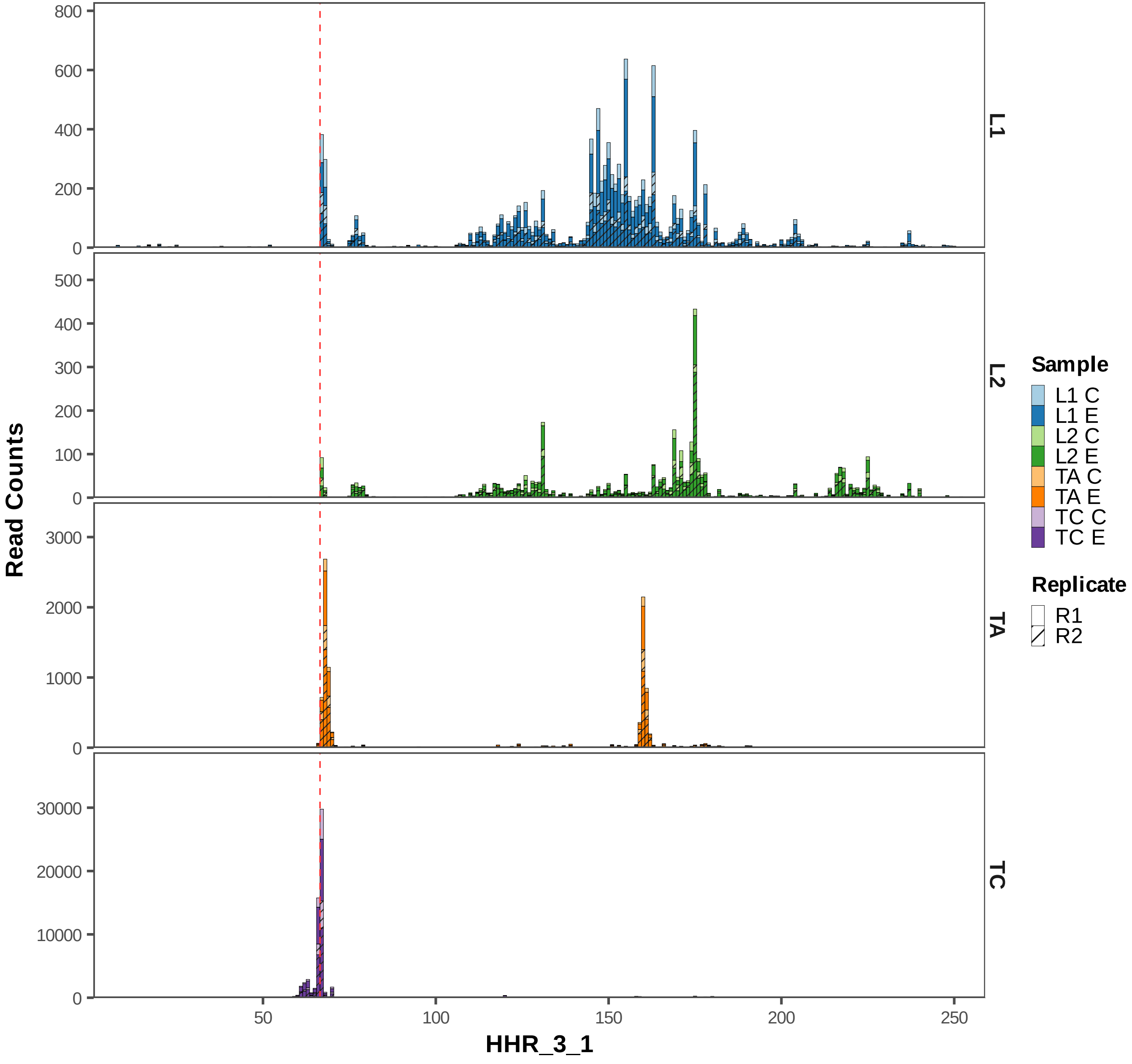

Read Counts
HHR_3_1
L1
L2
TA
TC
800
600
400
200
0
500
300
100
3000
2000
1000
30000
20000
10000
50
150
250
Sample
L1 C
L1 E
L2 C
L2 E
TA C
TA E
TC C
TC E
Replicate
R1
R2

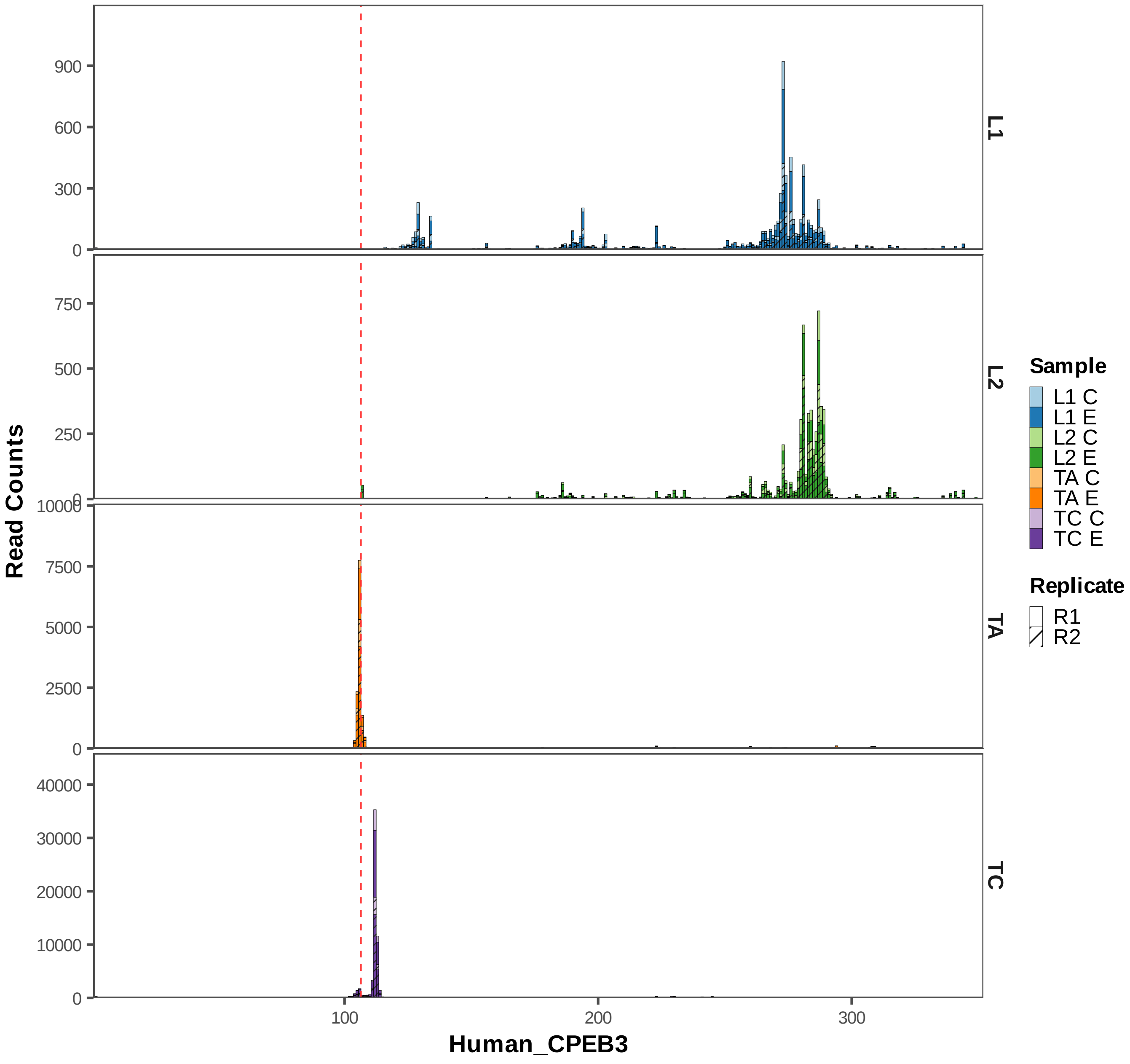

L1
L2
TA
TC
Read Counts
Human_CPEB3
900
600
300
0
750
500
250
10000
7500
5000
2500
40000
30000
20000
10000
100
200
300
Sample
L1 C
L1 E
L2 C
L2 E
TA C
TA E
TC C
TC E
Replicate
R1
R2

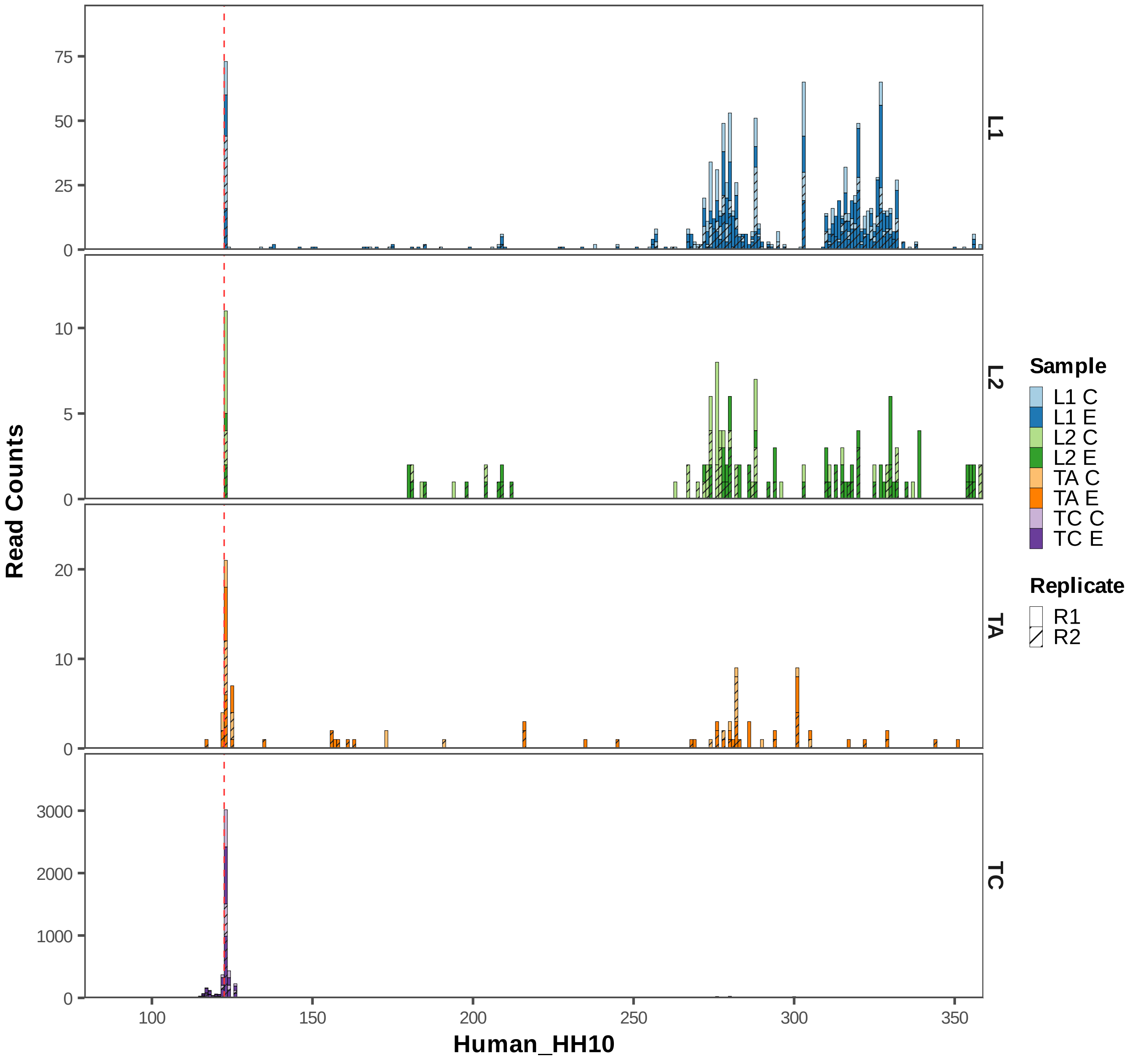
Read Counts
Human_HH10
L1
L2
TA
TC
Sample
L1 C
L1 E
L2 C
L2 E
TA C
TA E
TC C
TC E
Replicate
R1
R2
0
25
50
75
0
5
10
0
10
20
0
1000
2000
3000
100
150
200
250
300
350

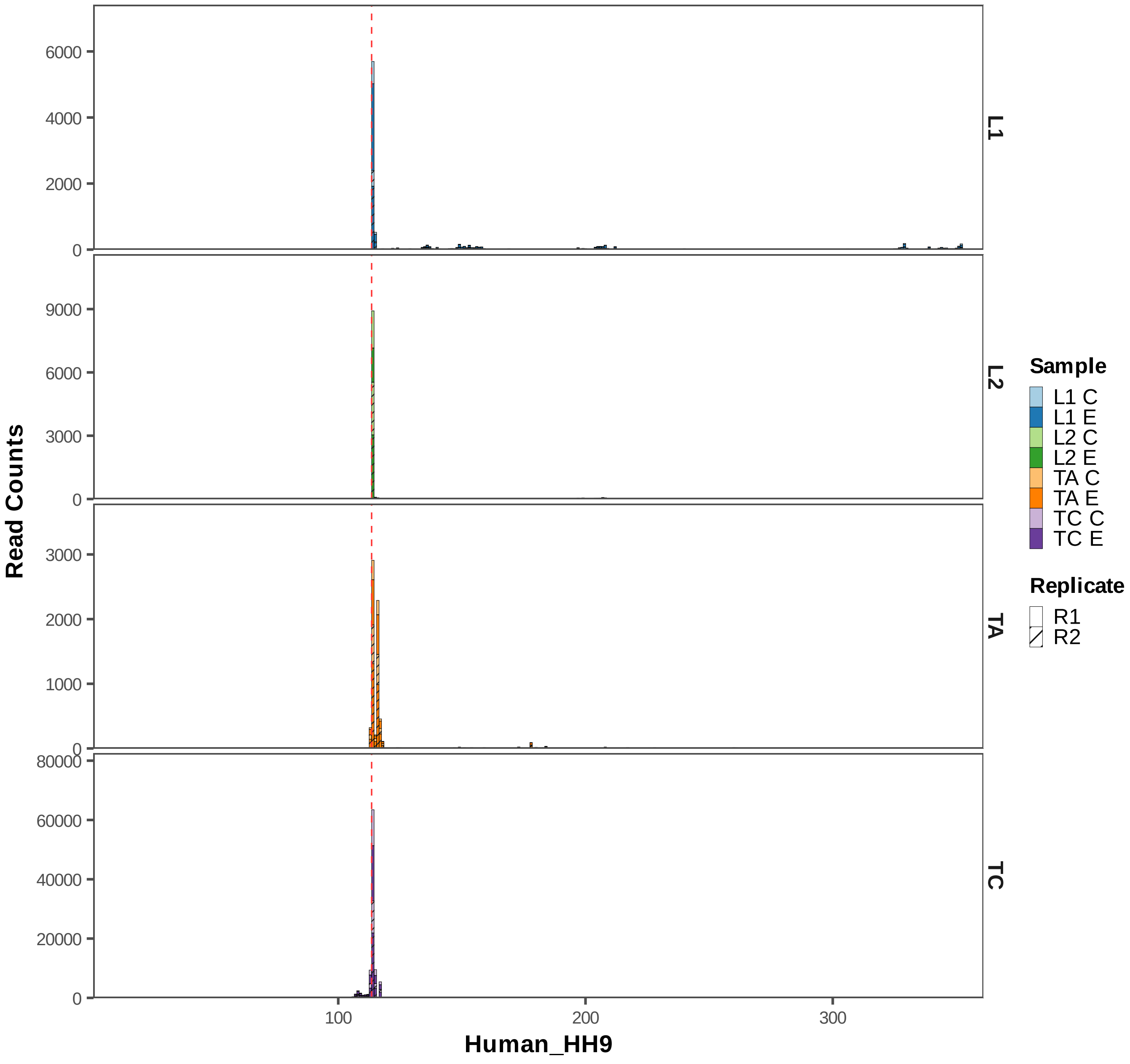

Read Counts
Human_HH9
L1
L2
TA
TC
6000
4000
2000
0
9000
6000
3000
0
3000
2000
1000
0
80000
60000
40000
20000
0
100
200
300
Sample
L1 C
L1 E
L2 C
L2 E
TA C
TA E
TC C
TC E
Replicate
R1
R2

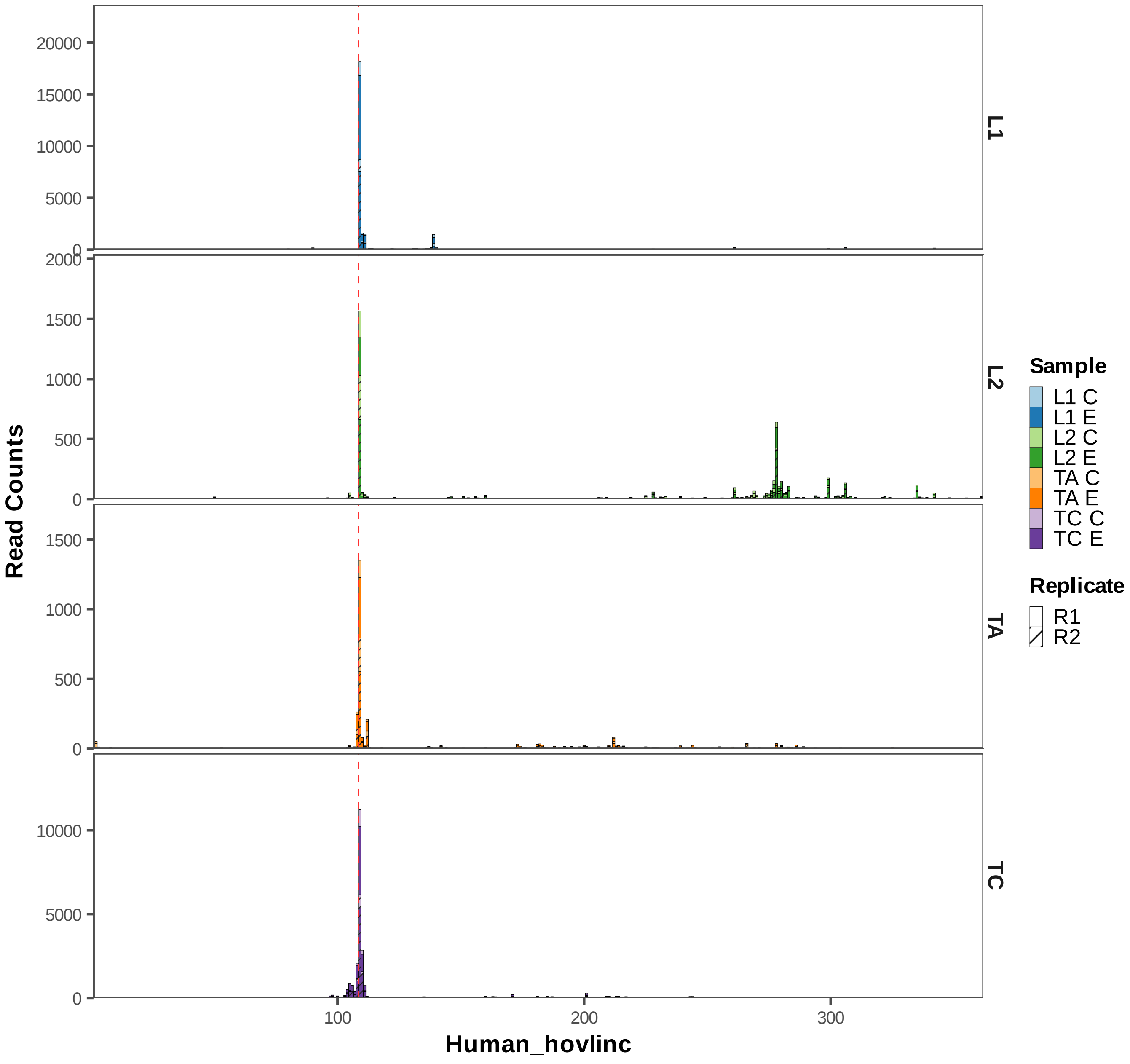
Read Counts
Human_hovlinc
L1
L2
TA
TC
Sample
L1 C
L1 E
L2 C
L2 E
TA C
TA E
TC C
TC E
Replicate
R1
R2
0
5000
10000
15000
20000
500
1000
1500
2000
100
200
300

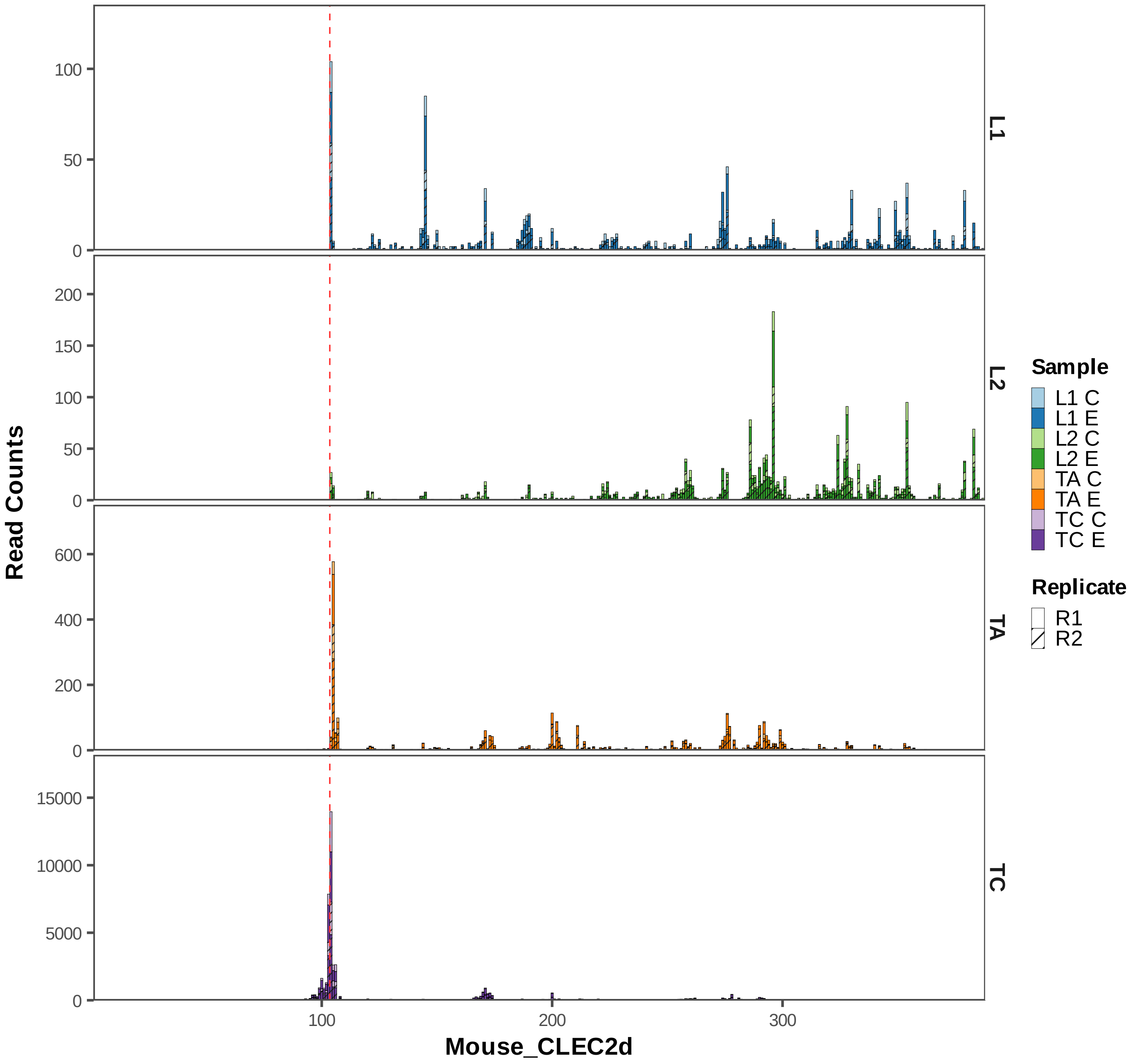

Read Counts
Mouse_CLEC2d
L1
L2
TA
TC
Sample
L1 C
L1 E
L2 C
L2 E
TA C
TA E
TC C
TC E
Replicate
R1
R2
100
200
300
0
50
100
150
200
400
600
5000
10000
15000

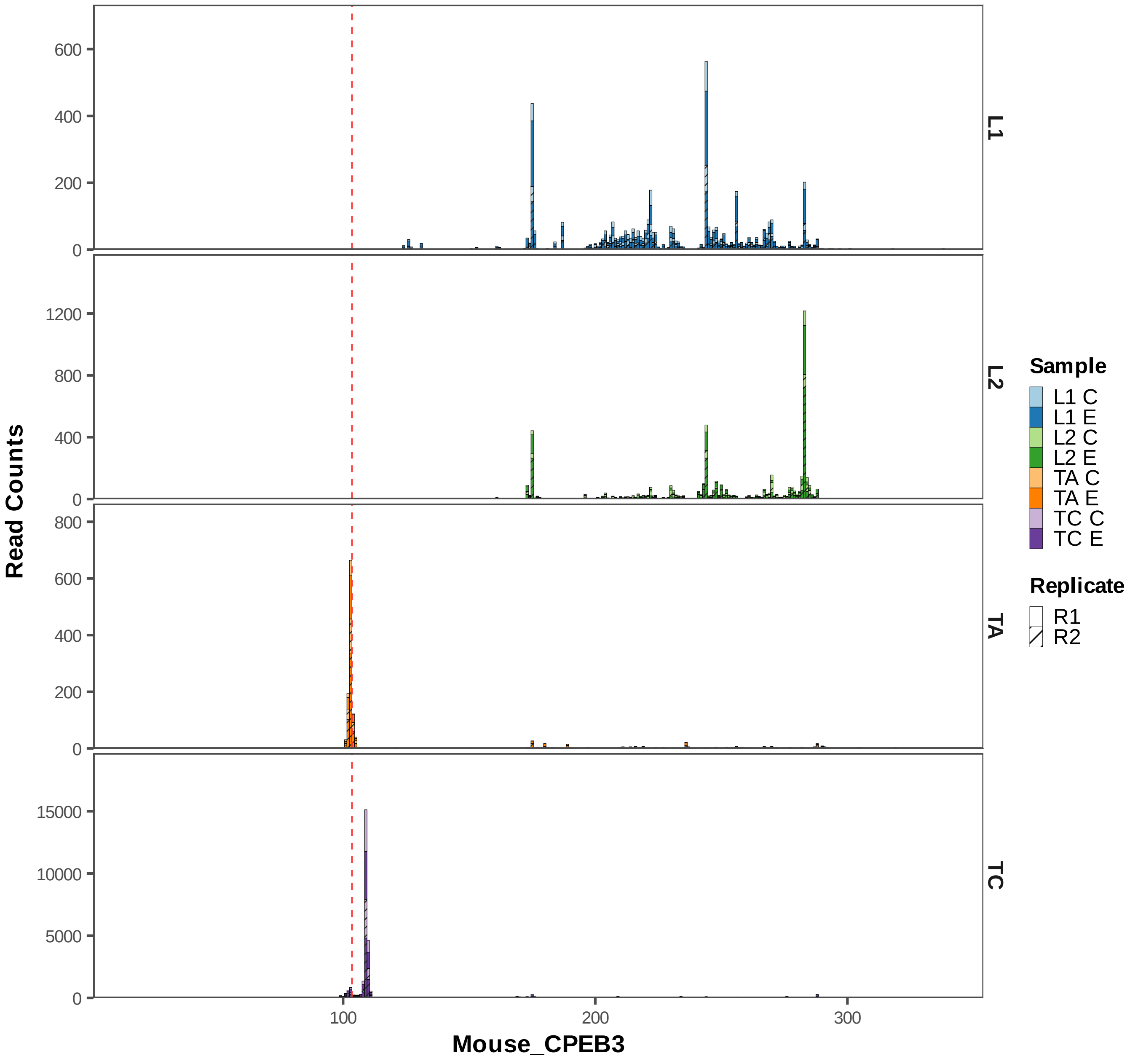

Read Counts
Mouse_CPEB3
L1
L2
TA
TC
Sample
L1 C
L1 E
L2 C
L2 E
TA C
TA E
TC C
TC E
Replicate
R1
R2

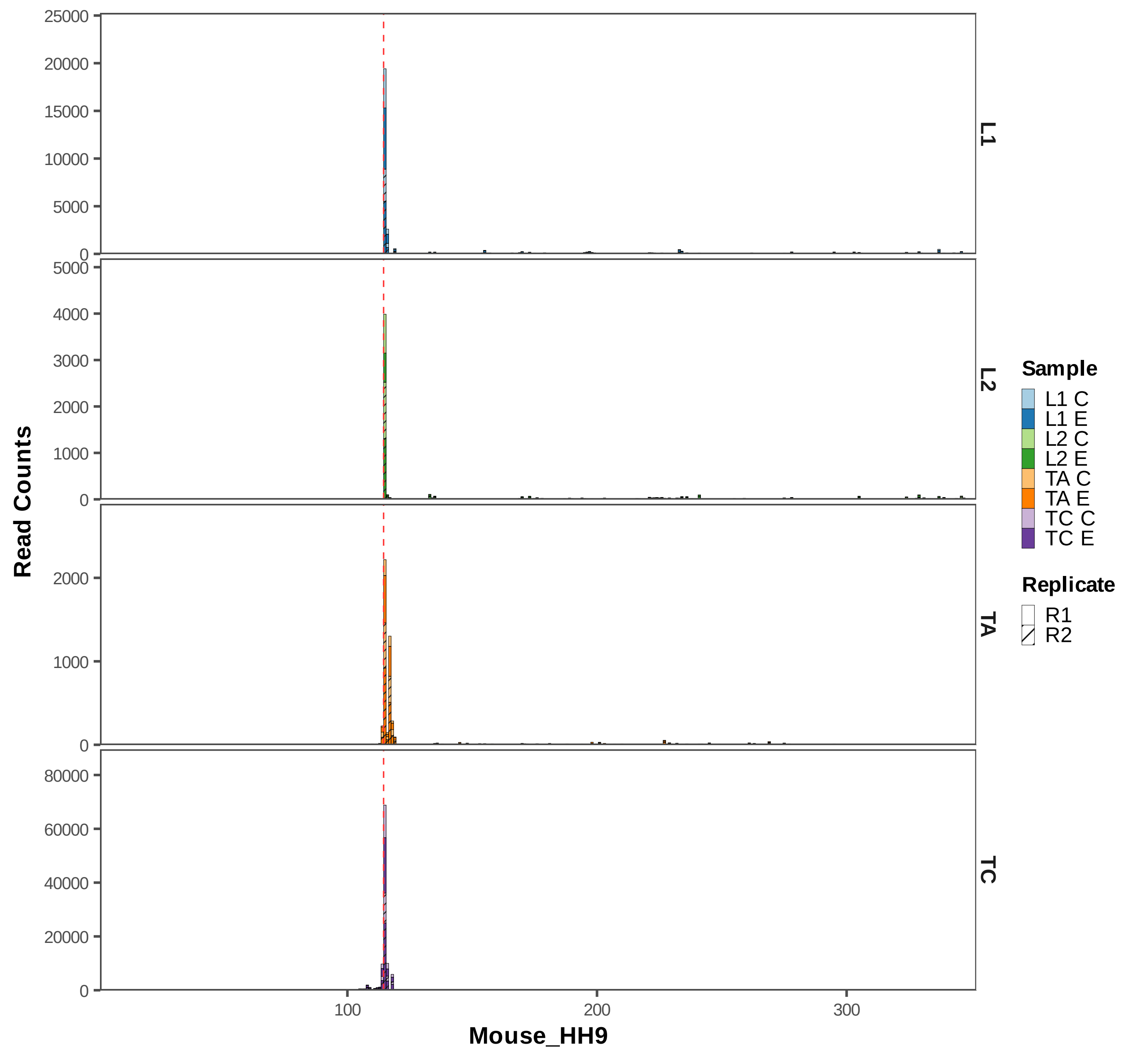
Read Counts
Mouse_HH9
L1
L2
TA
TC
0
5000
10000
15000
20000
25000
1000
2000
3000
4000
5000
20000
40000
60000
80000
100
200
300
Sample
L1 C
L1 E
L2 C
L2 E
TA C
TA E
TC C
TC E
Replicate
R1
R2

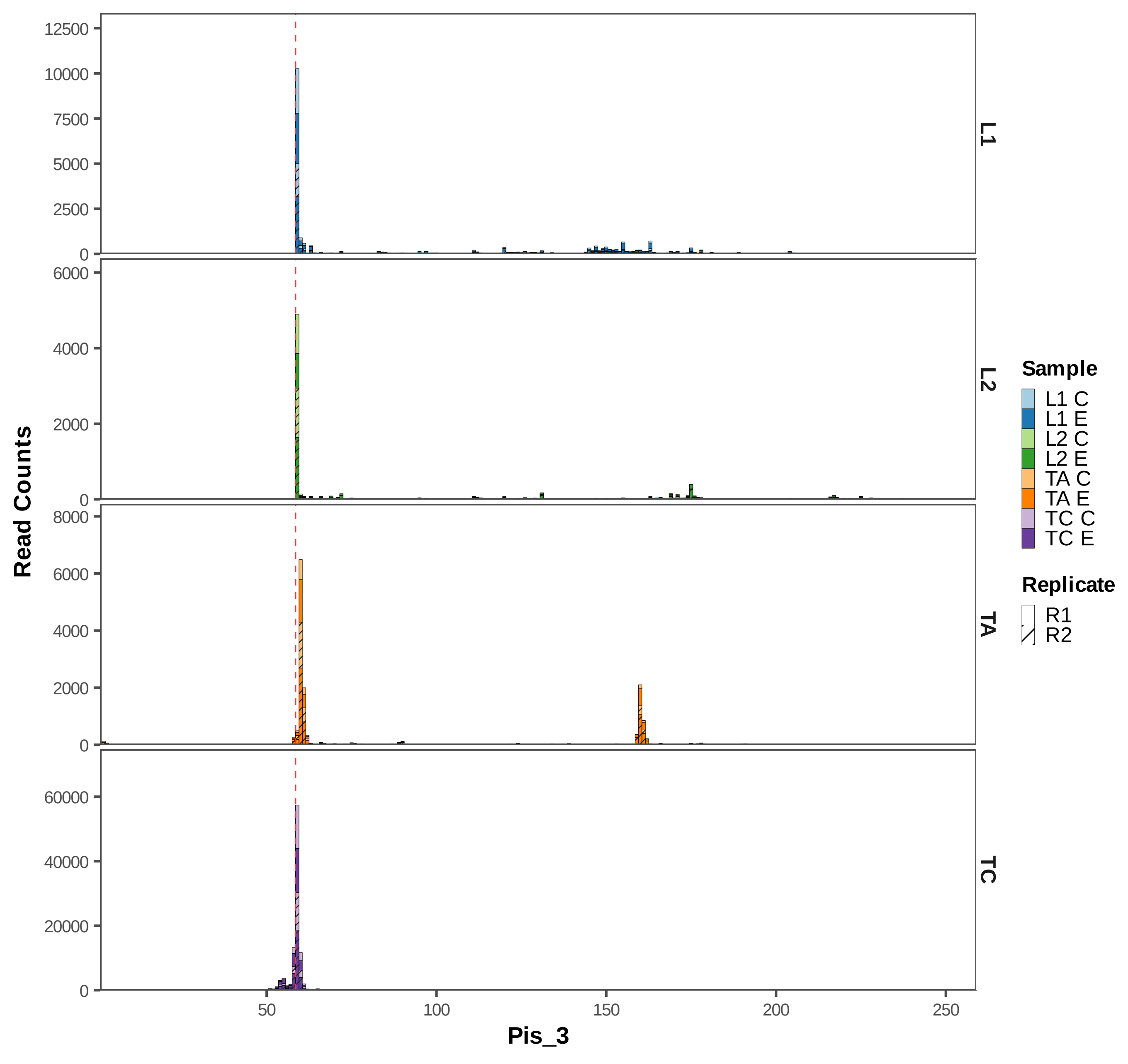
Read Counts
Pis_3
L1
L2
TA
TC
Sample
L1 C
L1 E
L2 C
L2 E
TA C
TA E
TC C
TC E
Replicate
R1
R2
12500
10000
7500
5000
2500
0
6000
4000
2000
0
8000
6000
4000
2000
0
60000
40000
20000
0
50
100
150
200
250

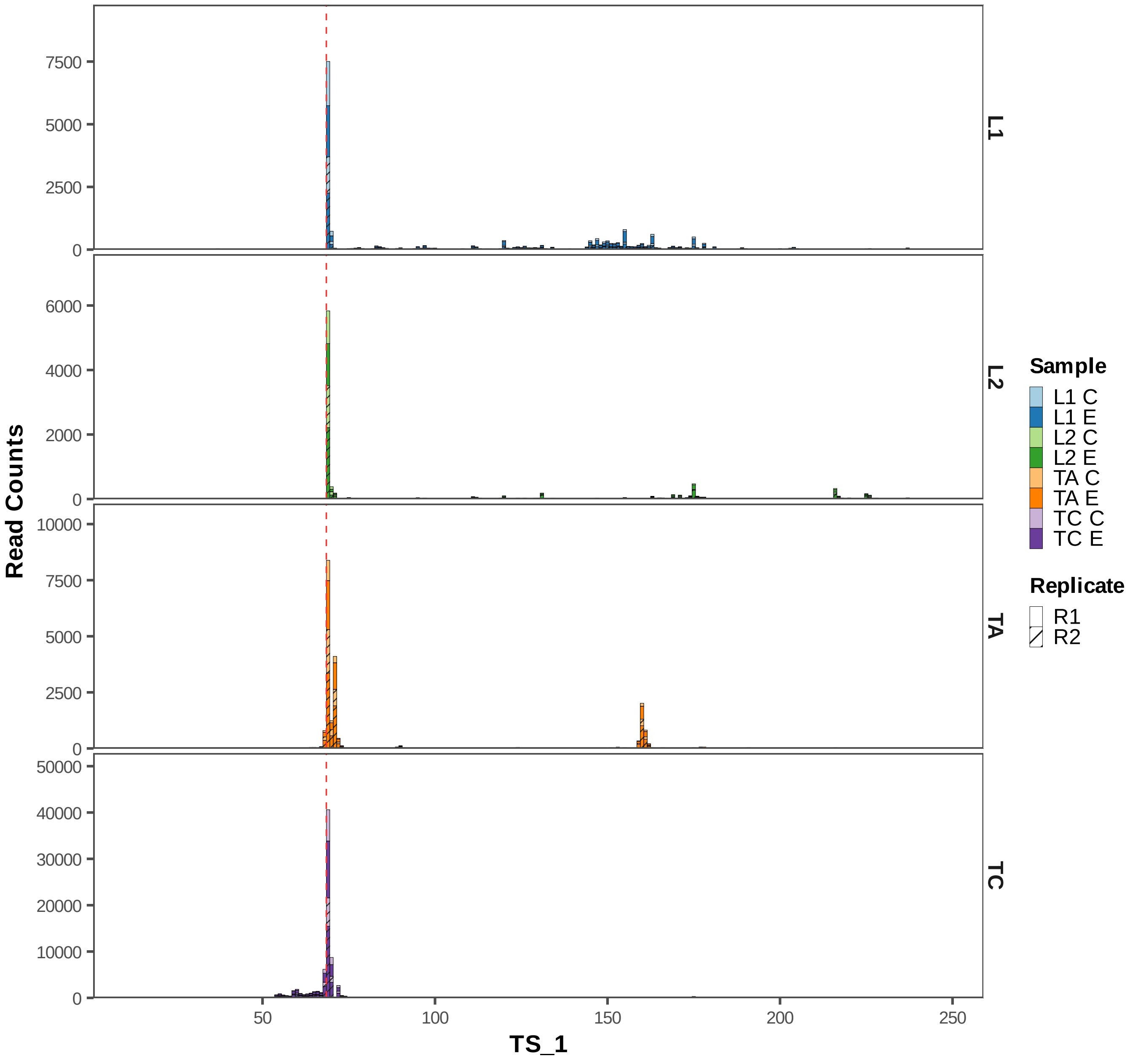

Read Counts
TS_1
L1
L2
TA
TC
0
2500
5000
7500
2000
4000
6000
10000
20000
30000
40000
50000
50
100
150
200
250
Sample
L1 C
L1 E
L2 C
L2 E
TA C
TA E
TC C
TC E
Replicate
R1
R2

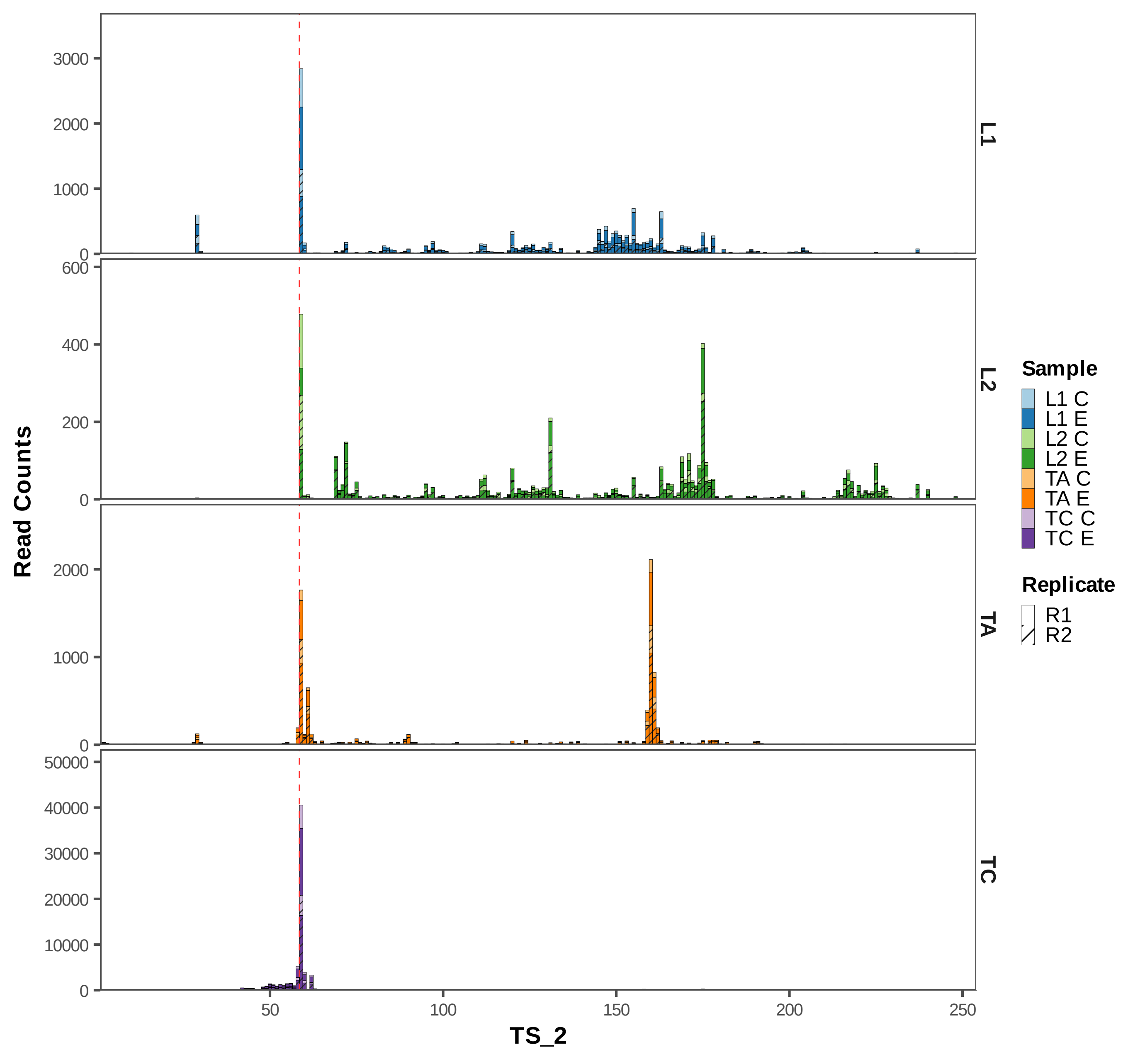

Read Counts
TS_2
L1
L2
TA
TC
3000
2000
1000
0
600
400
200
0
2000
1000
0
50000
40000
30000
20000
10000
0
50
100
150
200
250
Sample
L1 C
L1 E
L2 C
L2 E
TA C
TA E
TC C
TC E
Replicate
R1
R2

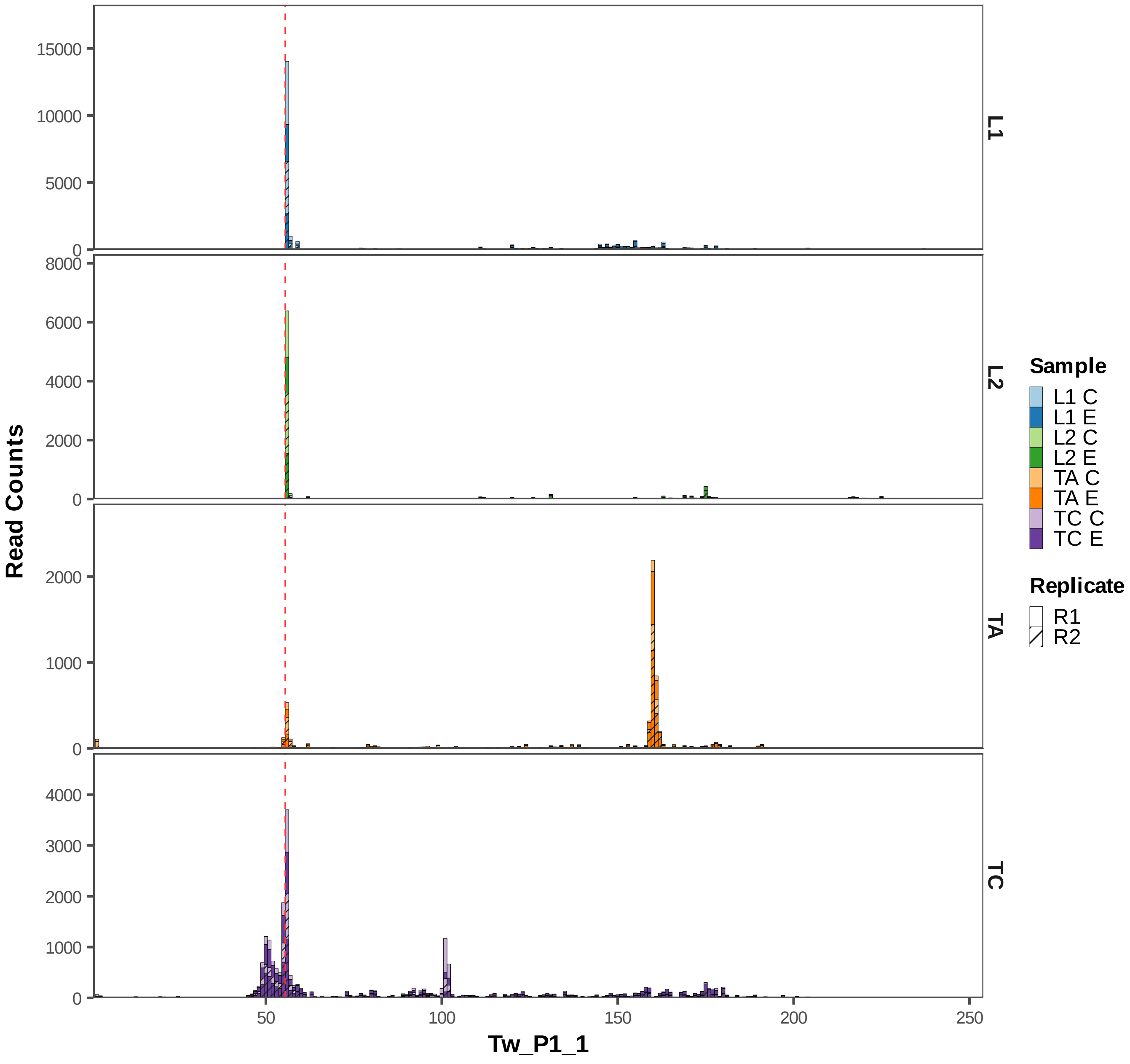

Read Counts
15000
10000
5000
0
8000
6000
4000
2000
0
2000
1000
0
4000
3000
2000
1000
0
L1
L2
TA
TC
50
100
150
200
250
Tw_P1_1
Sample
L1 C
L1 E
L2 C
L2 E
TA C
TA E
TC C
TC E
Replicate
R1
R2

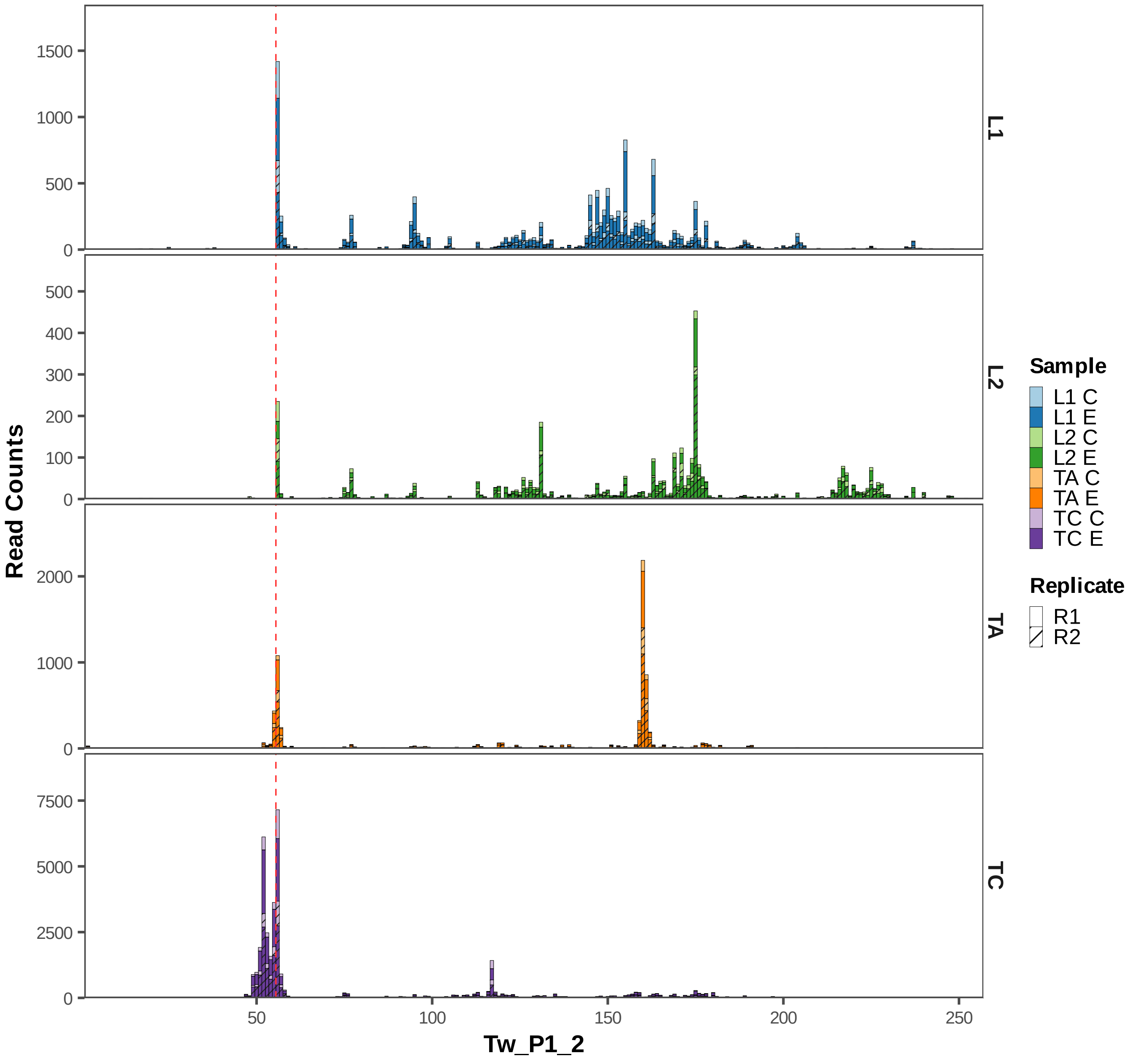

Read Counts
Tw_P1_2
L1
L2
TA
TC
Sample
L1 C
L1 E
L2 C
L2 E
TA C
TA E
TC C
TC E
Replicate
R1
R2

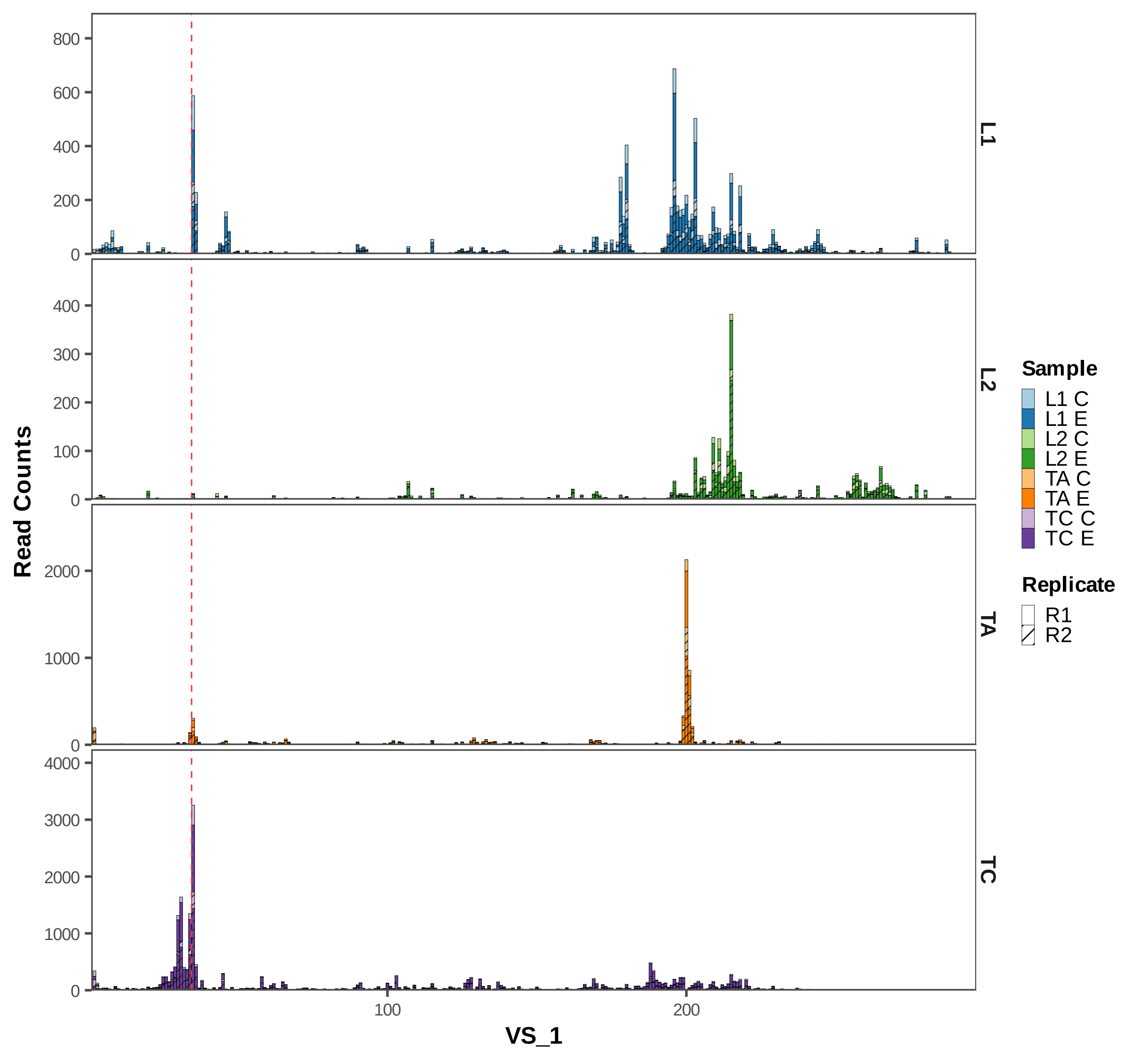
Read Counts
VS_1
L1
L2
TA
TC
Sample
L1 C
L1 E
L2 C
L2 E
TA C
TA E
TC C
TC E
Replicate
R1
R2
800
600
400
200
0
300
100
2000
1000
4000
3000
100
200
