## Supplementary Figures for "Widespread Presence of Catalytic RNAs in All Domains of Life": Figure S4.pdf

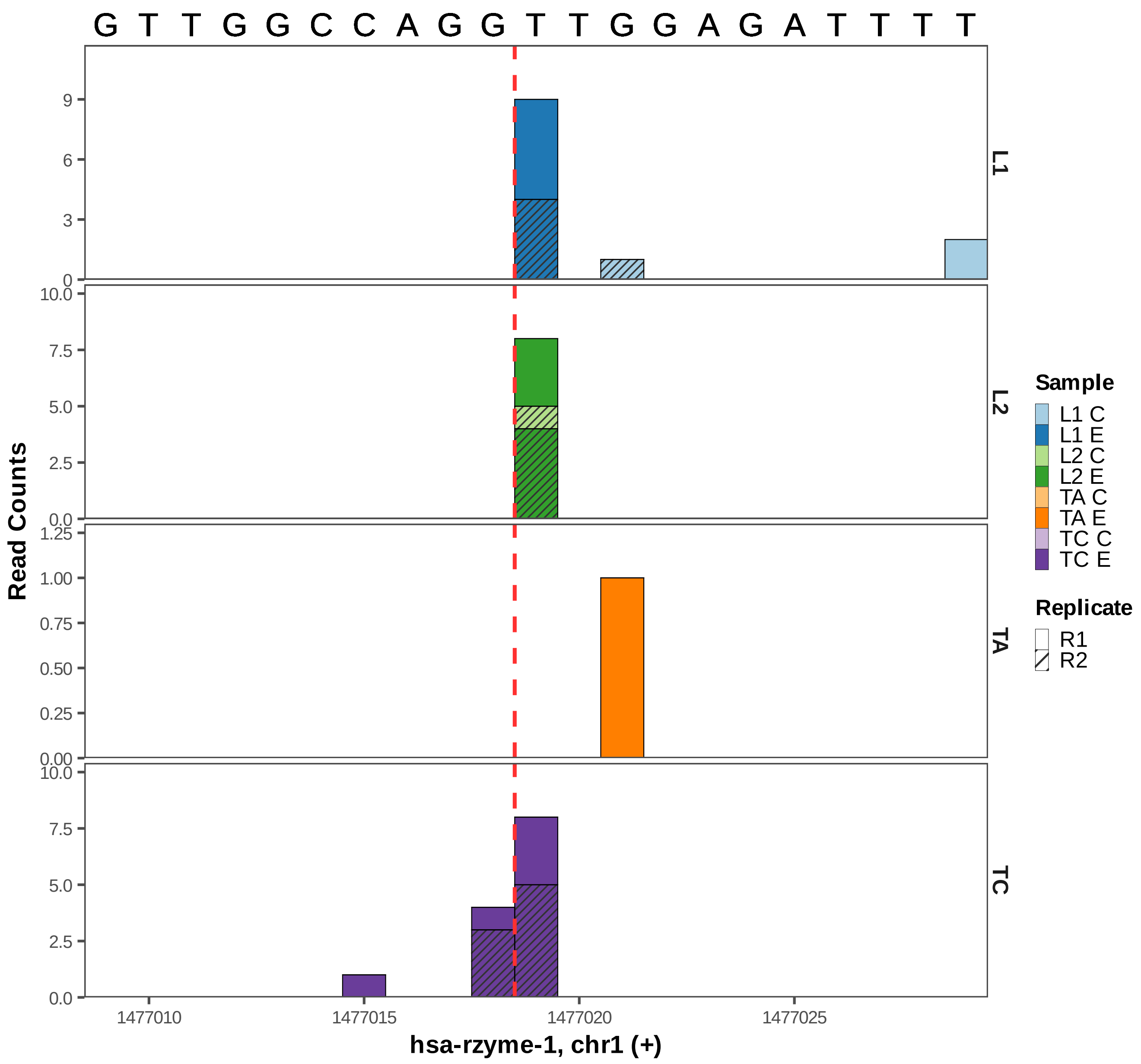
G T T G G C C A G G T T G G A G A T T T T
Read Counts
L1
L2
TA
TC
9
6
3
0
10.0
7.5
5.0
2.5
0.0
1.25
1.00
0.75
0.50
0.25
0.00
1477010
1477015
1477020
1477025
hsa-rzyme-1, chr1 (+)
Sample
L1 C
L1 E
L2 C
L2 E
TA C
TA E
TC C
TC E
Replicate
R1
R2

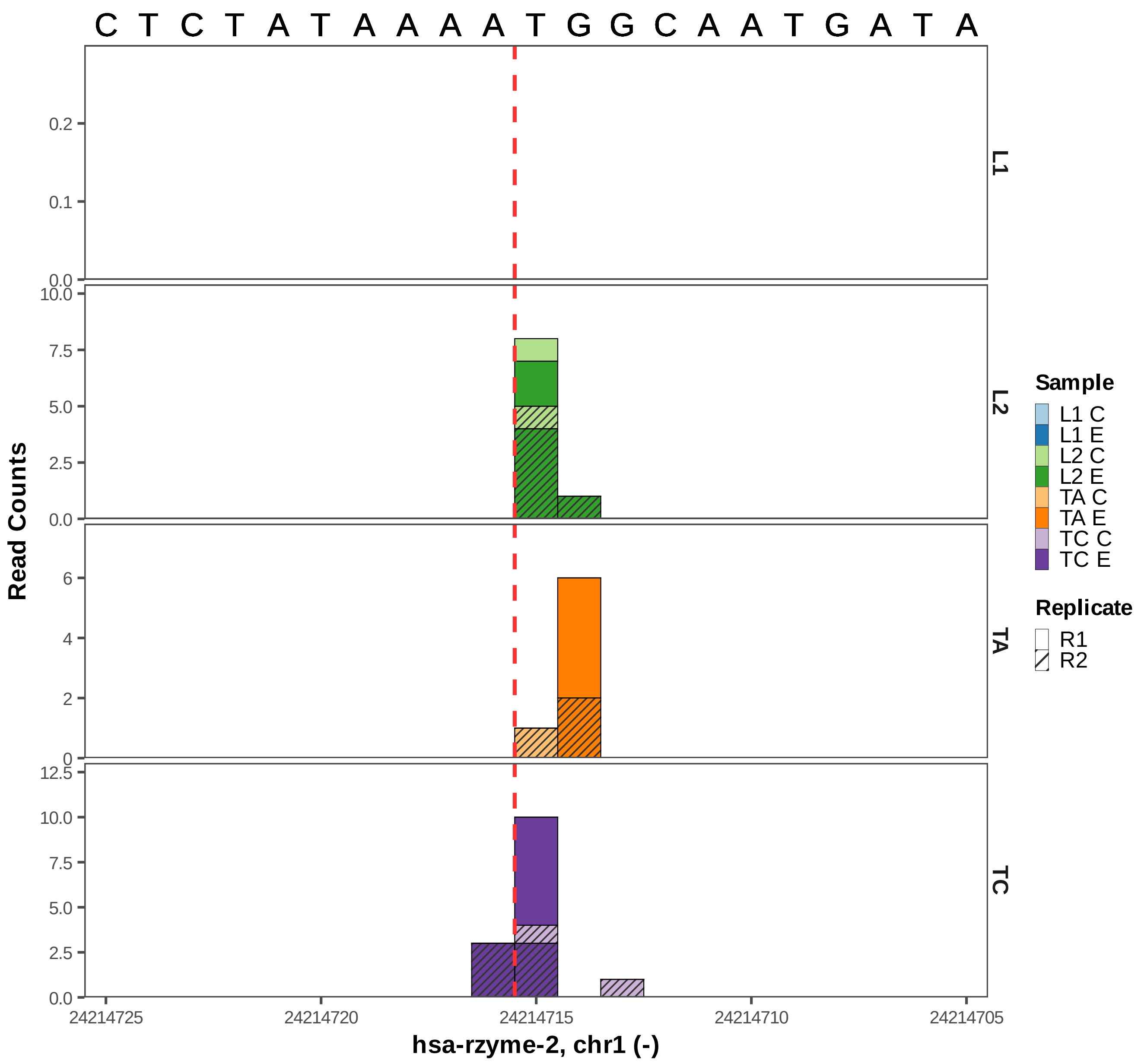
C T C T A T A A A A T G G C A A T G A T A
L1
L2
TA
TC
Read Counts
0.0
0.1
0.2
10.0
7.5
5.0
2.5
0
2
4
6
12.5
24214725
24214720
24214715
24214710
24214705
hsa-rzyme-2, chr1 (-)
Sample
L1 C
L1 E
L2 C
L2 E
TA C
TA E
TC C
TC E
Replicate
R1
R2

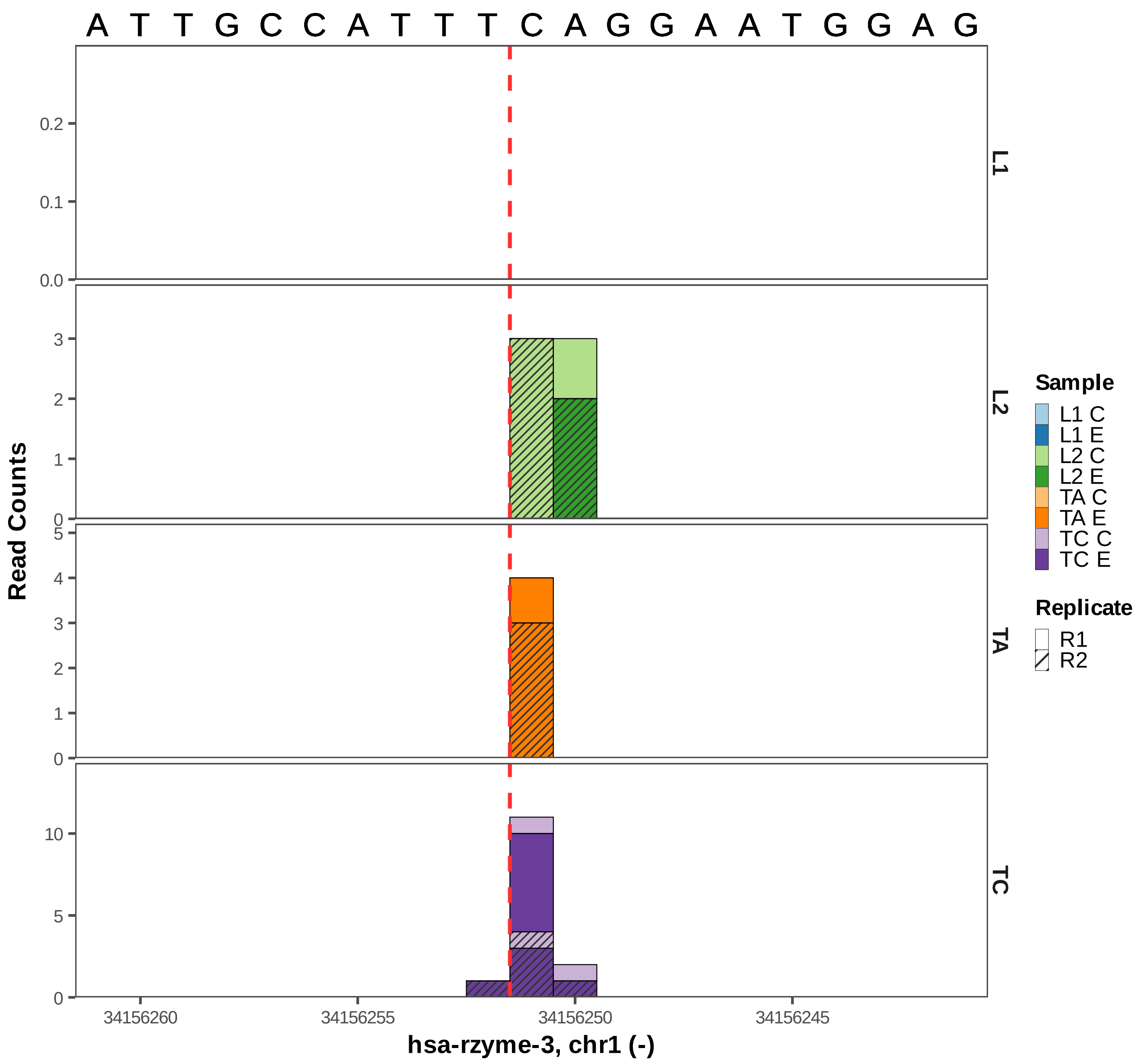

A T T G C C A T T T C A G G A A T G G A G
Read Counts
L1
L2
TA
TC
0.0
0.1
0.2
0
1
2
3
4
5
10
34156260
34156255
34156250
34156245
hsa-rzyme-3, chr1 (-)
Sample
L1 C
L1 E
L2 C
L2 E
TA C
TA E
TC C
TC E
Replicate
R1
R2

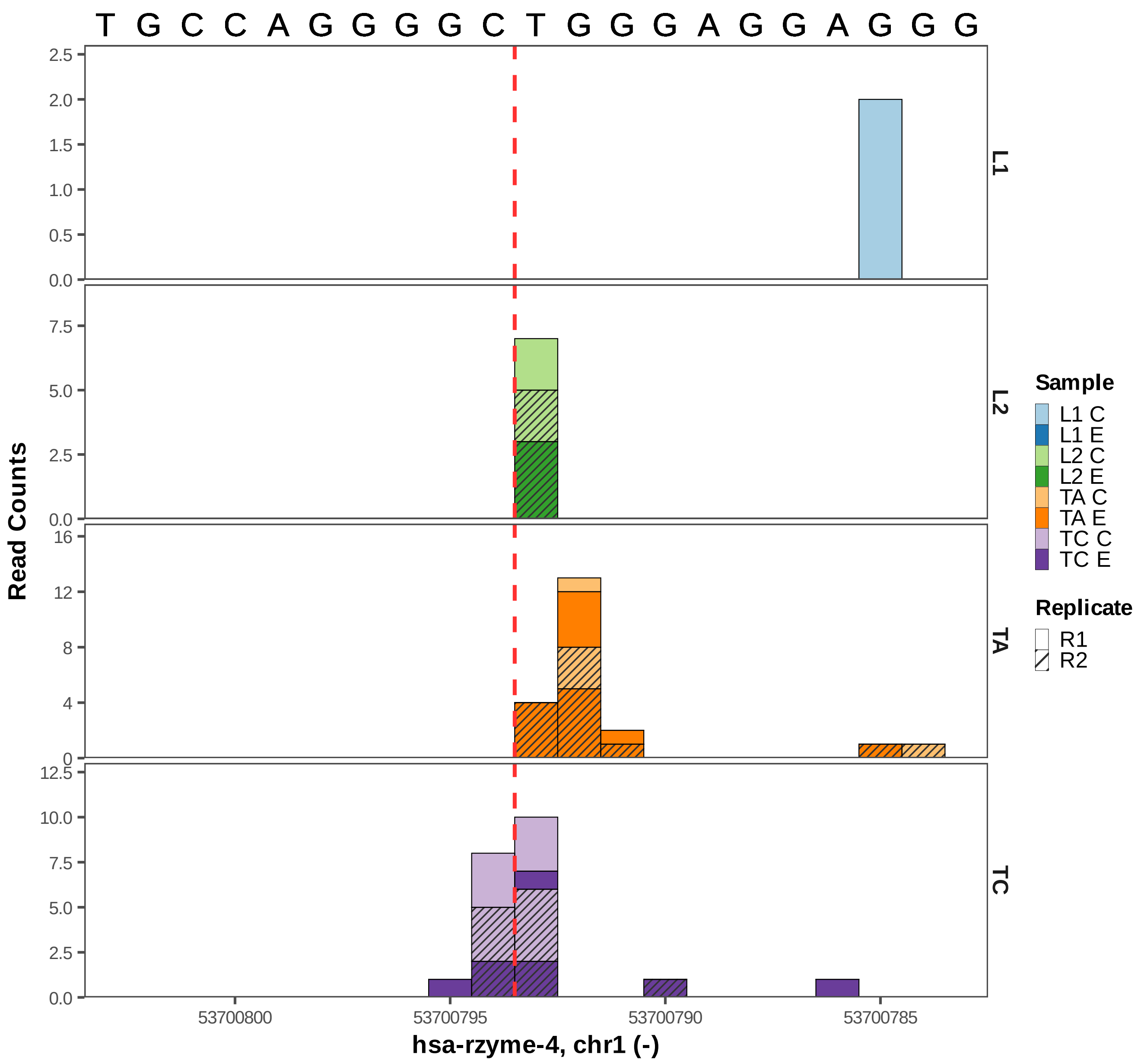

T G C C A G G G G C T G G G A G G A G G G
Read Counts
L1
L2
TA
TC
53700800
53700795
53700790
53700785
hsa-rzyme-4, chr1 (-)
Sample
L1 C
L1 E
L2 C
L2 E
TA C
TA E
TC C
TC E
Replicate
R1
R2

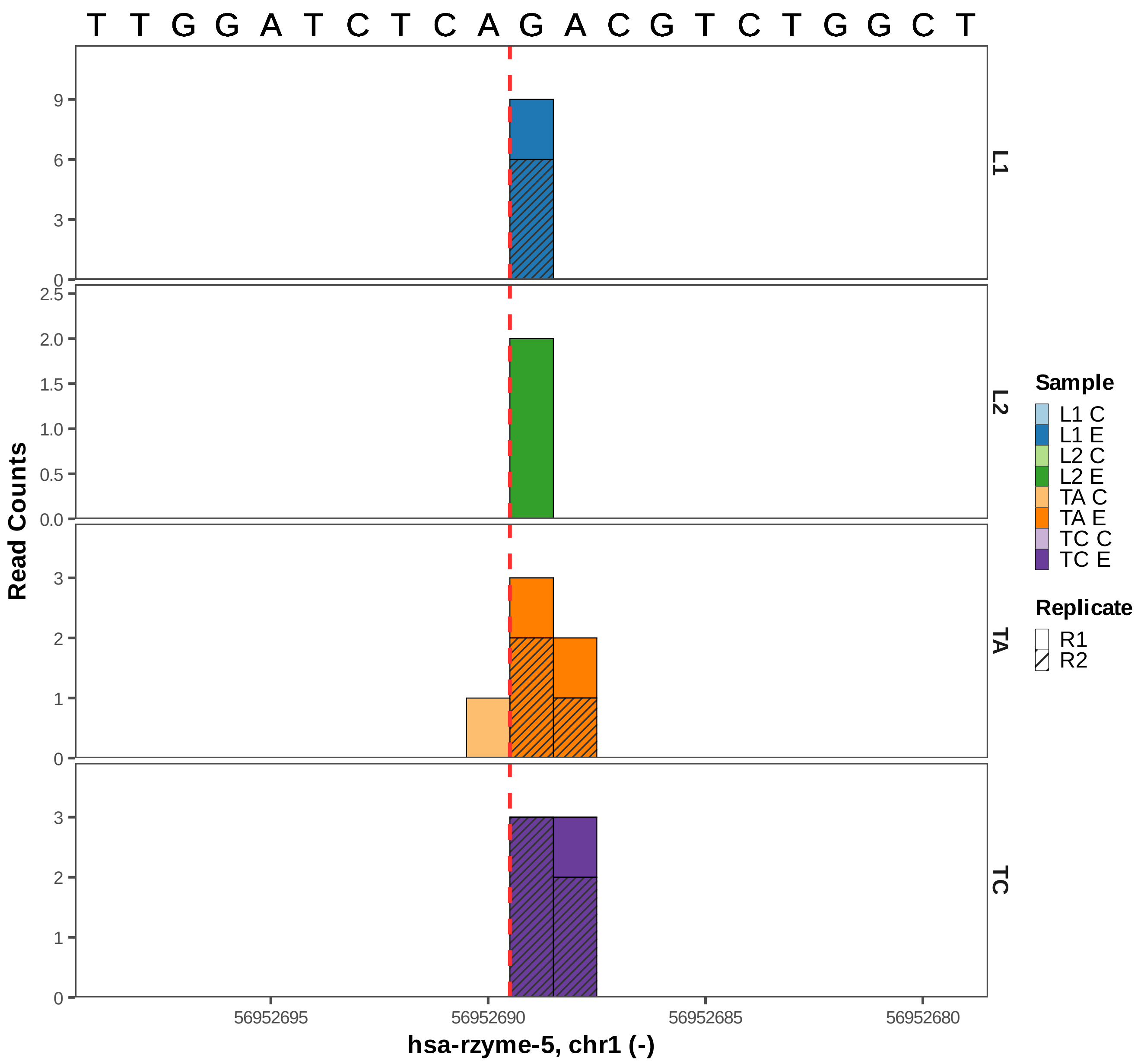

T T G G A T C T C A G A C G T C T G G C T
L1
L2
TA
TC
Read Counts
hsa-rzyme-5, chr1 (-)
56952695
56952690
56952685
56952680
Sample
L1 C
L1 E
L2 C
L2 E
TA C
TA E
TC C
TC E
Replicate
R1
R2

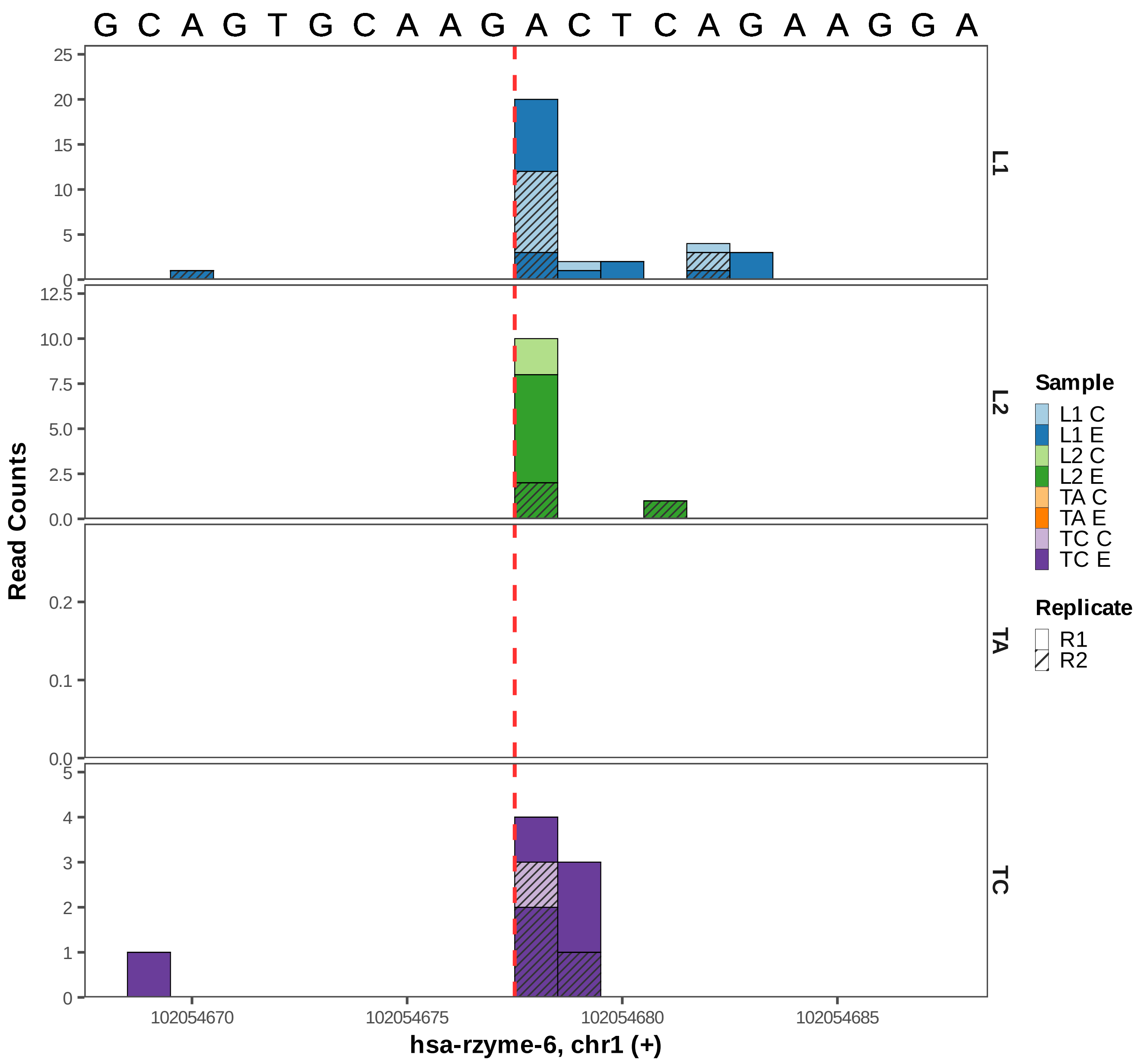

G C A G T G C A A G A C T C A G A A G G A
Read Counts
L1
L2
TA
TC
102054670
102054675
102054680
102054685
hsa-rzyme-6, chr1 (+)
Sample
L1 C
L1 E
L2 C
L2 E
TA C
TA E
TC C
TC E
Replicate
R1
R2

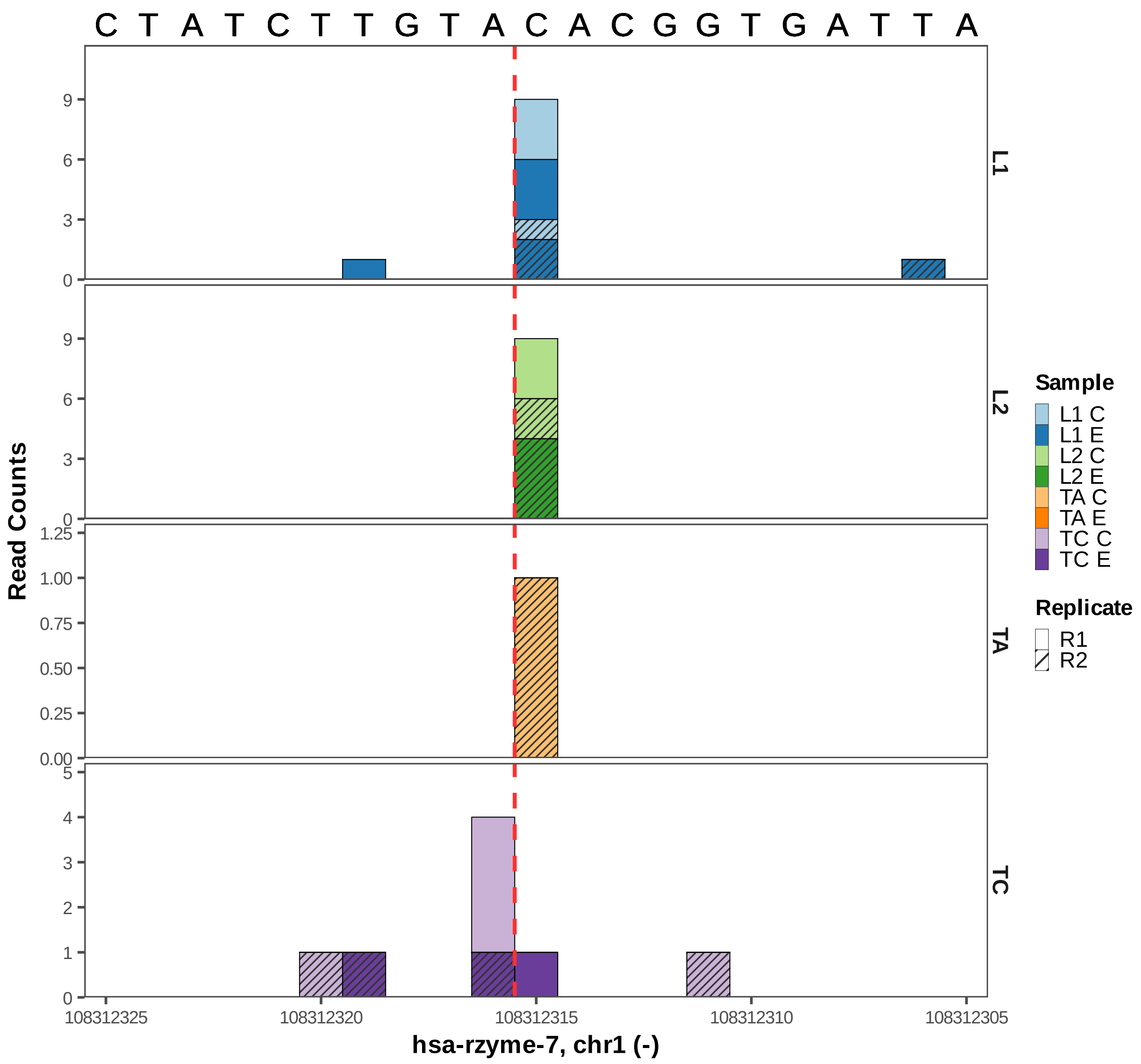

C T A T C T T G T A C A C G G T G A T T A
L1
L2
TA
TC
Read Counts
hsa-rzyme-7, chr1 (-)
108312325
108312320
108312315
108312310
108312305
Sample
L1 C
L1 E
L2 C
L2 E
TA C
TA E
TC C
TC E
Replicate
R1
R2

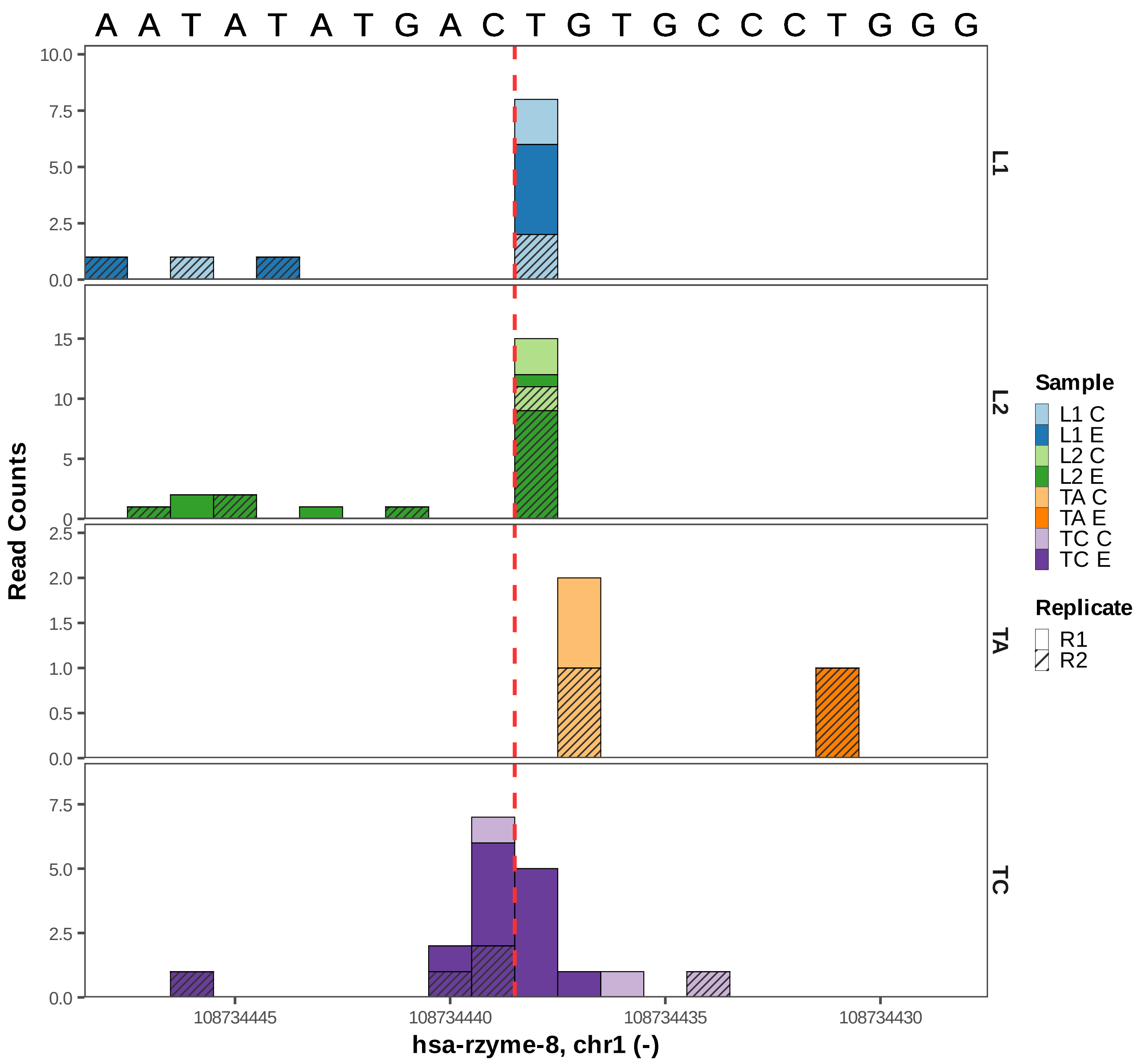
A A T A T A T G A C T G T G C C C T G G G
Read Counts
L1
L2
TA
TC
108734445
108734440
108734435
108734430
hsa-rzyme-8, chr1 (-)
Sample
L1 C
L1 E
L2 C
L2 E
TA C
TA E
TC C
TC E
Replicate
R1
R2

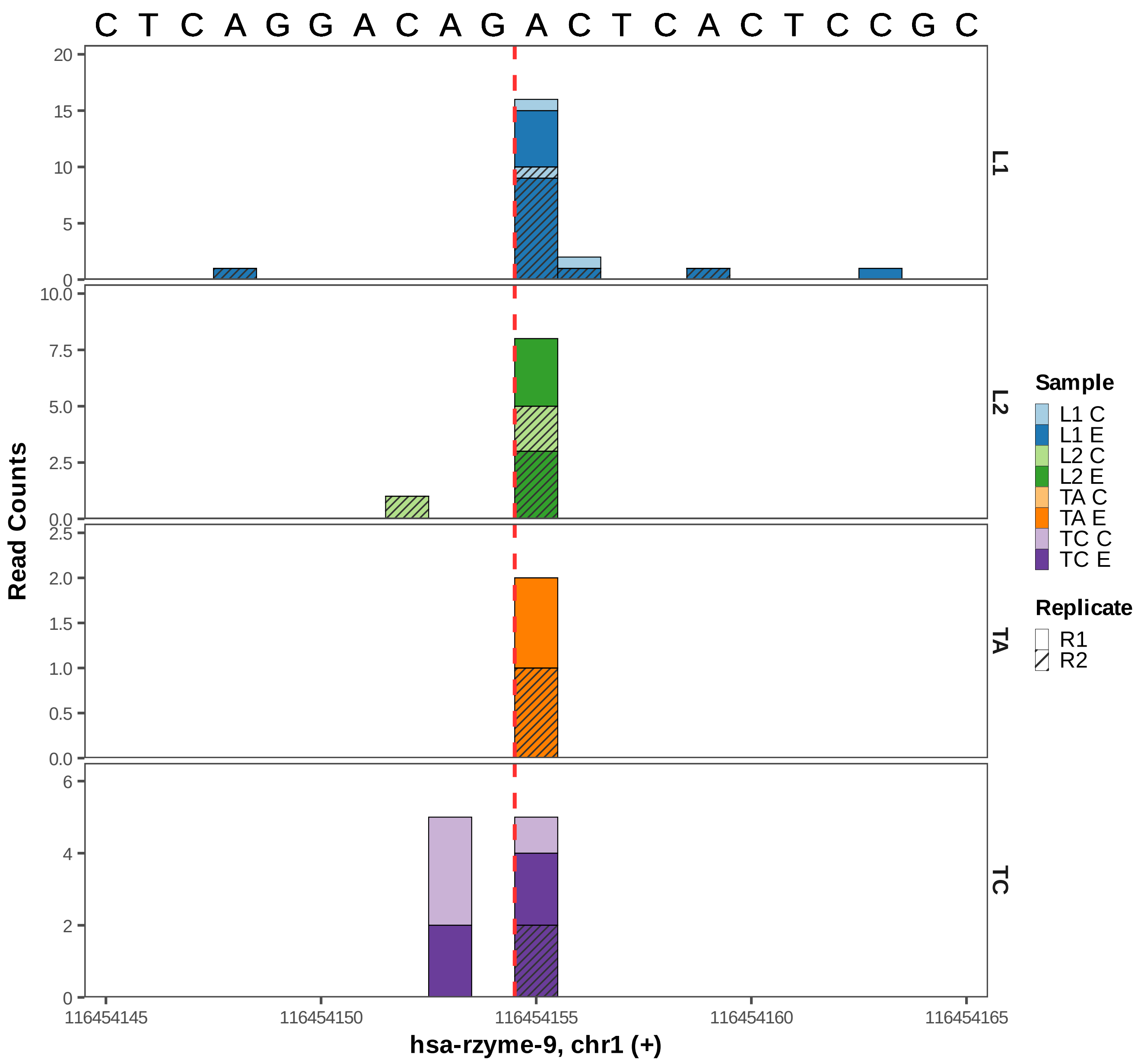

C T C A G G A C A G A C T C A C T C C G C
Read Counts
L1
L2
TA
TC
116454145
116454150
116454155
116454160
116454165
hsa-rzyme-9, chr1 (+)
Sample
L1 C
L1 E
L2 C
L2 E
TA C
TA E
TC C
TC E
Replicate
R1
R2

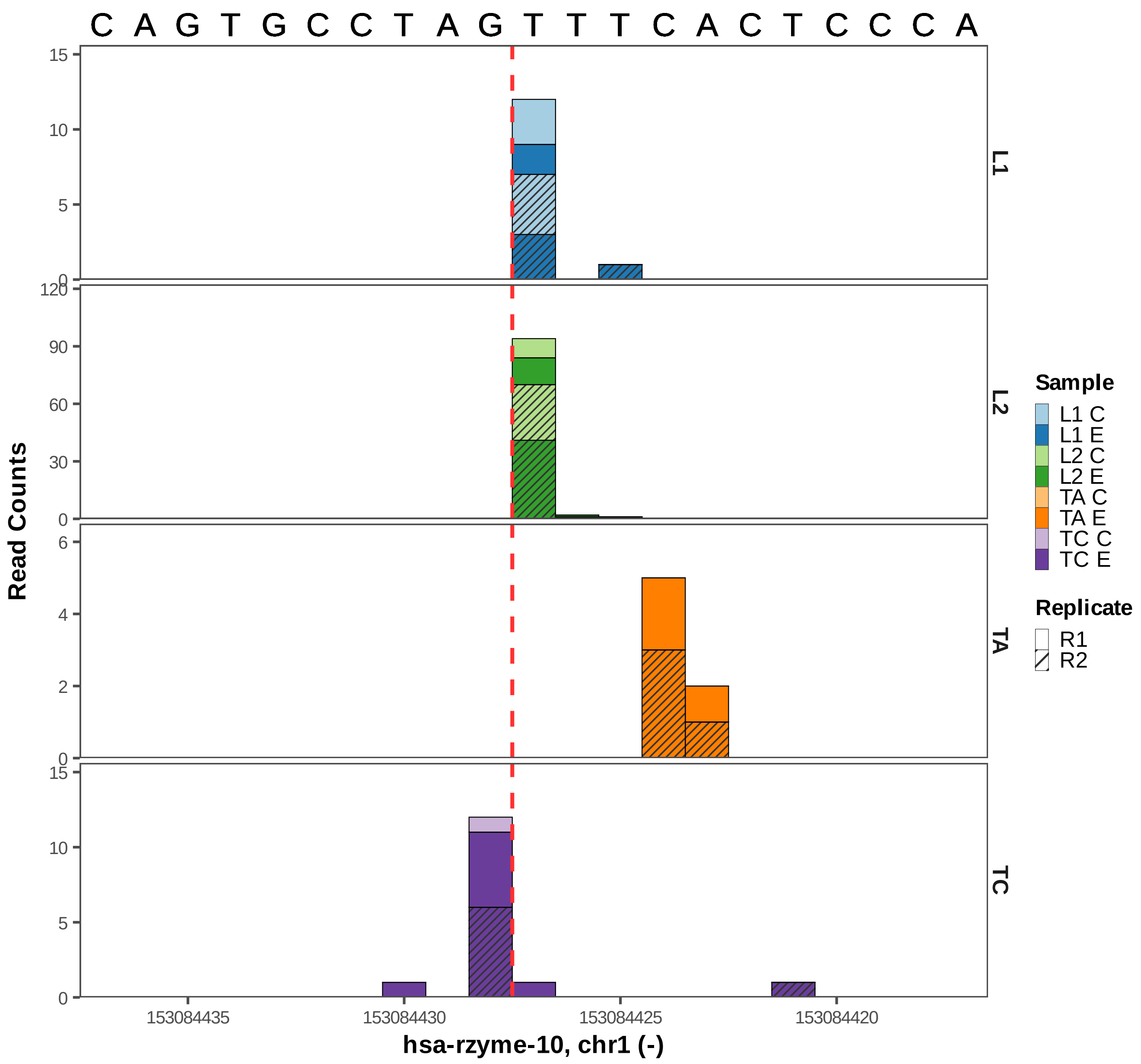

C A G T G C C T A G T T T C A C T C C C A
Read Counts
L1
L2
TA
TC
15
10
5
0
120
90
60
30
6
4
2
153084435
153084430
153084425
153084420
hsa-rzyme-10, chr1 (-)
Sample
L1 C
L1 E
L2 C
L2 E
TA C
TA E
TC C
TC E
Replicate
R1
R2

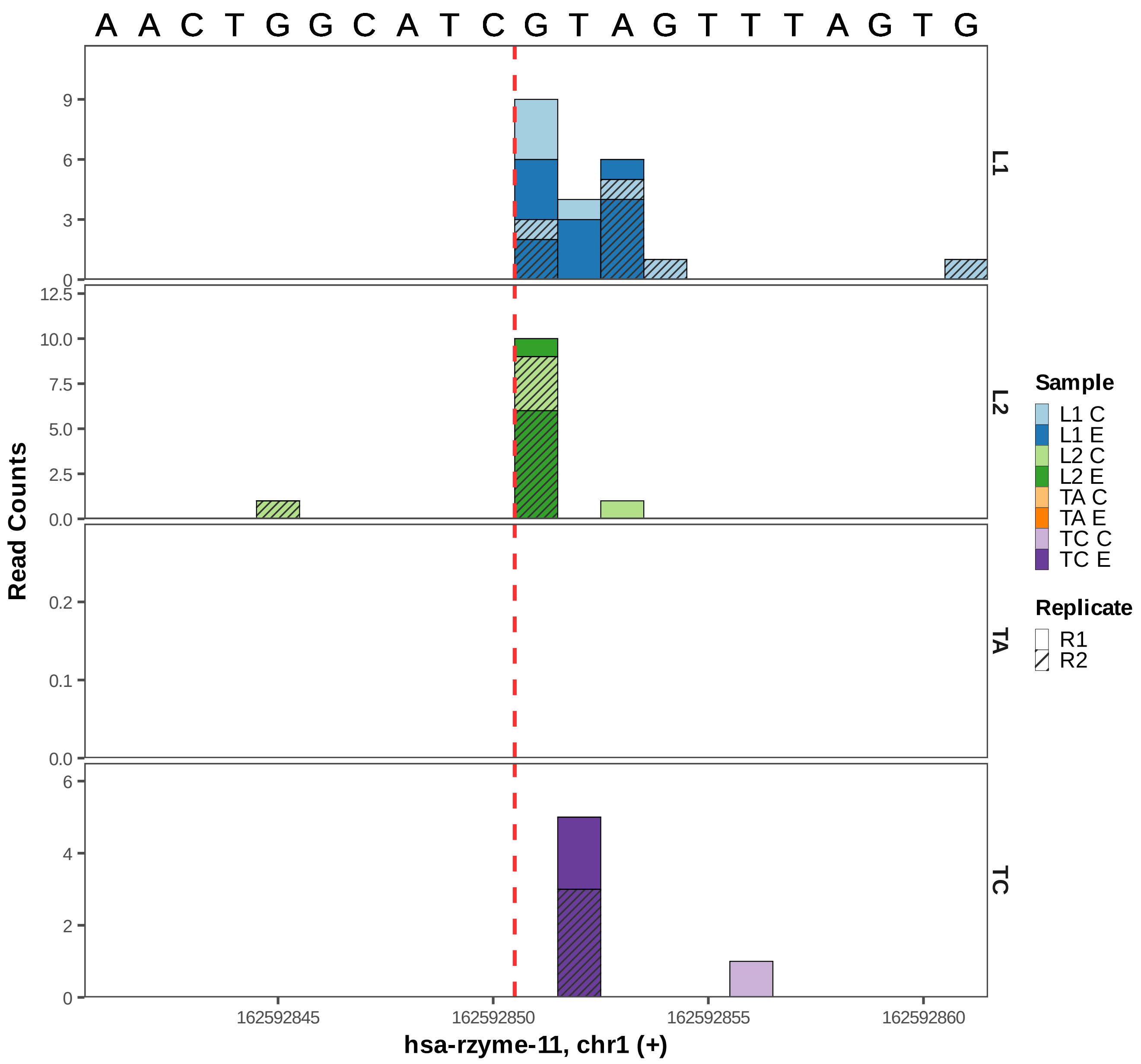

A A C T G G C A T C G T A G T T T A G T G
L1
L2
TA
TC
Read Counts
hsa-rzyme-11, chr1 (+)
162592845
162592850
162592855
162592860
Sample
L1 C
L1 E
L2 C
L2 E
TA C
TA E
TC C
TC E
Replicate
R1
R2

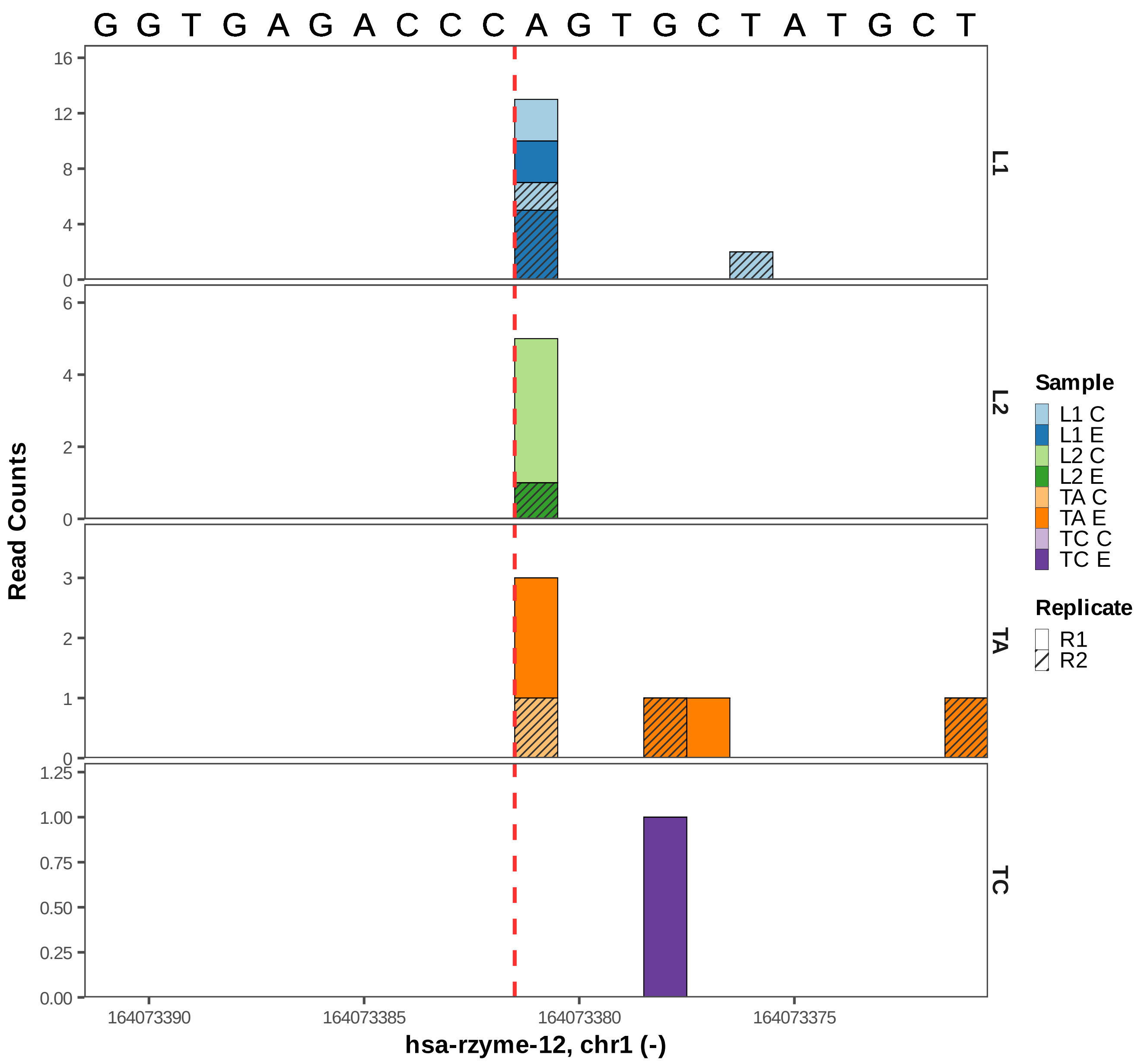
G G T G A G A C C C A G T G C T A T G C T
Read Counts
L1
L2
TA
TC
16
12
8
4
0
6
4
2
0
3
2
1
0
1.25
1.00
0.75
0.50
0.25
0.00
164073390
164073385
164073380
164073375
hsa-rzyme-12, chr1 (-)
Sample
L1 C
L1 E
L2 C
L2 E
TA C
TA E
TC C
TC E
Replicate
R1
R2

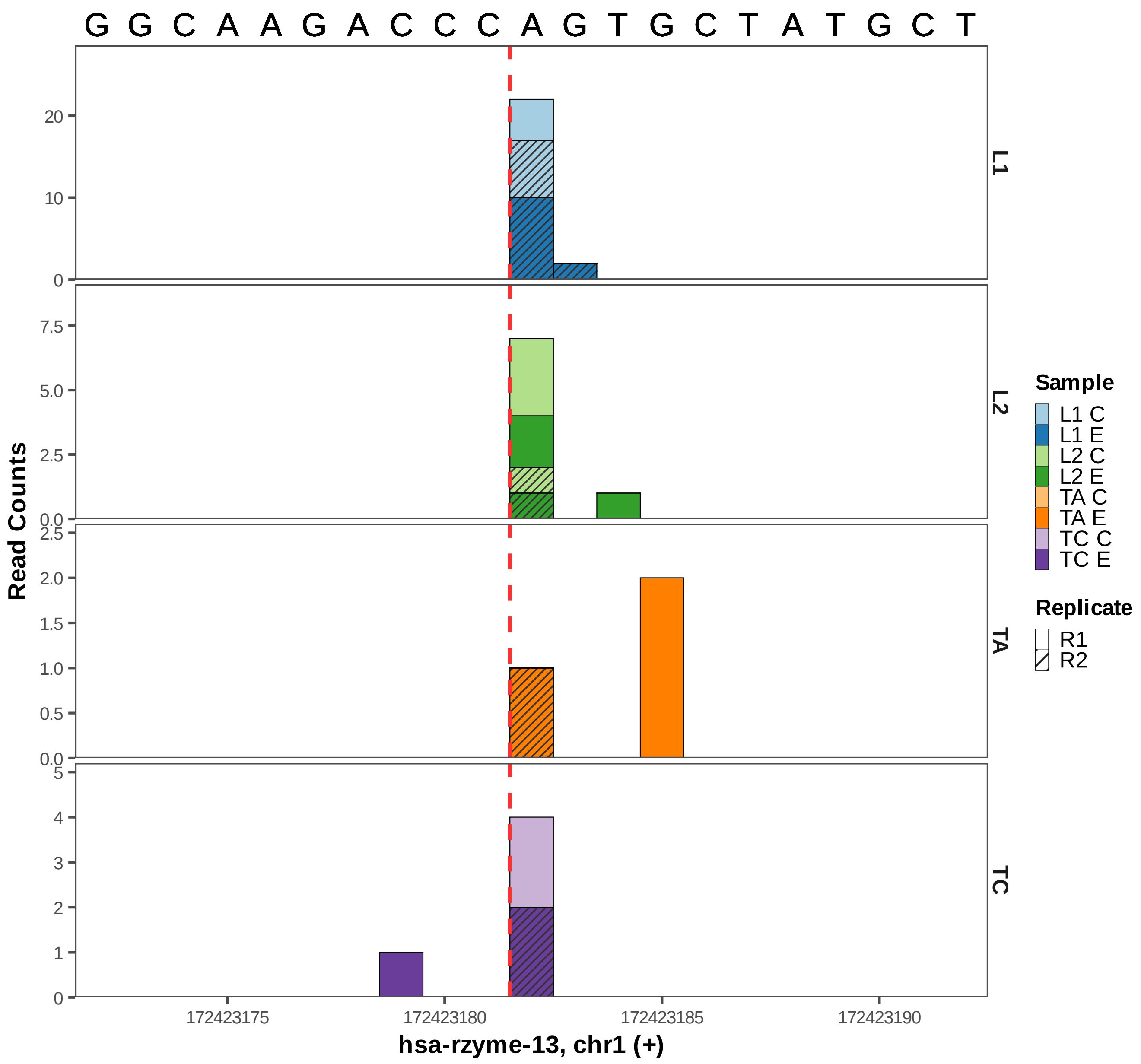

G G C A A G A C C C A G T G C T A T G C T
L1
L2
TA
TC
Read Counts
20
10
0
7.5
5.0
2.5
0.0
2.0
1.5
1.0
0.5
5
4
3
2
1
172423175
172423180
172423185
172423190
hsa-rzyme-13, chr1 (+)
Sample
L1 C
L1 E
L2 C
L2 E
TA C
TA E
TC C
TC E
Replicate
R1
R2

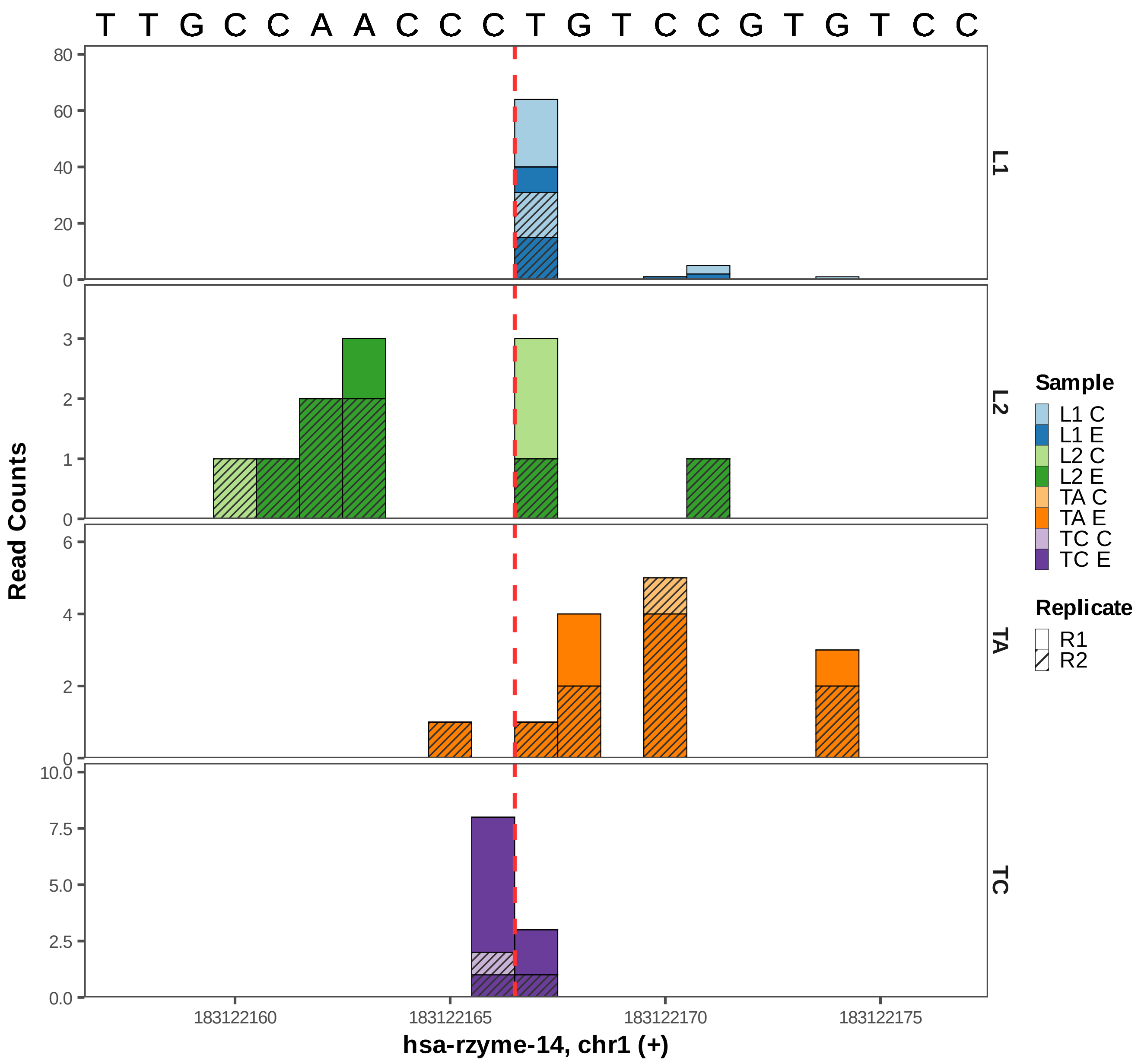

T T G C C A A C C C T G T C C G T G T C C
Read Counts
L1
L2
TA
TC
80
60
40
20
0
3
2
1
0
6
4
2
0
10.0
7.5
5.0
2.5
0.0
183122160
183122165
183122170
183122175
hsa-rzyme-14, chr1 (+)
Sample
L1 C
L1 E
L2 C
L2 E
TA C
TA E
TC C
TC E
Replicate
R1
R2

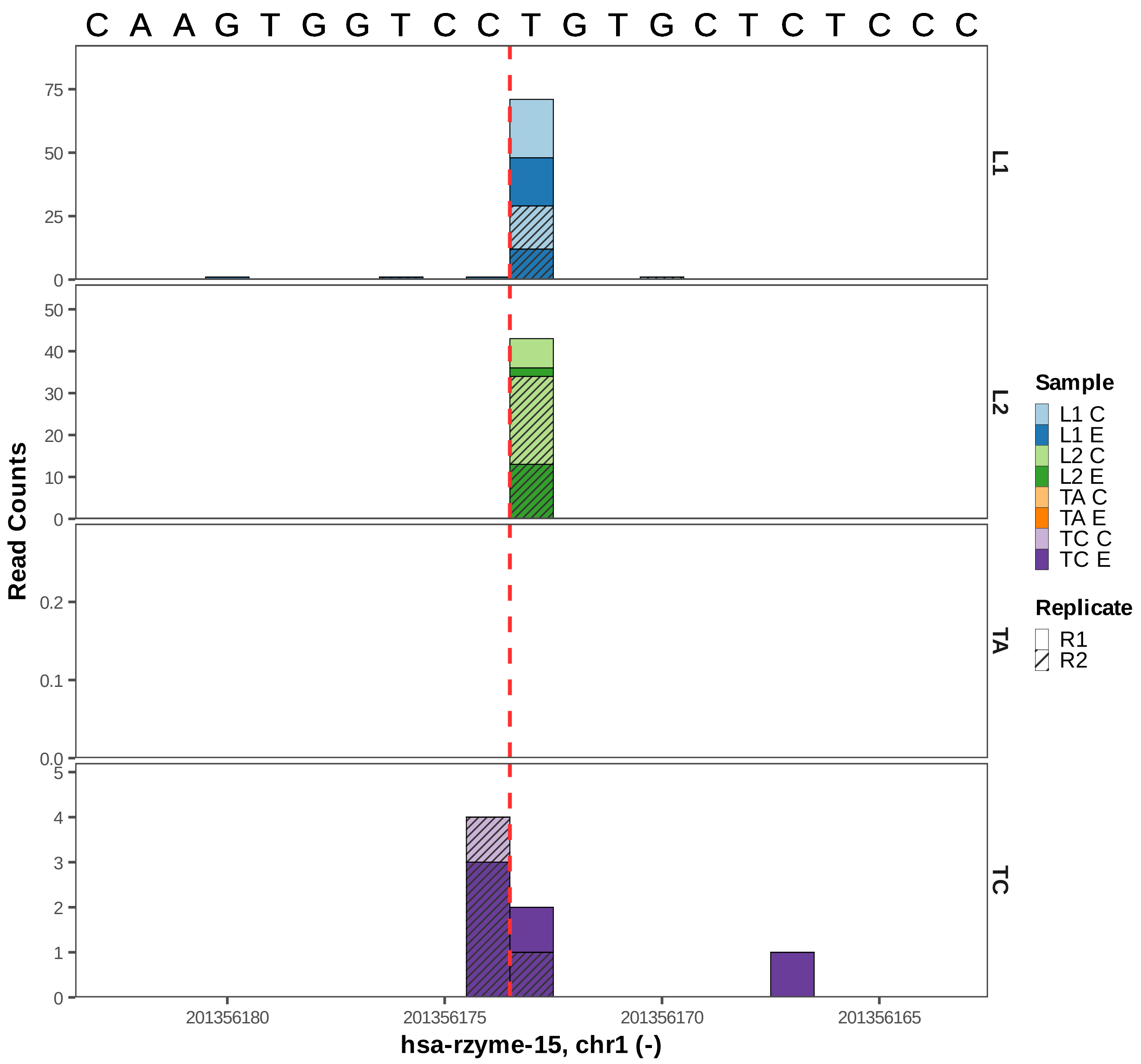
C A A G T G G T C C T G T G C T C T C C C
L1
L2
TA
TC
Read Counts
Sample
L1 C
L1 E
L2 C
L2 E
TA C
TA E
TC C
TC E
Replicate
R1
R2
201356180
201356175
201356170
201356165
hsa-rzyme-15, chr1 (-)

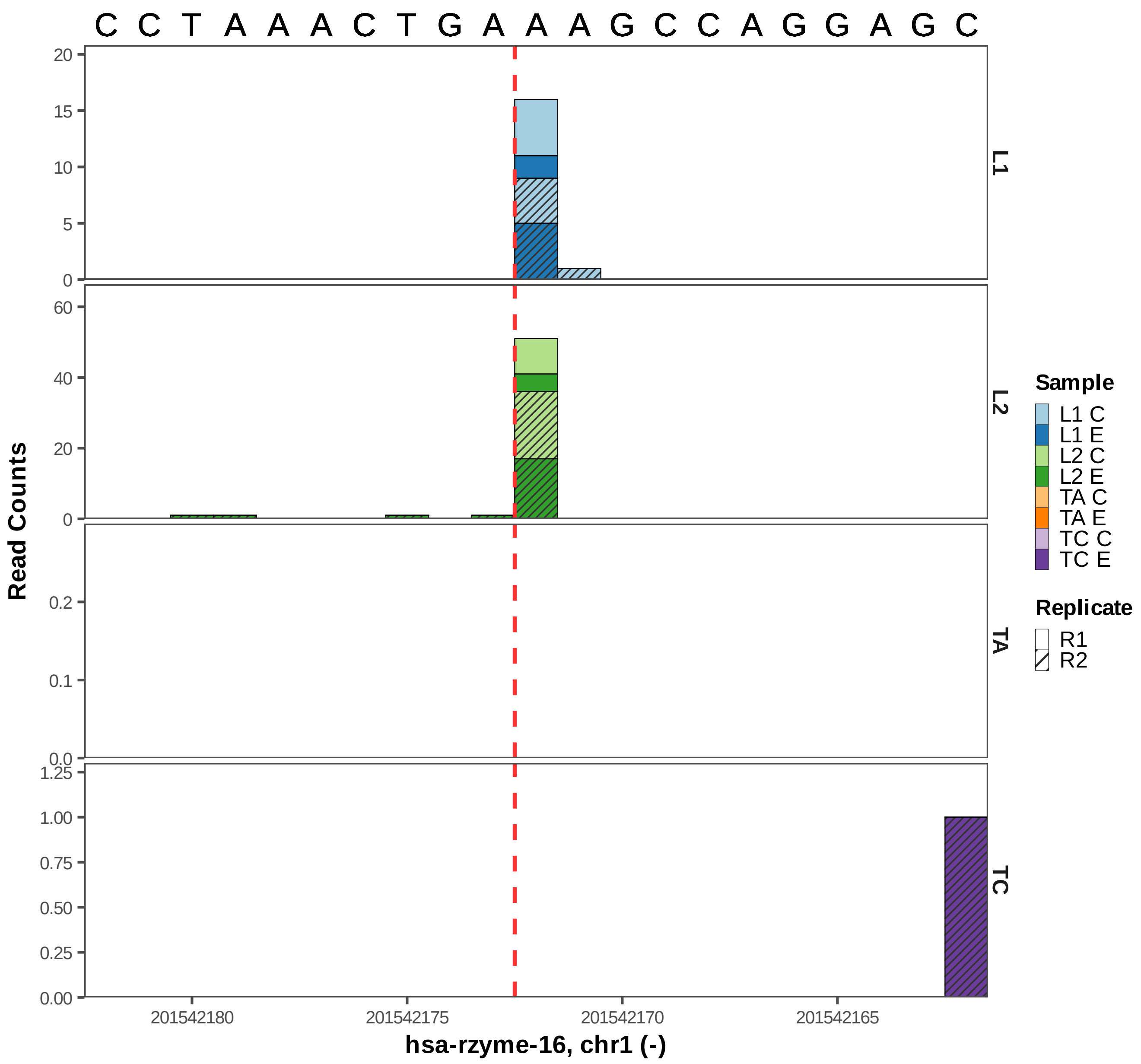
C C T A A A C T G A A A G C C A G G A G C
Read Counts
L1
L2
TA
TC
Sample
L1 C
L1 E
L2 C
L2 E
TA C
TA E
TC C
TC E
Replicate
R1
R2
201542180
201542175
201542170
201542165
hsa-rzyme-16, chr1 (-)

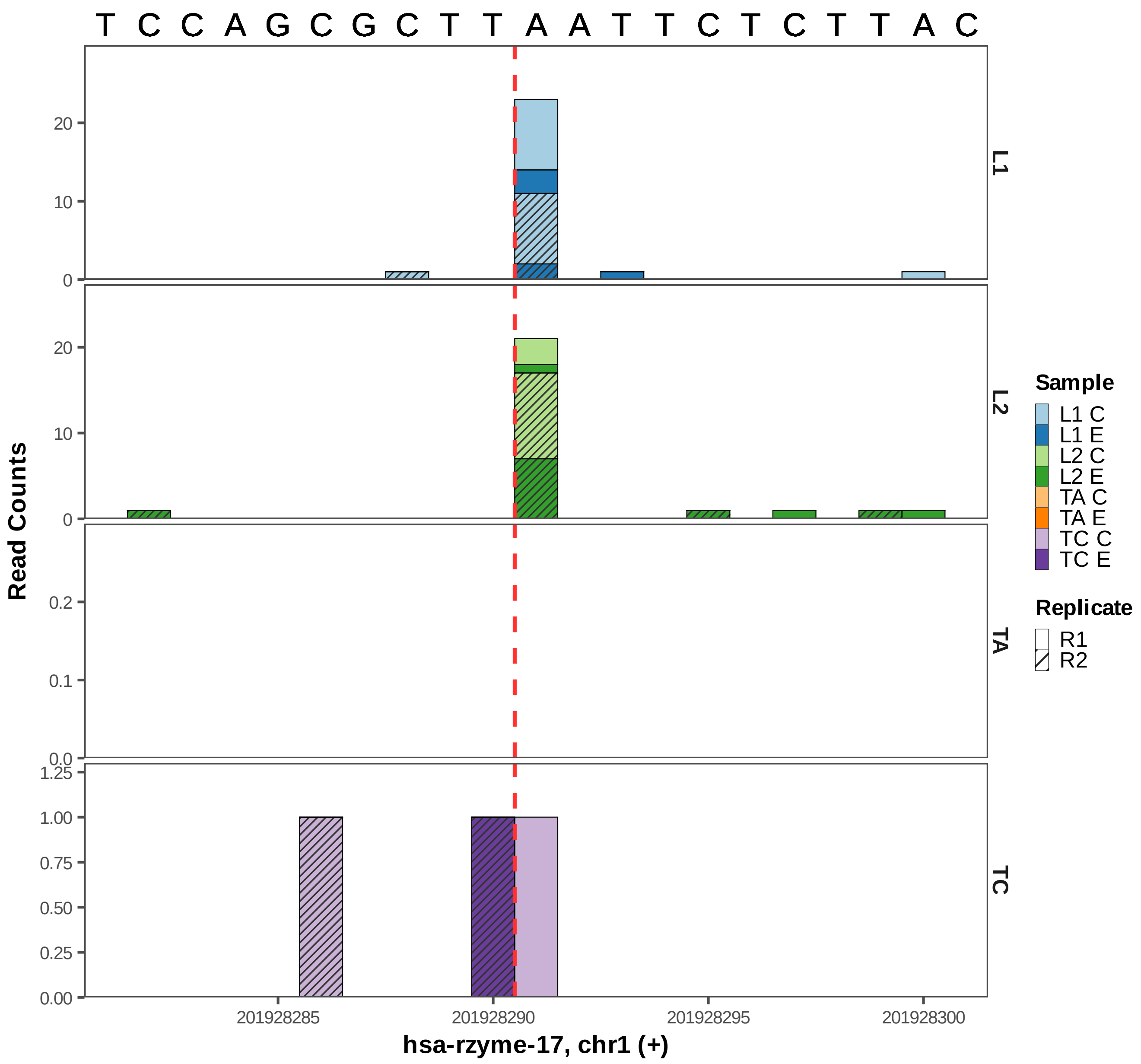

T C C A G C G C T T A A T T C T C T T A C
Read Counts
L1
L2
TA
TC
0
10
20
0.0
0.1
0.2
0.00
0.25
0.50
0.75
1.00
1.25
201928285
201928290
201928295
201928300
hsa-rzyme-17, chr1 (+)
Sample
L1 C
L1 E
L2 C
L2 E
TA C
TA E
TC C
TC E
Replicate
R1
R2

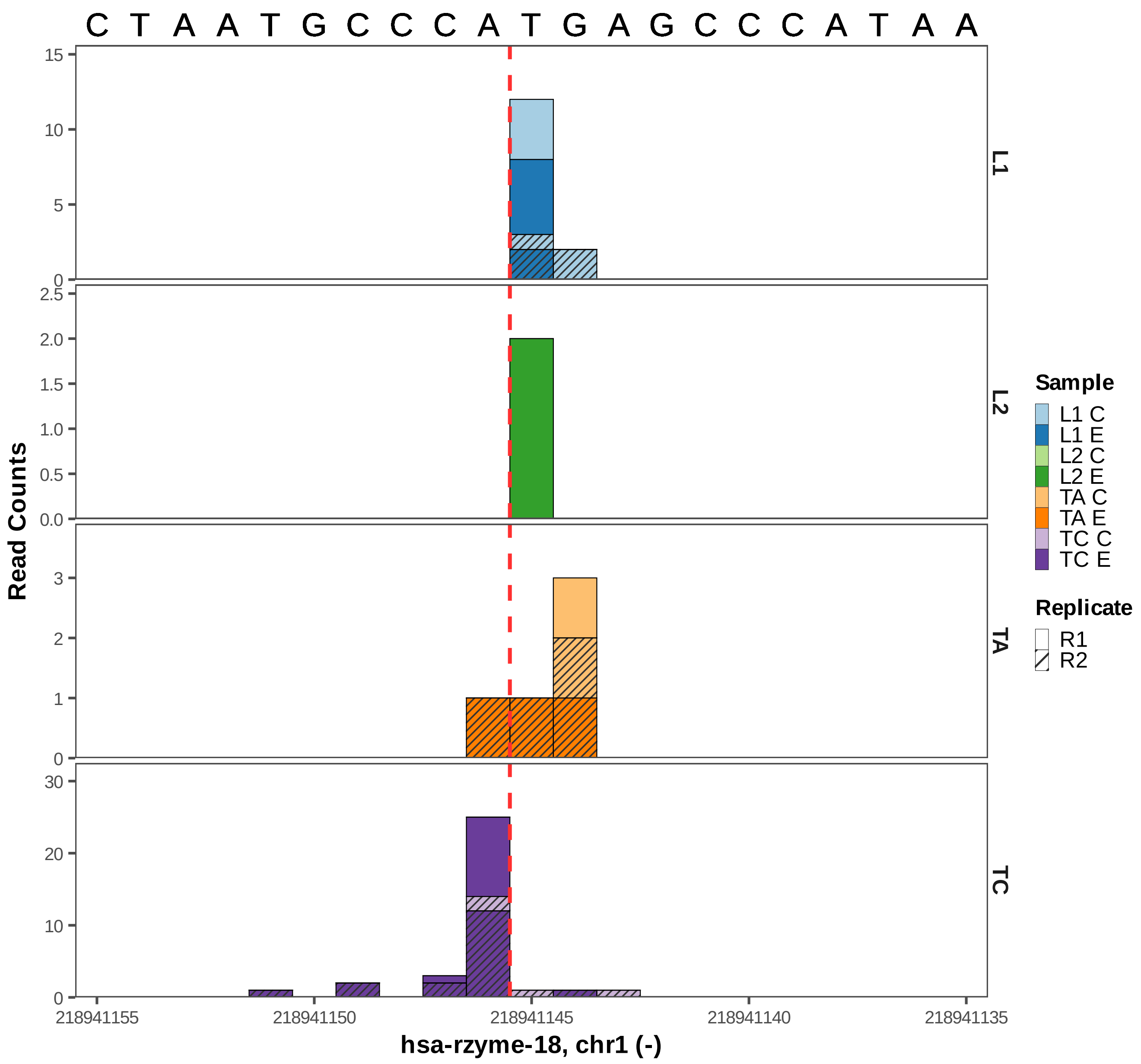

C T A A T G C C C A T G A G C C C A T A A
Read Counts
L1
L2
TA
TC
218941155
218941150
218941145
218941140
218941135
hsa-rzyme-18, chr1 (-)
Sample
L1 C
L1 E
L2 C
L2 E
TA C
TA E
TC C
TC E
Replicate
R1
R2

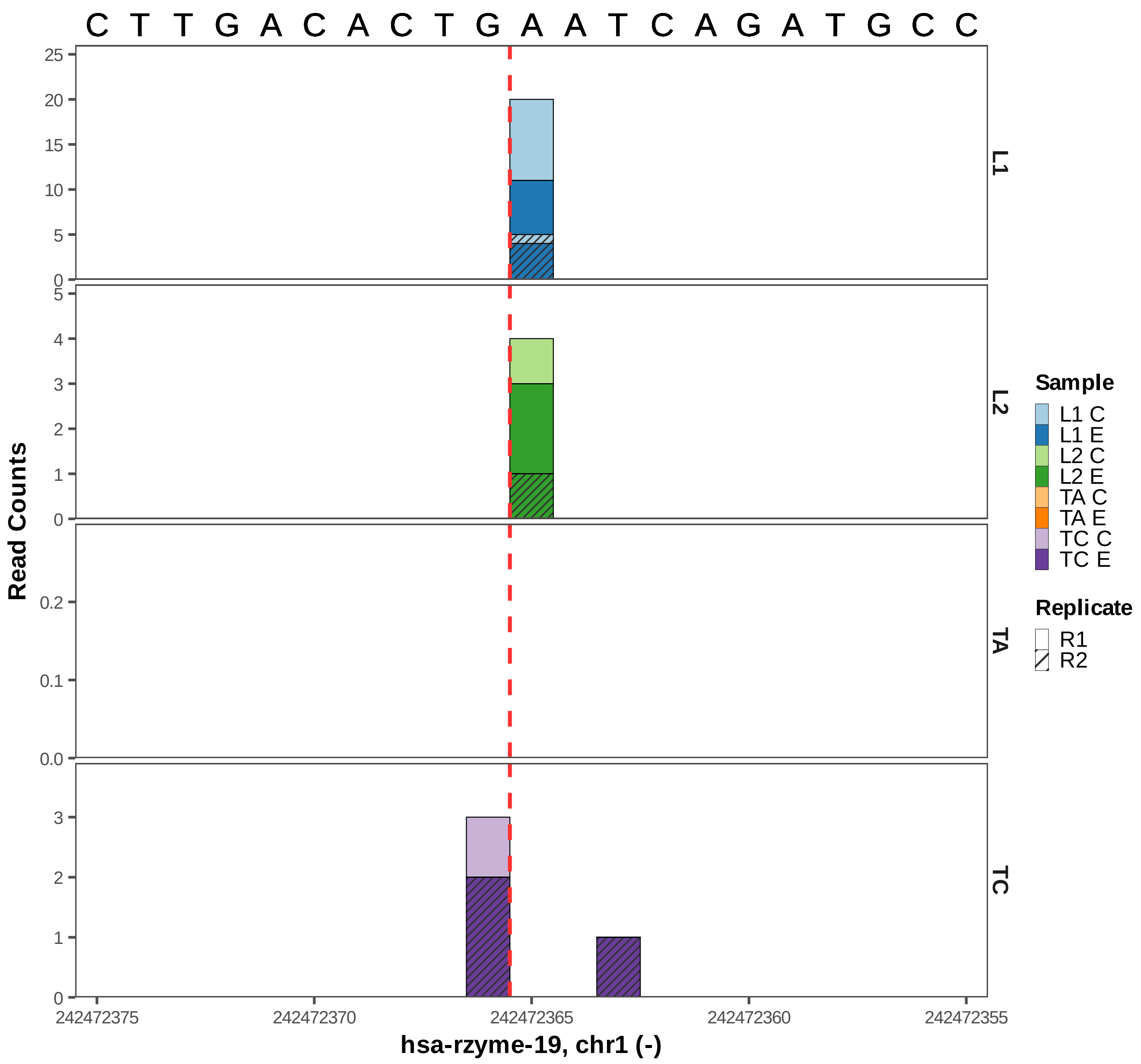

C T T G A C A C T G A A T C A G A T G C C
L1
L2
TA
TC
Read Counts
Sample
L1 C
L1 E
L2 C
L2 E
TA C
TA E
TC C
TC E
Replicate
R1
R2
242472375
242472370
242472365
242472360
242472355
hsa-rzyme-19, chr1 (-)

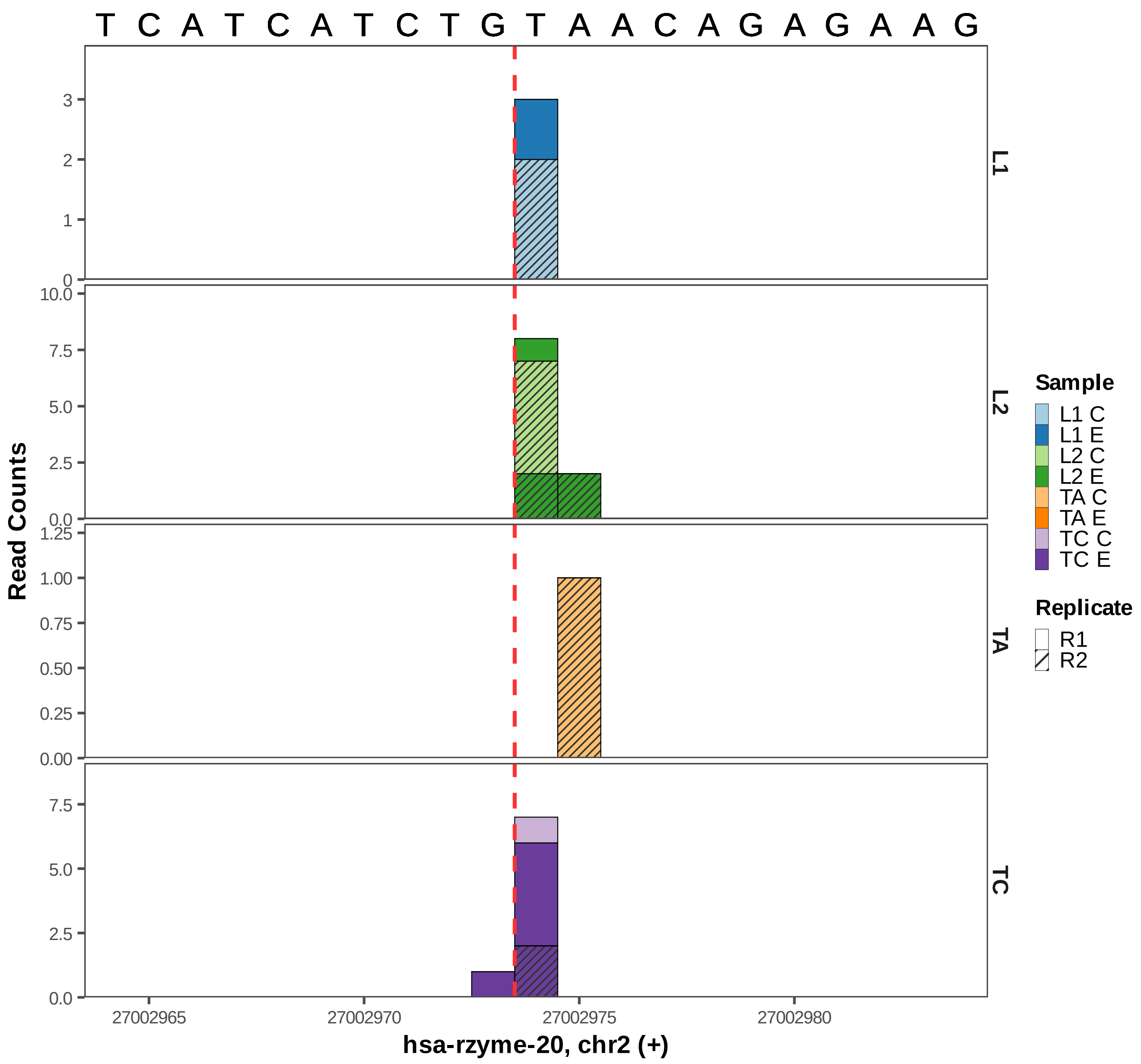
T C A T C A T C T G T A A C A G A G A A G
Read Counts
L1
L2
TA
TC
Sample
L1 C
L1 E
L2 C
L2 E
TA C
TA E
TC C
TC E
Replicate
R1
R2
27002965
27002970
27002975
27002980
hsa-rzyme-20, chr2 (+)

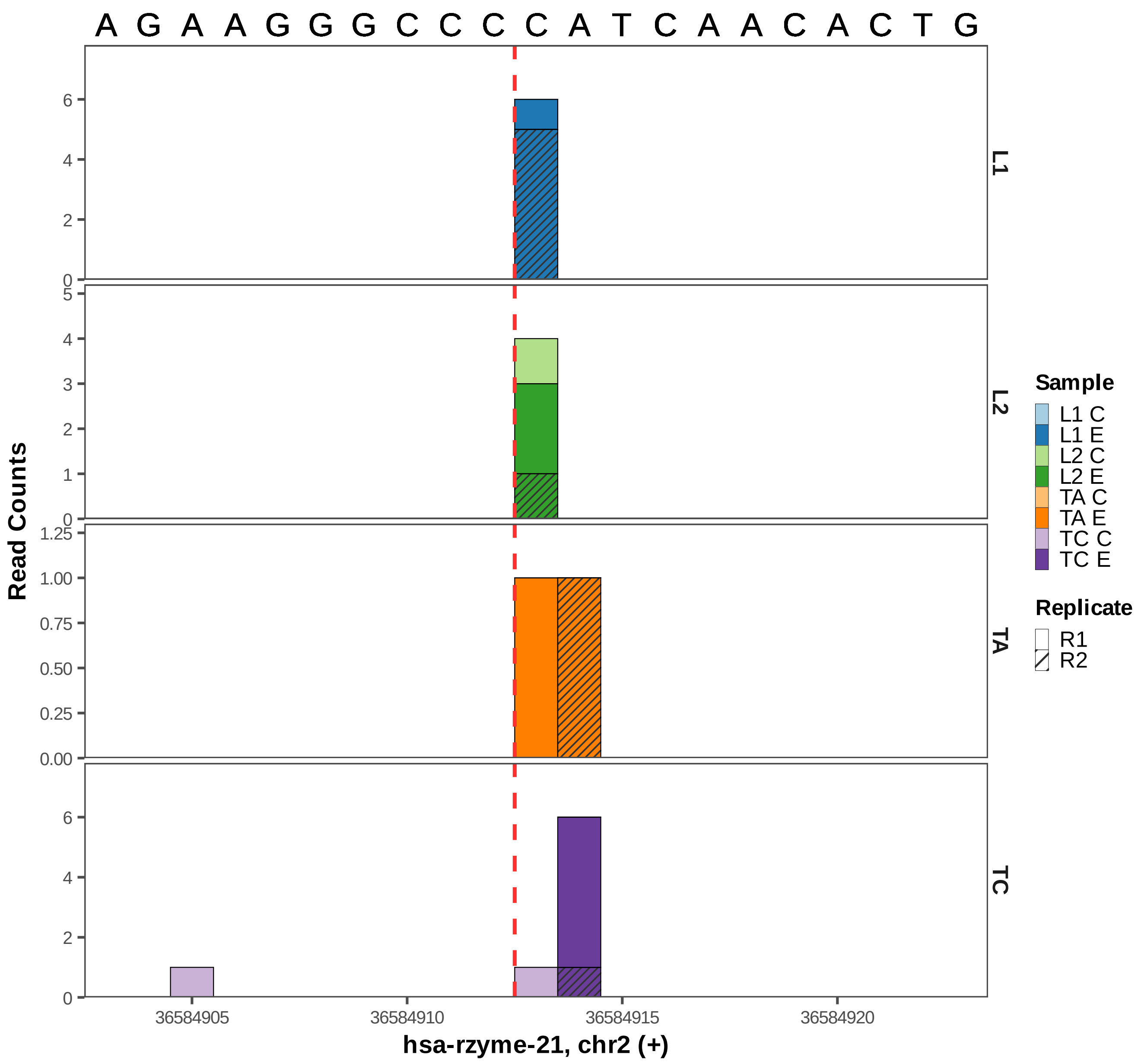

A G A A G G G C C C C A T C A A C A C T G
L1
L2
TA
TC
Read Counts
0
2
4
6
0
1
2
3
4
5
0.00
0.25
0.50
0.75
1.00
1.25
36584905
36584910
36584915
36584920
hsa-rzyme-21, chr2 (+)
Sample
L1 C
L1 E
L2 C
L2 E
TA C
TA E
TC C
TC E
Replicate
R1
R2

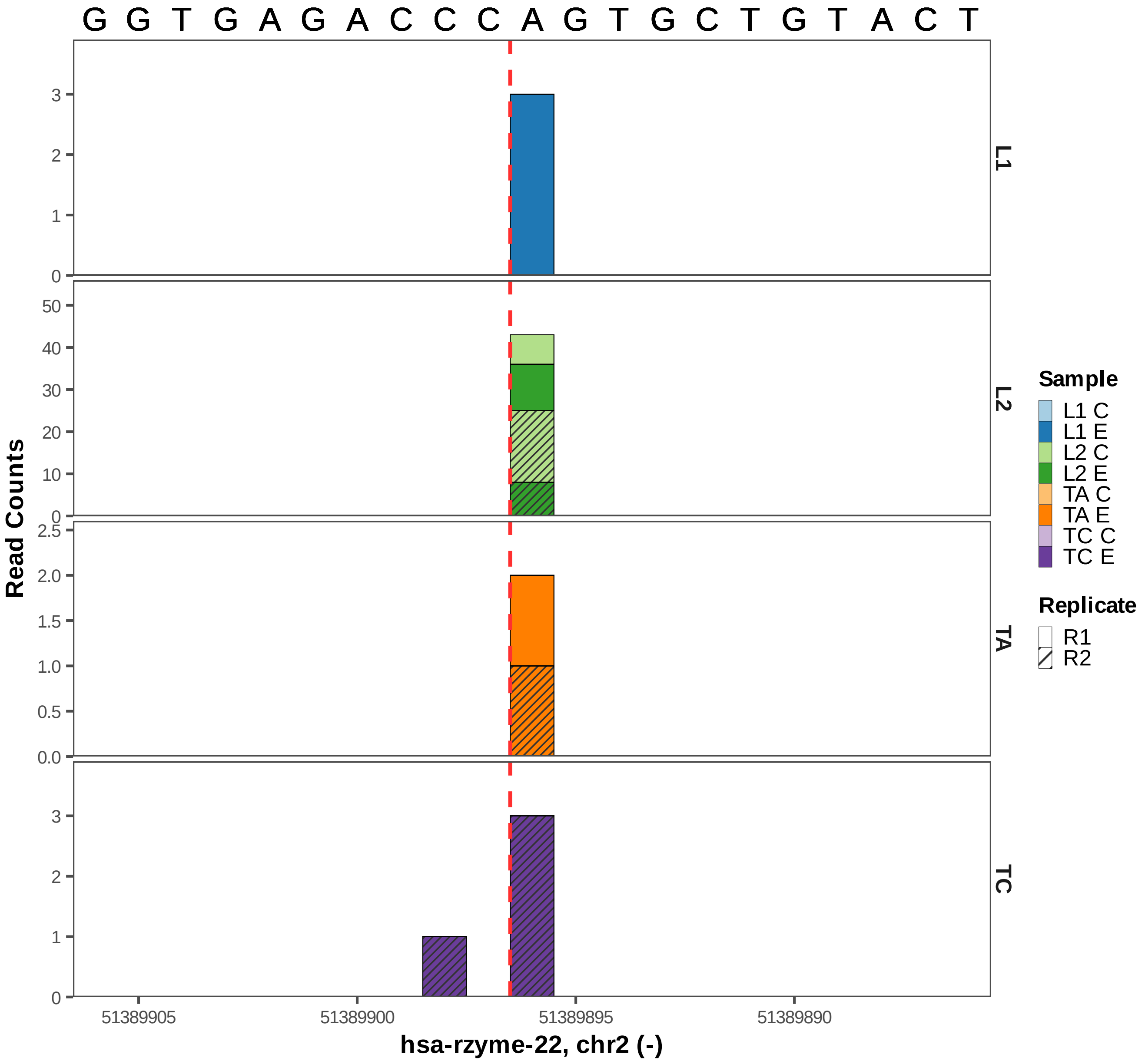
G G T G A G A C C C A G T G C T G T A C T
Read Counts
L1
L2
TA
TC
51389905
51389900
51389895
51389890
hsa-rzyme-22, chr2 (-)
Sample
L1 C
L1 E
L2 C
L2 E
TA C
TA E
TC C
TC E
Replicate
R1
R2

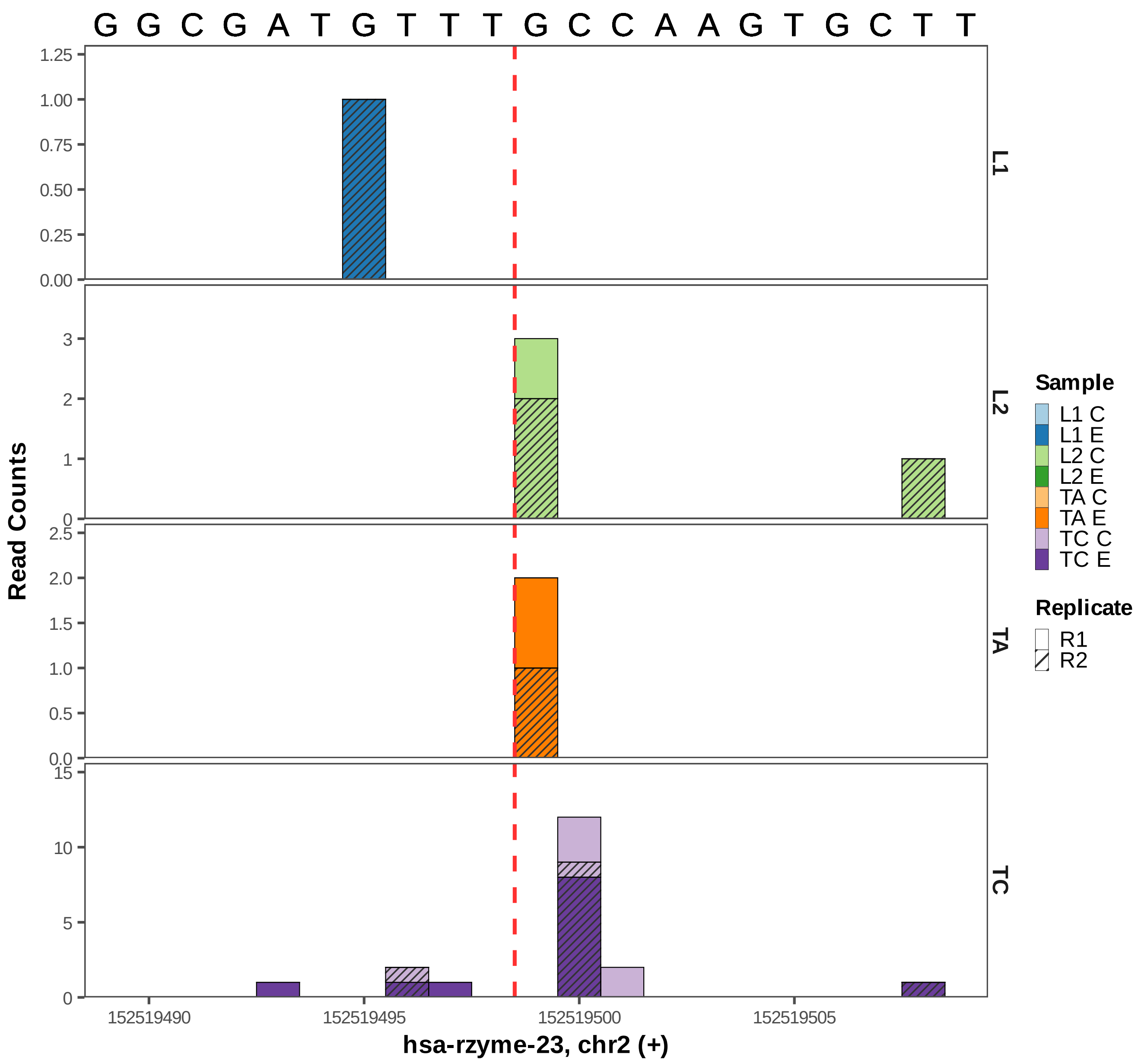

G G C G A T G T T T G C C A A G T G C T T
Read Counts
L1
L2
TA
TC
152519490
152519495
152519500
152519505
hsa-rzyme-23, chr2 (+)
Sample
L1 C
L1 E
L2 C
L2 E
TA C
TA E
TC C
TC E
Replicate
R1
R2

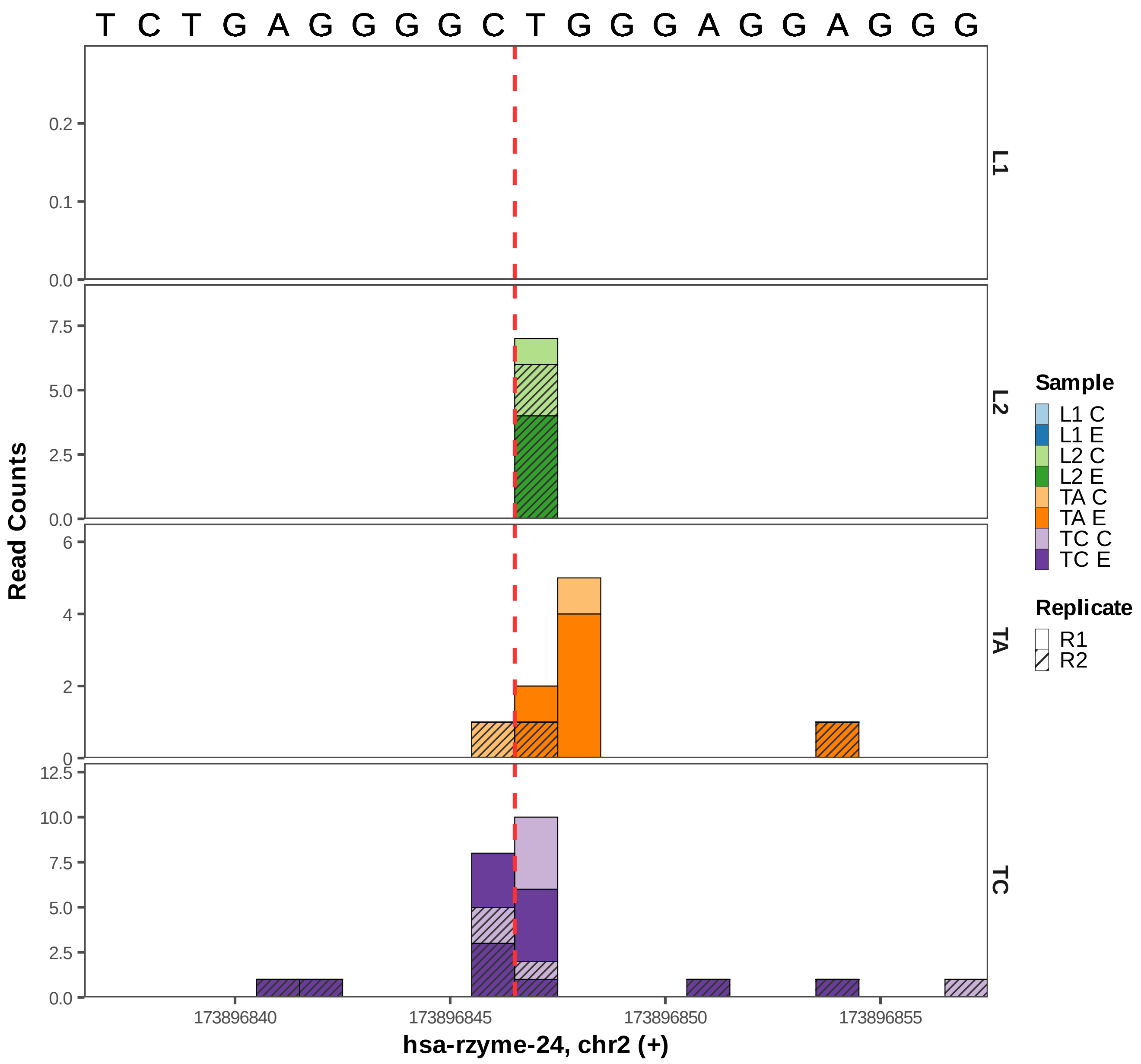
T C T G A G G G G C T G G G A G G A G G G
L1
L2
TA
TC
Read Counts
0.0
0.1
0.2
2.5
5.0
7.5
0
2
4
6
10.0
12.5
173896840
173896845
173896850
173896855
hsa-rzyme-24, chr2 (+)
Sample
L1 C
L1 E
L2 C
L2 E
TA C
TA E
TC C
TC E
Replicate
R1
R2

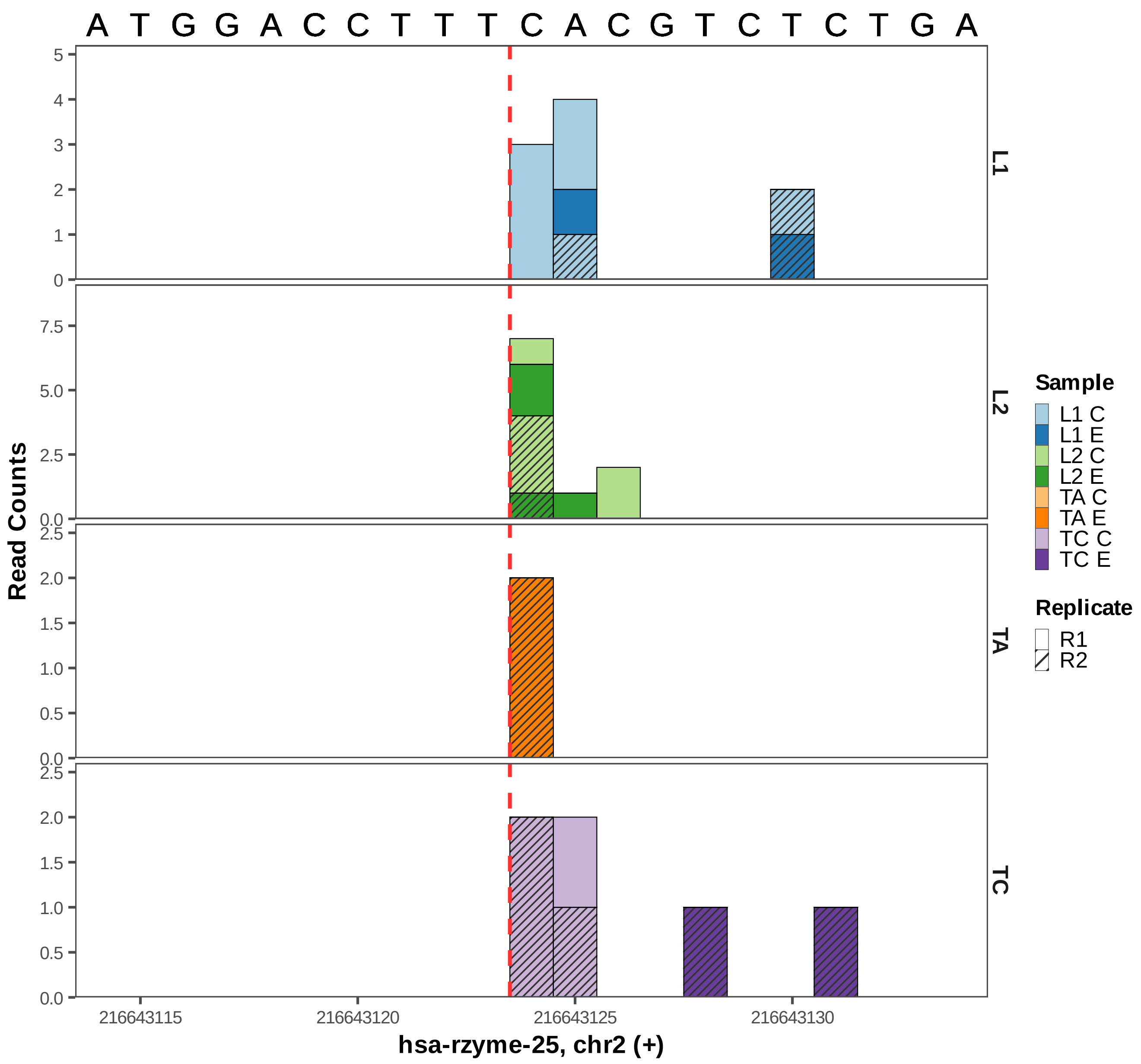
A T G G A C C T T T C A C G T C T C T G A
L1
L2
TA
TC
Read Counts
Sample
L1 C
L1 E
L2 C
L2 E
TA C
TA E
TC C
TC E
Replicate
R1
R2
216643115
216643120
216643125
216643130
hsa-rzyme-25, chr2 (+)

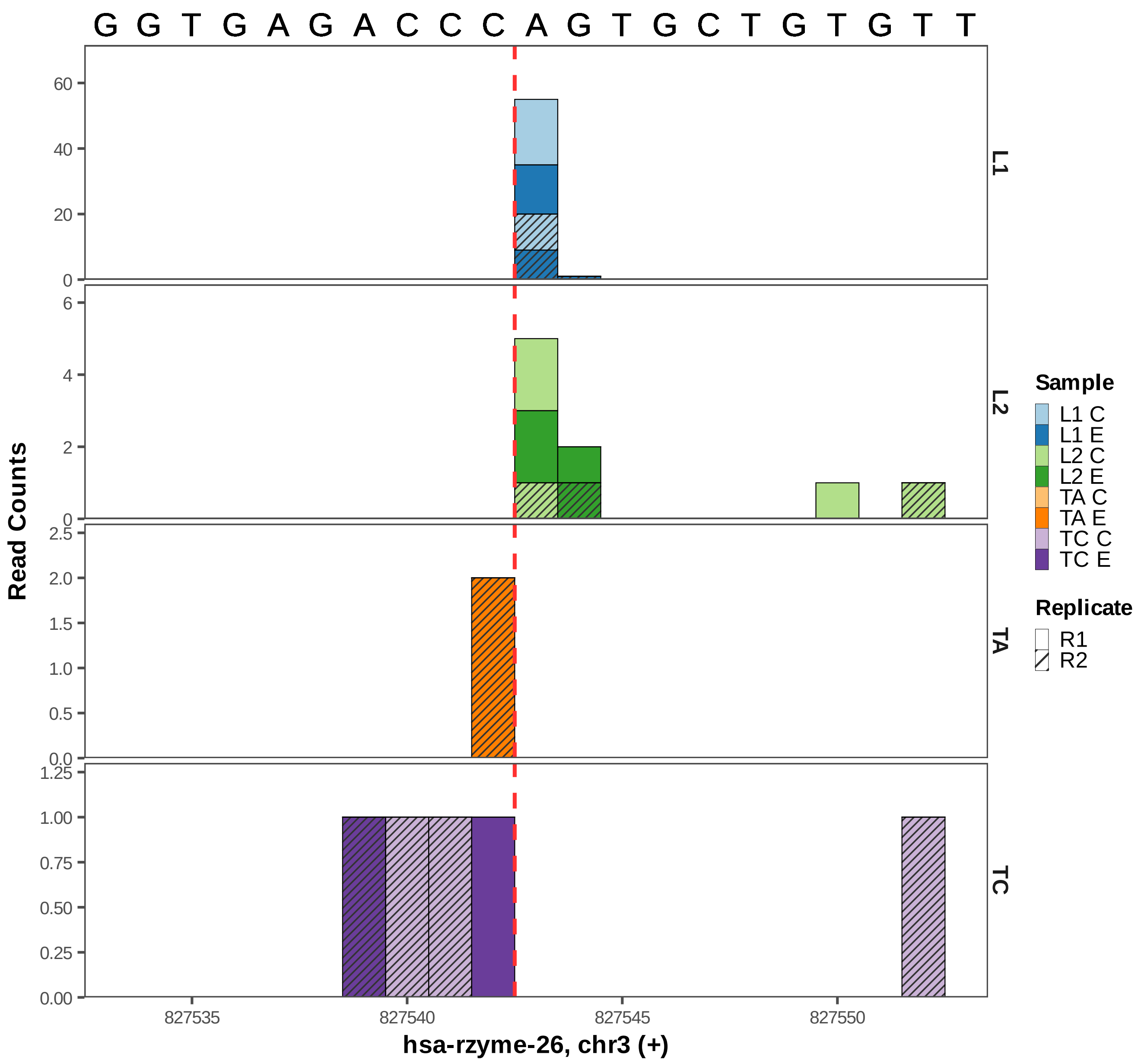

G G T G A G A C C C A G T G C T G T G T T
Read Counts
L1
L2
TA
TC
Sample
L1 C
L1 E
L2 C
L2 E
TA C
TA E
TC C
TC E
Replicate
R1
R2
827535
827540
827545
827550
hsa-rzyme-26, chr3 (+)

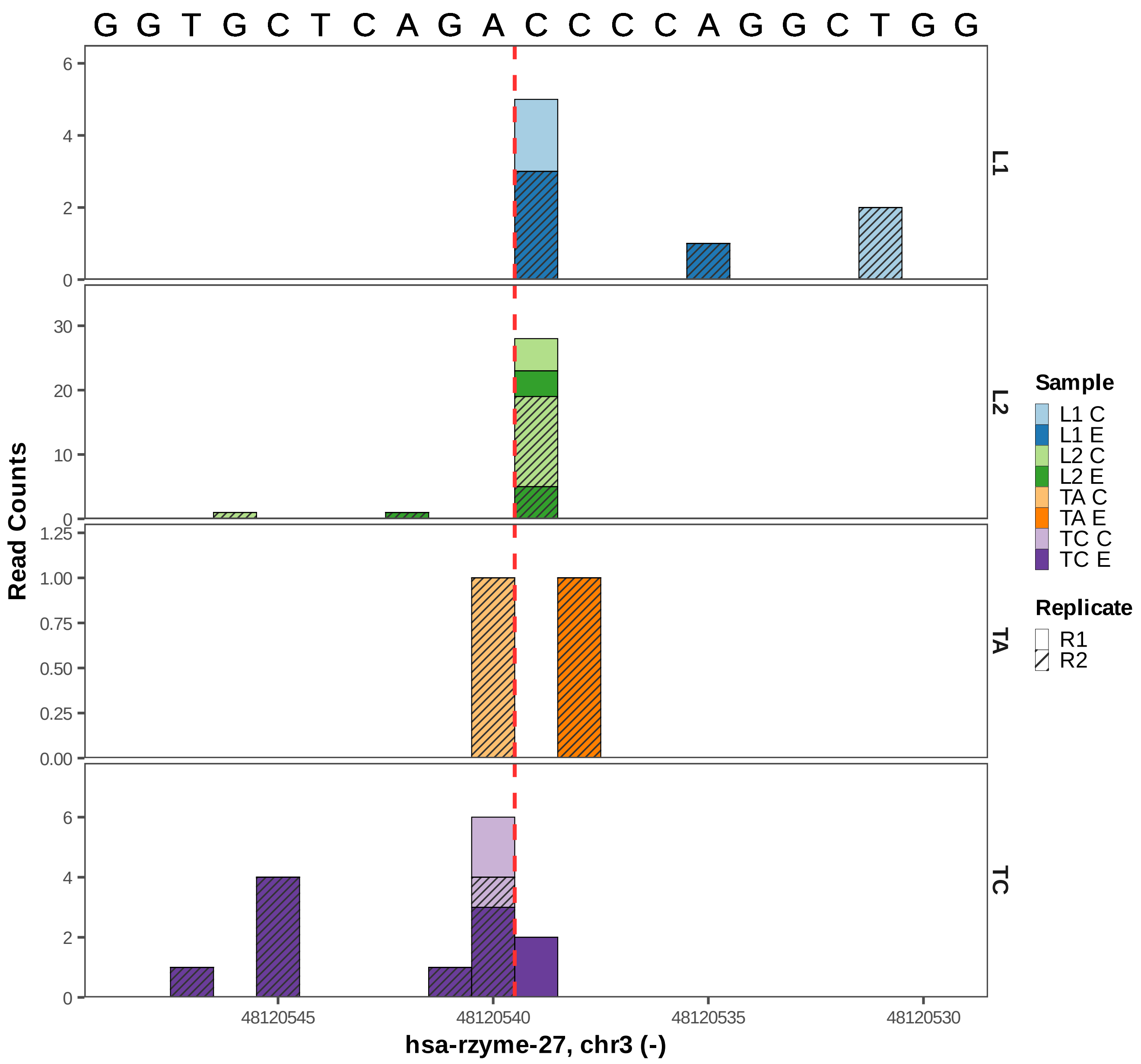
G G T G C T C A G A C C C C A G G C T G G
Read Counts
L1
L2
TA
TC
48120545
48120540
48120535
48120530
hsa-rzyme-27, chr3 (-)
Sample
L1 C
L1 E
L2 C
L2 E
TA C
TA E
TC C
TC E
Replicate
R1
R2

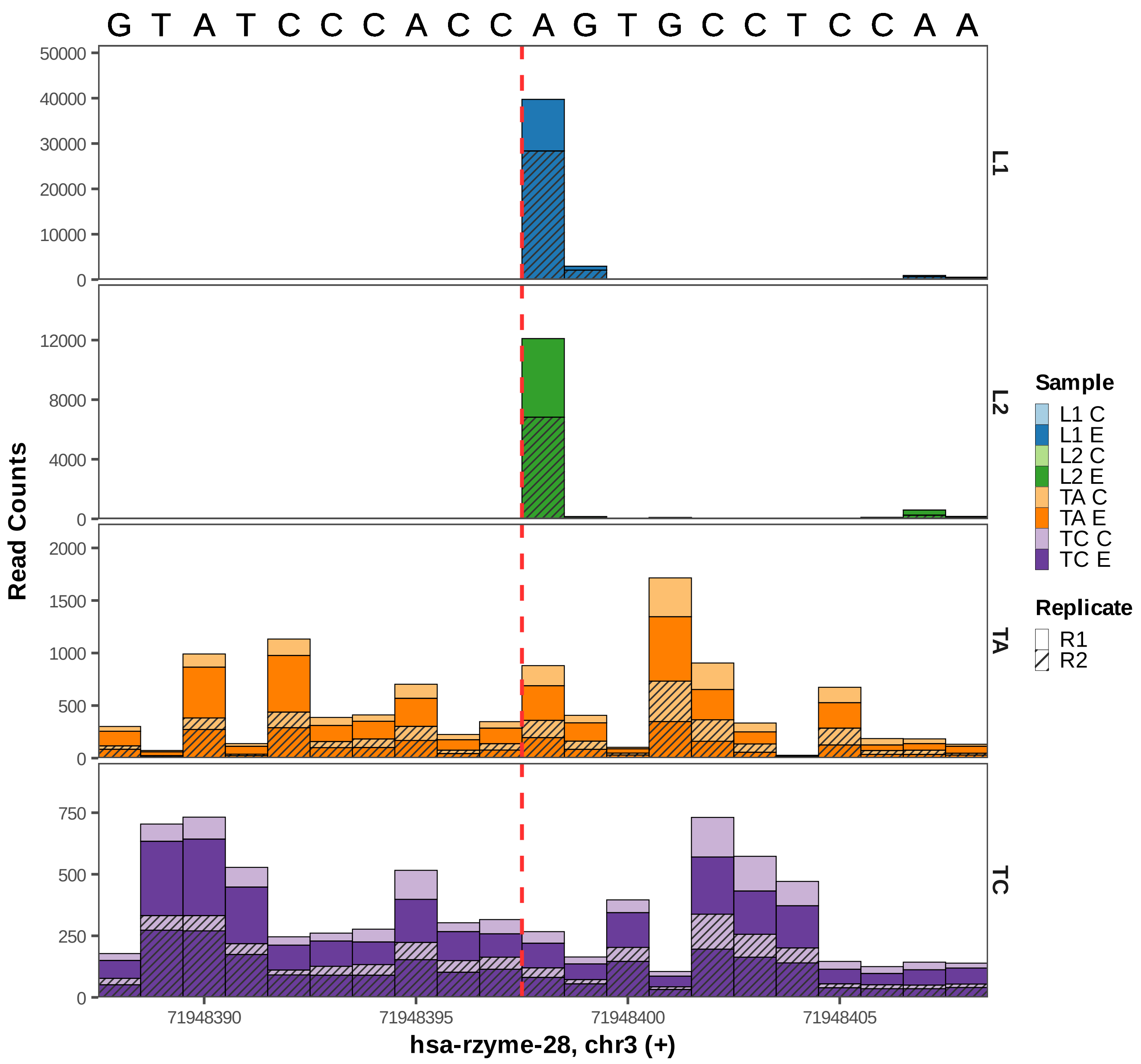
G T A T C C C A C C A G T G C C T C C A A
Read Counts
L1
L2
TA
TC
Sample
L1 C
L1 E
L2 C
L2 E
TA C
TA E
TC C
TC E
Replicate
R1
R2
71948390
71948395
71948400
71948405
hsa-rzyme-28, chr3 (+)

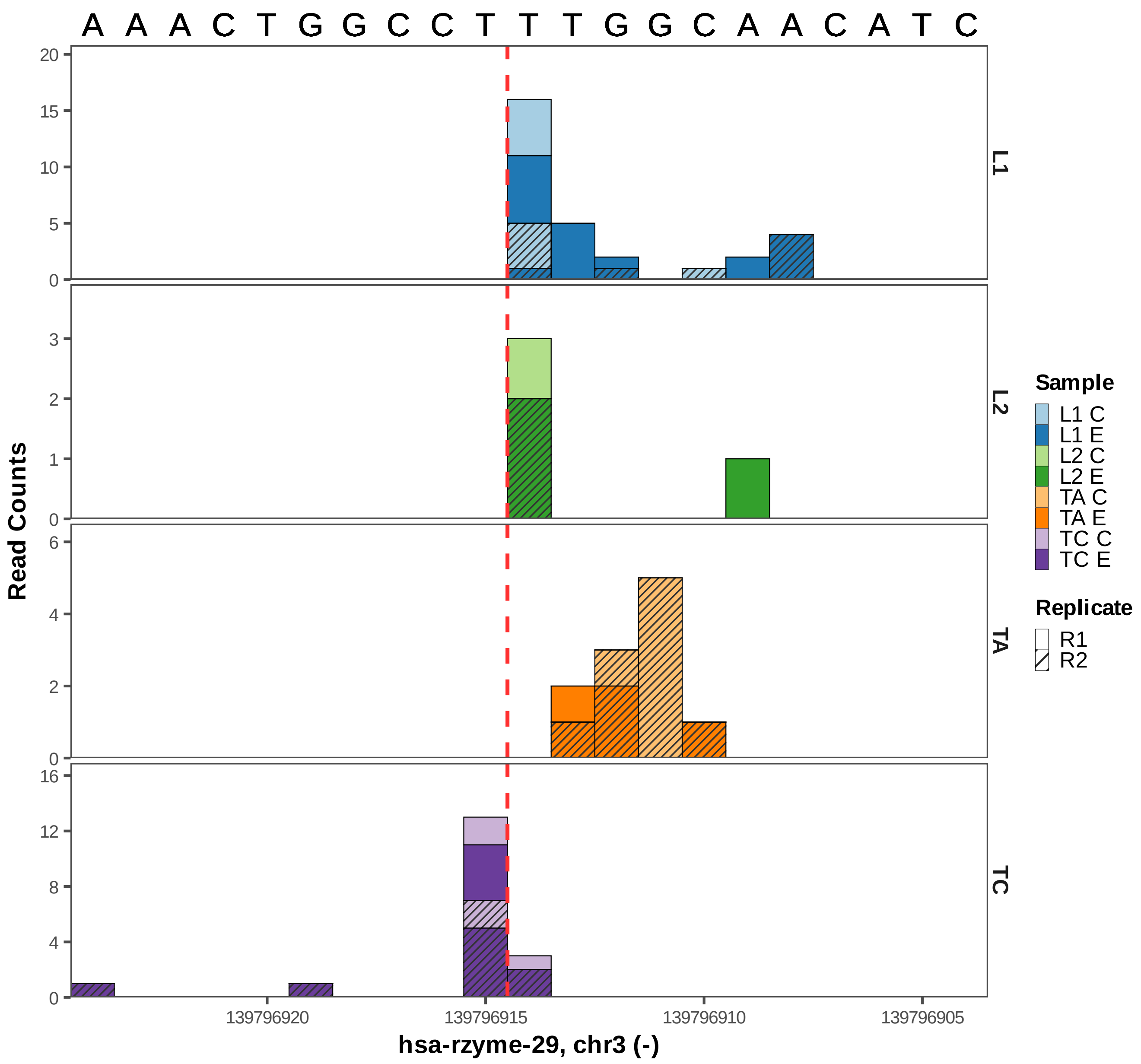

A A A C T G G C C T T T G G C A A C A T C
Read Counts
20
15
10
5
0
3
2
1
0
6
4
2
0
16
12
8
4
0
L1
L2
TA
TC
139796920
139796915
139796910
139796905
hsa-rzyme-29, chr3 (-)
Sample
L1 C
L1 E
L2 C
L2 E
TA C
TA E
TC C
TC E
Replicate
R1
R2

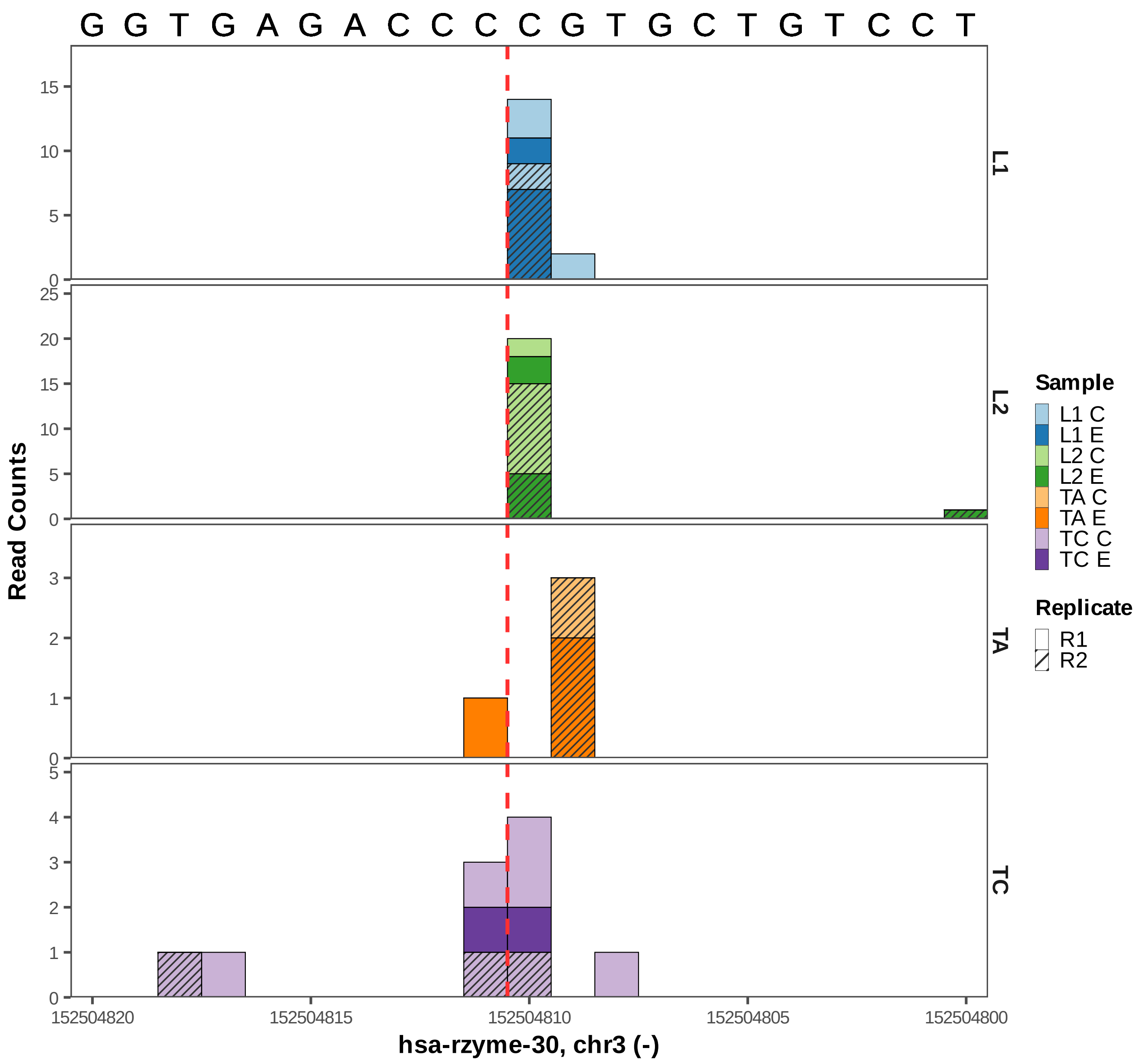

G G T G A G A C C C C G T G C T G T C C T
Read Counts
L1
L2
TA
TC
hsa-rzyme-30, chr3 (-)
152504820
152504815
152504810
152504805
152504800
Sample
L1 C
L1 E
L2 C
L2 E
TA C
TA E
TC C
TC E
Replicate
R1
R2

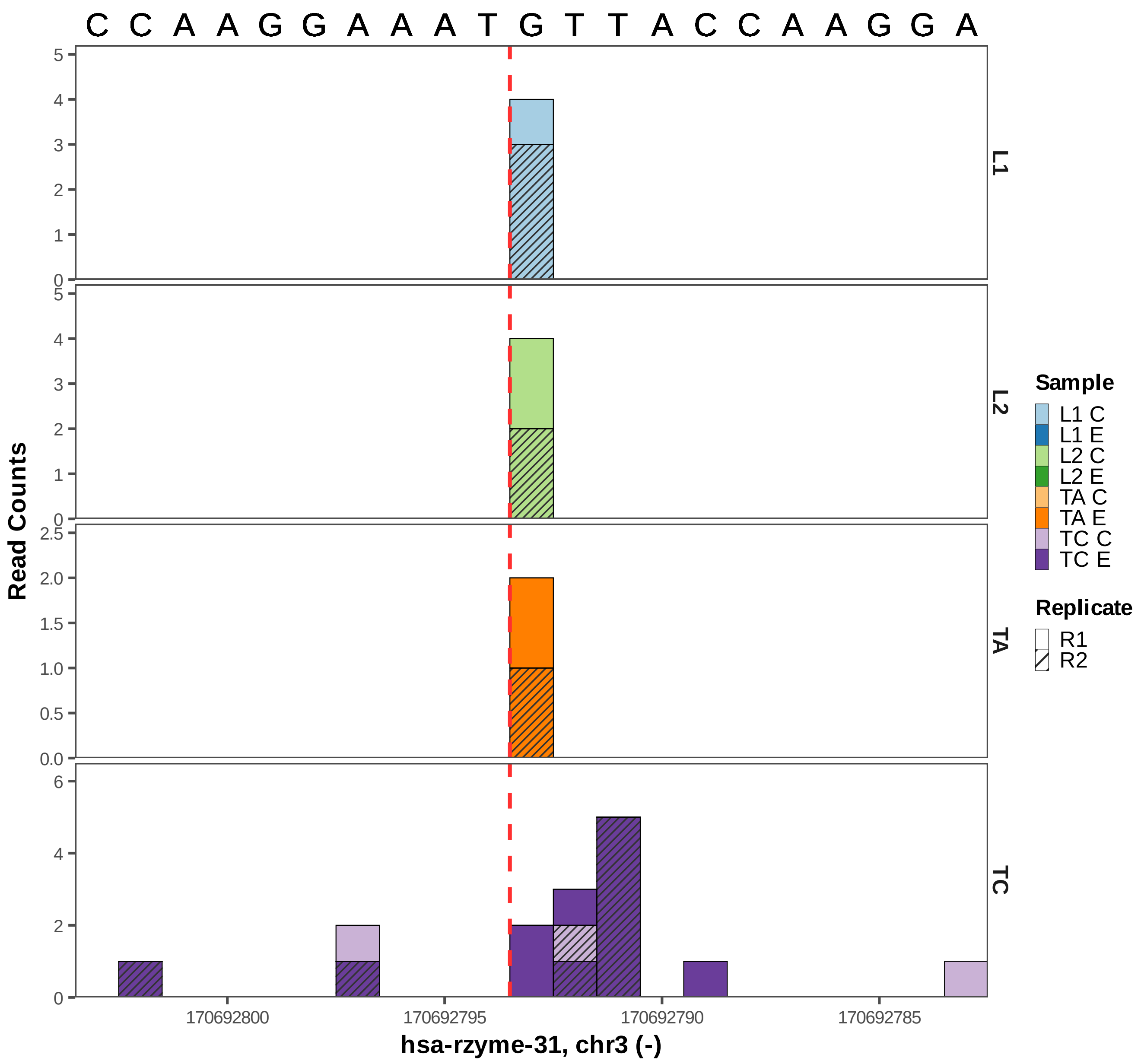

C C A A G G A A A T G T T A C C A A G G A
Read Counts
L1
L2
TA
TC
170692800
170692795
170692790
170692785
hsa-rzyme-31, chr3 (-)
Sample
L1 C
L1 E
L2 C
L2 E
TA C
TA E
TC C
TC E
Replicate
R1
R2

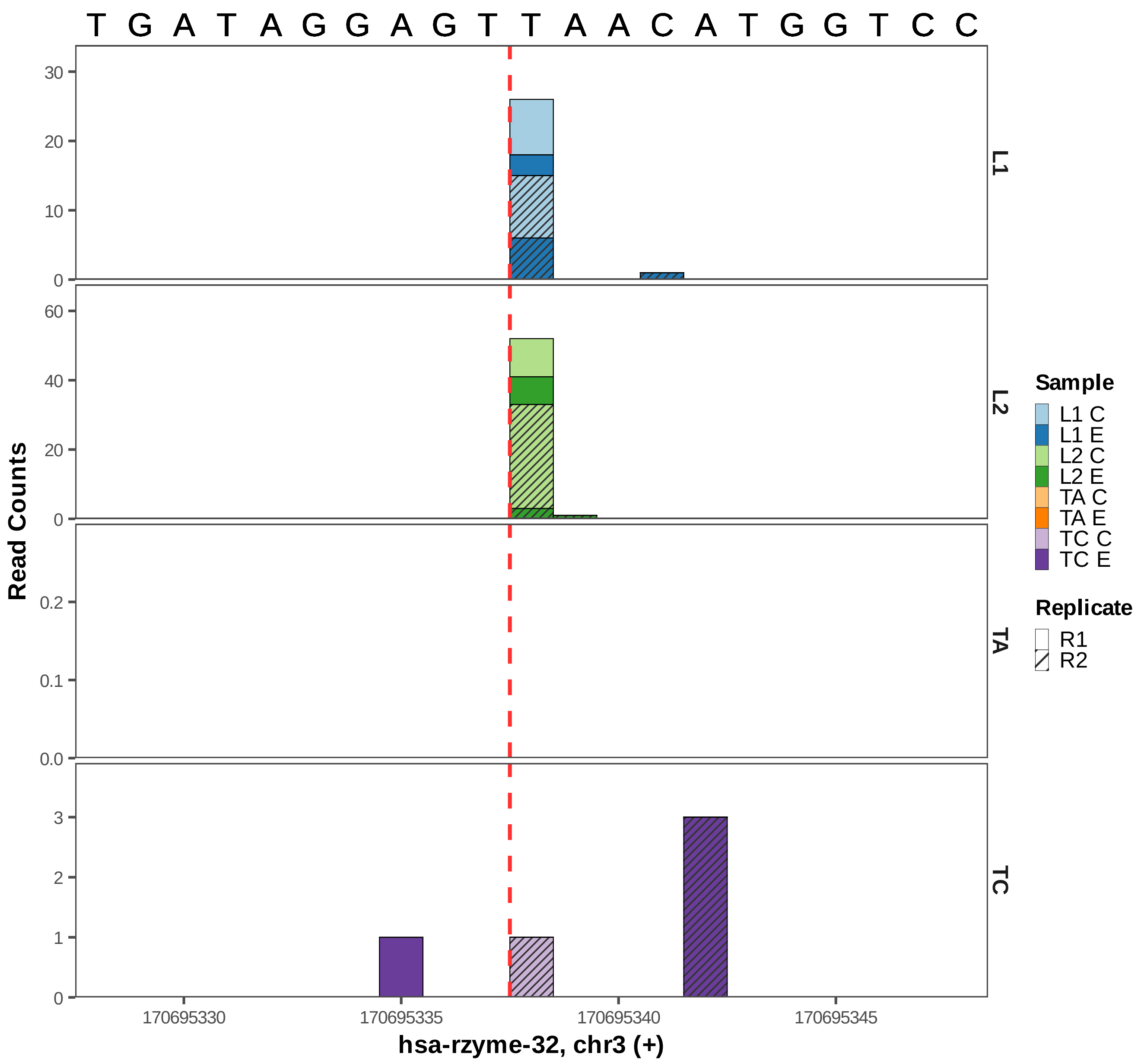

T G A T A G G A G T T A A C A T G G T C C
Read Counts
L1
L2
TA
TC
30
20
10
0
60
40
20
0
0.2
0.1
0.0
3
2
1
0
170695330
170695335
170695340
170695345
hsa-rzyme-32, chr3 (+)
Sample
L1 C
L1 E
L2 C
L2 E
TA C
TA E
TC C
TC E
Replicate
R1
R2

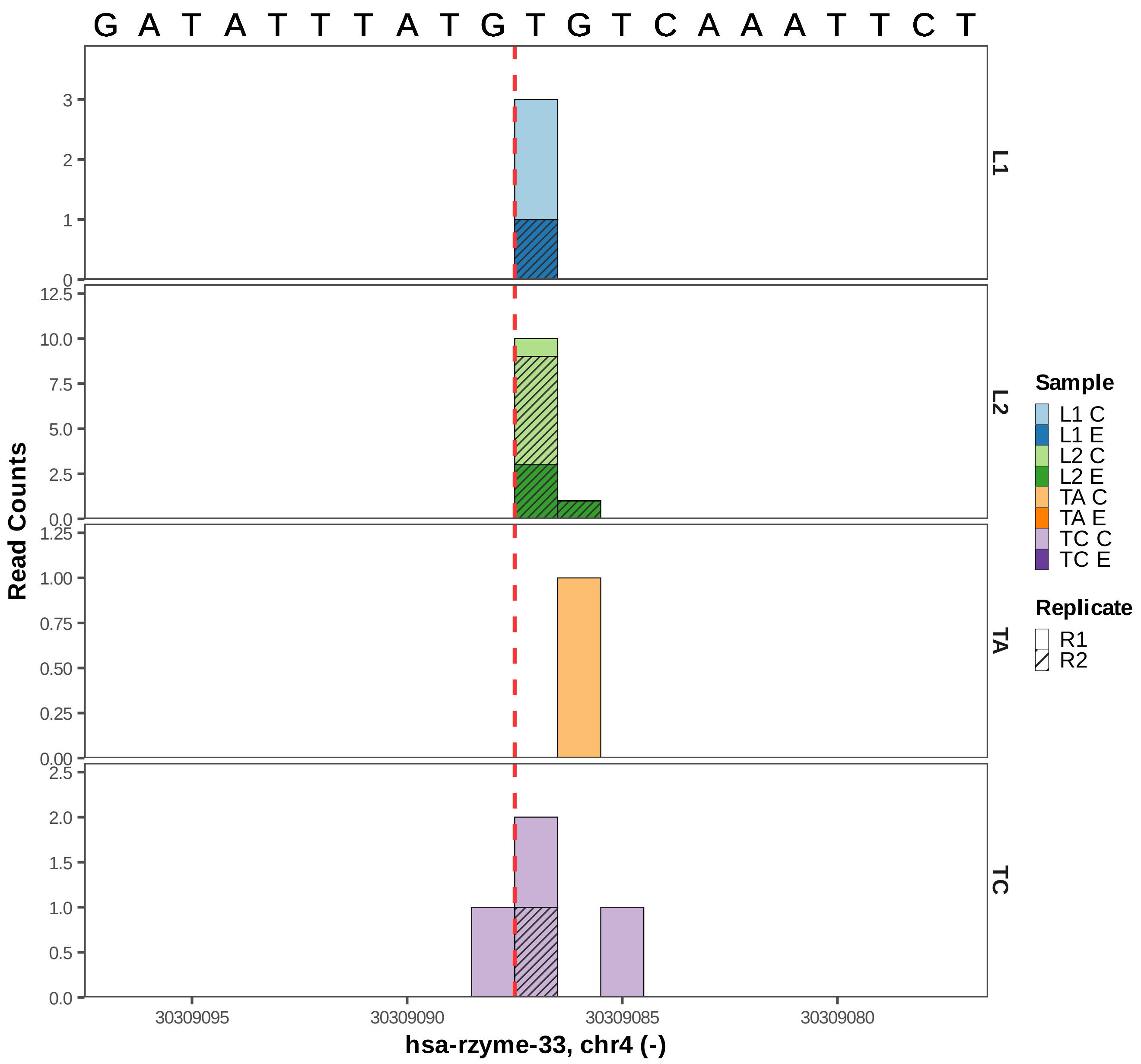
G A T A T T T A T G T G T C A A A T T C T
Read Counts
L1
L2
TA
TC
Sample
L1 C
L1 E
L2 C
L2 E
TA C
TA E
TC C
TC E
Replicate
R1
R2
30309095
30309090
30309085
30309080
hsa-rzyme-33, chr4 (-)

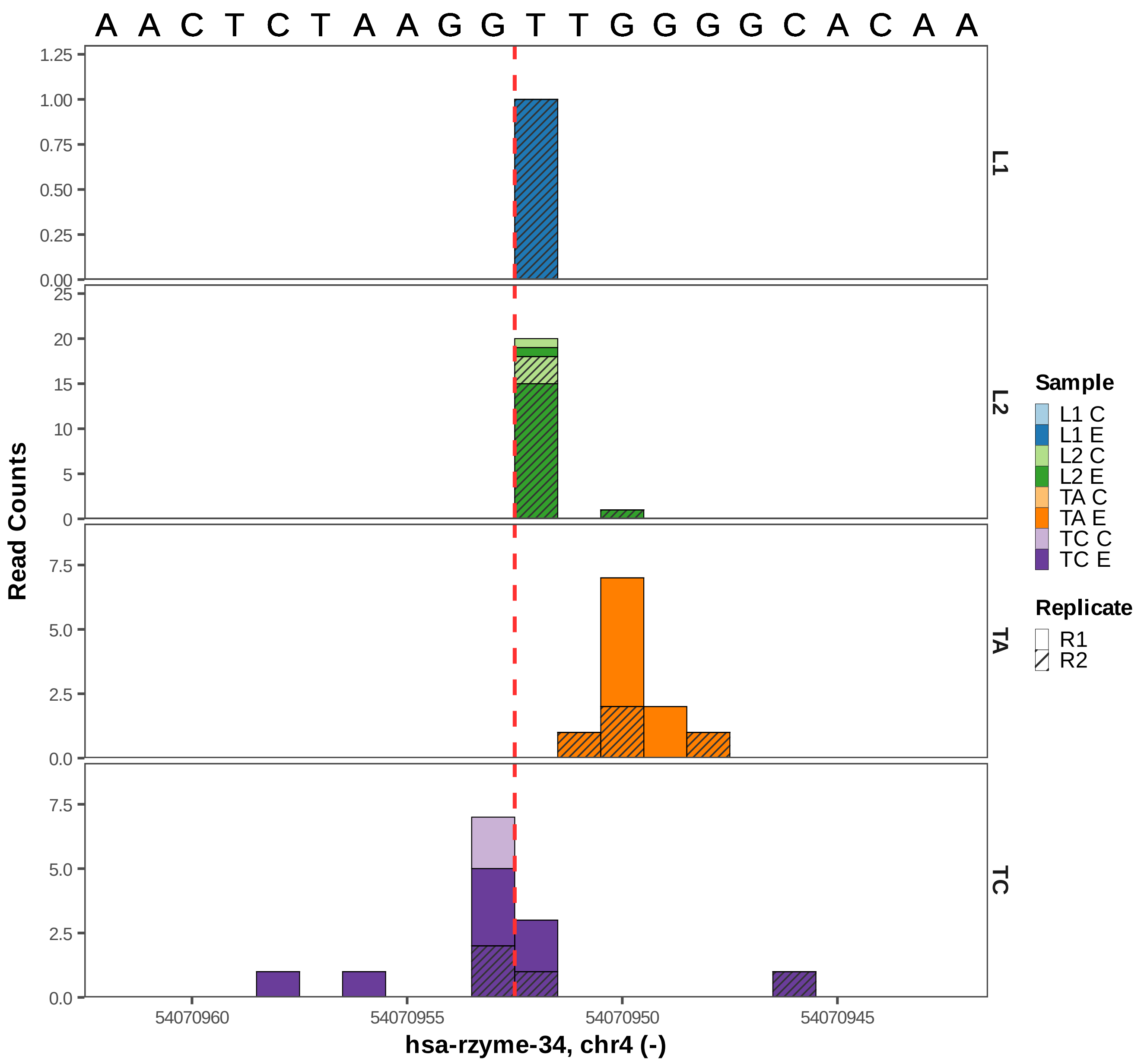

A A C T C T A A G G T T G G G G C A C A A
L1
L2
TA
TC
Read Counts
hsa-rzyme-34, chr4 (-)
54070960
54070955
54070950
54070945
Sample
L1 C
L1 E
L2 C
L2 E
TA C
TA E
TC C
TC E
Replicate
R1
R2

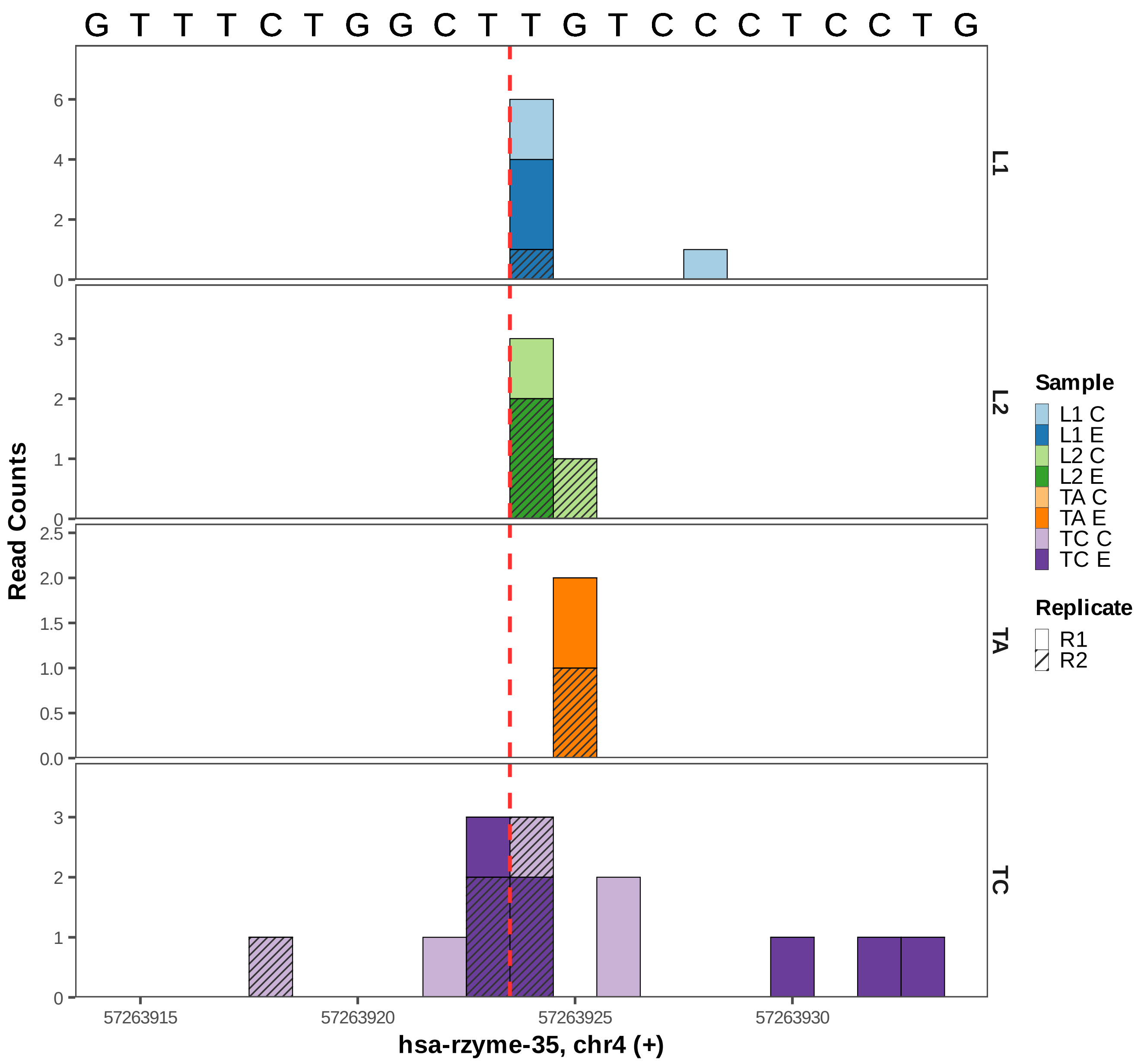

G T T T C T G G C T T G T C C C T C C T G
L1
L2
TA
TC
Read Counts
hsa-rzyme-35, chr4 (+)
57263915
57263920
57263925
57263930
Sample
L1 C
L1 E
L2 C
L2 E
TA C
TA E
TC C
TC E
Replicate
R1
R2

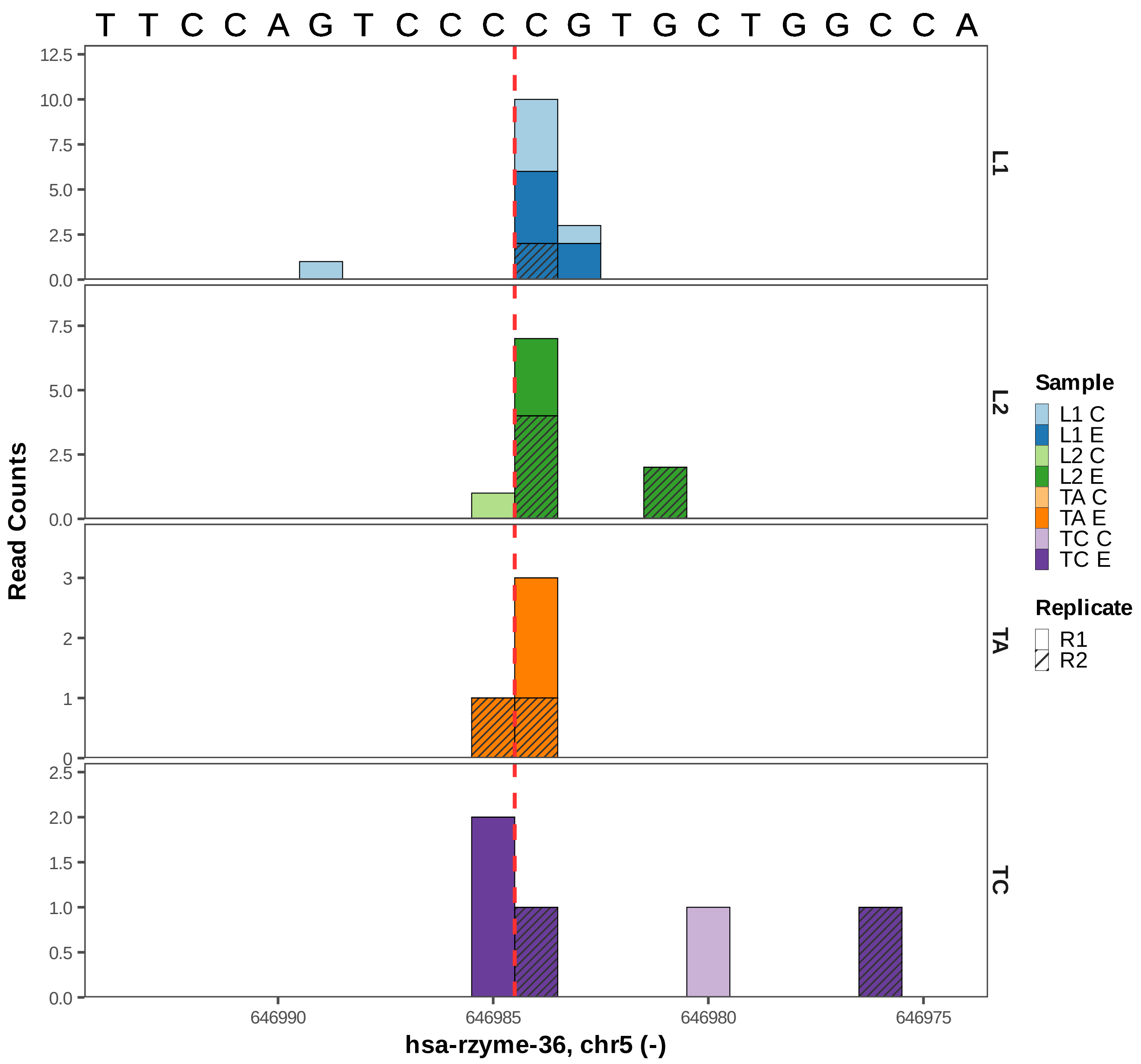

T T C C A G T C C C C G T G C T G G C C A
Read Counts
L1
L2
TA
TC
0.0
2.5
5.0
7.5
10.0
12.5
0
1
2
3
0.5
1.0
1.5
2.0
646990
646985
646980
646975
hsa-rzyme-36, chr5 (-)
Sample
L1 C
L1 E
L2 C
L2 E
TA C
TA E
TC C
TC E
Replicate
R1
R2

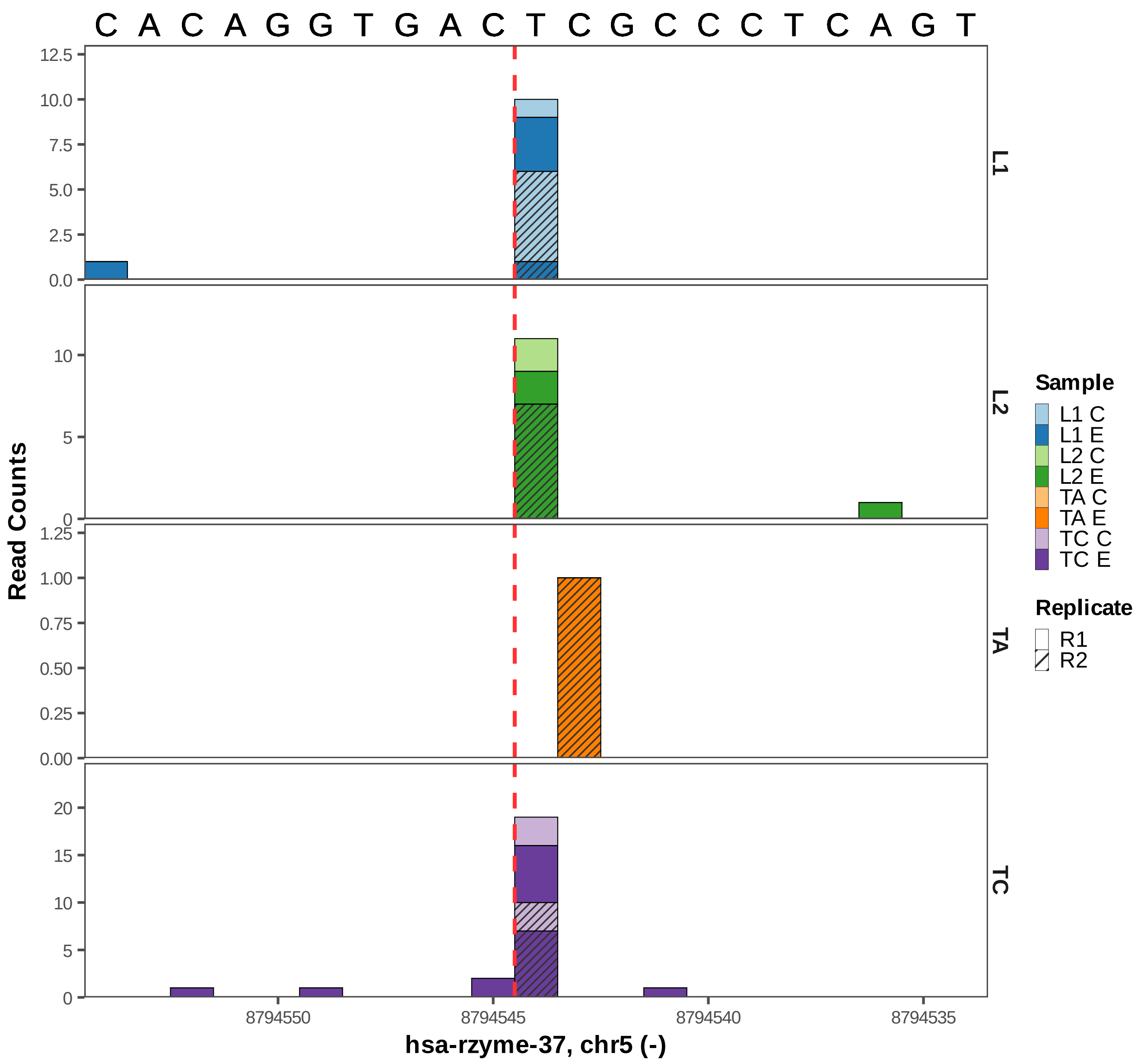

C A C A G G T G A C T C G C C C T C A G T
Read Counts
12.5
10.0
7.5
5.0
2.5
0.0
L1
10
5
0
L2
1.25
1.00
0.75
0.50
0.25
0.00
TA
20
15
10
5
0
TC
8794550
8794545
8794540
8794535
hsa-rzyme-37, chr5 (-)
Sample
L1 C
L1 E
L2 C
L2 E
TA C
TA E
TC C
TC E
Replicate
R1
R2

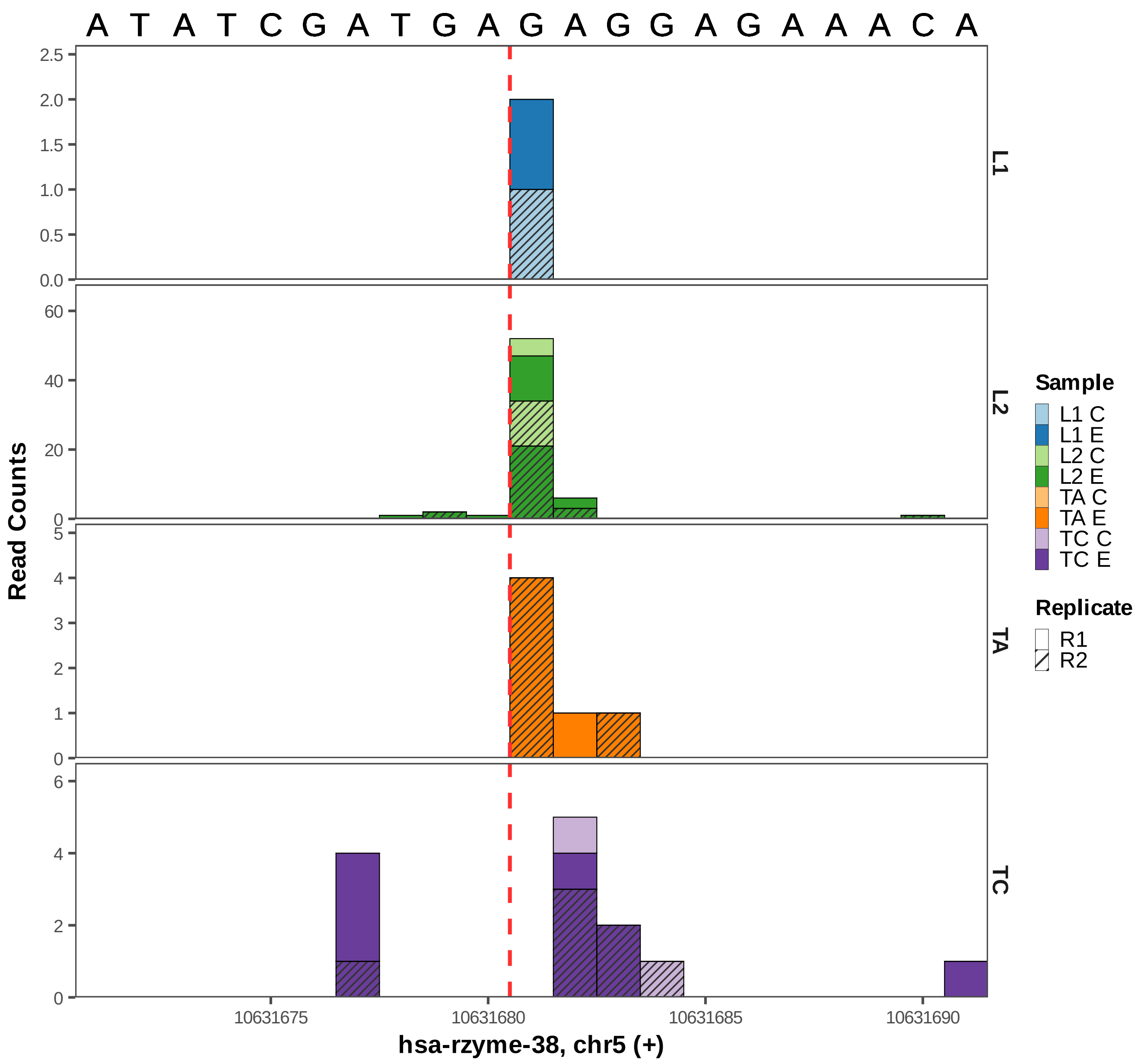
A T A T C G A T G A G A G G A G A A A C A
Read Counts
L1
L2
TA
TC
10631675
10631680
10631685
10631690
hsa-rzyme-38, chr5 (+)
Sample
L1 C
L1 E
L2 C
L2 E
TA C
TA E
TC C
TC E
Replicate
R1
R2

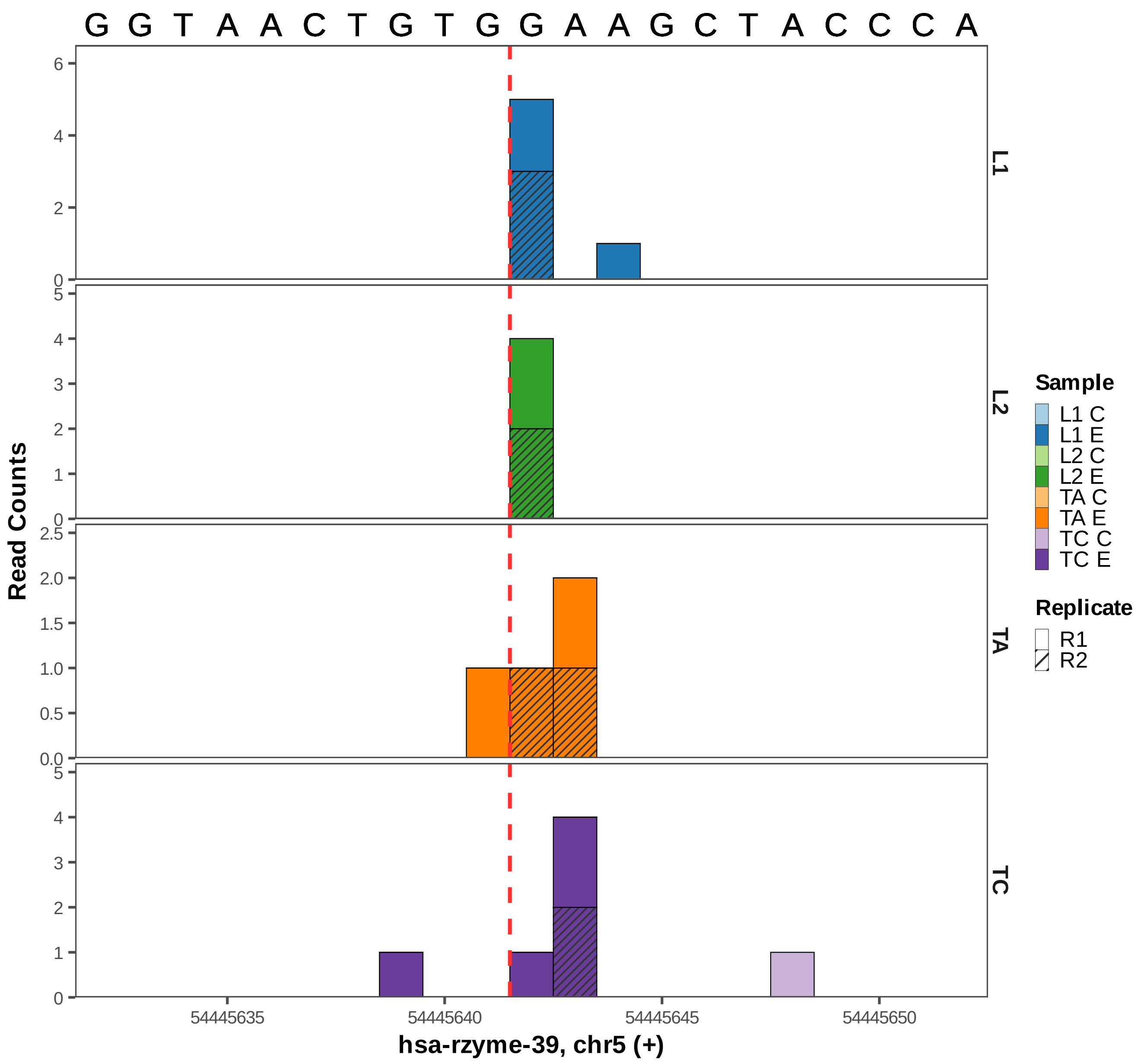

G G T A A C T G T G G A A G C T A C C C A
L1
L2
TA
TC
Read Counts
hsa-rzyme-39, chr5 (+)
54445635
54445640
54445645
54445650
Sample
L1 C
L1 E
L2 C
L2 E
TA C
TA E
TC C
TC E
Replicate
R1
R2

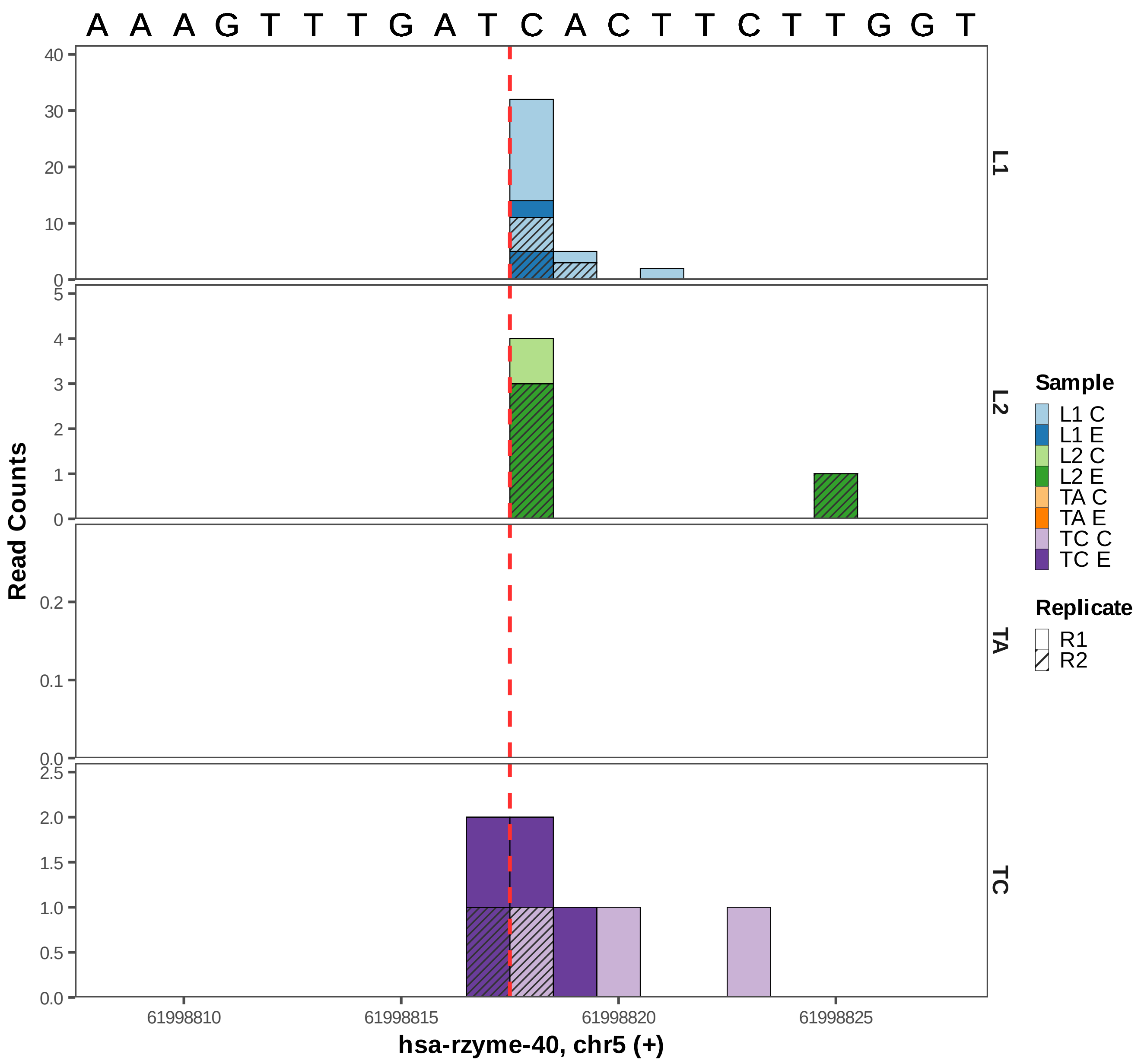
A A A G T T T G A T C A C T T C T T G G T
Read Counts
L1
L2
TA
TC
Sample
L1 C
L1 E
L2 C
L2 E
TA C
TA E
TC C
TC E
Replicate
R1
R2
61998810
61998815
61998820
61998825
hsa-rzyme-40, chr5 (+)

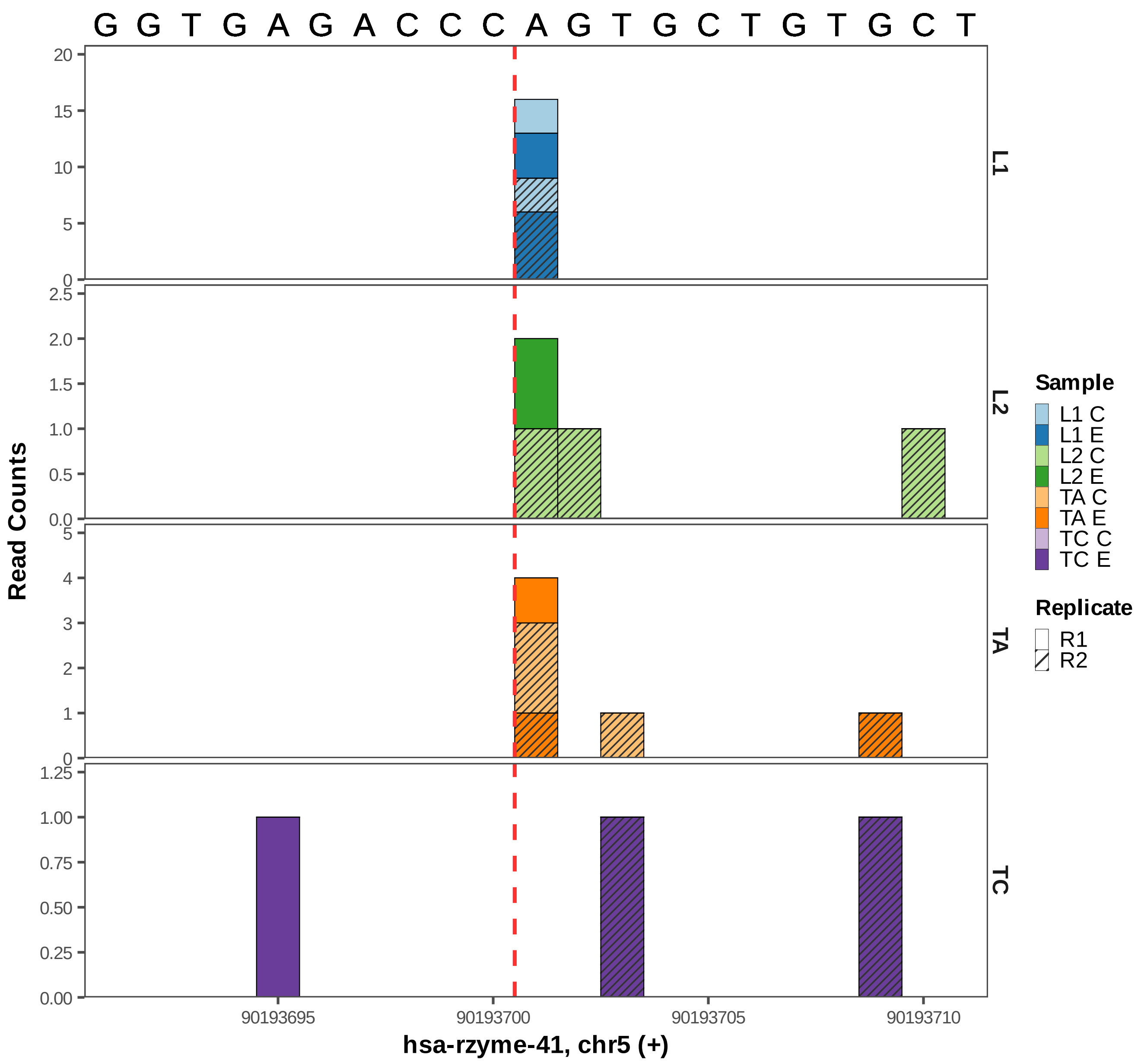

G G T G A G A C C C A G T G C T G T G C T
Read Counts
L1
L2
TA
TC
Sample
L1 C
L1 E
L2 C
L2 E
TA C
TA E
TC C
TC E
Replicate
R1
R2
90193695
90193700
90193705
90193710
hsa-rzyme-41, chr5 (+)

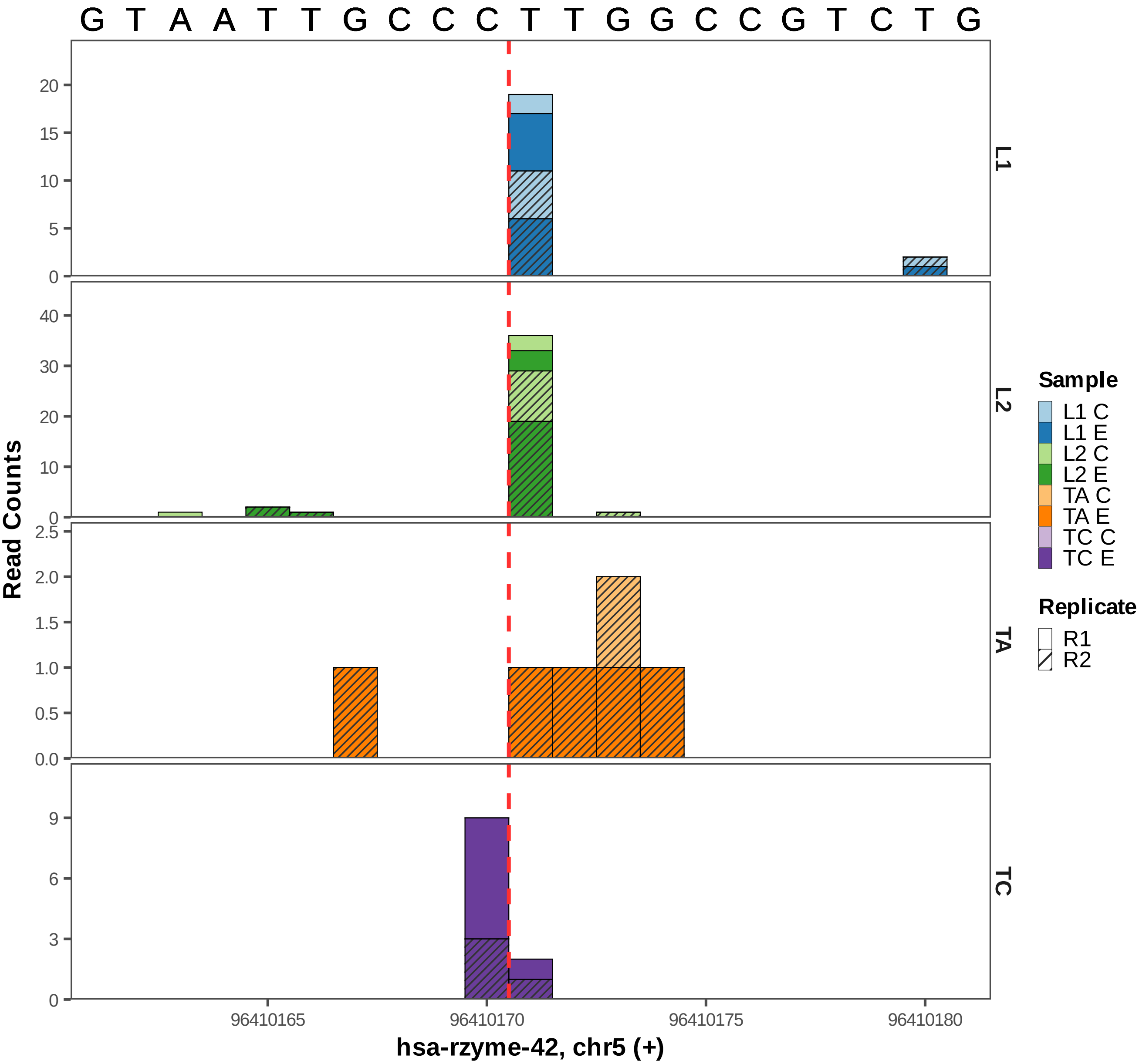

G T A A T T G C C C T T G G C C G T C T G
L1
L2
TA
TC
Read Counts
hsa-rzyme-42, chr5 (+)
96410165
96410170
96410175
96410180
Sample
L1 C
L1 E
L2 C
L2 E
TA C
TA E
TC C
TC E
Replicate
R1
R2

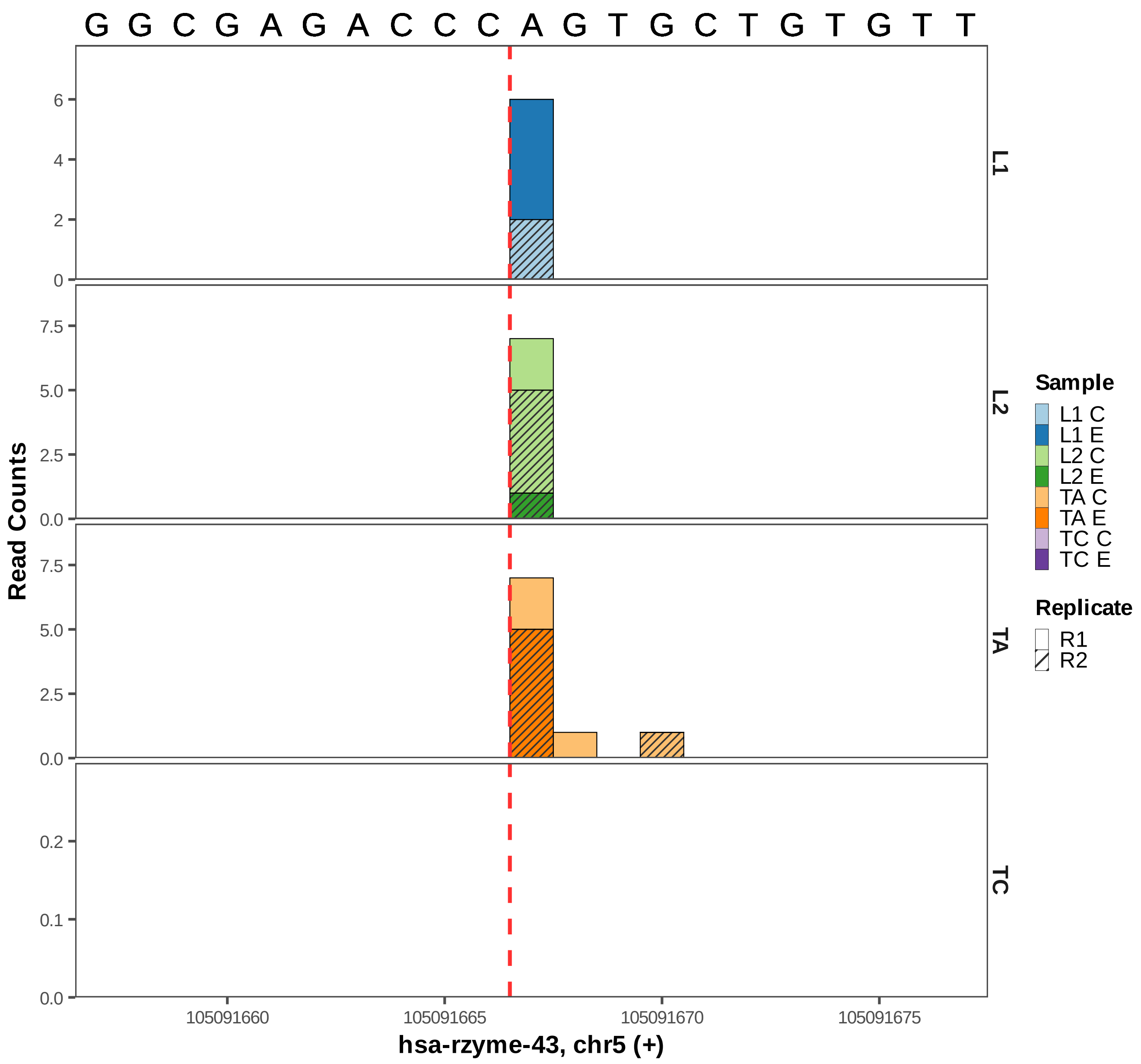

G G C G A G A C C C A G T G C T G T G T T
L1
L2
TA
TC
Read Counts
0
2
4
6
0.0
2.5
5.0
7.5
0.0
0.1
0.2
105091660
105091665
105091670
105091675
hsa-rzyme-43, chr5 (+)
Sample
L1 C
L1 E
L2 C
L2 E
TA C
TA E
TC C
TC E
Replicate
R1
R2

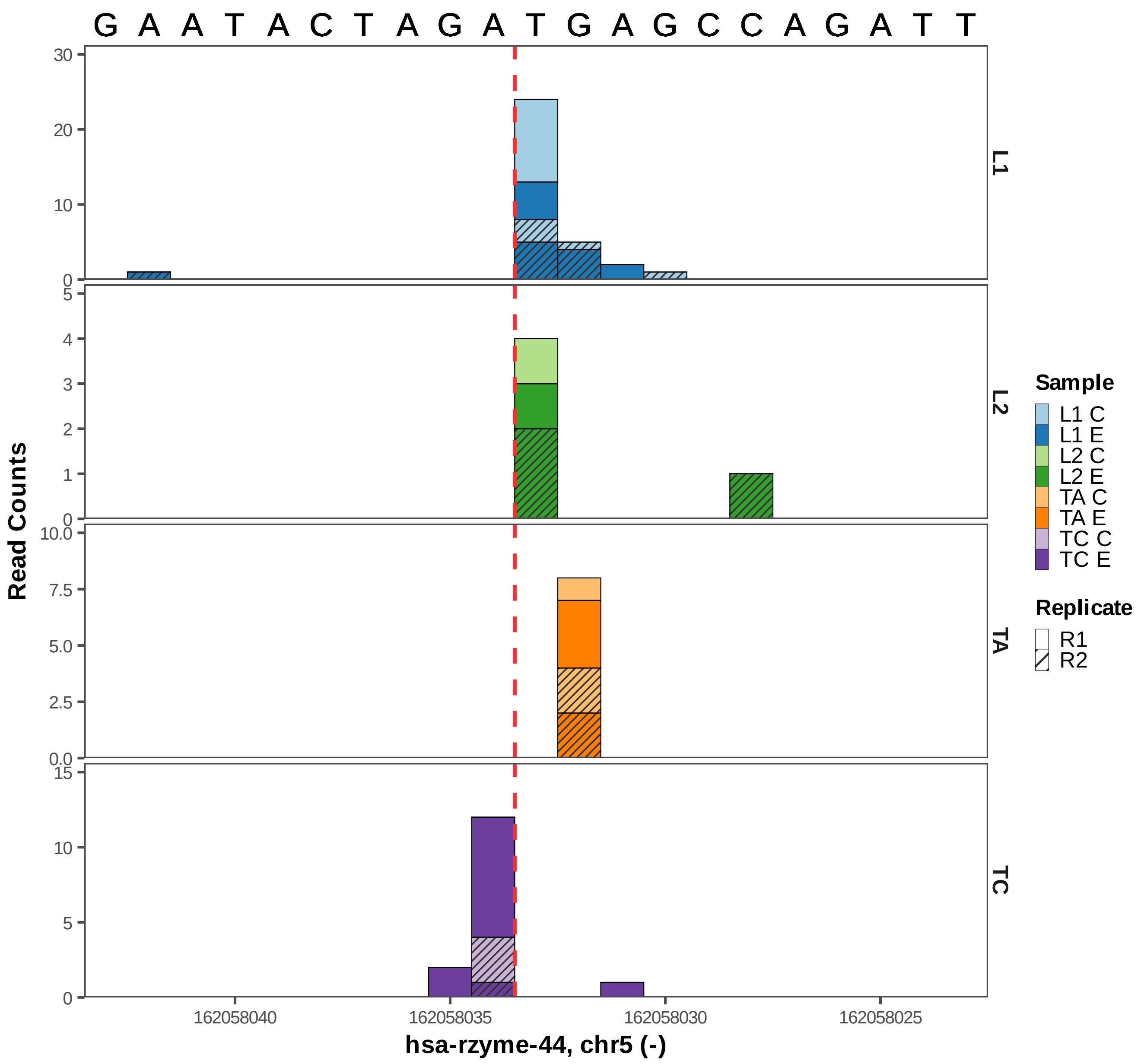

G A A T A C T A G A T G A G C C A G A T T
Read Counts
L1
L2
TA
TC
Sample
L1 C
L1 E
L2 C
L2 E
TA C
TA E
TC C
TC E
Replicate
R1
R2
162058040
162058035
162058030
162058025
hsa-rzyme-44, chr5 (-)

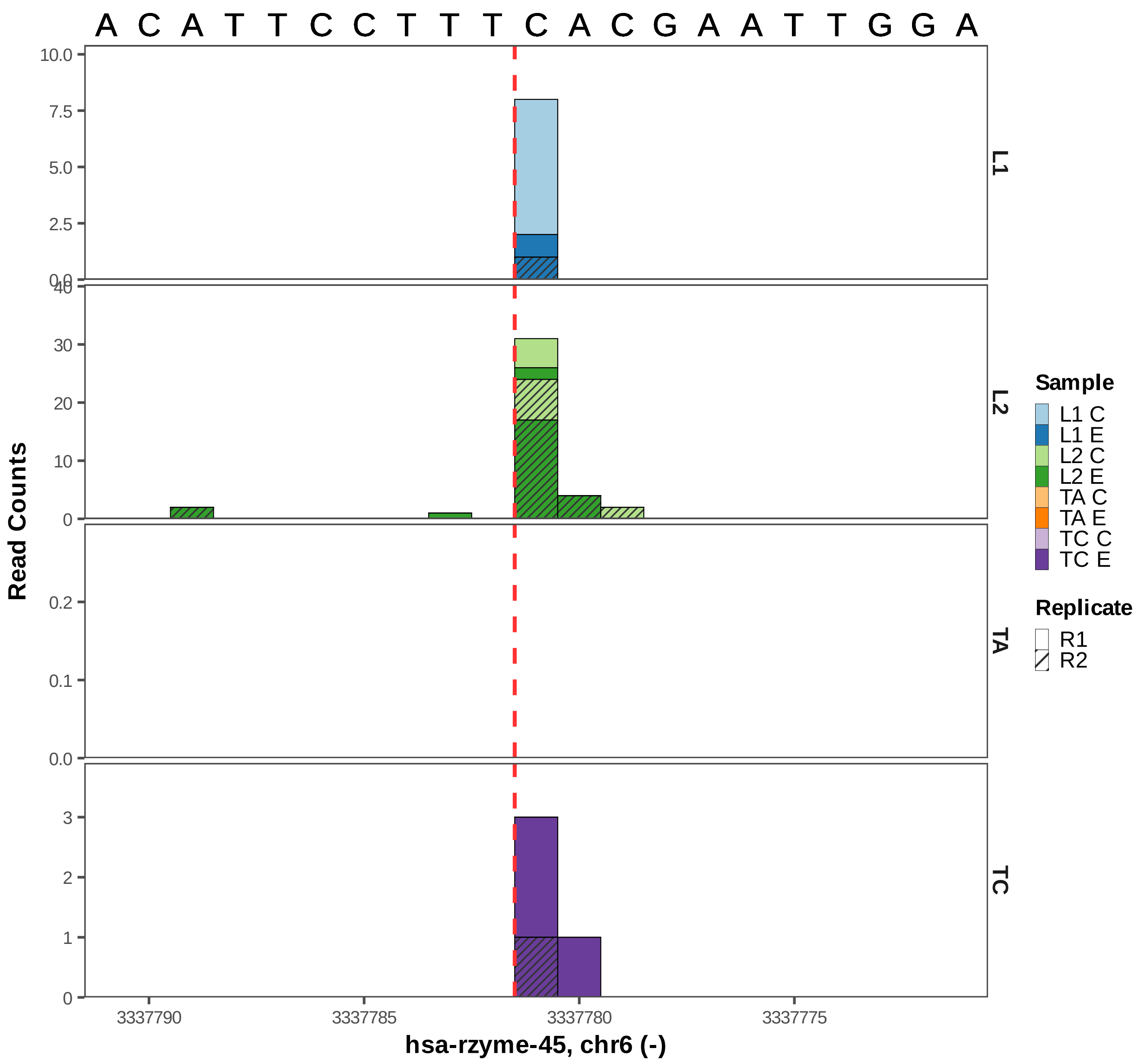

A C A T T C C T T T C A C G A A T T G G A
Read Counts
L1
L2
TA
TC
Sample
L1 C
L1 E
L2 C
L2 E
TA C
TA E
TC C
TC E
Replicate
R1
R2
3337790
3337785
3337780
3337775
hsa-rzyme-45, chr6 (-)

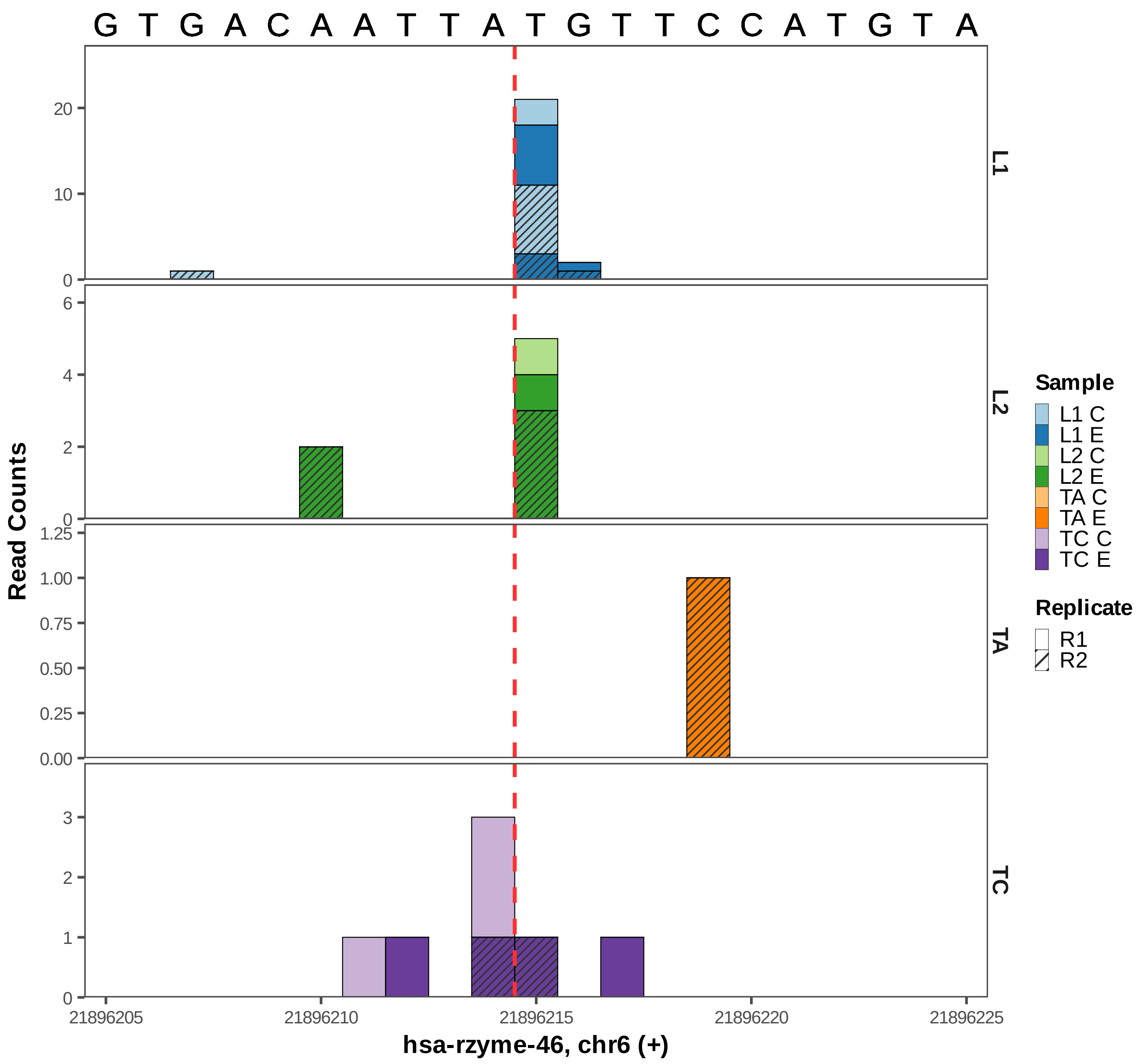
G T G A C A A T T A T G T T C C A T G T A
Read Counts
L1
L2
TA
TC
Sample
L1 C
L1 E
L2 C
L2 E
TA C
TA E
TC C
TC E
Replicate
R1
R2
21896205
21896210
21896215
21896220
21896225
hsa-rzyme-46, chr6 (+)

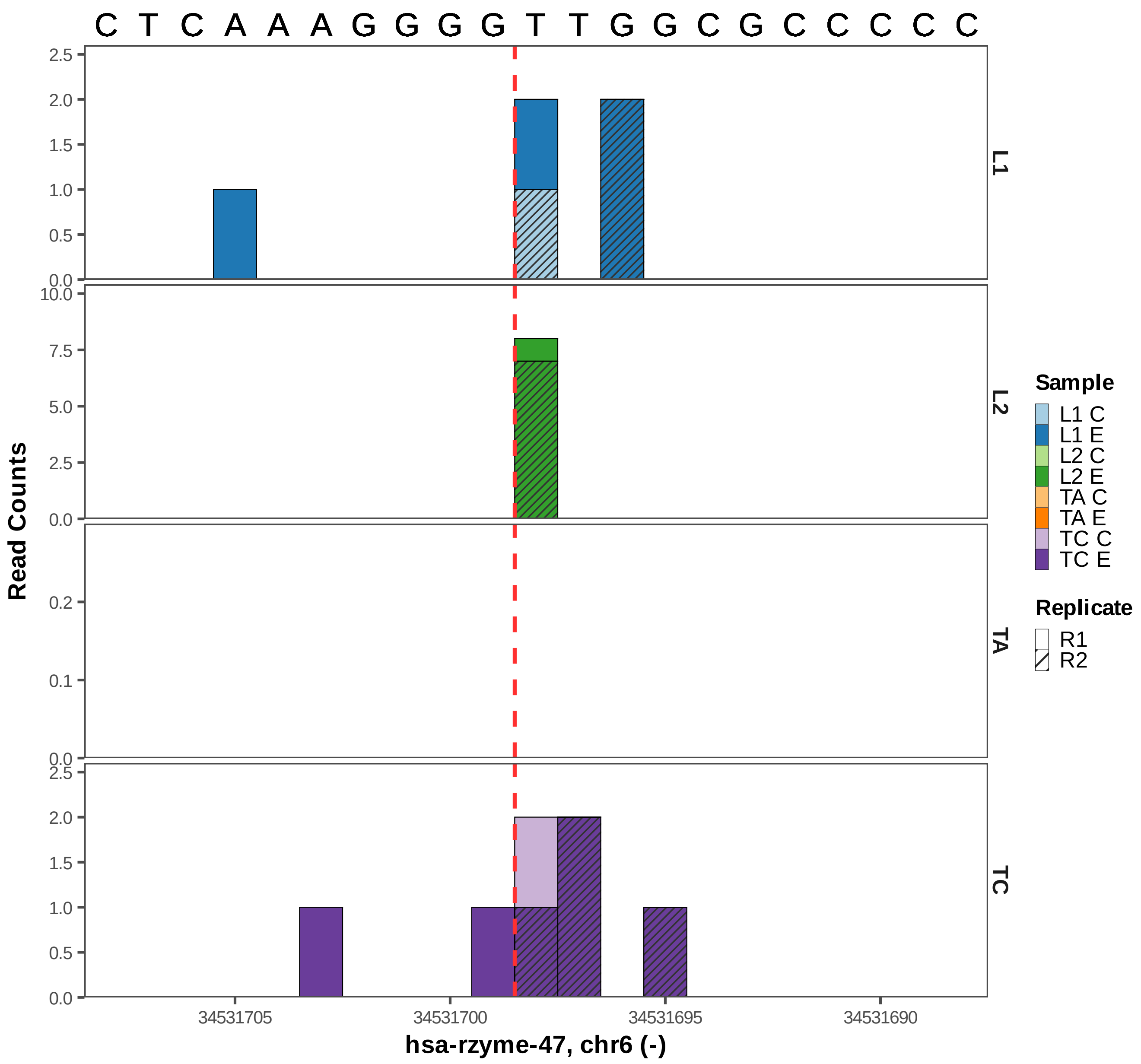

C T C A A A G G G G T T G G C G C C C C C
L1
L2
TA
TC
Read Counts
hsa-rzyme-47, chr6 (-)
34531705
34531700
34531695
34531690
Sample
L1 C
L1 E
L2 C
L2 E
TA C
TA E
TC C
TC E
Replicate
R1
R2

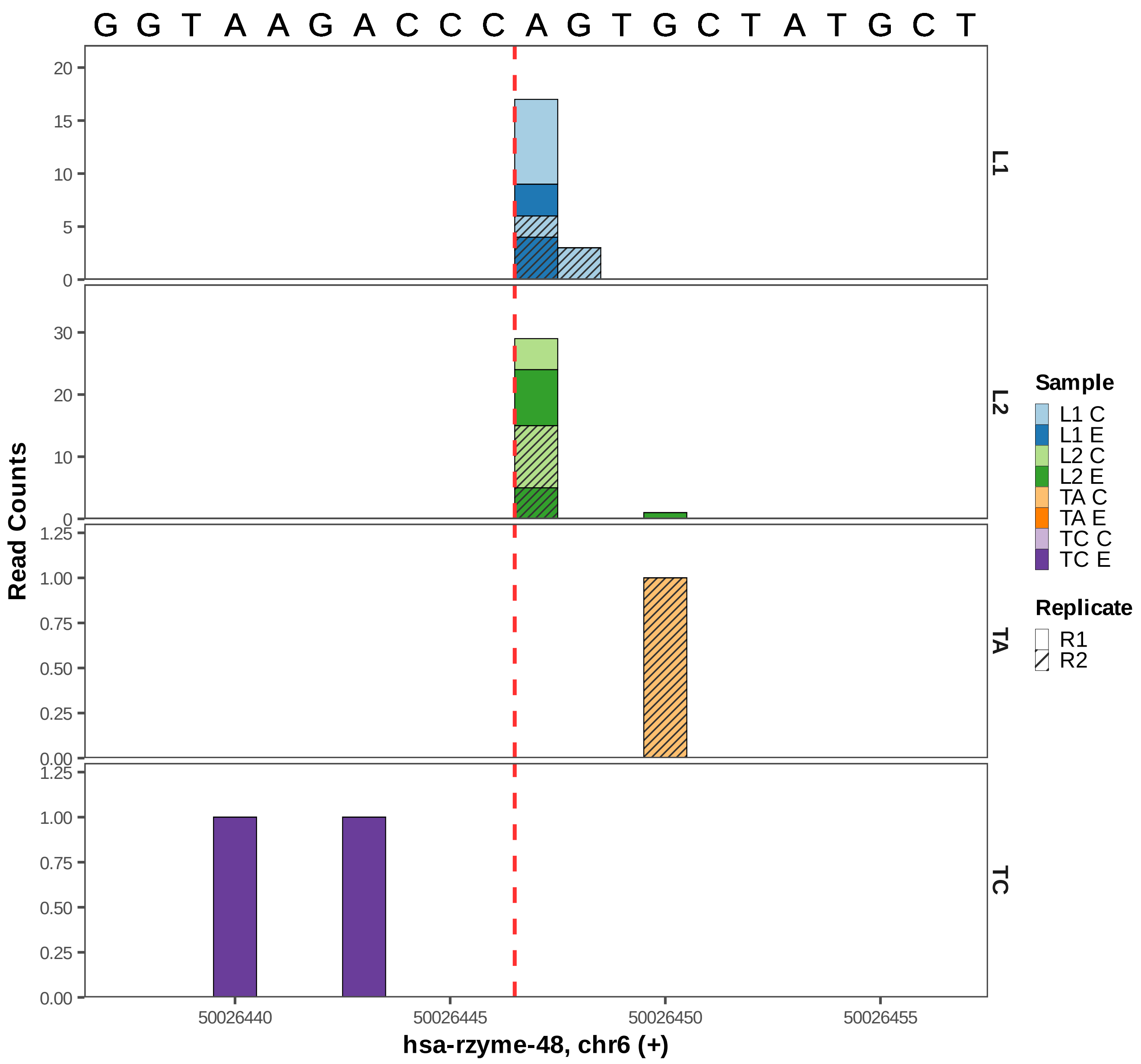

G G T A A G A C C C A G T G C T A T G C T
Read Counts
L1
L2
TA
TC
20
15
10
5
0
30
20
10
0
1.25
1.00
0.75
0.50
0.25
0.00
1.25
1.00
0.75
0.50
0.25
0.00
50026440
50026445
50026450
50026455
hsa-rzyme-48, chr6 (+)
Sample
L1 C
L1 E
L2 C
L2 E
TA C
TA E
TC C
TC E
Replicate
R1
R2

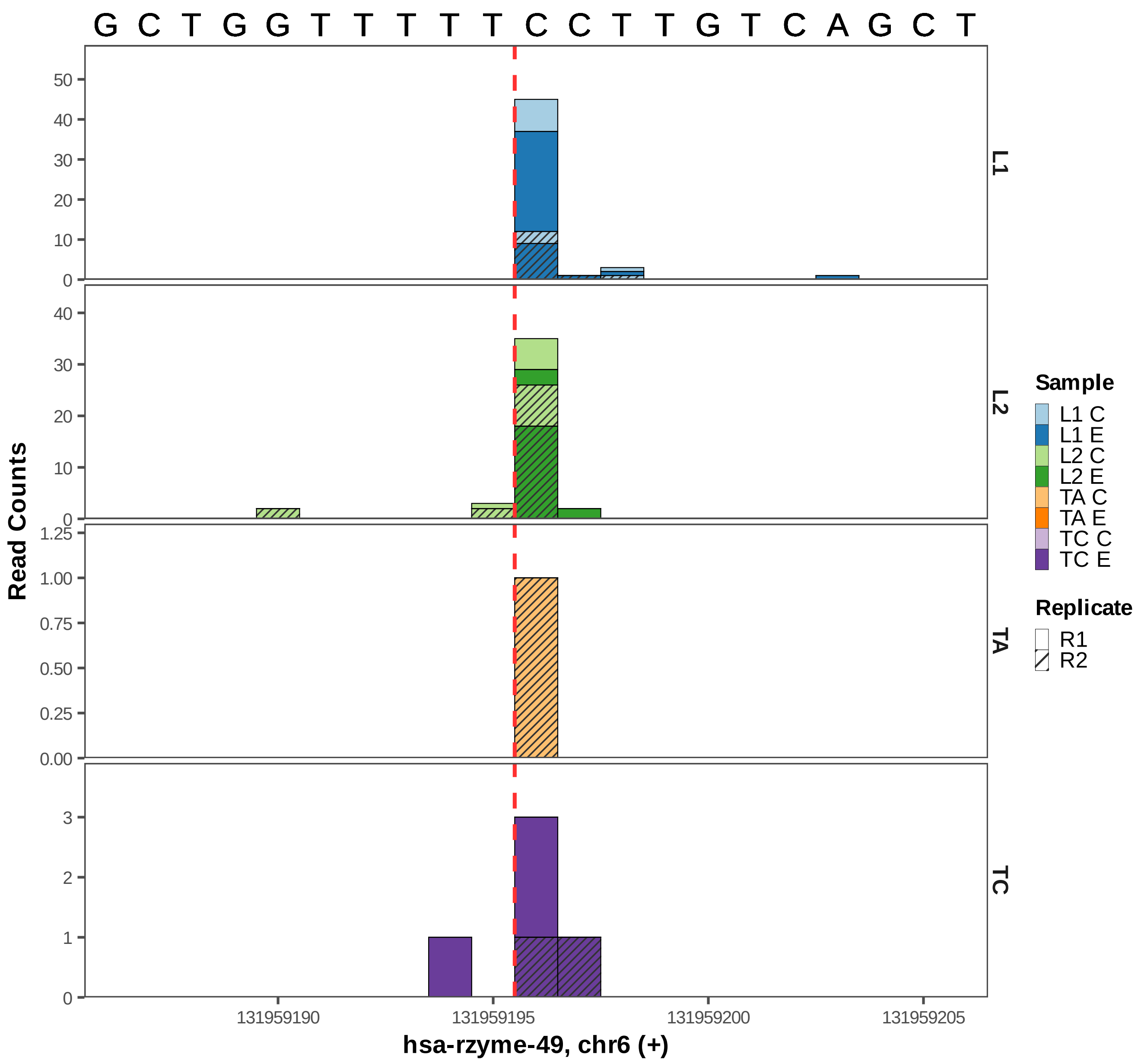

G C T G G T T T T T C C T T G T C A G C T
Read Counts
L1
L2
TA
TC
Sample
L1 C
L1 E
L2 C
L2 E
TA C
TA E
TC C
TC E
Replicate
R1
R2
131959190
131959195
131959200
131959205
hsa-rzyme-49, chr6 (+)

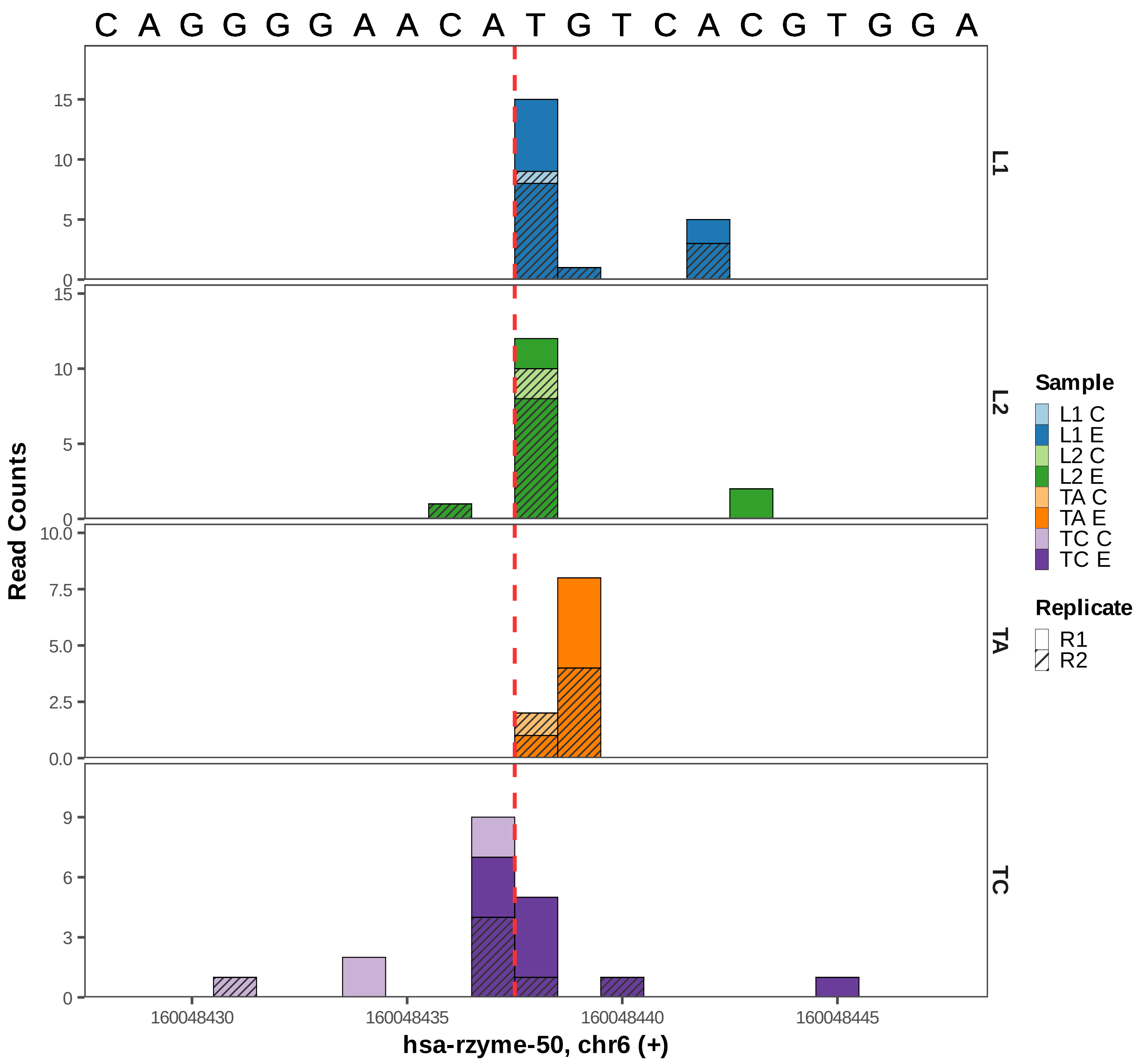
C A G G G G A A C A T G T C A C G T G G A
L1
L2
TA
TC
Read Counts
160048430
160048435
160048440
160048445
hsa-rzyme-50, chr6 (+)
Sample
L1 C
L1 E
L2 C
L2 E
TA C
TA E
TC C
TC E
Replicate
R1
R2

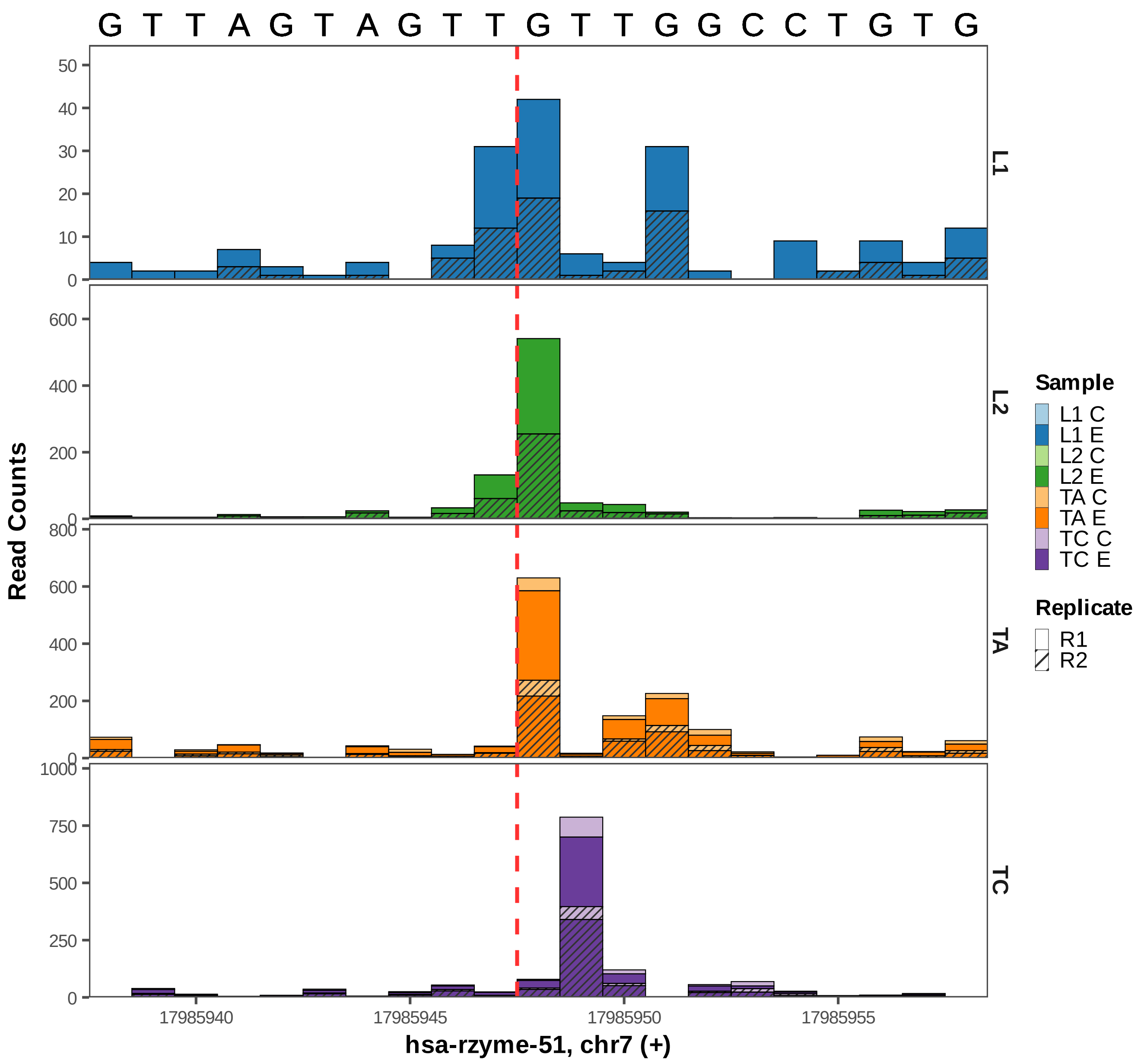

G T T A G T A G T T G T T G G C C T G T G
L1
L2
TA
TC
Read Counts
hsa-rzyme-51, chr7 (+)
17985940
17985945
17985950
17985955
Sample
L1 C
L1 E
L2 C
L2 E
TA C
TA E
TC C
TC E
Replicate
R1
R2

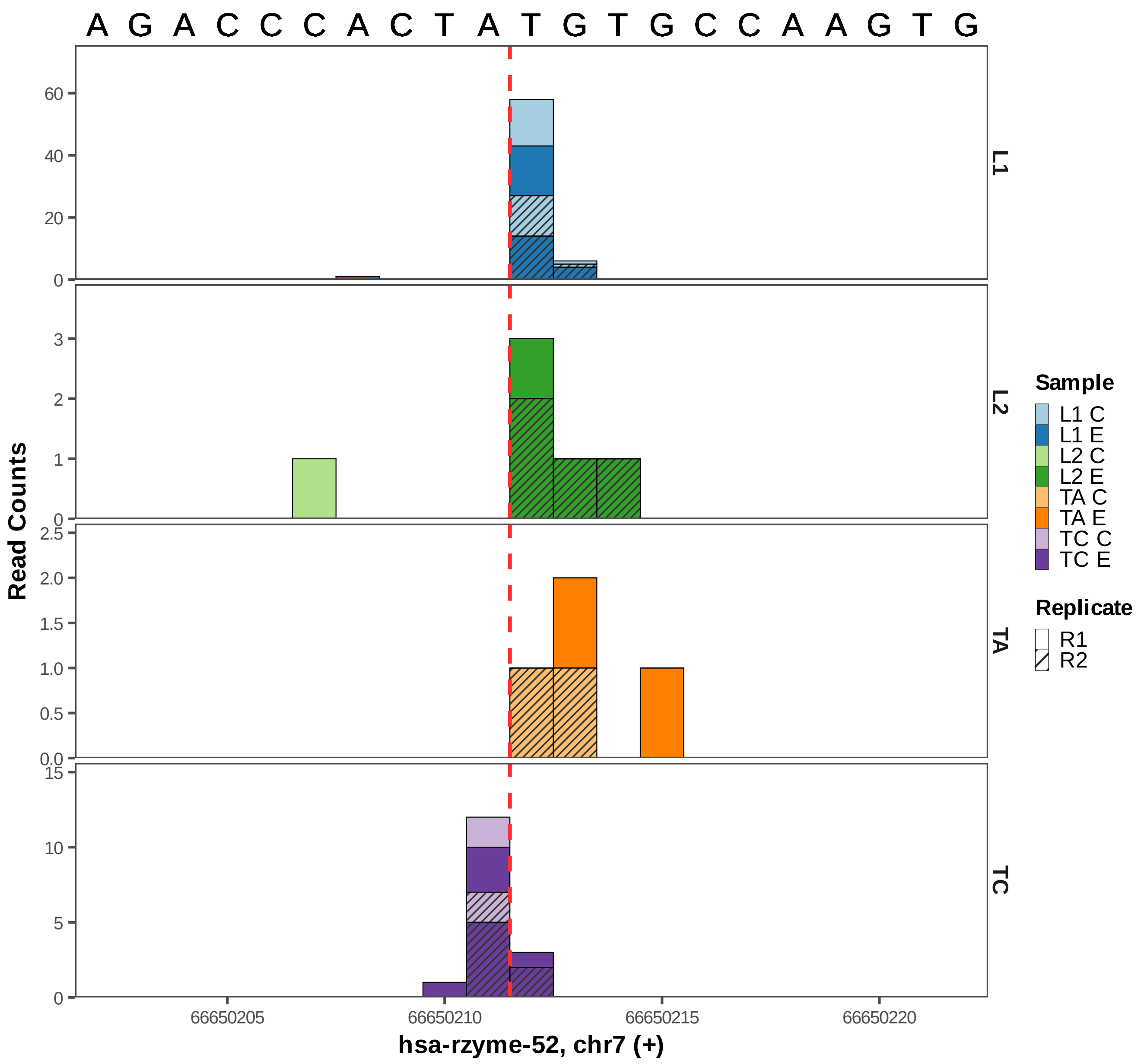
A G A C C C A C T A T G T G C C A A G T G
Read Counts
L1
L2
TA
TC
60
40
20
0
3
2
1
0
2.5
2.0
1.5
1.0
0.5
0.0
15
10
5
0
66650205
66650210
66650215
66650220
hsa-rzyme-52, chr7 (+)
Sample
L1 C
L1 E
L2 C
L2 E
TA C
TA E
TC C
TC E
Replicate
R1
R2

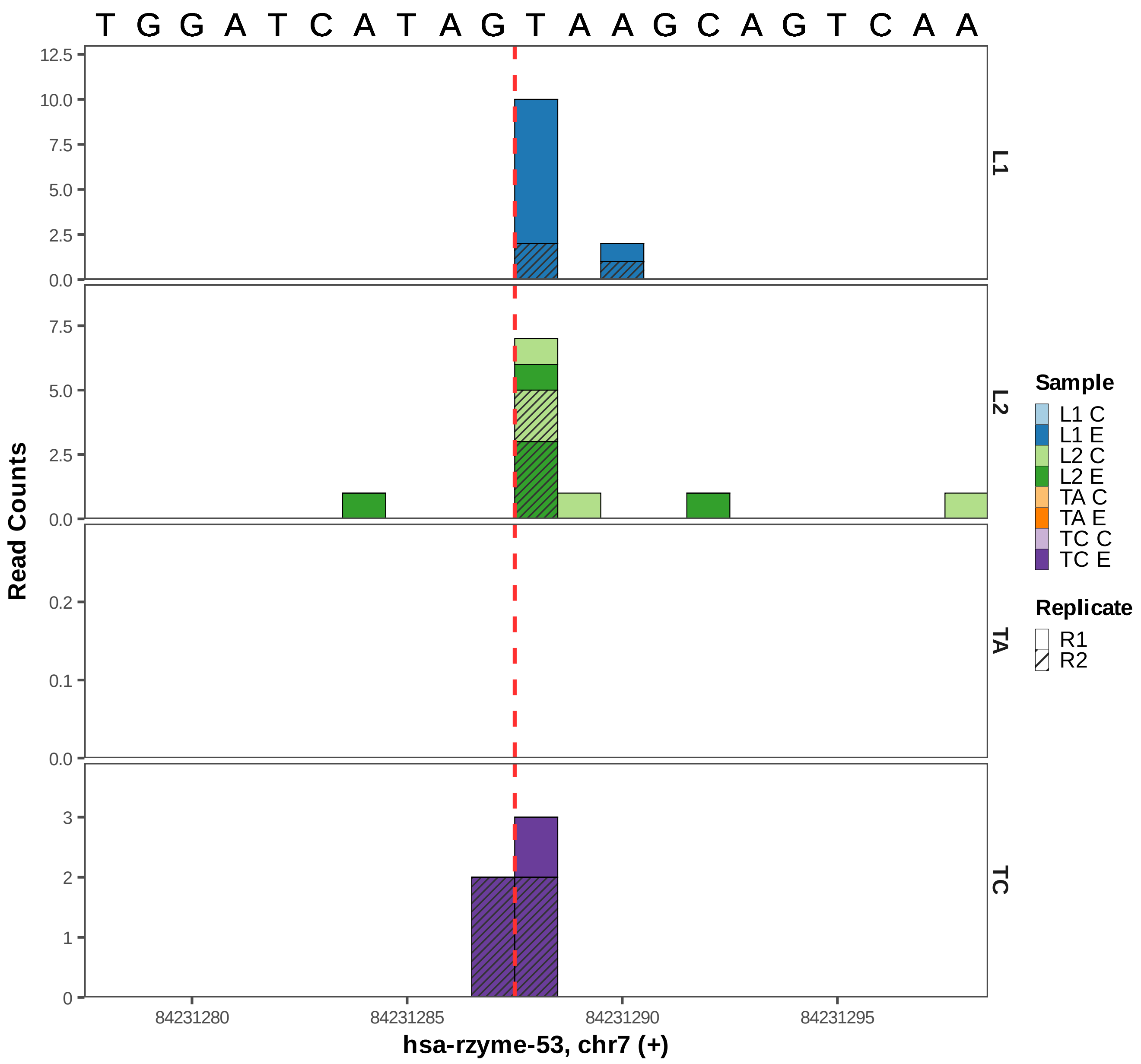
T G G A T C A T A G T A A G C A G T C A A
Read Counts
L1
L2
TA
TC
12.5
10.0
7.5
5.0
2.5
0.0
0.2
0.1
3
2
1
0
84231280
84231285
84231290
84231295
hsa-rzyme-53, chr7 (+)
Sample
L1 C
L1 E
L2 C
L2 E
TA C
TA E
TC C
TC E
Replicate
R1
R2

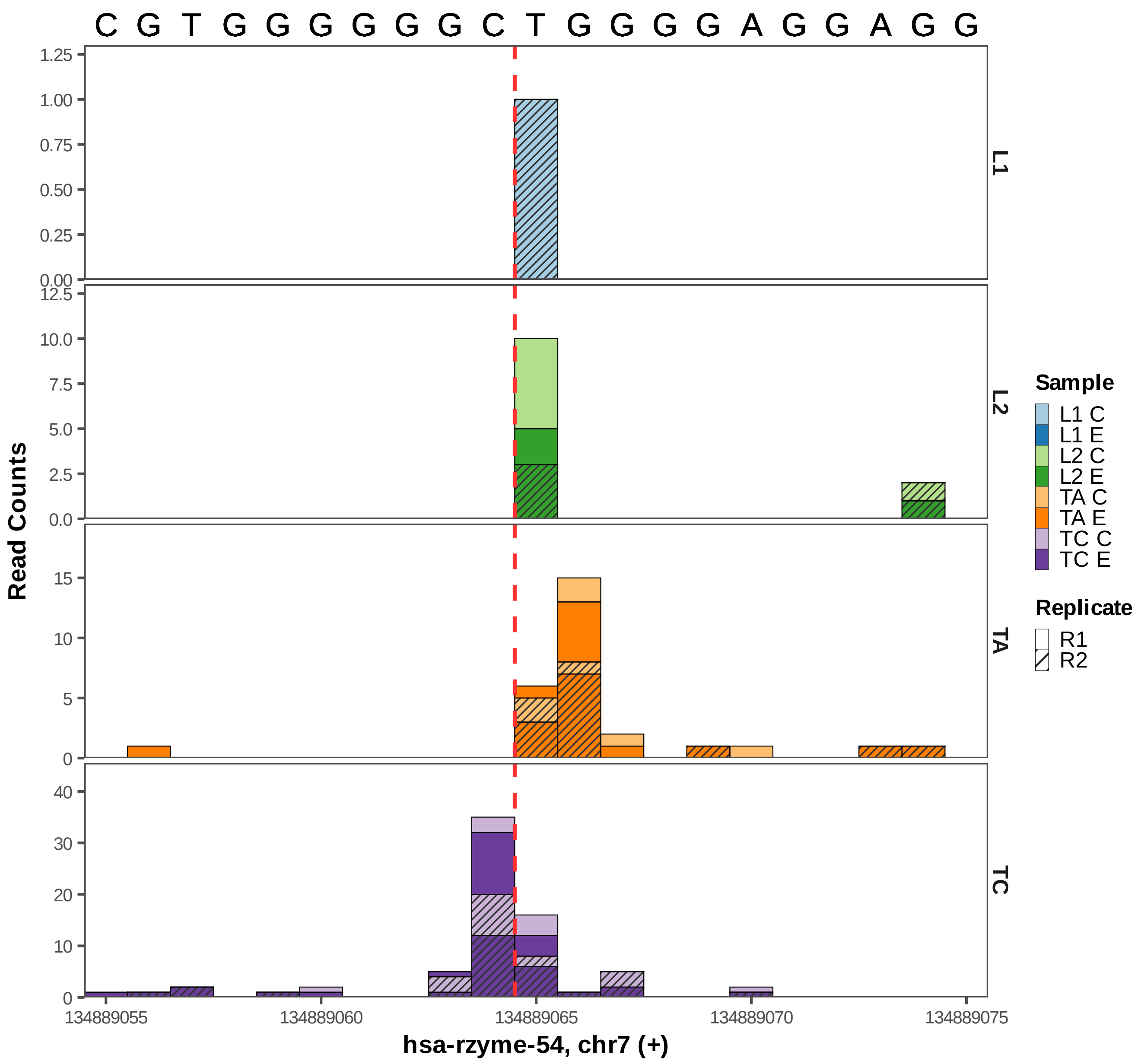

C G T G G G G G G C T G G G G A G G A G G
1.25
1.00
0.75
0.50
0.25
0.00
12.5
10.0
7.5
5.0
2.5
0.0
15
10
5
0
40
30
20
10
0
L1
L2
TA
TC
Read Counts
Sample
L1 C
L1 E
L2 C
L2 E
TA C
TA E
TC C
TC E
Replicate
R1
R2
134889055
134889060
134889065
134889070
134889075
hsa-rzyme-54, chr7 (+)

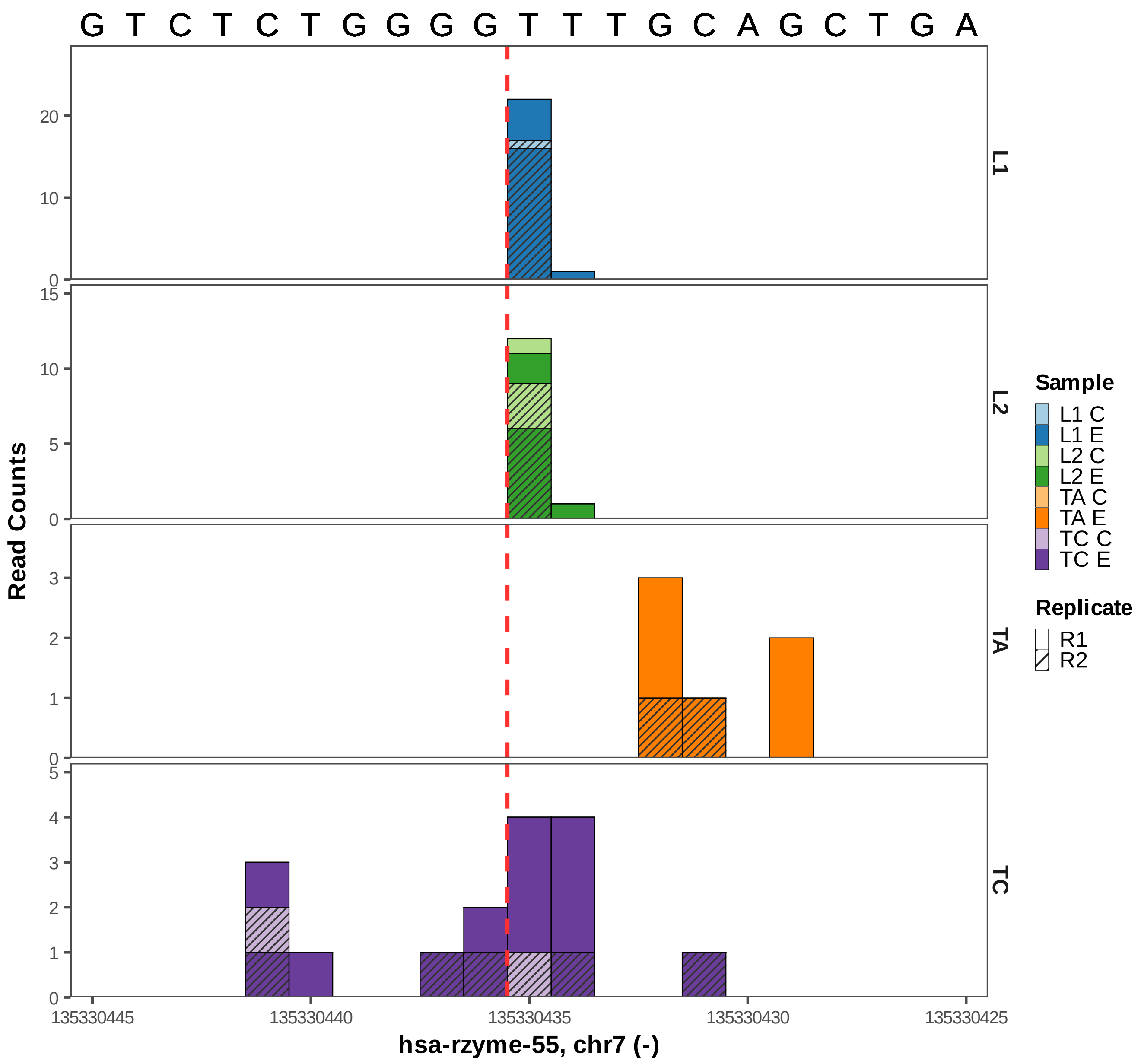
G T C T C T G G G G T T T G C A G C T G A
L1
L2
TA
TC
20
10
0
15
5
3
2
1
4
Read Counts
135330445
135330440
135330435
135330430
135330425
hsa-rzyme-55, chr7 (-)
Sample
L1 C
L1 E
L2 C
L2 E
TA C
TA E
TC C
TC E
Replicate
R1
R2

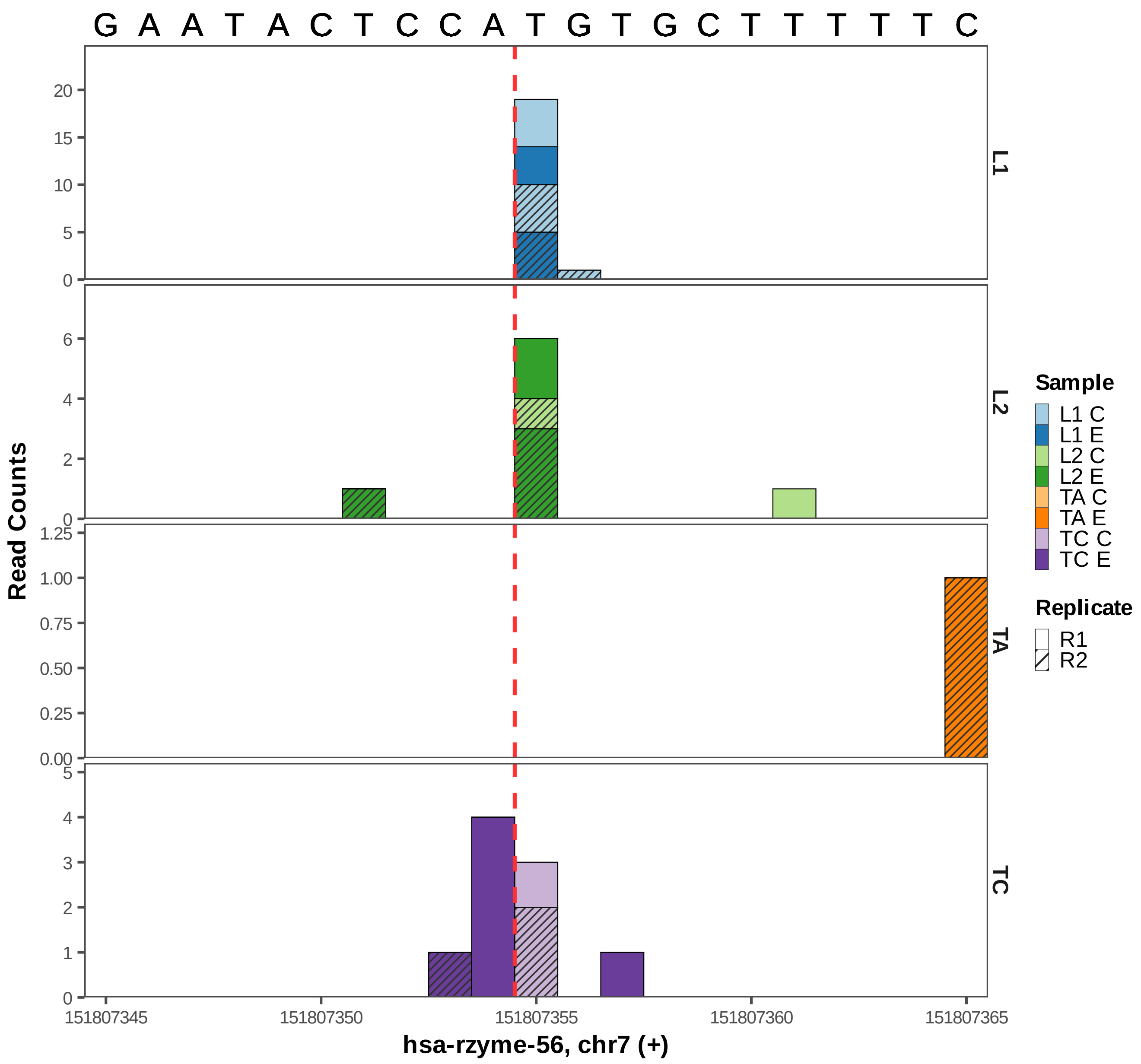

G A A T A C T C C A T G T G C T T T T T C
Read Counts
L1
L2
TA
TC
20
15
10
5
0
6
4
2
0
1.25
1.00
0.75
0.50
0.25
0.00
5
4
3
2
1
0
151807345
151807350
151807355
151807360
151807365
hsa-rzyme-56, chr7 (+)
Sample
L1 C
L1 E
L2 C
L2 E
TA C
TA E
TC C
TC E
Replicate
R1
R2

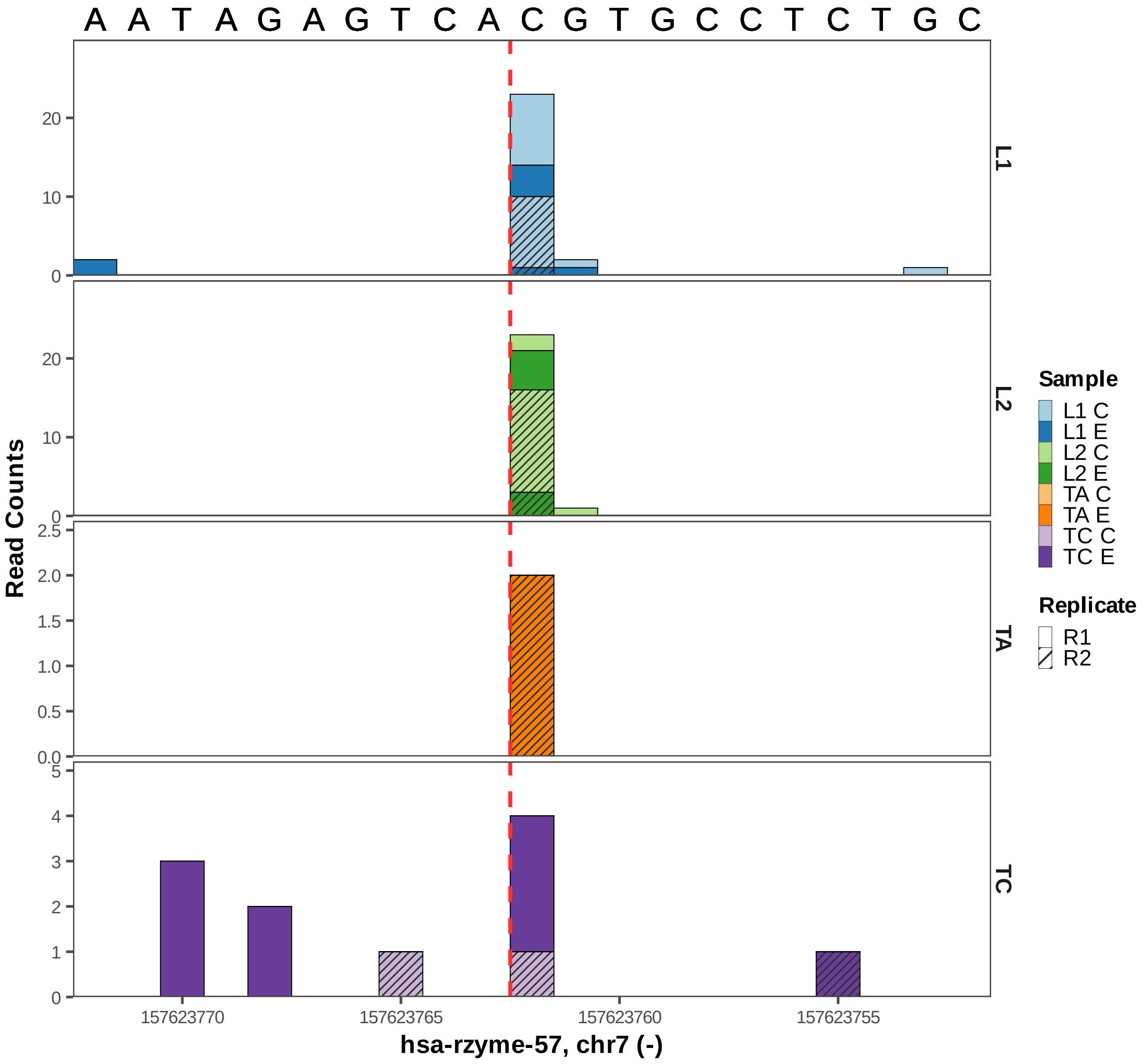

A A T A G A G T C A C G T G C C T C T G C
Read Counts
L1
L2
TA
TC
Sample
L1 C
L1 E
L2 C
L2 E
TA C
TA E
TC C
TC E
Replicate
R1
R2
157623770
157623765
157623760
157623755
hsa-rzyme-57, chr7 (-)

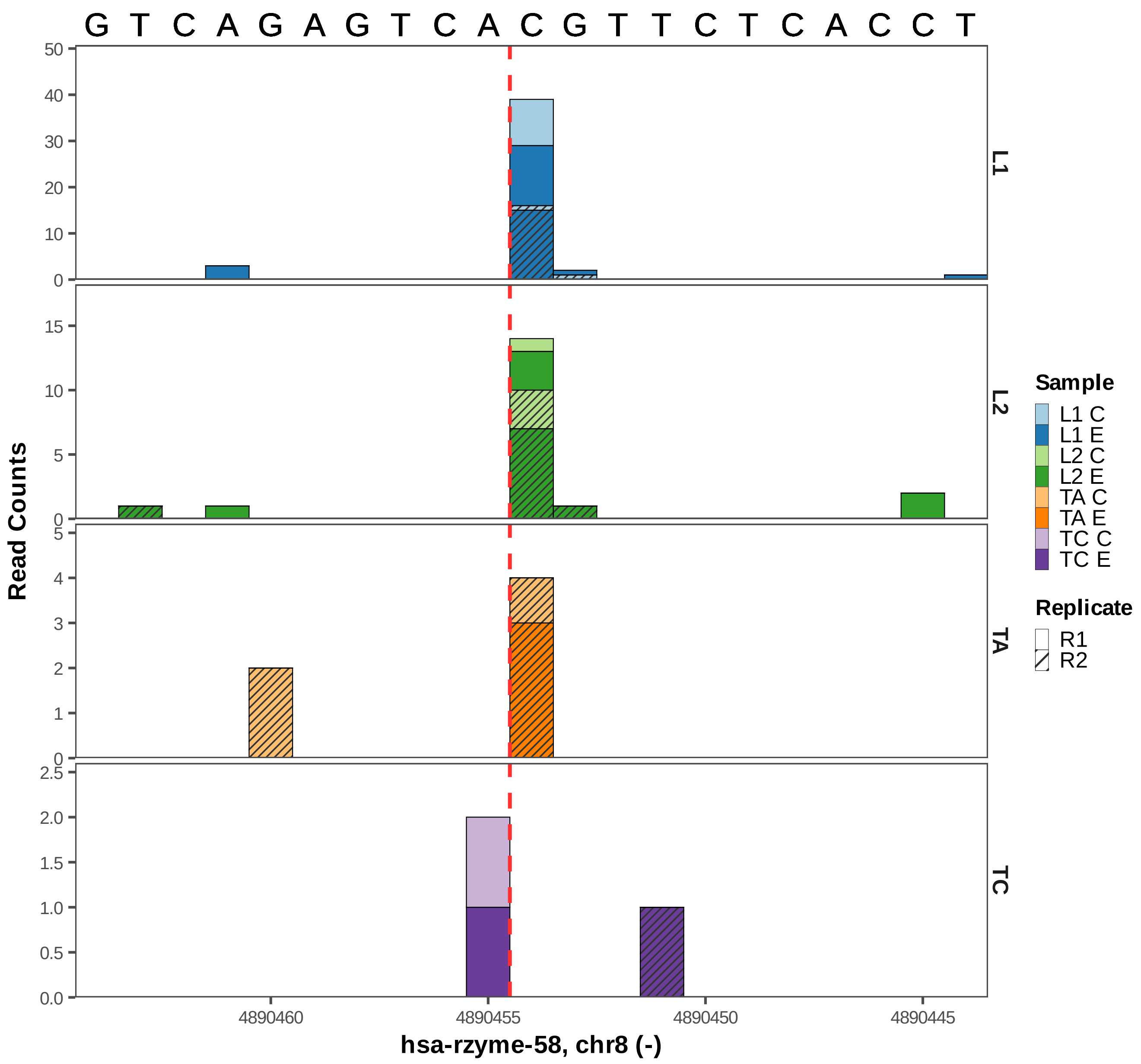

G T C A G A G T C A C G T T C T C A C C T
Read Counts
L1
L2
TA
TC
4890460
4890455
4890450
4890445
hsa-rzyme-58, chr8 (-)
Sample
L1 C
L1 E
L2 C
L2 E
TA C
TA E
TC C
TC E
Replicate
R1
R2

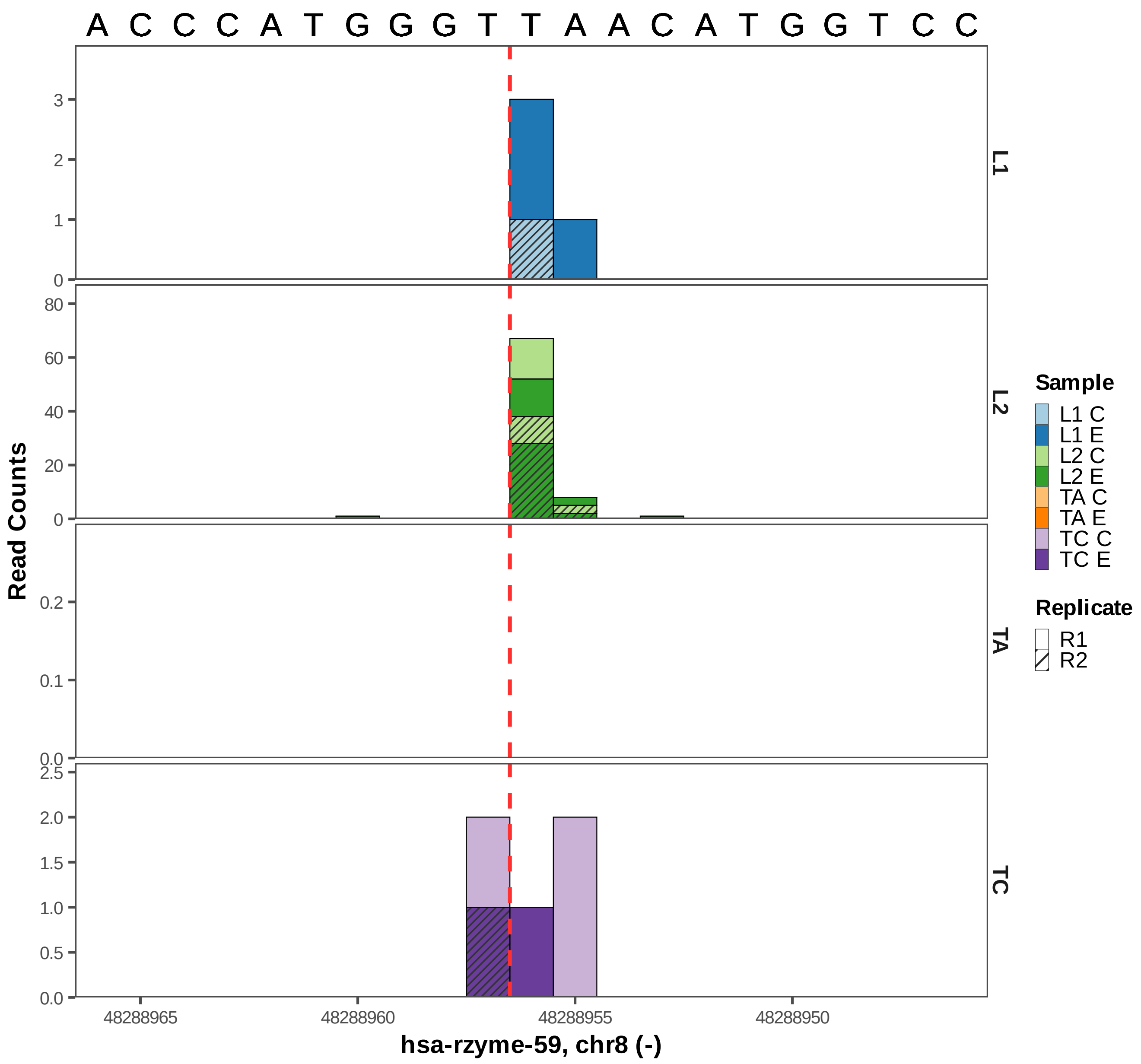

A C C C A T G G G T T A A C A T G G T C C
Read Counts
L1
L2
TA
TC
0
1
2
3
0
20
40
60
80
0.0
0.1
0.2
0.0
0.5
1.0
1.5
2.0
2.5
48288965
48288960
48288955
48288950
hsa-rzyme-59, chr8 (-)
Sample
L1 C
L1 E
L2 C
L2 E
TA C
TA E
TC C
TC E
Replicate
R1
R2

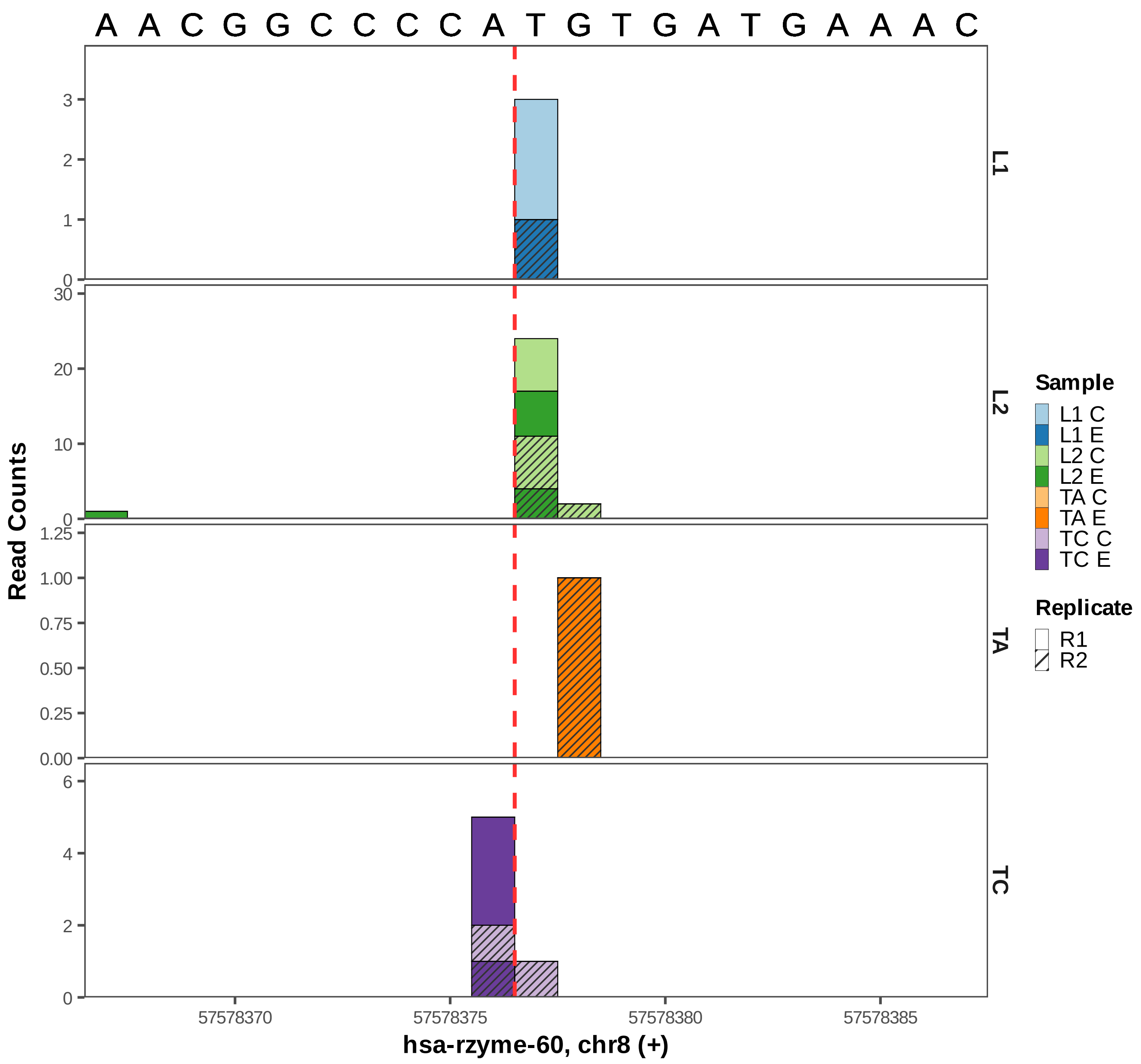
A A C G G C C C C A T G T G A T G A A A C
L1
L2
TA
TC
Read Counts
0
1
2
3
0
10
20
30
0.00
0.25
0.50
0.75
1.00
1.25
0
2
4
6
57578370
57578375
57578380
57578385
hsa-rzyme-60, chr8 (+)
Sample
L1 C
L1 E
L2 C
L2 E
TA C
TA E
TC C
TC E
Replicate
R1
R2

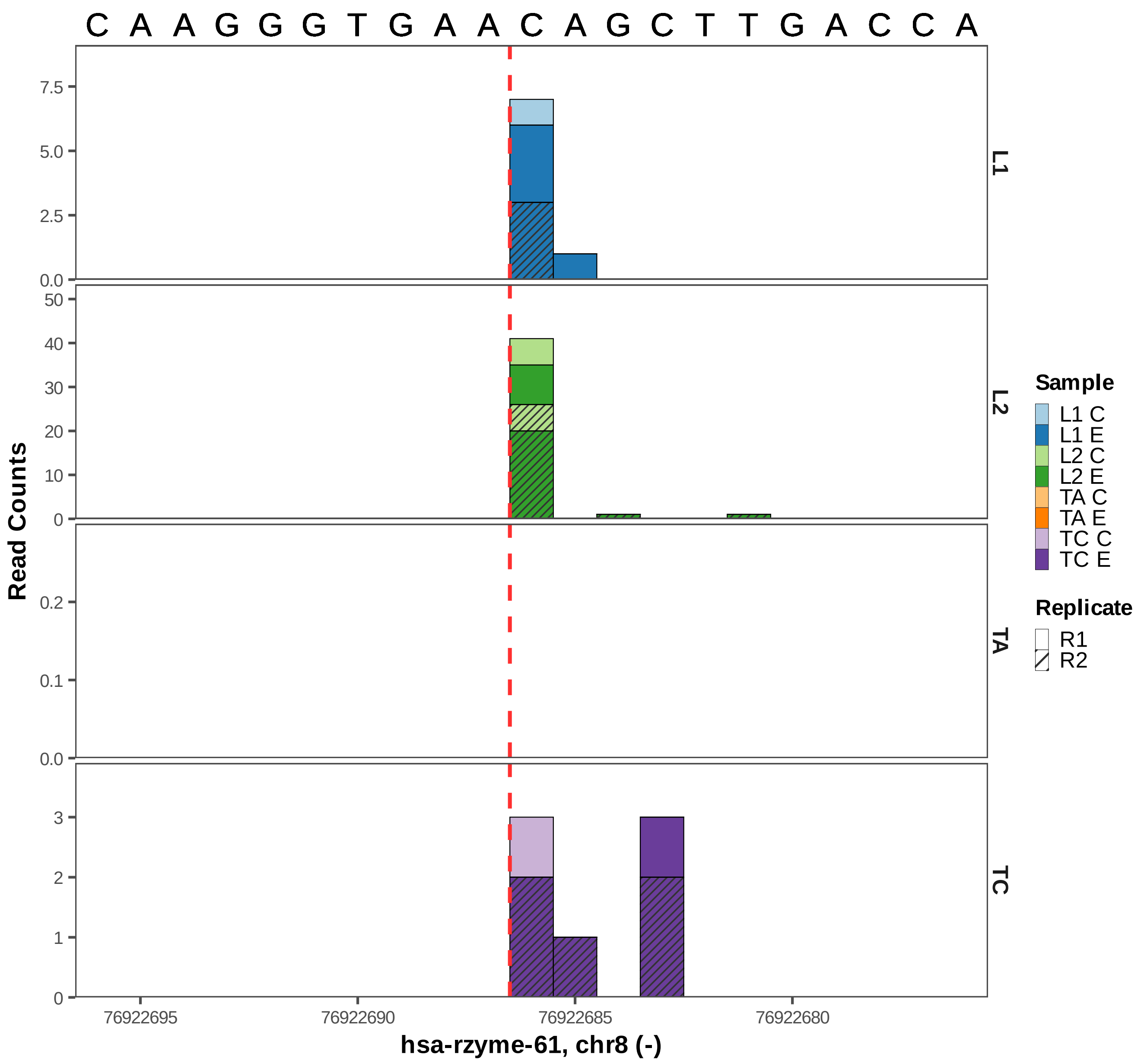

C A A G G G T G A A C A G C T T G A C C A
Read Counts
L1
L2
TA
TC
7.5
5.0
2.5
0.0
50
40
30
20
10
0
0.2
0.1
0.0
3
2
1
0
76922695
76922690
76922685
76922680
hsa-rzyme-61, chr8 (-)
Sample
L1 C
L1 E
L2 C
L2 E
TA C
TA E
TC C
TC E
Replicate
R1
R2

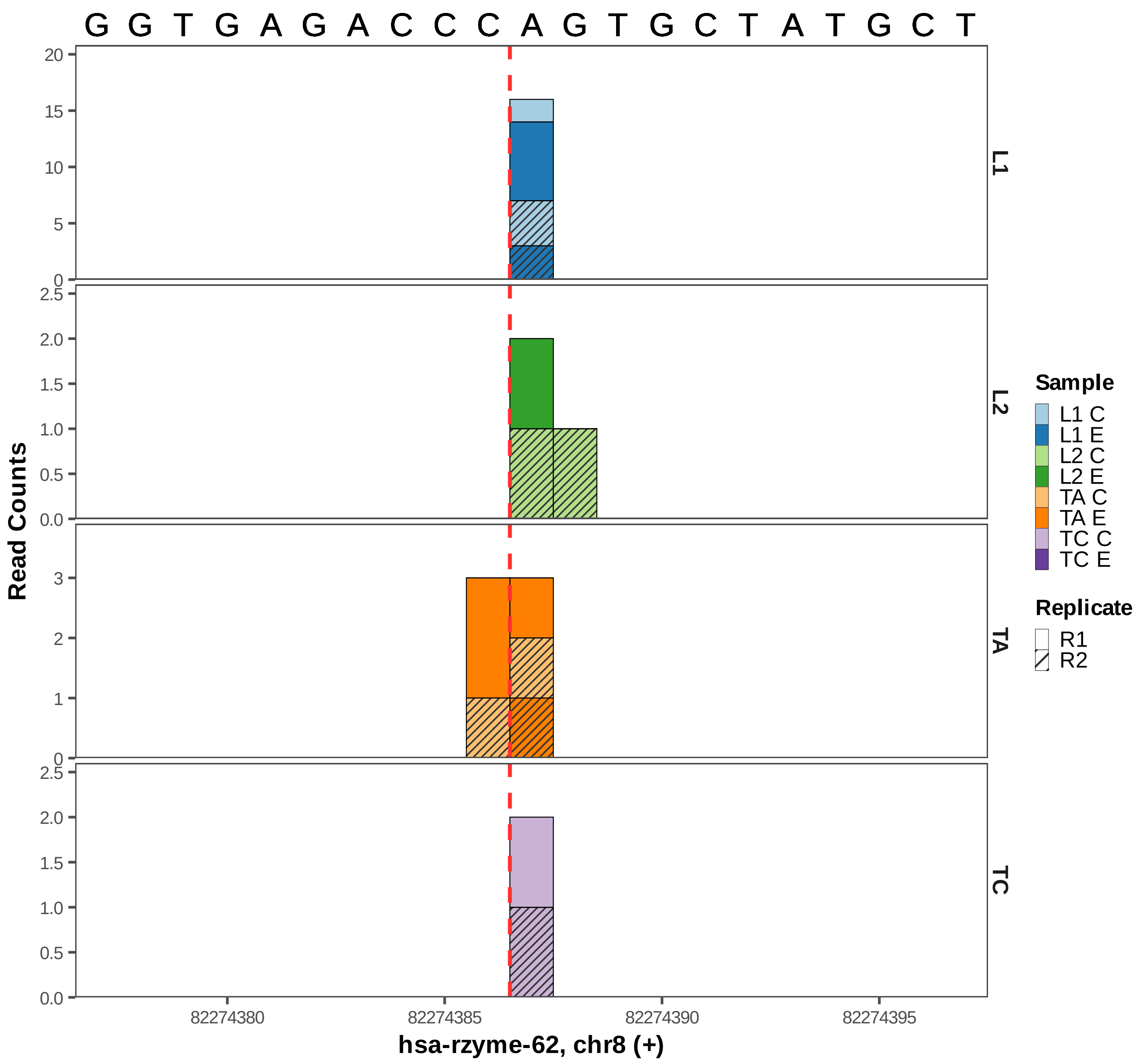

G G T G A G A C C C A G T G C T A T G C T
Read Counts
L1
L2
TA
TC
20
15
10
5
0
2.5
2.0
1.5
1.0
0.5
0.0
3
2
1
0
82274380
82274385
82274390
82274395
hsa-rzyme-62, chr8 (+)
Sample
L1 C
L1 E
L2 C
L2 E
TA C
TA E
TC C
TC E
Replicate
R1
R2

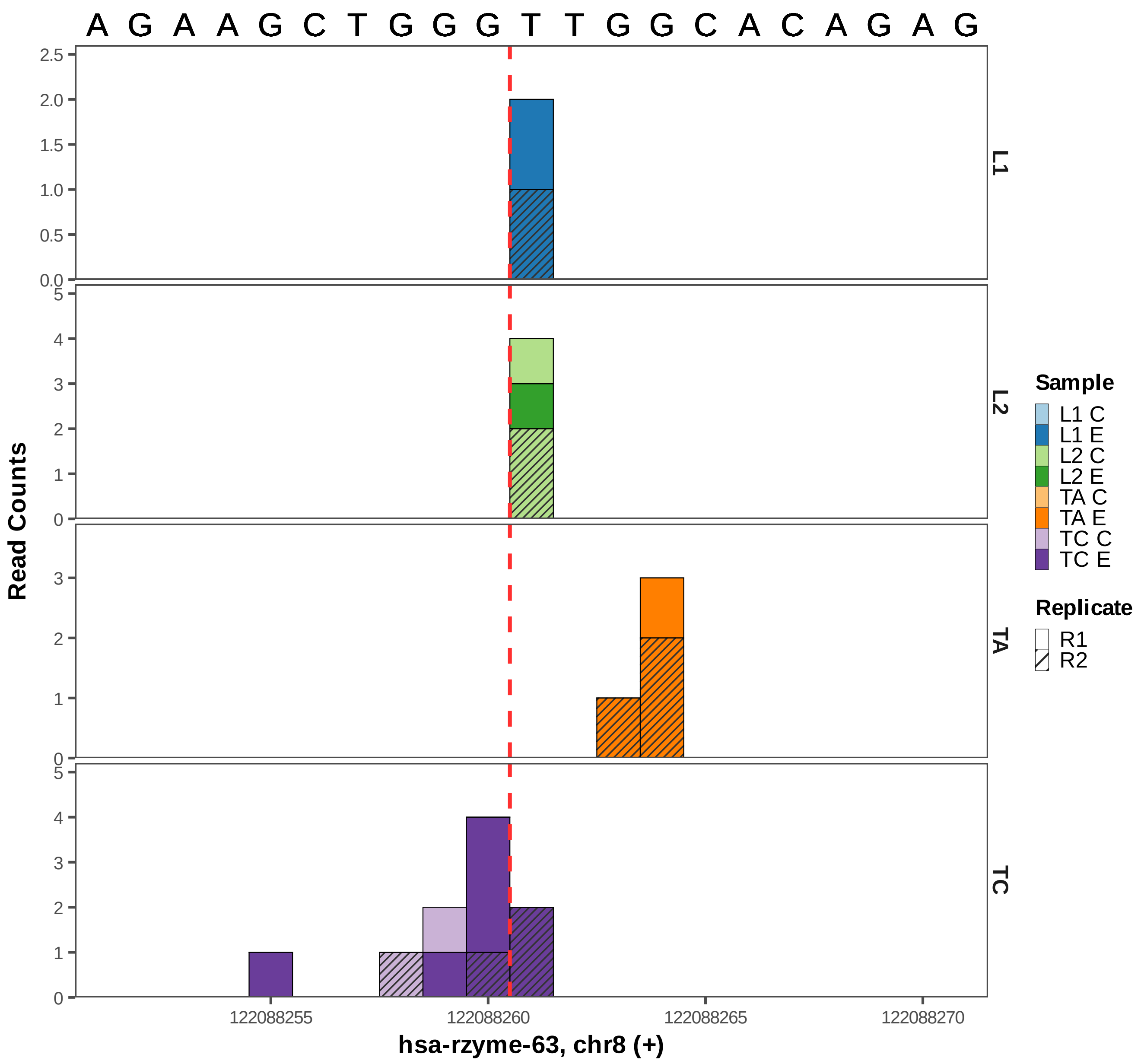

A G A A G C T G G G T T G G C A C A G A G
Read Counts
L1
L2
TA
TC
Sample
L1 C
L1 E
L2 C
L2 E
TA C
TA E
TC C
TC E
Replicate
R1
R2
122088255
122088260
122088265
122088270
hsa-rzyme-63, chr8 (+)

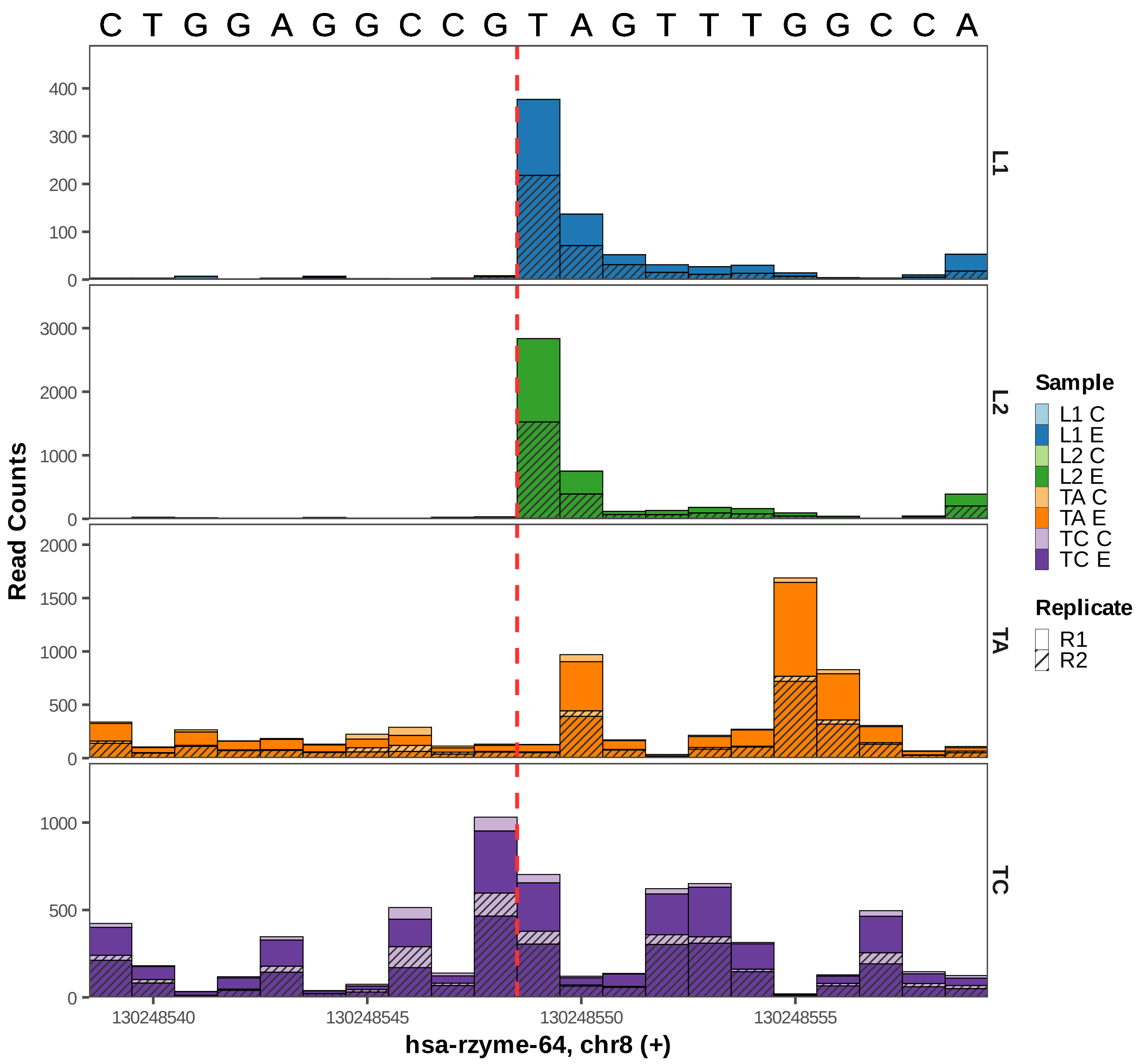

C T G G A G G C C G T A G T T T G G C C A
L1
L2
TA
TC
Read Counts
130248540
130248545
130248550
130248555
hsa-rzyme-64, chr8 (+)
Sample
L1 C
L1 E
L2 C
L2 E
TA C
TA E
TC C
TC E
Replicate
R1
R2

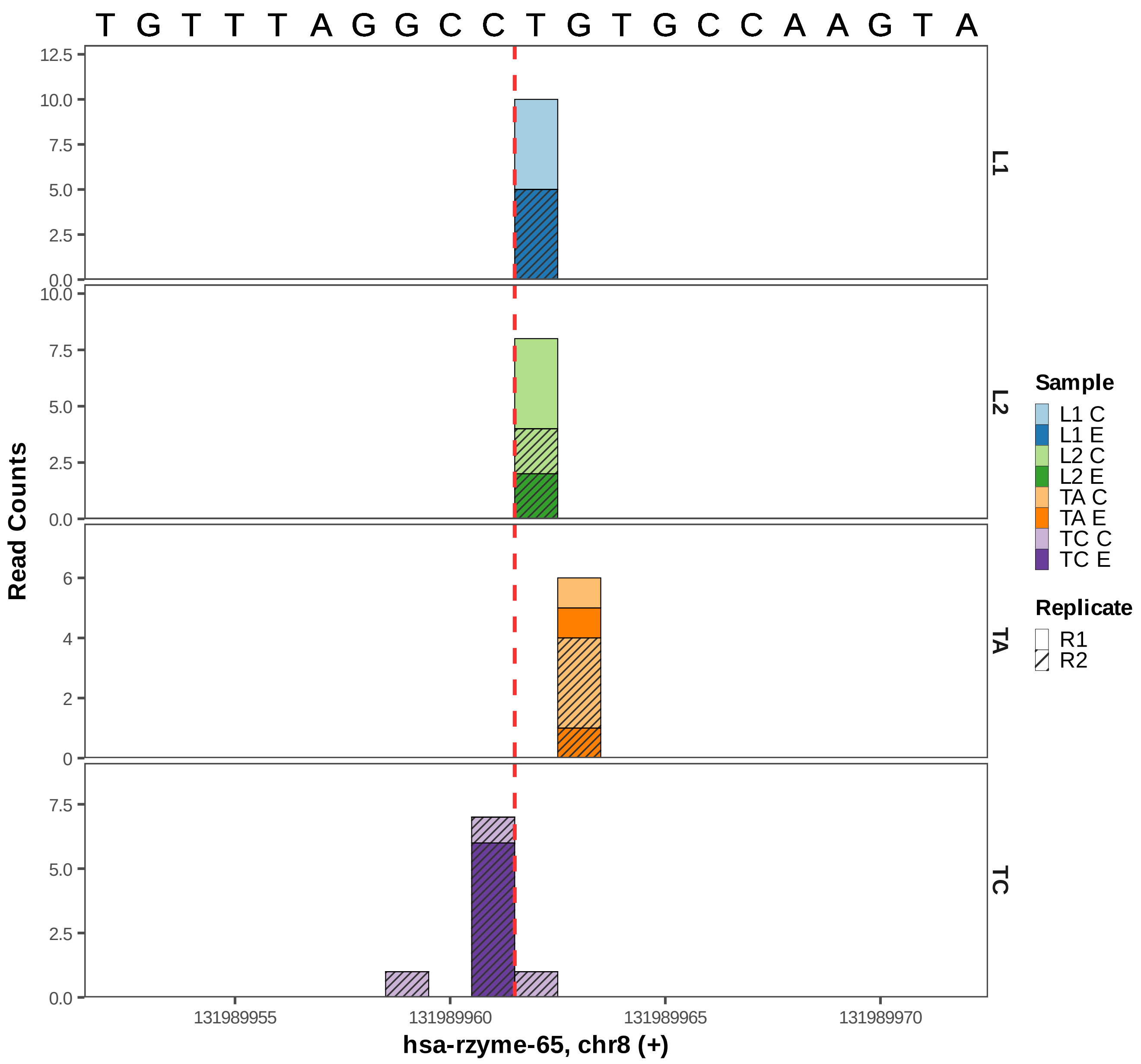
T G T T T A G G C C T G T G C C A A G T A
Read Counts
L1
L2
TA
TC
12.5
10.0
7.5
5.0
2.5
0.0
6
4
2
0
131989955
131989960
131989965
131989970
hsa-rzyme-65, chr8 (+)
Sample
L1 C
L1 E
L2 C
L2 E
TA C
TA E
TC C
TC E
Replicate
R1
R2

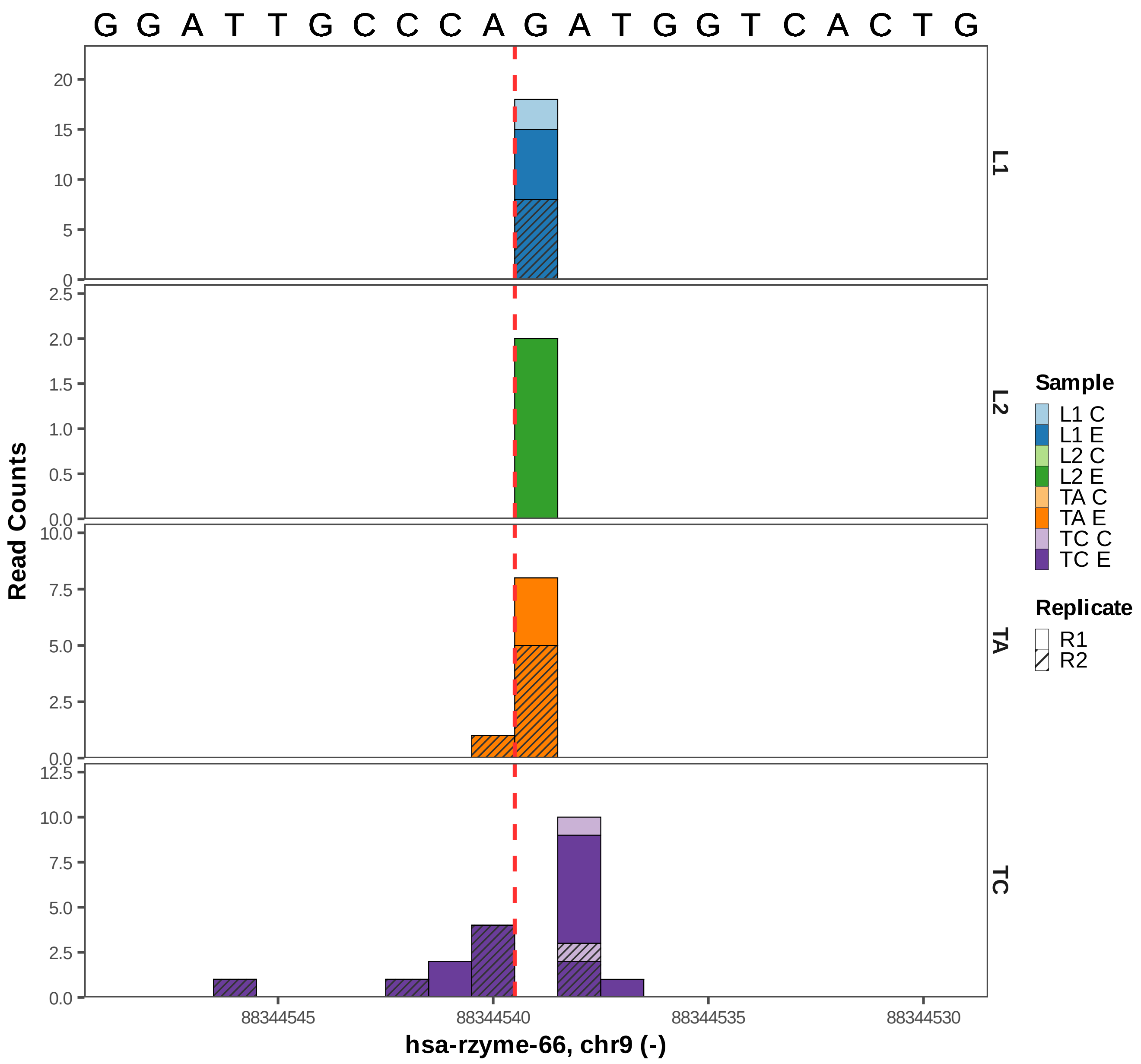

G G A T T G C C C A G A T G G T C A C T G
L1
L2
TA
TC
Read Counts
20
15
10
5
0
2.5
2.0
1.5
1.0
0.5
0.0
10.0
7.5
5.0
2.5
0.0
12.5
10.0
7.5
5.0
2.5
0.0
88344545
88344540
88344535
88344530
hsa-rzyme-66, chr9 (-)
Sample
L1 C
L1 E
L2 C
L2 E
TA C
TA E
TC C
TC E
Replicate
R1
R2

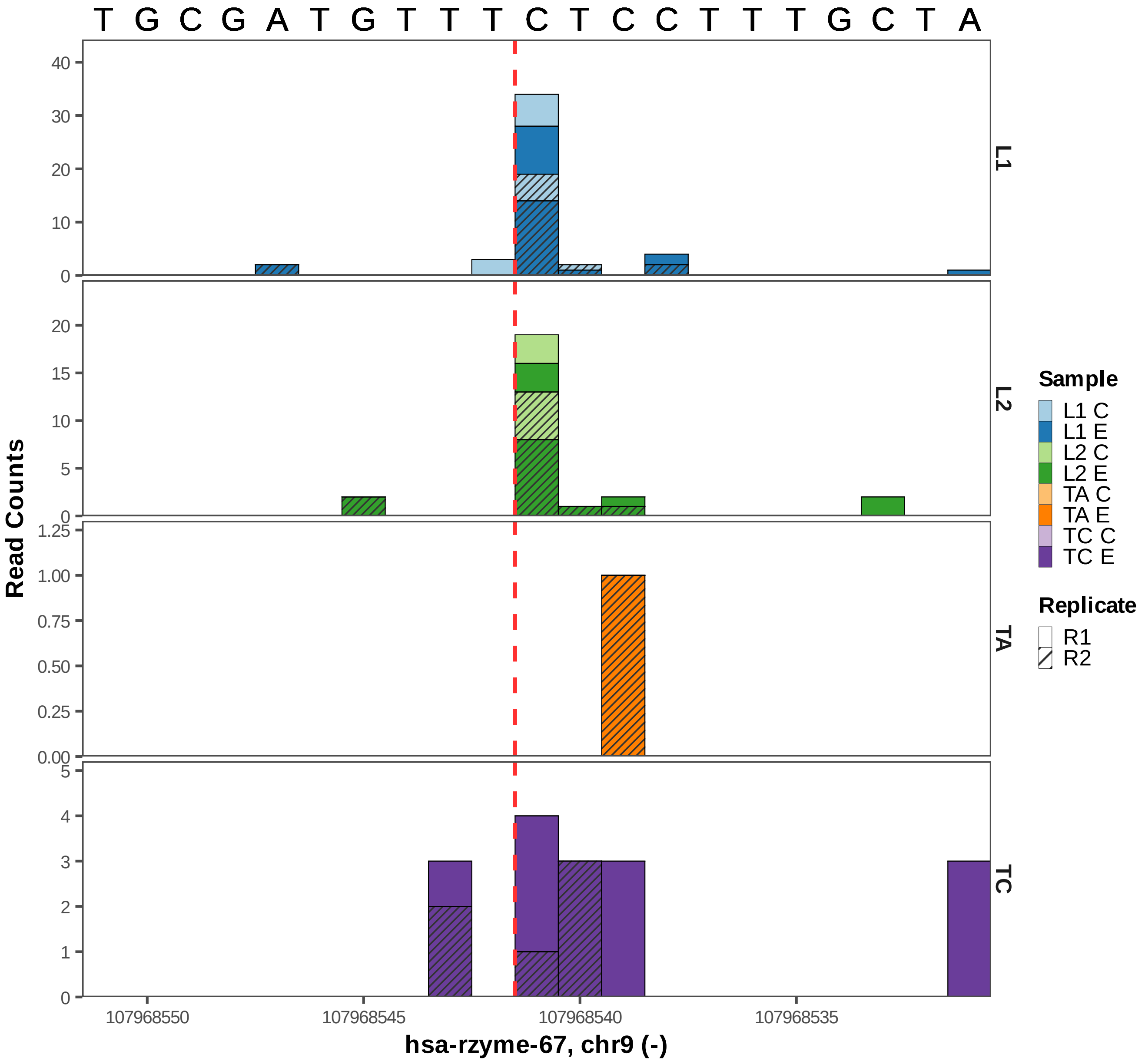

T G C G A T G T T T C T C C T T T G C T A
Read Counts
L1
L2
TA
TC
Sample
L1 C
L1 E
L2 C
L2 E
TA C
TA E
TC C
TC E
Replicate
R1
R2
107968550
107968545
107968540
107968535
hsa-rzyme-67, chr9 (-)

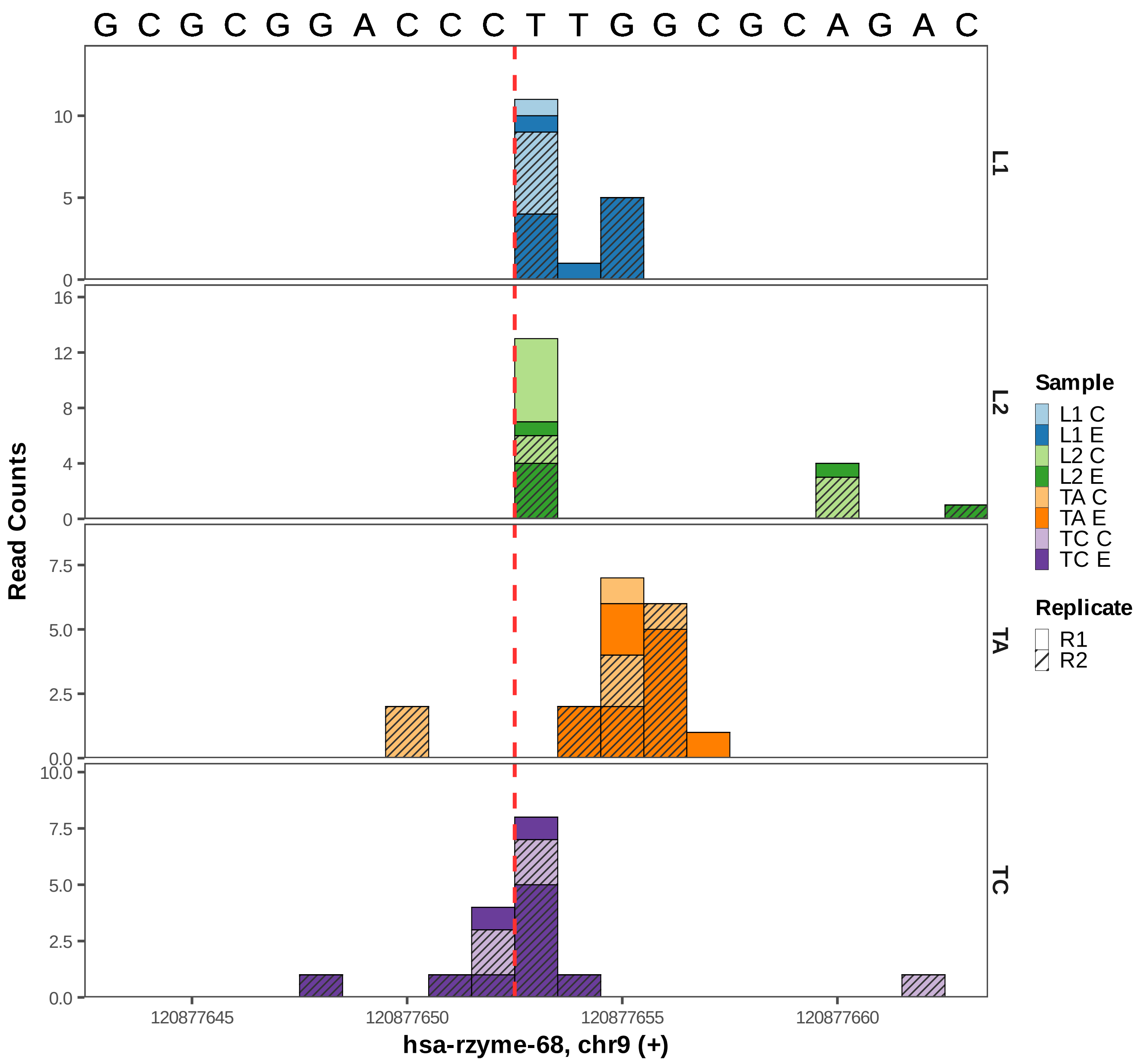
G C G C G G A C C C T T G G C G C A G A C
Read Counts
L1
L2
TA
TC
10
5
0
16
12
8
4
0
7.5
5.0
2.5
0.0
10.0
7.5
5.0
2.5
0.0
120877645
120877650
120877655
120877660
hsa-rzyme-68, chr9 (+)
Sample
L1 C
L1 E
L2 C
L2 E
TA C
TA E
TC C
TC E
Replicate
R1
R2

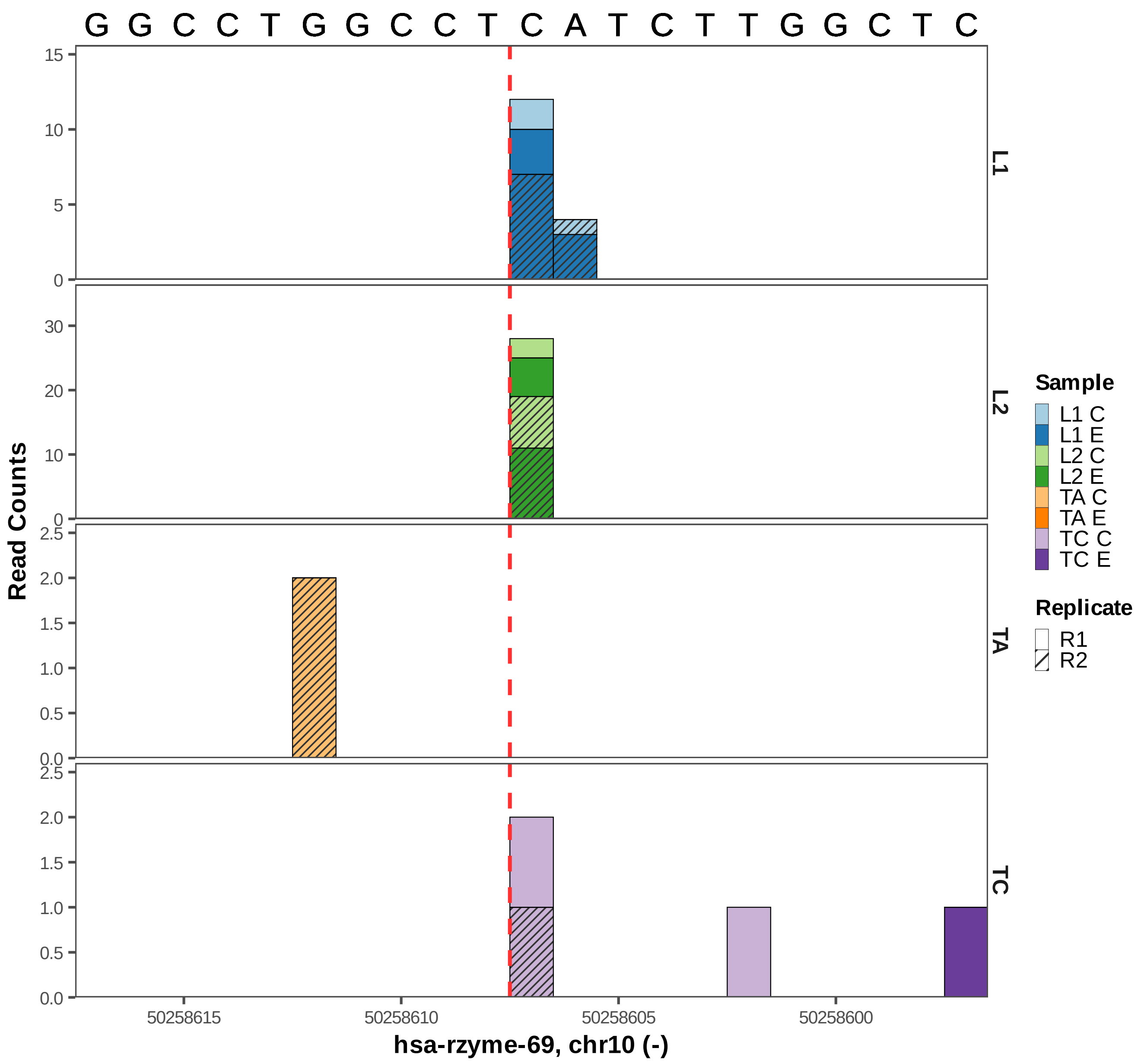
G G C C T G G C C T C A T C T T G G C T C
Read Counts
L1
L2
TA
TC
hsa-rzyme-69, chr10 (-)
50258615
50258610
50258605
50258600
Sample
L1 C
L1 E
L2 C
L2 E
TA C
TA E
TC C
TC E
Replicate
R1
R2

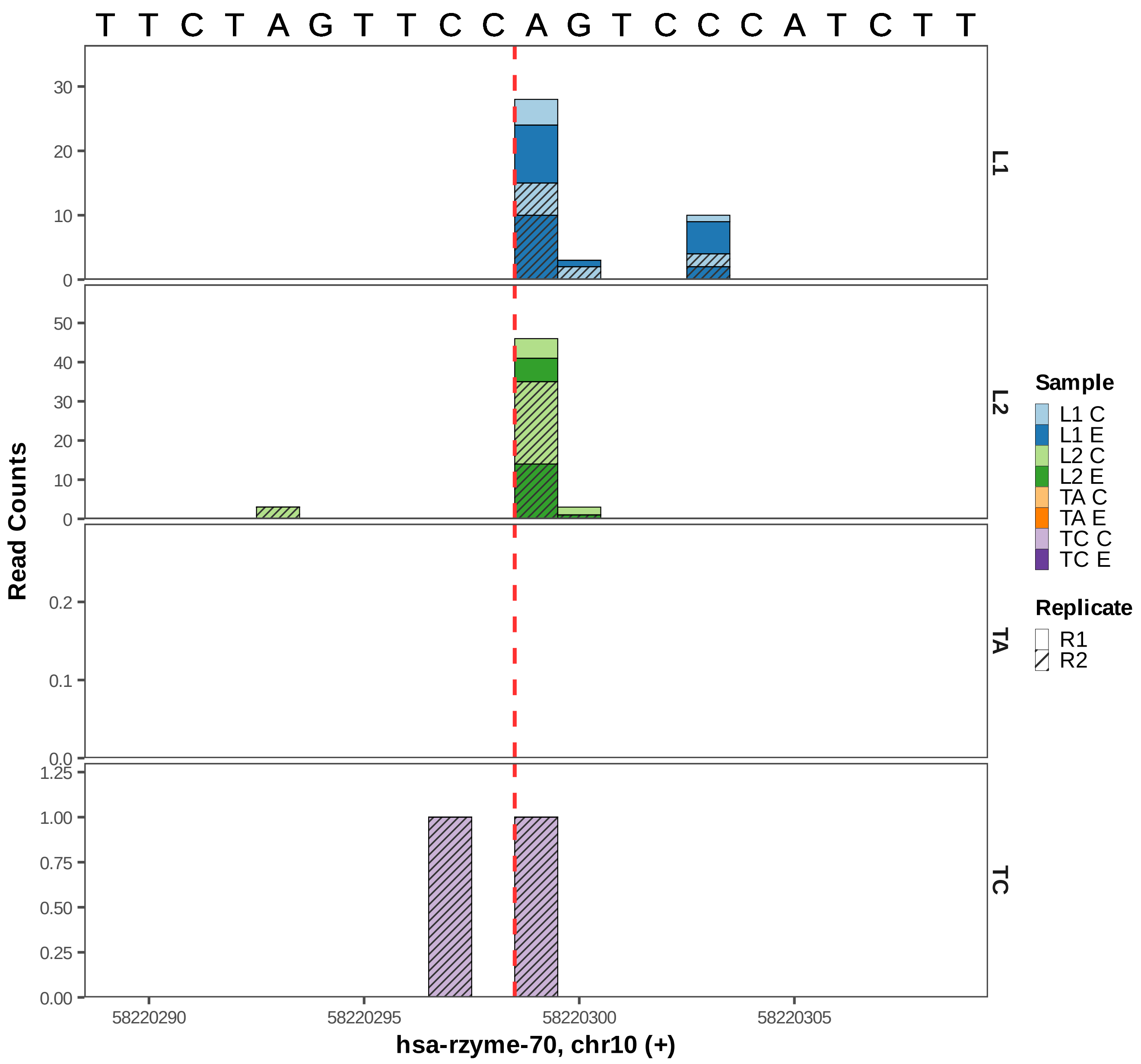

T T C T A G T T C C A G T C C C A T C T T
Read Counts
L1
L2
TA
TC
30
20
10
0
50
40
0.2
0.1
0.0
1.25
1.00
0.75
0.50
0.25
0.00
58220290
58220295
58220300
58220305
hsa-rzyme-70, chr10 (+)
Sample
L1 C
L1 E
L2 C
L2 E
TA C
TA E
TC C
TC E
Replicate
R1
R2

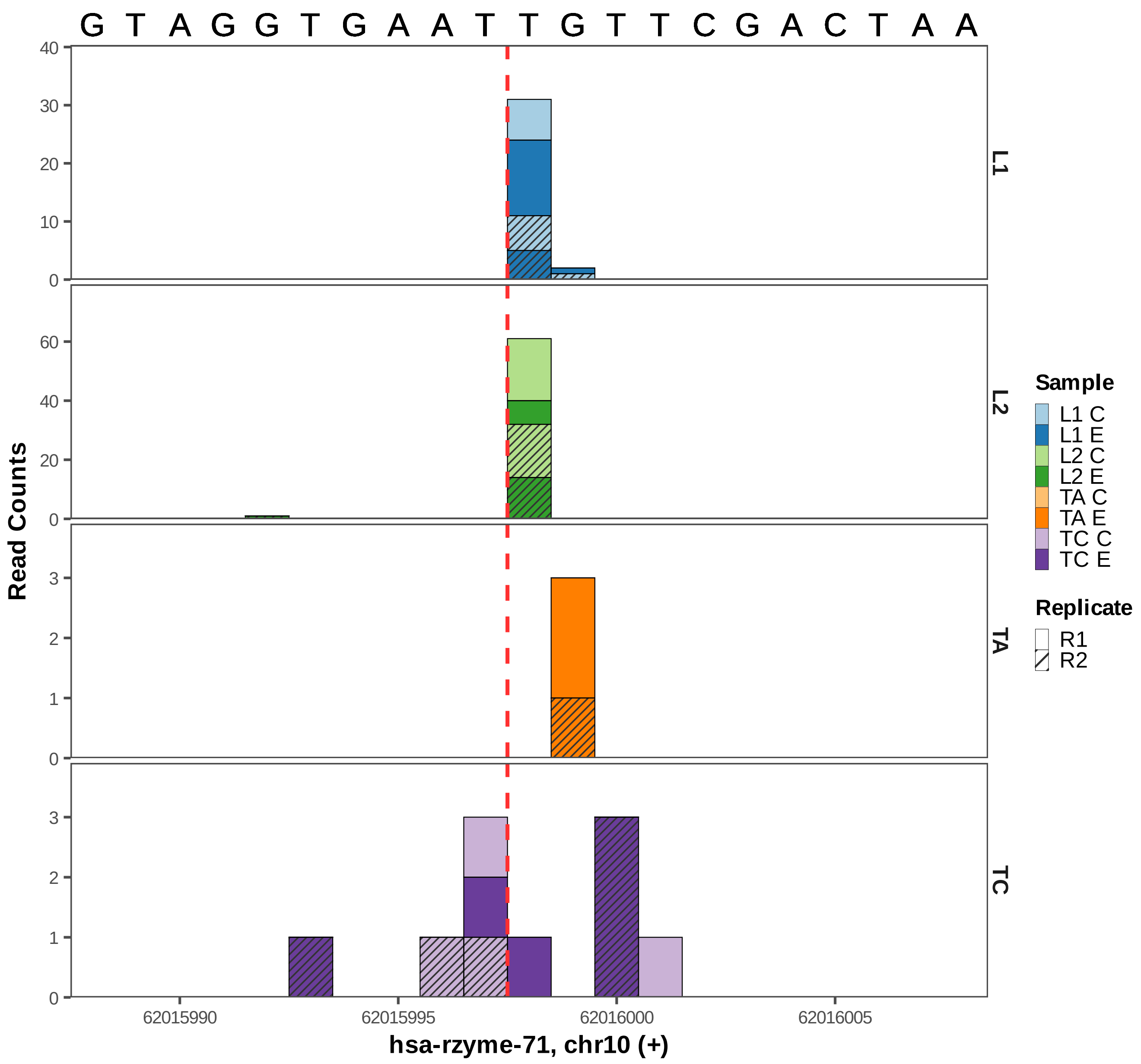
G T A G G T G A A T T G T T C G A C T A A
Read Counts
L1
L2
TA
TC
Sample
L1 C
L1 E
L2 C
L2 E
TA C
TA E
TC C
TC E
Replicate
R1
R2
62015990
62015995
62016000
62016005
hsa-rzyme-71, chr10 (+)

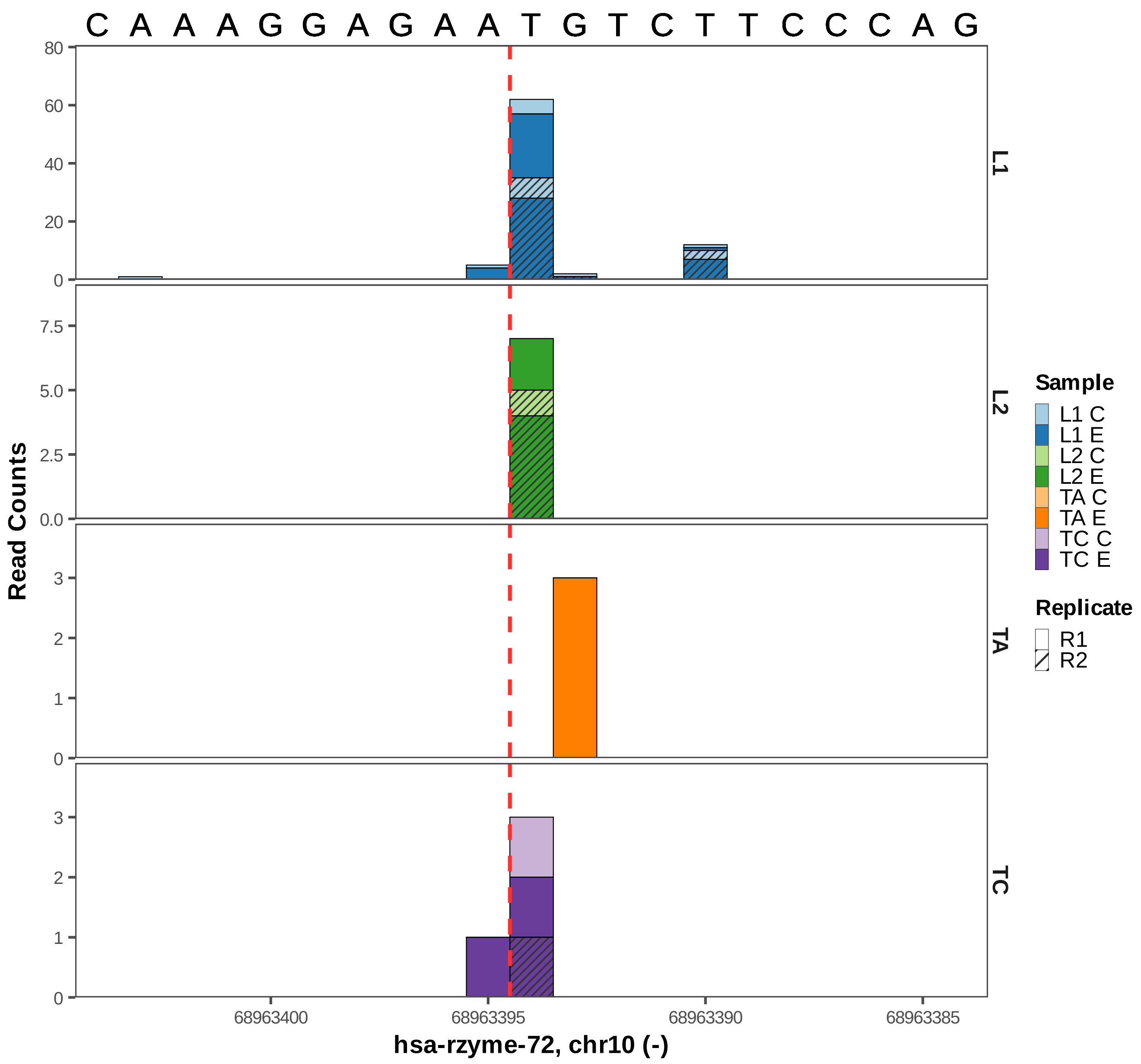

C A A A G G A G A A T G T C T T C C C A G
Read Counts
L1
L2
TA
TC
Sample
L1 C
L1 E
L2 C
L2 E
TA C
TA E
TC C
TC E
Replicate
R1
R2
68963400
68963395
68963390
68963385
hsa-rzyme-72, chr10 (-)

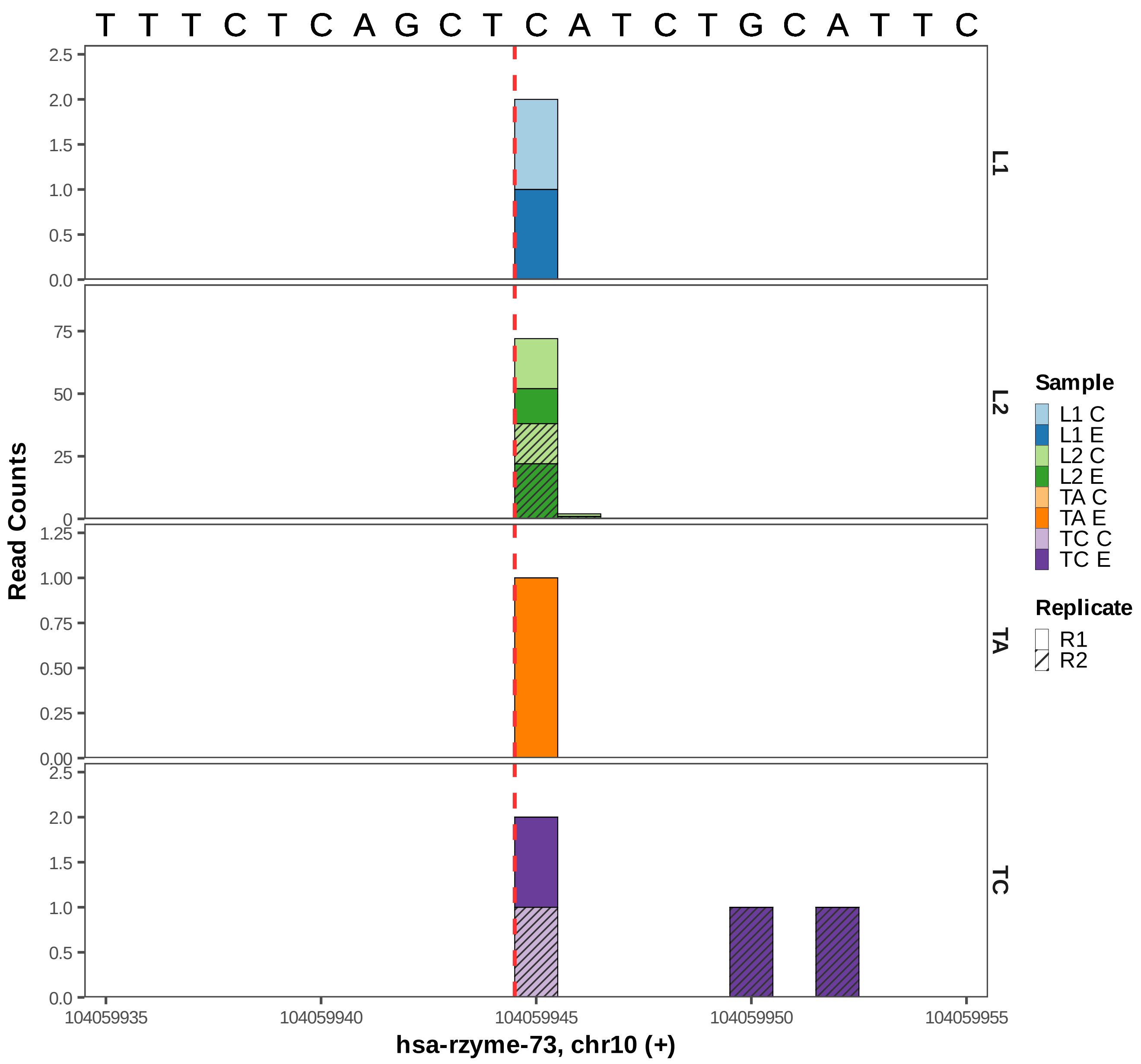

T T T C T C A G C T C A T C T G C A T T C
Read Counts
L1
L2
TA
TC
2.5
2.0
1.5
1.0
0.5
0.0
75
50
25
0
1.25
1.00
0.75
0.50
0.25
0.00
104059935
104059940
104059945
104059950
104059955
hsa-rzyme-73, chr10 (+)
Sample
L1 C
L1 E
L2 C
L2 E
TA C
TA E
TC C
TC E
Replicate
R1
R2

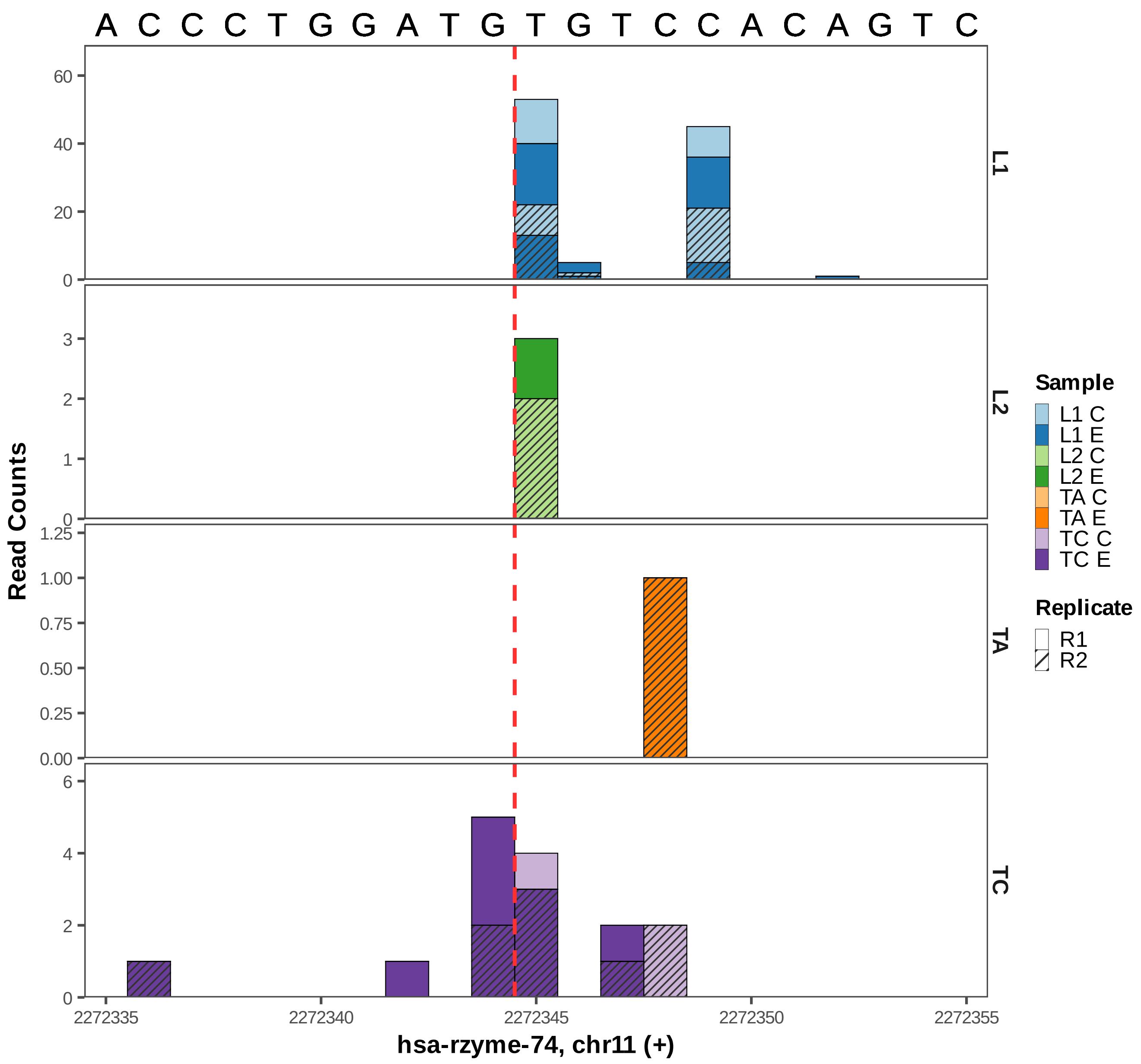

A C C C T G G A T G T G T C C A C A G T C
L1
L2
TA
TC
Read Counts
60
40
20
0
3
2
1
0
1.25
1.00
0.75
0.50
0.25
0.00
6
4
2
0
2272335
2272340
2272345
2272350
2272355
hsa-rzyme-74, chr11 (+)
Sample
L1 C
L1 E
L2 C
L2 E
TA C
TA E
TC C
TC E
Replicate
R1
R2

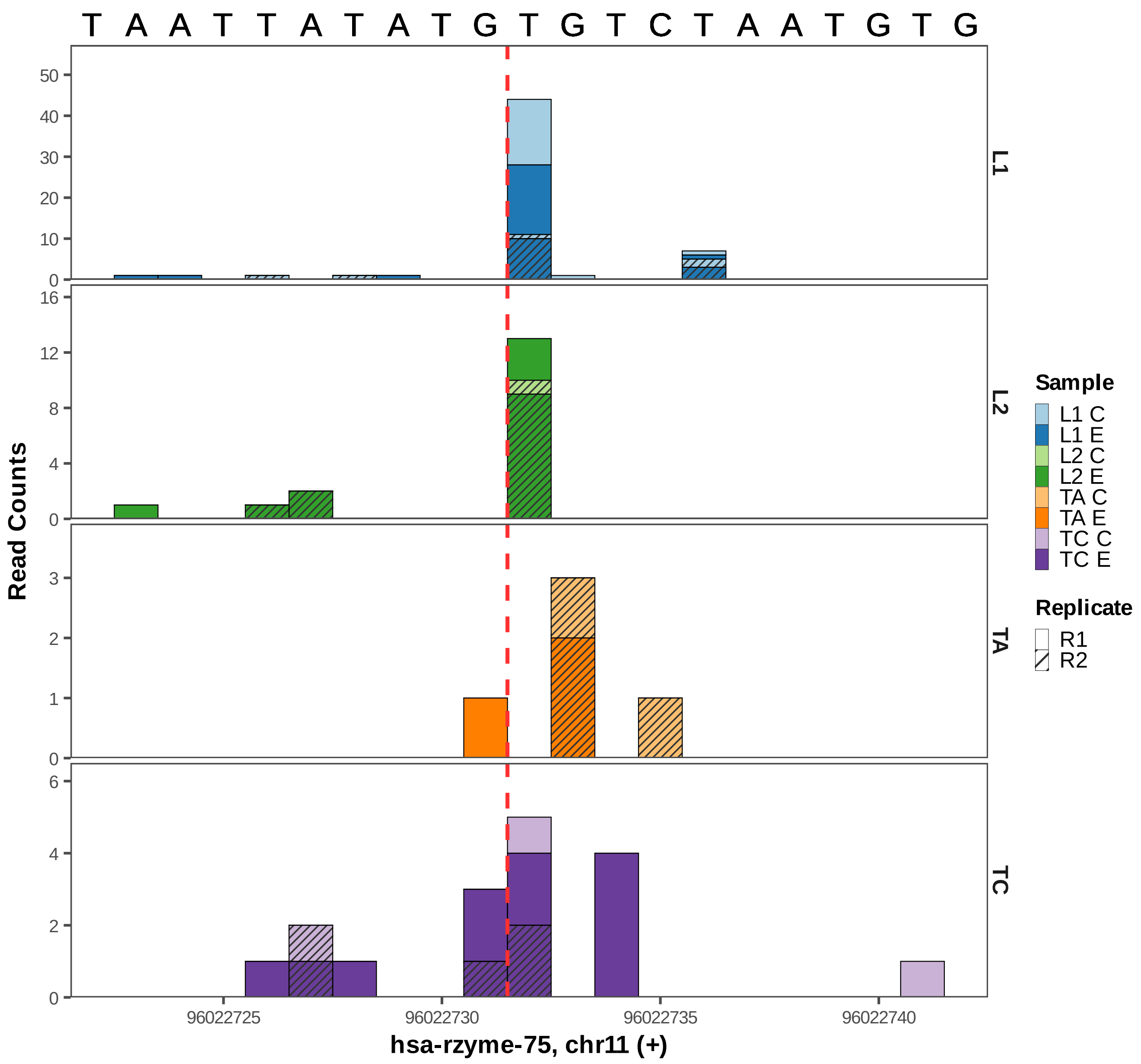
T A A T T A T A T G T G T C T A A T G T G
50
40
30
20
10
0
16
12
8
4
0
3
2
1
0
6
4
2
0
L1
L2
TA
TC
Read Counts
Sample
L1 C
L1 E
L2 C
L2 E
TA C
TA E
TC C
TC E
Replicate
R1
R2
96022725
96022730
96022735
96022740
hsa-rzyme-75, chr11 (+)

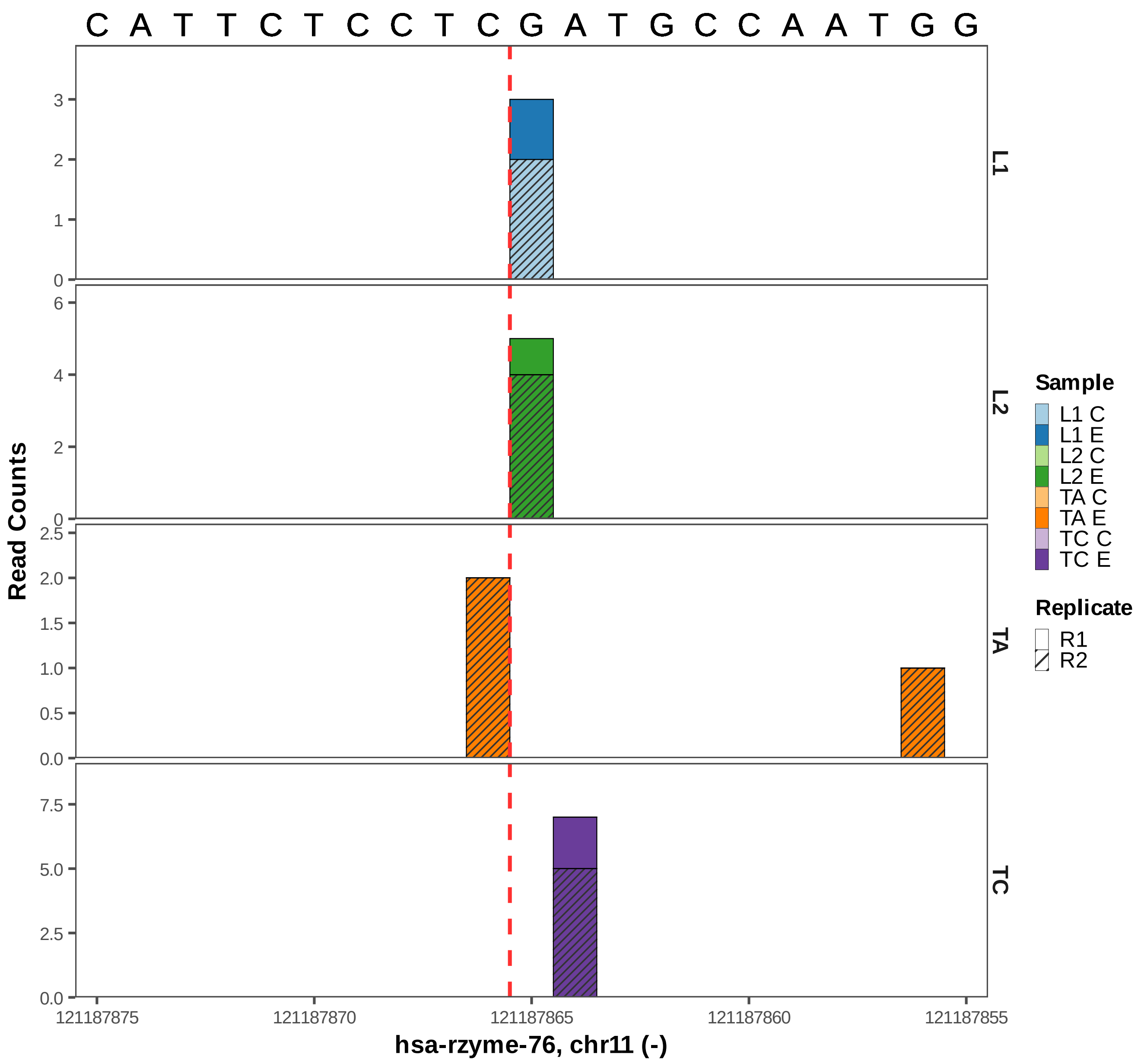

C A T T C T C C T C G A T G C C A A T G G
L1
L2
TA
TC
Read Counts
3
2
1
0
6
4
2
0
2.5
2.0
1.5
1.0
0.5
0.0
7.5
5.0
2.5
0.0
121187875
121187870
121187865
121187860
121187855
hsa-rzyme-76, chr11 (-)
Sample
L1 C
L1 E
L2 C
L2 E
TA C
TA E
TC C
TC E
Replicate
R1
R2

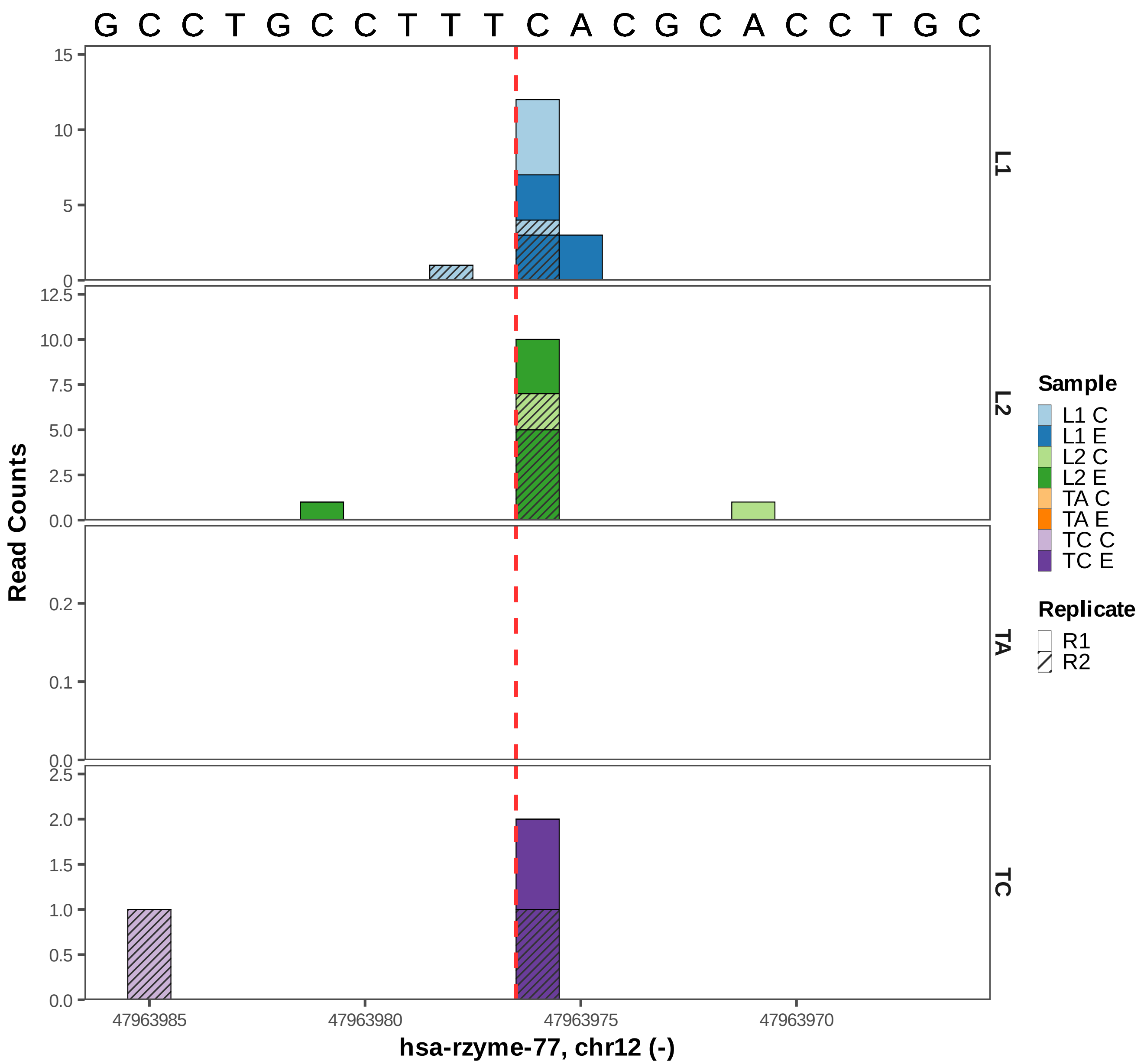

G C C T G C C T T T C A C G C A C C T G C
L1
L2
TA
TC
Read Counts
hsa-rzyme-77, chr12 (-)
47963985
47963980
47963975
47963970
Sample
L1 C
L1 E
L2 C
L2 E
TA C
TA E
TC C
TC E
Replicate
R1
R2

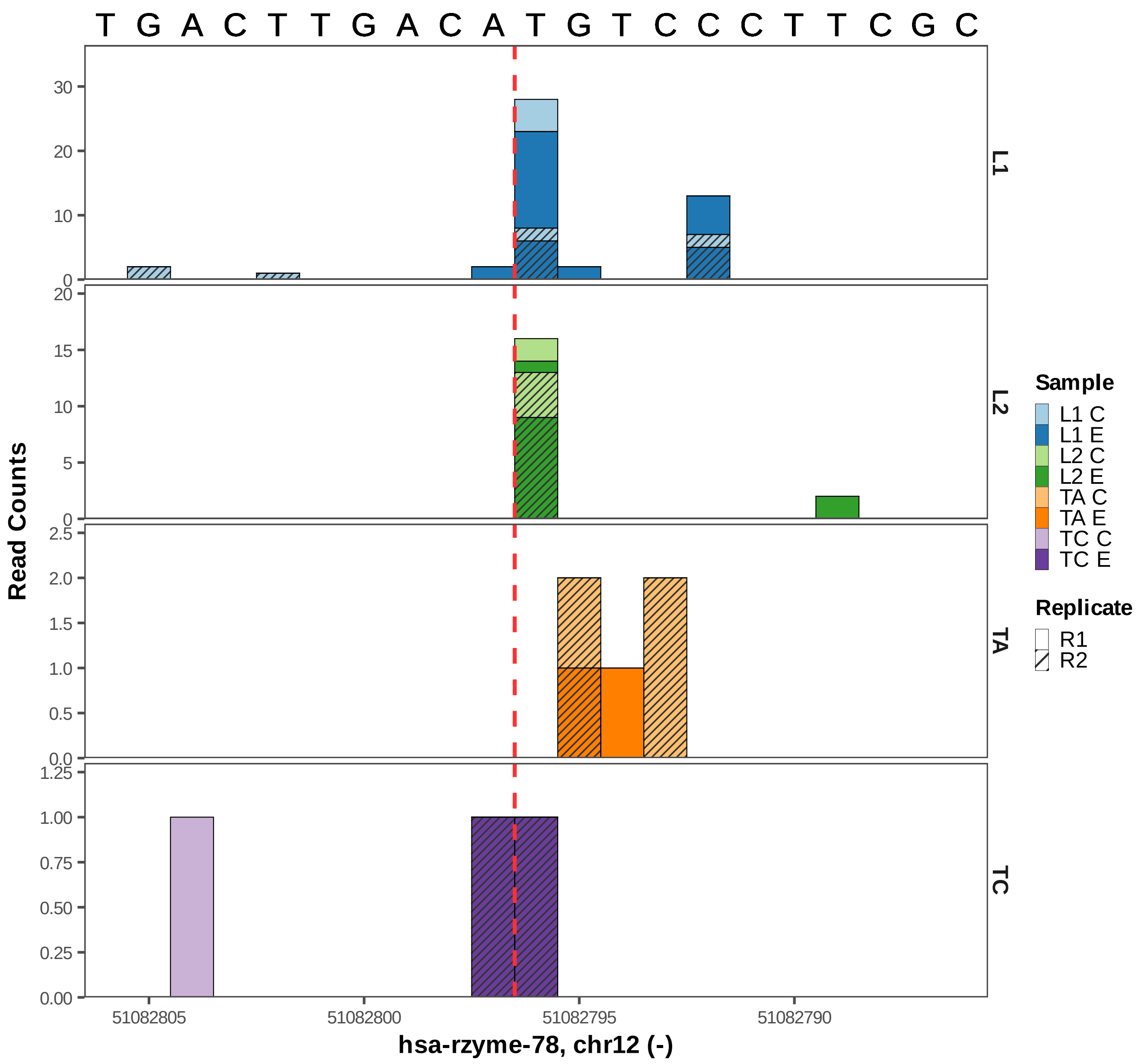

T G A C T T G A C A T G T C C C T T C G C
L1
L2
TA
TC
Read Counts
Sample
L1 C
L1 E
L2 C
L2 E
TA C
TA E
TC C
TC E
Replicate
R1
R2
51082805
51082800
51082795
51082790
hsa-rzyme-78, chr12 (-)

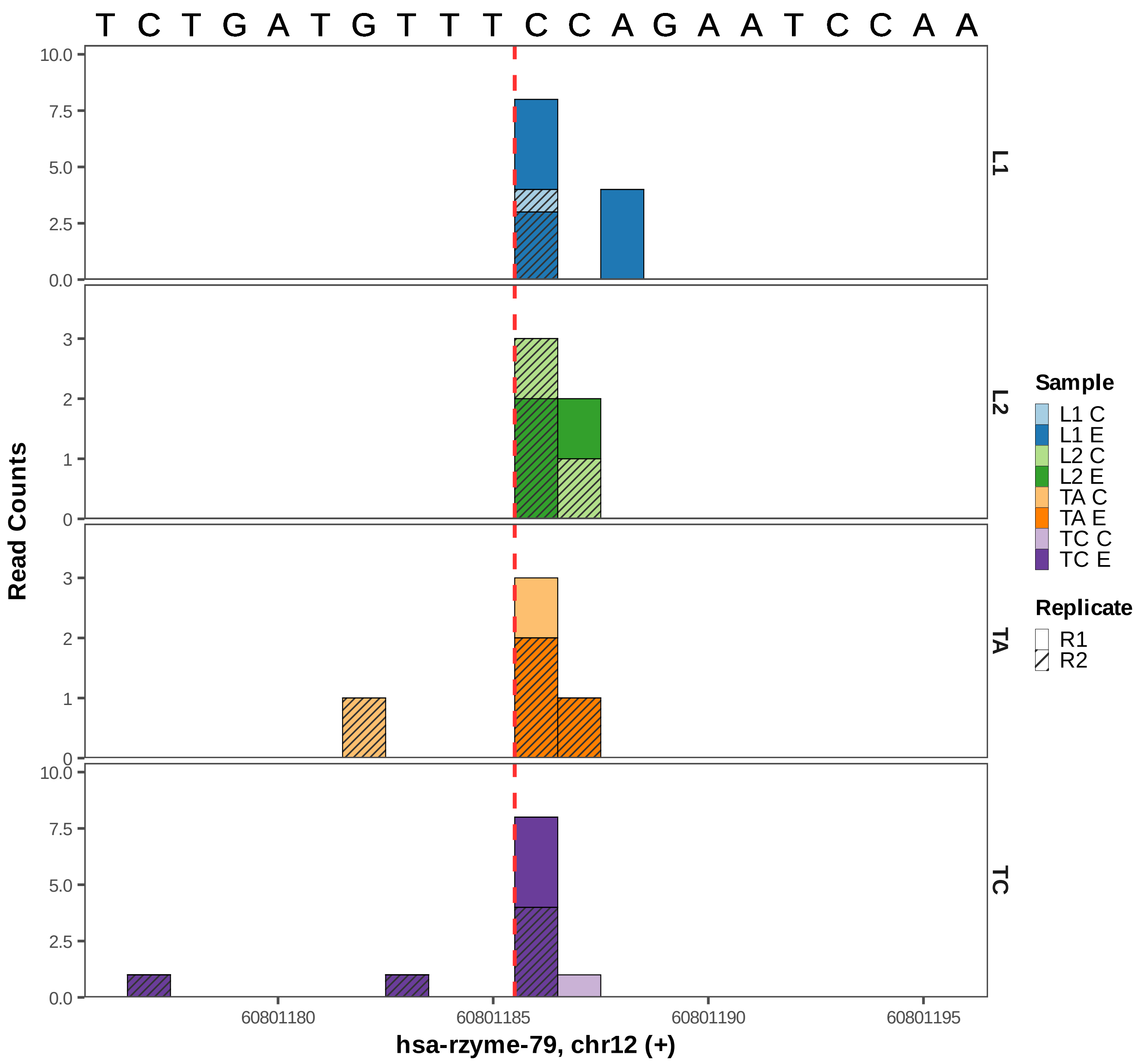

T C T G A T G T T T C C A G A A T C C A A
Read Counts
L1
L2
TA
TC
hsa-rzyme-79, chr12 (+)
60801180
60801185
60801190
60801195
Sample
L1 C
L1 E
L2 C
L2 E
TA C
TA E
TC C
TC E
Replicate
R1
R2

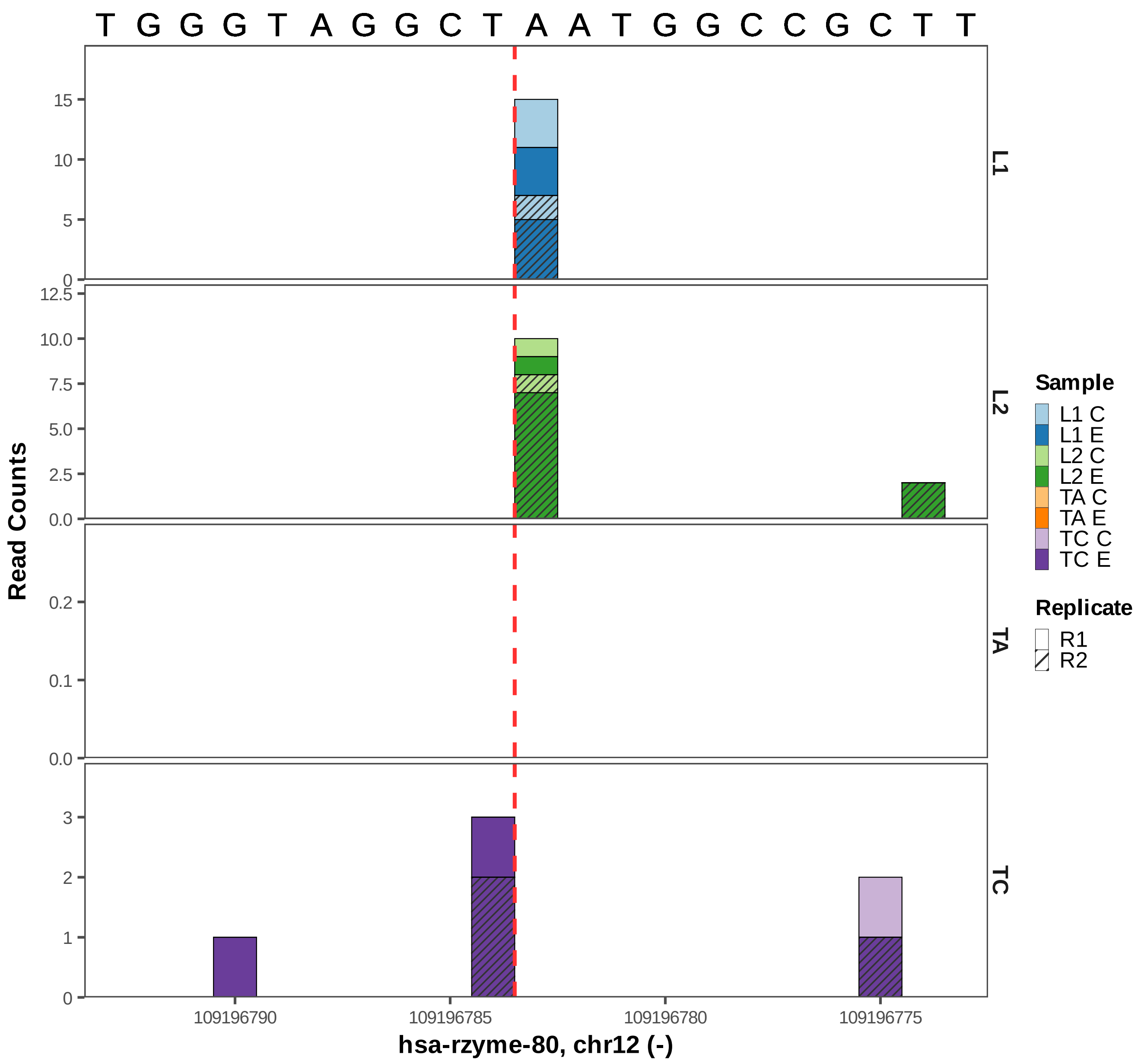
T G G G T A G G C T A A T G G C C G C T T
L1
L2
TA
TC
Read Counts
hsa-rzyme-80, chr12 (-)
109196790
109196785
109196780
109196775
Sample
L1 C
L1 E
L2 C
L2 E
TA C
TA E
TC C
TC E
Replicate
R1
R2

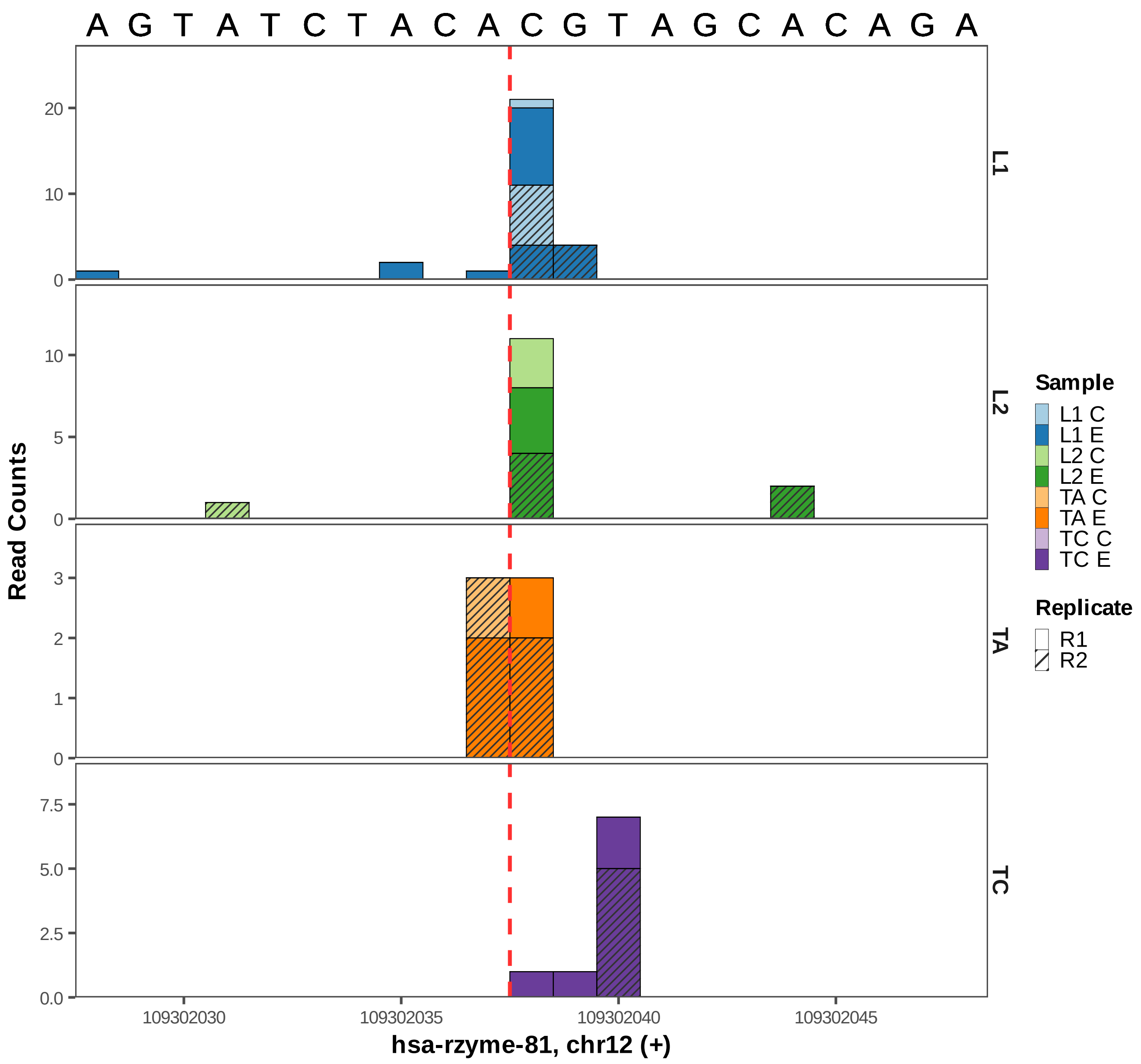

A G T A T C T A C A C G T A G C A C A G A
Read Counts
L1
L2
TA
TC
20
10
0
5
3
2
1
7.5
5.0
2.5
0.0
109302030
109302035
109302040
109302045
hsa-rzyme-81, chr12 (+)
Sample
L1 C
L1 E
L2 C
L2 E
TA C
TA E
TC C
TC E
Replicate
R1
R2

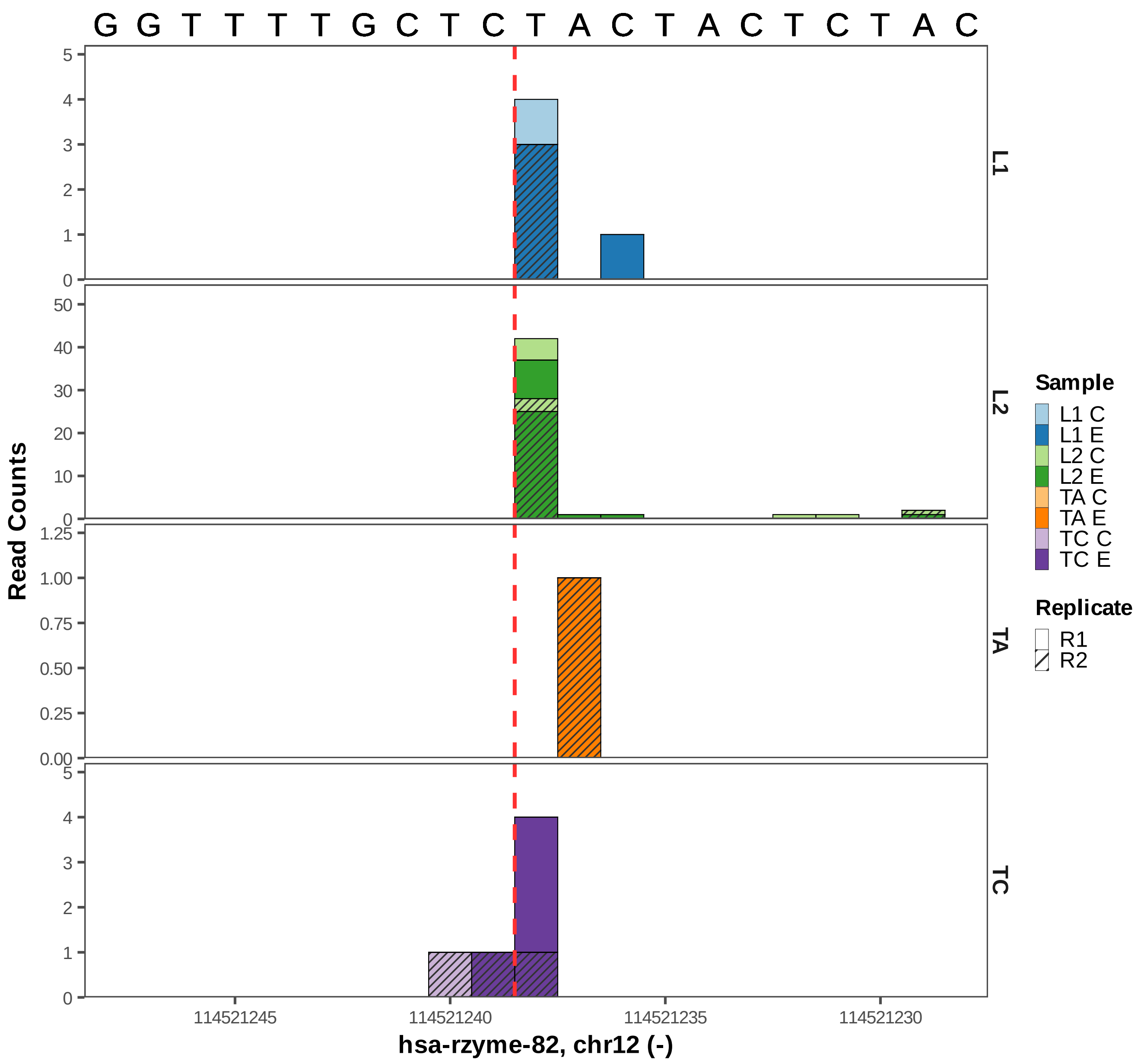
G G T T T T G C T C T A C T A C T C T A C
L1
L2
TA
TC
Read Counts
114521245
114521240
114521235
114521230
hsa-rzyme-82, chr12 (-)
Sample
L1 C
L1 E
L2 C
L2 E
TA C
TA E
TC C
TC E
Replicate
R1
R2

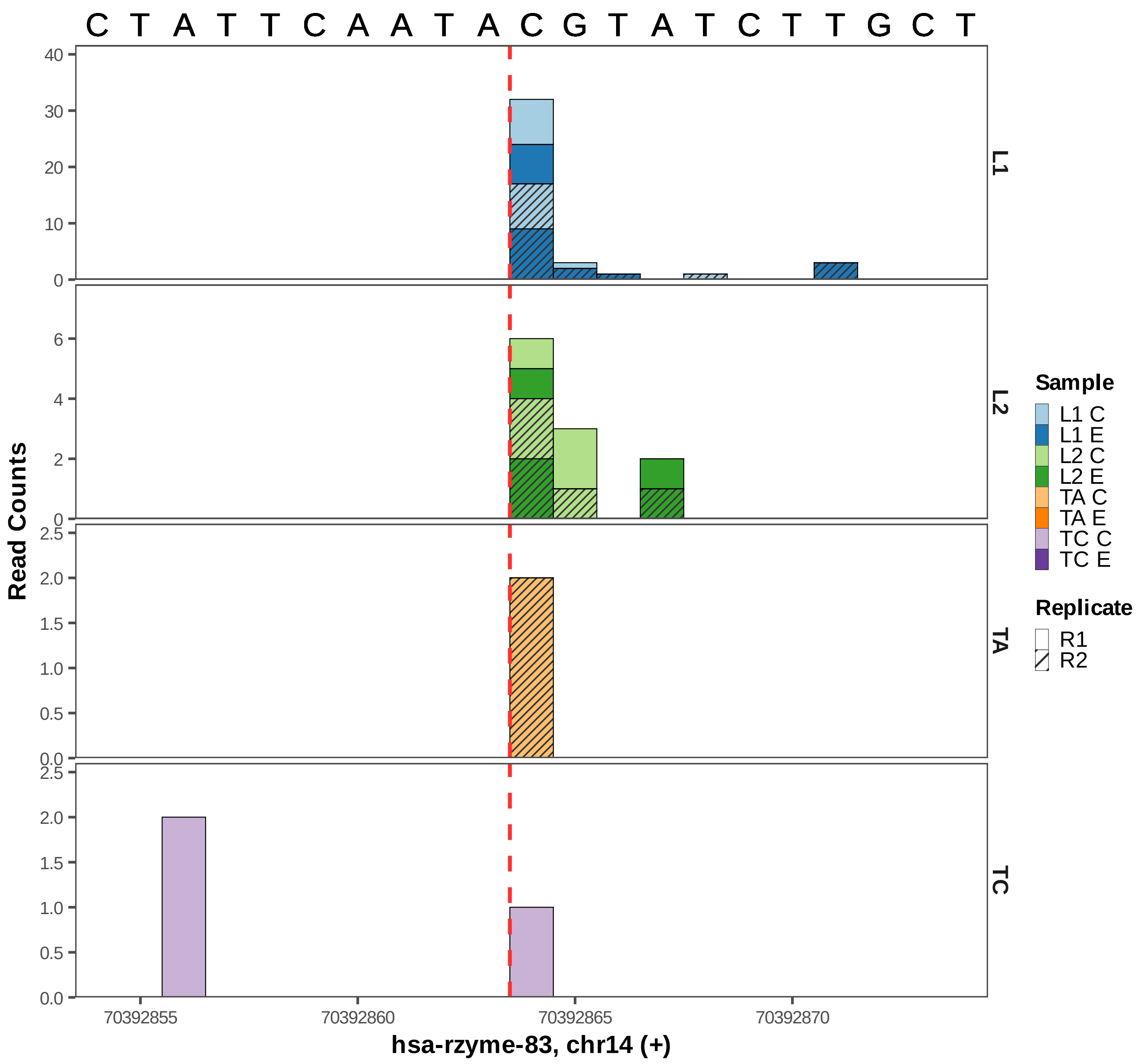

C T A T T C A A T A C G T A T C T T G C T
Read Counts
L1
L2
TA
TC
Sample
L1 C
L1 E
L2 C
L2 E
TA C
TA E
TC C
TC E
Replicate
R1
R2
70392855
70392860
70392865
70392870
hsa-rzyme-83, chr14 (+)

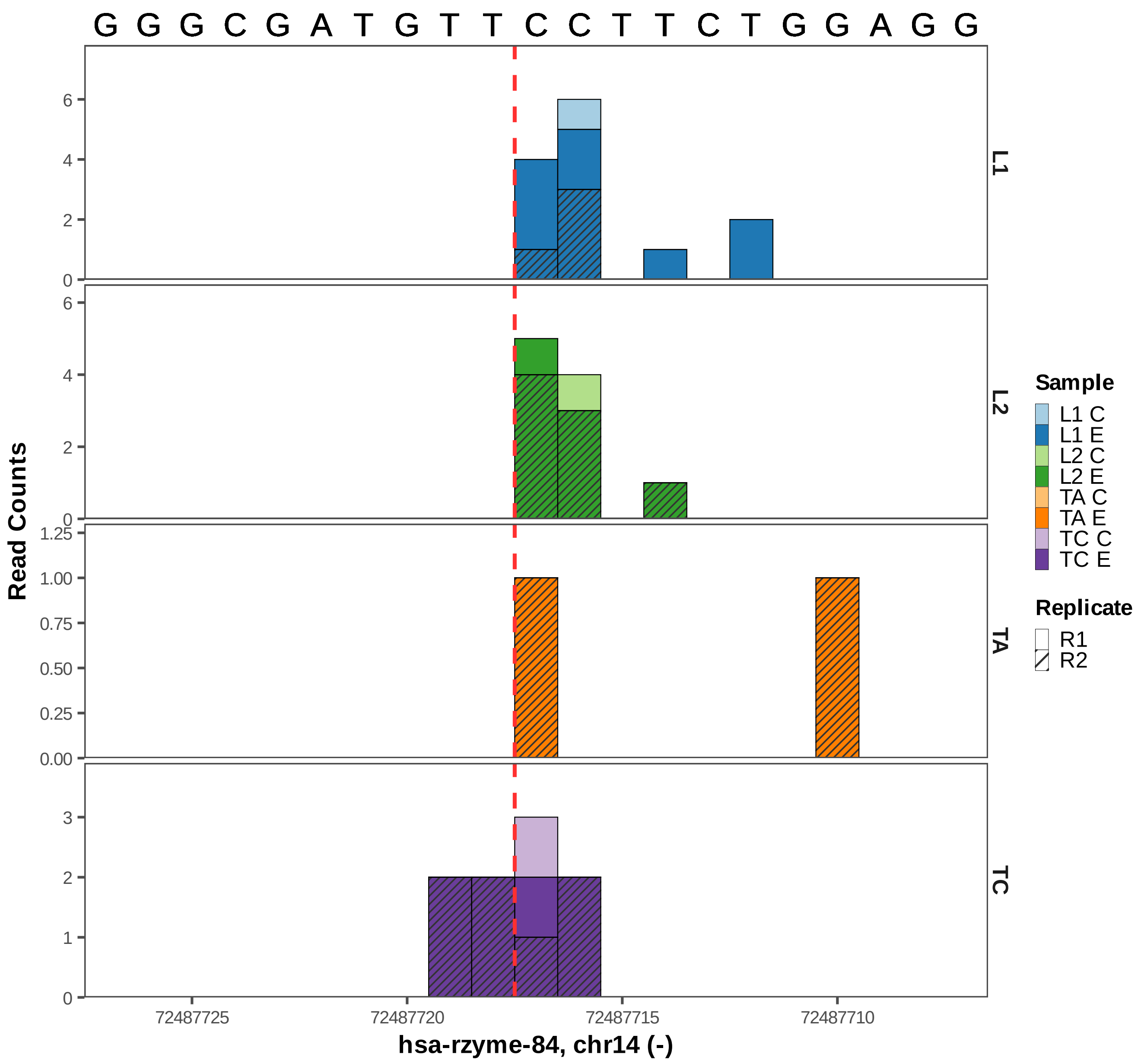

G G G C G A T G T T C C T T C T G G A G G
Read Counts
L1
L2
TA
TC
Sample
L1 C
L1 E
L2 C
L2 E
TA C
TA E
TC C
TC E
Replicate
R1
R2
72487725
72487720
72487715
72487710
hsa-rzyme-84, chr14 (-)

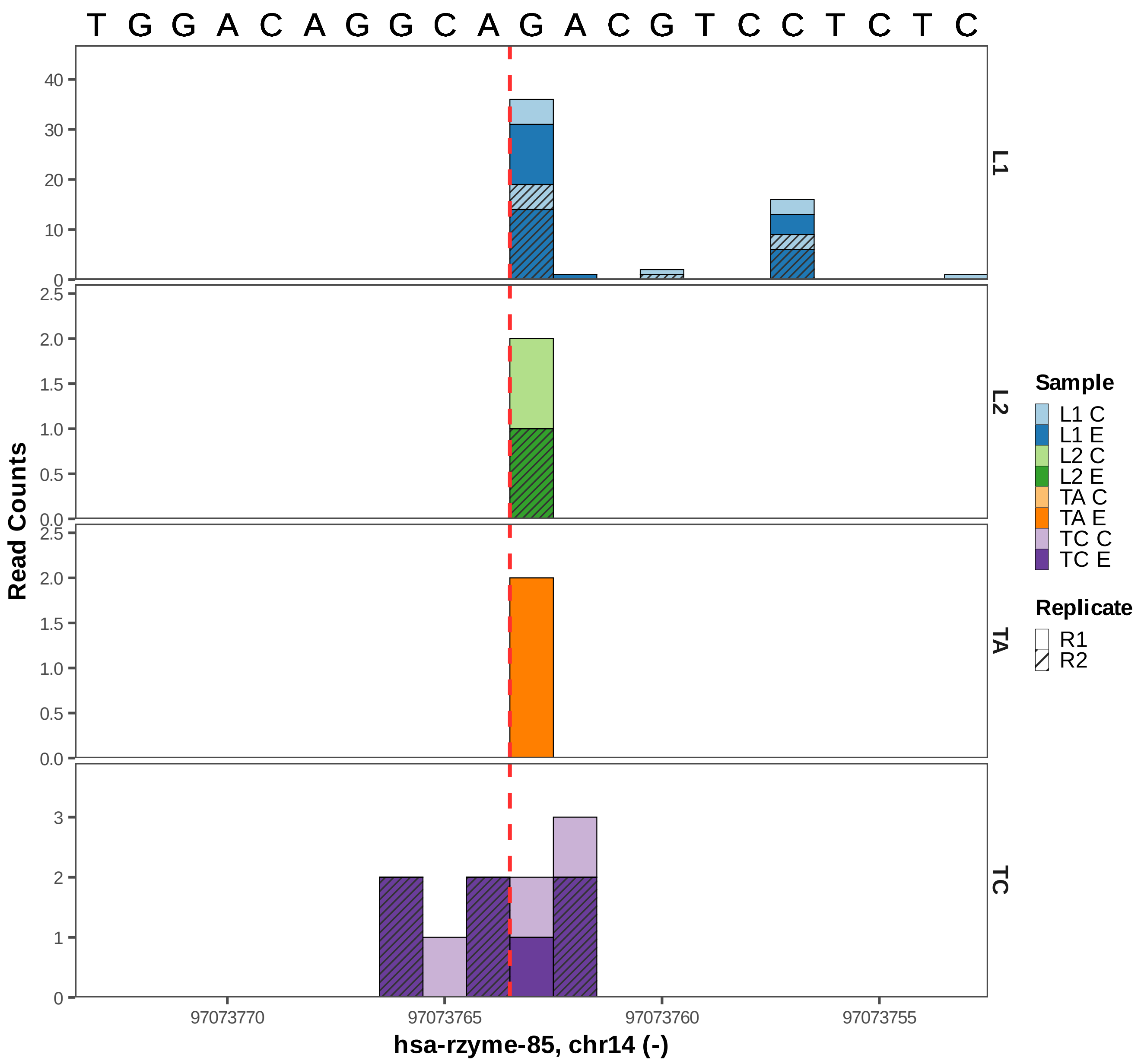

T G G A C A G G C A G A C G T C C T C T C
Read Counts
L1
L2
TA
TC
Sample
L1 C
L1 E
L2 C
L2 E
TA C
TA E
TC C
TC E
Replicate
R1
R2
97073770
97073765
97073760
97073755
hsa-rzyme-85, chr14 (-)

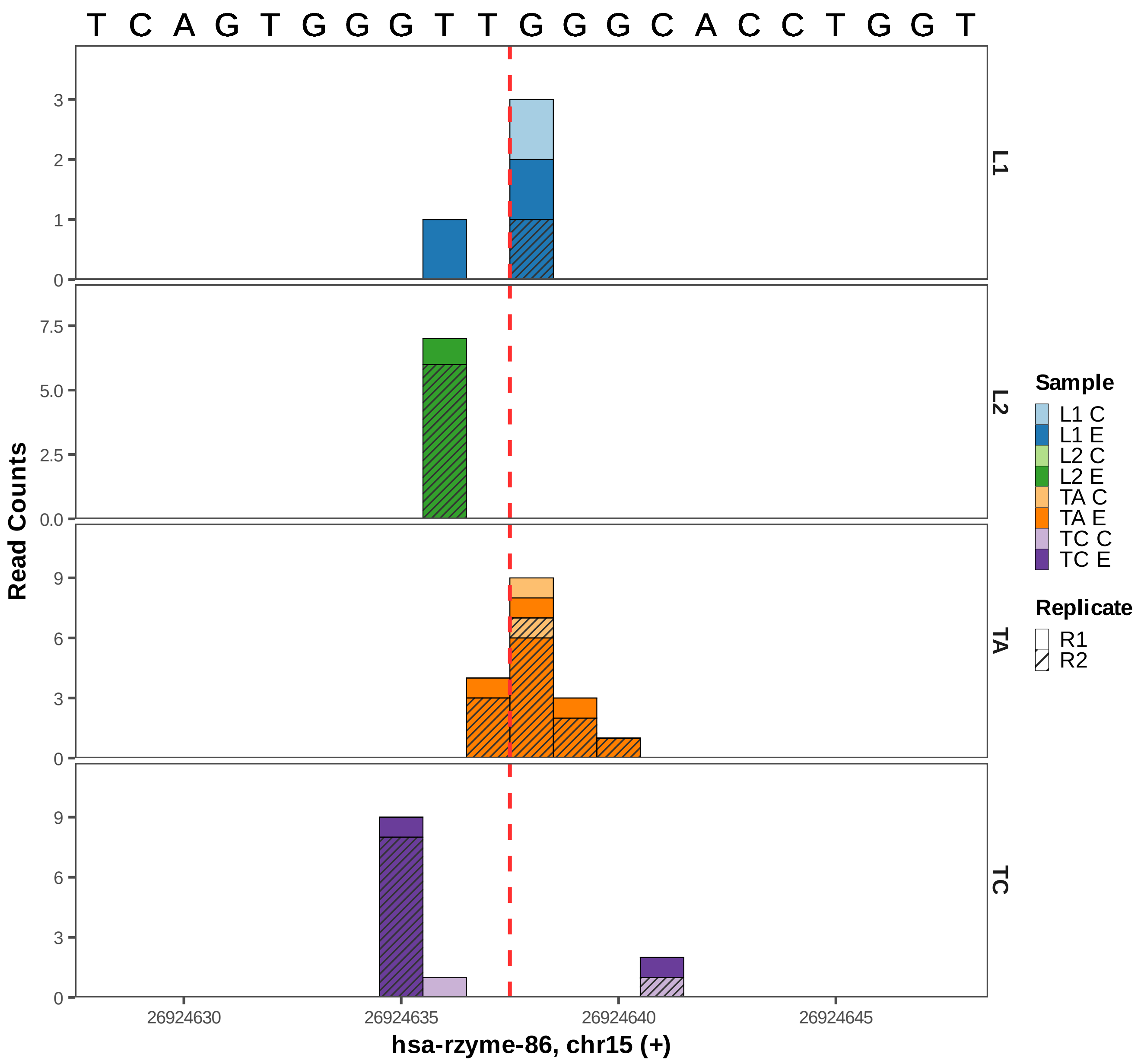
T C A G T G G G T T G G G C A C C T G G T
Read Counts
L1
L2
TA
TC
Sample
L1 C
L1 E
L2 C
L2 E
TA C
TA E
TC C
TC E
Replicate
R1
R2
26924630
26924635
26924640
26924645
hsa-rzyme-86, chr15 (+)

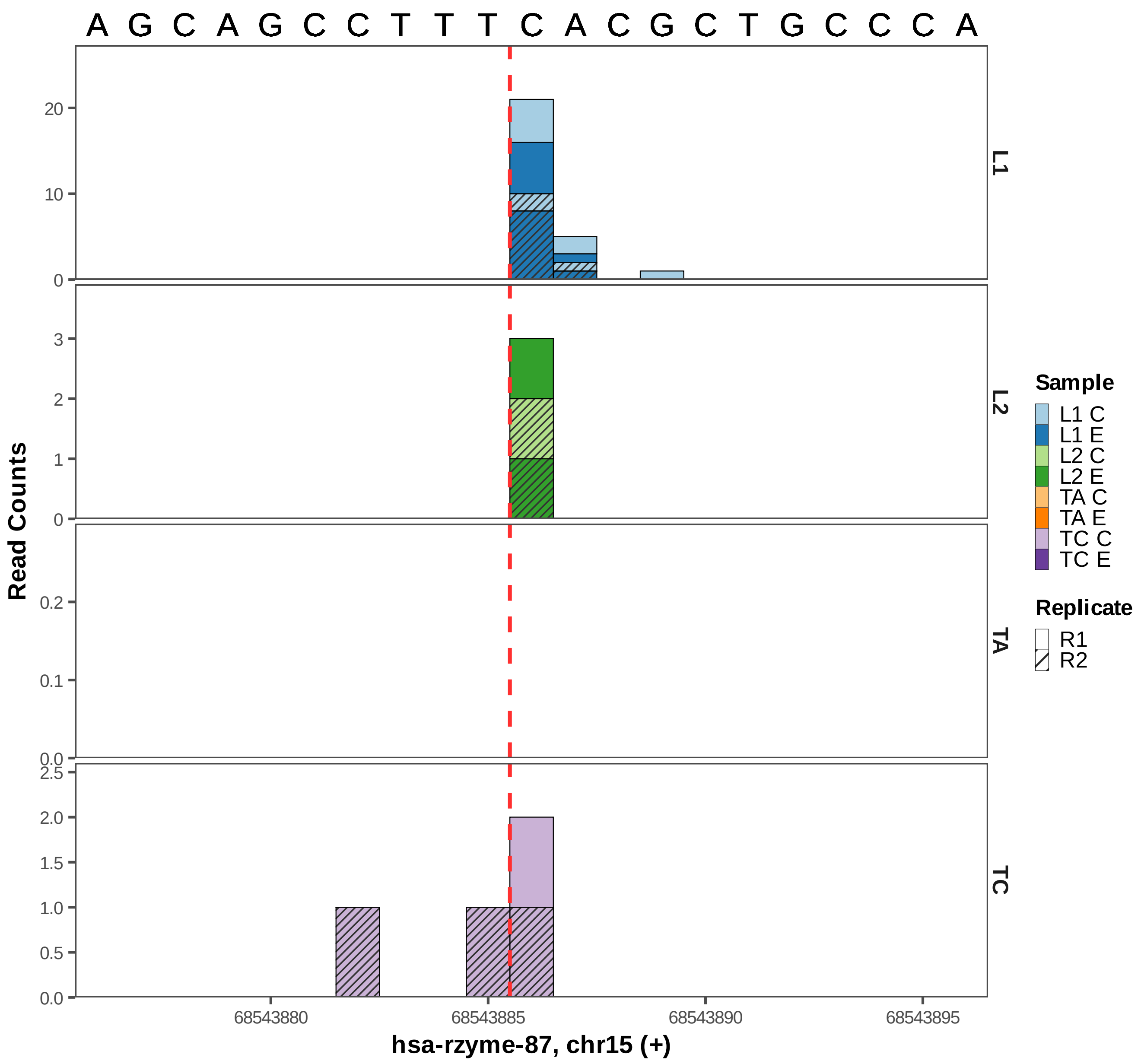

A G C A G C C T T T C A C G C T G C C C A
Read Counts
L1
L2
TA
TC
Sample
L1 C
L1 E
L2 C
L2 E
TA C
TA E
TC C
TC E
Replicate
R1
R2
68543880
68543885
68543890
68543895
hsa-rzyme-87, chr15 (+)

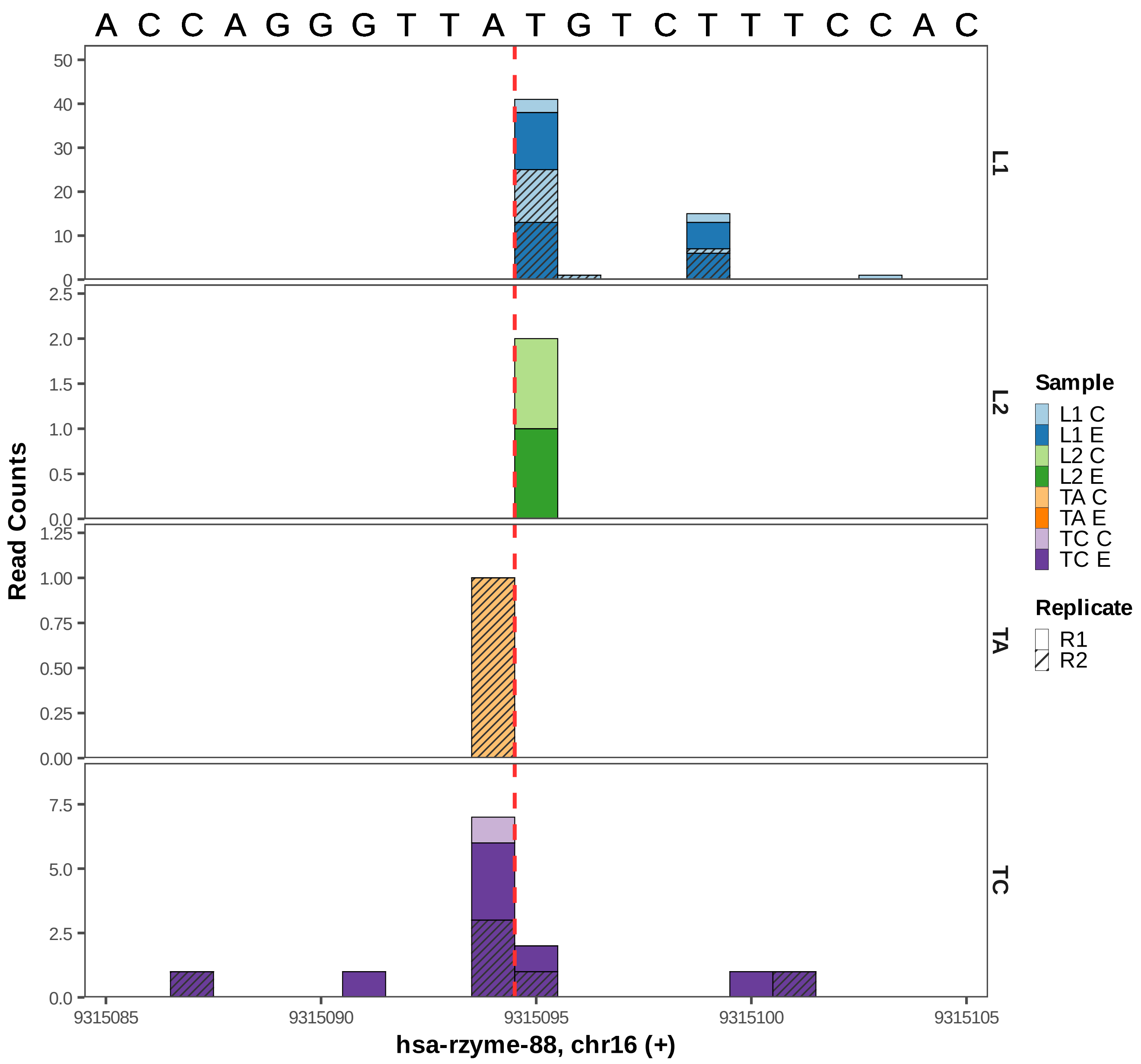

A C C A G G G T T A T G T C T T T C C A C
Read Counts
L1
L2
TA
TC
Sample
L1 C
L1 E
L2 C
L2 E
TA C
TA E
TC C
TC E
Replicate
R1
R2
9315085
9315090
9315095
9315100
9315105
hsa-rzyme-88, chr16 (+)

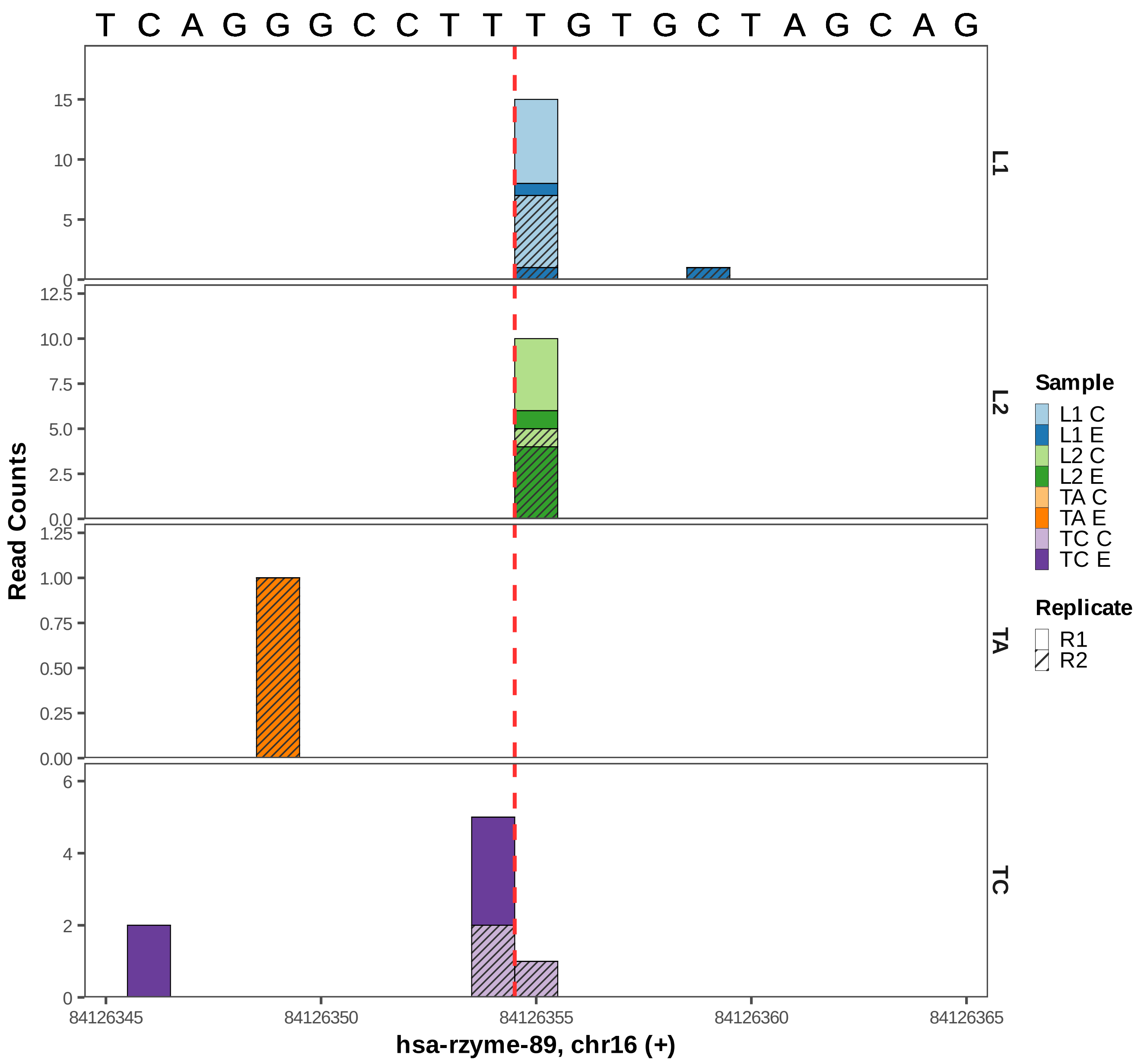

T C A G G G C C T T T G T G C T A G C A G
Read Counts
L1
L2
TA
TC
hsa-rzyme-89, chr16 (+)
84126345
84126350
84126355
84126360
84126365
Sample
L1 C
L1 E
L2 C
L2 E
TA C
TA E
TC C
TC E
Replicate
R1
R2

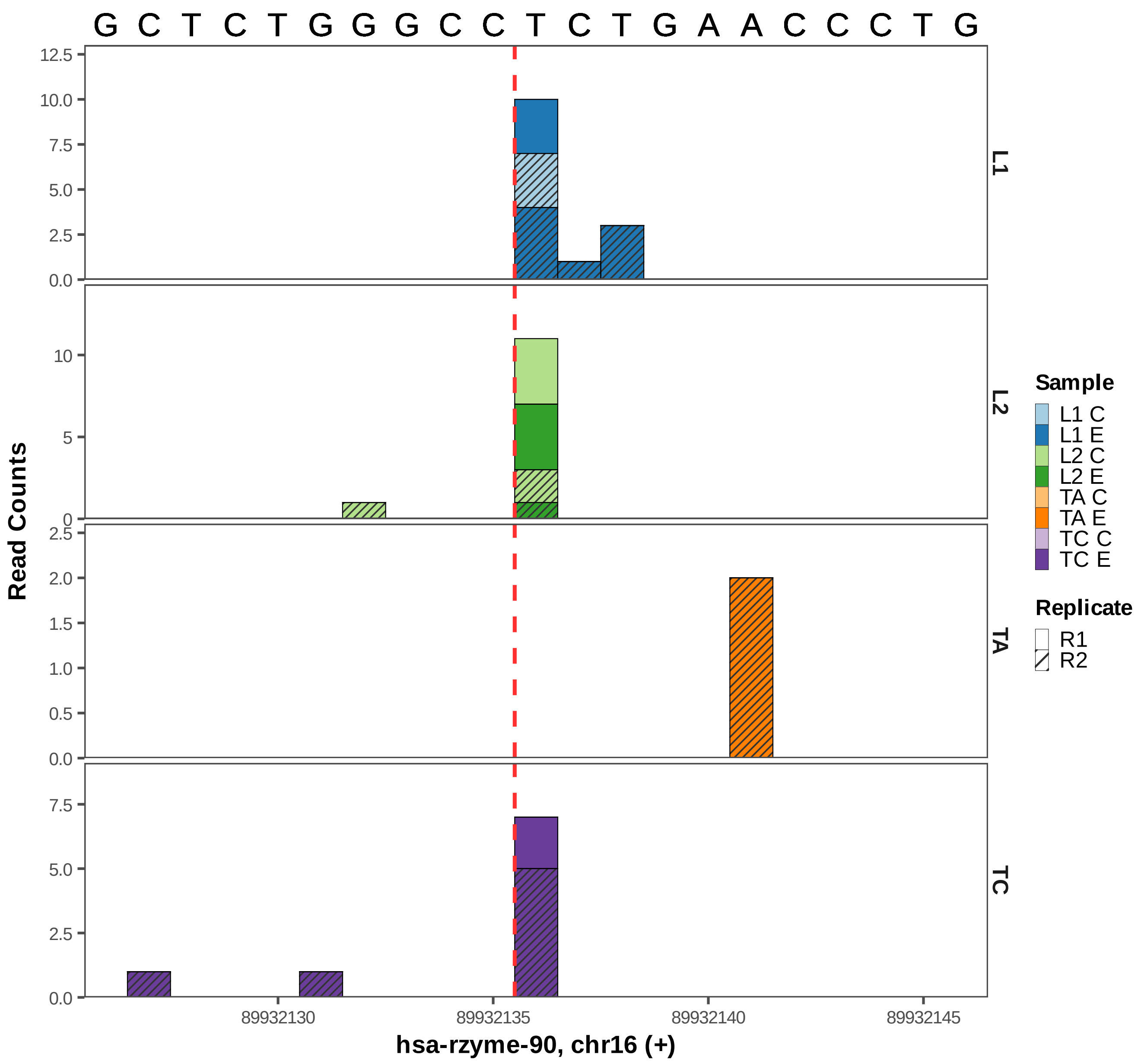

G C T C T G G G C C T C T G A A C C C T G
L1
L2
TA
TC
Read Counts
hsa-rzyme-90, chr16 (+)
89932130
89932135
89932140
89932145
Sample
L1 C
L1 E
L2 C
L2 E
TA C
TA E
TC C
TC E
Replicate
R1
R2

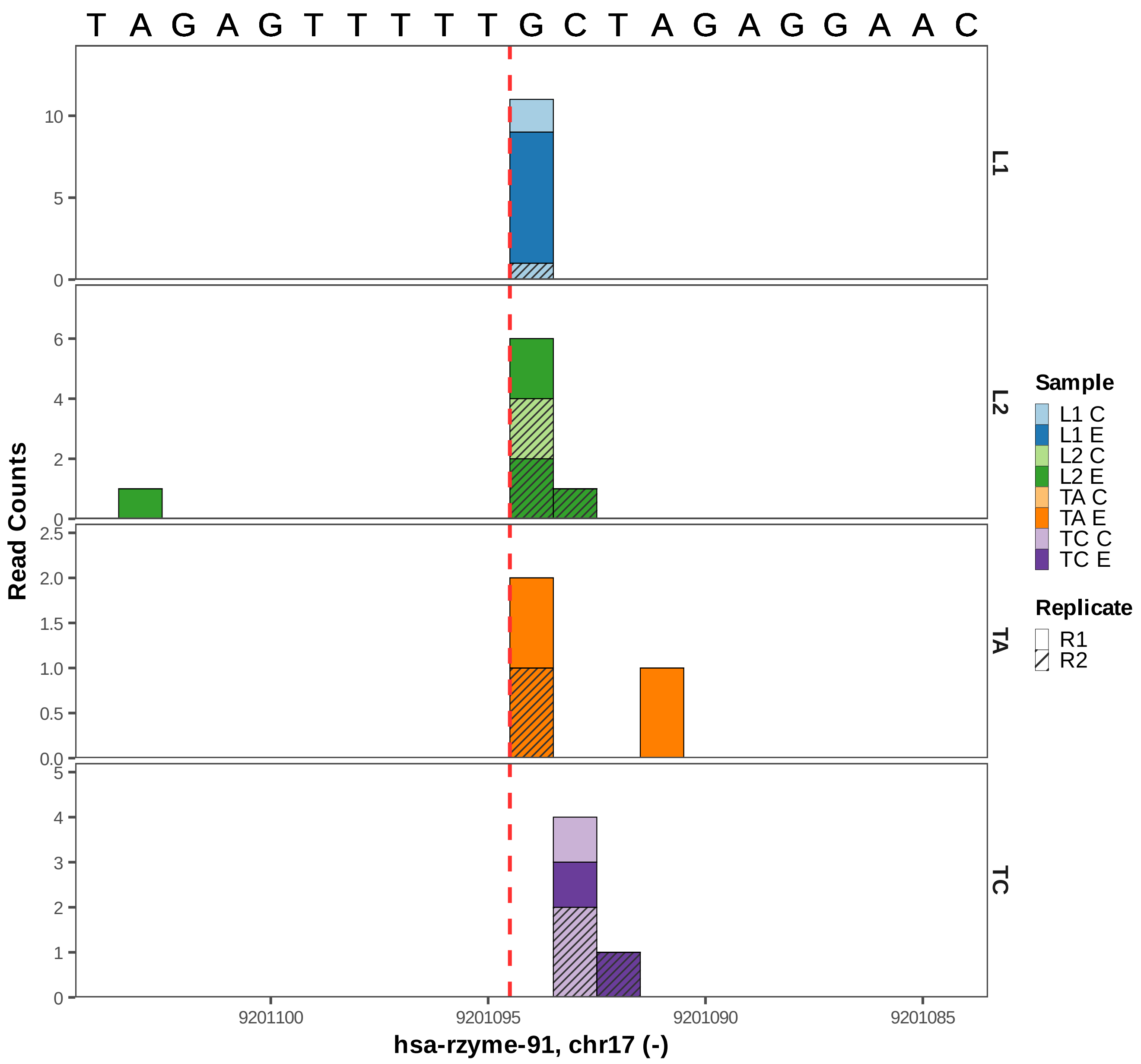

T A G A G T T T T T G C T A G A G G A A C
L1
L2
TA
TC
Read Counts
10
5
0
6
4
2
0
2.5
2.0
1.5
1.0
0.5
0.0
5
4
3
2
1
0
9201100
9201095
9201090
9201085
hsa-rzyme-91, chr17 (-)
Sample
L1 C
L1 E
L2 C
L2 E
TA C
TA E
TC C
TC E
Replicate
R1
R2

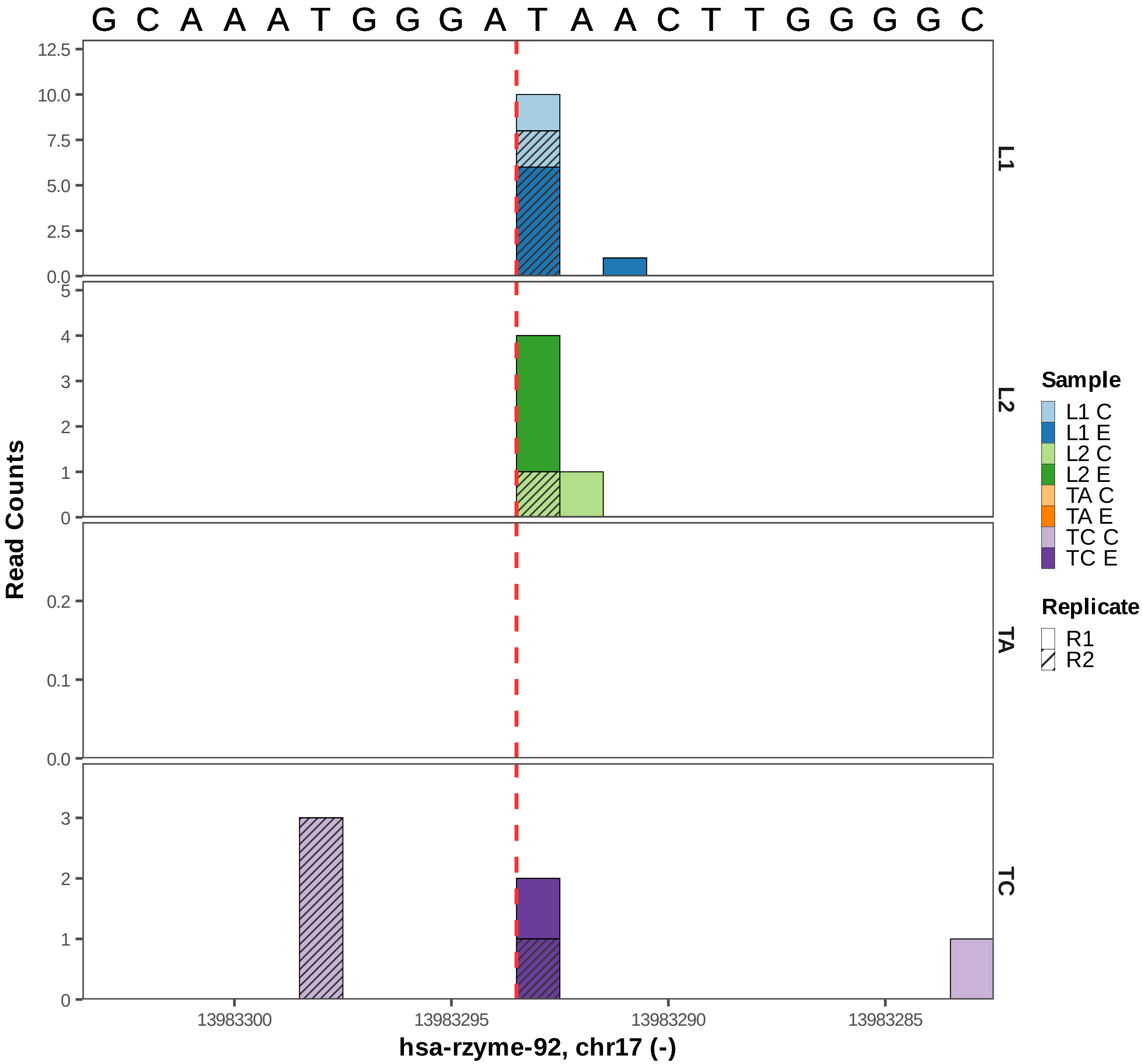

G C A A A T G G G A T A A C T T G G G G C
Read Counts
hsa-rzyme-92, chr17 (-)
13983300
13983295
13983290
13983285
L1
L2
TA
TC
Sample
L1 C
L1 E
L2 C
L2 E
TA C
TA E
TC C
TC E
Replicate
R1
R2

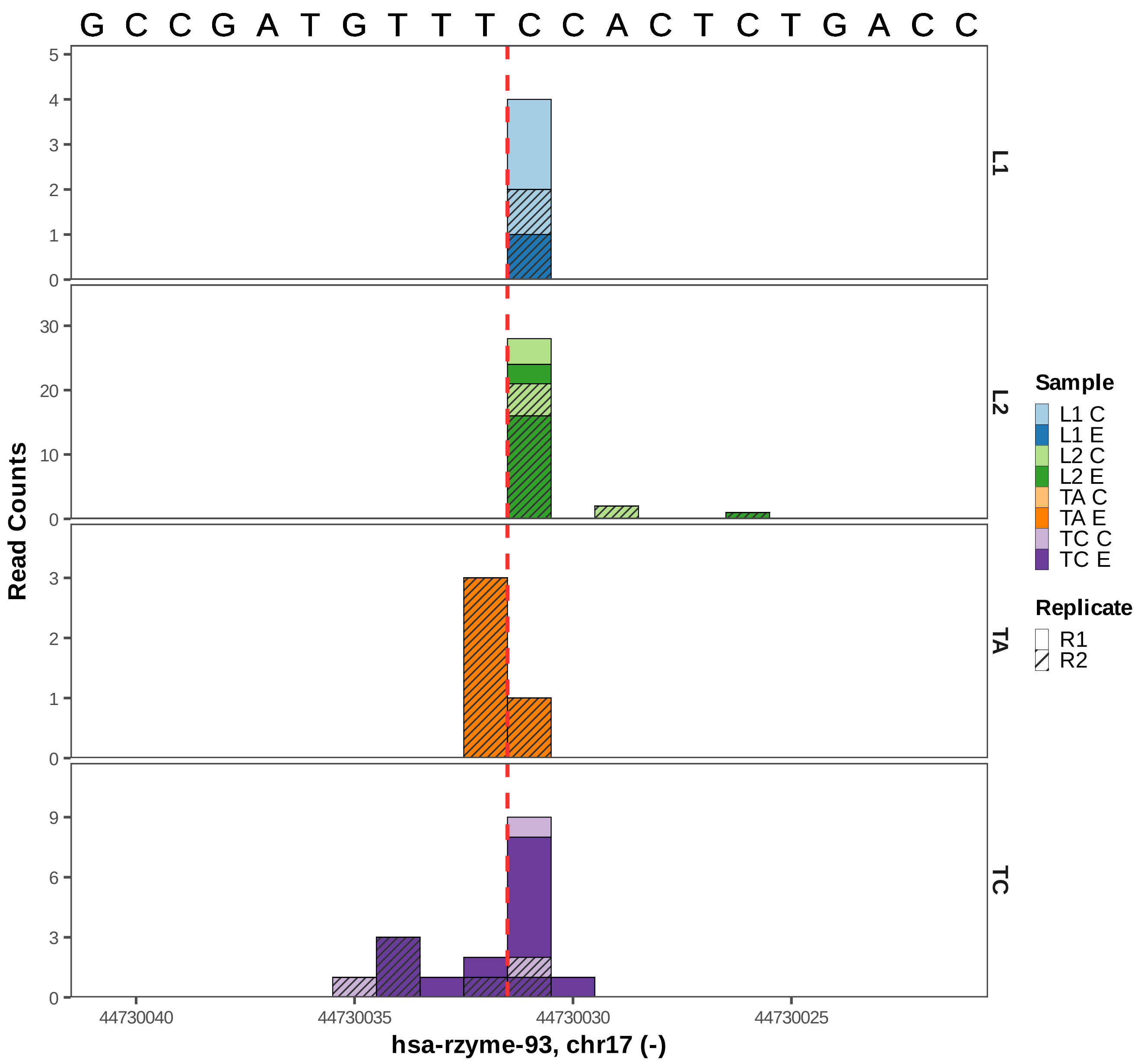
G C C G A T G T T T C C A C T C T G A C C
L1
L2
TA
TC
Read Counts
44730040
44730035
44730030
44730025
hsa-rzyme-93, chr17 (-)
Sample
L1 C
L1 E
L2 C
L2 E
TA C
TA E
TC C
TC E
Replicate
R1
R2

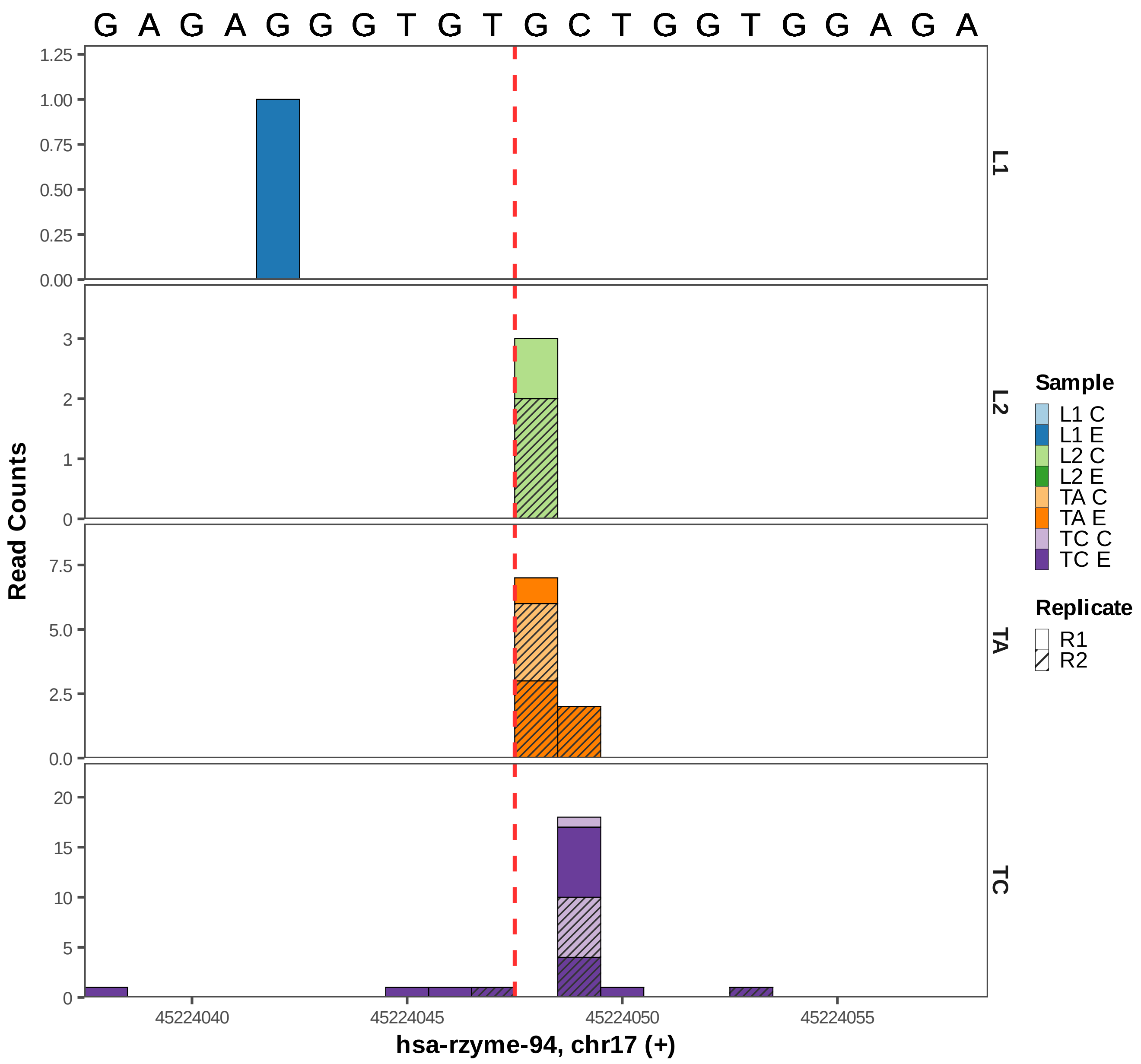

G A G A G G G T G T G C T G G T G G A G A
1.25
1.00
0.75
0.50
0.25
0.00
L1
3
2
1
0
L2
7.5
5.0
2.5
0.0
TA
20
15
10
5
0
TC
Read Counts
45224040
45224045
45224050
45224055
hsa-rzyme-94, chr17 (+)
Sample
L1 C
L1 E
L2 C
L2 E
TA C
TA E
TC C
TC E
Replicate
R1
R2

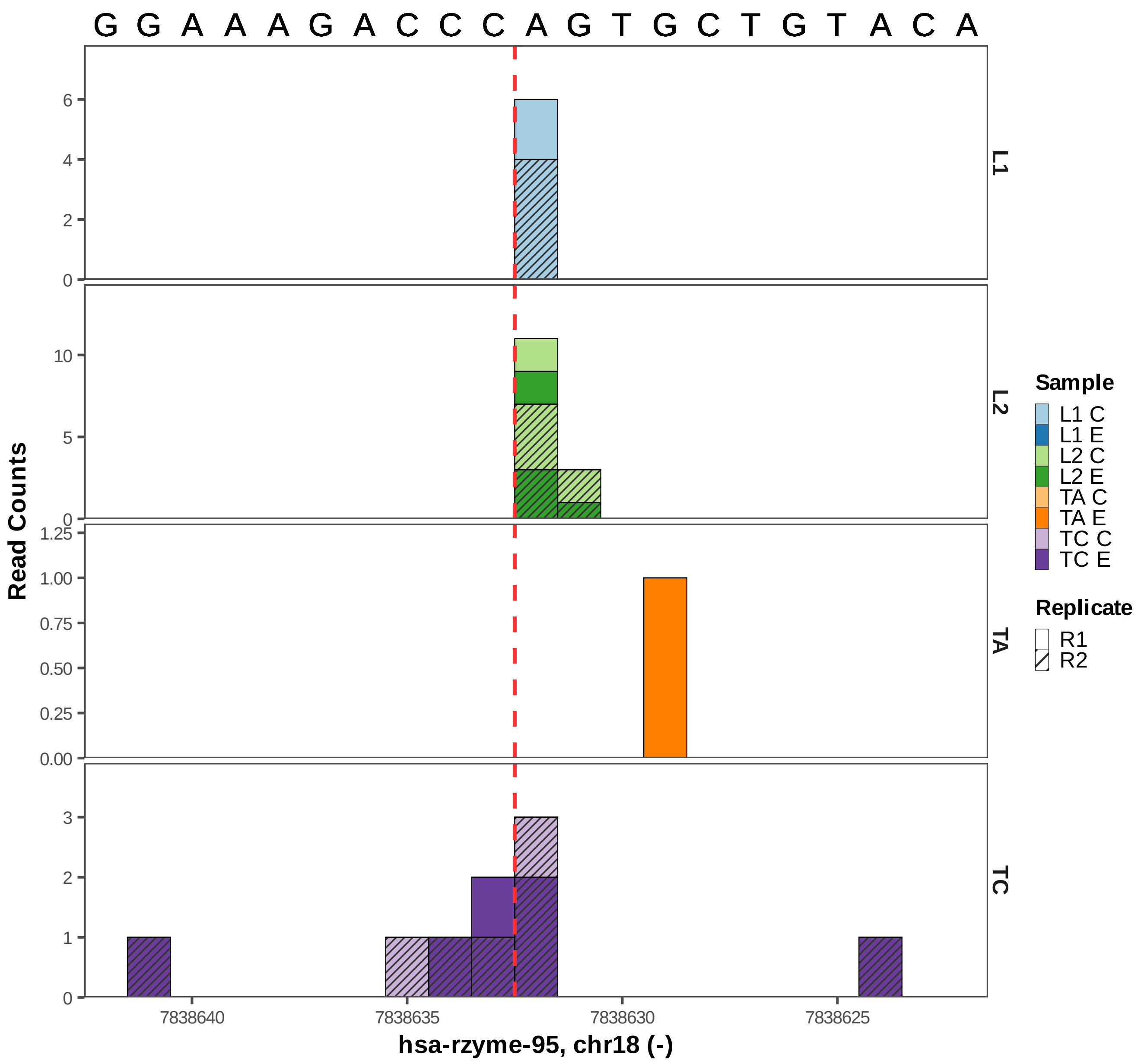

G G A A A G A C C C A G T G C T G T A C A
L1
L2
TA
TC
Read Counts
0
2
4
6
5
10
0.00
0.25
0.50
0.75
1.00
1.25
1
3
7838640
7838635
7838630
7838625
hsa-rzyme-95, chr18 (-)
Sample
L1 C
L1 E
L2 C
L2 E
TA C
TA E
TC C
TC E
Replicate
R1
R2

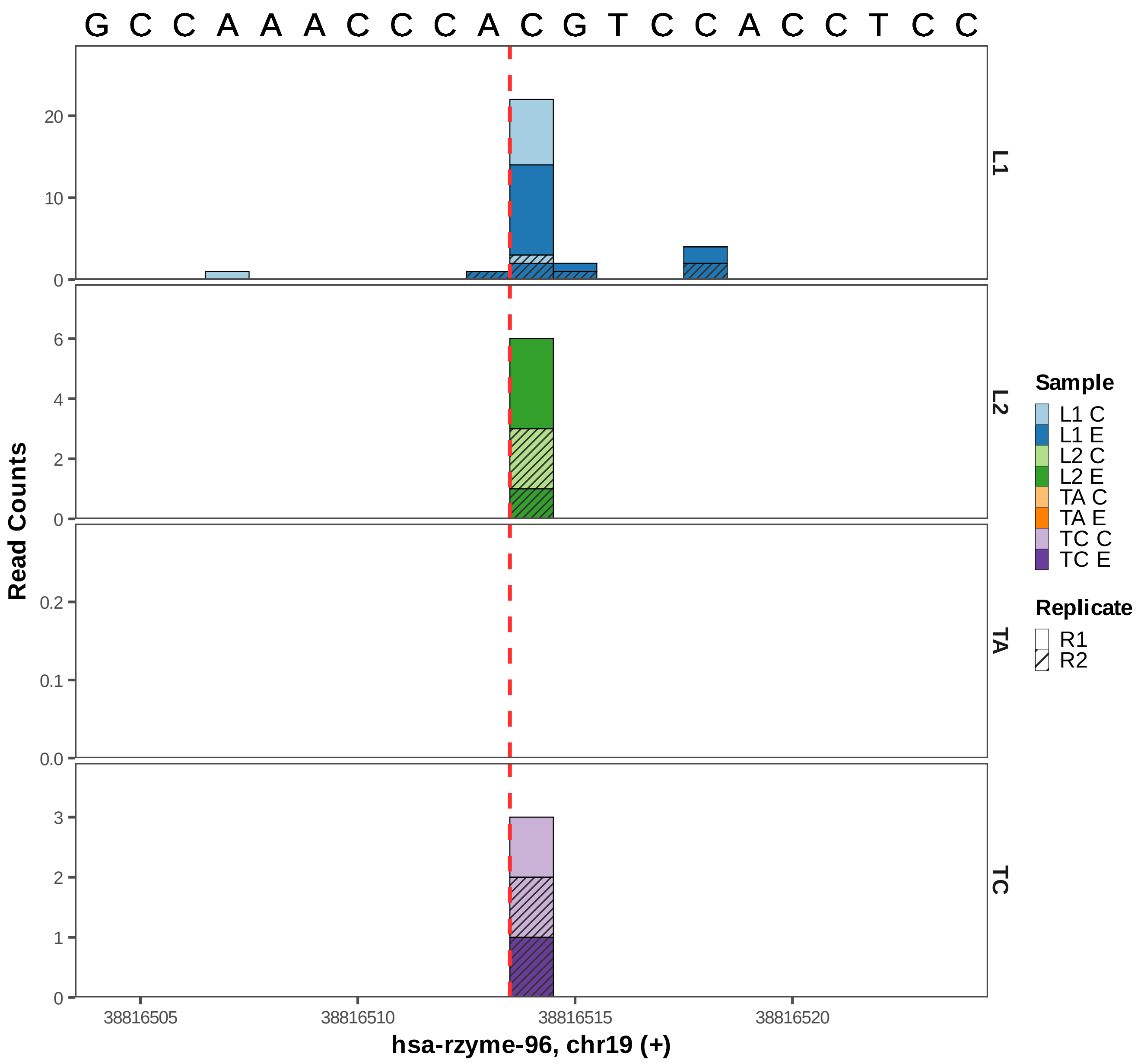

G C C A A A C C C A C G T C C A C C T C C
L1
L2
TA
TC
Read Counts
38816505
38816510
38816515
38816520
hsa-rzyme-96, chr19 (+)
Sample
L1 C
L1 E
L2 C
L2 E
TA C
TA E
TC C
TC E
Replicate
R1
R2

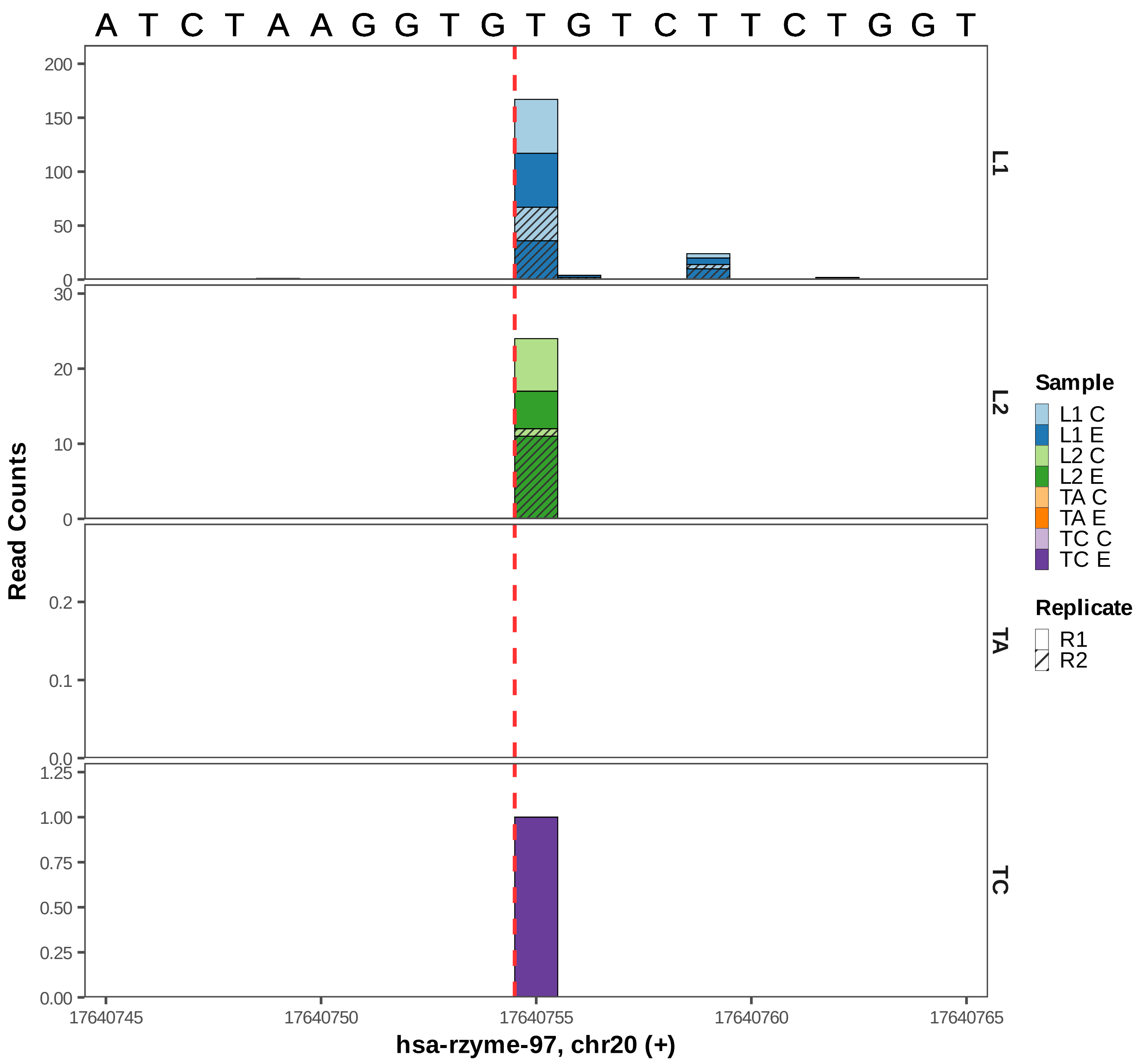

A T C T A A G G T G T G T C T T C T G G T
Read Counts
L1
L2
TA
TC
200
150
100
50
0
30
20
10
0
0.2
0.1
0.0
1.25
1.00
0.75
0.50
0.25
0.00
17640745
17640750
17640755
17640760
17640765
hsa-rzyme-97, chr20 (+)
Sample
L1 C
L1 E
L2 C
L2 E
TA C
TA E
TC C
TC E
Replicate
R1
R2

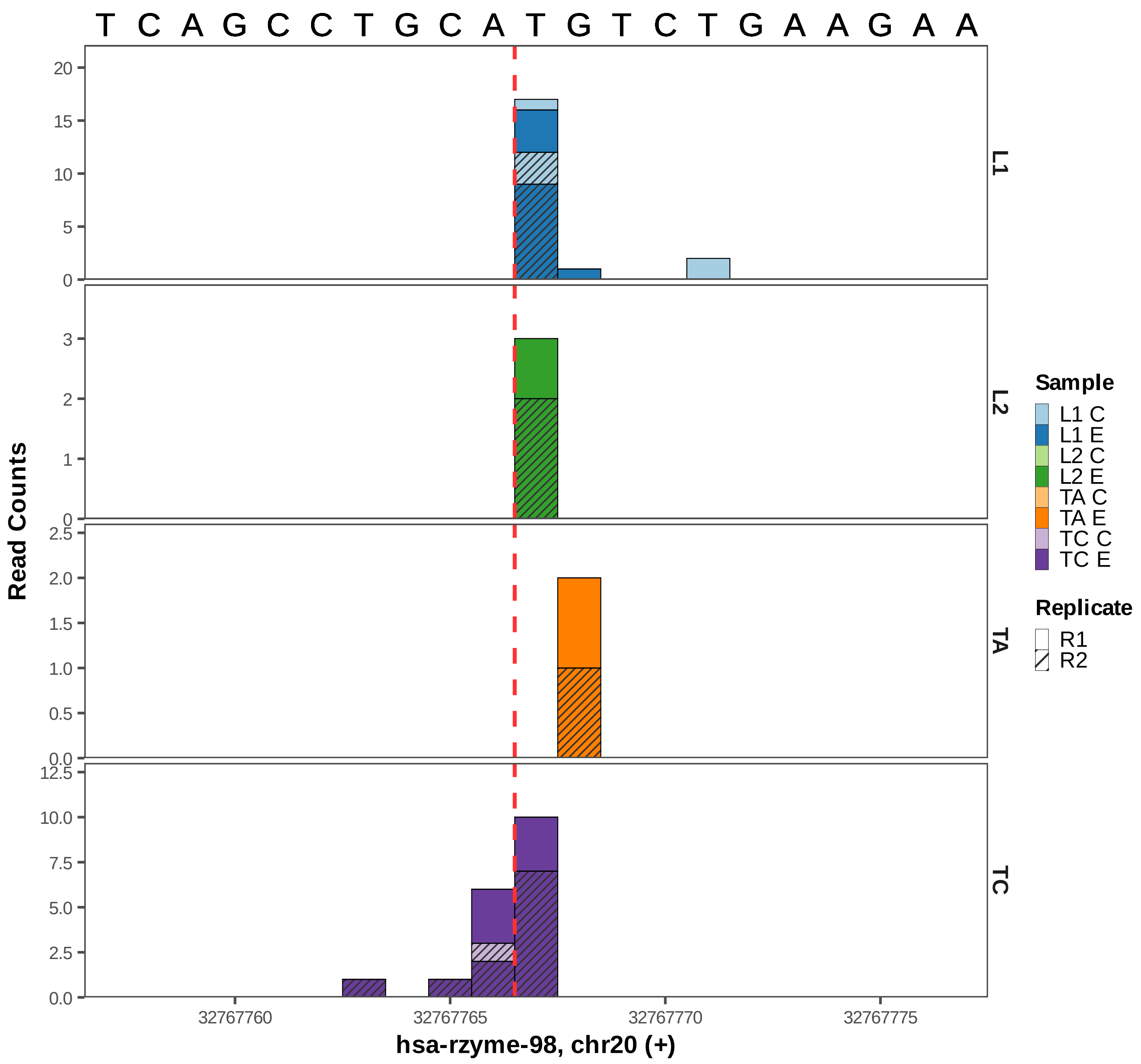

T C A G C C T G C A T G T C T G A A G A A
L1
L2
TA
TC
Read Counts
Sample
L1 C
L1 E
L2 C
L2 E
TA C
TA E
TC C
TC E
Replicate
R1
R2
32767760
32767765
32767770
32767775
hsa-rzyme-98, chr20 (+)

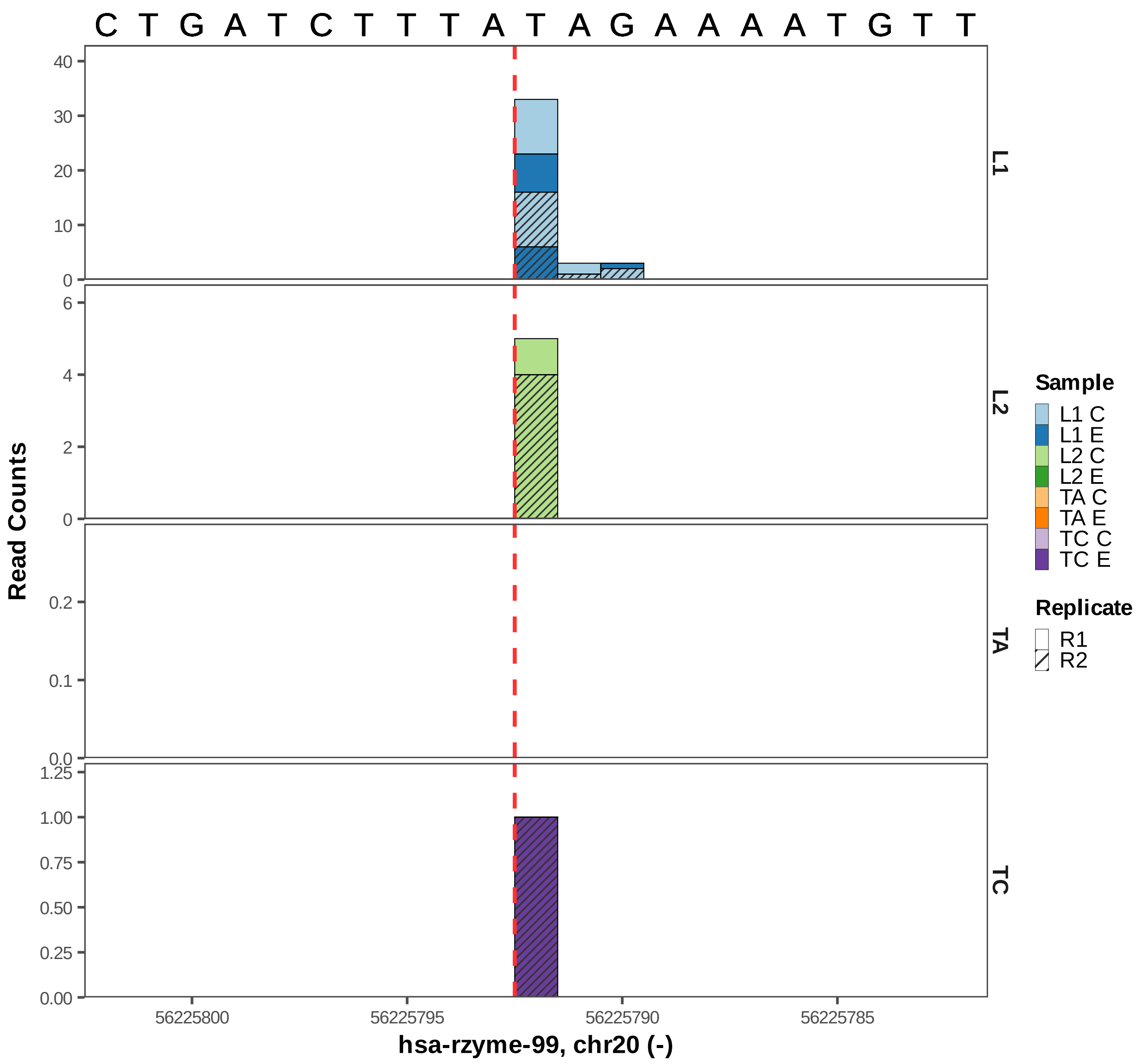

C T G A T C T T T A T A G A A A A T G T T
Read Counts
40
30
20
10
0
6
4
2
0
0.2
0.1
0.0
1.25
1.00
0.75
0.50
0.25
0.00
L1
L2
TA
TC
56225800
56225795
56225790
56225785
hsa-rzyme-99, chr20 (-)
Sample
L1 C
L1 E
L2 C
L2 E
TA C
TA E
TC C
TC E
Replicate
R1
R2

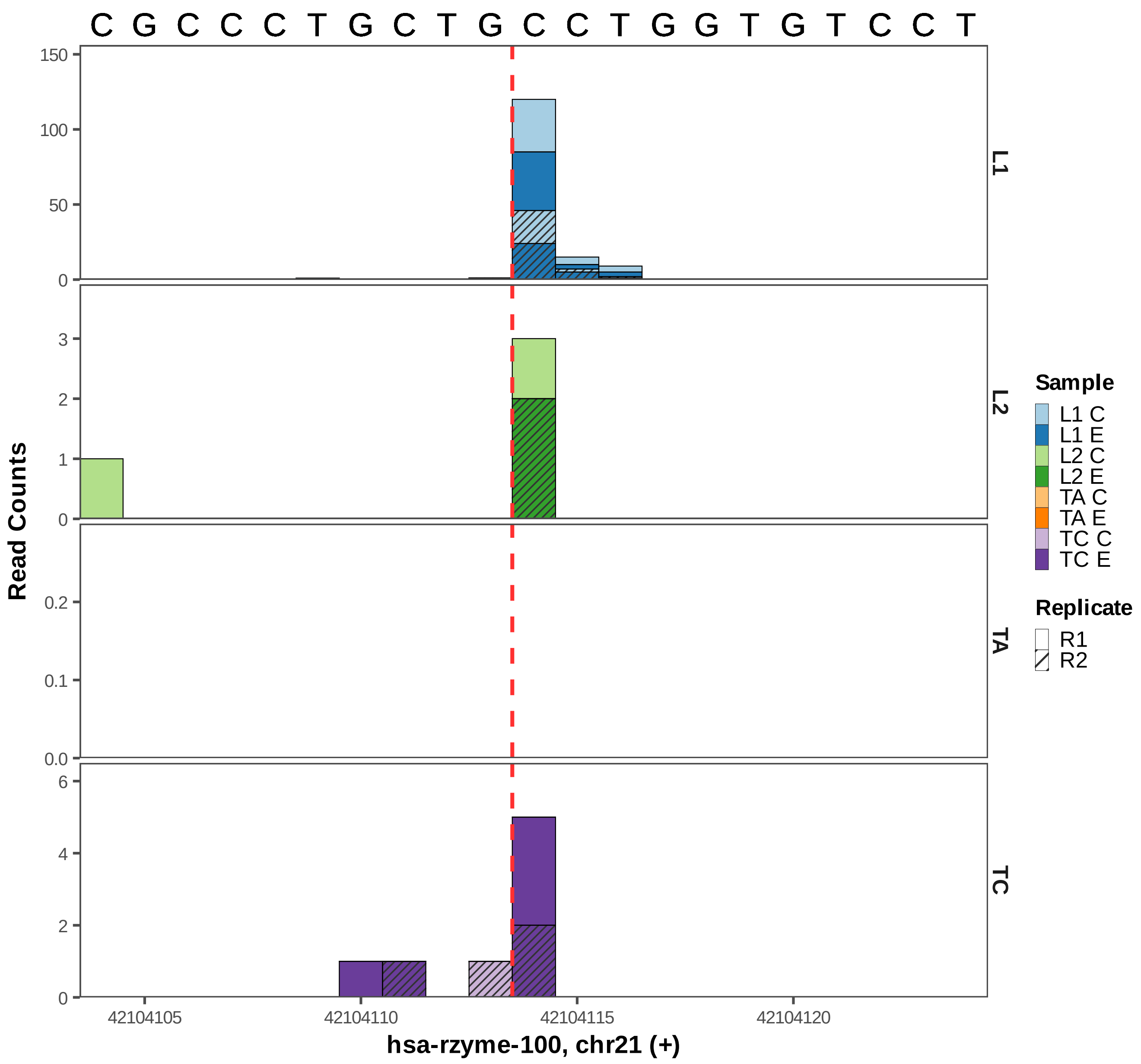
C G C C C T G C T G C C T G G T G T C C T
150
100
50
0
3
2
1
0
0.2
0.1
0.0
6
4
2
0
L1
L2
TA
TC
Read Counts
42104105
42104110
42104115
42104120
hsa-rzyme-100, chr21 (+)
Sample
L1 C
L1 E
L2 C
L2 E
TA C
TA E
TC C
TC E
Replicate
R1
R2

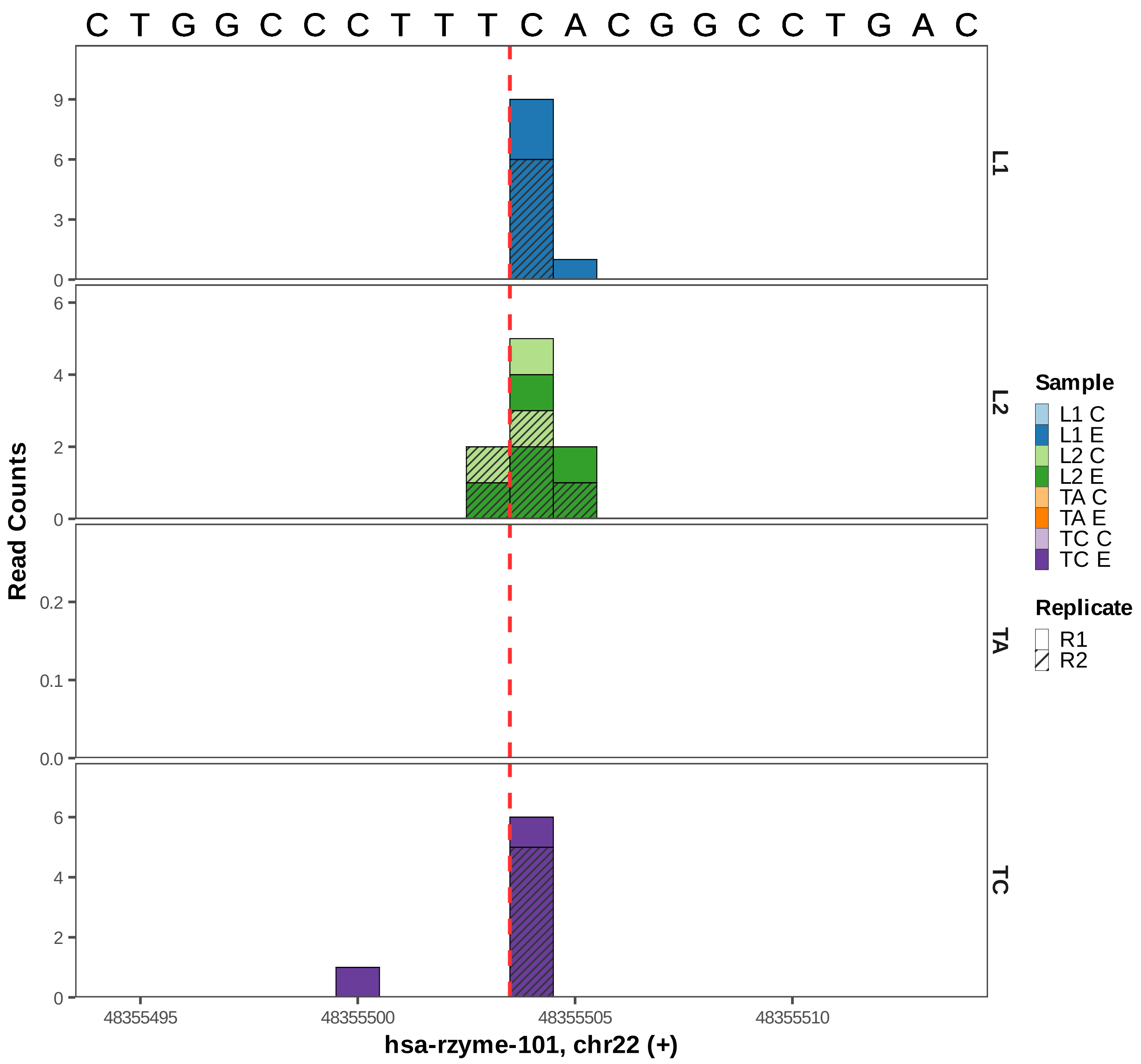
C T G G C C C T T T C A C G G C C T G A C
Read Counts
L1
L2
TA
TC
Sample
L1 C
L1 E
L2 C
L2 E
TA C
TA E
TC C
TC E
Replicate
R1
R2
48355495
48355500
48355505
48355510
hsa-rzyme-101, chr22 (+)

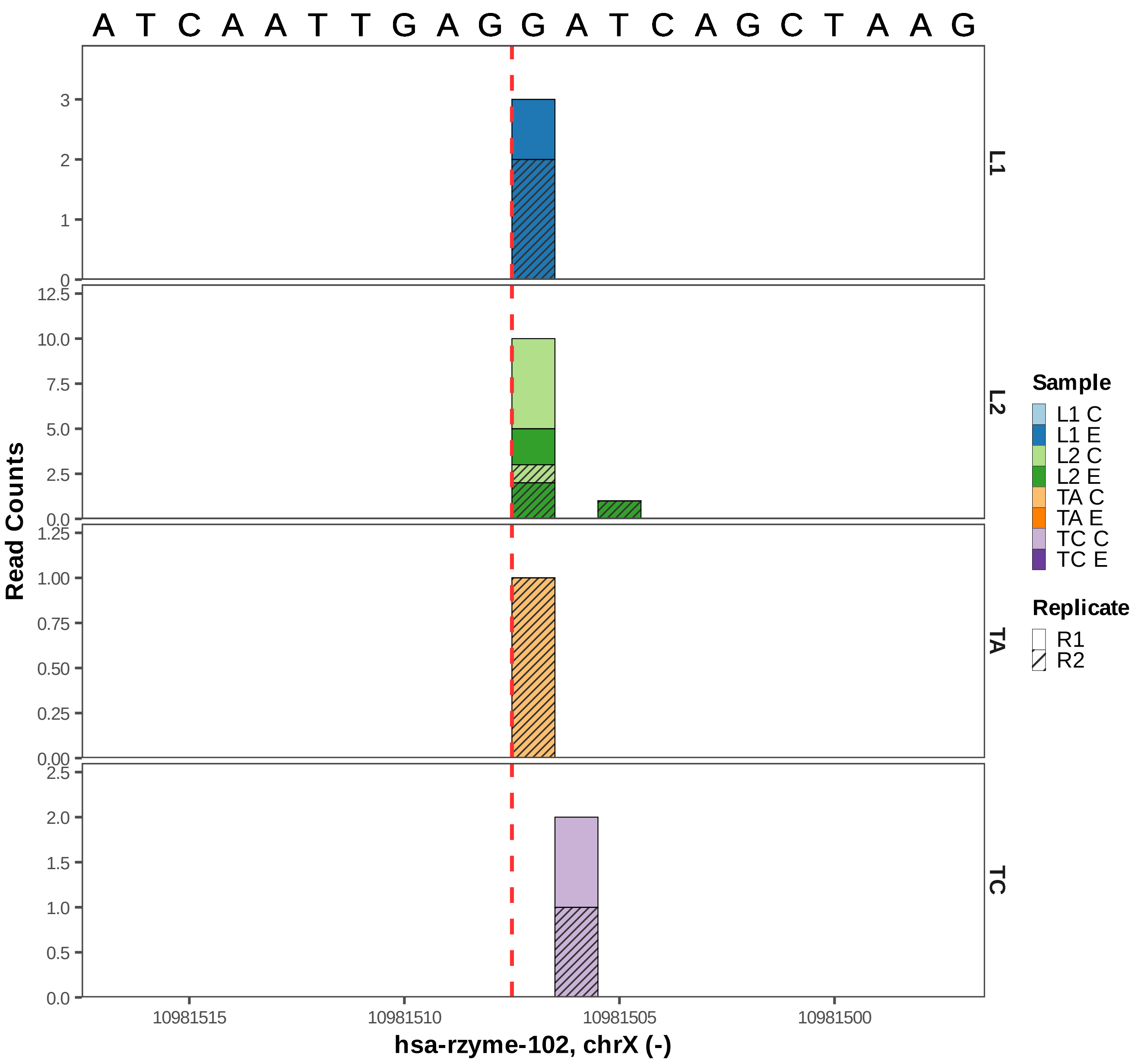

A T C A A T T G A G G A T C A G C T A A G
Read Counts
L1
L2
TA
TC
10981515
10981510
10981505
10981500
hsa-rzyme-102, chrX (-)
Sample
L1 C
L1 E
L2 C
L2 E
TA C
TA E
TC C
TC E
Replicate
R1
R2

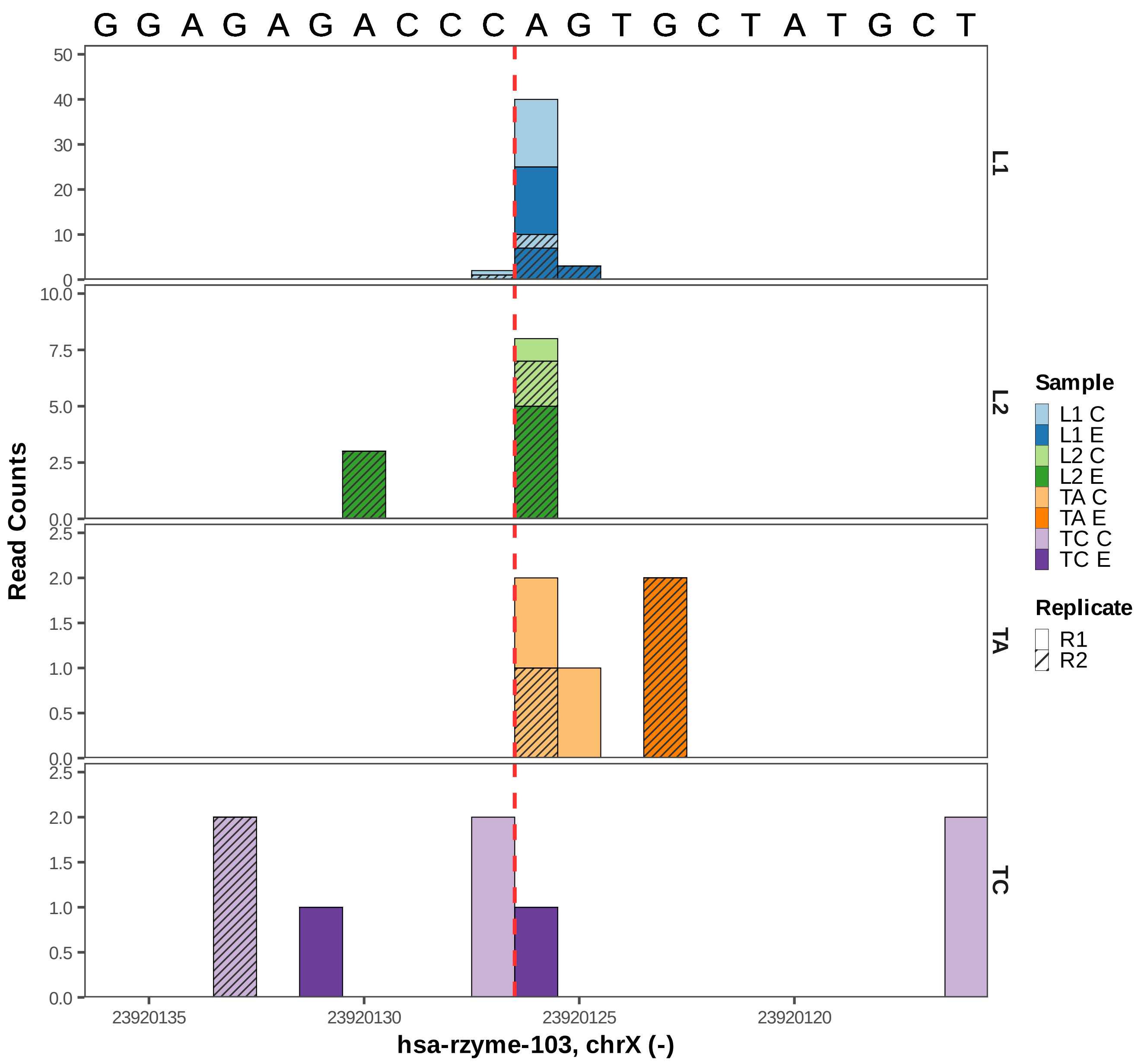
G G A G A G A C C C A G T G C T A T G C T
Read Counts
L1
L2
TA
TC
hsa-rzyme-103, chrX (-)
23920135
23920130
23920125
23920120
Sample
L1 C
L1 E
L2 C
L2 E
TA C
TA E
TC C
TC E
Replicate
R1
R2

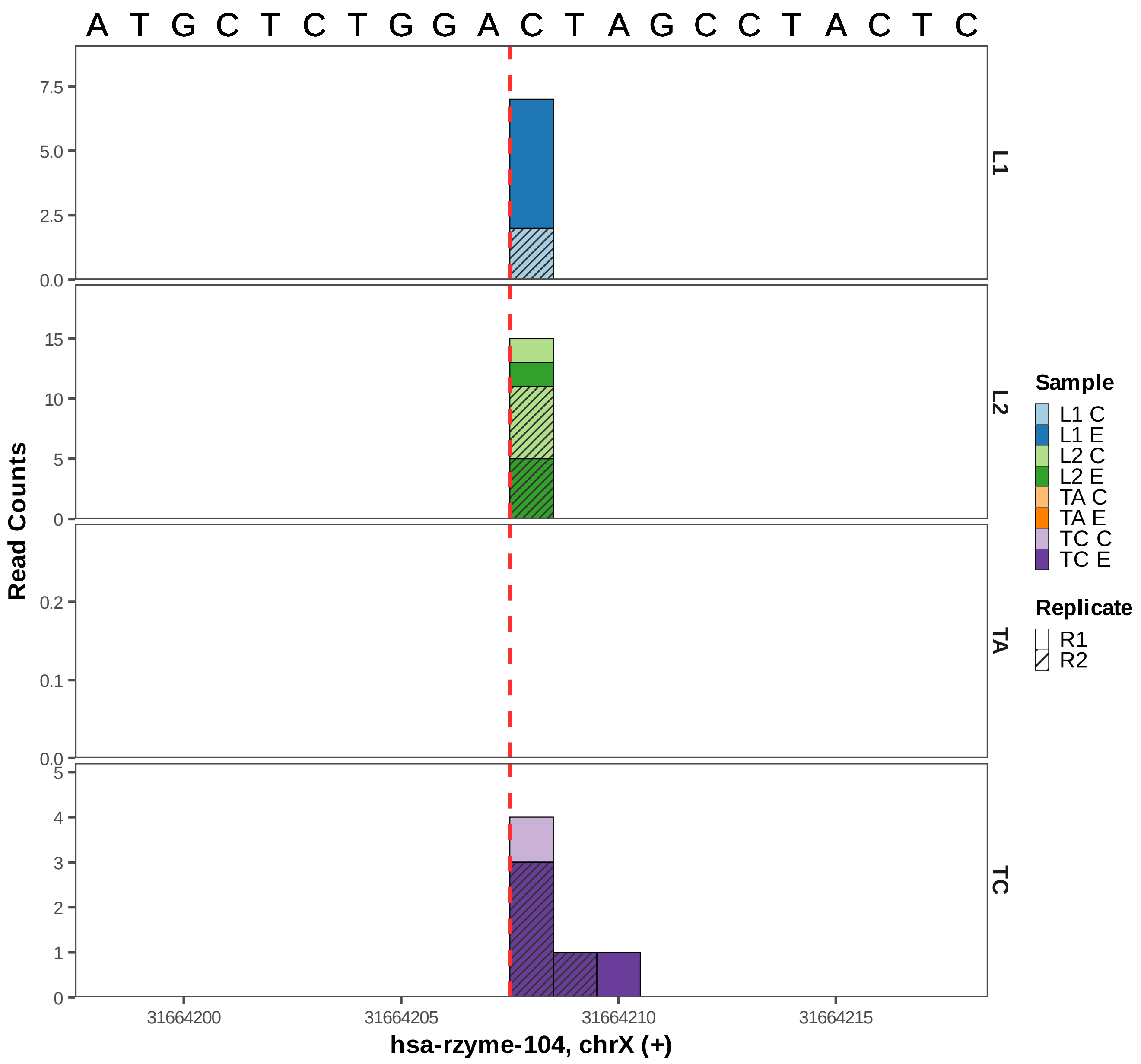

A T G C T C T G G A C T A G C C T A C T C
Read Counts
L1
L2
TA
TC
7.5
5.0
2.5
0.0
15
10
5
0
0.2
0.1
0.0
5
4
3
2
1
0
31664200
31664205
31664210
31664215
hsa-rzyme-104, chrX (+)
Sample
L1 C
L1 E
L2 C
L2 E
TA C
TA E
TC C
TC E
Replicate
R1
R2

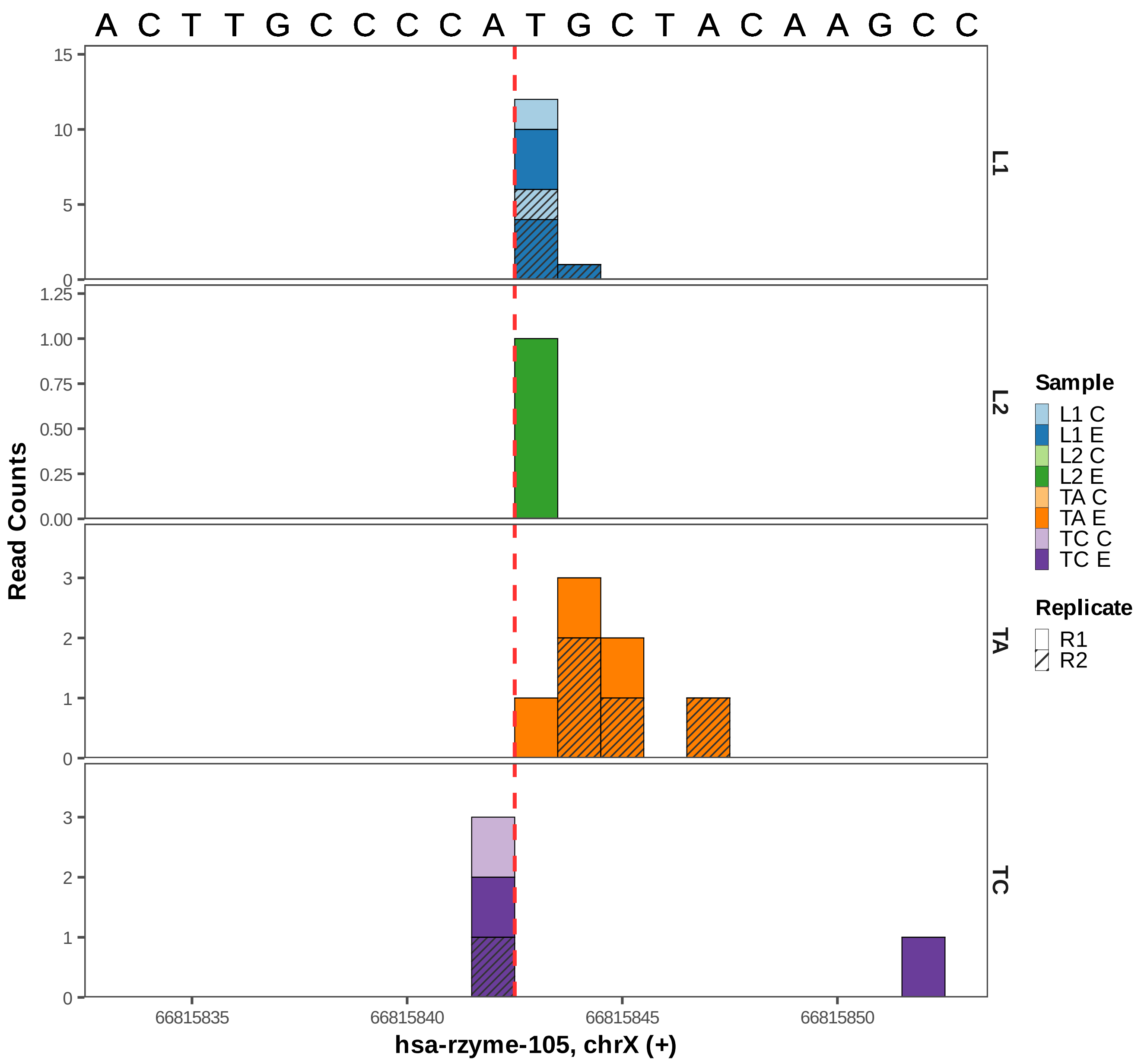

A C T T G C C C C A T G C T A C A A G C C
L1
L2
TA
TC
Read Counts
hsa-rzyme-105, chrX (+)
66815835
66815840
66815845
66815850
15
10
5
0
1.25
1.00
0.75
0.50
0.25
0.00
3
2
1
0
Sample
L1 C
L1 E
L2 C
L2 E
TA C
TA E
TC C
TC E
Replicate
R1
R2
