## Supplementary Figures for "Widespread Presence of Catalytic RNAs in All Domains of Life": Figure S7.pdf

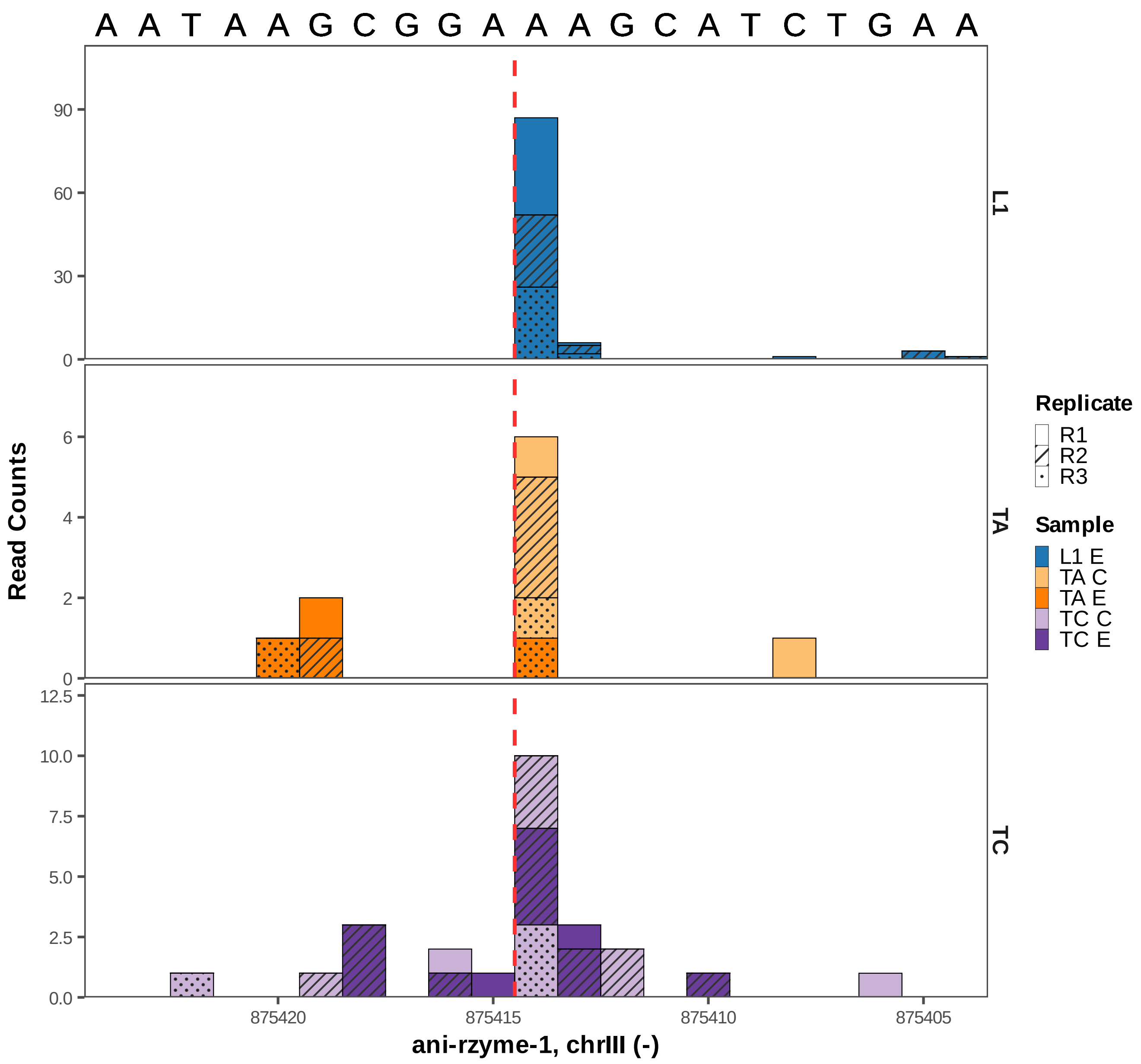

A A T A A G C G G A A A G C A T C T G A A
Read Counts
L1
TA
TC
Replicate
R1
R2
R3
Sample
L1 E
TA C
TA E
TC C
TC E
875420
875415
875410
875405
ani-rzyme-1, chrIII (-)

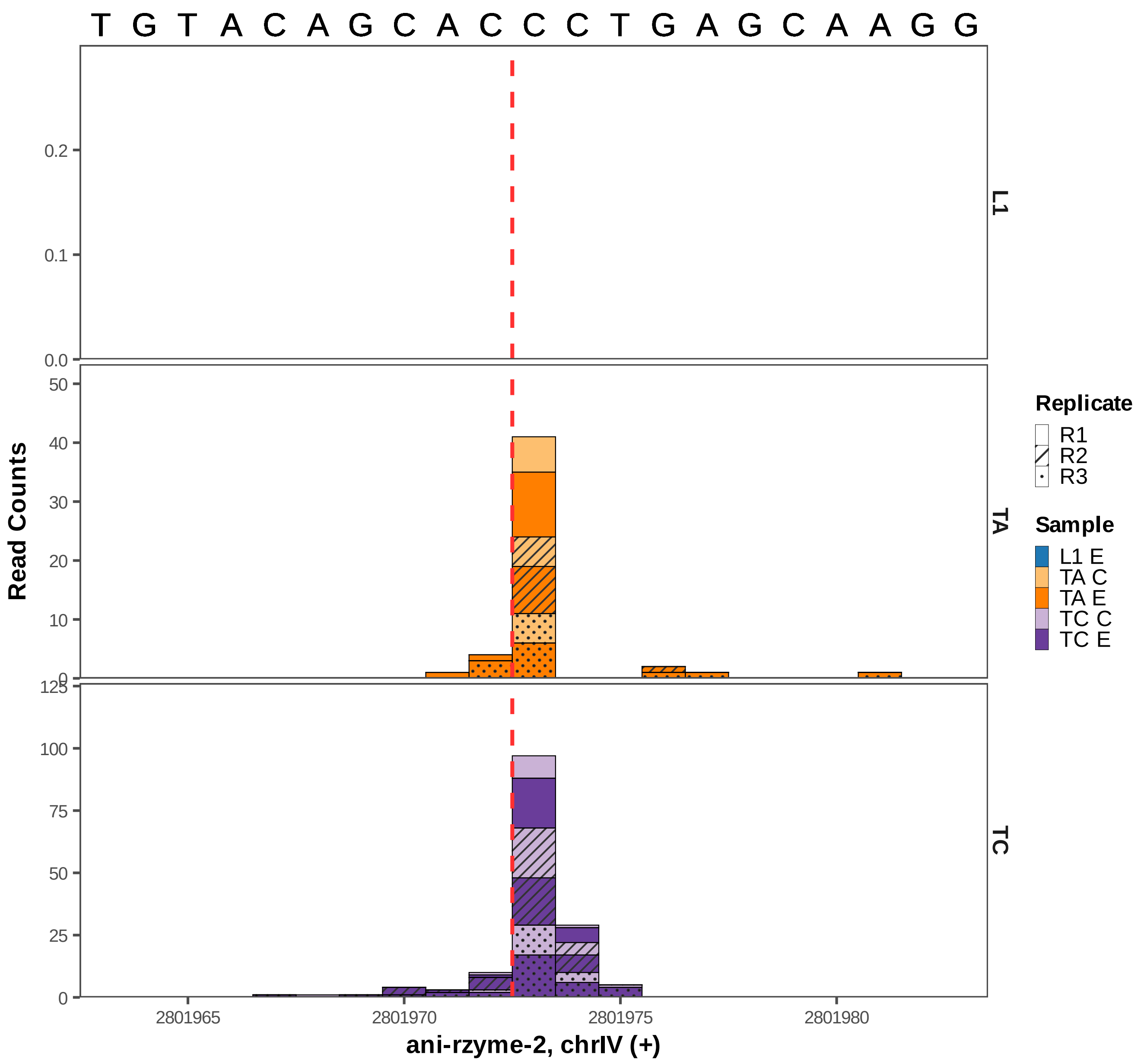

T G T A C A G C A C C C T G A G C A A G G
Read Counts
L1
TA
TC
0.0
0.1
0.2
0
10
20
30
40
50
0
25
50
75
100
125
2801965
2801970
2801975
2801980
ani-rzyme-2, chrIV (+)
Replicate
R1
R2
R3
Sample
L1 E
TA C
TA E
TC C
TC E

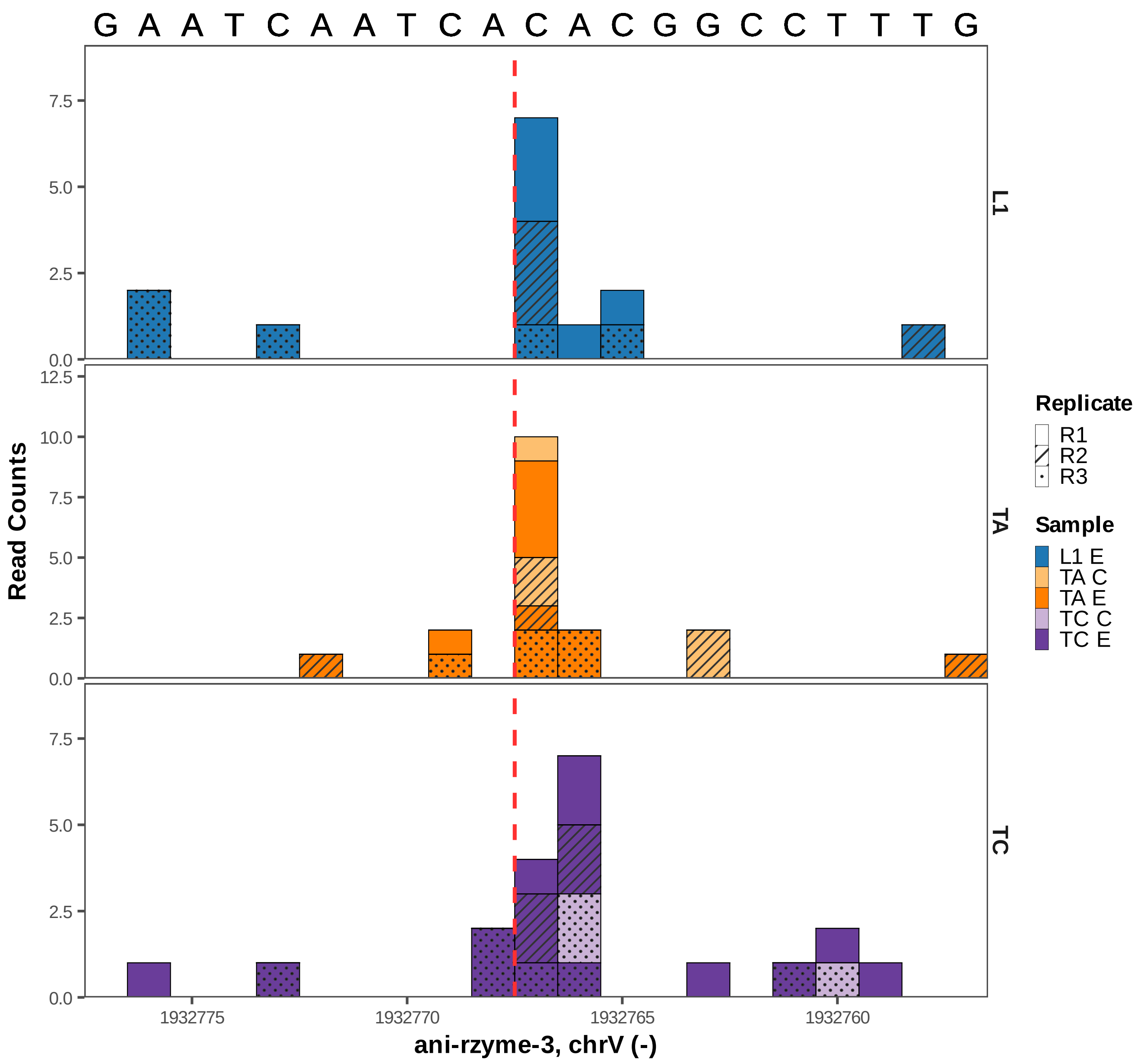

G A A T C A A T C A C A C G G C C T T T G
L1
TA
TC
Read Counts
ani-rzyme-3, chrV (-)
1932775
1932770
1932765
1932760
0.0
2.5
5.0
7.5
10.0
12.5
Replicate
R1
R2
R3
Sample
L1 E
TA C
TA E
TC C
TC E

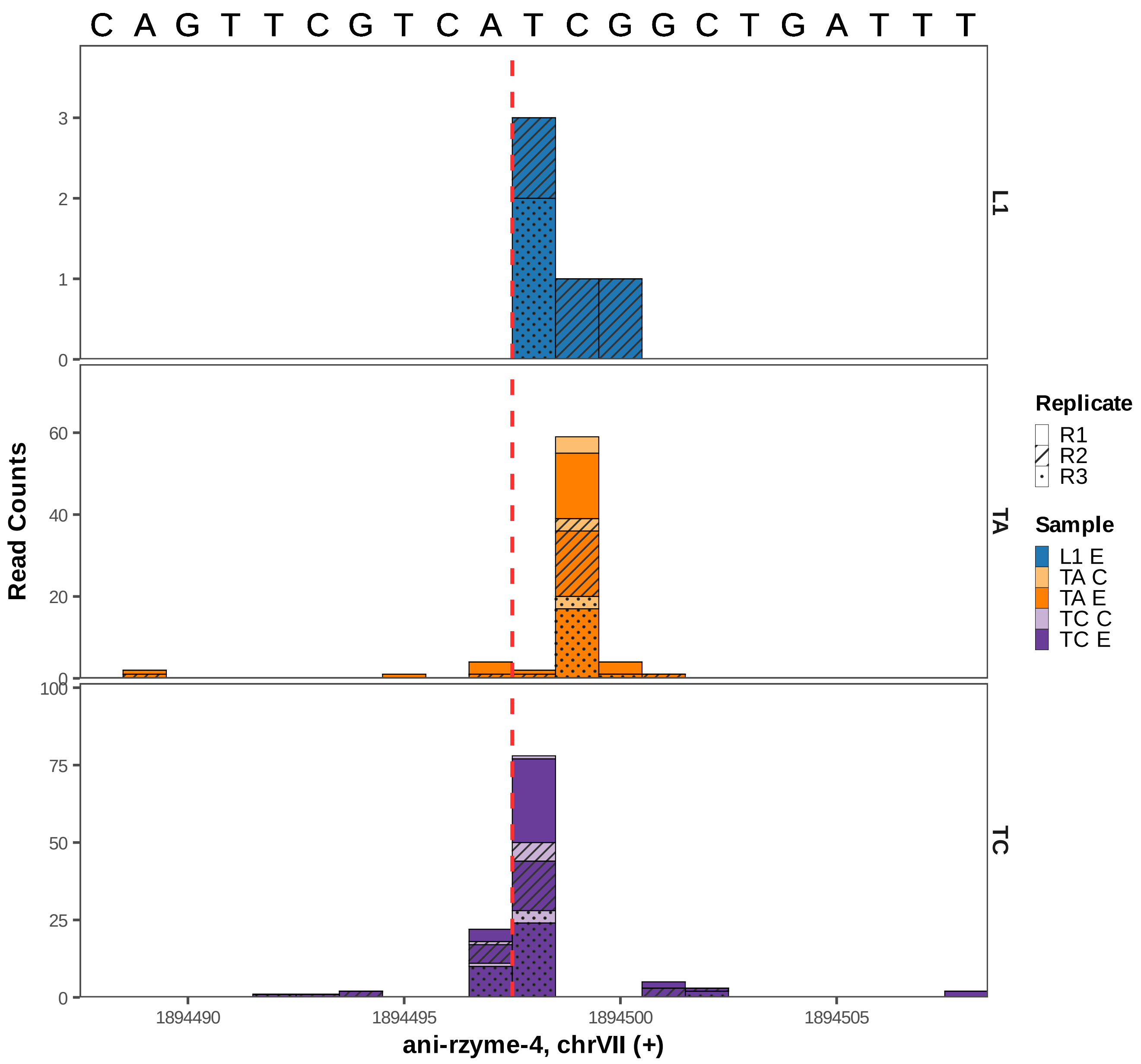

C A G T T C G T C A T C G G C T G A T T T
Read Counts
L1
TA
TC
0
1
2
3
20
40
60
25
50
75
100
1894490
1894495
1894500
1894505
ani-rzyme-4, chrVII (+)
Replicate
R1
R2
R3
Sample
L1 E
TA C
TA E
TC C
TC E

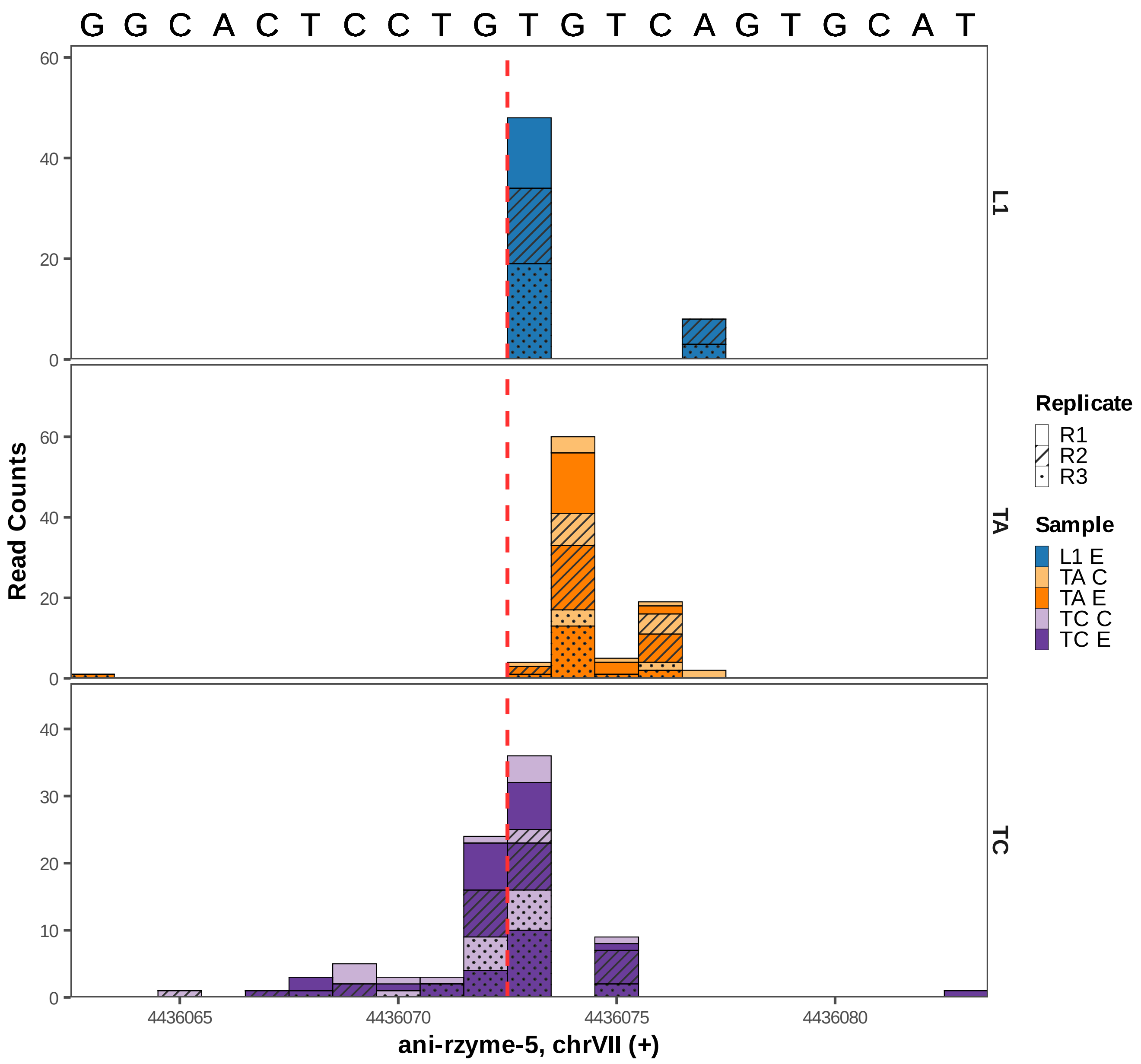

G G C A C T C C T G T G T C A G T G C A T
Read Counts
L1
TA
TC
60
40
20
0
10
30
4436065
4436070
4436075
4436080
ani-rzyme-5, chrVII (+)
Replicate
R1
R2
R3
Sample
L1 E
TA C
TA E
TC C
TC E

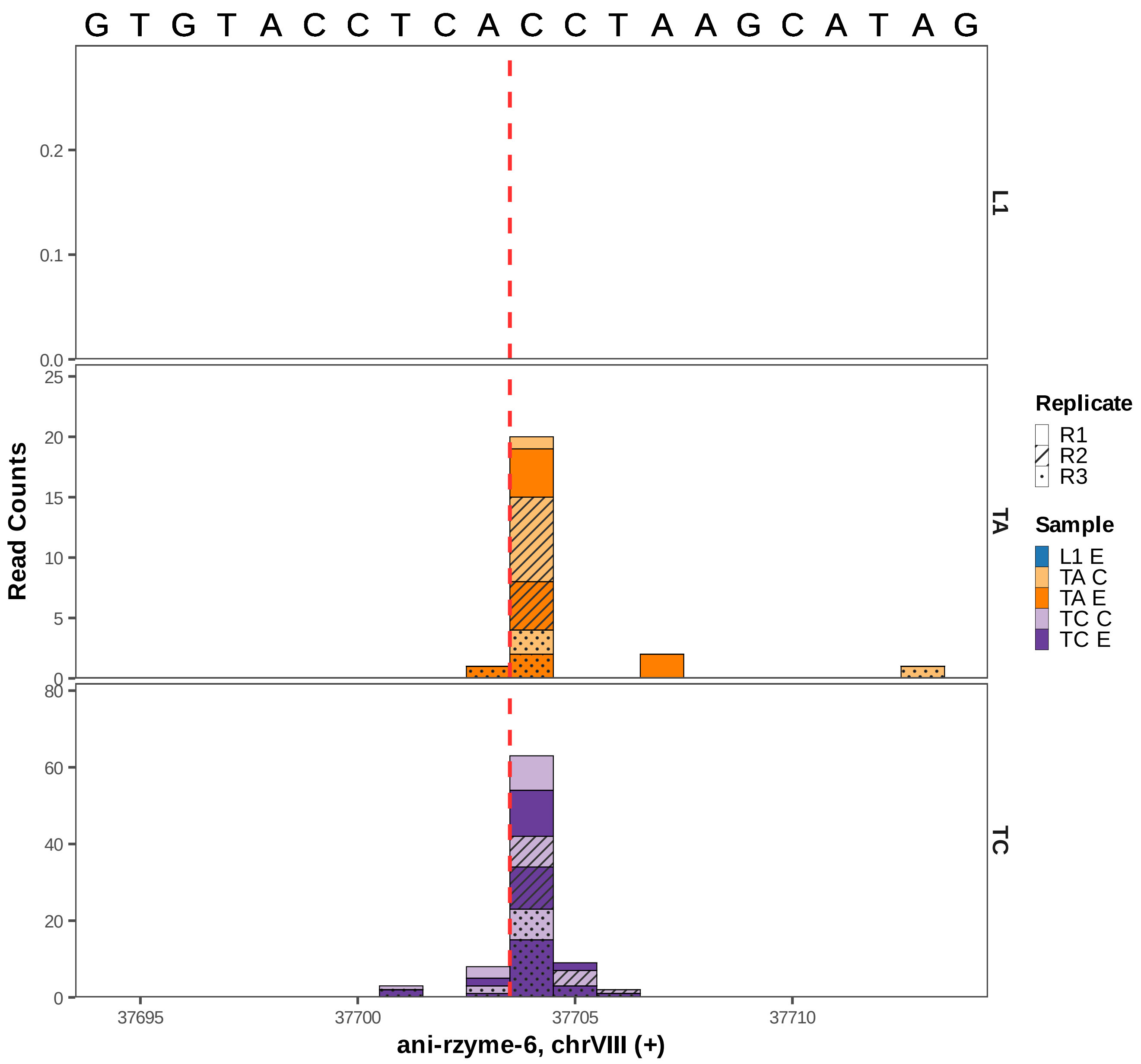

G T G T A C C T C A C C T A A G C A T A G
L1
TA
TC
Read Counts
ani-rzyme-6, chrVIII (+)
37695
37700
37705
37710
0.0
0.1
0.2
0
5
10
15
20
25
0
20
40
60
80
Replicate
R1
R2
R3
Sample
L1 E
TA C
TA E
TC C
TC E

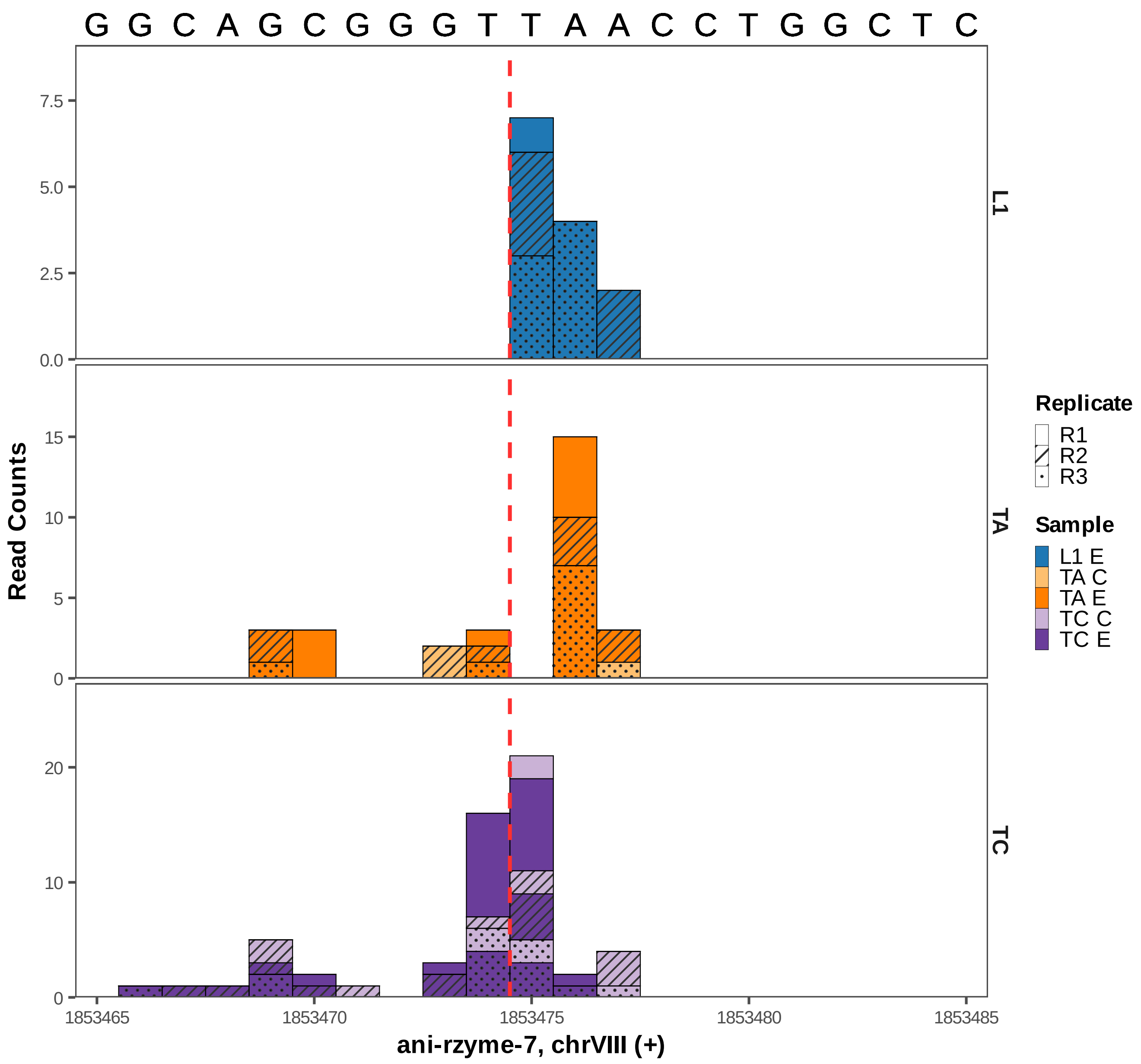

G G C A G C G G G T T A A C C T G G C T C
L1
TA
TC
Read Counts
ani-rzyme-7, chrVIII (+)
1853465
1853470
1853475
1853480
1853485
Replicate
R1
R2
R3
Sample
L1 E
TA C
TA E
TC C
TC E

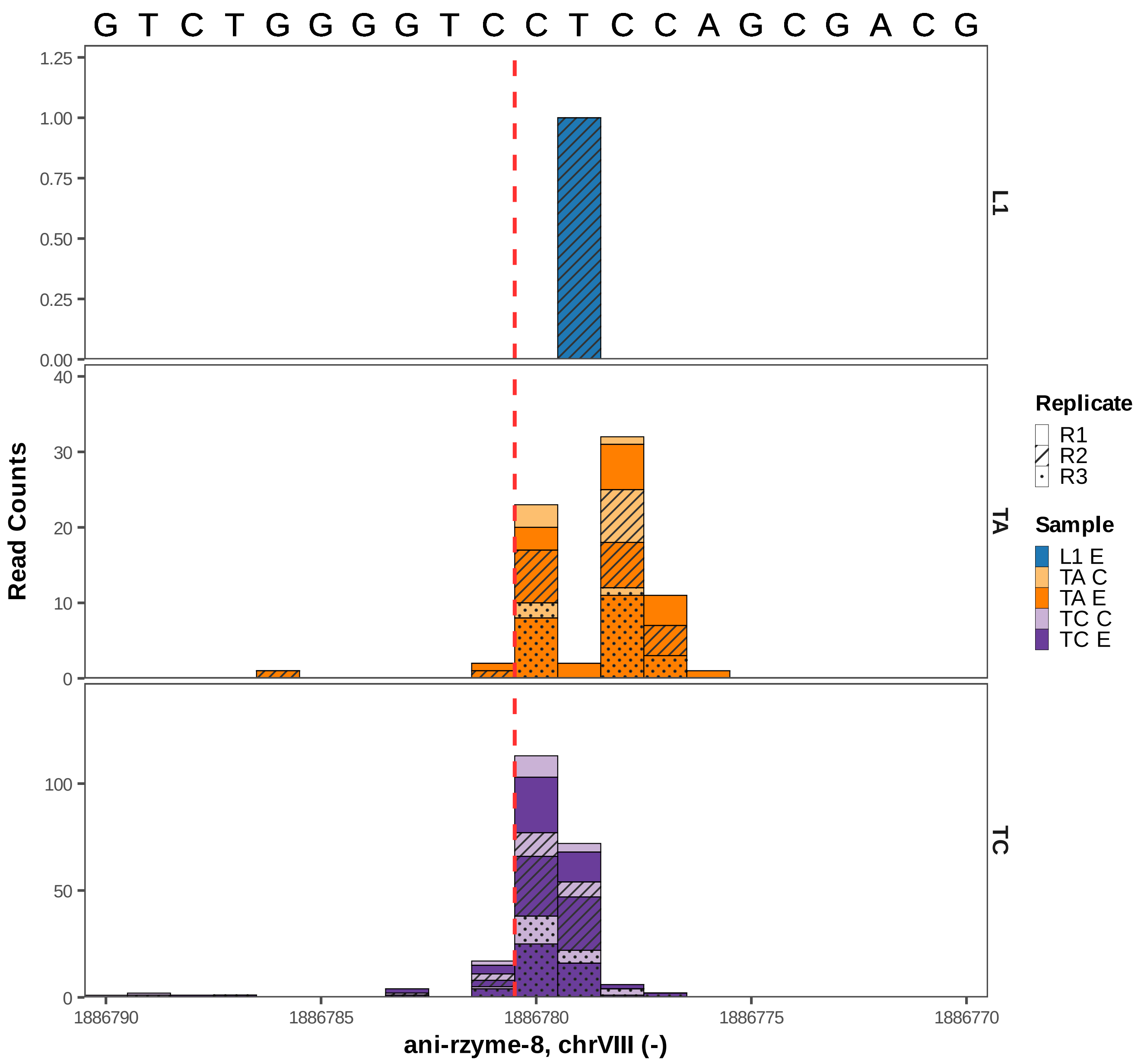

G T C T G G G G T C C T C C A G C G A C G
1.25
1.00
0.75
0.50
0.25
0.00
L1
40
30
20
10
0
TA
100
50
0
TC
Read Counts
1886790
1886785
1886780
1886775
1886770
ani-rzyme-8, chrVIII (-)
Replicate
R1
R2
R3
Sample
L1 E
TA C
TA E
TC C
TC E

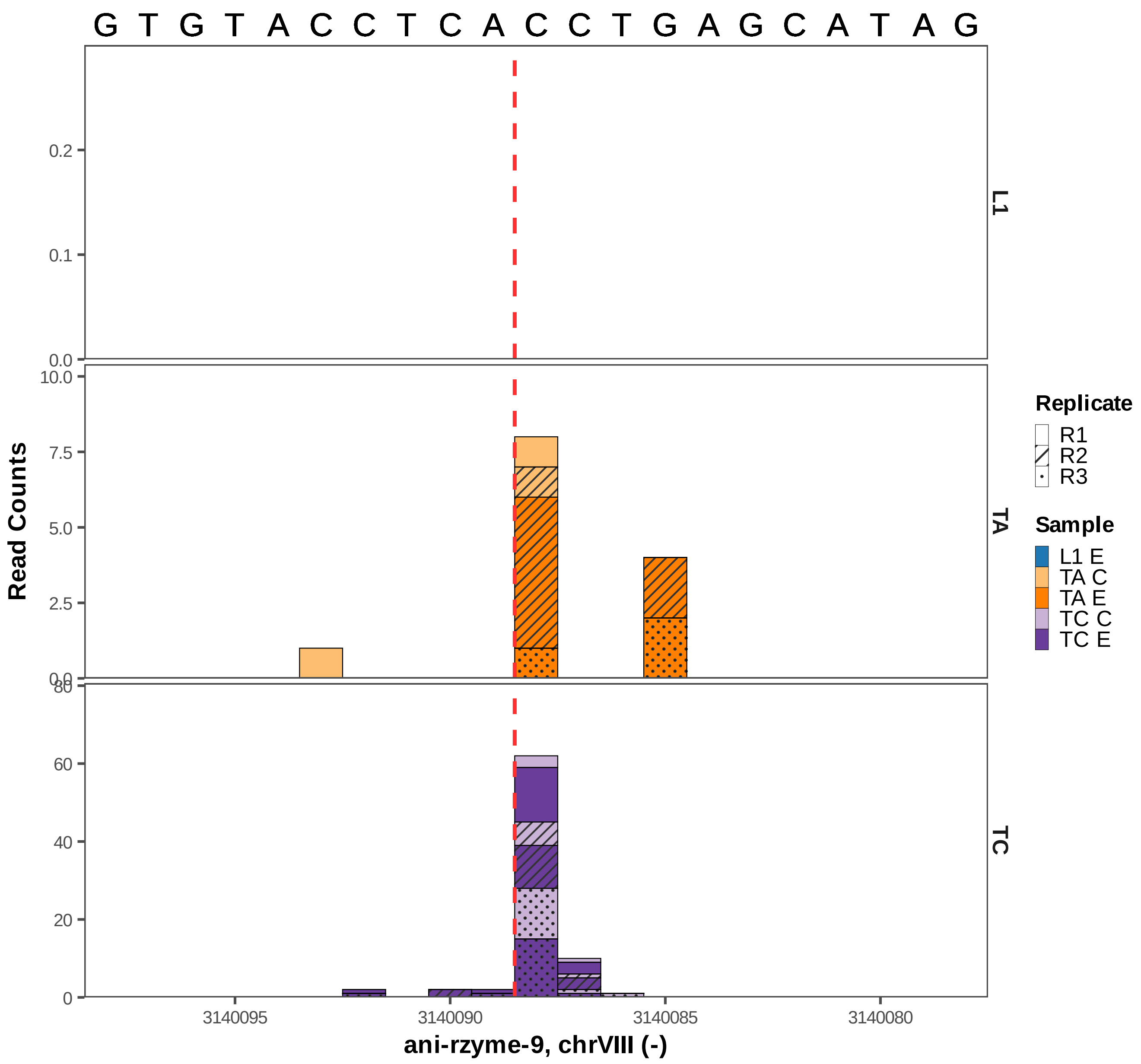

G T G T A C C T C A C C T G A G C A T A G
L1
TA
TC
Read Counts
0.0
0.1
0.2
0.0
2.5
5.0
7.5
10.0
0
20
40
60
80
3140095
3140090
3140085
3140080
ani-rzyme-9, chrVIII (-)
Replicate
R1
R2
R3
Sample
L1 E
TA C
TA E
TC C
TC E

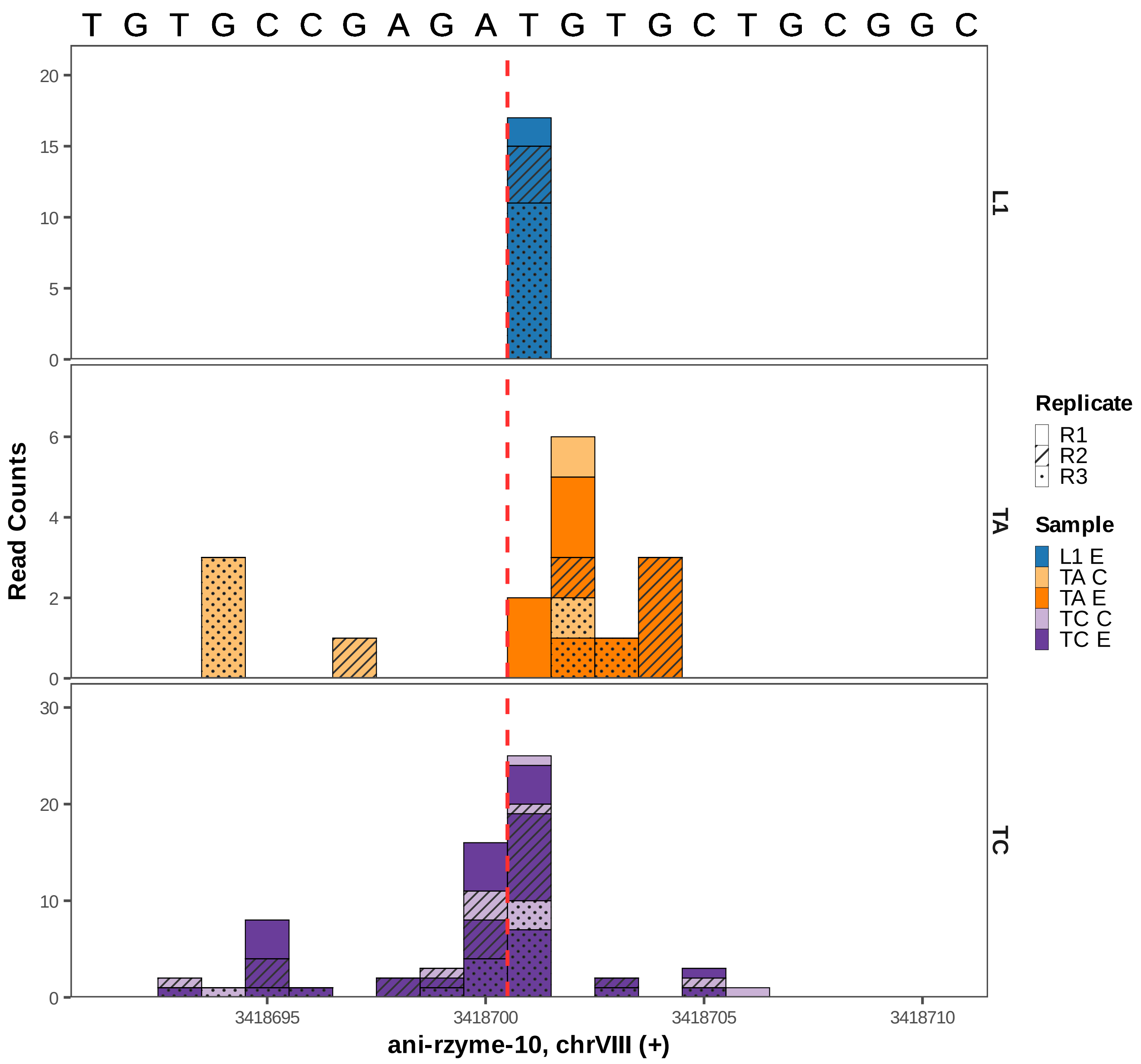

T G T G C C G A G A T G T G C T G C G G C
Read Counts
L1
TA
TC
Replicate
R1
R2
R3
Sample
L1 E
TA C
TA E
TC C
TC E
0
5
10
15
20
0
2
4
6
0
10
20
30
3418695
3418700
3418705
3418710
ani-rzyme-10, chrVIII (+)

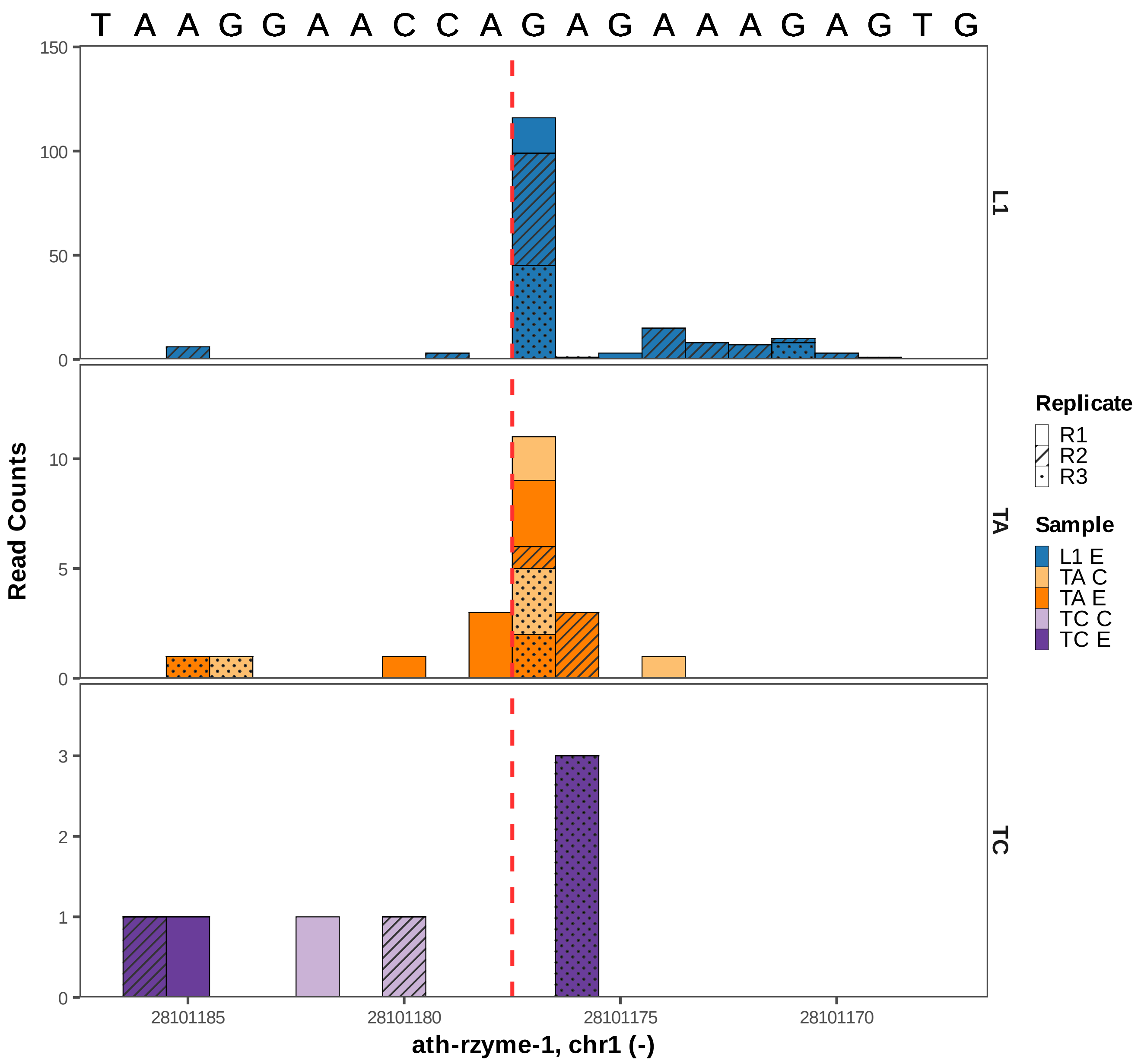

T A A G G A A C C A G A G A A A G A G T G
150
100
50
0
10
5
0
3
2
1
0
Read Counts
L1
TA
TC
Replicate
R1
R2
R3
Sample
L1 E
TA C
TA E
TC C
TC E
28101185
28101180
28101175
28101170
ath-rzyme-1, chr1 (-)

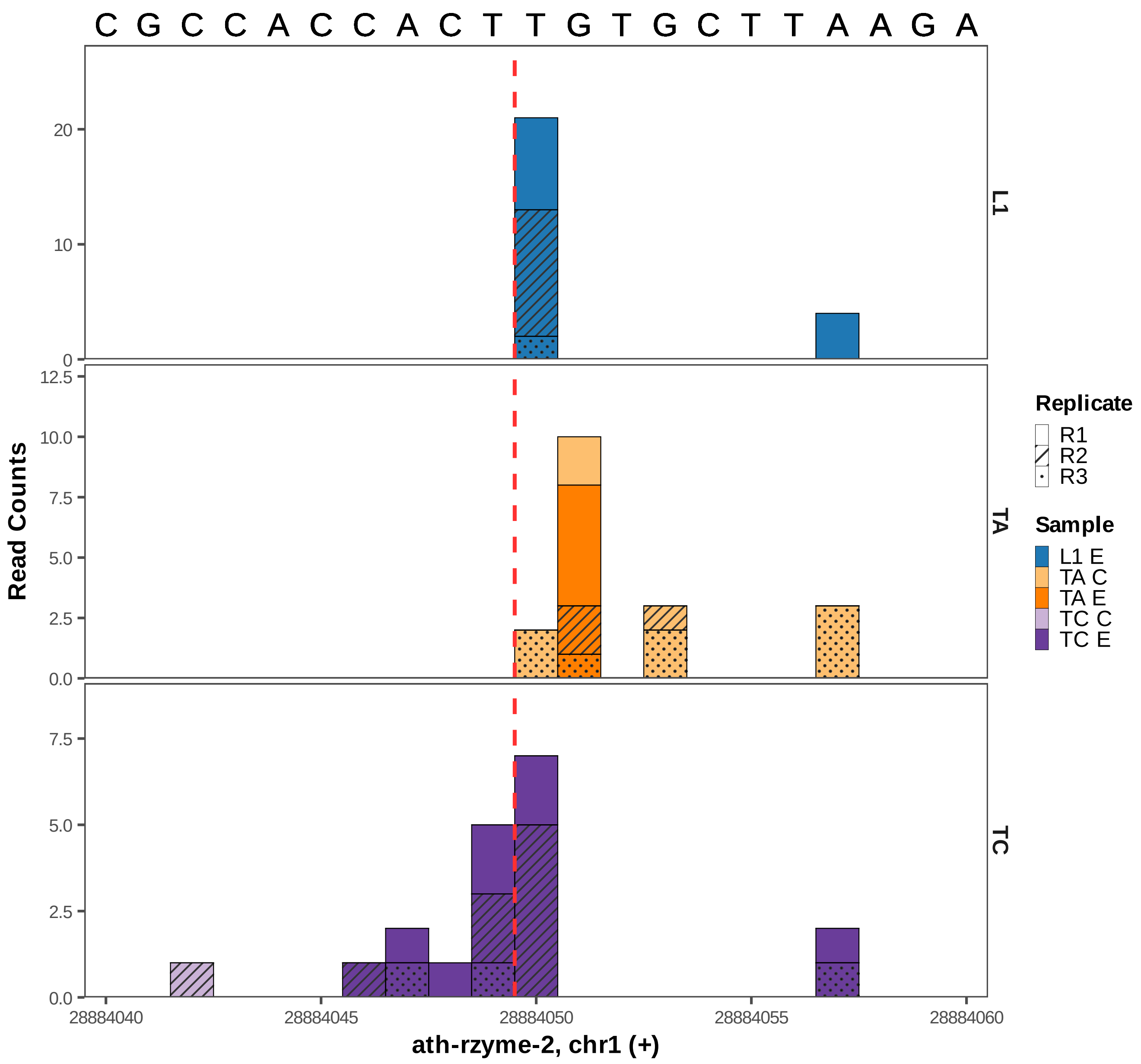

C G C C A C C A C T T G T G C T T A A G A
Read Counts
L1
TA
TC
Replicate
R1
R2
R3
Sample
L1 E
TA C
TA E
TC C
TC E
28884040
28884045
28884050
28884055
28884060
ath-rzyme-2, chr1 (+)

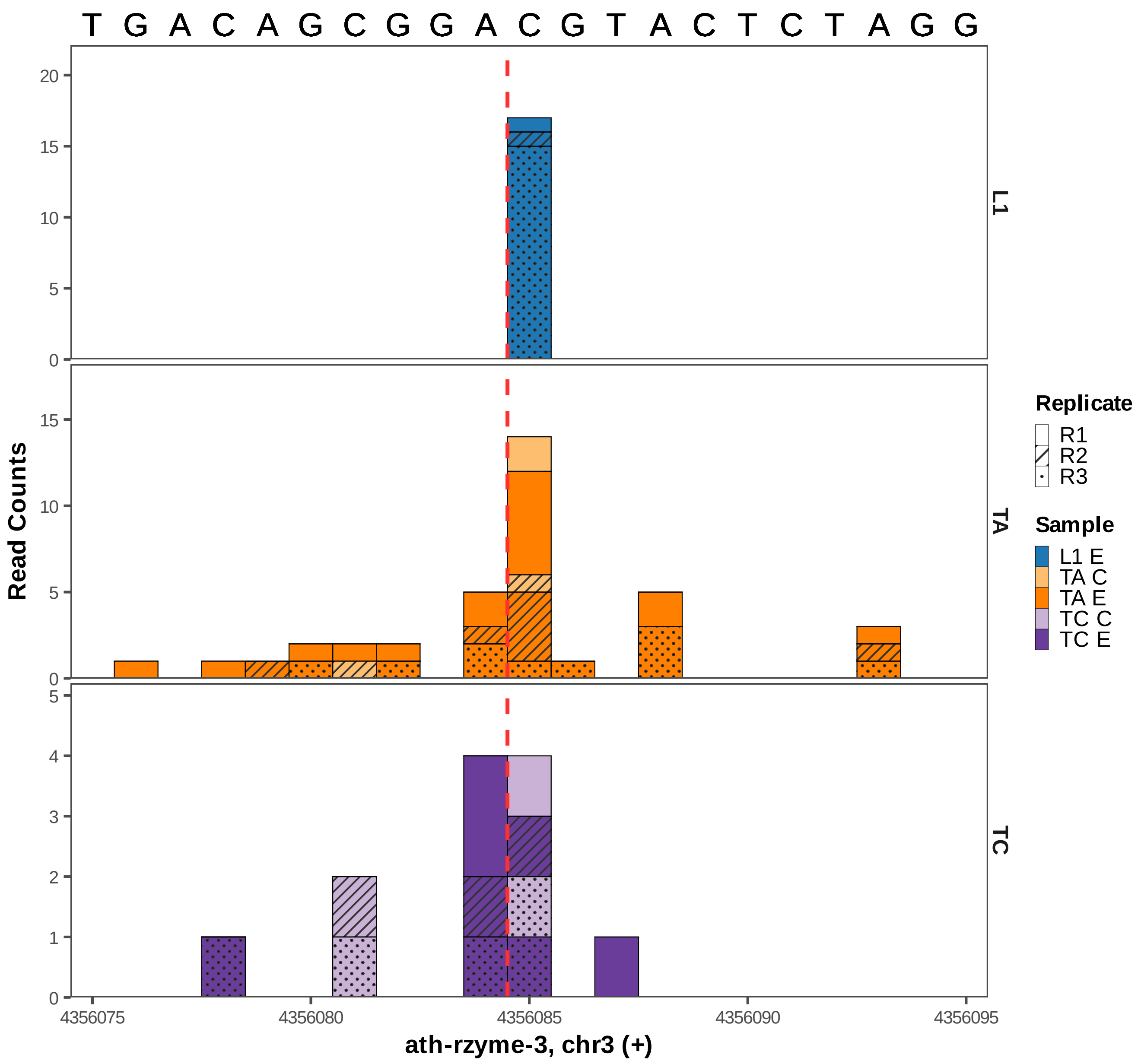

T G A C A G C G G A C G T A C T C T A G G
Read Counts
L1
TA
TC
Replicate
R1
R2
R3
Sample
L1 E
TA C
TA E
TC C
TC E
4356075
4356080
4356085
4356090
4356095
ath-rzyme-3, chr3 (+)

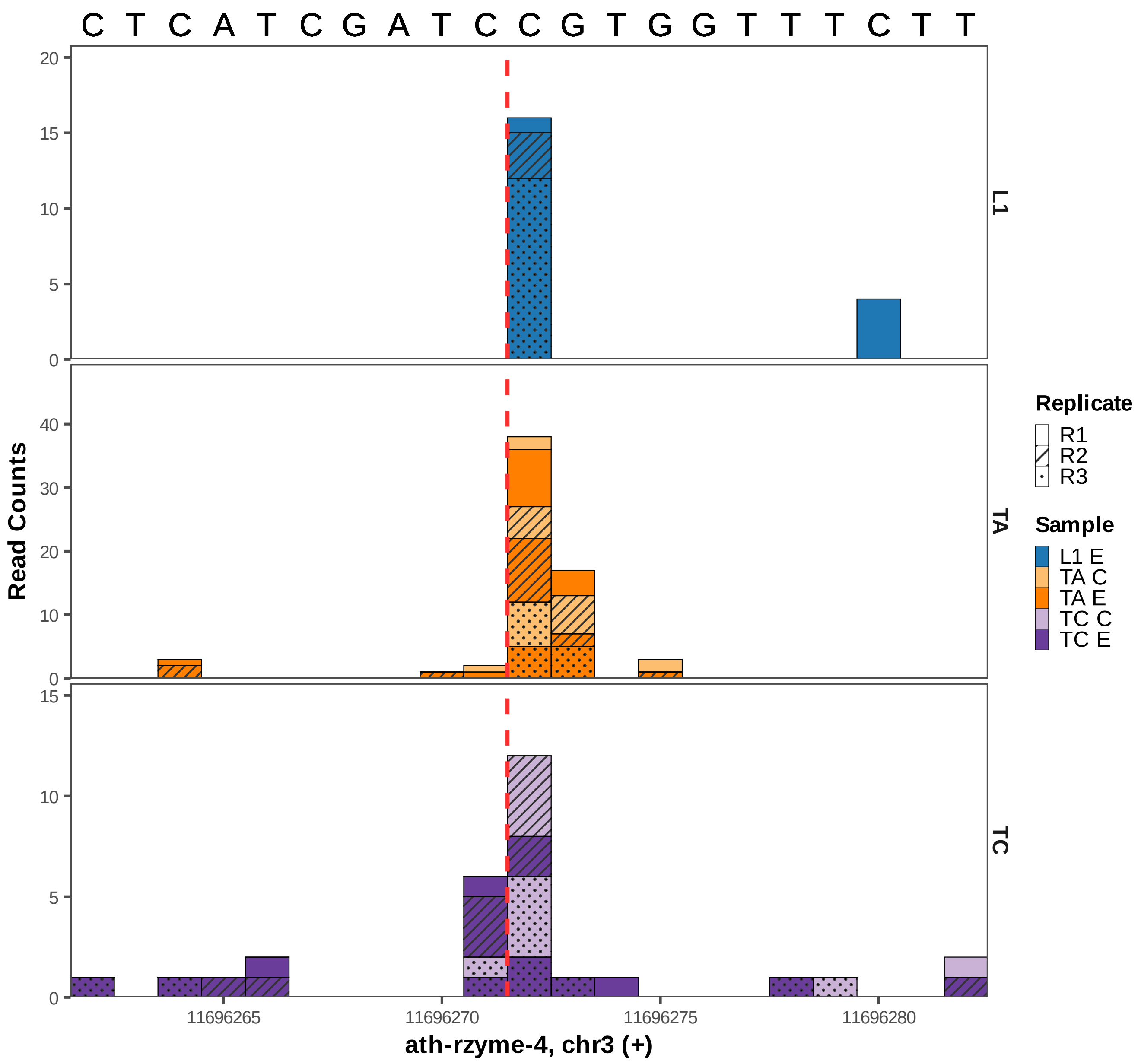

C T C A T C G A T C C G T G G T T T C T T
Read Counts
L1
TA
TC
0
5
10
15
20
0
10
20
30
40
0
5
10
15
11696265
11696270
11696275
11696280
ath-rzyme-4, chr3 (+)
Replicate
R1
R2
R3
Sample
L1 E
TA C
TA E
TC C
TC E

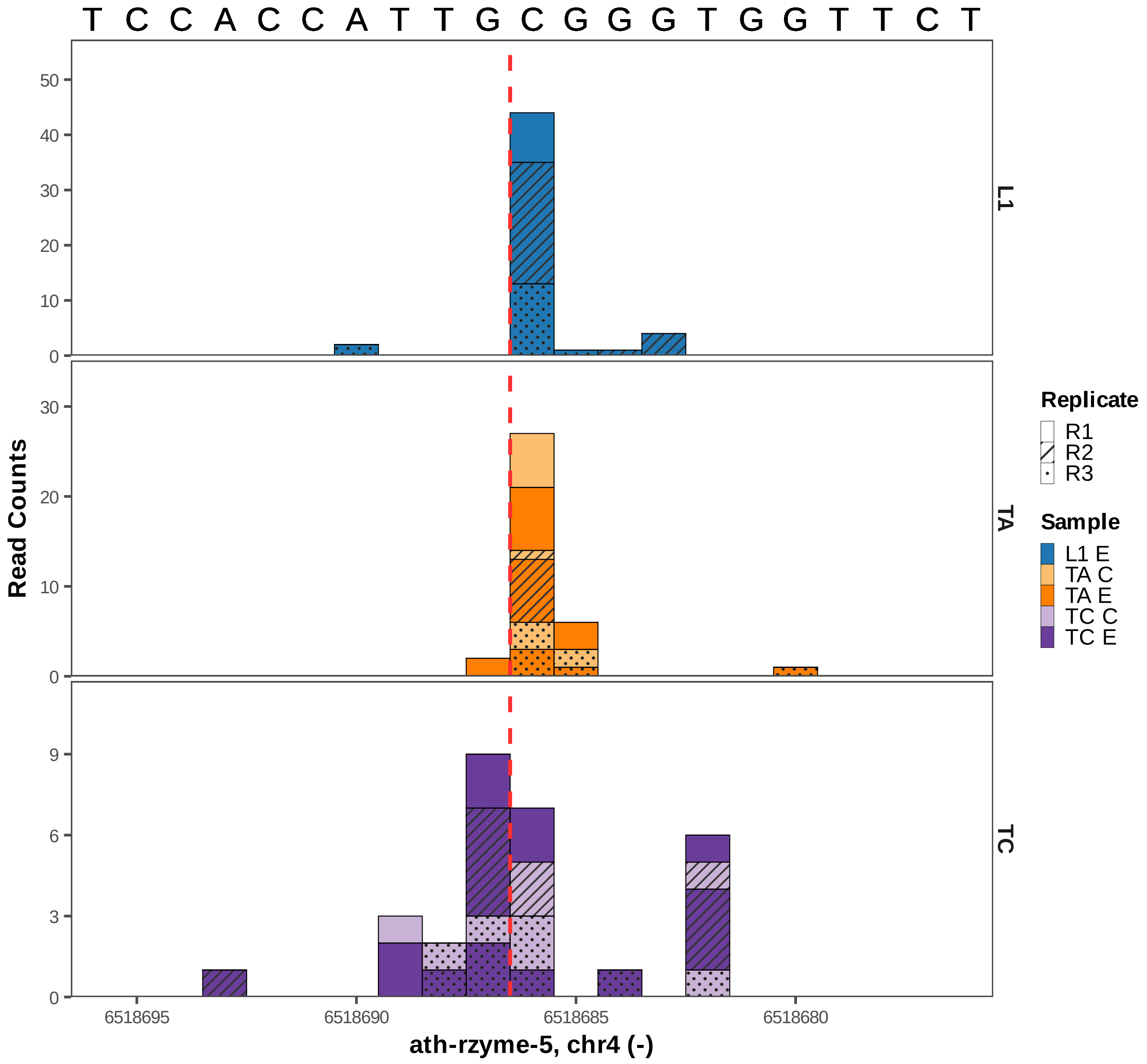

T C C A C C A T T G C G G G T G G T T C T
L1
TA
TC
Read Counts
Replicate
R1
R2
R3
Sample
L1 E
TA C
TA E
TC C
TC E
6518695
6518690
6518685
6518680
ath-rzyme-5, chr4 (-)

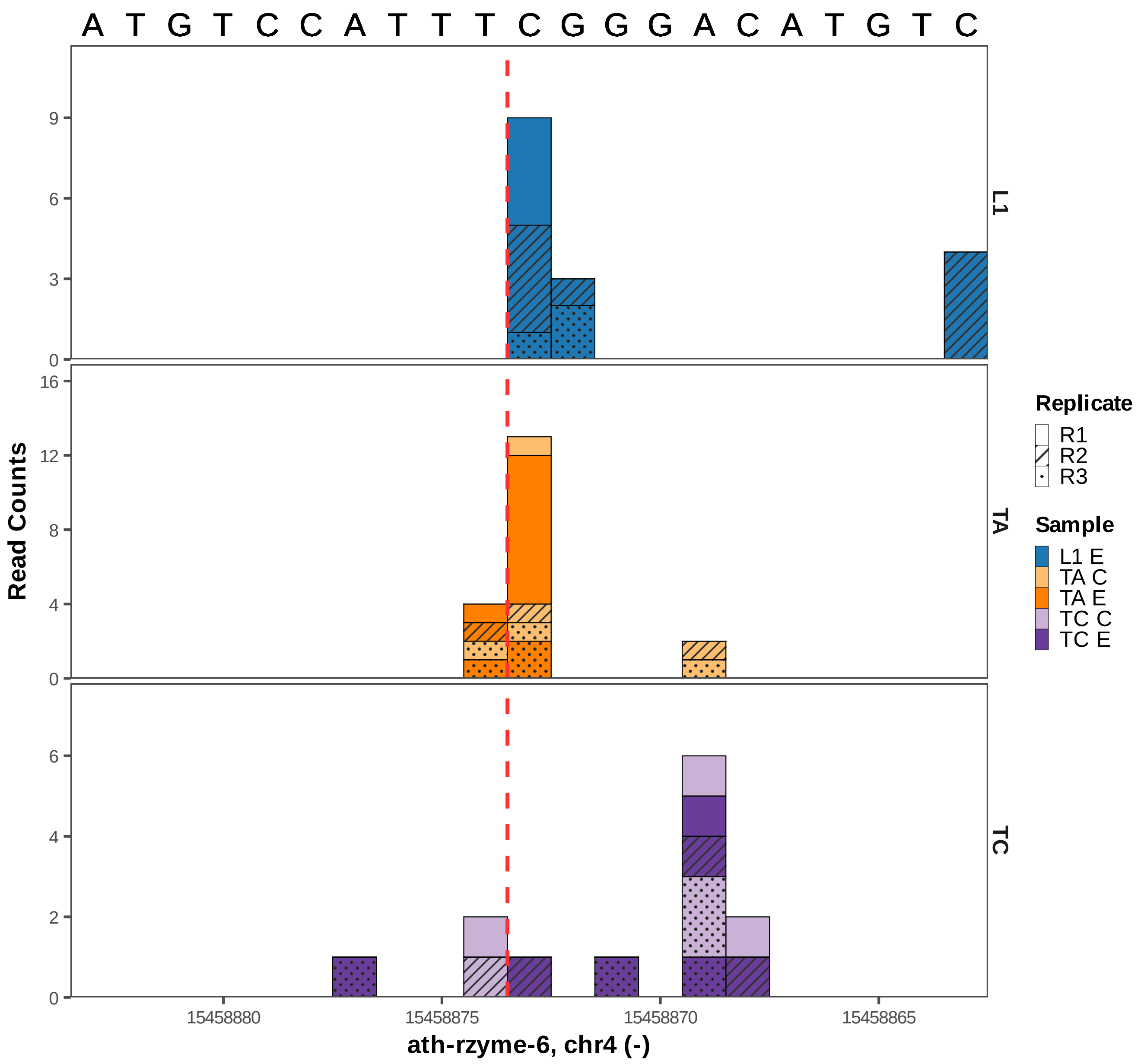

A T G T C C A T T T C G G G A C A T G T C
Read Counts
L1
TA
TC
0
3
6
9
0
4
8
12
16
0
2
4
6
15458880
15458875
15458870
15458865
ath-rzyme-6, chr4 (-)
Replicate
R1
R2
R3
Sample
L1 E
TA C
TA E
TC C
TC E

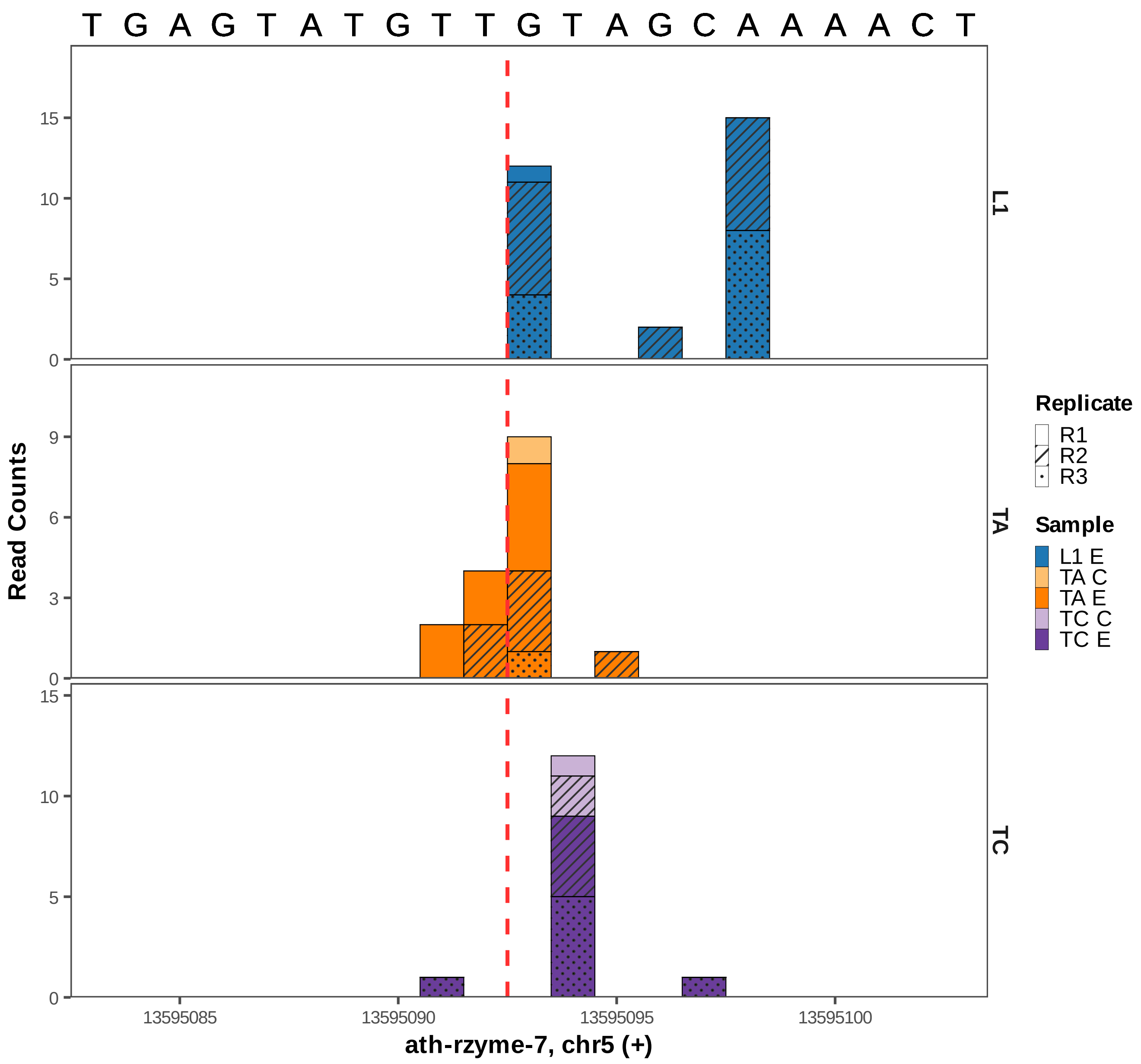

T G A G T A T G T T G T A G C A A A A C T
Read Counts
L1
TA
TC
15
10
5
0
9
6
3
0
15
10
5
0
13595085
13595090
13595095
13595100
ath-rzyme-7, chr5 (+)
Replicate
R1
R2
R3
Sample
L1 E
TA C
TA E
TC C
TC E

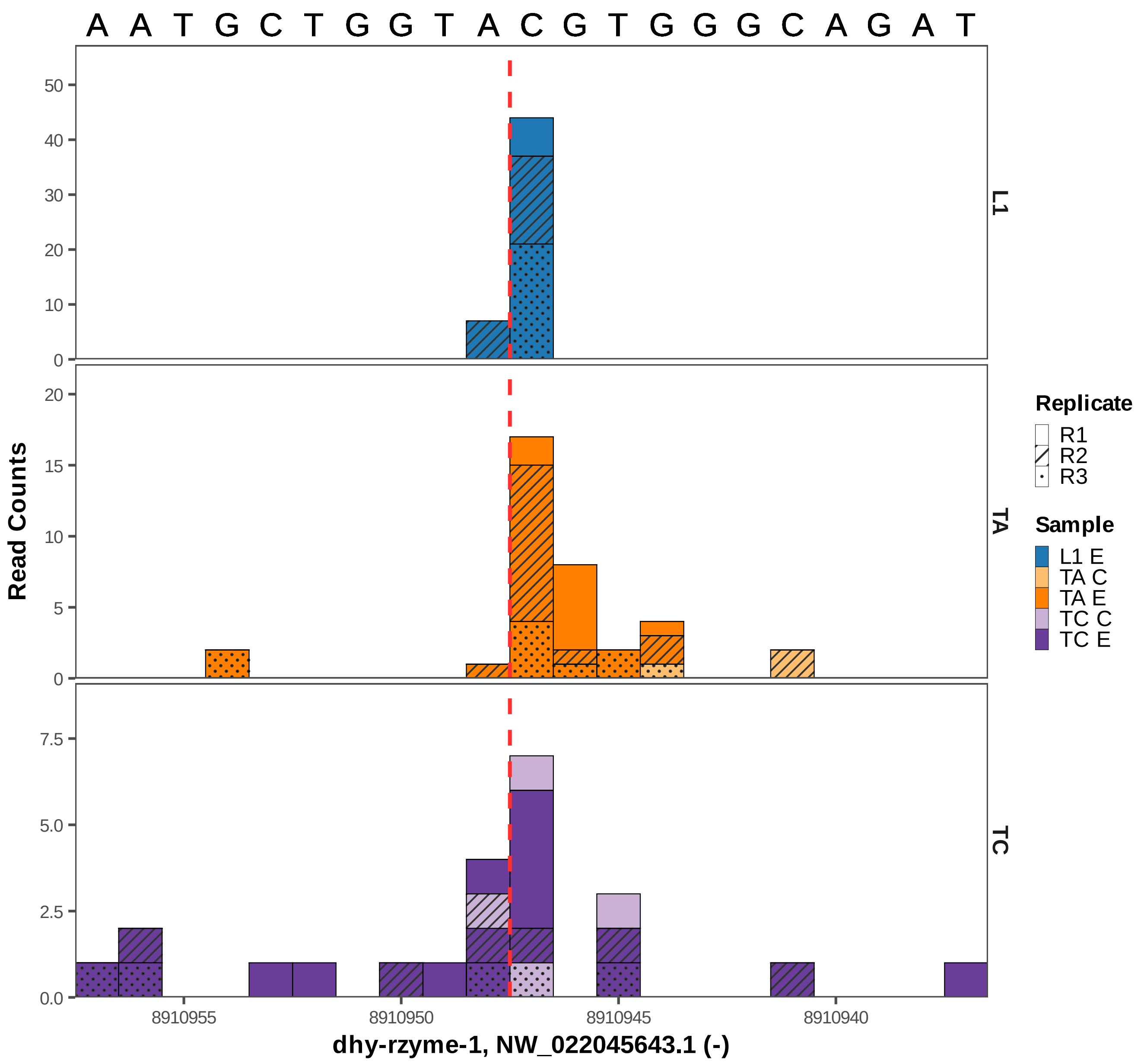

A A T G C T G G T A C G T G G G C A G A T
Read Counts
L1
TA
TC
Replicate
R1
R2
R3
Sample
L1 E
TA C
TA E
TC C
TC E
8910955
8910950
8910945
8910940
dhy-rzyme-1, NW_022045643.1 (-)

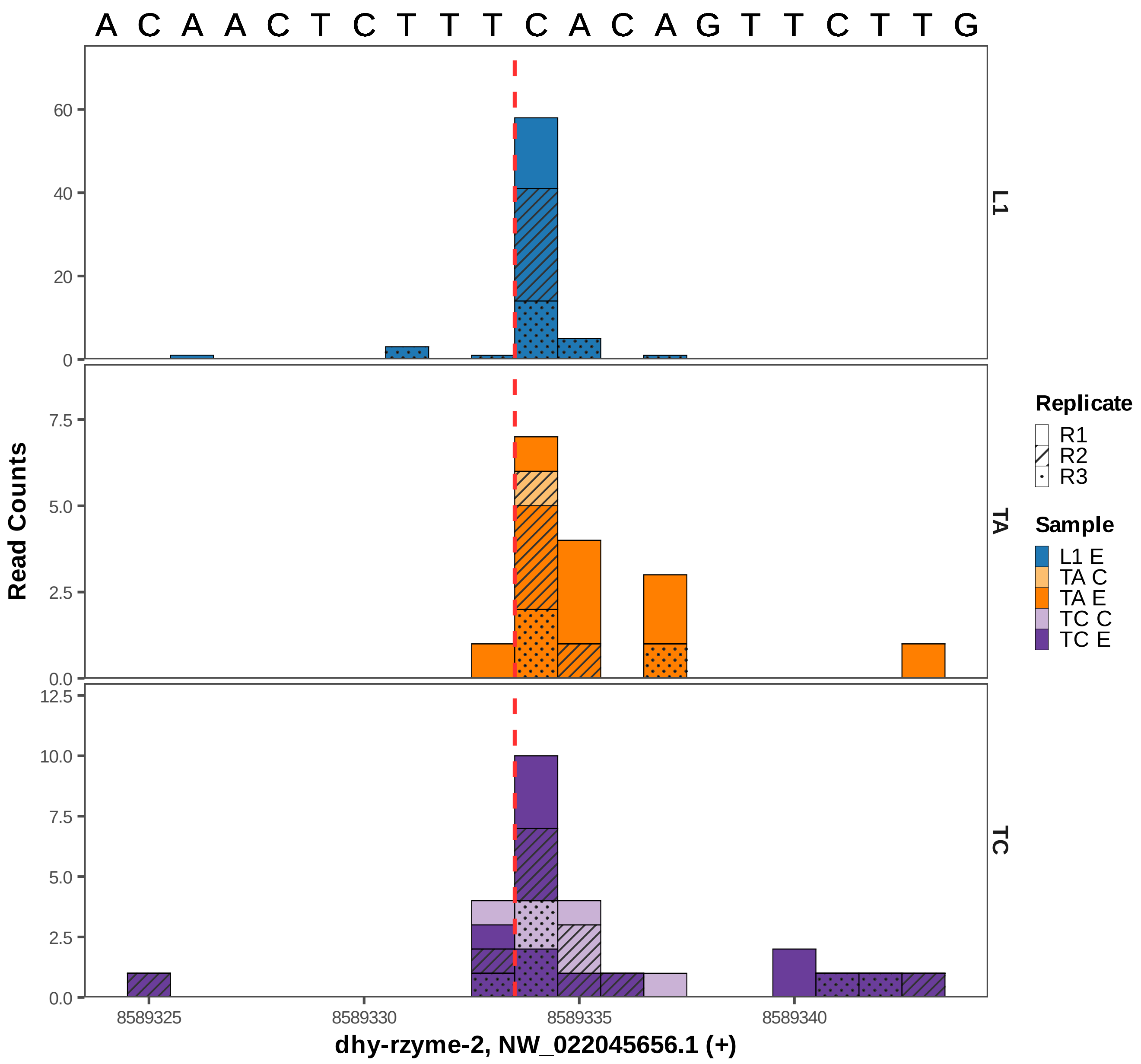

A C A A C T C T T T C A C A G T T C T T G
Read Counts
L1
TA
TC
Replicate
R1
R2
R3
Sample
L1 E
TA C
TA E
TC C
TC E
0
20
40
60
0.0
2.5
5.0
7.5
0.0
2.5
5.0
7.5
10.0
12.5
8589325
8589330
8589335
8589340
dhy-rzyme-2, NW_022045656.1 (+)

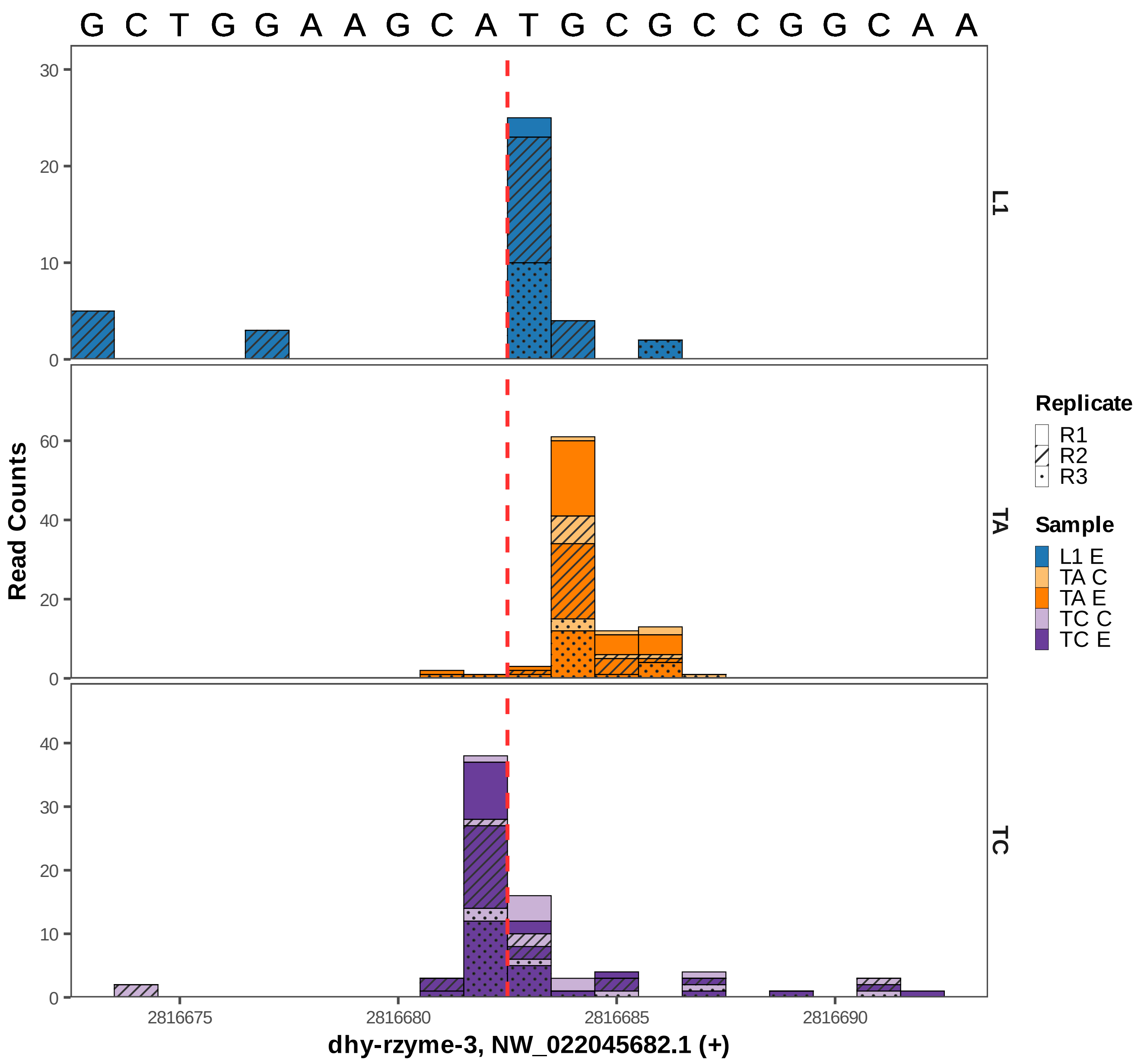

G C T G G A A G C A T G C G C C G G C A A
L1
TA
TC
Read Counts
Replicate
R1
R2
R3
Sample
L1 E
TA C
TA E
TC C
TC E
2816675
2816680
2816685
2816690
dhy-rzyme-3, NW_022045682.1 (+)

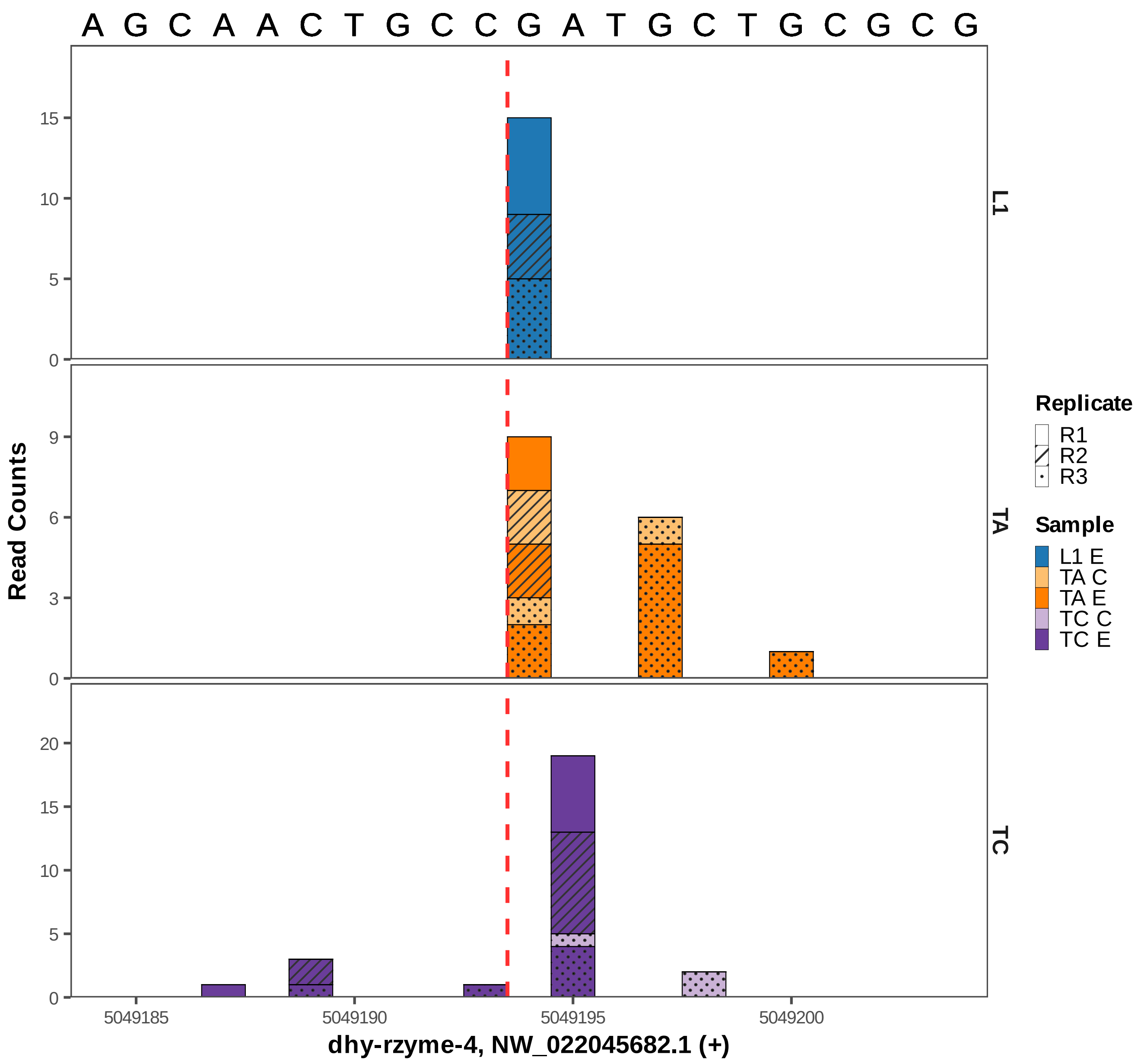

A G C A A C T G C C G A T G C T G C G C G
L1
TA
TC
Read Counts
0
5
10
15
0
3
6
9
0
5
10
15
20
5049185
5049190
5049195
5049200
dhy-rzyme-4, NW_022045682.1 (+)
Replicate
R1
R2
R3
Sample
L1 E
TA C
TA E
TC C
TC E

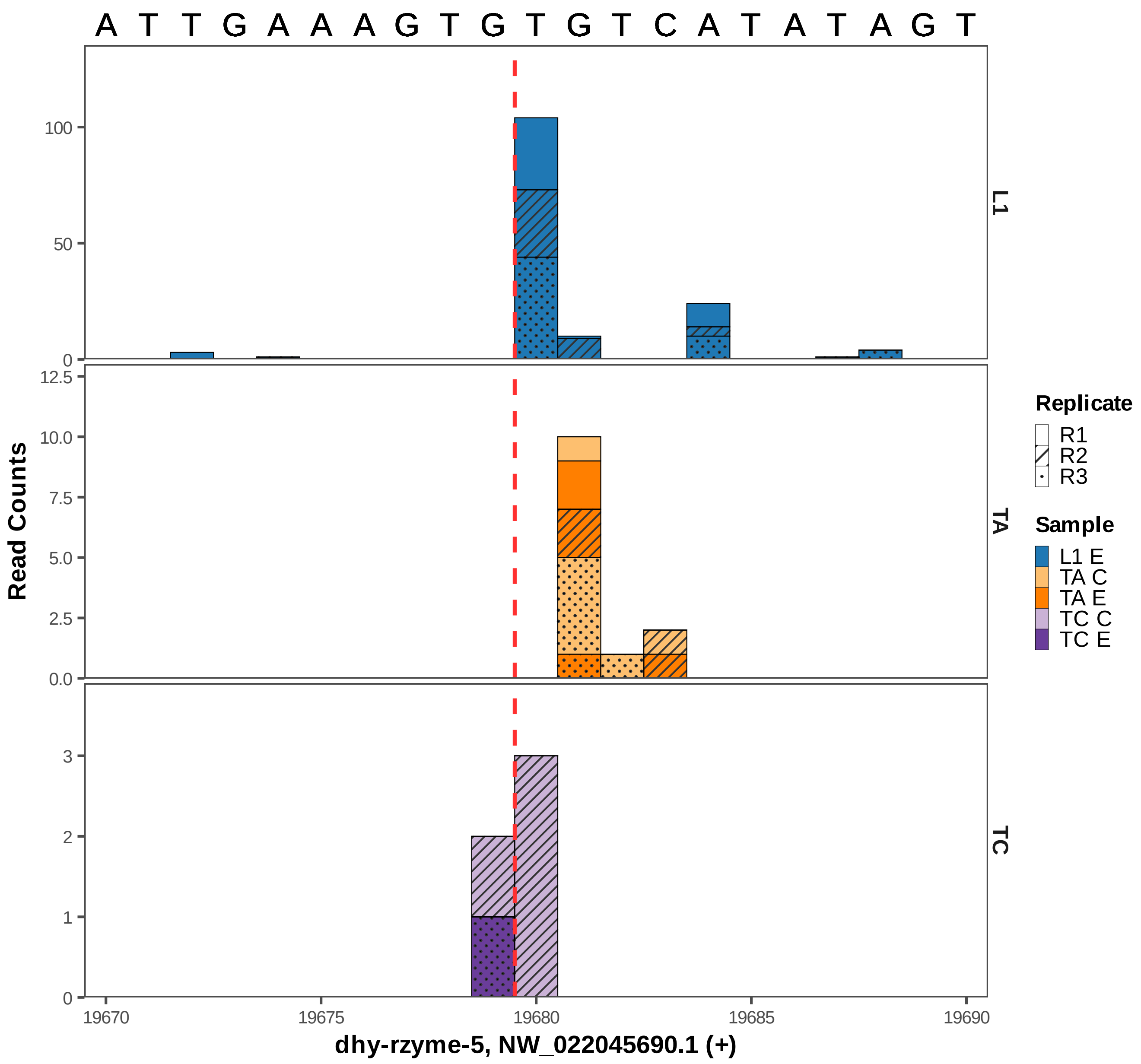

A T T G A A A G T G T G T C A T A T A G T
L1
TA
TC
Read Counts
0
50
100
0.0
2.5
5.0
7.5
10.0
12.5
0
1
2
3
19670
19675
19680
19685
19690
dhy-rzyme-5, NW_022045690.1 (+)
Replicate
R1
R2
R3
Sample
L1 E
TA C
TA E
TC C
TC E

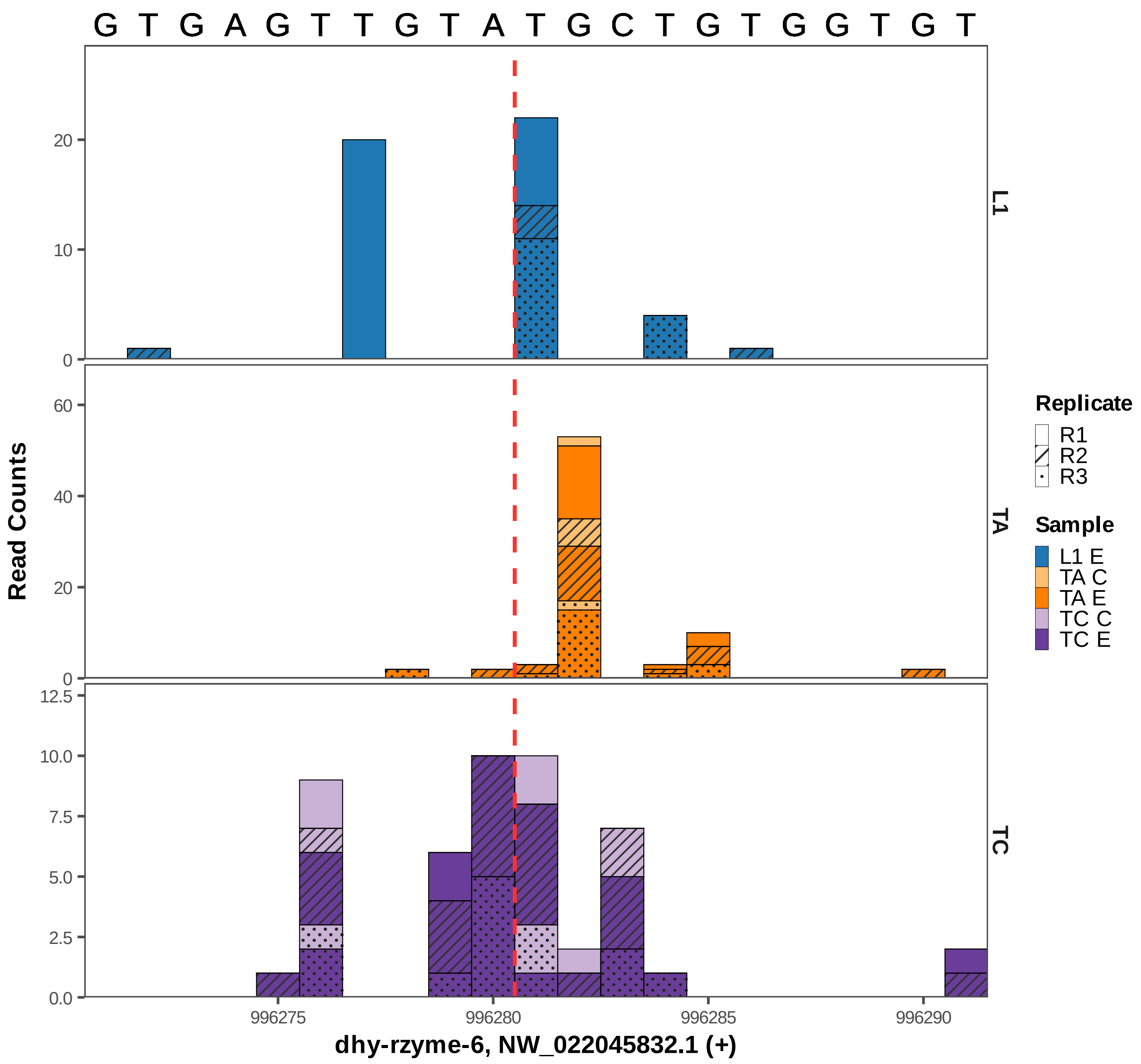

G T G A G T T G T A T G C T G T G G T G T
Read Counts
L1
TA
TC
0
10
20
0
20
40
60
0.0
2.5
5.0
7.5
10.0
12.5
996275
996280
996285
996290
dhy-rzyme-6, NW_022045832.1 (+)
Replicate
R1
R2
R3
Sample
L1 E
TA C
TA E
TC C
TC E

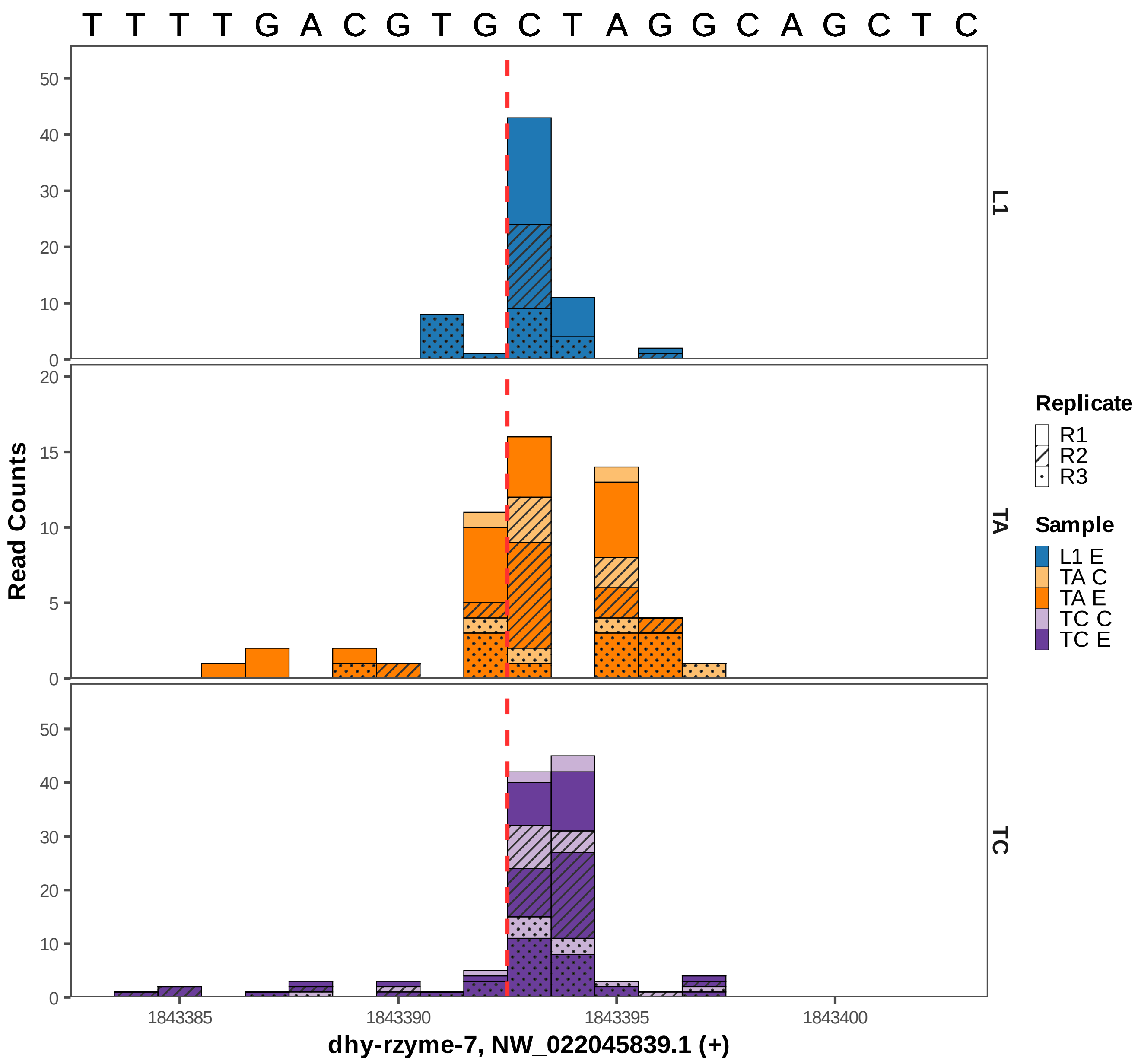

T T T T G A C G T G C T A G G C A G C T C
L1
TA
TC
Read Counts
0
10
20
30
40
50
5
15
1843385
1843390
1843395
1843400
dhy-rzyme-7, NW_022045839.1 (+)
Replicate
R1
R2
R3
Sample
L1 E
TA C
TA E
TC C
TC E

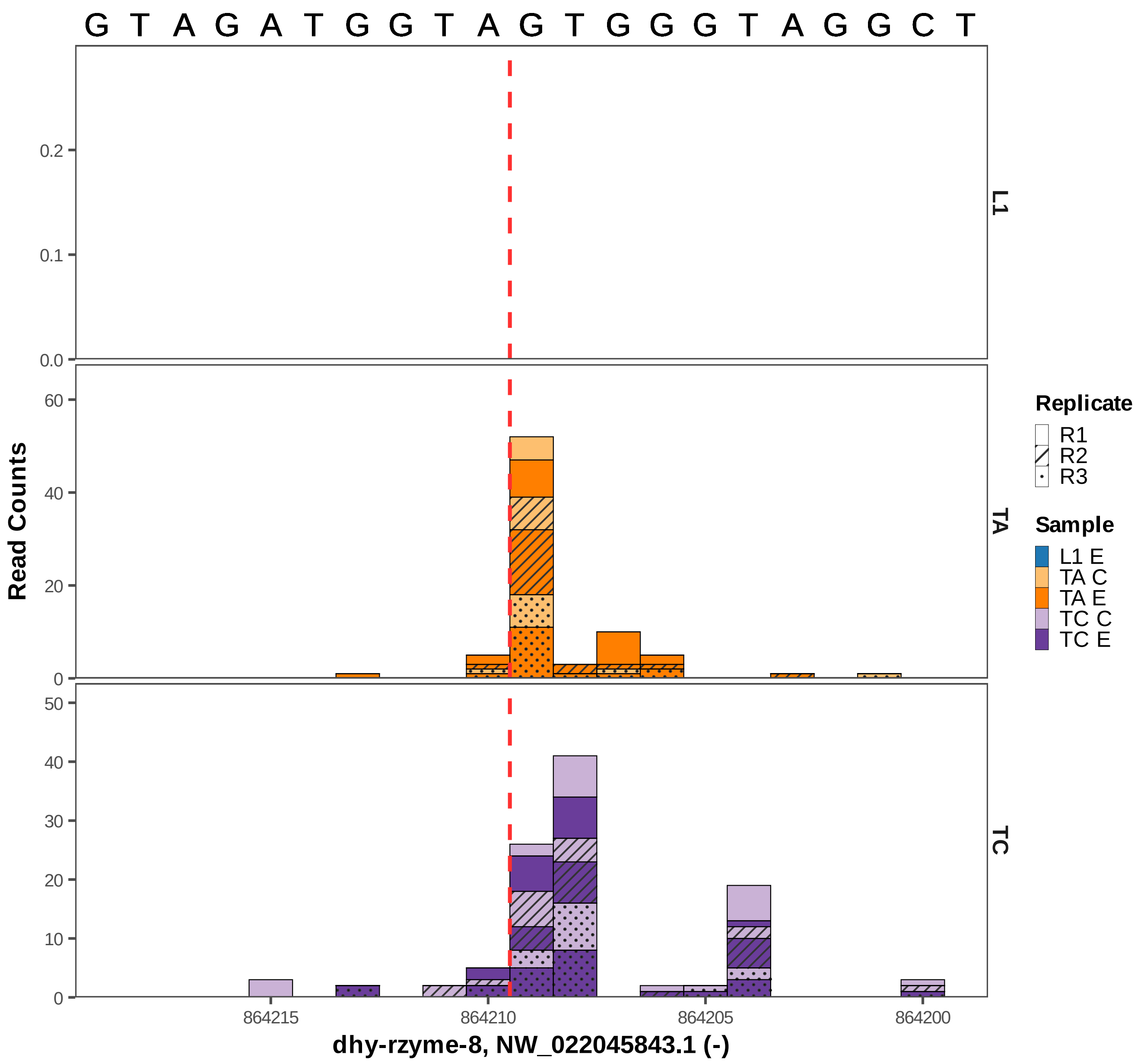

G T A G A T G G T A G T G G G T A G G C T
L1
TA
TC
Read Counts
0.0
0.1
0.2
0
20
40
60
0
10
20
30
40
50
864215
864210
864205
864200
dhy-rzyme-8, NW_022045843.1 (-)
Replicate
R1
R2
R3
Sample
L1 E
TA C
TA E
TC C
TC E

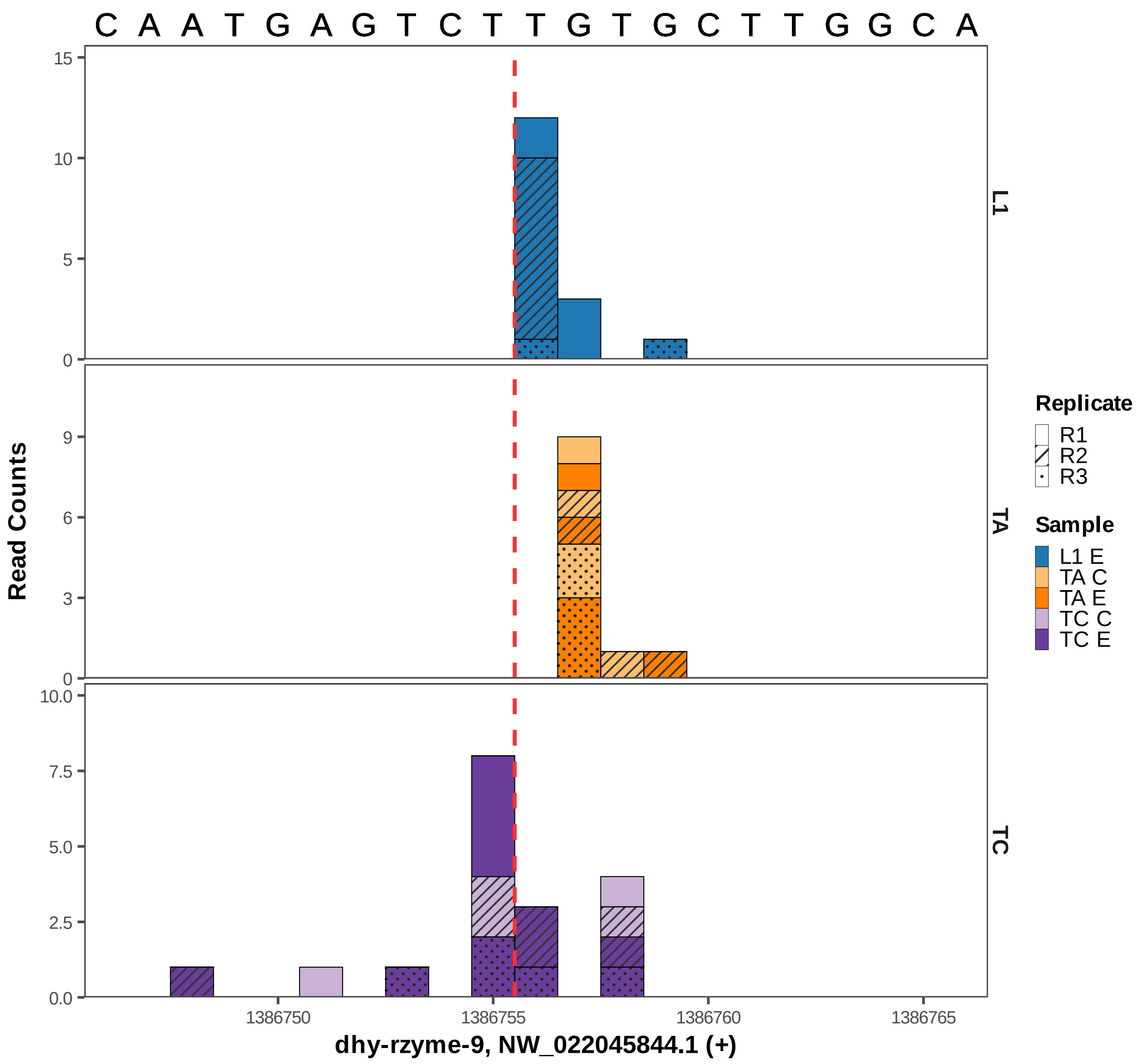

C A A T G A G T C T T G T G C T T G G C A
Read Counts
L1
TA
TC
Replicate
R1
R2
R3
Sample
L1 E
TA C
TA E
TC C
TC E
1386750
1386755
1386760
1386765
dhy-rzyme-9, NW_022045844.1 (+)

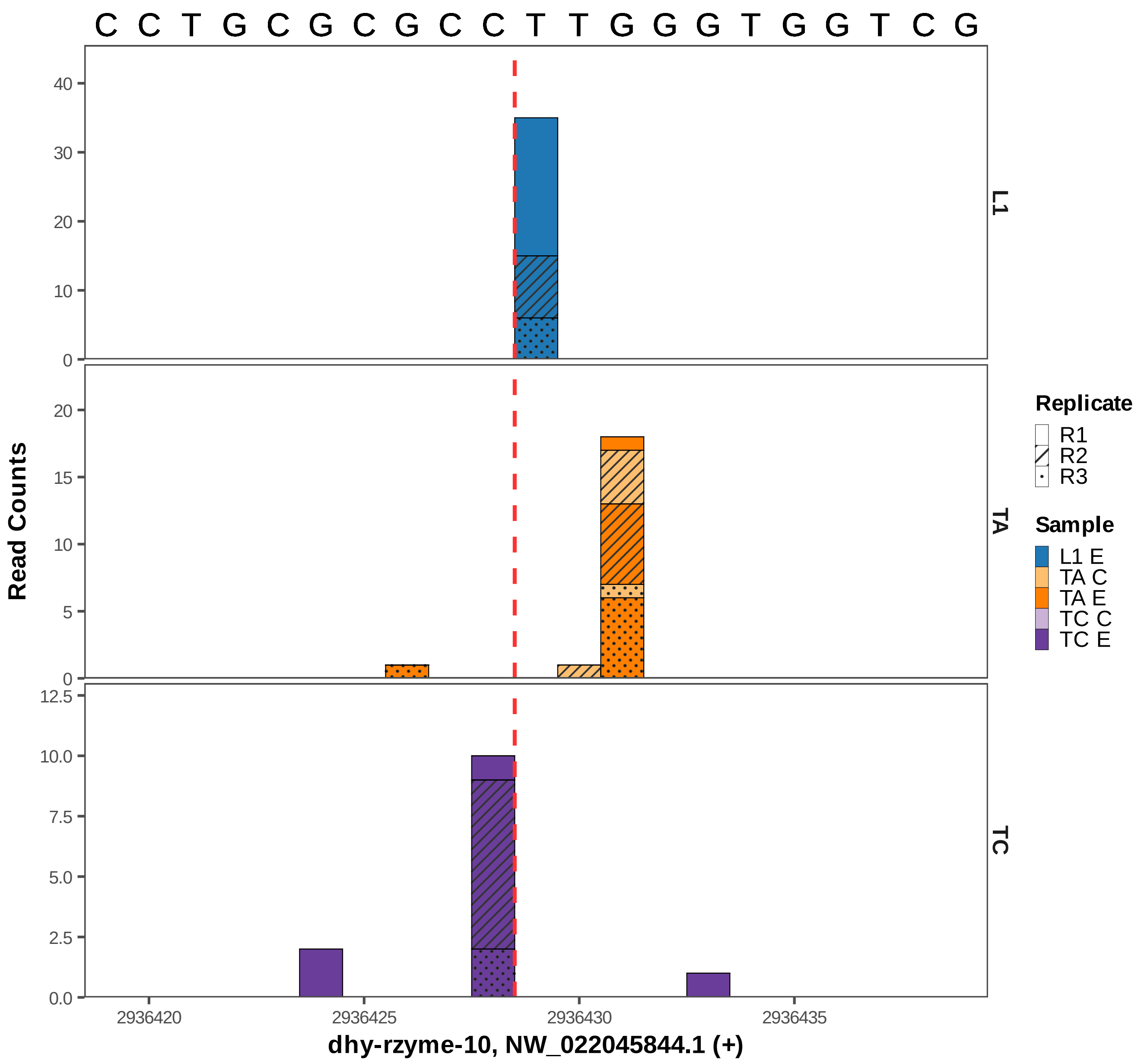

C C T G C G C G C C T T G G G T G G T C G
Read Counts
L1
TA
TC
Replicate
R1
R2
R3
Sample
L1 E
TA C
TA E
TC C
TC E
2936420
2936425
2936430
2936435
dhy-rzyme-10, NW_022045844.1 (+)

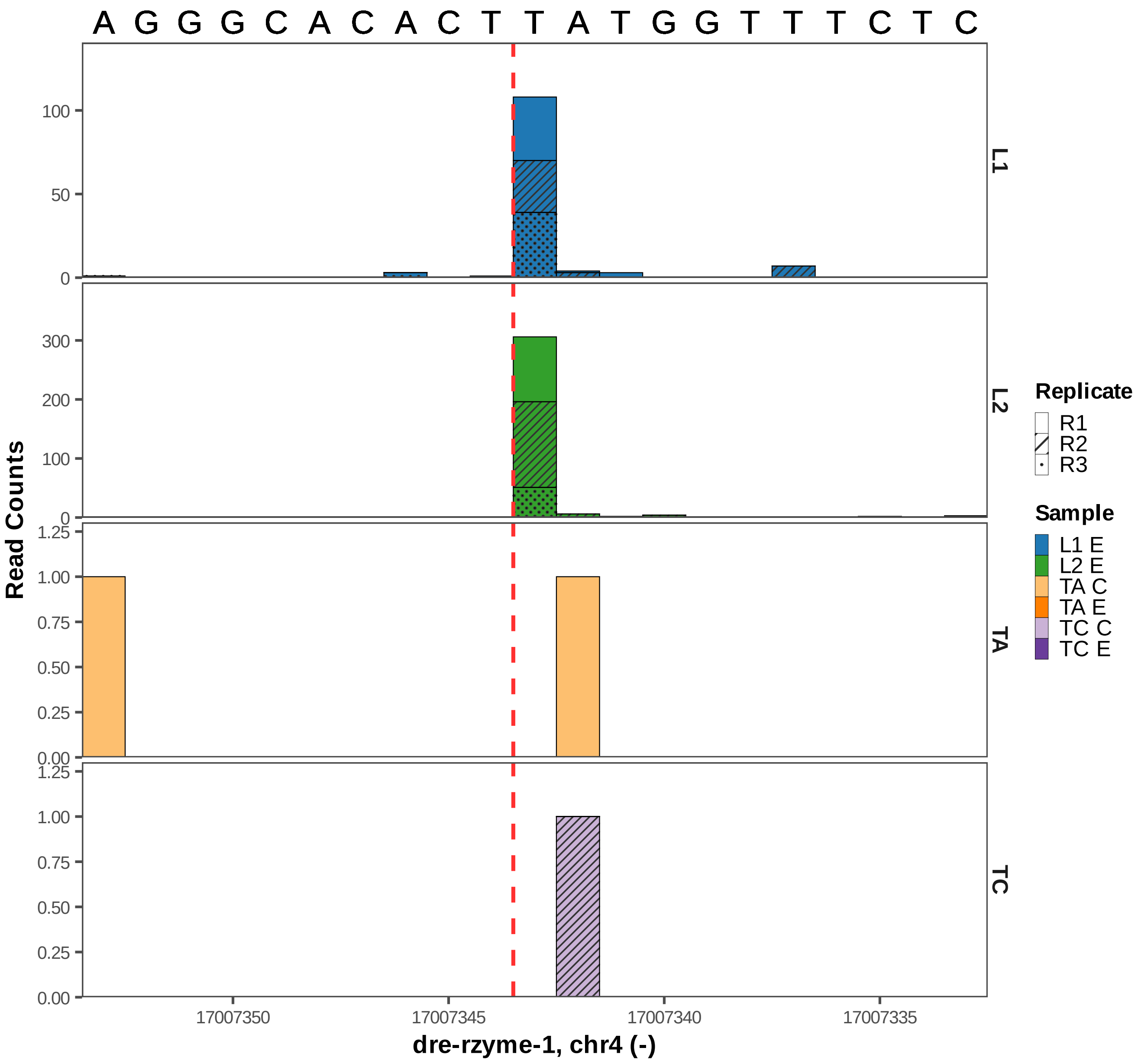

A G G G C A C A C T T A T G G T T T C T C
L1
L2
TA
TC
Read Counts
100
50
0
300
200
100
0
1.25
1.00
0.75
0.50
0.25
0.00
17007350
17007345
17007340
17007335
dre-rzyme-1, chr4 (-)
Replicate
R1
R2
R3
Sample
L1 E
L2 E
TA C
TA E
TC C
TC E

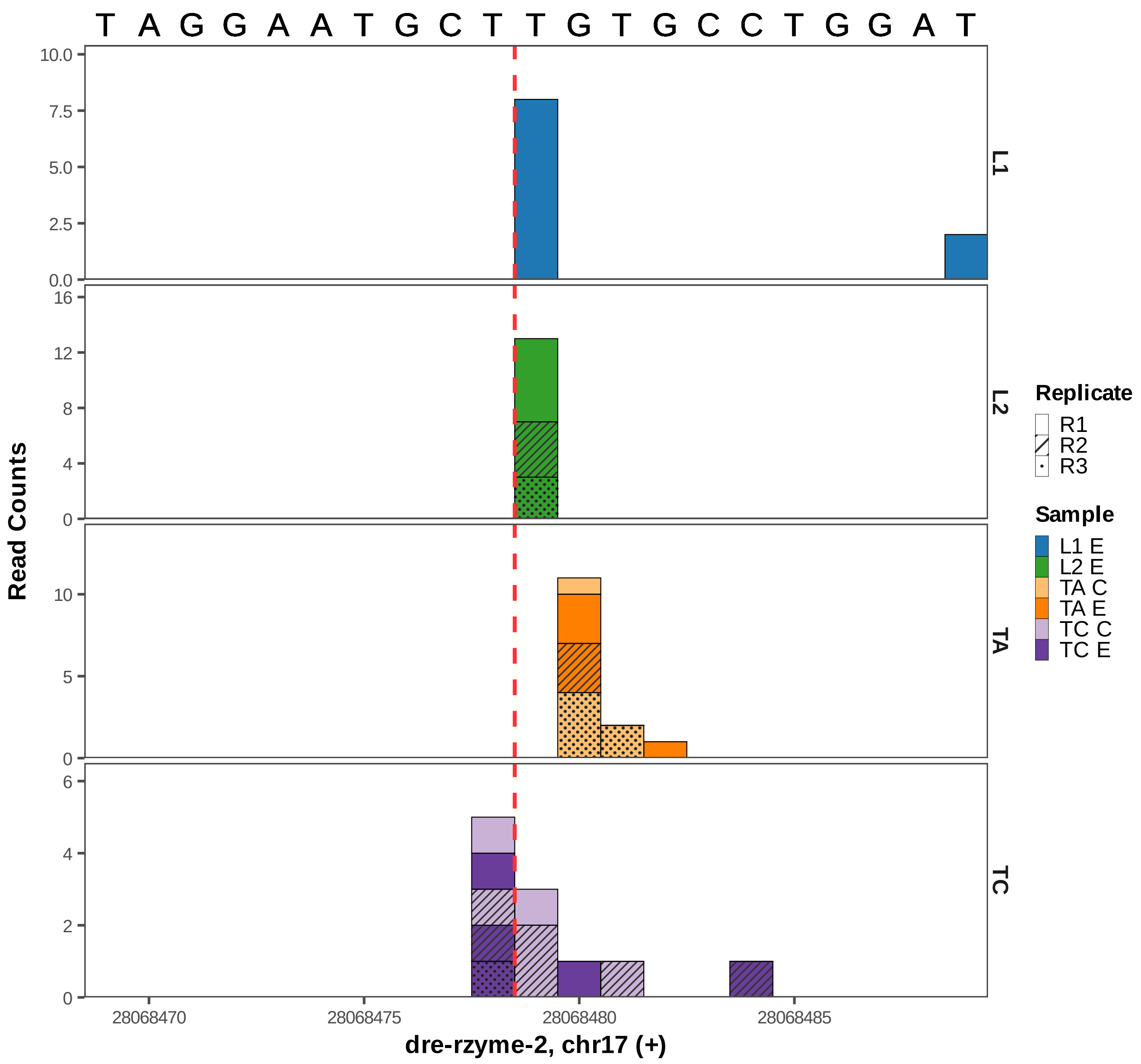

T A G G A A T G C T T G T G C C T G G A T
Read Counts
L1
L2
TA
TC
10.0
7.5
5.0
2.5
0.0
16
12
8
4
0
10
5
0
6
4
2
0
28068470
28068475
28068480
28068485
dre-rzyme-2, chr17 (+)
Replicate
R1
R2
R3
Sample
L1 E
L2 E
TA C
TA E
TC C
TC E

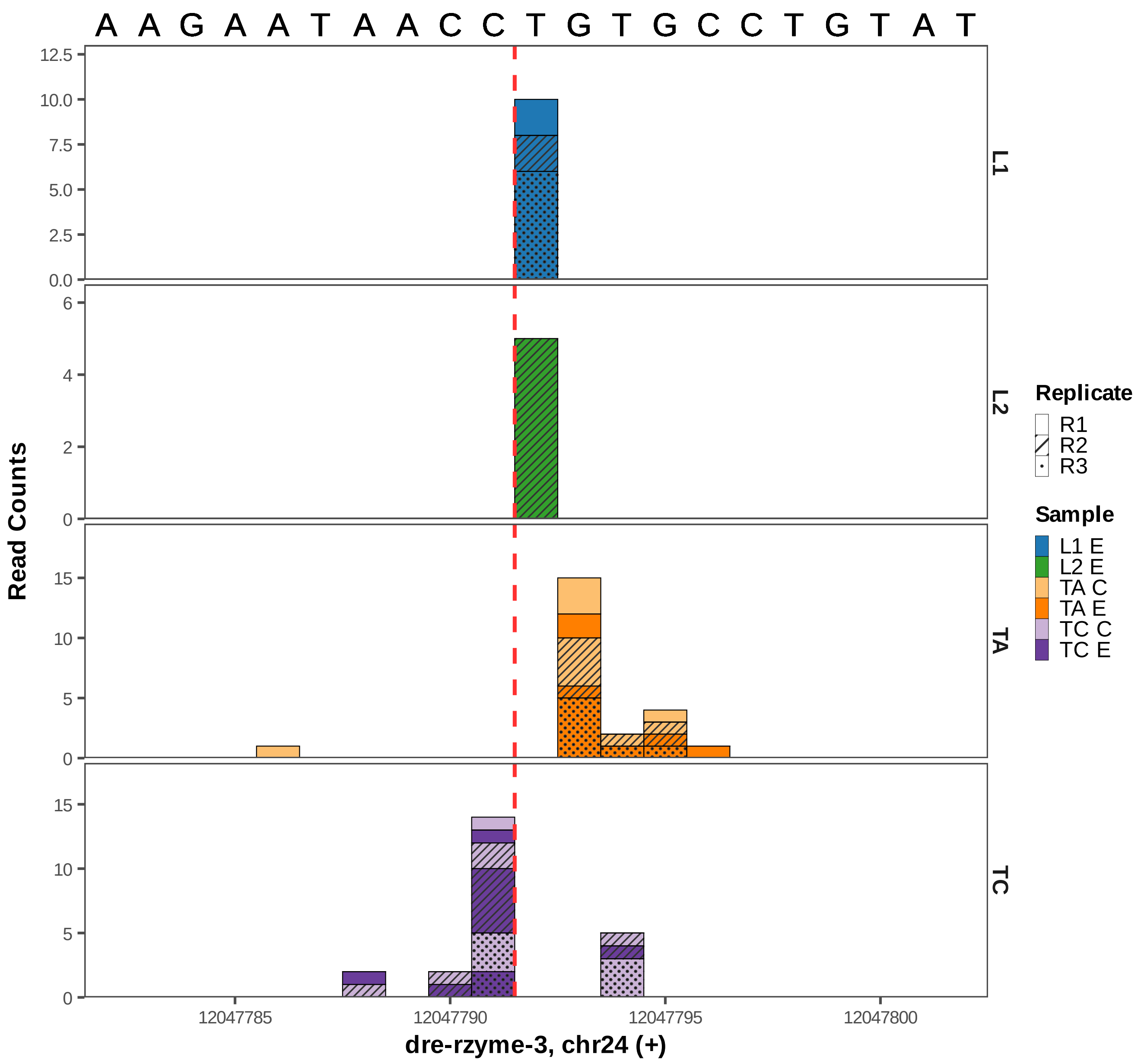
A A G A A T A A C C T G T G C C T G T A T
Read Counts
L1
L2
TA
TC
12.5
10.0
7.5
5.0
2.5
0.0
6
4
2
0
15
10
5
0
15
10
5
0
12047785
12047790
12047795
12047800
dre-rzyme-3, chr24 (+)
Replicate
R1
R2
R3
Sample
L1 E
L2 E
TA C
TA E
TC C
TC E

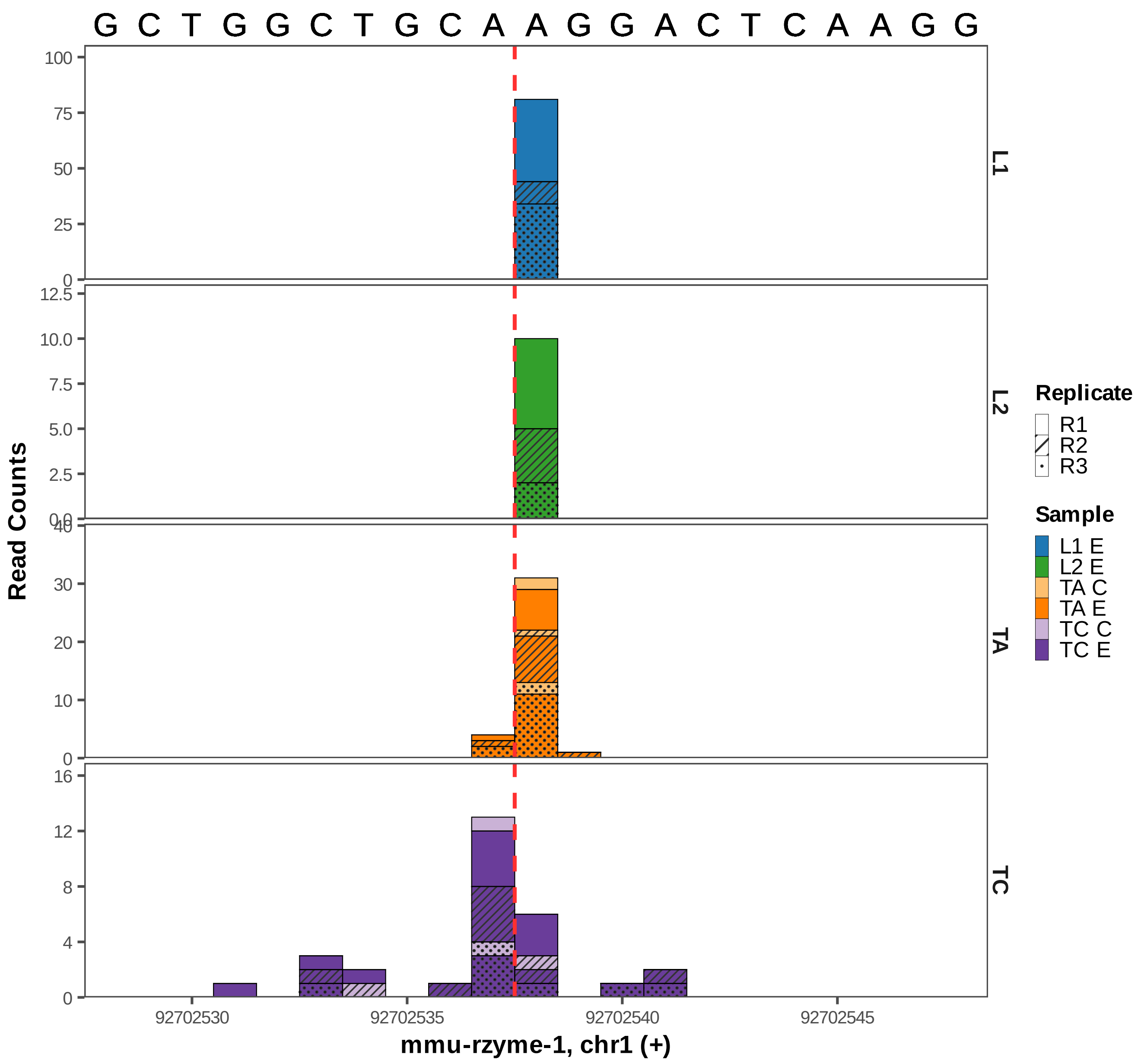

G C T G G C T G C A A G G A C T C A A G G
Read Counts
L1
L2
TA
TC
92702530
92702535
92702540
92702545
mmu-rzyme-1, chr1 (+)
Replicate
R1
R2
R3
Sample
L1 E
L2 E
TA C
TA E
TC C
TC E

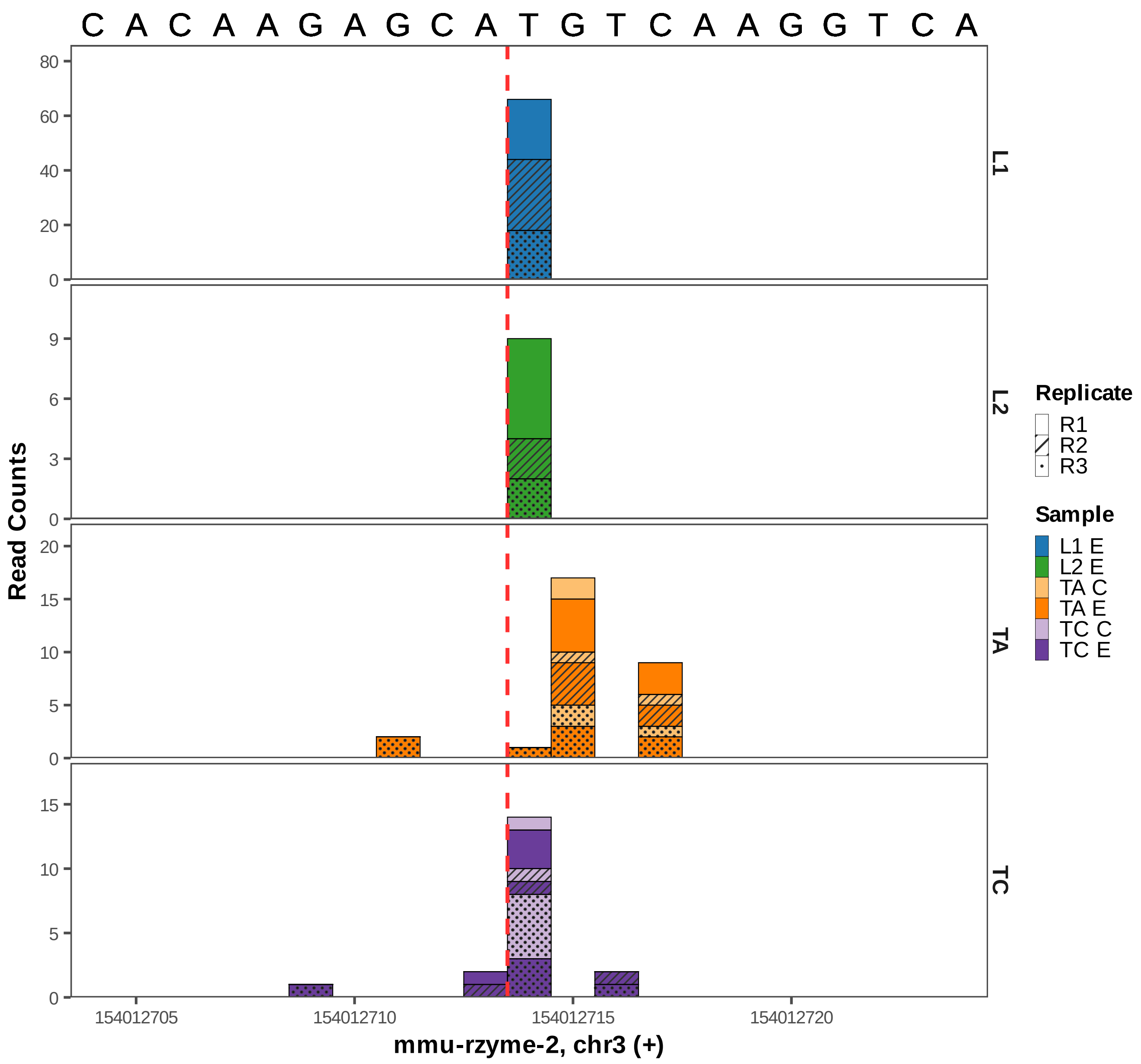
C A C A A G A G C A T G T C A A G G T C A
Read Counts
L1
L2
TA
TC
154012705
154012710
154012715
154012720
mmu-rzyme-2, chr3 (+)
Replicate
R1
R2
R3
Sample
L1 E
L2 E
TA C
TA E
TC C
TC E

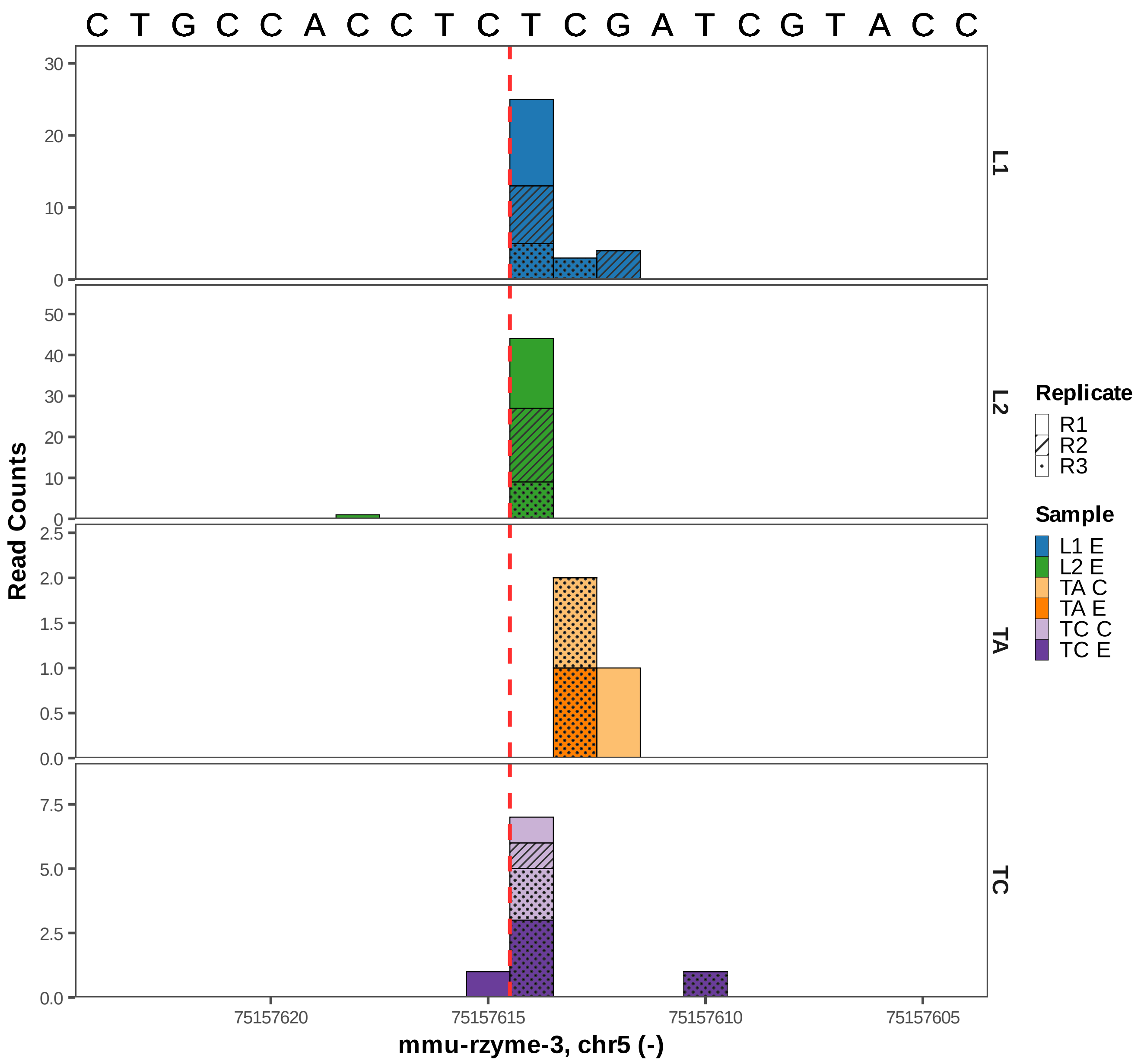

C T G C C A C C T C T C G A T C G T A C C
Read Counts
L1
L2
TA
TC
75157620
75157615
75157610
75157605
mmu-rzyme-3, chr5 (-)
Replicate
R1
R2
R3
Sample
L1 E
L2 E
TA C
TA E
TC C
TC E

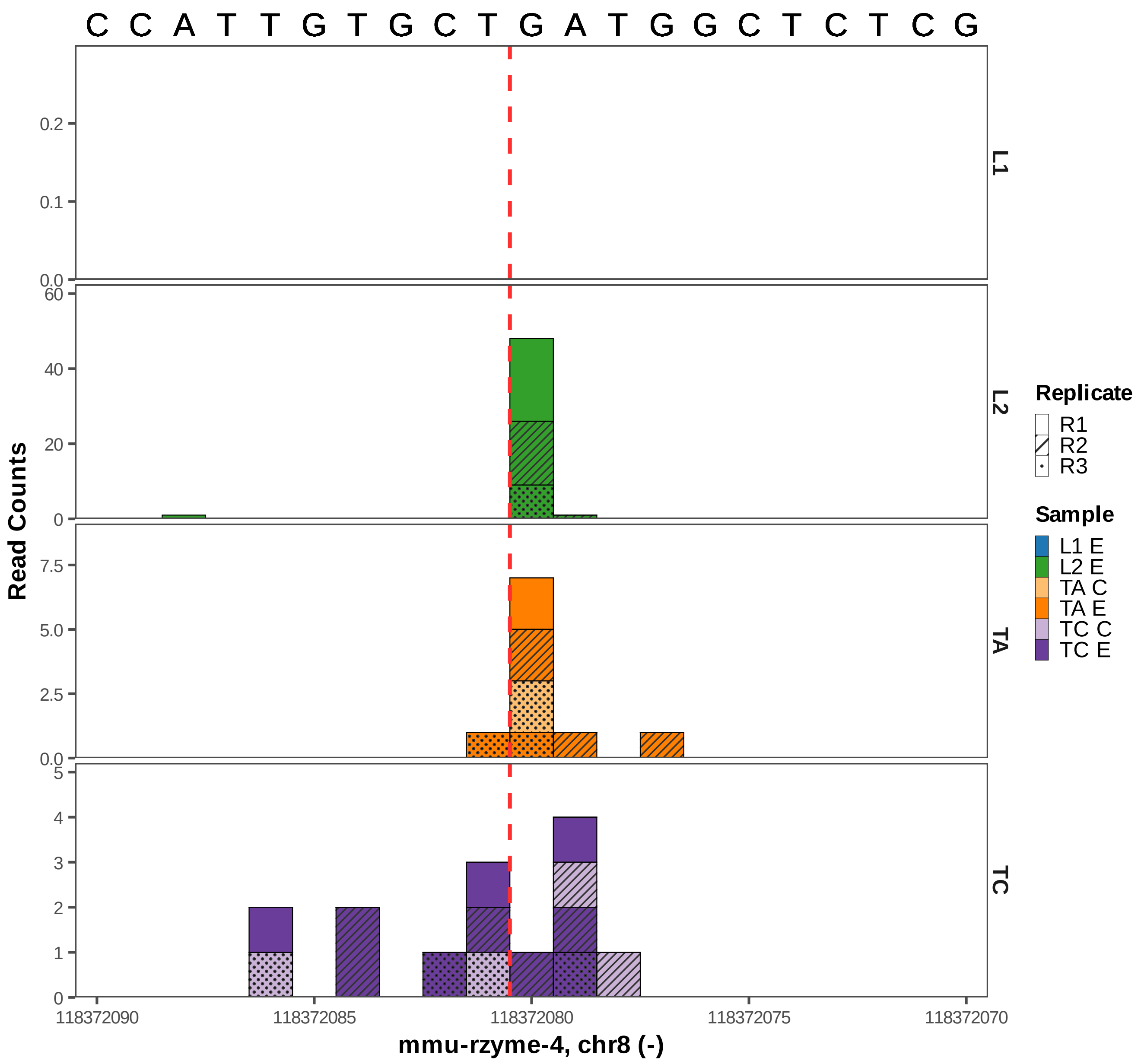

C C A T T G T G C T G A T G G C T C T C G
L1
L2
TA
TC
Read Counts
0.0
0.1
0.2
0
20
40
60
2.5
5.0
7.5
1
2
3
4
5
118372090
118372085
118372080
118372075
118372070
mmu-rzyme-4, chr8 (-)
Replicate
R1
R2
R3
Sample
L1 E
L2 E
TA C
TA E
TC C
TC E

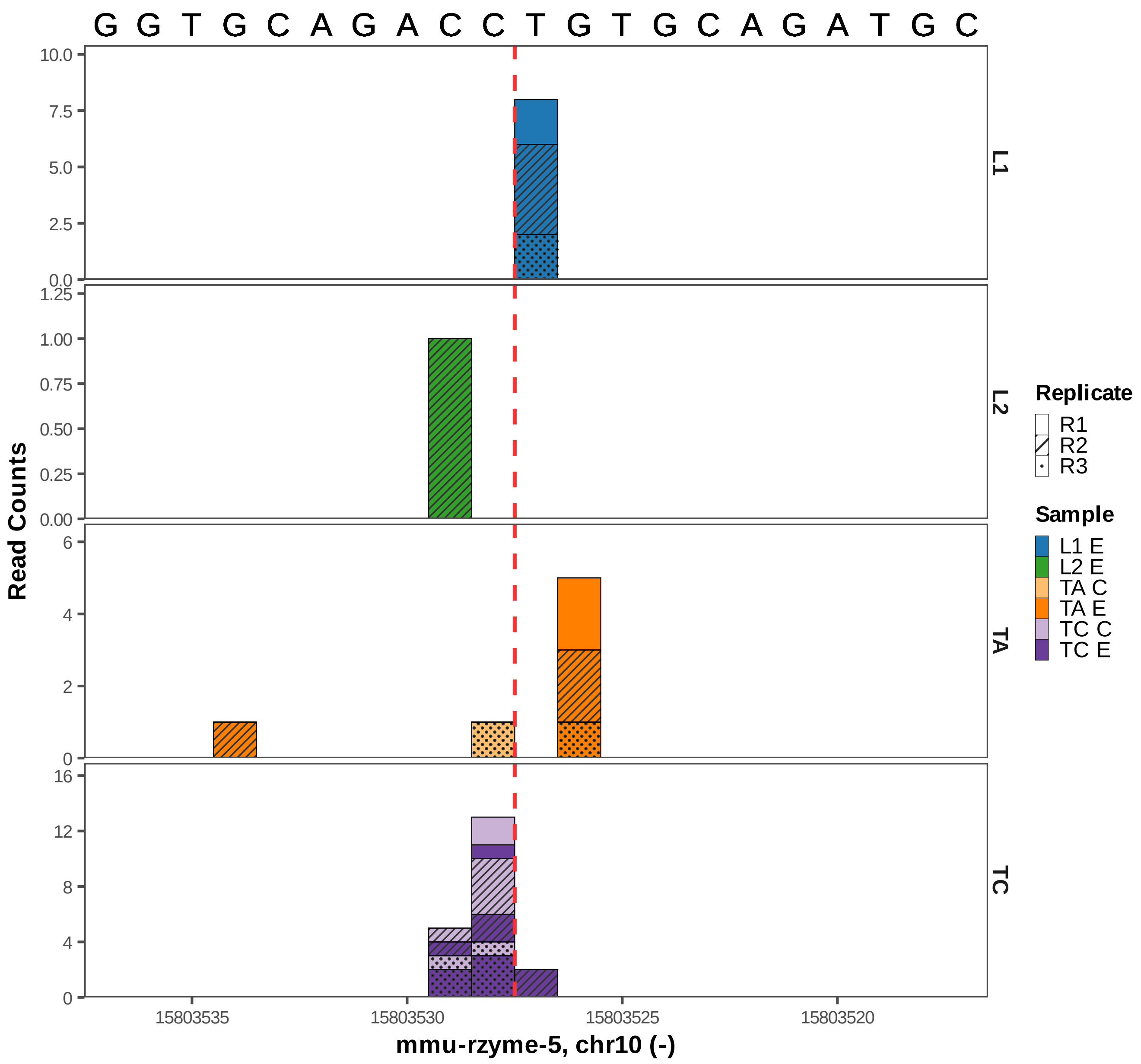

G G T G C A G A C C T G T G C A G A T G C
L1
L2
TA
TC
Read Counts
mmu-rzyme-5, chr10 (-)
15803535
15803530
15803525
15803520
Replicate
R1
R2
R3
Sample
L1 E
L2 E
TA C
TA E
TC C
TC E

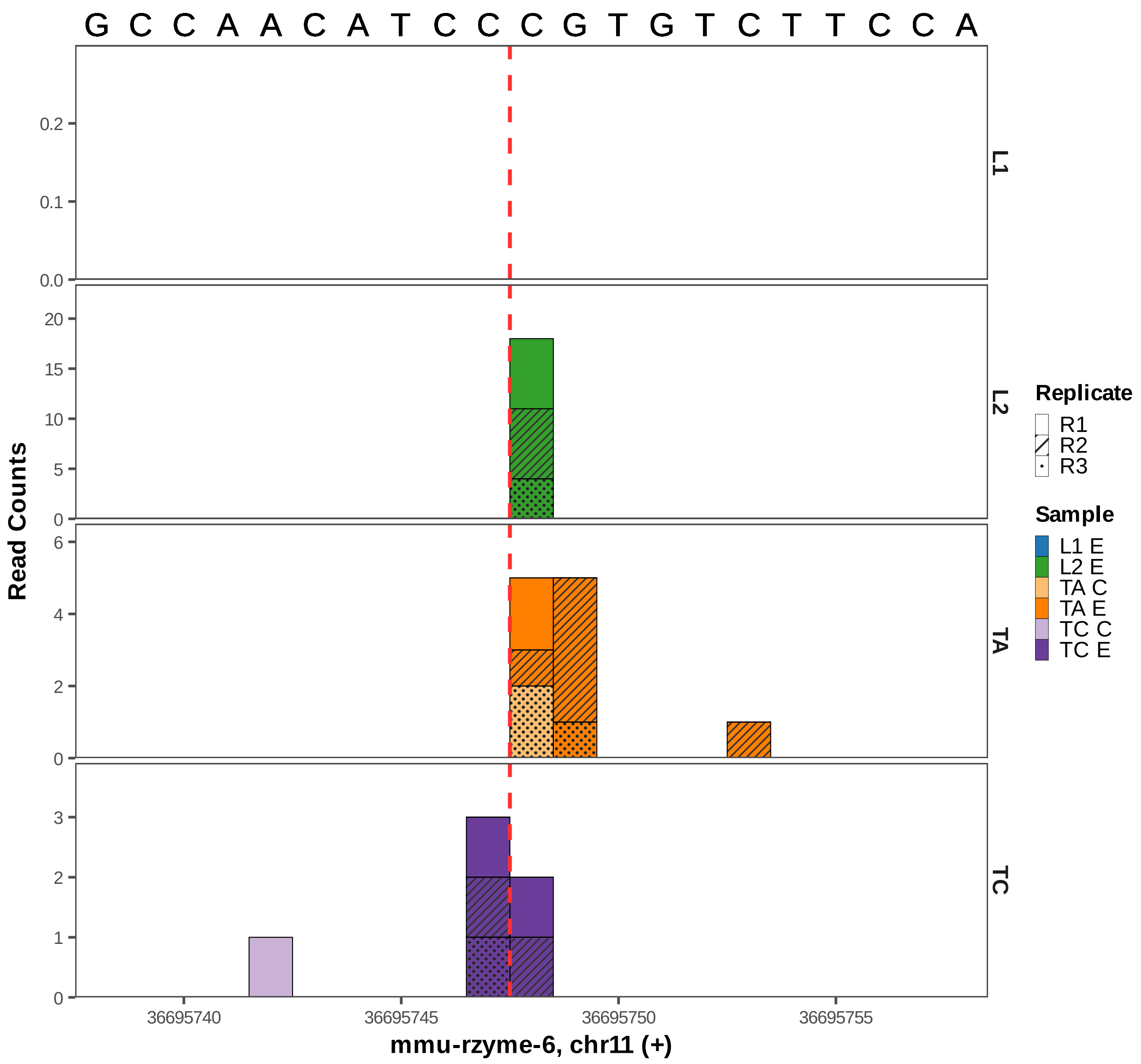
G C C A A C A T C C C G T G T C T T C C A
L1
L2
TA
TC
Read Counts
0.0
0.1
0.2
0
5
10
15
20
0
2
4
6
0
1
2
3
36695740
36695745
36695750
36695755
mmu-rzyme-6, chr11 (+)
Replicate
R1
R2
R3
Sample
L1 E
L2 E
TA C
TA E
TC C
TC E

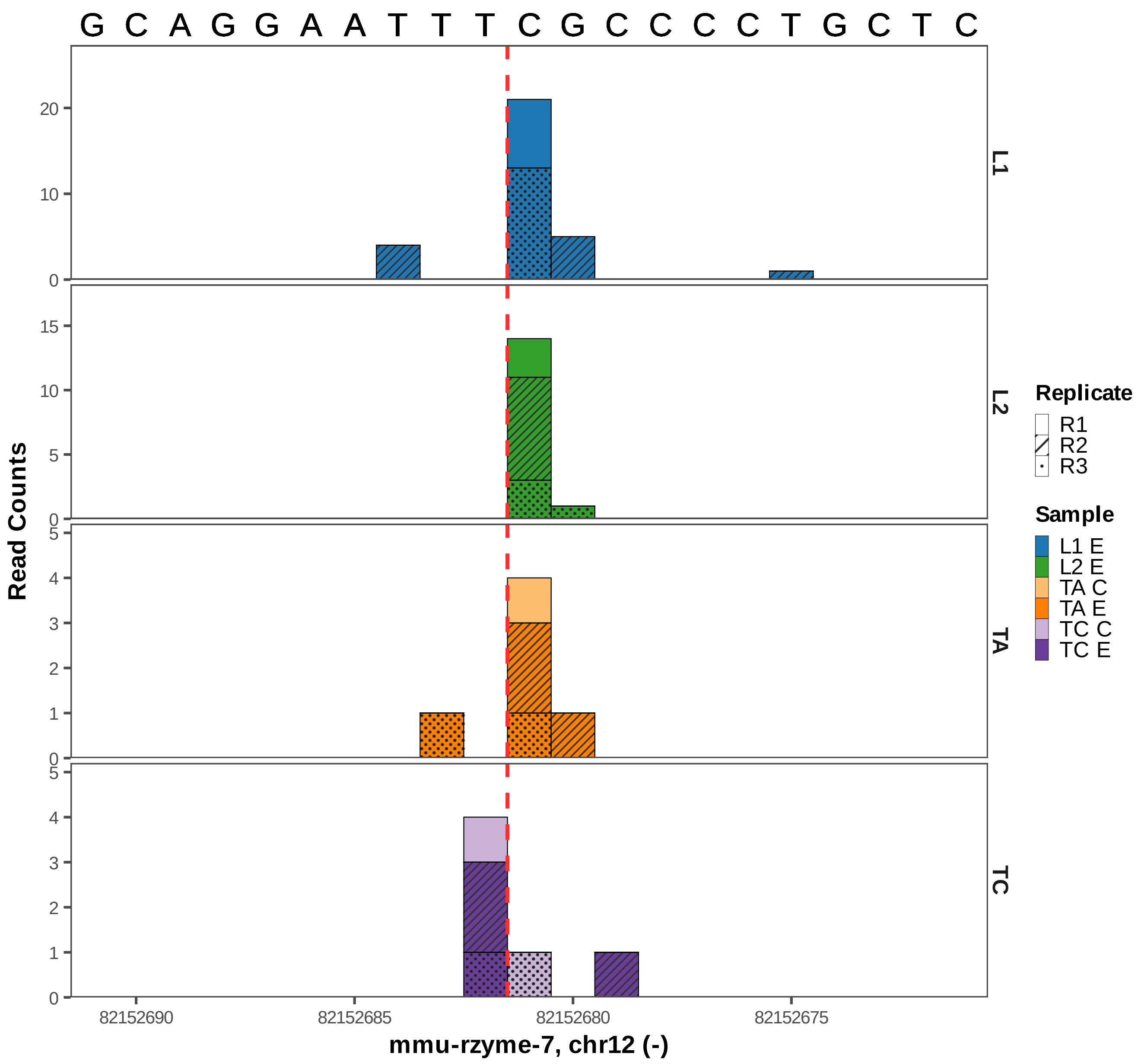

G C A G G A A T T T C G C C C C T G C T C
L1
L2
TA
TC
Read Counts
20
10
15
5
0
4
3
2
1
82152690
82152685
82152680
82152675
mmu-rzyme-7, chr12 (-)
Replicate
R1
R2
R3
Sample
L1 E
L2 E
TA C
TA E
TC C
TC E

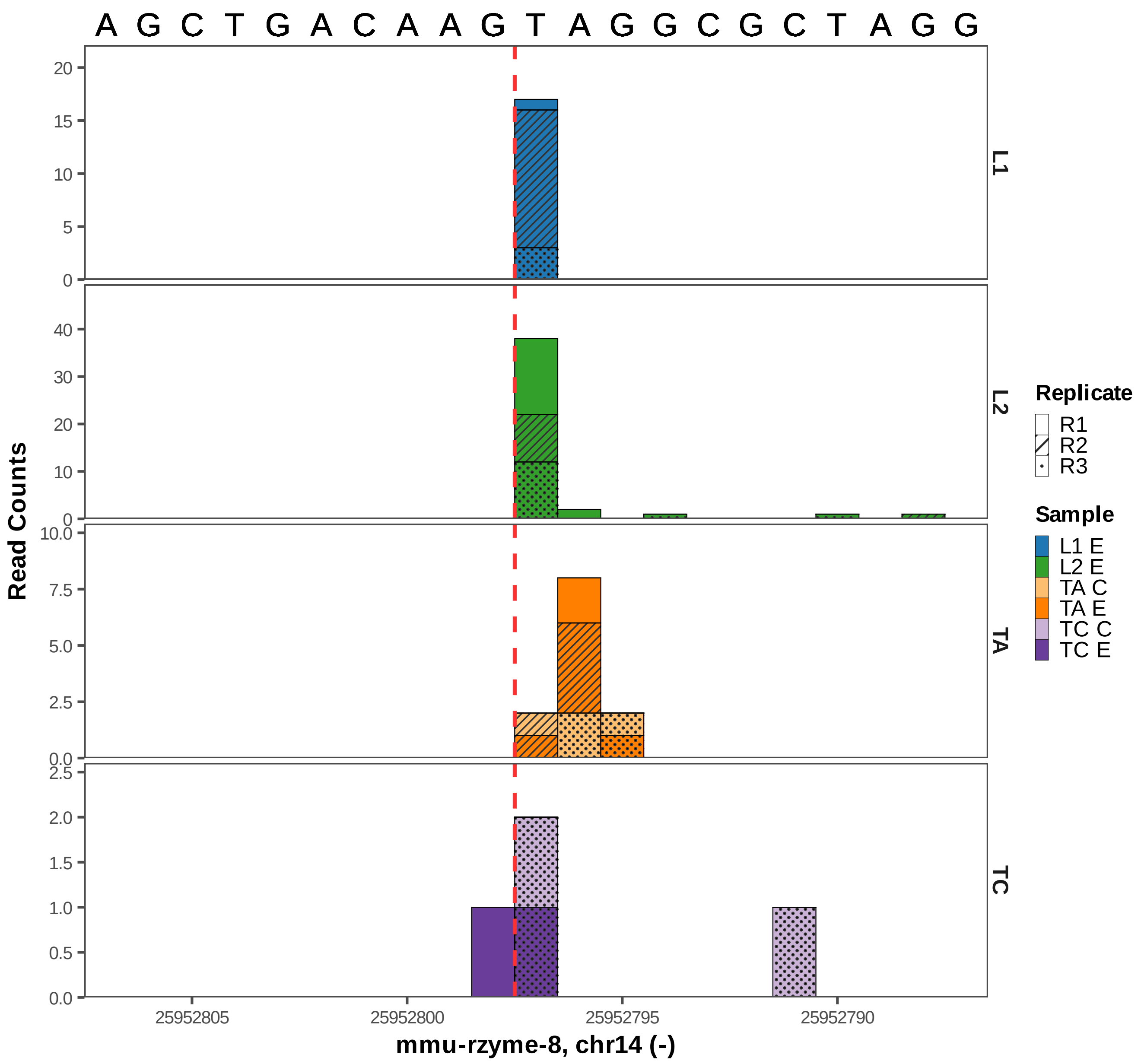

A G C T G A C A A G T A G G C G C T A G G
Read Counts
L1
L2
TA
TC
25952805
25952800
25952795
25952790
mmu-rzyme-8, chr14 (-)
Replicate
R1
R2
R3
Sample
L1 E
L2 E
TA C
TA E
TC C
TC E

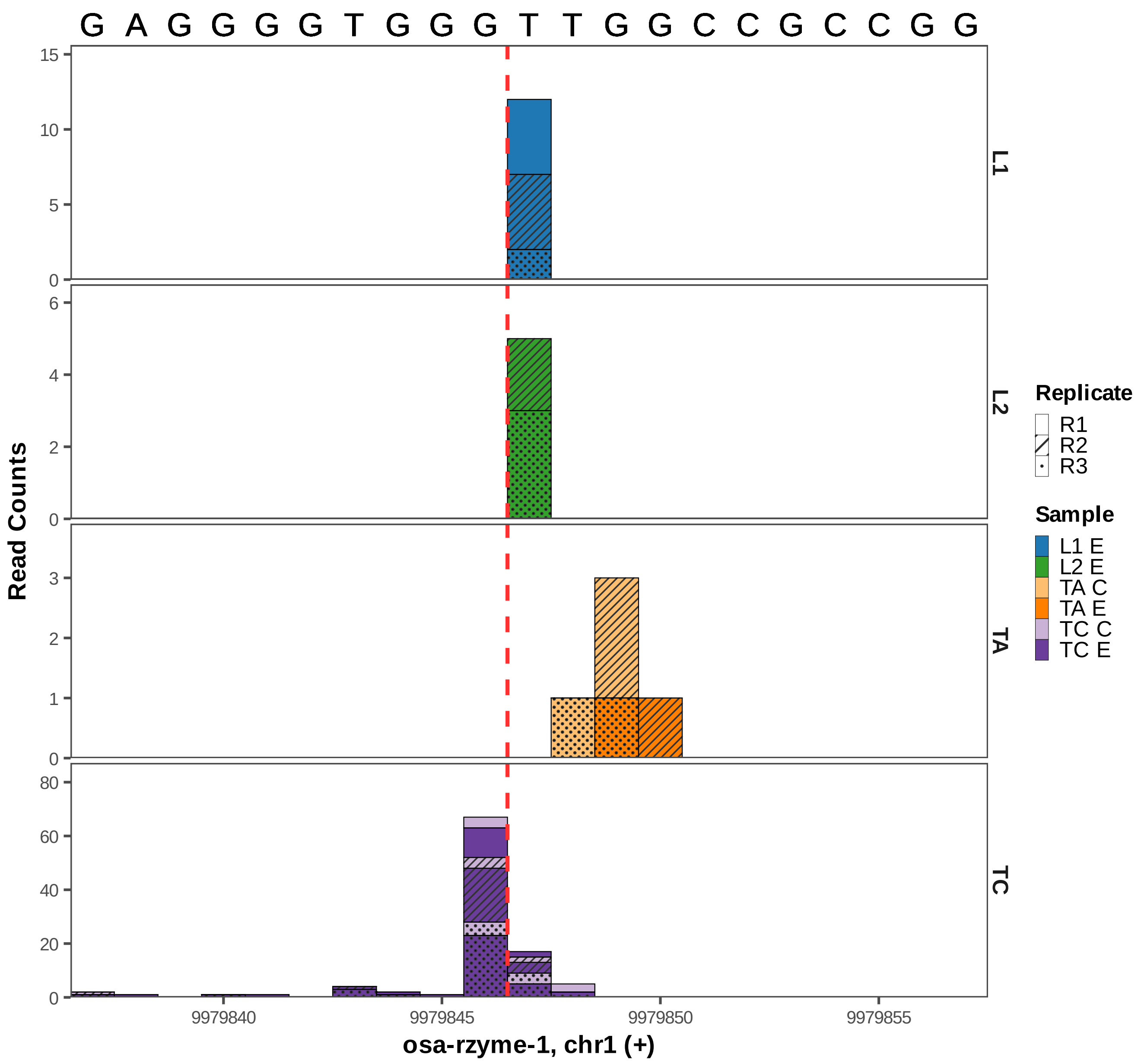

G A G G G G T G G G T T G G C C G C C G G
L1
L2
TA
TC
Read Counts
osa-rzyme-1, chr1 (+)
9979840
9979845
9979850
9979855
Replicate
R1
R2
R3
Sample
L1 E
L2 E
TA C
TA E
TC C
TC E

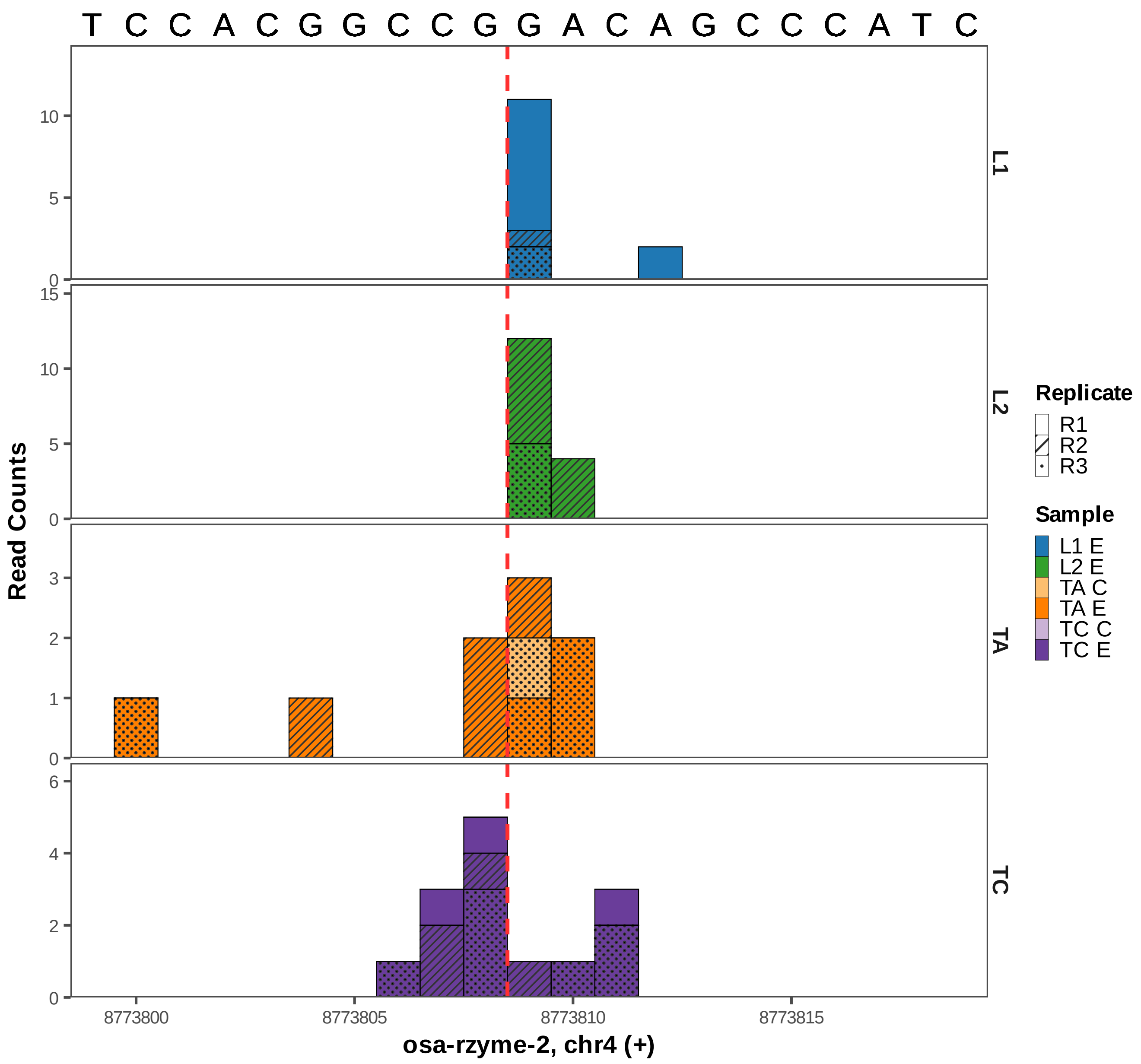
T C C A C G G C C G G A C A G C C C A T C
Read Counts
L1
L2
TA
TC
8773800
8773805
8773810
8773815
osa-rzyme-2, chr4 (+)
Replicate
R1
R2
R3
Sample
L1 E
L2 E
TA C
TA E
TC C
TC E

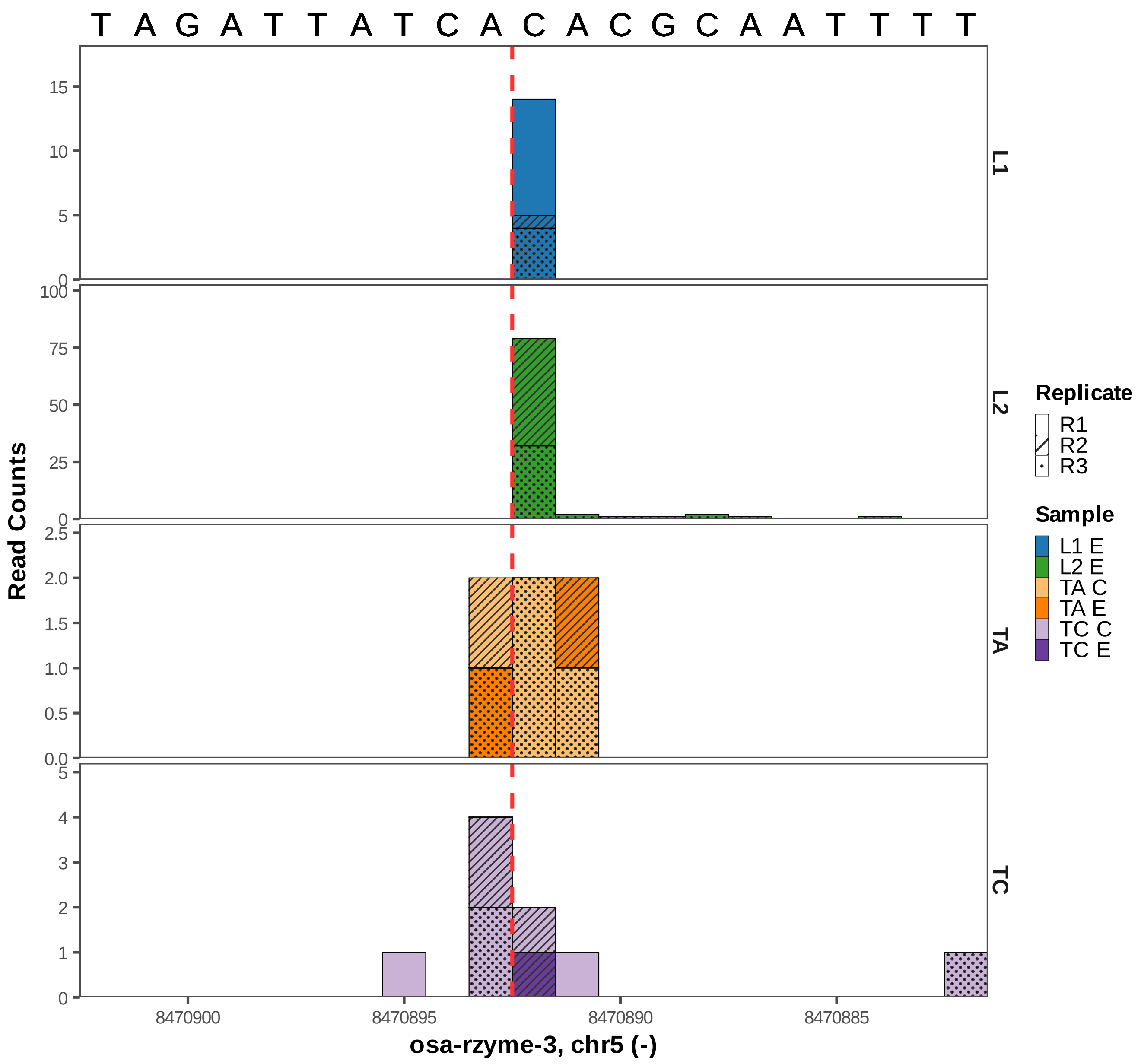

T A G A T T A T C A C A C G C A A T T T T
Read Counts
L1
L2
TA
TC
osa-rzyme-3, chr5 (-)
8470900
8470895
8470890
8470885
Replicate
R1
R2
R3
Sample
L1 E
L2 E
TA C
TA E
TC C
TC E

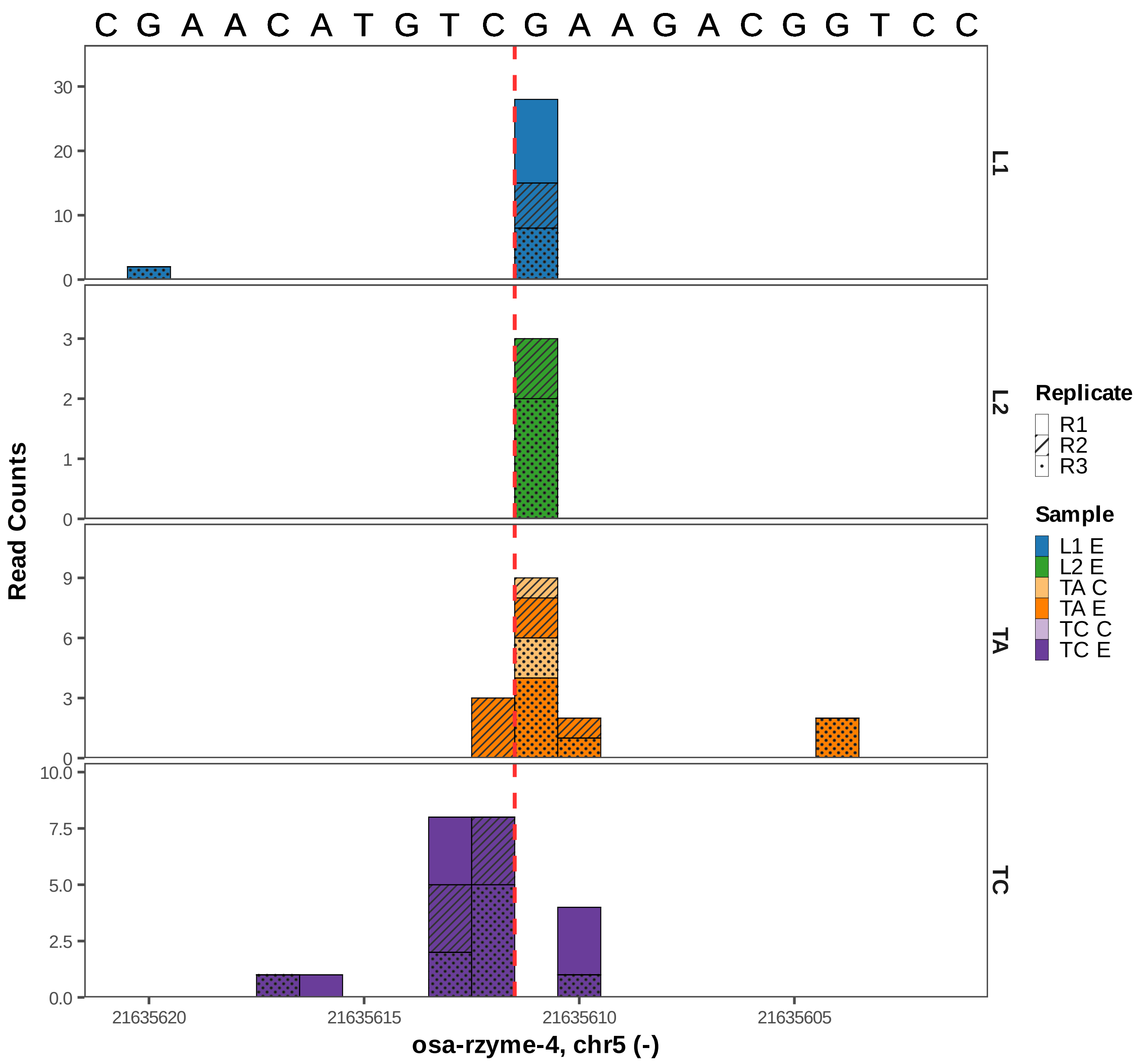

C G A A C A T G T C G A A G A C G G T C C
Read Counts
L1
L2
TA
TC
30
20
10
0
3
2
1
0
9
6
3
0
10.0
7.5
5.0
2.5
0.0
21635620
21635615
21635610
21635605
osa-rzyme-4, chr5 (-)
Replicate
R1
R2
R3
Sample
L1 E
L2 E
TA C
TA E
TC C
TC E

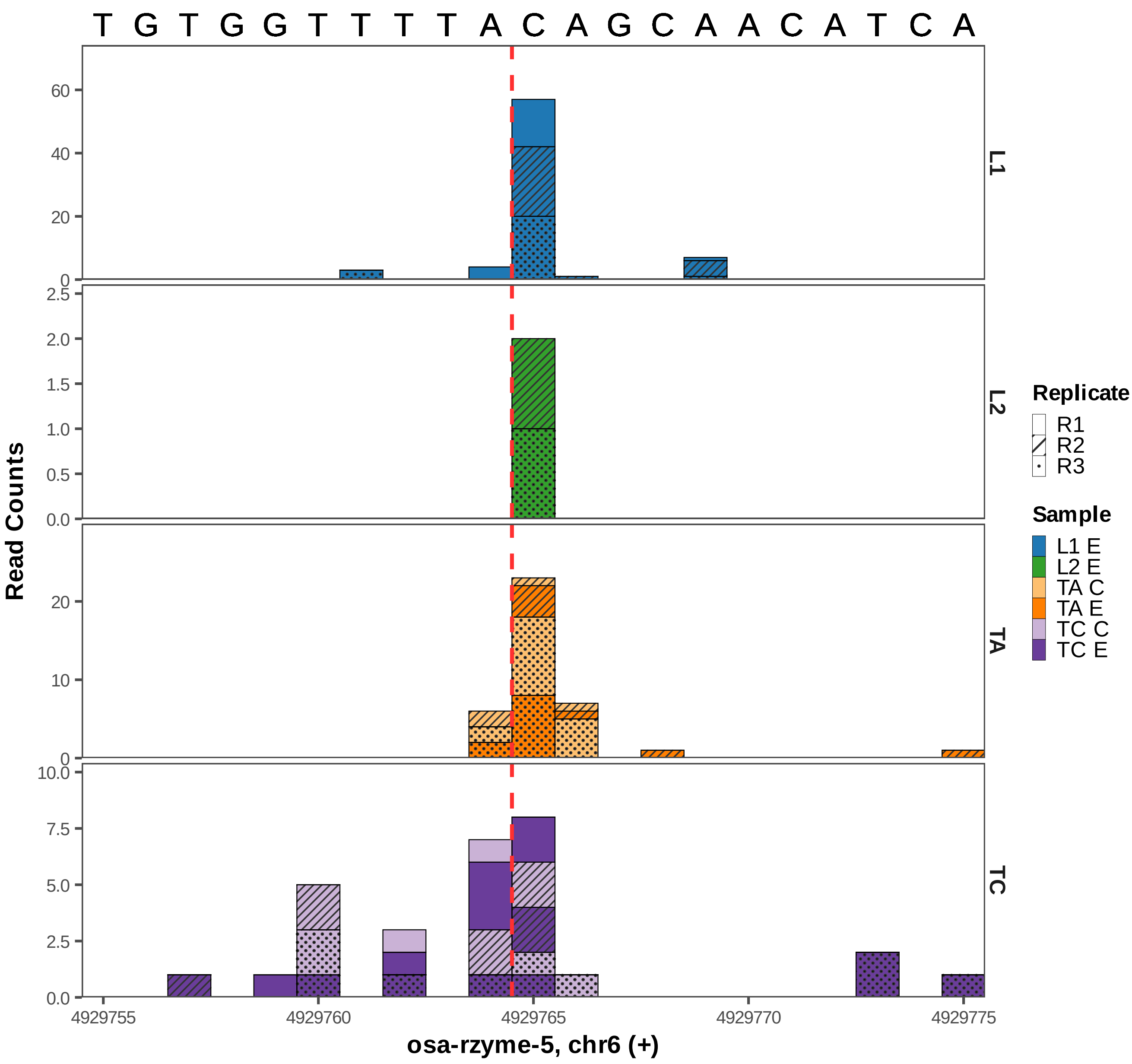

T G T G G T T T T A C A G C A A C A T C A
Read Counts
L1
L2
TA
TC
Replicate
R1
R2
R3
Sample
L1 E
L2 E
TA C
TA E
TC C
TC E
4929755
4929760
4929765
4929770
4929775
osa-rzyme-5, chr6 (+)

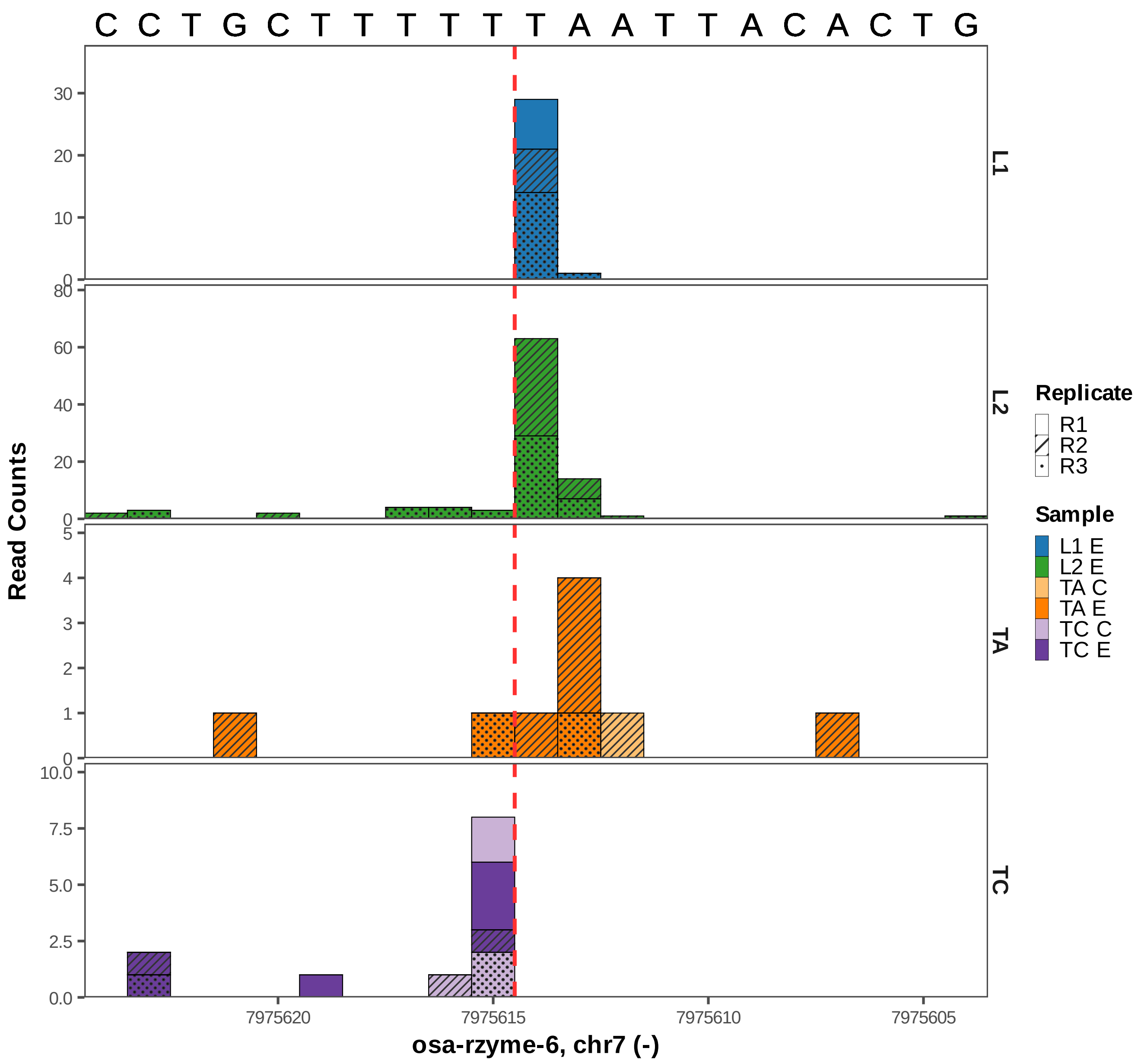
C C T G C T T T T T T A A T T A C A C T G
L1
L2
TA
TC
Read Counts
7975620
7975615
7975610
7975605
osa-rzyme-6, chr7 (-)
Replicate
R1
R2
R3
Sample
L1 E
L2 E
TA C
TA E
TC C
TC E

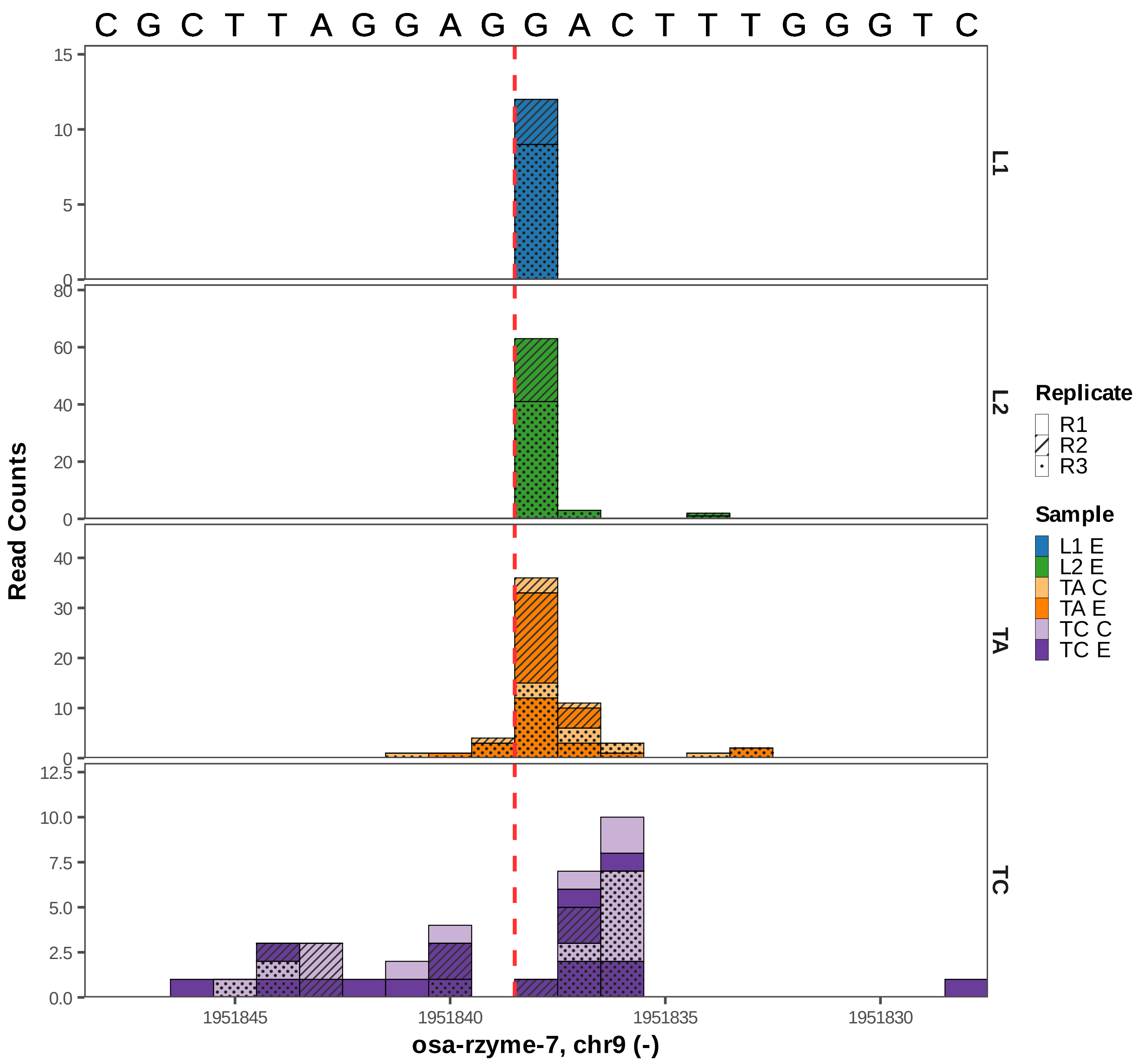

C G C T T A G G A G G A C T T T G G G T C
L1
L2
TA
TC
Read Counts
osa-rzyme-7, chr9 (-)
1951845
1951840
1951835
1951830
Replicate
R1
R2
R3
Sample
L1 E
L2 E
TA C
TA E
TC C
TC E

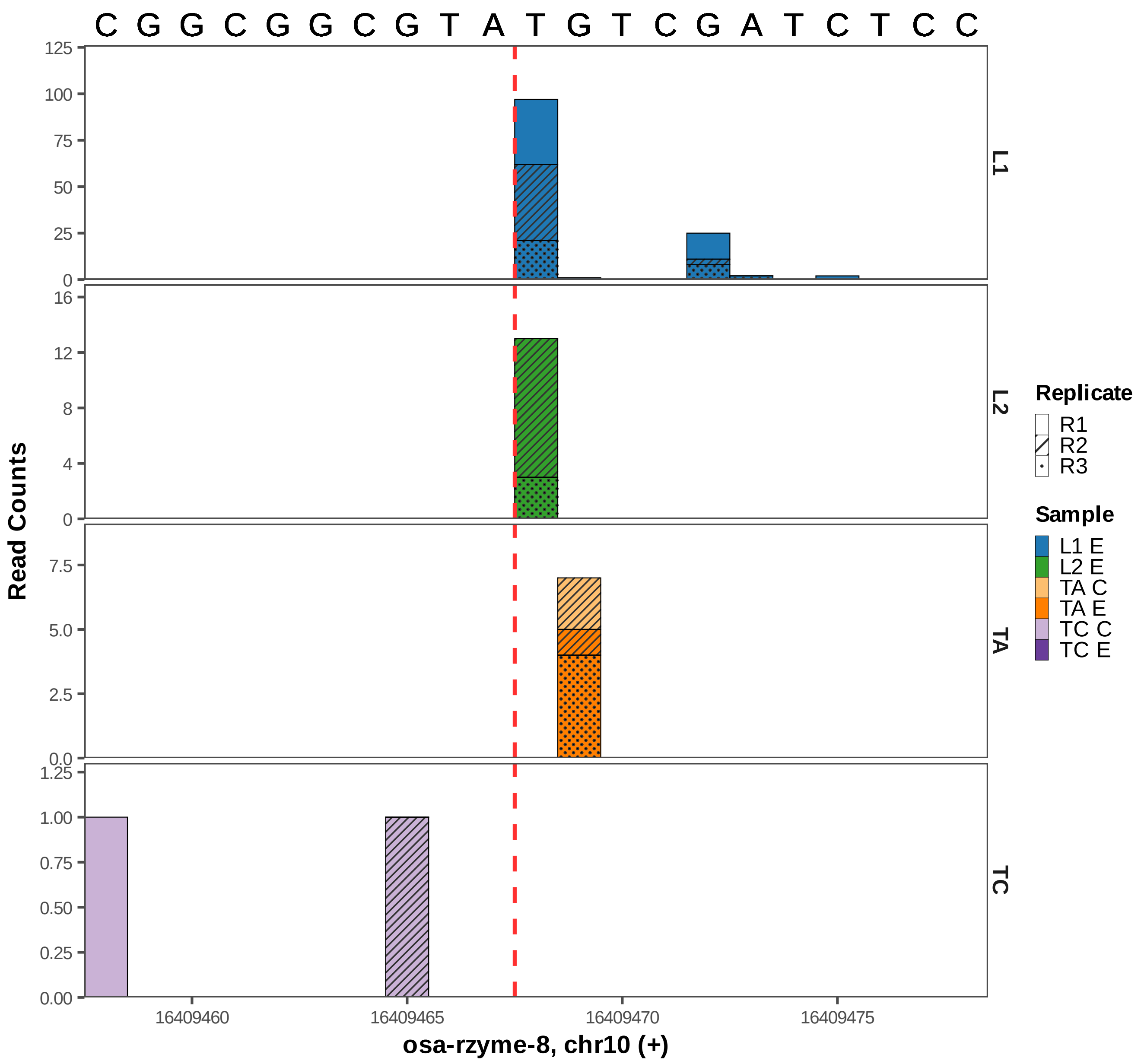

C G G C G G C G T A T G T C G A T C T C C
Read Counts
L1
L2
TA
TC
125
100
75
50
25
0
16
12
8
4
0
7.5
5.0
2.5
0.0
1.25
1.00
0.75
0.50
0.25
0.00
16409460
16409465
16409470
16409475
osa-rzyme-8, chr10 (+)
Replicate
R1
R2
R3
Sample
L1 E
L2 E
TA C
TA E
TC C
TC E

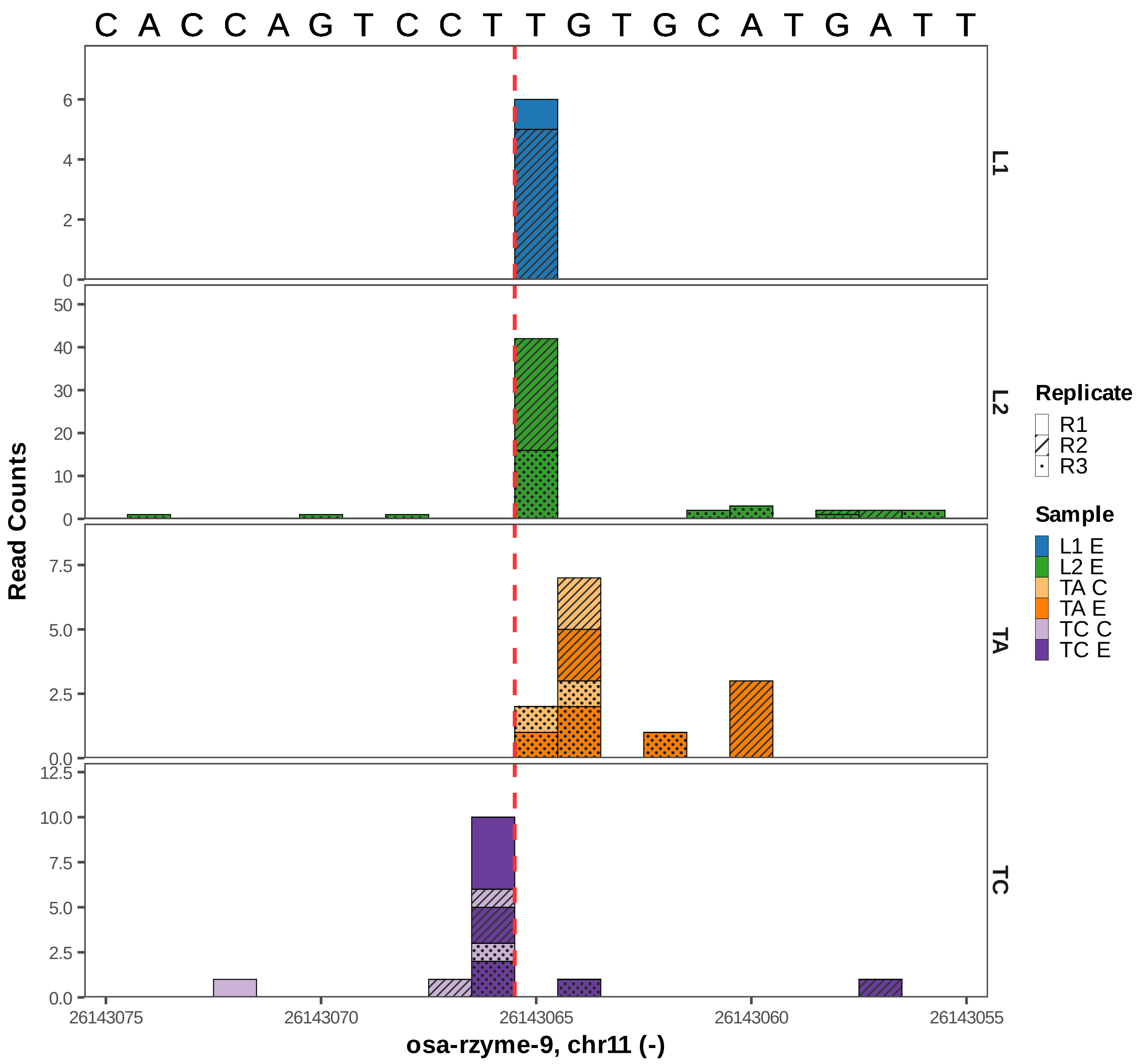

C A C C A G T C C T T G T G C A T G A T T
Read Counts
L1
L2
TA
TC
26143075
26143070
26143065
26143060
26143055
osa-rzyme-9, chr11 (-)
Replicate
R1
R2
R3
Sample
L1 E
L2 E
TA C
TA E
TC C
TC E

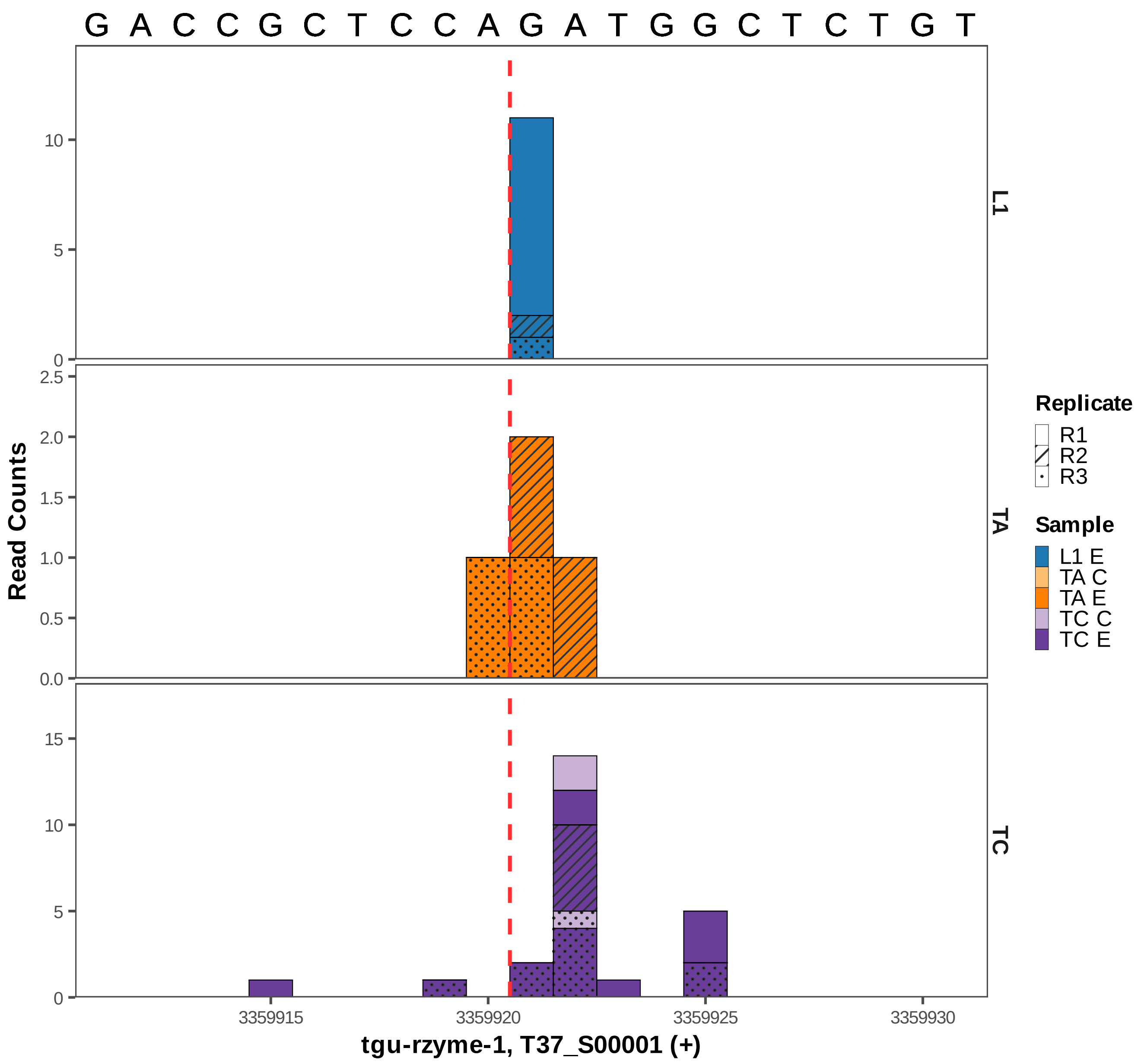
G A C C G C T C C A G A T G G C T C T G T
L1
TA
TC
Read Counts
tgu-rzyme-1, T37_S00001 (+)
3359915
3359920
3359925
3359930
Replicate
R1
R2
R3
Sample
L1 E
TA C
TA E
TC C
TC E

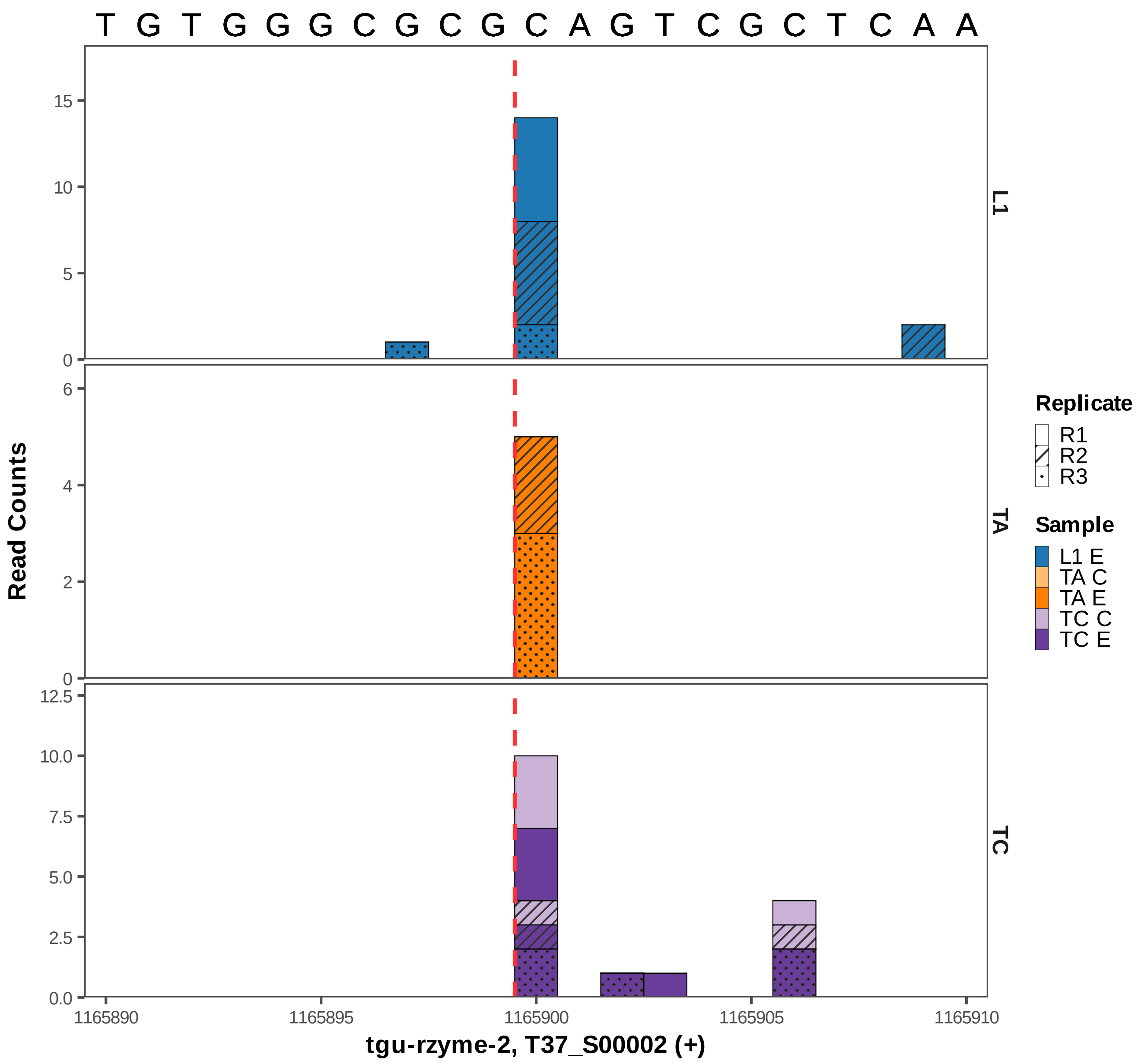
T G T G G G C G C G C A G T C G C T C A A
Read Counts
L1
TA
TC
Replicate
R1
R2
R3
Sample
L1 E
TA C
TA E
TC C
TC E
0
5
10
15
0
2
4
6
0.0
2.5
5.0
7.5
10.0
12.5
1165890
1165895
1165900
1165905
1165910
tgu-rzyme-2, T37_S00002 (+)

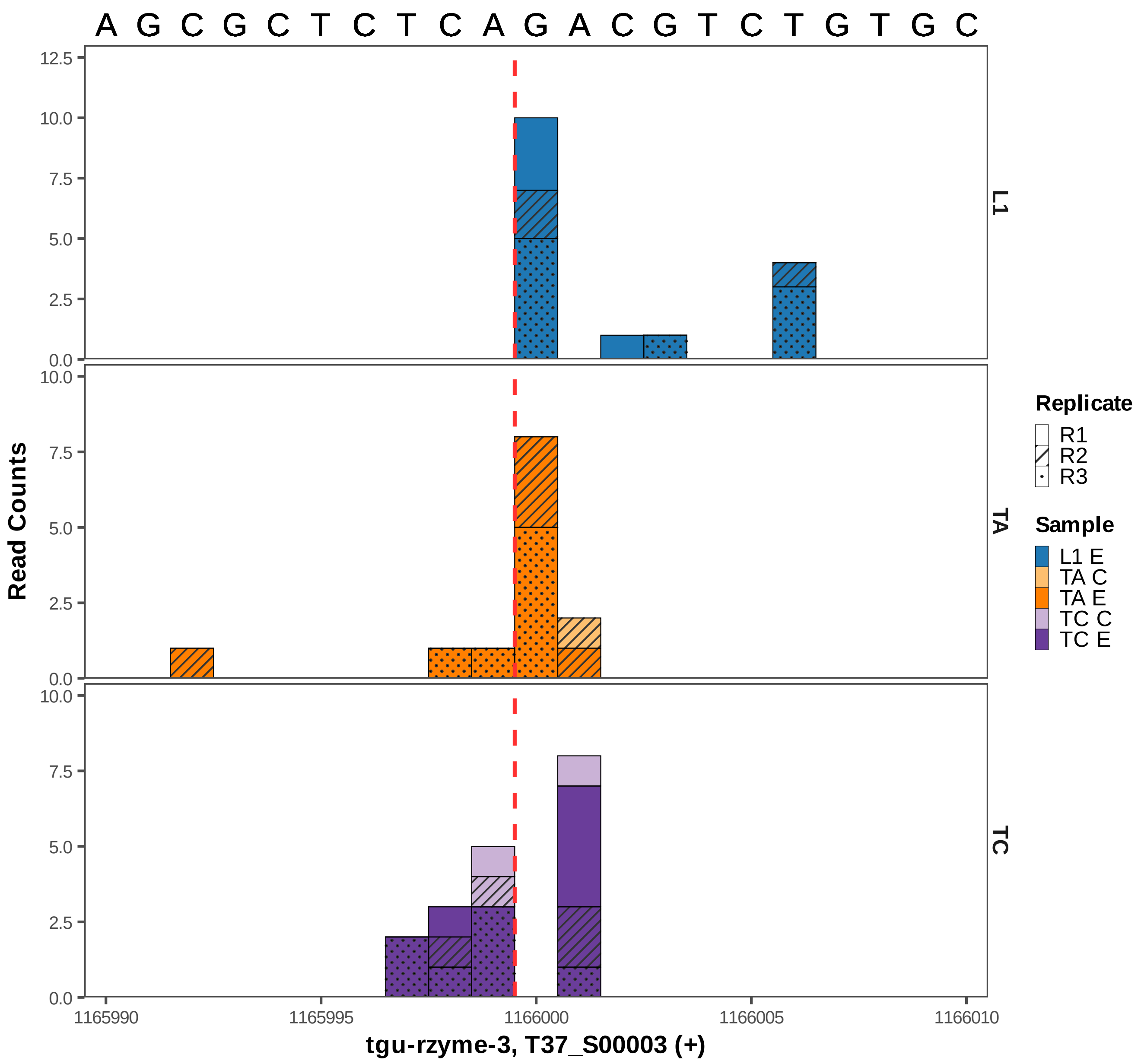
A G C G C T C T C A G A C G T C T G T G C
12.5
10.0
7.5
5.0
2.5
0.0
L1
TA
TC
Read Counts
Replicate
R1
R2
R3
Sample
L1 E
TA C
TA E
TC C
TC E
1165990
1165995
1166000
1166005
1166010
tgu-rzyme-3, T37_S00003 (+)

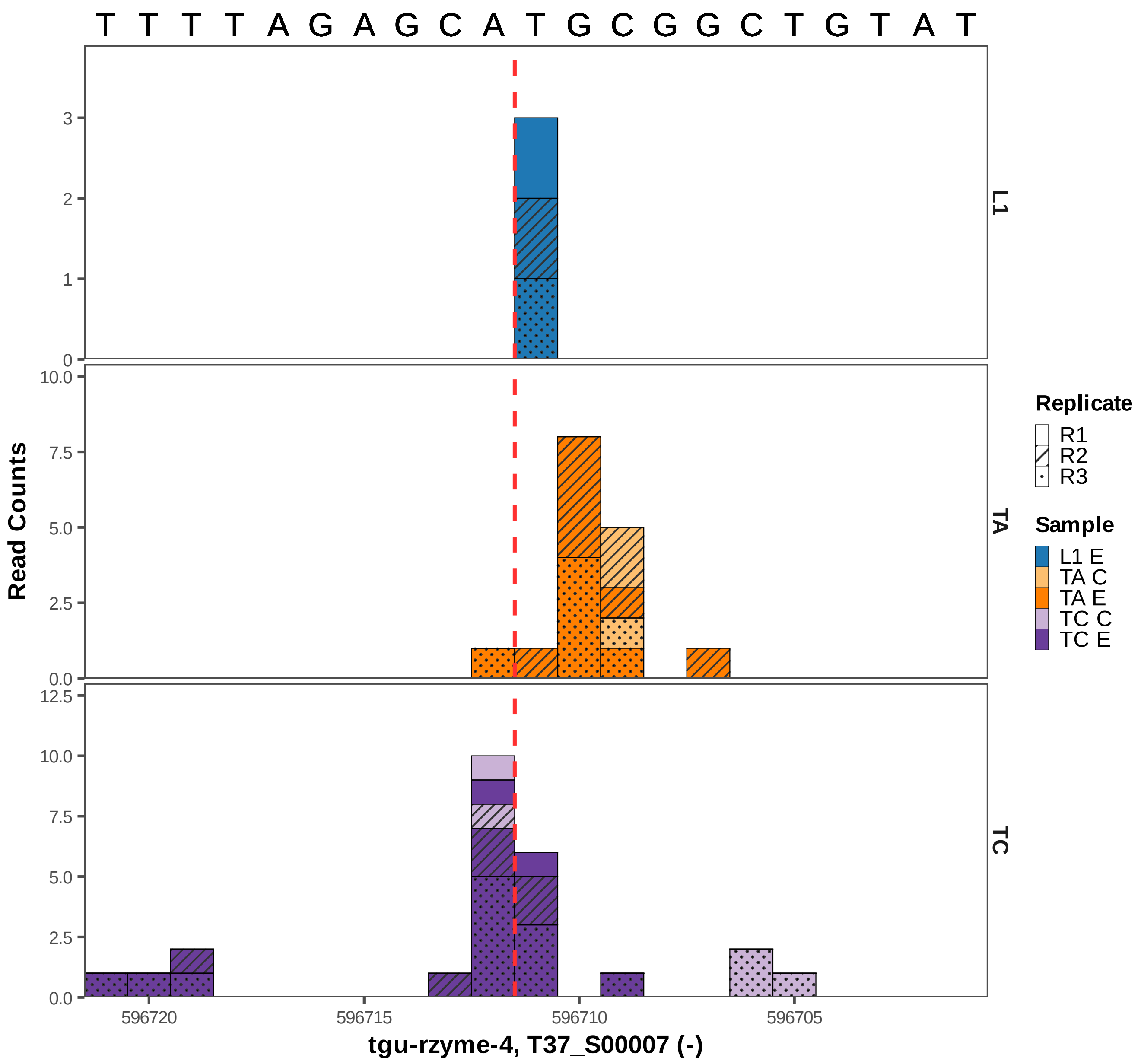

T T T T A G A G C A T G C G G C T G T A T
Read Counts
L1
TA
TC
Replicate
R1
R2
R3
Sample
L1 E
TA C
TA E
TC C
TC E
596720
596715
596710
596705
tgu-rzyme-4, T37_S00007 (-)

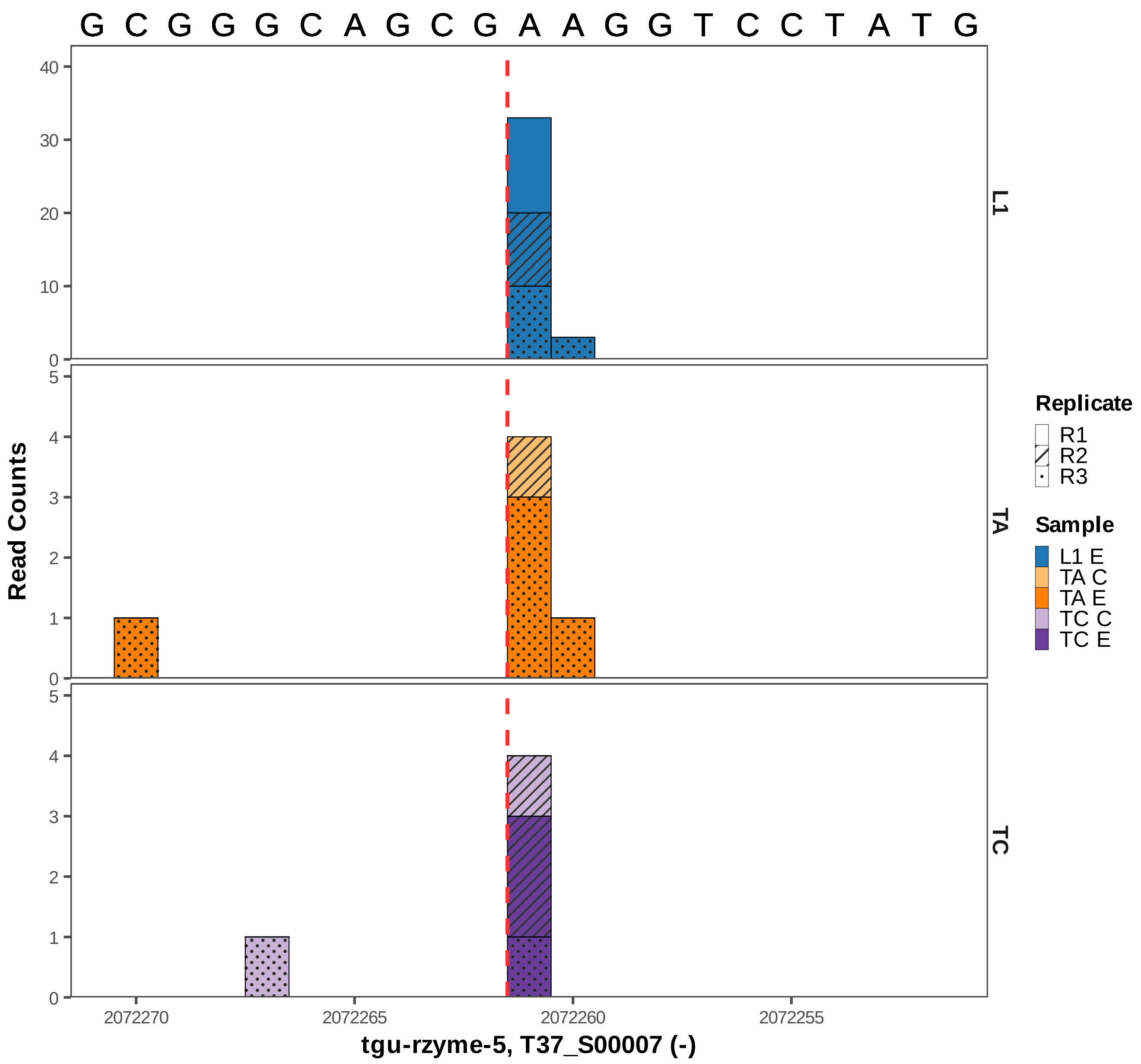

G C G G G C A G C G A A G G T C C T A T G
Read Counts
L1
TA
TC
Replicate
R1
R2
R3
Sample
L1 E
TA C
TA E
TC C
TC E
2072270
2072265
2072260
2072255
tgu-rzyme-5, T37_S00007 (-)

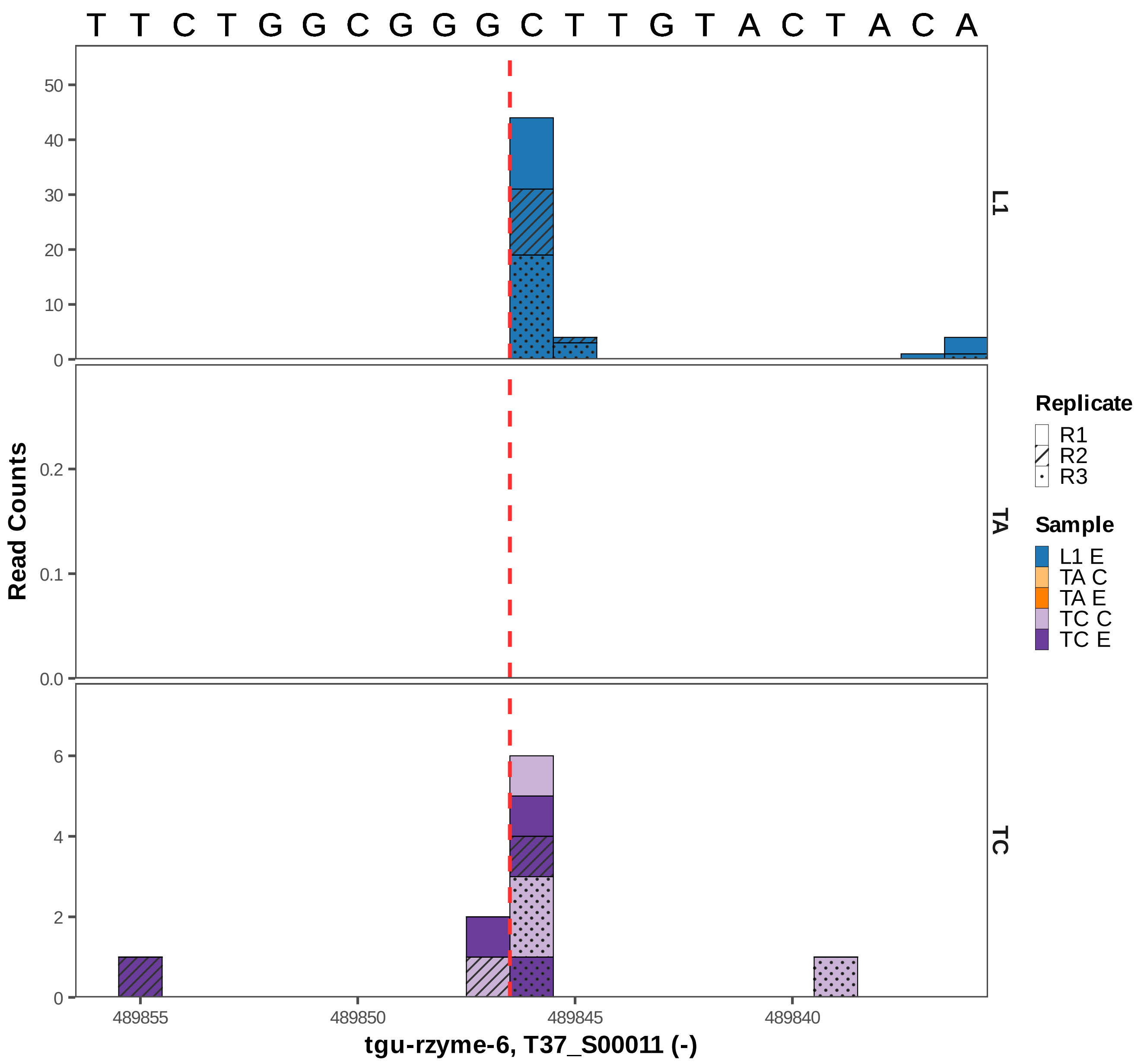

T T C T G G C G G G C T T G T A C T A C A
Read Counts
L1
TA
TC
Replicate
R1
R2
R3
Sample
L1 E
TA C
TA E
TC C
TC E
489855
489850
489845
489840
tgu-rzyme-6, T37_S00011 (-)

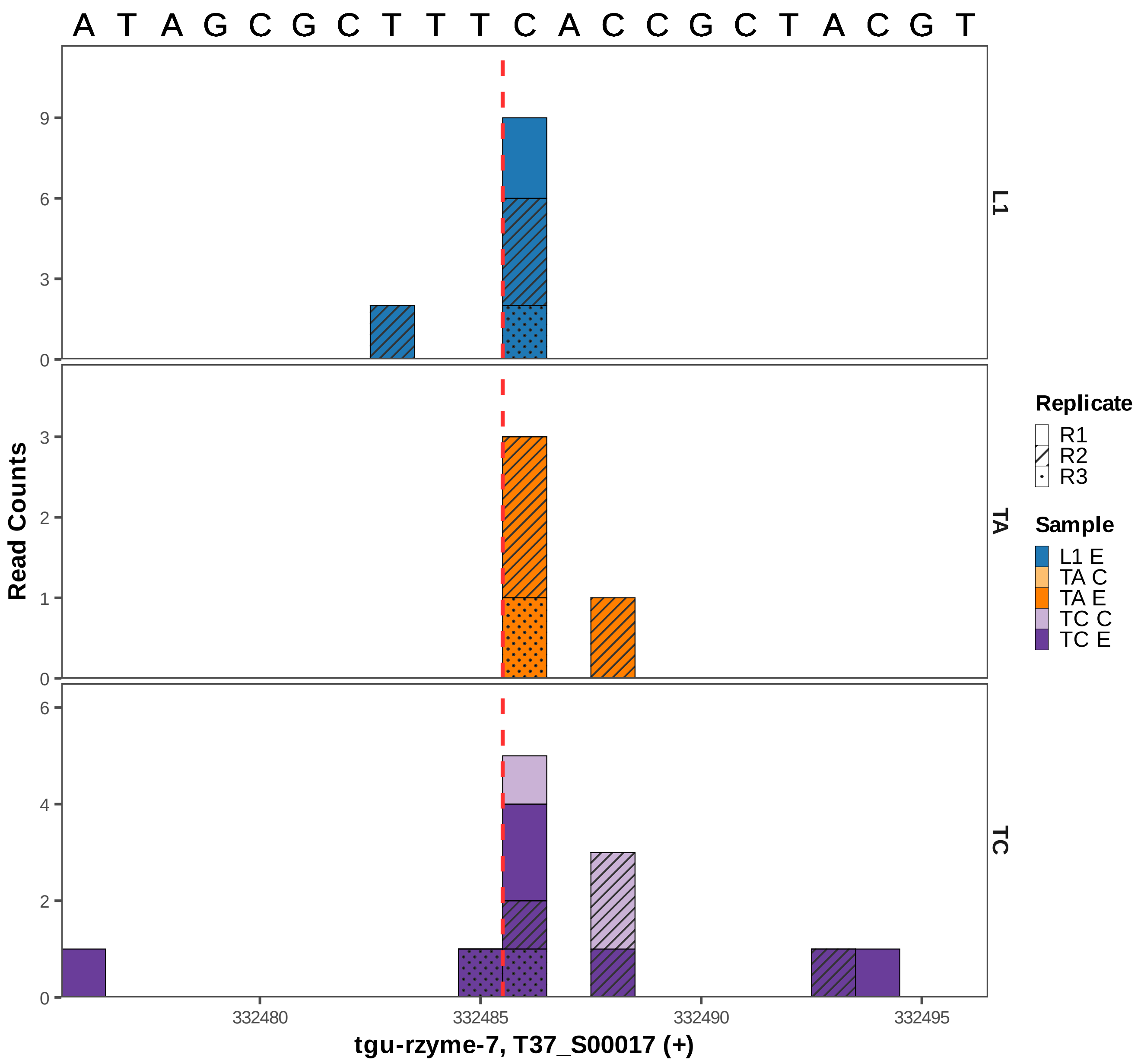

A T A G C G C T T T C A C C G C T A C G T
L1
TA
TC
Read Counts
0
3
6
9
0
1
2
3
0
2
4
6
332480
332485
332490
332495
tgu-rzyme-7, T37_S00017 (+)
Replicate
R1
R2
R3
Sample
L1 E
TA C
TA E
TC C
TC E
