## Supplementary Figures for "Widespread Presence of Catalytic RNAs in All Domains of Life": Supplementary Figure Guide.docx

**Figure S1. Detailed schematic representation of the key molecular biological steps of the AbRiDGed method.**

The method detects the genomic position of the first nucleotide of the downstream cleavage product. RT, reverse transcription; IVT, *in vitro* transcription.

**Figure S2. Detection of the known human ribozymes by the AbRiDGed method.**

The read counts (Y-axes) were obtained from the high-throughput sequencing data of each sample. The red dashed lines mark the known cleavage sites, and the X-axes denote the genomic coordinates (from 5′ end to 3′ end of each ribozyme). “L1”, “L2”, “TA” and “TC” denote the library types; “R1” and “R2” denote the two replicates; “E” and “C” denote samples that were respectively treated or not with the RppH and XRN-1 enzymes.

**Figure S3. Detection of the cleavage sites of the spike-in ribozymes via AbRiDGed.**

The annotations are as described in the legend to Figure S2 with the exception that the X-axes represent coordinates within the spike-in sequences that are listed in Table S1.

**Figure S4. Cleavage sites of the novel human ribozymes detected by AbRiDGed.**

The annotations are as described in the legend to the top panel of Figure 1B.

**Figure S5. Validation gels for the novel human ribozymes from the Tier 1 candidates identified via AbRiDGed.**

Two replicates of validation gels are shown for each ribozyme. The validation gels annotations are as described in the legend to Figures 1B and 1C. The names of the ribozymes are shown on the top of the gels with the expected sizes of the uncleaved RNAs (“unclv”) and the upstream and downstream cleavage products (“5′-clv” and “3′-clv”, respectively) annotated in the parenthesis respectively from left to right.

**Figure S6. Validation gels for the novel human ribozymes from the Tiers 2 and 3 candidates identified via AbRiDGed.**

The annotations are as described in the legend to Figure S5.

**Figure S7. Cleavage sites of novel ribozymes from the 7 eukaryotes detected by AbRiDGed.**

The annotations are as described in the legend to the top panel of Figure 1B.

**Figure S8. Validation gels for the novel ribozymes from the 7 eukaryotes.**

The annotations are as described in the legend to Figure S5.

**Figure S9. Evolutionary analysis of the novel human self-cleaving ribozymes.**

(A) Evolutionary histories of the novel ribozymes. Each column of the heatmap represents a ribozyme and the color of the cells represents the maximum sequence identity to the human sequence among the homologs found in the corresponding clade shown in the evolutionary tree. The numbers of species included in the analysis are in the parentheses.

(B) Evolutionary histories of the novel ribozymes in primates. Red names of groups of species or individual species in the evolutionary tree (left) indicate the presence of the self-cleavage activity for at least one of the 7 selected ribozymes as shown in detail in the right panel. NA, no homologous sequence was found. Middle, the sequence identity heatmap as described in (A).

(C) Evolutionary history of the hsa-rzyme-97 ribozyme in mammals (left) and primates (right). The red names represent the presence of either the homologous sequences (left) or self-cleavage activity (right). The red dot represents the deduced first appearance of the self-cleavage activity which was subsequently lost in the gorilla branch (blue branch). The colored boxes indicate the branches that contain sequences that are either homologous (left) or highly similar (>90% sequence identity; right) to the human hsa-rzyme-97 ribozyme, respectively.

(B and C) The numbers in the parentheses represent respectively from left to right the numbers of species tested for the self-cleavage activity, possessing homologous sequences and included in the analysis.

**Figure S10. Detection of self-cleavage activity in various primate homologs of novel human ribozymes.** The positions of uncleaved RNAs and the upstream and downstream cleavage products are shown by respectively black, blue and red arrows. Note, when calculating the expected sizes, the 1−3 guanosines potentially added during sequence construction were included. The “E” lanes contain the control reactions with 6 mM EDTA and no Mg^2+^; the “M” lanes contain the size standards.

**Figure S11. Single nucleotide substitutions (nearly) abolish activities of novel self-cleaving ribozymes.**

(A) Schematics of the single nucleotide substitutions that affect the activities of the novel ribozymes.

(B) The corresponding gels showing the products of the cleavage reactions (performed in 30 mM Tris-HCl pH 8, 20 mM MgCl_2_ and 150 mM KCl for 6 h at 37°C) of the wild-type human (WT) and the mutant (MT) ribozyme sequences. The gel annotations are as described in the legend to Figures 1B and 1C. The WT and MT sequences are shown in Table S13.

**Figure S12. Biochemical characterization** **of the hsa-rzyme-97 ribozyme.**

(A) The cleavage time course (left) and the first-order rate constant (*k*_obs_, right) calculated based on the cleavage fractions (*F_t_*) obtained from the time course.

(B and C) Dependency of *k*_obs_ on pH (B) and Mg^2+^ concentration (C).

(D) Activity of the hsa-zyme-97 ribozyme in different cations.

(E) Mass spectrum analysis of the cleavage products of a 34-nt hsa-rzyme-97 sequence (Table S22).

(A–C) The orange and blue dots represent 2 replicates.

(A and D) The annotations of the gels are as described in the legend to Figures 1B and 1C.

**Figure S13.** **Comparison of the self-cleavage kinetics of self-cleaving ribozymes and the *metY*** **SAM-V riboswitch sequence.**

(A) Detection of the SAM-V cleavage using the published condition in which its cleavage was originally found (50 mM Tris-HCl pH 8.3, 20 mM MgCl_2_ and 100 mM KCl at 23°C).

(B) The cleavage fractions (Y-axis) of the known (orange) and novel (blue) ribozymes as well as the SAM-V sequence (magenta) as a function of the reaction time (X-axis) under the same condition as in (A). The entire time course data are shown — the data of 0−48 hours are the same as those shown in the left plot on the Figure 5E.

(C) SAM-V cleavage cannot be detected under the conditions used in this study to measure the ribozyme activity.

(A and C) The “E” lanes contain the control reactions with 6 mM EDTA, no Mg^2+^ and no SAM; the “Mg^2+^” lanes contain control reactions with no SAM; the “M” lanes contain the size standards; the “R” lanes contain the synthetic RNA identical to the expected 48-nt downstream cleavage product and used as the size reference. The positions of the 62-nt uncleaved RNAs (“unclv”) and the 48-nt downstream cleavage product (“3′-clv”) are marked.

**Figure S14. MINDS ribozymes are active in low concentrations of various monovalent cations.**

(A) The initial discovery of the MINDS ribozymes by virtue of their activity in either the presence (the “Mg^2+^” lanes) or absence (the “E” lanes) of Mg^2+^ as exemplified by the validation gels for the hsa-MINDS-9 and hsa-MINDS-1 ribozymes. The gel annotations are as described in the legend to Figures 1B and 1C.

(B) Activity of the MINDS ribozymes (Table S17) in 150 mM of various monovalent cations in the presence of 30 mM Tris-HCl pH 8 for 24 hours at 37°C. The gel annotations are as described in the legend to Figure 6A.

**Figure S15. Known classes of self-cleaving ribozymes are mostly inactive in low concentrations of various monovalent cations.**

(A and B) The ribozyme sequences corresponding to the known classes (listed in Table S1) are active in Mg^2+^ (A), yet mostly inactive in 150 mM of various monovalent cations under the same conditions as in Figure S14B (B). The “M” lanes contain the size standards. The “R” lanes contain the IVT products of the ribozymes without any incubation. Since some ribozymes self-cleave during the IVT and purification steps, a ribozyme is considered active in a particular cation only if the corresponding cleavage fraction is higher than the one in the “R” lane.

(B) The positions of uncleaved RNAs (“unclv”) and the upstream and downstream cleavage products (“5′-clv” and “3′-clv”, respectively) are shown by respectively black, blue and red arrows in the lanes where self-cleavage activity was observed. Note, the 13-nt upstream cleavage product of the HPR_1 ribozyme is too short for detection.

**Figure S16. The short stem-loop structures of the MINDS ribozymes and their activities.**

(A) The short stem-loop structures of the MINDS ribozymes. The red triangles mark the cleavage sites.

(B) Activities of the short MINDS ribozyme sequences in K^+^. The “M” lanes contain the size standards; the “R” lanes contain the intact synthetic RNAs used as the control. The positions of uncleaved RNAs (“unclv”) and the upstream and downstream cleavage products (“5′-clv” and “3′-clv”, respectively) are shown by respectively black, blue and red arrows. Note, the downstream cleavage products of several ribozymes are too short for detection.

**Figure S17. Biochemical characterization** **of the hsa-MINDS-1 ribozyme.**

(A) Left, the short stem-loop structure of the 25-nt hsa-MINDS-1 sequence (Table S22). The red triangle marks the cleavage site; “M1” denotes the “U>dU” substitution. Right-top, the “U>dU” substitution (“M1”; Table S22) abolished the activity of the hsa-MINDS-1 ribozyme compared to the original sequence (“WT”). Right-bottom, mass spectrum analysis of the cleavage products of the 25-nt hsa-MINDS-1 sequence.

(B) Activity of the hsa-MINDS-1 ribozyme in various cations.

(C) The *F_t_* and *k*_obs_ of the hsa-MINDS-1 ribozyme in 150 mM K^+^, 6 mM Mg^2+^ or 150 mM K^+^ and 6 mM Mg^2+^.

**(**D−F**)** Dependency of the *k*_obs_ of the hsa-MINDS-1 ribozyme on Mg^2+^ concentration (D), K^+^ concentration (E) and pH (F).

(G) Comparison of the self-cleavage kinetics of the hsa-MINDS-1 ribozyme and the *metY* SAM-V riboswitch sequence.

(A and B) the “M” lanes contain the size standards; the “R” lanes contain the intact 25-nt RNAs as the control; the 8-nt downstream cleavage product is too short for detection.

**Figure S18. In-depth sequence-structure analysis of the MINDS ribozymes.**

(A and B) The synthetic sequence variants of the 15-nt hsa-MINDS-5 ribozyme and their self-cleavage activities. Left, the stem-loop structure of the 15-nt hsa-MINDS-5 ribozyme with the various elements subjected to the mutagenesis analysis demarcated by the dashed lines to test the importance of specific nucleotides and base pairs (A) and MFE (B) for the activity. For the panels B, the corresponding mutated sequences are shown in boxes with the resulting MFE of the stem-loop structures shown below each box in kcal/mol. The red triangles mark the cleavage site. Right, the activity tests for the various sequence variants shown in the left panels. The text above each lane refers to single- or di-nucleotide substitutions at various positions relative to the cleavage site (A), and multi-nucleotide substitutions that change MFE (B). The “M” lanes contain the size standards; the “R” lanes contain the intact 15-nt RNAs as a control; the “K^+^” and the “-” lanes contain the RNAs incubated 24 h at 37°C in 30 mM Tris-HCl pH 8 with and without 150 mM K^+^, respectively. The positions of the uncleaved RNAs and the upstream cleavage products are marked in lanes showing self-cleavage activity by respectively black and red arrows. Note, the downstream cleavage products are too short for detection.

(C) Plot of the cleavage fraction of the MINDS ribozymes and the sequence variants after incubation in 30 mM Tris-HCl pH 8, 150 mM KCl for 24 h at 37°C versus their MFE. The line represents the fitted curve, and the dashed line indicates the MFE of 2 kcal/mol which was used as the threshold in the identification of the MINDS ribozymes.
