## Supplementary figures and images for "Widespread Presence of Catalytic RNAs in All Domains of Life"

### Figure S1.pdf

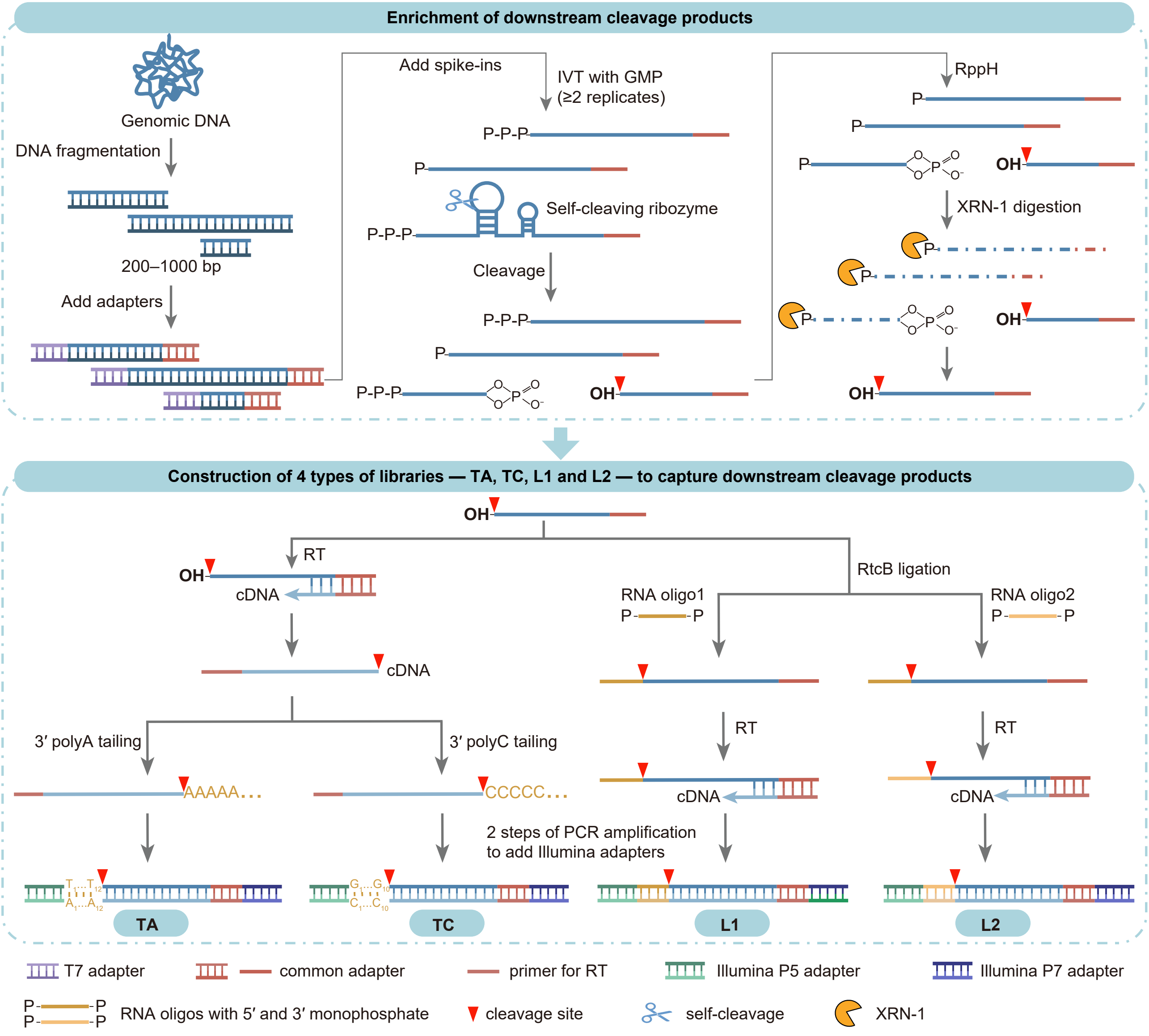

### Figure S2.pdf

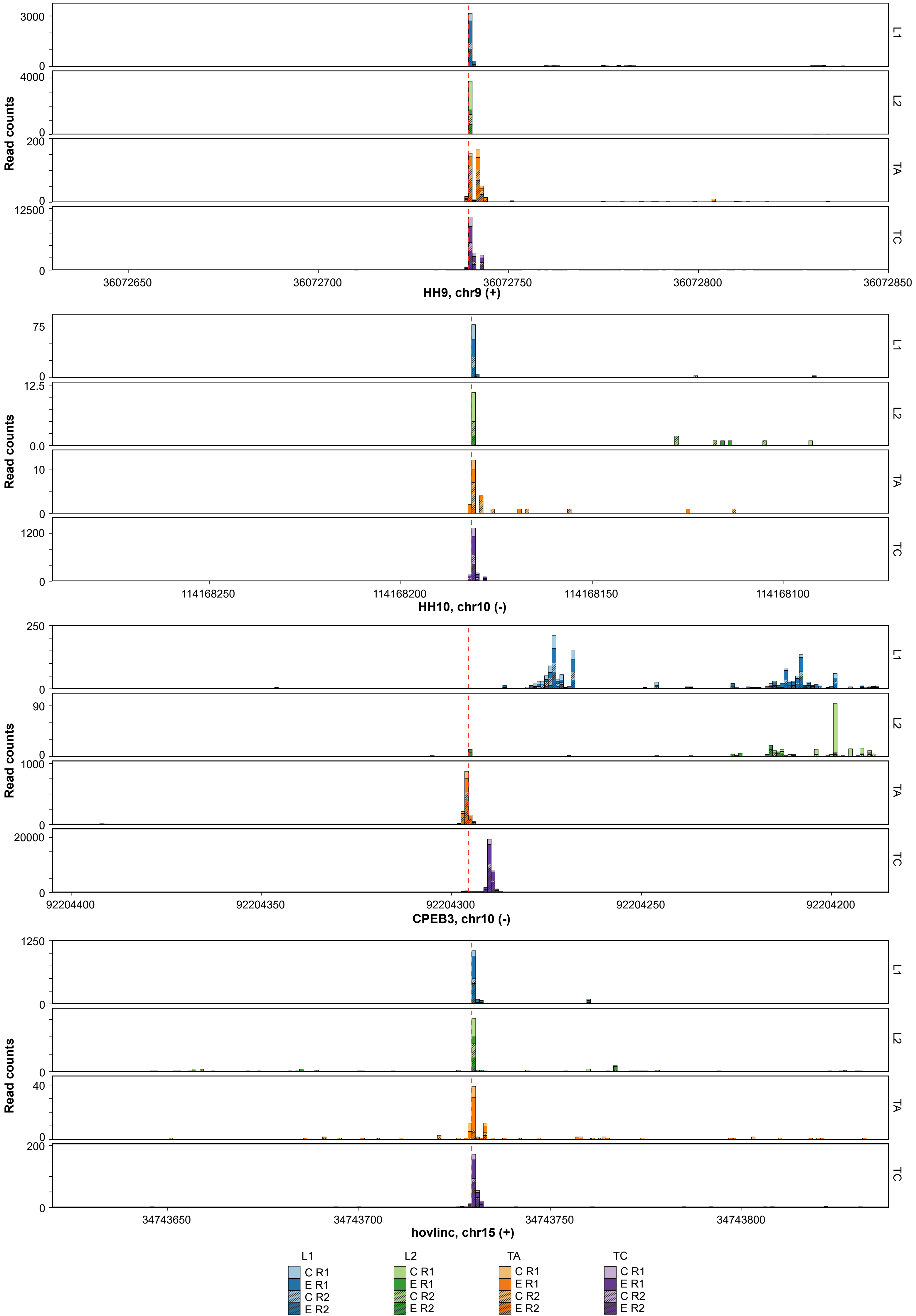

### Figure S5.pdf

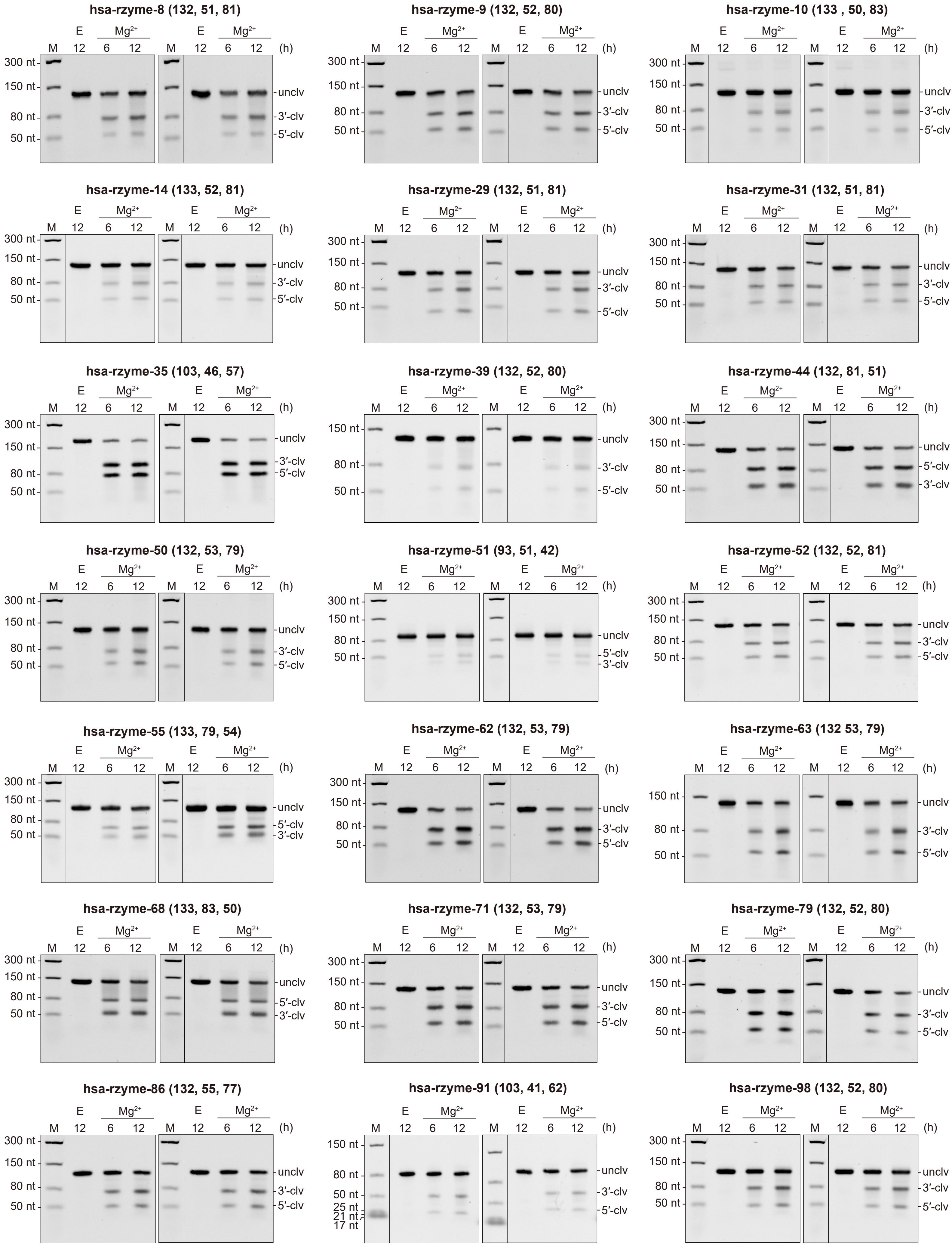

### Figure S6.pdf

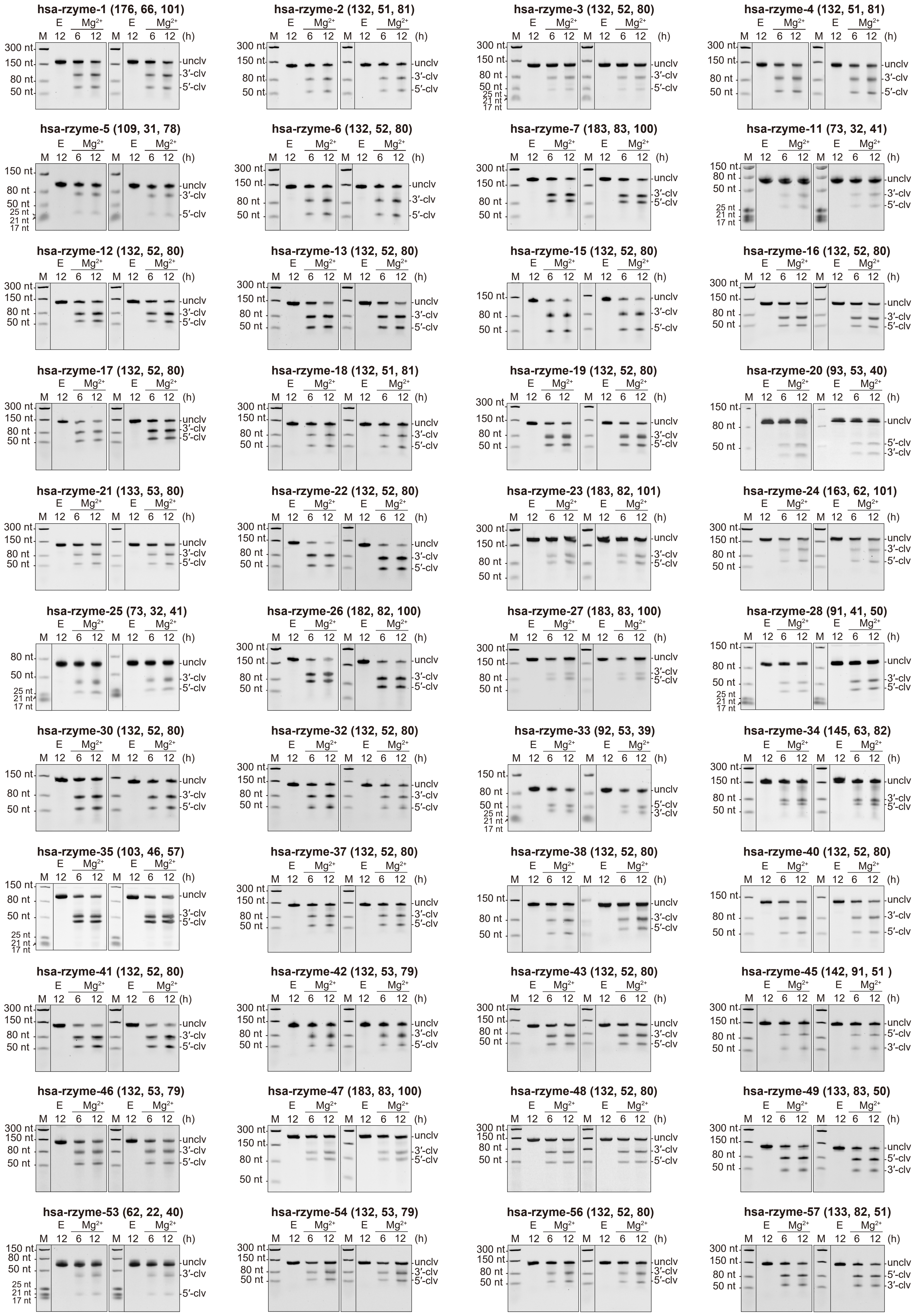

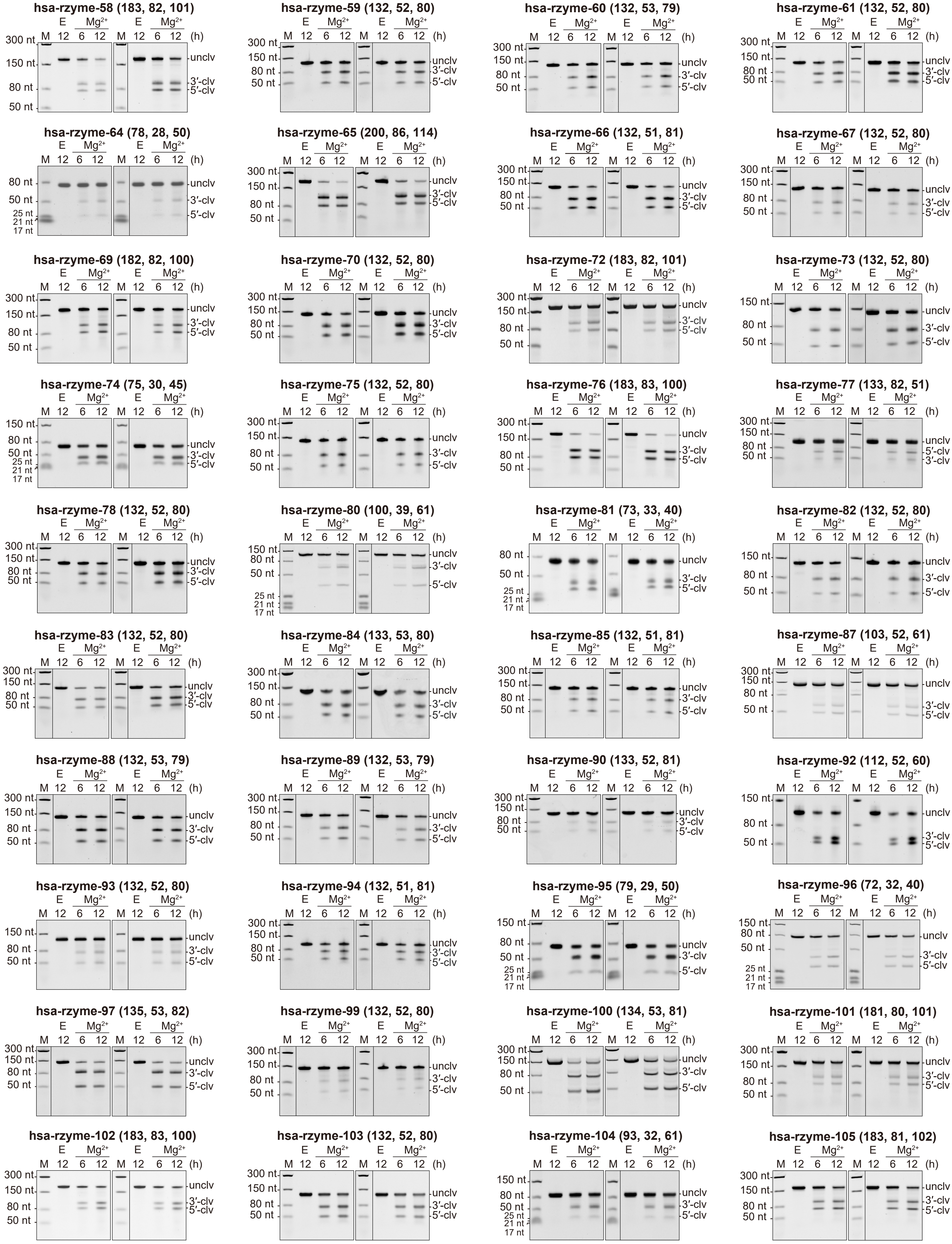

### Figure S8.pdf

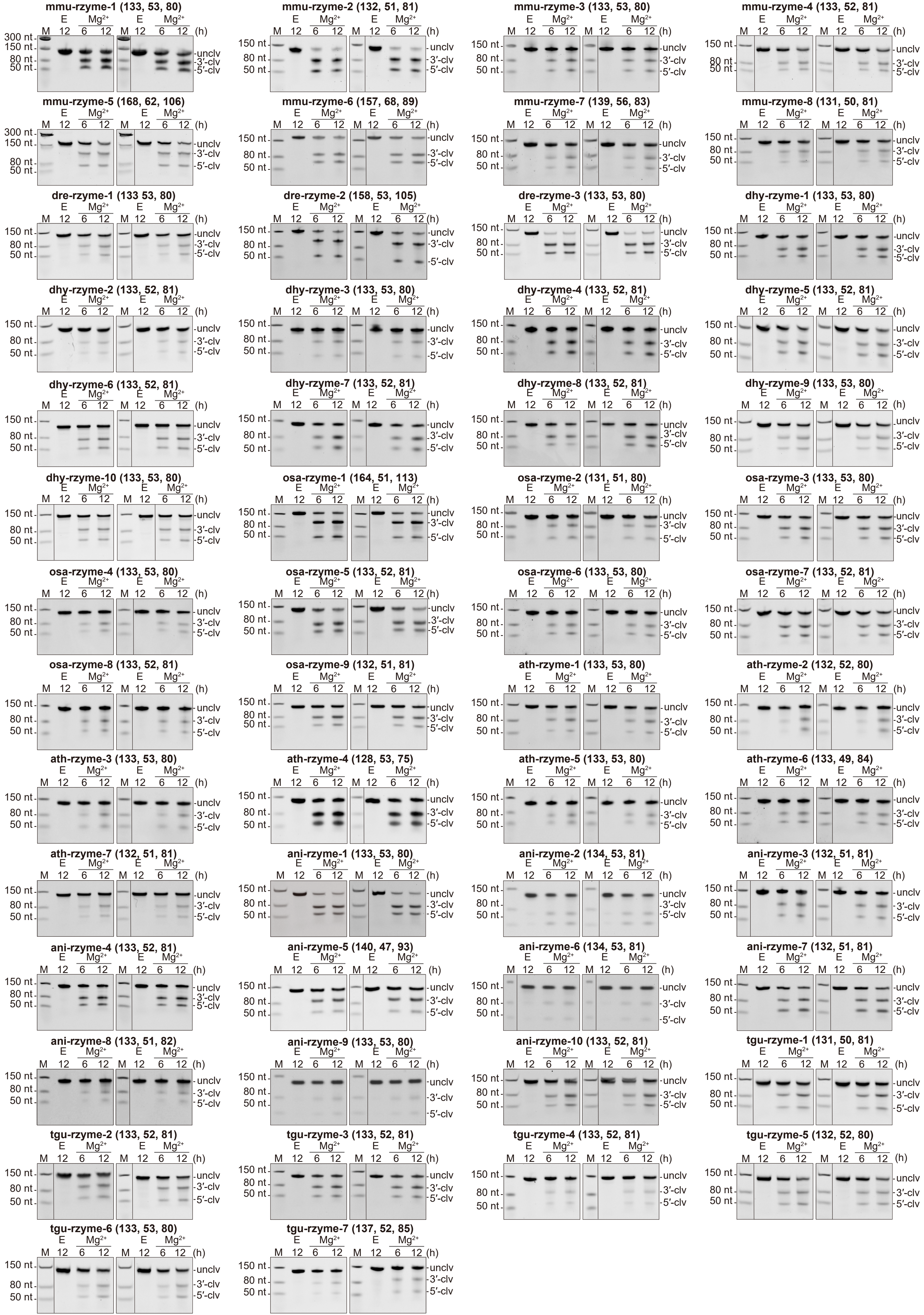

### Figure S9.pdf

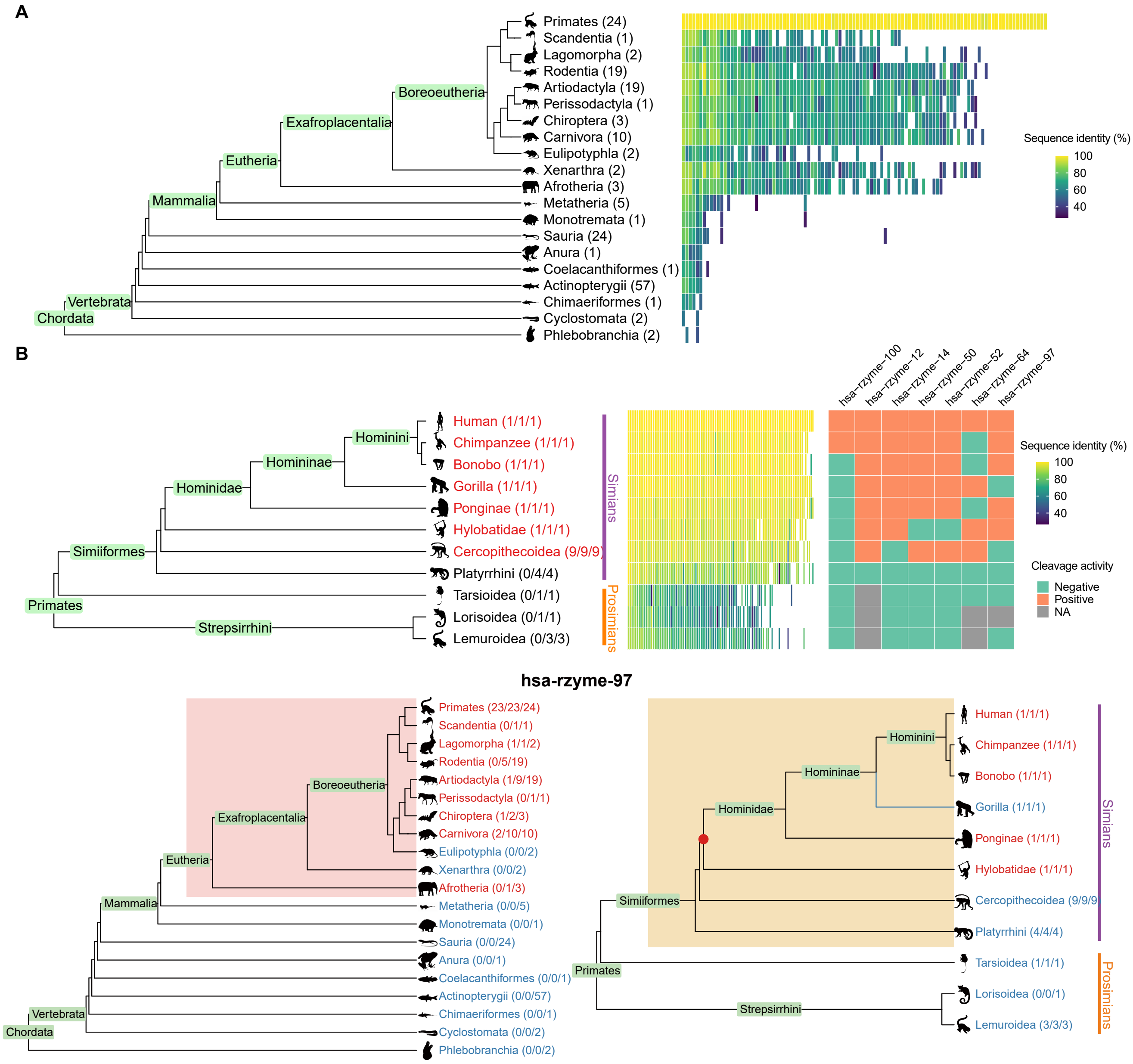

### Figure S10.pdf

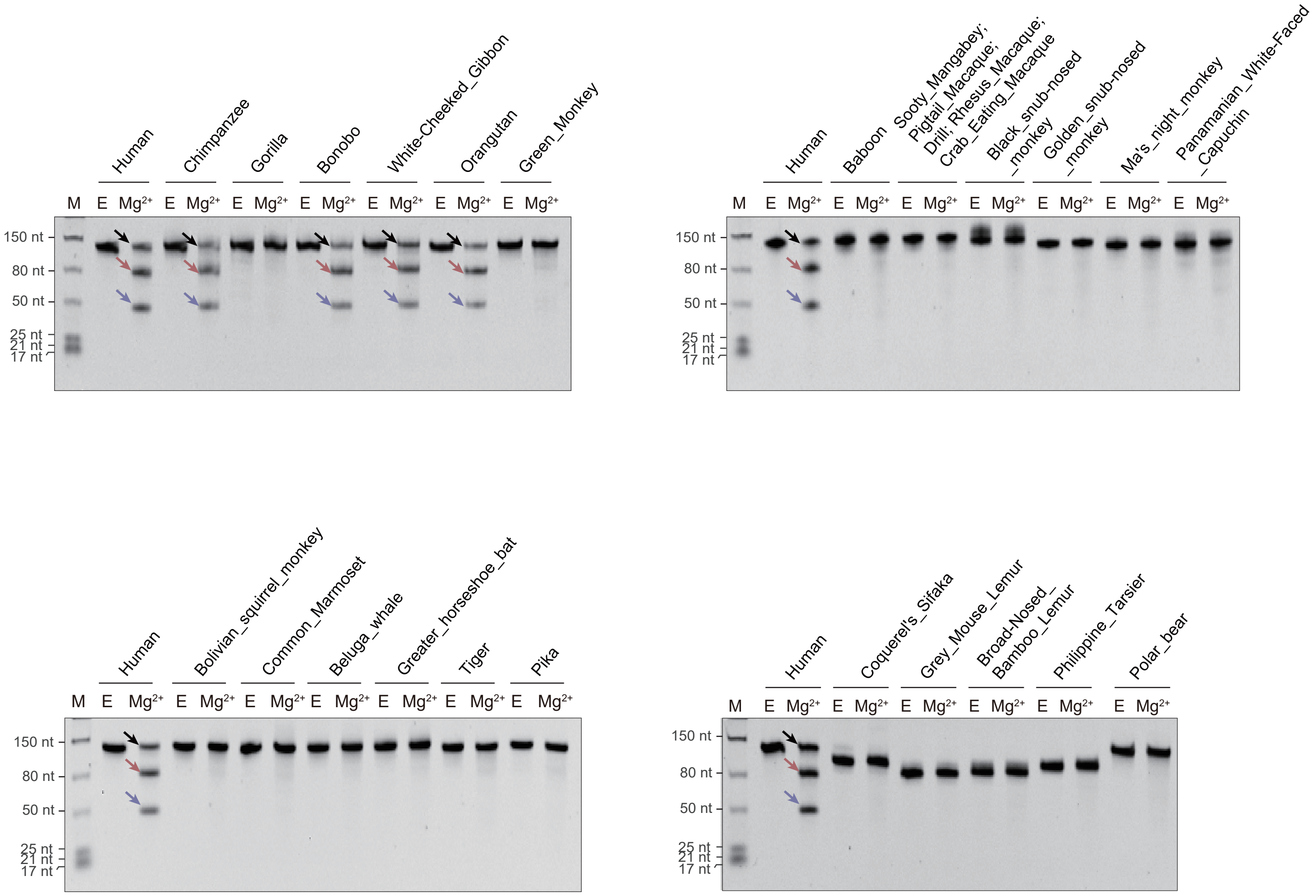

**hsa−rzyme−100**

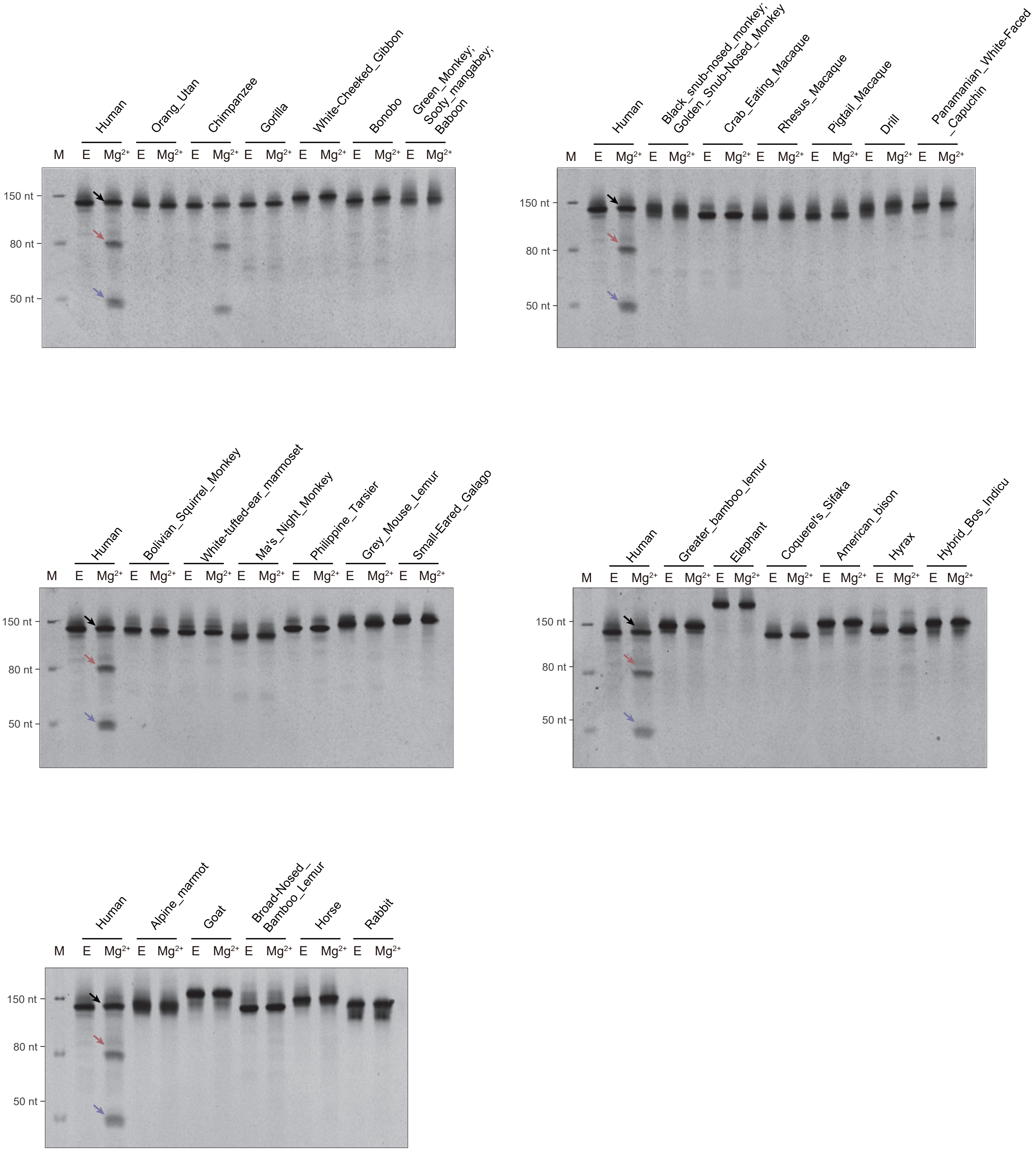

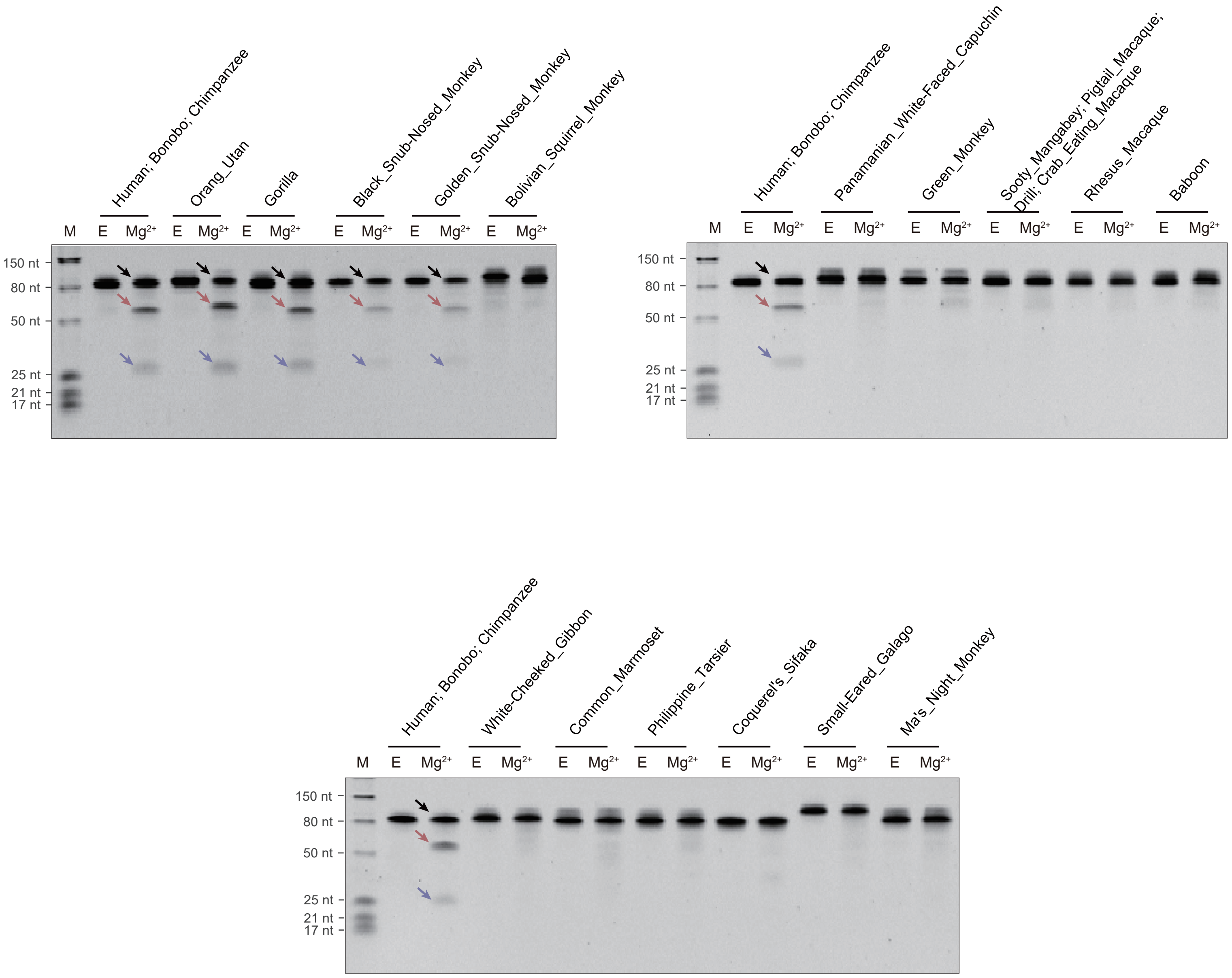

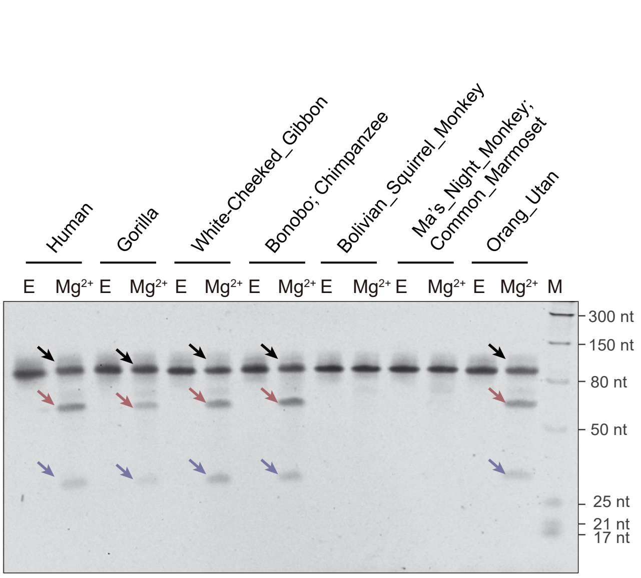

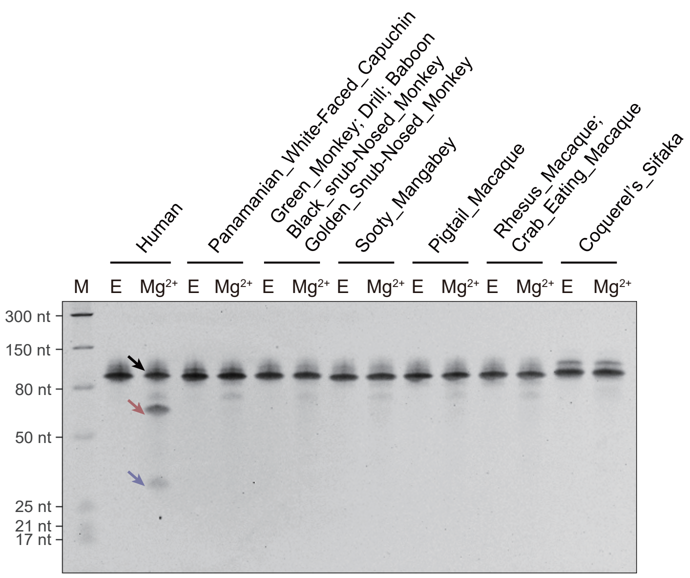

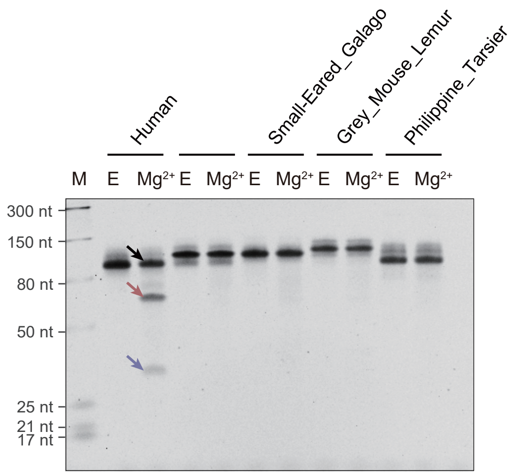

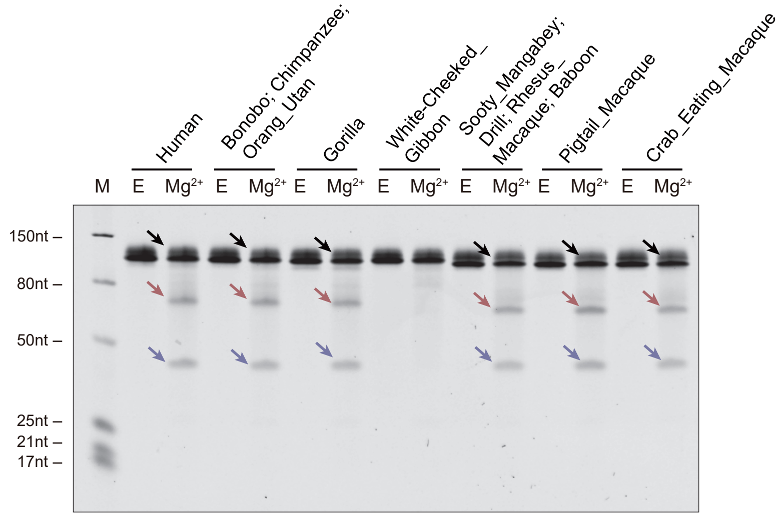

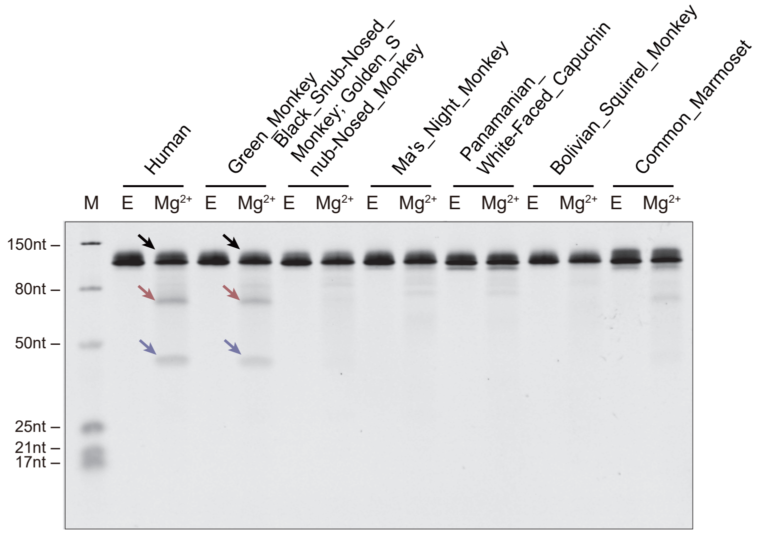

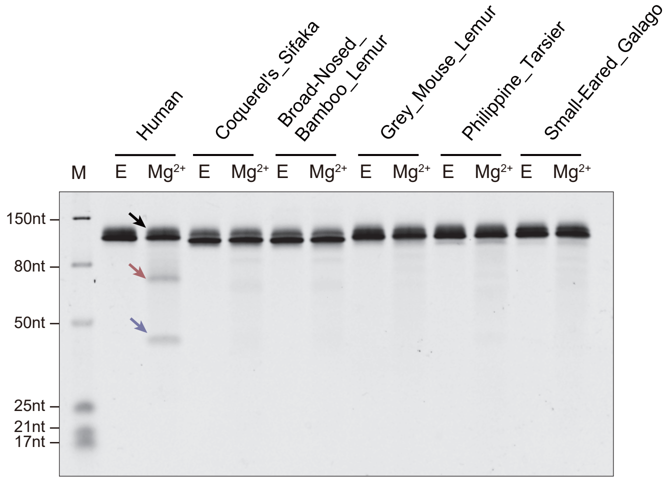

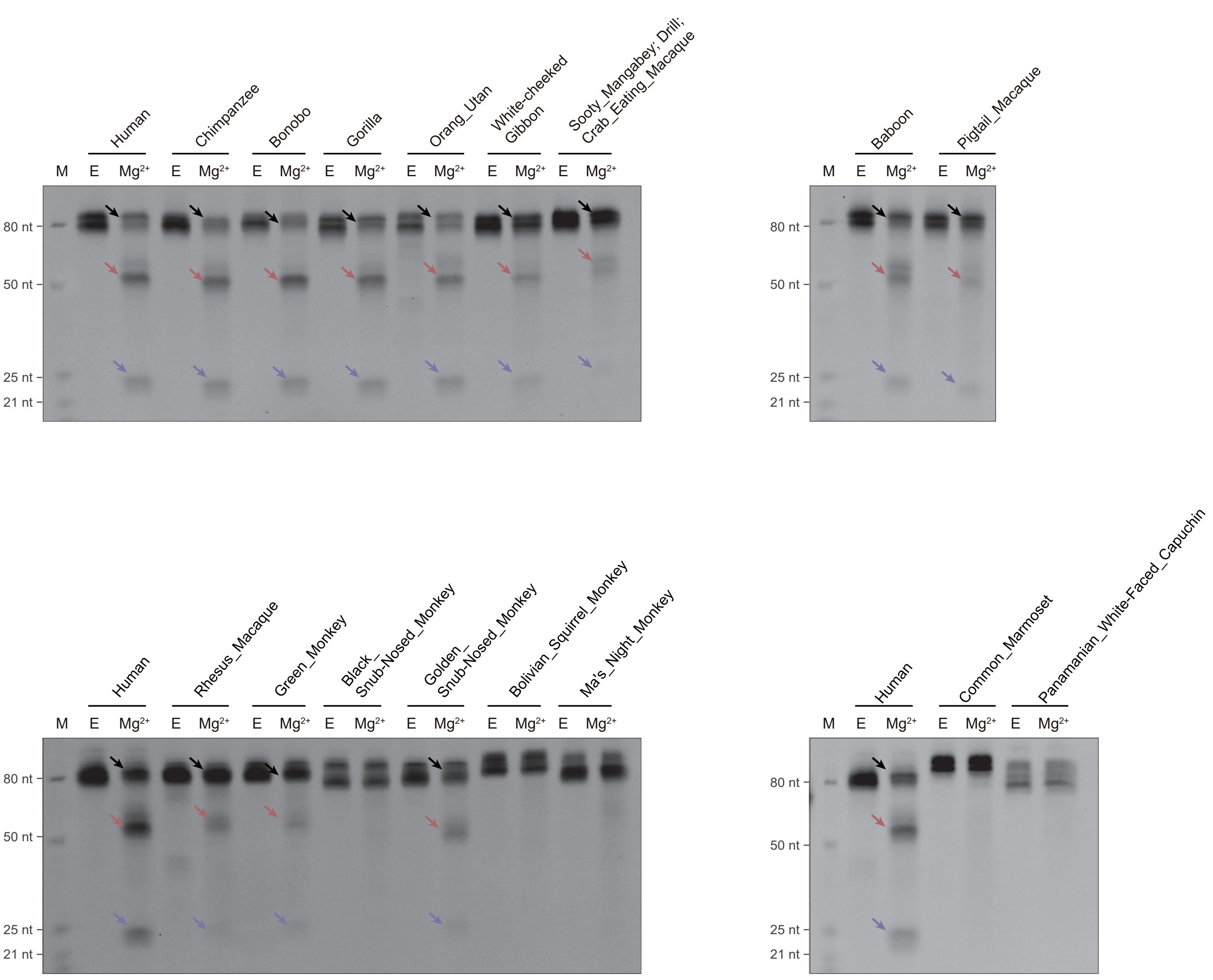

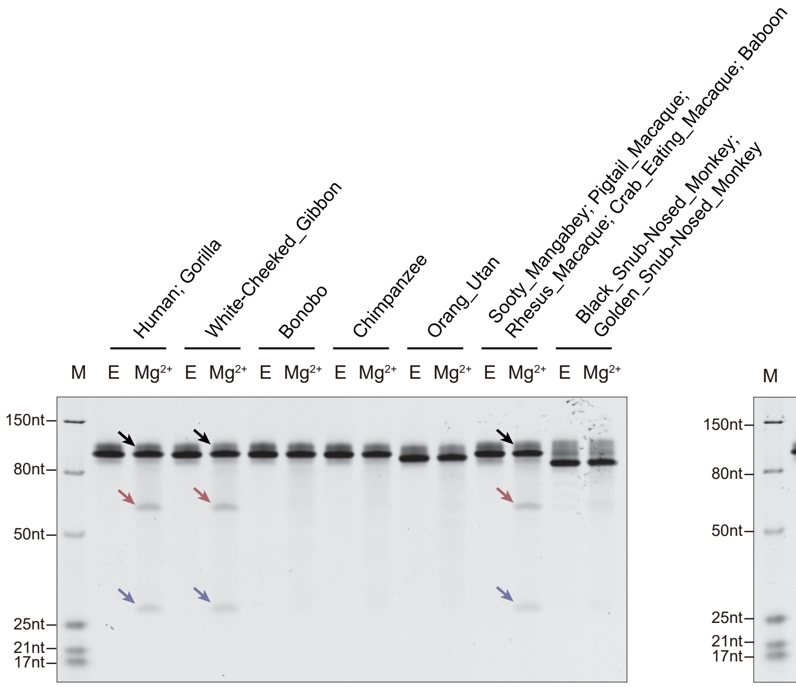

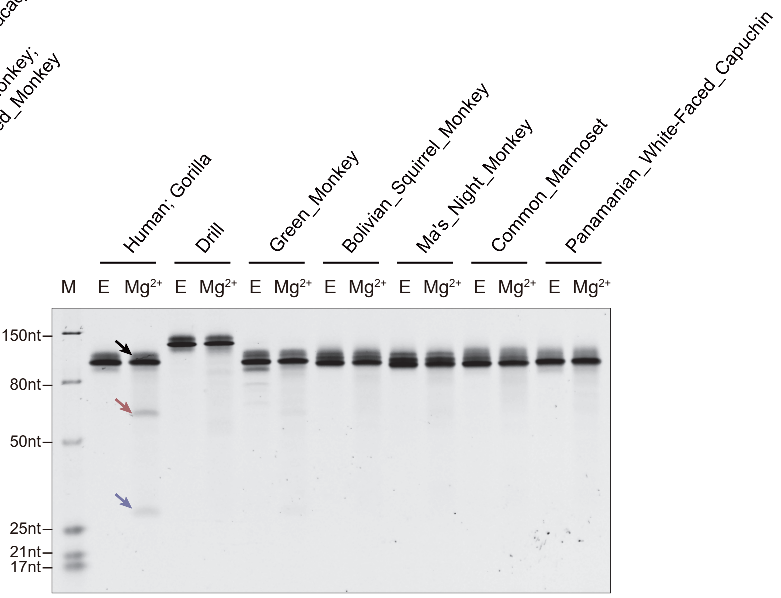

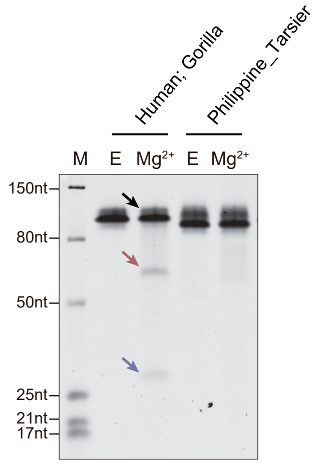

### Figure S11.pdf

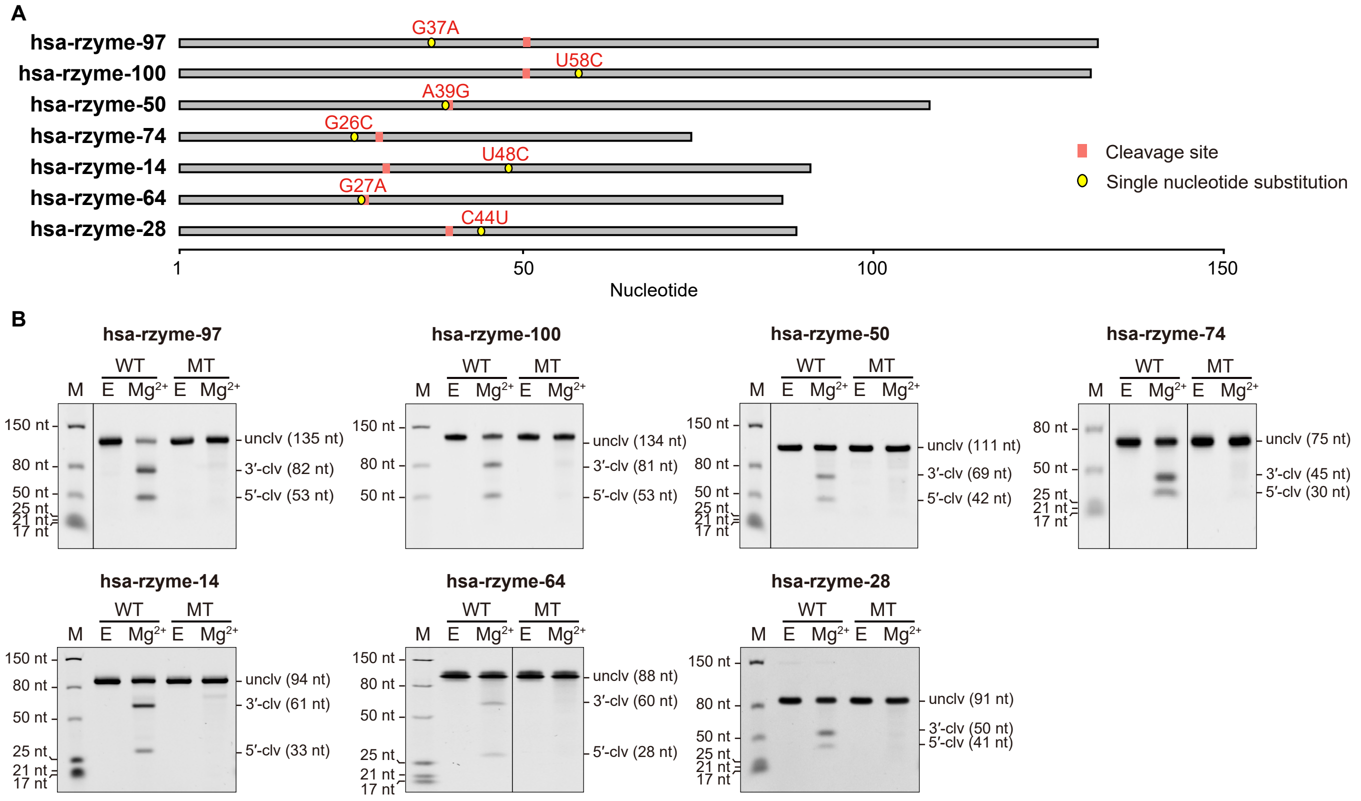

### Figure S12.pdf

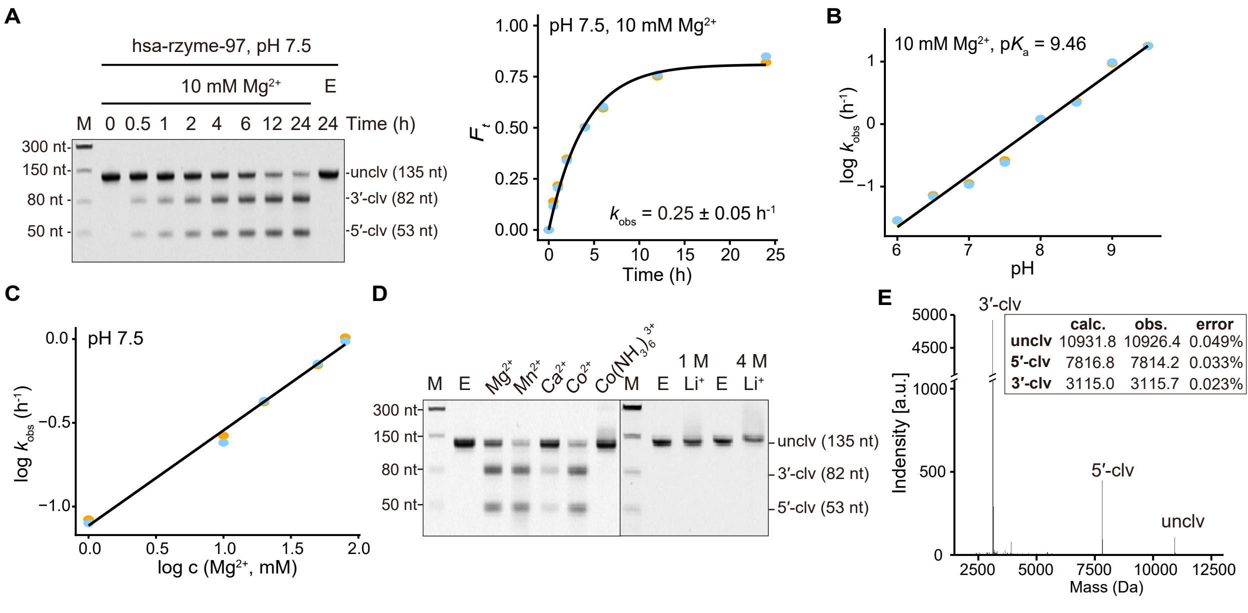

### Figure S13.pdf

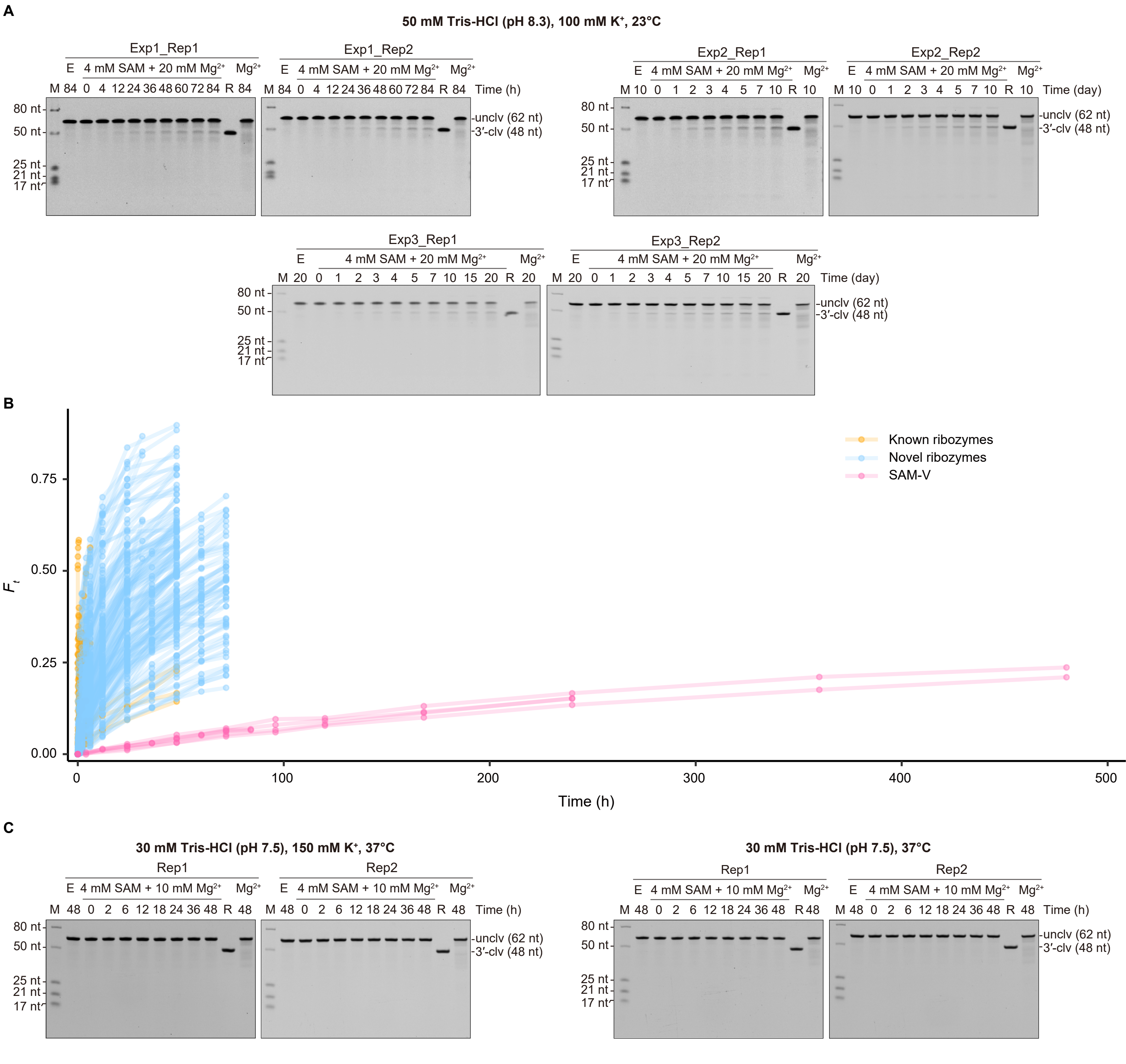

### Figure S14.pdf

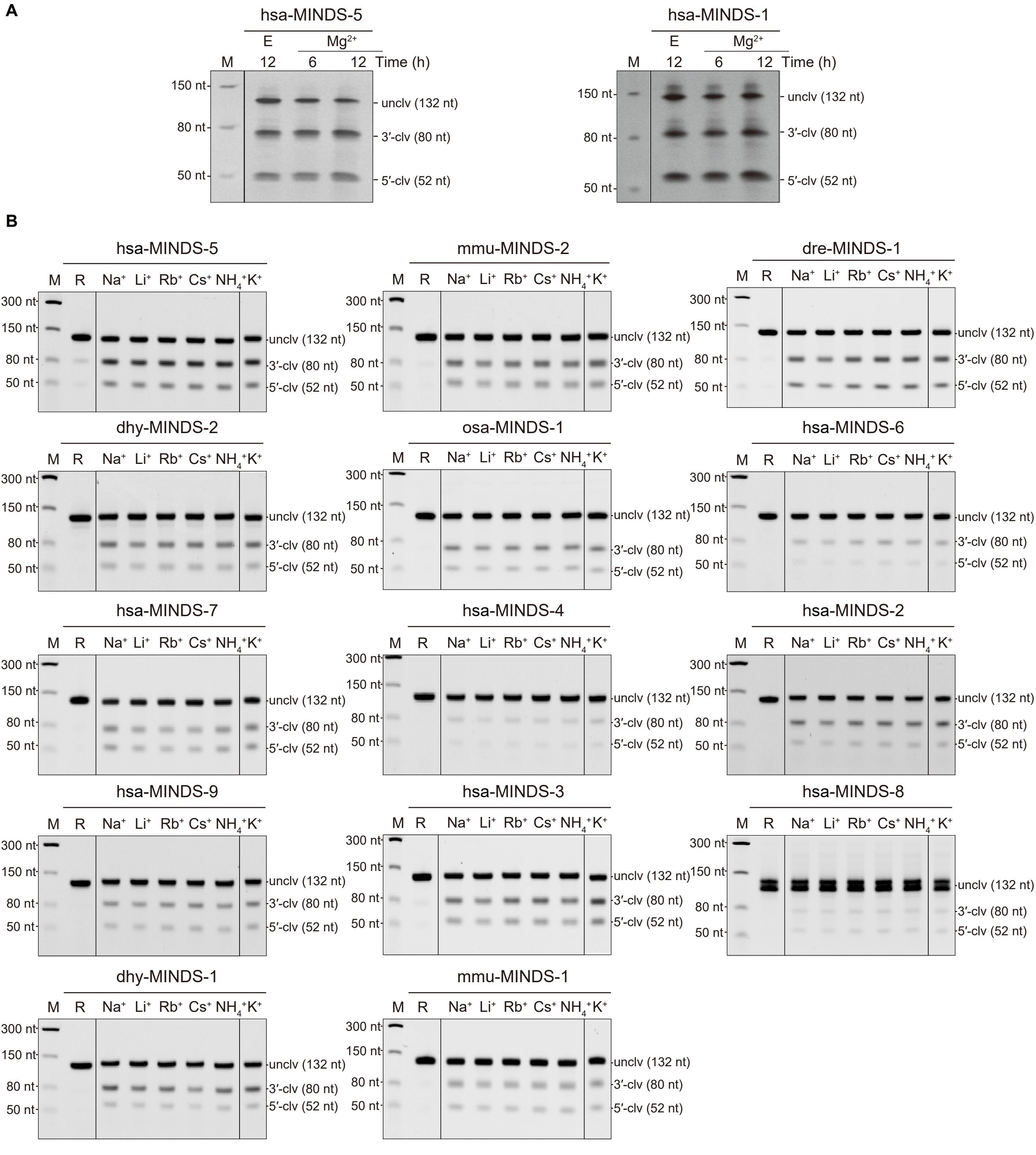

### Figure S15.pdf

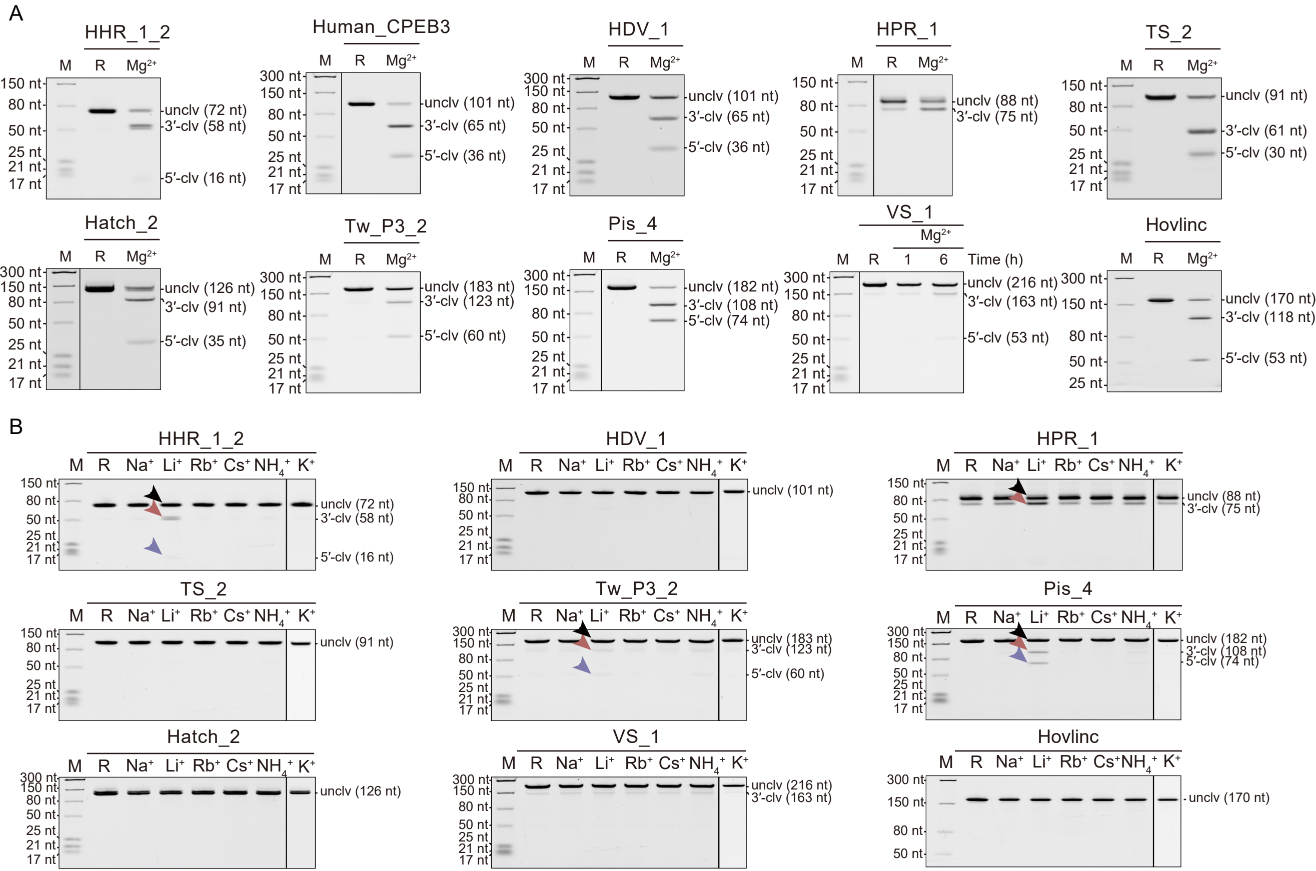

### Figure S16.pdf

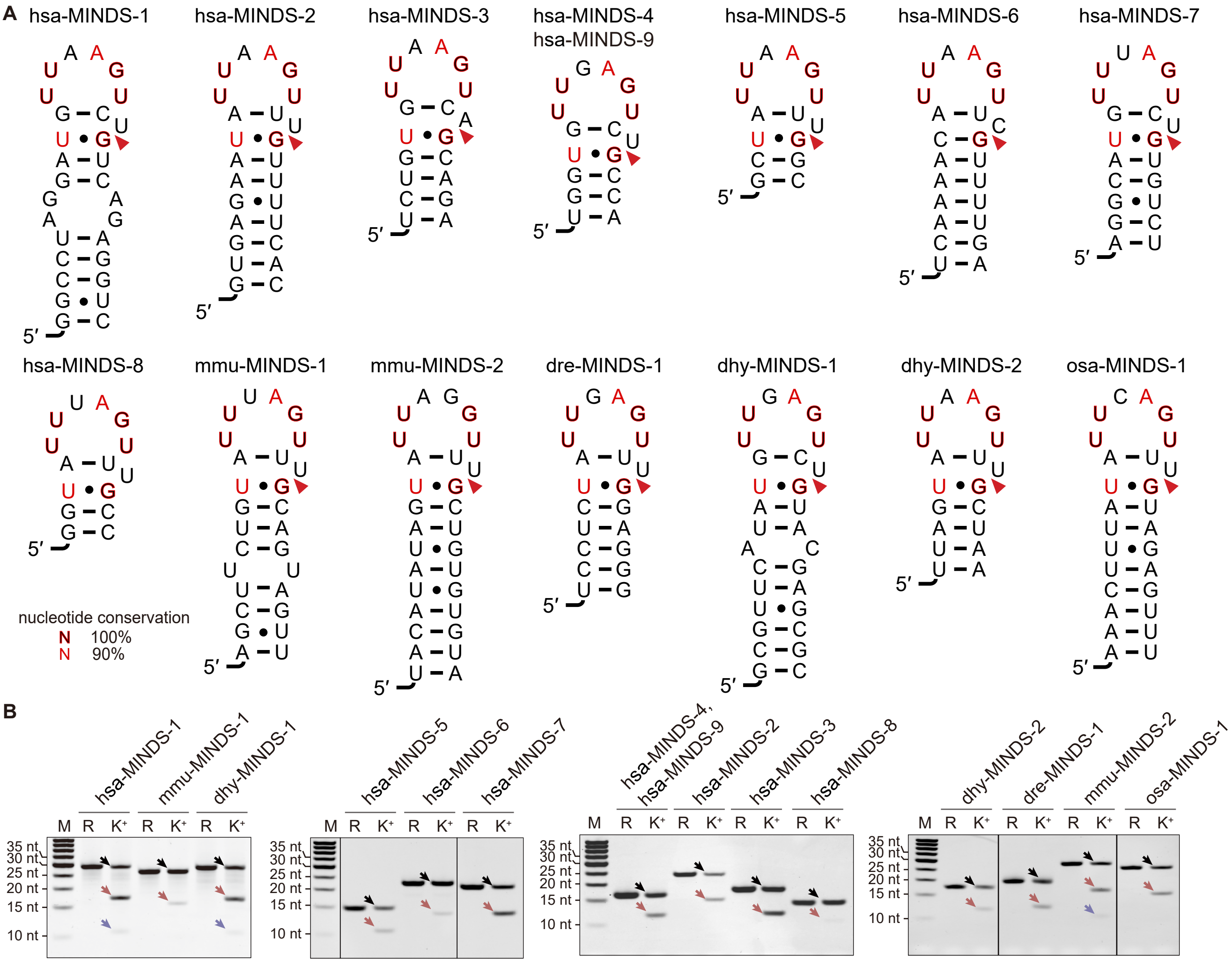

### Figure S17.pdf

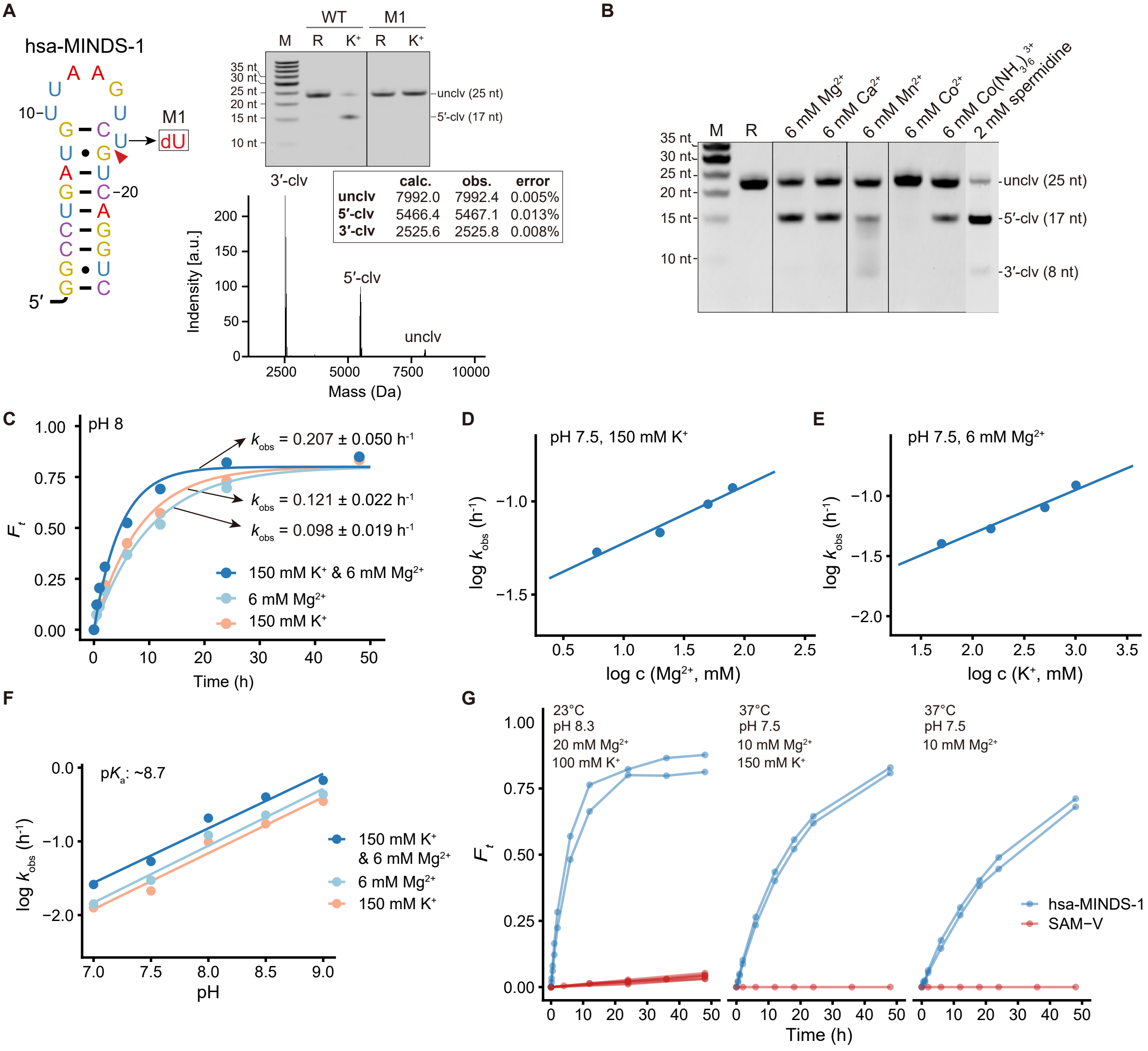

### Figure S18.pdf

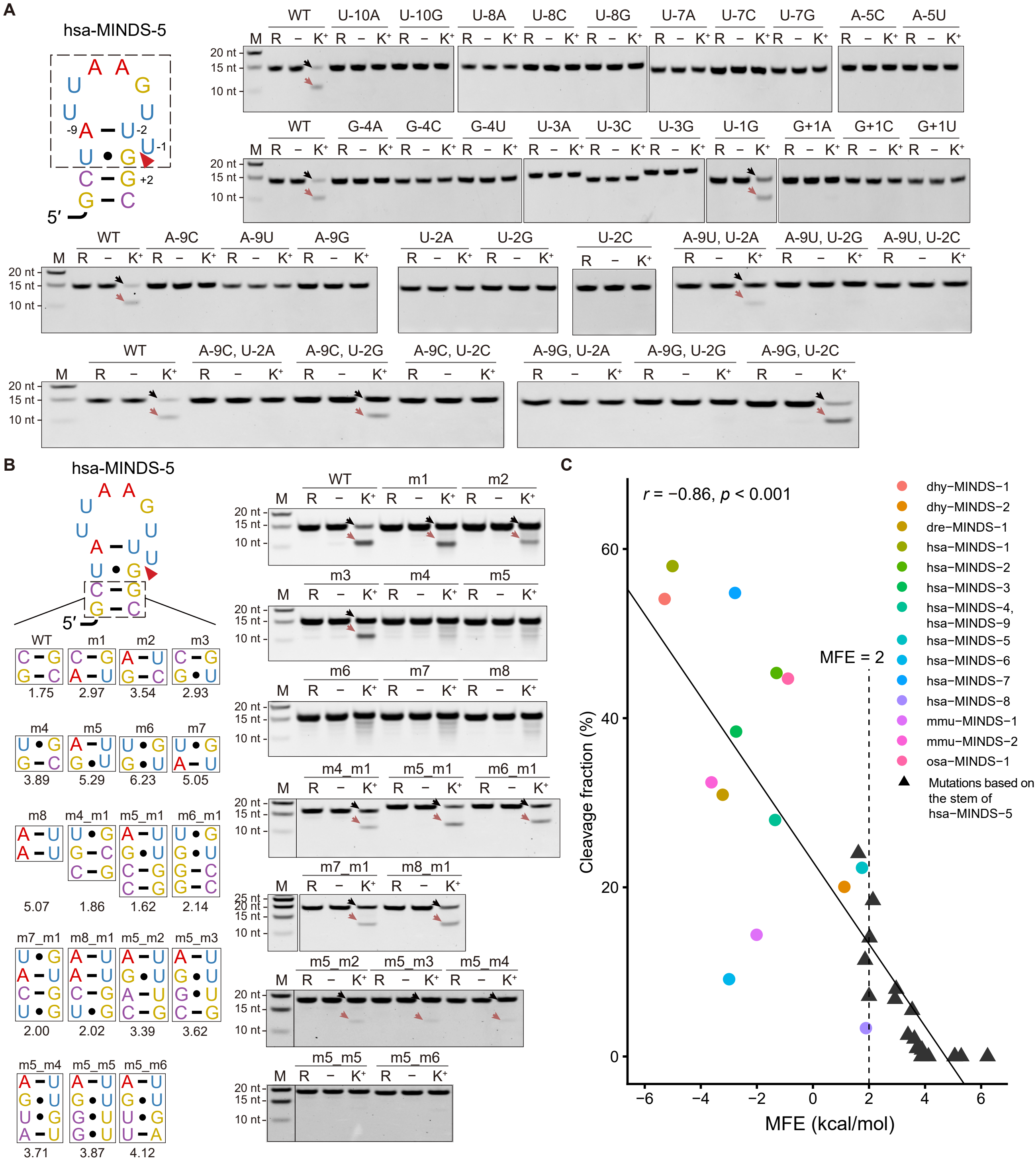
