## Supplementary Tables for "Widespread Presence of Catalytic RNAs in All Domains of Life": Supplementary Table Guide.docx

| Table S1 | All the spike-ins and known ribozyme sequences used in this study. |
| --- | --- |
| Table S2 | All the human candidate ribozyme cleavage sites in the Tiers 1–3 obtained with the AbRiDGed method in the genome-wide survey. The candidates that were subjected to the validation and the ones proven to be novel human Mg^2+^-dependent ribozymes are marked. The “cleavage sites” represent the genomic positions of the 5′ ends of the downstream cleavage products. |
| Table S3 | All of the human Tier 4 candidate ribozyme cleavage sites obtained with the AbRiDGed method in the genome-wide survey. The 50 candidates that were subjected to the validation are marked. |
| Table S4 | All the randomly selected human genomic regions and the candidate ribozyme cleavage sites in the Tiers 1–3 obtained with the AbRiDGed method. The candidates that were subjected to the validation and the ones proven to be positive are marked. The “cleavage sites” represent the genomic positions of the 5′ ends of the downstream cleavage products. |
| Table S5 | List of all the 230 candidate ribozyme cleavage sites from the 7 species in the Tiers 1–3 obtained with the AbRiDGed method that were subjected to the validation. The ones proven to be positive are marked. Each species-Tier combination is presented in a separate sheet. The “cleavage sites” represent the genomic positions of the 5′ ends of the downstream cleavage products. |
| Table S6 | The results of the clustering analysis on the known and novel ribozyme sequences. |
| Table S7 | The results of the search against the Rfam and ZWD databases using the sequences of the known and newly discovered self-cleaving ribozymes. |
| Table S8 | Genomic locations of the human self-cleaving ribozymes and their overlaps with protein-coding and lncRNA genes. The overlaps between the same ribozyme and multiple annotations are shown in separate rows. The genic and intergenic ribozymes are listed in separate sheets. The “cleavage sites” represent the genomic positions of the 5′ ends of the downstream cleavage products. |
| Table S9 | The overlaps between the human self-cleaving ribozymes and vlincRNAs. The “cleavage sites” represent the genomic positions of the 5′ ends of the downstream cleavage products. |
| Table S10 | The overlaps between the human self-cleaving ribozymes and repeats. The “cleavage sites” represent the genomic positions of the 5′ ends of the downstream cleavage products. |
| Table S11 | All the species used for the evolutionary analysis. |
| Table S12 | Summary of the evolutionary data for each novel human self-cleaving ribozyme. For the ribozymes shown in the top-right panel of Figure S9B, the primate homologs with self-cleavage activities are also indicated. |
| Table S13 | The single nucleotide substitutions that (nearly) abolished activities of the 7 novel self-cleaving ribozymes. The original (WT) and mutated (MT) sequences for each of the 7 ribozymes are shown. |
| Table S14 | Results of the genome-wide search for the human L1MA homologs. |
| Table S15 | The list of known ribozymes used for measuring the *k*_obs_ values. |
| Table S16 | The *k*_obs_ values of the novel and known ribozymes measured in this study under the two cleavage conditions with and without K^+^ |
| Table S17 | The genomic locations of the 15 MINDS ribozymes found with the AbRiDGed method in the genome-wide surveys. The “cleavage sites” represent the genomic positions of the 5′ ends of the downstream cleavage products. |
| Table S18 | The species included in this study and the accession of their genomes, as well as the number of MINDS ribozymes found in each species. |
| Table S19 | The results of the analysis for enrichment or depletion of the MINDS ribozymes in various organisms. |
| Table S20 | The overlaps between the cleavage sites of the human Mg^2+^-dependent self-cleaving ribozymes and the transcripts boundaries from the CAGE and NAP-seq datasets. The “cleavage sites” represent the genomic positions of the 5′ ends of the downstream cleavage products. |
| Table S21 | The overlaps between the cleavage sites of the human MINDS self-cleaving ribozymes and the transcripts boundaries from the CAGE and NAP-seq datasets. The “cleavage sites” represent the genomic positions of the 5′ ends of the downstream cleavage products. |
| Table S22 | The human genomic variants mapping to MINDS ribozymes —their effects on the ribozyme activity and the associated clinical traits. |
| Table S23 | All primers and oligonucleotides used in this study. |
